# Evolutionary genomics of structural variation in Asian rice (*Oryza sativa*) and its wild progenitor (*O. rufipogon*)

**DOI:** 10.1101/2019.12.19.883231

**Authors:** Yixuan Kou, Yi Liao, Tuomas Toivainen, Yuanda Lv, Xinmin Tian, J.J Emerson, Brandon S. Gaut, Yongfeng Zhou

**Affiliations:** Department of Ecology and Evolutionary Biology, UC Irvine, Irvine, CA, USA; Laboratory of Subtropical Biodiversity, Jiangxi Agricultural University, Nanchang, Jiangxi, 330045, China; Department of Biological Sciences, College of Life Science and Technology, Xinjiang University, Urumqi, China

## Abstract

Structural variants (SVs) are a largely unstudied feature of plant genome evolution, despite the fact that SVs contribute substantially to phenotypes. In this study, we discovered structural variants (SVs) across a population sample of 358 high-coverage, resequenced genomes of Asian rice (*Oryza sativa*) and its wild ancestor (*O. rufipogon*). In addition to this short-read dataset, we also inferred SVs from whole-genome assemblies and long-read data. Comparisons among datasets revealed different features of genome variability. For example, genome alignment identified a large (~4.3 Mb) inversion in indica rice varieties relative to an outgroup, and long-read analyses suggest that ~9% of genes from this outgroup are hemizygous. We focused, however, on the resequencing sample to investigate the population genomics of SVs. Clustering analyses with SVs recapitulated the rice cultivar groups that were also inferred from SNPs. However, the site-frequency spectrum of each SV type -- which included inversions, duplications, deletions, translocations and mobile element insertions -- was skewed toward lower frequency variants than synonymous SNPs, suggesting that SVs are predominantly deleterious. The strength of these deleterious effects varied among SV types, with inversions especially deleterious, and across transposable element (TE) families. Among TEs SINE and *mariner* insertions were especially deleterious, due to stronger selection against their insertions. We also used SVs to study domestication by contrasting between rice and *O. rufipogon*. Cultivated genomes contained ~25% more derived SVs than *O. rufipogon*, suggesting these deleterious SVs contribute to the cost of domestication. We also used SVs to study the effects of positive selection on the rice genome. Generally, the search for domestication genes were enriched for known candidates, suggesting some utility for SVs towards this purpose. More importantly, we detected hundreds to thousands of genes gained and lost during domestication, many of which are predicted to contribute to traits of agronomic interest.

## INTRODUCTION

Asian rice (*Oryza sativa*) is a staple food for more than one-third of the world’s population (1, 2), but the domestication history of the two main varieties of Asian rice (*Oryza sativa* ssp. *japonica* and ssp. *indica*; hereafter japonica and indica) was enigmatic until the emergence of a recent consensus. Under this consensus, japonica was domesticated from its wild progenitor (*O. rufipogon*; hereafter rufipogon) in Southern China, ~10,000 years ago (ya) and perhaps earlier (3, 4). This primary event was facilitated by selection for agronomic phenotypes that shifted allele frequencies at domestication genes, such as *sh4*, which contributes to a non-shattering phenotype (5), and *qSW5*, which affects grain width (6). While the origin of japonica has been reasonably clear, the uncertainty came primarily from the indica variety, which arose in the Asian sub-continent more recently and perhaps as late as ~4,500ya (7). For nearly a decade, evolutionary analyses came to disparate conclusions as to whether indica was domesticated completely independently (8, 9), whether the two varieties represented a single domestication event with subsequent divergence between them (10, 11), or whether the domestication of indica was geographically separate but was facilitated by the introgression of beneficial domestication alleles from japonica (11–14). This last scenario is the basis for the recent consensus (15–17).

Several lines of evidence have led to this consensus, but it has been fueled primarily by the study of population genetics across large samples of resequenced rice genomes. These studies have not only substantiated that japonica and indica are genetically distinct, but they have also supported other discernible rice groups within domesticated rice, such as the aus and aromatic varieties and temperate vs. tropical japonica (12, 18–20). Population genomic studies have also documented complex relationships between cultivated rice and *O. rufipogon*, because the latter commonly – and perhaps usually - bears the signature of introgression events from cultivated accessions (21). Population studies have also established that nucleotide diversity varies among varietal types, with diversity ~2 to 3-fold higher in indica than japonica (12, 22), implying a more severe domestication bottleneck for the latter. Finally, population genomics have been an important source for identifying putative domestication genes -- such as *Sh4*, *qSW5, qSH1*, *prog1*, *sd1*, *Wx*, *Badh2*, *Rc* and others -- via the identification of selective sweep regions (12, 18).

One major and surprising omission in these studies has been investigation of the evolutionary genomics of structural variants (or SVs) in rice genomes. SVs are commonly defined as differences between individuals in genome order or DNA content that span > 50 bp in length (23, 24). Differences in DNA order occur via inversion events, but differences in content can be attributed to a wider range of mechanisms, including deletions, duplications, translocations, and mobile element insertions (MEIs). In this context, it is important to convey that the *number* of SVs have been estimated in rice [and also in other crops (25)] based on large resequencing data sets (18, 20, 26). For example, a resequencing study reported the identification of 93,683 SVs from 453 rice accessions (20). However, to date, few studies have reported the *frequencies* of SVs within crops or contrasted SV frequencies between crops and their wild relatives (27).

The study of SV frequencies is critical for at least three reasons. First, SV frequencies can, like SNP frequencies, inform about population history, subdivision and introgression. Second, frequencies are necessary for inferring the strength and direction of evolutionary forces that act on SVs. For example, specific MEIs are often found in a low proportion of individuals within plant populations (28, 29), suggesting that they are generally deleterious and restricted from reaching high frequency by natural selection. In this context, it is interesting to note that patterns of selection can vary among transposable element (TE) families (30, 31). Finally, SV frequencies are necessary to assess whether SVs can be effectively tagged by SNPs in association analyses. There is remarkably little information on this point, but thus far it seems as if SVs are usually not in high LD with SNPs. For example, 20% of maize copy number variants (32), 27% of maize SVs (33) and ~70% of arabidopsis MEIs cannot be anchored by nearby SNPs (31). Understanding associations between SNPs and SVs is important because SVs frequently cause phenotypic change; SVs are the causative variant in at least one-third of known domestication alleles (27) and also have substantial explanatory power when they are included in GWAS analyses (26, 32, 34).

The first genome-wide characterization of SV frequencies between a crop and its wild relatives was in grapevine (*Vitis vinifera* ssp. *sativa*) (35). Zhou et al. (2019) utilized genome alignments, long-read (Pacific Biosciences SMRT) sequence data and short-read (Illumina) sequence data to identify SVs. They ultimately analyzed a set of > 400,000 highly curated SVs in a population sample of ~80 wild and cultivated individuals. They found all SV types at lower population frequencies than synonymous SNPs (sSNPs), suggesting that all SV types are deleterious, on average. The pattern did differ among SV types, however. Duplications, deletions and MEIs had estimated fitness effects that were similar to nonsynonymous SNPs (nSNPs), but inversion and translocations events were more highly deleterious. Even so, it is also clear that SVs can be beneficial. For example, duplication events have an estimated proportion of adaptive variants (a) of 25% in wild grapevines (35), and a ~4Mb inversion on chromosome 2 was likely selected by humans because it mediated a phenotypic shift from dark to white berries (35, 36).

Another interesting feature of grapevines is that there were no substantive differences in SV frequencies between wild and cultivated grapes, likely reflecting the fact that grapes did not undergo a strong bottleneck during domestication (37, 38). However, other crops, like rice (13, 22) and maize (39, 40), have experienced bottlenecks that decrease genetic diversity and repattern the frequencies of genetic variants (41). For this reason, the analysis of SVs in crops with a bottlenecked history may differ from inferences based on grapevines. Furthermore, evidence from rice (42–44) and maize (45) suggest that these bottlenecks contribute to a ‘cost of domestication’, as reflected in an increased burden of deleterious mutations. This cost has not been evaluated with respect to SVs in rice or any other grain.

Here our goal is to investigate the population genomics of SVs in cultivated Asian rice relative to its progenitor rufipogon. To do so, we take advantage of a remarkable array of publicly-available genomic resources to analyze population genomic data from 404 high-coverage resequenced genomes, including individuals from the major groups of Asian rice. With SV frequencies inferred from these short-read (Illumina) resequencing data, we address three sets of questions. First, do inferred SVs provide population genetic information that is consistent with SNPs? Second, to what extent are SVs inferred from short-read data comparable to those from other data types, such as whole-genome alignment and long-read data? Third, what do SV population frequencies suggest about the evolutionary processes that act on SVs? Finally, do SVs yield insights beyond SNPs into the cost of domestication and the genomic regions targeted by positive selection?

## RESULTS

### The population dataset

We inferred SVs from publicly-available, high-coverage, short-read resequencing data (see Methods), targeting accessions with > 15x genome coverage. The initial dataset included 404 individuals representing the five major rice subpopulations (19) (temperate japonica, tropical japonica, indica, aus and aromatic), two wild relatives (*O. rufipogon* and *O. nivara*, which is often considered an annual form of rufipogon), and two outgroup species (*O. meridionalis* and *O. longistaminata*). We mapped these resequencing data to an updated version of the Nipponbare genome (46), called SNPs, and then subjected the sample to clustering analyses (see Methods). From this analysis, we detected 46 individuals (11.39%) that did not cluster with their reported group of origin, suggesting they were misidentified in public databases (**Table S1**). After their removal, the curated dataset consisted of 358 accessions: 244 cultivated rice representing indica (*n*=96), japonica (*n*=106), aus (n=24) and aromatic (n=18) varieties; 97 wild rice, including *O. rufipogon* (n=90) and *O. nivara* (n=7) accessions; and outgroup accessions from *O. meridionalis* (n=3) and *O. longistaminata* (n=14) (**Table S2**). The mean coverage among the 358 accessions was 51.6x, with a range of 15x to 333x (**Table S2**), providing sufficient coverage for high-sensitivity SV calls (47).

Given this short-read dataset, we first focused on SNPs. SNP-calling identified 38,717,560 SNPs and small variants (i.e., indels < 50 bp). As expected (12, 18–20), phylogenetic analyses based on SNPs revealed clear group-specific clades, with the aus-indica clade separated from the japonica-aromatic clade (Fig. 1a). The exception to clear separation was *O. nivara*, which nested within rufipogon clades, but this result is consistent with previous analyses based on wide sampling of *O. nivara* and rufipogon (48). In the phylogeny, rufipogon was represented by two major clades: one that branched next to the outgroups (*O. meridionalis* and *O. longistaminata*) and a second that rooted between indica and japonica. The first clade included accessions that derive primarily from India and Southeast Asia; the second included accessions primarily from China. We used SNPs to infer genome-wide nucleotide diversity, which recapitulated the well-substantiated hierarchy of diversity within *Oryza* -i.e., rufipogon had higher diversity (⍰_w_ = 0.0213 ± 0.0022, ⍰ = 0.0129 ± 0.0015, n = 90) than indica (⍰_w_ = 0.0100 ± 0.0017, ⍰ = 0.0094 ± 0.0018, n=96), which was more diverse than japonica (⍰_w_ = 0.0057 ± 0.0012, ⍰ = 0.0039 ± 0.0012, n=106). Note that our estimates of genetic diversity were based on genomic likelihoods and thus were several times higher, and likely more accurate (49), than previous genome-wide reports based on filtered SNPs [e.g. (12)].

**Fig. 1.**
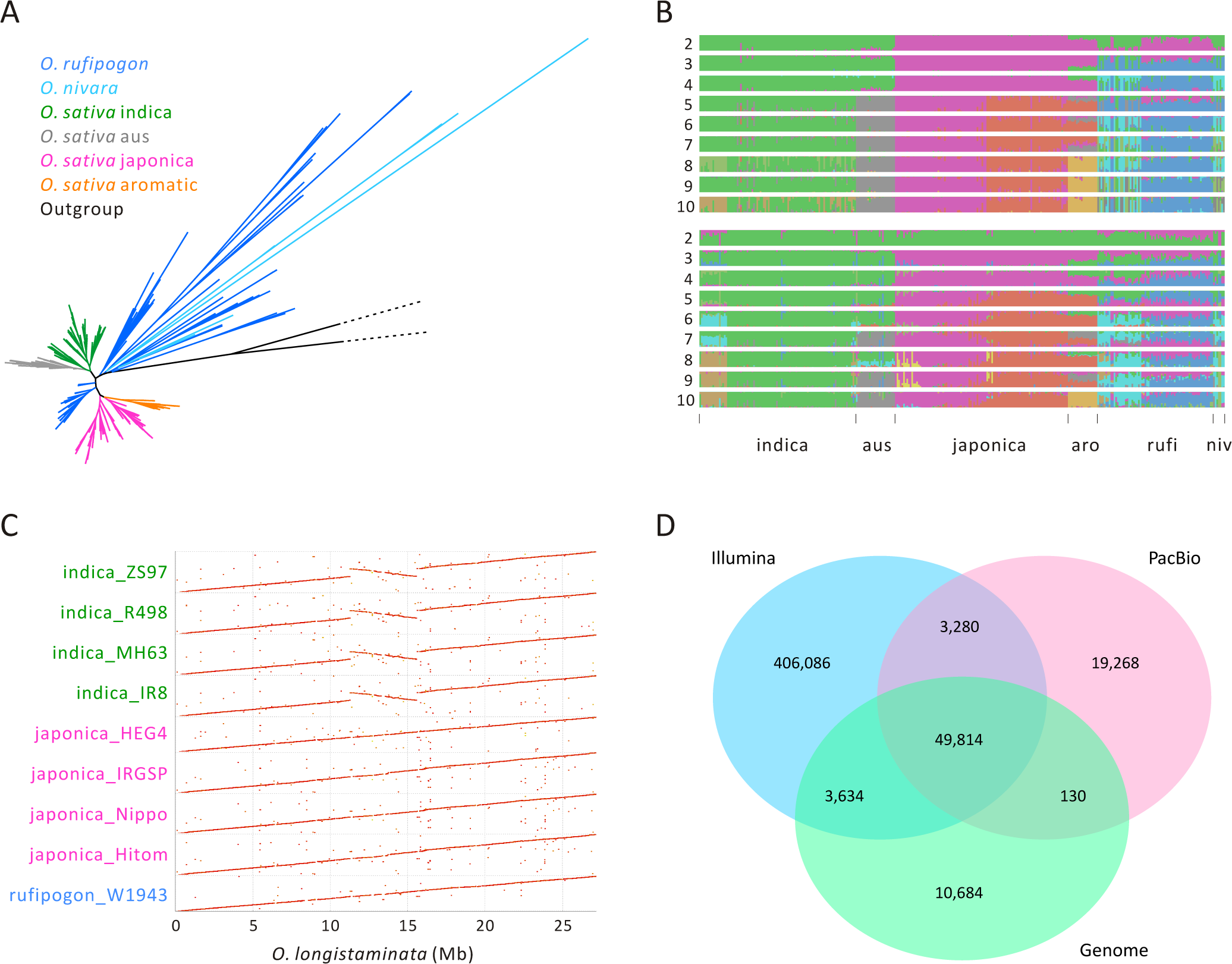
Features of the data and SV datasets (a) A phylogeny based on SNPs of n=358 accessions of Asian rice with outgroups *O. meridionalis* and *O. glaberrima*. (b) Population structure inference based on SNPs (top) and SVs for the short-read dataset of 358 individuals. The accessions are arranged in the same order for the SNP and SV plots, the x-axis labels denote the different groups, with “aro”, “rufi” and “niv” referring to aromatic, rufipogon and *O. nivara*. (c) A dotplot of chromosome 6 showing the large (~4.3 Mb) inversion in indica accessions relative to the *O. longistaminata* outgroup. The inversion is not shared with the japonica accessions in our sample. (d) A Venn diagram based on the combined results from three SV types (DEL, DUP, and INV) that compares SVs among three datasets based on short-reads (Illumina, n=358), long reads (SMRT, n = 10) and genome alignments (n= 15). Results for each SV type separately are available in **Figure S3**.

SVs were called from the set of 358 individuals using two approaches, following (35). First, to identify deletion (DEL), duplication (DUP), insertion (INS), inversion (INV) and translocation (TRA) events, we combined population calls from Delly (50) and Lumpy (47). The combined SVs were then filtered based on minimal quality scores and also on the requirement that a single SV had exact breakpoints shared among accessions (see Methods). Second, MEIs were inferred using a TE-specific platform (51) and then assigned to specific TE families (see Methods). Altogether, these approaches identified a highly curated set of 824,390 SVs across the entire data set of 358 accessions, including 72,930 DELs, 48,132 DUPs, 341,752 INVs, 284,741 MEIs and 76,835 TRAs (Table 1 **& S3**).

**Table 1.** The number of variants detected by Illumina short reads, PacBio long reads, and genome alignment.

| Variant type | Illumina | PacBio | Genome alignment |  |
| --- | --- | --- | --- | --- |
|  |  |  | LASTZ | MUMmer |
| Number of samples | 358 | 10 | 15 | 15 |
| TRA | 76,835 | 85,279 |  | 1,332 |
| DUP | 48,132 | 213 | 3,700 | 2 |
| DEL | 72,930 | 60,747 | 62,953 | 48,153 |
| INV | 341,752 | 11,532 | 159 | 343 |
| INS |  | 103,447 | 70,065 | 113 |
| MEI | 284,741 | 270,708 |  | 246,340 |
| TOTAL | 824,748 | 531,936 |  | 296,298 |

### Assessing SV calls

We examined our short-read SV calls by first comparing population information between SVs and SNPs; we then related the SVs to those inferred from other data types.

#### Comparing SVs to SNPs

We compared SNP and SV calls from the same data - i.e., the 358 resequenced individuals - to see if they provided similar insights into population structure and history (35, 52). To do so, we applied NGSadmix to the SNP data, allowing the number of groups (*k*) to range from 2 to 10. At *k* =10, the SNP data recapitulated the expected population groupings, in that: *i*) cultivated individuals differed from wild individuals, *ii*) indica and japonica were separated from each other, and *iii*) further expected subgroupings (Fig. 1a) were recapitulated, such as temperate vs. tropical japonica and the two distinct clades of rufipogon (Fig. 1b). We then applied ADMIXTURE to the full dataset of SVs, which identified the same major groups (Fig. 1b). Per-individual assignments were strongly correlated between SV and SNP results (Pearson’s *r* =0.853; *p* < 2.2 × 10^−16^). This concordance between population information based on SNPs and SVs supports the suitability of SV calls for population genetic analyses.

#### Comparisons across datasets

To further evaluate our SVs, we compared them to SVs inferred from two other types of sequence data: whole-genome assemblies and SMRT reads (**Table S4**). SVs were inferred from genome assemblies based on pairwise genome alignments using MUMmer (53) and LASTZ (54) (see Methods). Across the pairwise comparison of 15 genomes, including *O. glaberrima* (55) and *O. longistaminata (56)* outgroups, we detected a total of 390,823 SVs across 15 assemblies (Table 1 **& S3)**. Notably, the SVs included a ~4.3 Mb (coordinates: 11296942-15576712 bp in the *O. longistaminata* reference) homozygous inversion that spanned the centromere of chromosome 6 and differed between the four indica assemblies and assemblies from other *Oryza* species (Fig. 1c; **Figs. S1 and S2**).

In addition to whole-genome assemblies, we collected raw SMRT reads from the subset of six indica, three japonica and one *O. longistaminata* accession that were used in genome alignment (**Table S4**) and then mapped SMRT reads to Nipponbare genome (Zhang et al. 2018) using Minimap2 (57). SVs were called across all ten samples using the Sniffles pipeline (58), identifying 60,747 DEL, 213 DUP, 11,532 INV, 103,447 INS, 85,279 TRA and 270,708 MEIs (**Table S3**). SMRT reads also provide the opportunity to investigate within-genome hemizygosity, based on the presence of alternative reads that do and do not span a genomic region (35). Applying this approach to a japonica individual (Nipponbare), an indica individual (93-11) and a wild outcrossing relative *O. longistaminata*, we detected similar numbers of presence-absence variants (PAVs) in Nipponbare (257 DELs and 31 DUPs) and 93-11 (306 DELs and 18 DUPs). For these two accessions, only 0.73% (385) and 0.35% (171) of genes were hemizygous. In contrast, 56-fold more PAVs were found in the outcrossing *O. longistaminata* accession, resulting in hemizygosity for 8.89% (or 3895) genes.

Finally, we compared the three datasets for three SV categories that were most comparable across methods: DEL, DUP and INV events. We first contrasted the genome and SMRT-read datasets. For the three categories, more SVs were detected with SMRT-reads (72,492) than with genome assemblies (64,262), despite the fact that data for the former (*n* = 10) were a subset of the latter (n=15; **Table S4**). Notably, the pipeline for genome alignments detected far 10-fold fewer INV events, while SMRT reads yielded 10-fold fewer DUP events (**Fig. S3**). Altogether, however, the two datasets had 57% of DEL, DUP and INV events in common (Fig. 1d **& S3**). We then compared short-read SVs to the two other datasets, but this was an inherently biased process because the sample size was much larger for short read data, yielding ~5x more SVs despite extensive filtering. Given this bias, we examined the overlap among data sets in a directional manner, asking: How often do Illumina SV calls identify SVs found within the other two datasets? The Illumina dataset included 70.01% of DEL events identified within the whole genome and SMRT-read datasets, 75.23% of DUP events, and a lower proportion (43.21%) of INV events. Summing across the three SV types, the short read data identified 64.35% of SVs found in the two other datasets (Fig. 1d). Notably, this level of correspondence (64.35%) exceeded that between the genome and SMRT-read datasets (57%), again suggesting the short-read population calls are reasonable.

### Population properties of SVs

We investigated the population frequencies and dynamics of SVs from the short-read data. For this purpose, we focused on the three groups with the largest population samples: rufipogon, indica (*n*=96) and japonica (*n*=106). For rufipogon, we examined only the clade of *n*=40 accessions that appeared to be truly wild, based on their phylogenetic position (Fig. 1a), further reasoning that combining the two distinct clades would produce skewed population statistics.

We first characterized chromosomal positions of SVs, plotting SV diversity using sliding window analyses for each taxon (e.g., **Figs. S4-S6**). Visually, there were no compelling patterns that suggested particular regions were more prone to specific SV events. However, SV and SNP diversity were slightly but significantly correlated across chromosomal windows in all three population groups (*r* = 0.0332, *p* = 1.07 × 10^−5^ for rufipogon; *r* = 0.0637, *p* < 2.2 × 10^−16^ for indica; *r* = 0.0494, *p* = 1.07 × 10^−10^ for japonica; Figs. 2a & **S7**).

**Fig. 2.**
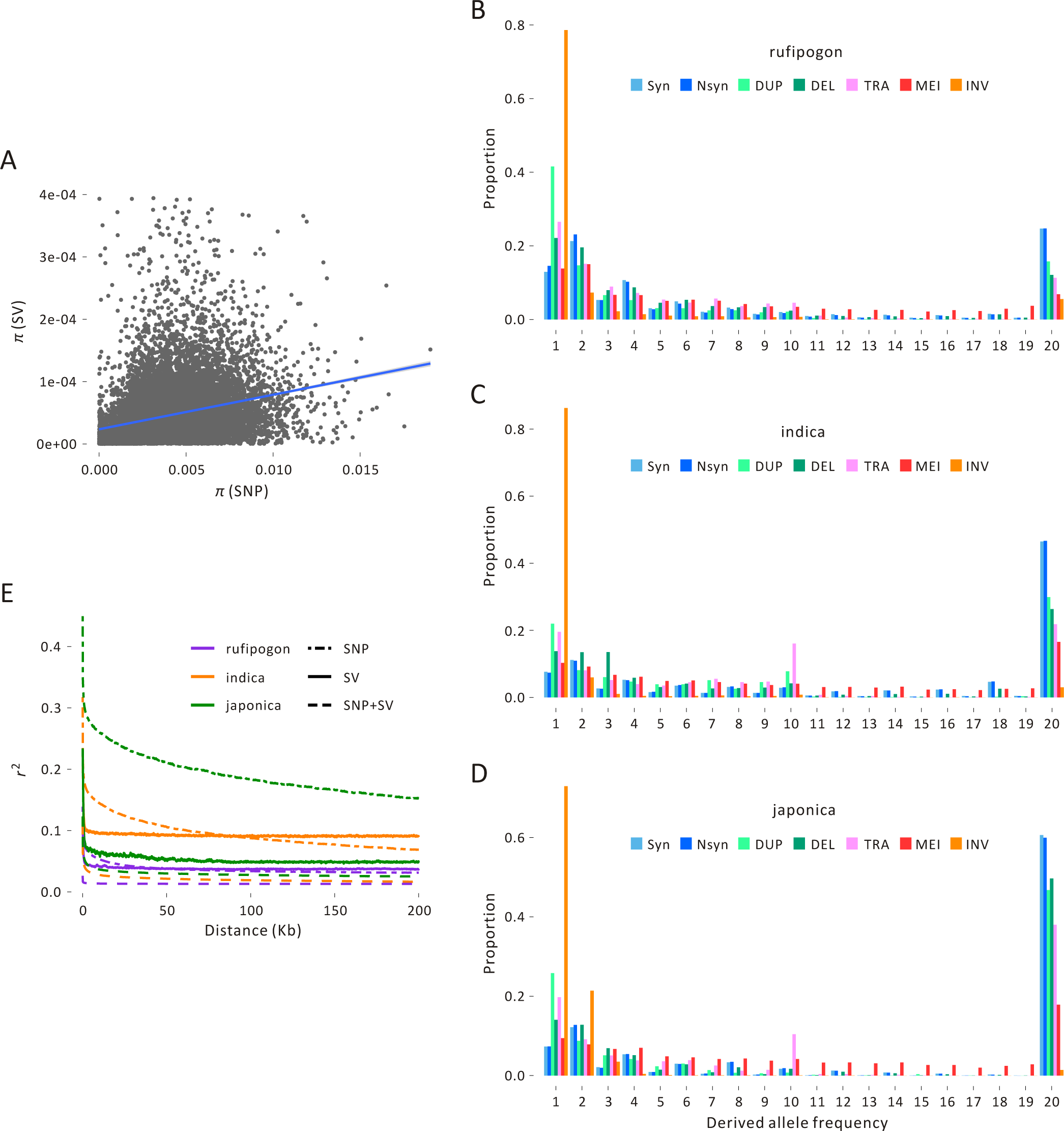
Population information about SVs. (a) The plot graphs SNP and SV average pairwise diversity (π) for the rufipogon sample across 20kb windows of the genome, with the line indicating the correlation, which is weakly positive but significant (*r* = 0.0332, *P* = 1.07e-5). Similar graphs for the japonica and indica samples are in **Figure S7**. Plots (b), (c) and (d) show the unfolded SFS of different types of SVs in (b) rufipogon, (c) indica, and (d) japonica. Each SFS contains synonymous SNPs (Syn), nonsynonymous SNPs (Nsyn), and SV events that fall into the duplication (DUP), deletion (DEL) translocation (TRA), mobile element insertion (MEI) and inversion (INV) categories. (e) The decay of linkage disequilibrium (LD) of SNPs and SVs measured by *r*^2^ for the three population groups based on SNPs, SVs and SNPs+SVs.

To illustrate the population frequencies of SNPs and SVs, we calculated the unfolded site-frequency spectra (SFS) of the three taxa for a sample of ten individuals with high coverage and little missing data (Fig. 2b-d). Each SFS included five SV types (DUP, DEL, TRA, MEI and INV), along with sSNPs and nSNPs. Altogether, the SFSs reveal three salient features of SV polymorphism. First, there were demonstrable differences among taxa, because there was a higher proportion of fixed variants (and fewer intermediate variants) in cultivated rice compared to rufipogon. The U-shape of the SFS from cultivated rice had been noted previously and is consistent with both enhanced genetic drift during a domestication bottleneck (13) and a shift in mating system. Second, in all three taxa, there was a lower proportion of fixed SVs than fixed sSNPs and nSNPs. The distributions for each SV type were significantly different from the sSNP distribution in all three taxa (*P* < 0.01, Kolmogorov-Smirnov test). Assuming sSNPs provide a reasonable “neutral” control, the leftward shift in the SFS suggests that SV variants were deleterious, on average. Finally, the SFS varied among SV types. In all three taxa, INV events had the most extreme SFS; in each group, >90% of INV events were identified in three or fewer individuals, suggesting either strong selection or perhaps detection biases (see Discussion).

The SFSs suggest that all SV types have lower population frequencies, on average, than sSNPs in all three taxa. As a consequence, SVs may have generally lower LD values than SNPs, with the potential for a faster decay of LD over physical distance. We calculated LD in each of the three taxa based on SNPs, SVs and both SNPs+SVs, using the squared correlation coefficients (*r*^2^). The SNP data confirmed previous observations that LD decays more slowly in japonica than either indica or rufipogon (59) (Fig. 2e). For example, *r*^2^ for SNPs remained ~0.2 over a distance of ~100kb for japonica, but was ~0.1 for indica and < 0.05 for rufipogon over the same physical distance. Note, however, that SVs had lower *r*^2^ values than SNPs for all taxa, with a value that exceeded 0.1 only over very short (<15kb) distances. The *r*^2^ values were even lower when based on both SNP+SV data.

### MEIs for specific TE families

MEIs represent a special SV category, because TEs are frequently implicated in phenotypic change (60). The SFSs in Fig. 2 show that MEIs have a distribution similar to that of nSNPs, suggesting that as a whole they are mildly deleterious. However, the MEI category combines information across many TE families. To better investigate the evolutionary dynamics of TEs, we separated MEIs into ten distinct TE families - *Gypsy*, *Copia*, LINE, SINE, CACTA, *hAT*, Mutator, *Harbinger*, *Mariner* and *Helitron* elements - and calculated their separate SFS. Focusing on the results from rufipogon for simplicity (Fig. 3a), but with similar results for indica and rufipogon (**Fig. S8**), all TE families had fixation frequencies lower than sSNPs, with each distribution significantly different from sSNPs (*P* < 0.01, Kolmogorov-Smirnov test). However, there were also marked differences among TE families (Fig. 3a **& S8**). The most obvious deviation was for SINE and *Mariner* elements, for which only a small proportion of MEIs were fixed; *hAT* and *Harbinger* elements also demonstrated a substantial leftward trend relative to *Gypsy*, *Copia* and other elements types. Consistent with this observation, estimation of the distribution of fitness effects (DFEs) suggest that selection was more severe against these four TE families (Fig. 3b), with the lowest proportion of putatively adaptive variants (α) for SINEs (α = 0.12%) followed by Mariner (α = 1.52%), Harbinger (α = 6.32%), and hAT (α = 5.91%) (Fig. 3c).

**Fig. 3.**
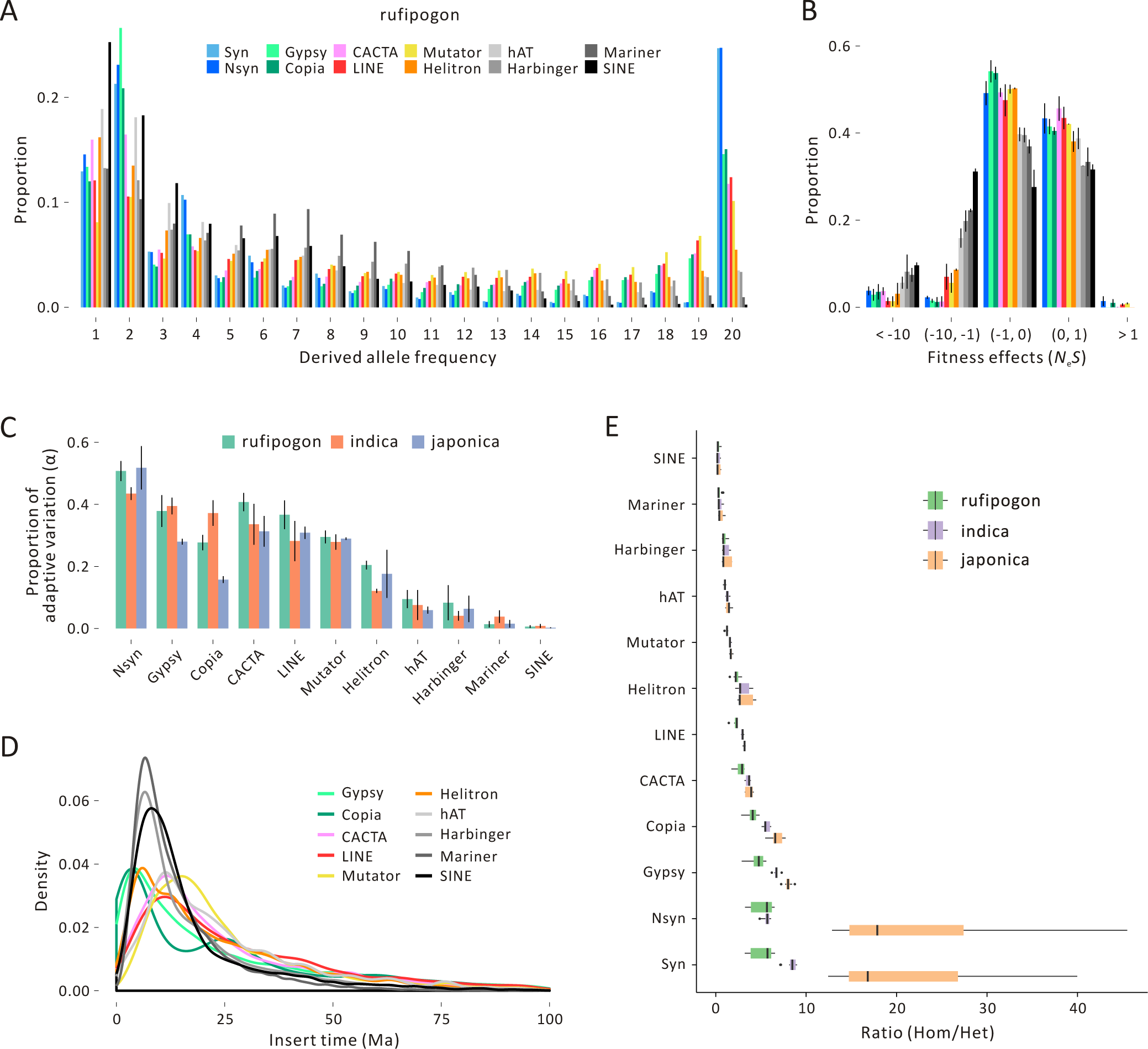
Features of TE diversity in rice. Plot (a) provides the SFS for ten element types along with synonymous SNPs (Syn) and nonsynonymous SNPs (Nsyn). This plot is for rufipogon; analogous plots for japonica and indica are provided in **Figure S8**. (b) The inferred distribution of fitness effects (DFE) in rufipogon relative to nSNPs. The y-axis provides the proportion of TE insertions, and the x-axis reports *N_e_s.* The color scheme for TE families is the same as (a). (c) The estimated proportion of adaptive variation (α) for each TE family and each of the three taxonomic groups. (d) Distributions of inferred insertion times for TE families in the Nipponbare reference. (e) The ratio of homozygous to heterozygous MEI variants in the three taxa for each TE family, which shows that the families under strong selection have relatively homozygous variants.

What might cause apparent differences in population dynamics among TE families? One explanation concerns the timing of TE insertion; if SINEs have been active more recently, it is possible that the SFS reflects a lack of sufficient time for insertions to reach fixation. To test this idea, we estimated the insertion time of individual elements within the japonica reference, producing a distribution of insertion times for all ten families (Fig. 3d). The distribution of insertion times was similar among TE families (t-tests, *P* > 0.05) with *Gypsy* and *Copia* elements (but not SINEs) biased toward slightly more recent insertions. Hence, more recent activity does not seem to be an adequate explanation for the SFS of SINE and *Mariner* elements. Another explanation is stronger purifying selection against some element families. To assess this idea, we examined the distribution of MEIs relative to genes in the japonica reference. We found that a lower proportion of SINE, *Mariner*, *Harbinger*, and *hAT* MEIs were inserted within exons relative to other TE families (**Fig. S8**). This observation could be fueled by insertion biases, but they may also point to stronger selection against these four families. Consistent with the latter interpretation, the ratio of homozygous to heterozygous variants was lower for these four families than for the other families (Fig. 3e), suggesting stronger selection when these elements are uncovered from a heterozygous state and experience recessive selection.

### SVs and the domestication process

Given insights into the frequencies of SVs in Asian rice and rufipogon, we examined SV population frequencies relative to rice domestication. Domestication repatterns population frequencies due to both positive selection on agronomic traits and enhanced genetic drift during a domestication bottleneck (41), which can drive an increased load of deleterious variants (42, 61, 62). These features of rice domestication have been investigated thoroughly with SNPs but not with SVs.

#### SVs and the cost of domestication

Because the SFS of SVs imply they are deleterious, we predicted that they contribute to the deleterious load, reflecting the cost of domestication in rice (42–44). We evaluated cost by calculating the additive SV burden per individual, which is the number of derived heterozygous sites (the heterozygote burden) plus two times the number of homozygous SVs (the recessive burden) (63). Comparing the additive SV burden across the three taxa, it was 35% and 25% higher on average for japonica and indica relative to rufipogon (Fig. 4a; t-test, *p* < 0.005 for both contrasts). Given the differences in mating system between cultivated and wild rice, we also expected the recessive burden to be the primary contributor to differences in the additive burden. This expectation held, because the recessive burden was >72% of the additive burden for both cultivars, but the proportion was lower (67%) for rufipogon (Fig. 4a). These patterns - i.e., higher additive and especially recessive burdens - held across SV types, with the apparent exception of DEL events (**Fig. S9**).

**Fig. 4.**
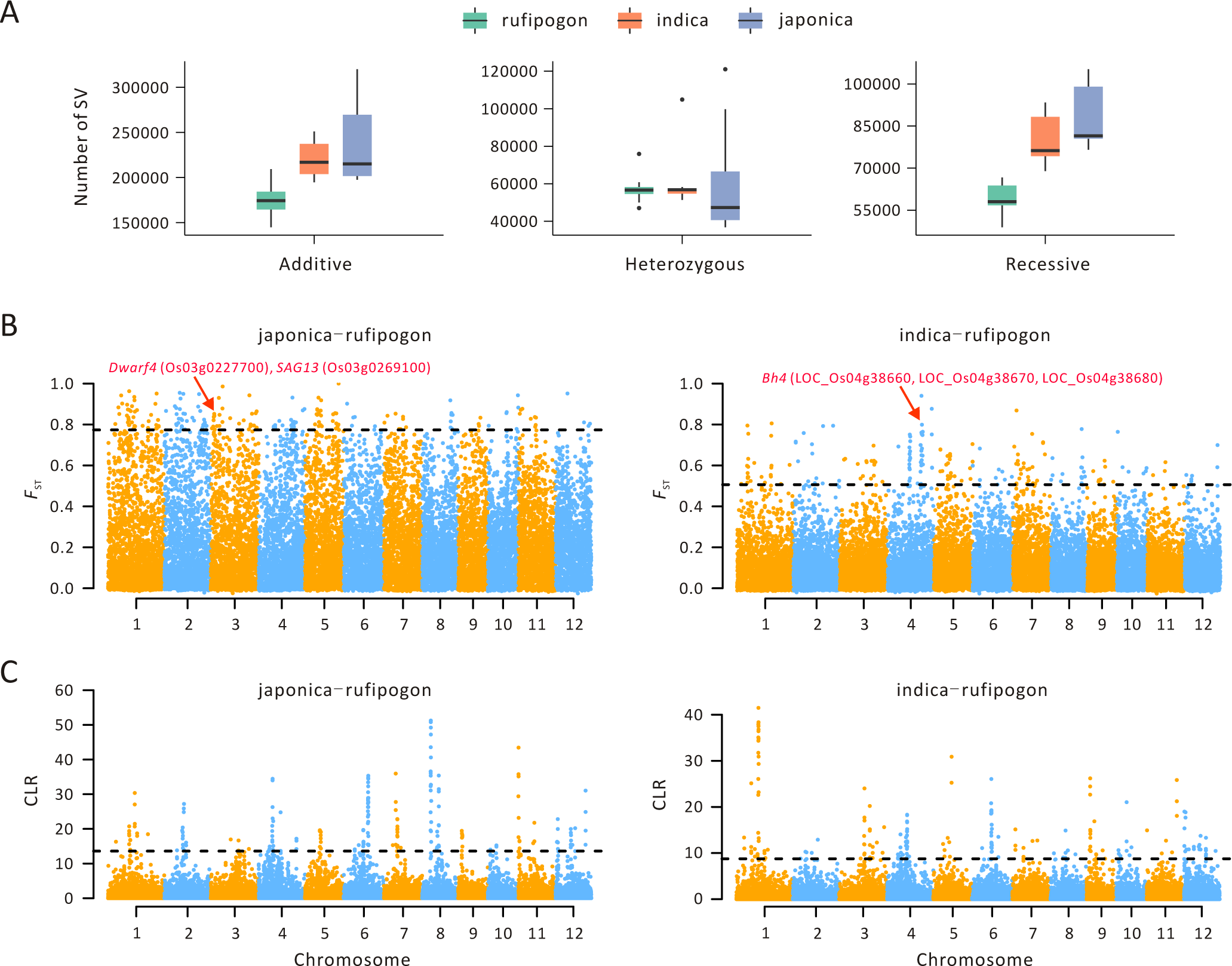
Feature of SVs associated with domestication. (a) The genetic load of SVs for rufipogon, japonica and indica. The selfing cultivars have a higher recessive (homozygous) load and correspondingly larger total SV burden, suggesting a cost of domestication. (b) Manhattan plots of F_ST_ values between indica and rufipogon, based on SVs within 20kb windows, with japonica on the left and indica on the right. The corresponding Manhattan plots for SNPs are provided in **Figure S11**. (c) Manhattan plots of CLR values for japonica (left) and rufipogon (right), based on SVs within 20kb windows. The corresponding Manhattan plots for SNPs are provided in **Figure S12**.

#### Genome-wide F_ST_ between taxa

In theory, the genes that contribute to domestication can be identified as regions of marked chromosomal divergence between wild and cultivated samples. We compared rufipogon to both indica and japonica by estimating SNP and SV divergence in fixed 20 kb windows across the genome. We calculated divergence with two measures (*F*_ST_ and *D*_*xy*_; **Fig. S10**) but focus on *F*_ST_ results here for simplicity. Across the entire genome, mean *F*_ST_ estimates were substantially higher for SNPs (indica-rufipogon 0.293 ± 0.134; japonica-rufipogon 0.485 ± 0.181) than for SVs (indica-rufipogon 0.122 ± 0.079; japonica-rufipogon 0.259 ± 0.141), reflecting the fact that SVs were typically at lower population frequencies than SNPs (Fig. 2a).

We contrasted the two cultivars to rufipogon and ranked the top 1% *F*_ST_ windows (or 187 of 18,654 windows throughout the genome) for both SNPs and SVs (Figs. 4b **& S11**). Only a small number of *F*_ST_ windows were within the top 1% for *both* SNPs and SVs (**Fig. S12**); for example, we detected 26 such windows for the indica-rufipogon comparison. Although a small number, 26 is far more than the ~2 windows expected at random (permutation; *p*<10^−3^), suggesting that the SNP and SV data do capture some common signatures (as is expected, given that they are in the same genomic window). Similarly, we detected 12 such windows in the japonica-rufipogon comparison, again representing an enrichment over a random draw (permutation; *p*<10^−3^). Of these, only one window overlapped between japonica and indica; it contained a gene (LOC_Os02g43800) that was annotated as a retrotransposon protein. Thus, arguably the strongest signal of positive selection based on *F*_ST_ - i.e., in a window identified from both SNPs and SVs across both cultivated taxa - contained a gene without obvious agronomic implications.

We examined the shared SNP-SV windows for potential candidate genes (**Table S5**). For example, of the 82 genes contained in the 26 windows for the indica-rufipogon comparison, 31 were annotated as expressed or hypothetical proteins, and 14 were annotated as TE-related; neither category were obvious candidates to contribute to agronomic phenotypes. Most of the remaining 37 genes were assigned putative functions, including a ribosomal protein, a male sterility protein, small auxin up-regulated genes, receptor like-kinase genes, and genes with other functions. GO analyses indicated that the 82 genes were enriched for a variety of putative functions, including cellular components extrinsic to membranes and biological processes related to superoxides (**Table S6**). Similarly, the shared SNP-SV peaks between japonica and rufipogon contained 36 genes in 12 windows, with 15 genes assigned putative functions and GO enrichment in DNA replication and other functions. For completeness, we also analyzed the set of genes identified in SNP-only (489) or SV-only (374) peaks for each taxon (**Table S5**). Not surprisingly, the genes were enriched for a variety of GO-based functions (**Table S6**).

We also took a candidate gene approach to assess whether SVs enhanced their identification. To do this, we focused on a set of 15 known domestication and improvement genes (12, 20) to ascertain whether they were identified in *F*_ST_ scans more often than expected at random (Table 2). Among the 15, six were within the top 1% of Fst windows between rufipogon and either indica or japonica (Table 2); three of these were found with SNPs alone (including *TAC1*, a gene implicated in tillering, and shattering genes *Sh1* and *Sh4*) and three more with SVs alone (*Bh4*, *Dwarf4* and *SAG13*) (Table 2). Overall, the set of 15 genes was highly enriched to be within *F*_ST_ peaks at the 1% and 10% levels for both SNPs (Wilcoxson-Mann-Whitney; p < 1.19 × 10^−15^ for 1% peaks, p< 6.67 × 10^−8^ for 10% peaks) and SVs (Wilcoxson-Mann-Whitney; p < 2.20 × 10^−16^ for 1% peaks, p< 2.54 × 10^−14^ for 10% peaks).

**Table 2:** Previously identified putative domestication or improvement genes with their ID, function and % ranking in Fst windows. F_ST_(i-r) refers to F_ST_ between indica and rufipogon, with additional columns for japonica-rufipogon (j-r). Cells are colored as to whether the gene is in a 1% peak (dark grey), a 10% peak (medium gray), or an extreme valley (≥90%) (light grey) based on either SNPs or SVs.

| Locus | MSU or RAP ID | Function | $F_{ST}(i-r)$ | | $F_{ST}(j-r)$ | |
| --- | --- | --- | --- | --- | --- | --- |
|  |  |  | SNPs | SVs | SNPs | SVs |
| Sh4 | LOC_Os04g57530 | Shattering | 0.18 | 17.05 | 5.08 | 81.02 |
| qSW5 | Os05g0187500 | Grain width | 97.09 | 1.44 | 84.78 | 68 |
| qSH1 | LOC_Os01g62920 | Shattering | 58.21 | 94.51 | 5.06 | 96.65 |
| Sd1 | LOC_Os01g66100 | Semi-dwarfing | 31.78 | 51.96 | 5.64 | 5.99 |
| Wx | Os06g0133000 | Grain quality | 32.07 | 42.17 | 27.02 | 58.23 |
| Badh2.1 | LOC_Os04g39020 | Flavor or fragrance | 84.57 | 70.14 | 7.46 | 64.06 |
| Rc | LOC_Os07g11020 | Red pericarp | 47.21 | 52.79 | 7.01 | 55.86 |
| Bh4 | LOC_Os04g38660,<br>LOC_Os04g38670,<br>LOC_Os04g38680 | Hull color | 3.18 | 0.95 | 6.51 | 4.67 |
| PROG1 | LOC_Os07g05900 | Plant Architecture, tiller angle | 1.79 | 2.16 | 14.18 | 6.17 |
| OsCl | Os06g0205100 | Leaf sheath color & apiculus color | 26.58 | 75.83 | 14.01 | 73.61 |
| TAC1 | Os09g0529300 | Tiller angle | 77.05 | 31.3 | 0.85 | 54.12 |
| Dwarf4 | Os03g0227700 | Leaf architecture | 9.2 | 80.98 | 1.53 | 0.35 |
| SAG13 | Os03g0269100 | Senescence | 48.3 | 47.08 | 30.39 | 0.56 |
| TB1 | LOC_Os03g49880 | Tillering | 2.22 | 81.45 | 5.23 | 78.2 |
| Sh1 | LOC_Os03g44710 | Shattering | 0.8 | 4.46 | 23.75 | 18.3 |

#### Selective Sweeps

Instead of relying on divergence between populations, we also searched for population-specific signals of selective sweeps using the composite likelihood method (64). We again investigated the top 1% of 20kb windows for both SVs and SNPs (Fig. 4c **& S13**). Qualitatively the results exhibited fewer but clearer peaks compared to *F*_ST_ (Fig. 4b) or *D*_*xy*_ analyses (**Fig. S10**). However, there was little correspondence between the top 1% windows based on SVs and SNPs (**Fig. S14**); the two data types shared two windows in common for rufipogon, only one for indica, and ten for japonica, which is the only taxon that had shared windows more often than expected by chance (permutation; *p*<10^−3^). None of these windows contained any obvious candidate genes (**Table S7**) or the previously identified set of 15 domestication/improvement genes. For completeness, we have listed all of the genes within CLR peaks (**Table S7**) with their GO enrichment categories (**Table S8**).

#### Gene Gain and Loss

The identification of genes gained and lost during domestication is an area of growing emphasis, because such genes may contribute to agronomic phenotypes (27, 65). We focused on the subset of SVs that included genes and determined whether they were private (i.e. variable within only one taxon) or fixed (i.e., in alternative states) between taxa. For example, between rufipogon and japonica we detected 114,840 SVs that were private in rufipogon; 144,600 that were private in japonica; 180,798 that were shared SVs between taxa; and only one fixed SV, corresponding to one gene that was annotated as related to retrotransposition. Focusing on the subset of SVs that include genic DUP and DEL events, we found 148 genes gained and 138 lost in our japonica sample relative to the rufipogon sample. Similarly, we detected 3,410 genes gained and 181 lost in indica relative to rufipogon (**Table S9**). Some of the genes lost during domestication had validated functions related to physiological and morphological traits, such as sterility (LOC_Os01g11054), culm leaf (LOC_Os04g39780), flowering (LOC_Os03g05680), and tolerance (LOC_Os01g64970). In addition to these functions, genes inferred to be gained during domestication involved functions that contribute to eating quality, including starch storage and biosynthesis (*e.g.*, LOC_Os01g65670, LOC_Os02g32350, LOC_Os03g09250, LOC_Os03g49350, LOC_Os03g52340, and LOC_Os09g26880) (20). A complete list of private genes and their GO enrichments are listed in **Tables S9 & S10**; the important point is that SV analyses between Asian rice and its wild relative yield reasonable candidate genes for traits involved in domestication or improvement.

## DISCUSSION

Structural variants (SV) are prevalent in plant genomes. They explain substantial trait variation in the few cases they have been included in GWAS analyses, and they contribute to important agronomic phenotypes (27). Yet, they remain unexplored for most crops, and this is particularly true with respect to their population frequencies in crops and their wild relatives. Here we have performed a genome-wide analysis of SVs in Asian rice and its wild progenitor *O. rufipogon*, with the goal of understanding more about the evolutionary processes that act on them and their fate during domestication.

Most of our inferences are based on SVs called from a large (358 accession) dataset consisting of high-coverage (average 51.6x, median 28x), short-read data. Given these calls, it is important to note two important caveats. First, we have focused on SVs that were useful for population genetic inference, meaning that the SVs were filtered both for quality and to avoid complex events, such as overlapping SVs. Given this curation, it is important to convey that we did not expect, nor intend, our set of SVs to represent a comprehensive catalog, as was reported for previously for 3,000 rice genomes without rufipogon (26). Second, all of our inferences rely on mapping to the Nipponbare genome, which may introduce a reference bias that may make it more difficult to identify novel SVs from non-japonica samples. Nonetheless, the SVs from short-read data do provide population structure information that is highly concordant to (and strongly correlated with) information from SNPs (Fig. 1b), suggesting their suitability for population analyses.

### SVs across datasets

We compared SVs from short-read data to SVs inferred from two additional data types: whole genome alignment and SMRT read analysis. The different SV datasets clearly have different advantages and disadvantages. For example, genome alignments can be poor for detecting heterozygous SVs, because primary assemblies ignore alternative haplotypes (66); they tend to be best at identifying large SV events, such as large insertions (67). In contrast, SMRT-read mapping should be efficient for most SV detection, outperforming short-read data (58, 68). However, the development of SMRT-read methods is still nascent (66) and by no means perfected, as perhaps evidenced by the very low number of DUP events that we detected with SMRT reads (**Table S3**). Finally, paired-end short-reads are accurate for SV calls, given sufficient coverage (47), but they often miss large and complex SV events (58). Given the strength and limitations of different approaches, our comparative data sets provide different insights into rice SVs.

For example, whole-genome alignments indicate the presence of a large centromeric inversion on chromosome 6 that differentiates indica from japonica and the outgroups in our sample (Fig. 1c). This inversion has been reported previously to contain 404 genes (69), and it was also identified in *O. brachyantha* compared to indica assemblies (70). Another SV feature is genic hemizygosity, which can be estimated reasonably from SMRT reads (35). Based on remapping SMRT reads to the Nipponbare reference, we have inferred the number and percentage of hemizygous genes in an indica (93-11), japonica (Nipponbare) and rufipogon individual based on split reads. Within Nipponbare and 93-11, we estimate that <1% of genes are hemizygous, which is consistent with the expectation of high homozygosity for predominantly selfed lineages. These estimates are low enough that they may reflect the false-positive rate of the method. In contrast, ~9% of genes are hemizygous for the *O. longistaminata* individual. Superficially 9% seems high, but it is similar to the ~10% PAV differences between inbred lines of maize (72) and lower than the ~10 to 14% genic hemizygosity of grapes (35, 73). This observation adds to a growing appreciation that outcrossed plants harbor a substantial fraction of SVs that lead to gene hemizygosity.

Our filtering of short-read SVs was (purposefully) biased against the detection of complex, overlapping SV regions, because we sought to identify discrete loci for population genetic analysis. It is worth emphasizing, however, that complex SV regions do exist. For example, we used genome alignments to investigate SVs in the region surrounding one of the genes (*LOC_Os01g05600*, MSU7) that was detected as gained during domestication, based on short-read SVs. The gene is a member of the NBS-LRR family, which is known for copy number turnover and as targets of positive selection (74). In this region the cultivated individuals in our sample have an additional NBS-LRR gene relative to the wild individuals; among cultivars, the region is marked by the movement of both DNA and RNA transposons that alter distances among genes (Fig. 5a). Similarly, we examined the *Submergence1* (*SUB1*) region (Fig. 5b), which contains CNVs that affect flooding tolerance (75, 76). This locus contains a cluster of three ethylene response factor (ERF) genes, *Sub1a*, *Sub1b* and *Sub1c*. Among them, only the *Sub1a-1* (a allele of the *Sub1a* gene) confers flooding tolerance and it was found to only present in a few indica cultivars (Fig. 5b). Among the cultivars investigated, the region has expanded nearly 2-fold in 93-11 and Tetep due to transposon element insertions and a genic copy number variant (*Sub1a-2*). Altogether, these analyses accentuate the prevalence of SVs in rice (20, 26) and the fluidity of *Oryza* genomes (18, 77).

**Fig. 5.**
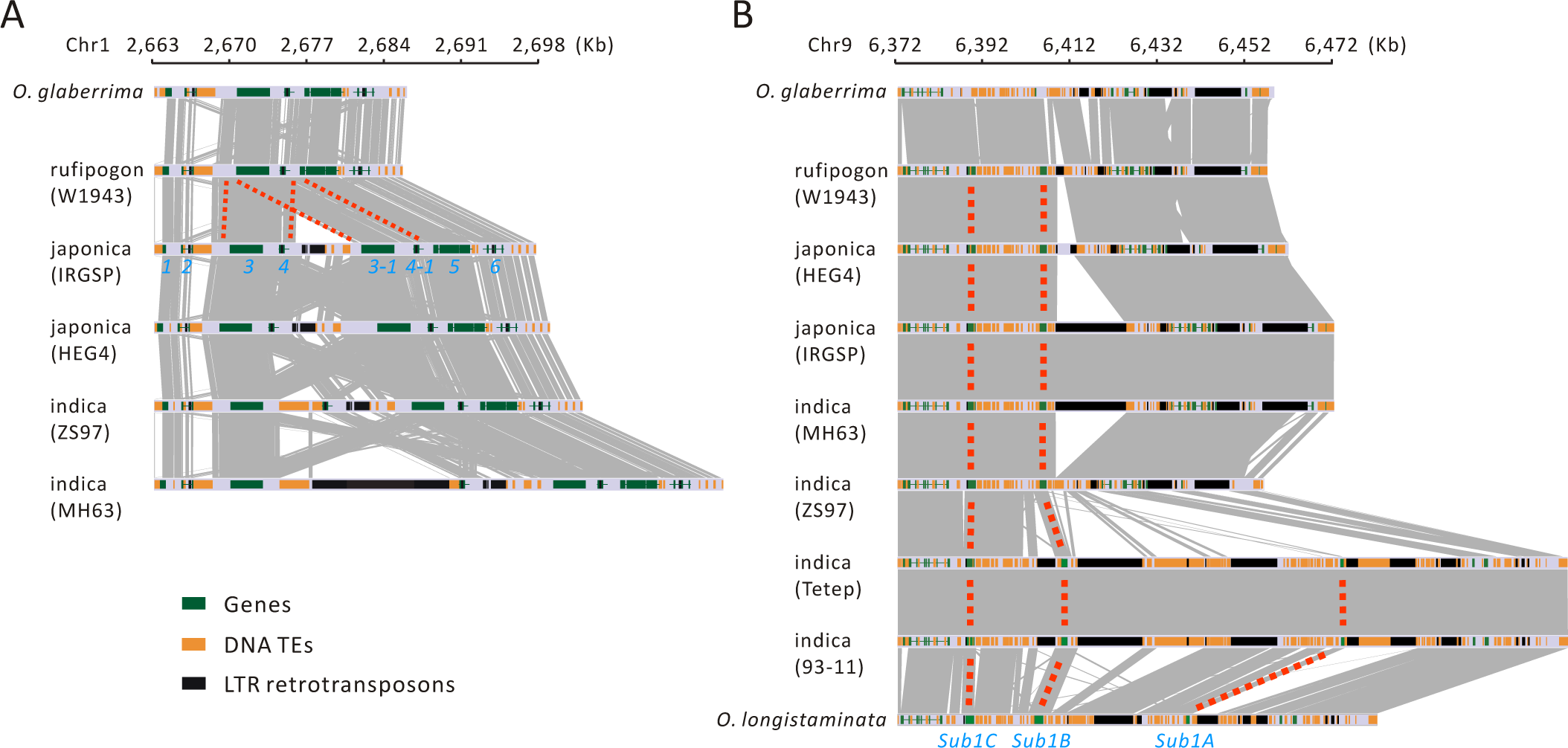
Two example regions of complex SVs across *Oryza* taxa. (a) A segmental tandem duplication of an NBS-LRR-encoding gene (*LOC_Os01g05600*, MSU7) that was found to be gained in indica and japonica relative to rufipogon. The synteny map is shown for a region corresponding to a 35-kb region in japonica (Nipponbare). (b) Gene and TE copy number variation in a 100 kb region of chromosome 9 that includes the *Sub1A* gene, which confers flooding tolerance. Both an indica accession and O. longistaminata contain three copies of genes. For both (a) and (b) gene copies are tracked by dotted red lines.

### SVs tend to be deleterious

For population genetic analyses, we focused on population samples of rufipogon, japonica and indica. We first constructed the SFS for SVs and compared them to sSNPs and nSNPs. Both SVs and SNPs recovered the well-documented ‘U-shaped’ SFS for Asian rice (Fig. 2cd), for which the fixed variants reflect the effects of the strong domestication bottleneck (13). The observed U-shape for rufipogon has been hinted at previously but to our knowledge has not been this pronounced (Fig. 2b). Our U-shaped SFS for rufipogon probably reflects two features of our sample and analyses. First, we limited the sample to a single rufipogon clade (Fig. 1a), whereas many and perhaps most previous studies combined rufipogon groups, which will tend to underestimate the fixed class of variants. Second, we used several outgroups to infer derived states, which improves accuracy (78). Note, however, that the U-shape is more right-leaning for the two cultivated rice varieties than for rufipogon (Fig. 2b-d), again suggesting that domestication and/or the shift in mating system led to the accelerated fixation of derived variants. Previous work has shown that plant SVs tend to be deleterious (27, 35, 65, 79), and the SFS of SVs support that view. In each of the three taxa, the SFS of each SV type differs significantly from the SFS of “neutral” sSNPs. However, the SFSs also suggests heterogeneity among SVs, because they follow an apparent hierarchy in which INV events are most deleterious, followed by MEI, TRA, DEL and DUP events. One pertinent question is whether detection biases somehow fuel this heterogeneity because, for example, INV events have a lower percentage of overlap among datasets than DEL and DUP events (Fig. 1d **& S3**). There are systematic biases for all SV types - e.g., ascertaiment biases affect the SFS, as to false negative and false positives - that tend to skew the SFS leftward (Emerson et al. 2008). The question here is whether INVs are particularly prone to these biases. We do not believe that this is the case for three reasons: *i*) the SFS are based on the individuals with highest coverage, which limits false negative results (Cridland and Thornton 2010); *ii*) SMRT-read analyses also indicate that INV events are found at low frequency, because 86% of INVs are found in only one indica, and this number is significantly higher than for other SV types (*P* < 0.05); *iii*) some studies suggest similar false discovery rate across SV types (47). Interestingly, the hierarchy of SV types differs somewhat for grapevines (35), where INV events were again inferred to be the most deleterious but TRA events were more deleterious than MEIs.

Just as we have identified putative differences in the deleterious effects of SV types, we have also examined the SFS for potential differences in the population dynamics of MEIs from different TE families. Several element families, but particularly SINE elements, have dramatically different population frequencies than other element families. Relatively few SINEs were fixed, and most were found at low frequency. This finding does not appear to be due to more recent activity (Fig. 3d) but rather to stronger selection, as implied by the low frequency of SINEs in and near genes (**Fig. S8**) and the relative dearth of homozygous variants (Fig. 3e). This last observation supports the growing consensus that deleterious variants can accrue as heterozygotes because they are typically under recessive selection (35, 37, 80), but the mating system of rice ensures that new, heterozygous TE insertions do not remain heterozygous for long. Because SVs tend to be deleterious, they are typically found at lower population frequencies than SNPs. This fact provides challenges for GWAS analyses, for two reasons. First, if SVs commonly underlie phenotypic traits, then our population genetic results imply that GWAS assumptions about common variants do not hold. Second, GWAS based on SNPs is unlikely to anchor SVs, given low LD between SNPs and SVs (Fig. 2e). Practically, this implies that causative SVs will not be easily tagged by anchoring SNPs in rice, as has been shown in Arabidopsis (31). Finally, in the context of GWAS it is worth briefly considering the fact that domesticates have a higher burden of fixed, derived SVs (Fig. 2b-d). Fixed SVs contribute to the cost of domestication (Fig 3a), but some subset may be advantageous because they contribute to agronomic traits that differentiate domesticated rice from rufipogon. Unfortunately, these fixed variants also cannot be identified with GWAS based only on cultivated samples (81), highlighting one of the limits of cultivar-specific collections (20).

### SVs and domestication

Domestication bottlenecks accelerate genetic drift, which can contribute to a cost of domestication. An appropriate measure of cost is the average number of deleterious variants per genome (*d*_*g*_) (82). Interestingly, *d*_*g*_ is not expected to vary substantially before and after a demographic shift under some conditions, such a strict genetic bottleneck with outcrossing and additive (*h*=0.5) variants (83). However, it can vary substantially with deviations from these conditions (63). For example, clonal variants tend to accrue recessive deleterious variants over time, because these variants are under recessive selection and permanently held in a heterozygous state (37). Similarly, forward simulations have shown that moderately and weakly deleterious variants accumulate under various demographic regimes (84). Here we have shown that the SV burden is elevated by 25% to 35% in our japonica and indica samples relative to rufipogon (Fig. 3a). While the estimated increase of *d*_*g*_ undoubtedly depends on the composition of the samples under comparison, this observation is consistent with previous studies suggesting a cost of domestication in Asian rice (42–44).

Positive selection can also be pervasive during domestication (39, 41, 85). We examined the SV and SNP data for signals of positive selection throughout the genome, relying either on divergence between rufipogon or signals of selective sweeps within cultivars. For divergence, as measured by F_ST_, only a small number of windows fell within the top 1% of peaks for both SVs and SNPs, with 23 windows for japonica-rufipogon and 33 for indica-rufipogon. Although small, the numbers are enriched relative to random expectation. None contain obvious candidate genes, at least under our examination, but the lists of genes within *F*_ST_ peaks may prove useful for researchers studying domestication and improvement traits in indica and japonica (**Tables S5 & S6**). The shared windows between SNPs and SVs also do not include any of our set of 15 domestication genes (Table 2), but these genes are enriched significantly in high-ranking F_ST_ peaks when SNPs and SVs are considered separately, which lends some credibility to the basic approach. Importantly, *F*_ST_ analyses suggest that SV calls aid the detection of divergent genomic regions, because some of the 15 genes were detected with SVs only (Table 2).

Given the consensus that domestication genes were introgressed into indica from japonica (15), a previous study remarked that domestication genes should be enriched in regions of low divergence between the two cultivars (12). We tested this notion by examining the candidate set and their corresponding window rankings in *F*_ST_ windows calculated between indica and japonica. None of the 15 genes is located in an *F*_ST_ trough, as defined by windows ranking in the lowest 99% percentile, but three of the genes (*Wx*, *Bh4*, and *PROG1*) had either SNP or SV values > 90% (**Table S12**), which is a significant enrichment (WMW, p < 0.05). In contrast to genes within *F*_ST_ troughs, several of the 15 genes are located in *F*_ST_ *peaks* between indica and japonica, including *Dwarf4, Sd1* and *TB1* (**Table S12**). Altogether, these analyses do support the consensus view that some domestication genes were introgressed between japonica and indica, as reflected by *F*_ST_ troughs, but still other domestication genes are in *F*_ST_ peaks, suggesting they differentiate japonica and indica and were not introgressed.

We also searched for signals of positive selection using population-specific signals of selective sweeps. One expects *a priori* that SVs have lower power to detect selection, given that there are fewer of them and that they tend to be at lower standing population frequencies than SNPs (so that the relative effect of sweeps is less readily detectable). Consistent with this premise, few windows of putative selection overlapped between SNPs and SVs, although there was a slight enrichment for overlapping windows in *japonica*. These regions again yielded no obvious candidate genes for contributing to domestication traits, but the full list of genes may prove useful (**Table S7 & S8**). In contrast, our analysis of gene gains and losses in the cultivars yielded

Overall, this study has provided unique insights into structural variants and genome evolution in *Oryza* and Asian rice. First, it provides a glimpse into some of the advantages and disadvantages of different SV-detection methods, although we were limited by publicly available data for genome alignments and long-read data. A fuller comparison will require study of the same samples with different data types and methods; such a comparison is not yet available for any crop. Second, by studying the population genetics of a highly curated set of SVs, we have shown that SVs are generally selectively deleterious, a conclusion that has been cited previously (27, 35, 65) but thus far based on few genome-wide analyses for plants. Third, just as we have suggested that different SV types have different selective effects, so do different TE families. SINE and mariner elements are especially deleterious, but the mechanisms underlying selection against them are as yet unknown. Finally, we have investigated features of rice domestication using SV markers and find that SVs can help identify domestication genes, particularly by defining regions of crop-ancestor diverges. However, SVs are particularly useful for documenting gene gains and losses during domestication. Many of the genes that we find have. SVs contribute to the cost of domestication.

## MATERIALS AND METHODS

### Data samples and pre-processing

We collected three kinds of data to detect SVs: whole-genome assemblies, PacBio SMRT reads, and paired-end short-read (Illumina) data. For the first, we downloaded 15 assemblies from previous publications (**Table S2**), including outgroups assemblies of *O. glaberrima* (55) and *O. longistaminata (56)* that were used as to infer the ancestral state of structural variants. For the second dataset, we gathered SMRT reads for ten accessions that were a subset of the whole-genome dataset (**Table S2**) and included data from 6 indica, 3 japonica, and one *O. longistaminata* accession. For the third dataset, we compiled paired-end, short-read resequencing data, requiring a minimum of 15x coverage per genome. We downloaded an initial set of genome sequences representing 404 individuals, called SNPs based on these individuals and then subjected them to population clustering using NGSadmix (86). After culling 46 individuals that were apparently misclassified (Table S1), the final dataset consisted of 358 individuals with a mean coverage of 51.6x (Table S2).

Both SMRT and Illumina reads were preprocessed. SMRT reads were extracted and filtered from h5 files using Dextractor v1.0 (https://github.com/thegenemyers/DEXTRACTOR) with a minimum length 1000 and a minimum quality score 750. Paired-end Illumina reads were trimmed to remove adapters and low quality bases (< 20) and filtered for reads < 40 bp using Trimmomatic 0.36 (87). The quality of raw and filtered reads was computed using FastQC 1.0.0 (https://www.bioinformatics.babraham.ac.uk/projects/fastqc/). Filtered short-reads were then mapped to the reference genome japonica Nipponbare (Zhang et al. 2018) using BWA-MEM (88). Reads with mapping qualities < 10 were filtered to remove non-uniquely mapped reads using SAMtools 1.9 (89). To account for the occurrence of PCR duplicates introduced during library construction, we used MarkDuplicates in the picard-tools v1.119 (https://github.com/broadinstitute/picard) to remove reads with identical external coordinates and insert lengths. The bam files were then sorted and indexed using samtools for downstream analyses.

### Variant calls

#### SNP calling with short-read data

SNPs and short (< 50 bp) indels were called for the entire dataset of 404 individuals using the HaplotypeCaller in GATK v4.1.2.0. SNPs were filtered using the VariationFiltration in GATK v4.1.2.0, according to the following criteria: variant quality (QD) > 2.0, quality score (QUAL) > 40.0, mapping quality (MQ) > 30.0, and < 80% missing genotypes across all samples. SNP variants were then annotated to be synonymous or nonsynonymous according to the gene annotation from MSU7 Rice Genome Annotation Project (http://rice.plantbiology.msu.edu/) (90) using the SnpEff v4.0 (91) with the structural annotation of the reference based on Maker Version 2.31.8. These SNP variants were used for population assignment and inference but not for estimates of nucleotide diversity (see below).

#### SV discovery

We used separate methods to infer SVs in the three different datasets. For genome alignment, we performed comparisons between pairs of assemblies with MUMmer v4 (Marçais et al. 2018). The minimum length of a single exact match (-l 1000) and a cluster of matches (-c 1000), and the maximum diagonal difference between two adjacent anchors in a cluster (-D 5) were set using the nucmer program (nucmer -maxmatch -noextend). Dot plots were generated to visualize chromosomal collinearity and large SVs, using mummerplot. SVs between pairs of assemblies were discovered using the show-diff program in MUMmer v4; collinear regions and SV breakpoints were shown using the show-coords program. The chromosome 6 inversion and other major SVs were verified in the IGV browser (92) from bam files mapped the SMRT and/or Illumina reads of both accessions to both assemblies. Genome assembly based SV discovery was performed using the pipeline previously described (77). Briefly, soft masked target and query genomes were first aligned using LASTZ (54, 93), and then processed with CHAIN/NET/NETSYNTENY tools (94) to construct the syntenic blocks. SV calling and further filtering were processed using custom perl scripts which are available at https://github.com/yiliao1022/LASTZ_SV_pipeline.

To infer SVs from SMRT reads, we mapped SMRT reads from each accession to the Nipponbare reference genome (Zhang et al. 2018) using minimap2 V2.15 (57). The population calling model of the Sniffles pipeline (58) was used to genotype SV across all ten accessions (Table S4). The SVs calls were then filtered following (35) by removing SVs with: *i*) flags “IMPRECISE” and “UNRESOLVED”, *ii*) length < 50 bp, and iii) support by fewer than four SMRT reads.

SV calls from paired-end Illumina reads were based on two methods: DELLY2 (50) and LUMPY 0.2.13 (47). Both programs were used to call and genotype SVs across the 358 accessions as a single sample. Technically, this means that SVs were called with ~18,000 X coverage of the reference genome. For DELLY, SV calling was performed with the recommended workflow (50). For LUMPY, read lengths and insert sizes were extracted from bam files for each sample using SAMtools 1.9 (89), and the SVs were genotyped using SVTyper (47). Both DELLY and LUMPY SV calls were filtered following Zhou et al. (35). SV calls from DELLY and LUMPY were merged using SURVIVOR v1.0.3 (95). We excluded SVs that overlapped existing TE annotations, based on the RepeatMasker Version 1.332 (http://www.repeatmasker.org) output using a curated rice TE library (96). The final SV calls were filtered with the additional criteria, including length > 50 bp, missing genotype < 80% and identical breakpoints across all 358 individuals.

#### Mobile element insertions (MEIs)

We used separate approaches to examine MEIs, because these are often large enough that they are called incorrectly by short-read SV callers. Insertion frequencies of mobile elements in population samples were detected using PoPoolationTE2 (51) using Illumina PE reads across the 358 accessions using four steps. First, the sequences of all TEs with length > 50bp were extracted from an existing TE annotation across all major TE families of reference genome japonica Nipponbare (46). These TE regions were then masked in the reference. The TE-merged-reference was generated by merging TE sequences and the masked reference genome (97). Second, illumina PE reads were mapped to the TE-merged-reference using BWA-MEM (88). Third, the insertion frequencies of TEs across population samples were identified, using the recommended workflow of PoPoolationTE2 with the joint algorithm and default parameters. Finally, the MEIs were genotyped for each individual based on the number of supporting reads; an MEI or non-MEI allele were genotyped as missing when there were < 4 reads at the breakpoints that support either allele. An MEI supported by < 4 reads were genotyped as 0/0 (homozygous non MEI); an MEI supported by most of the reads (<4 reads support non-MEI) were genotyped as 1/1 (homozygous MEI); and the remaining cases were genotyped as 0/1 (heterozygous MEI).

### Population genetic analyses

Our variant calls resulted in filtered bam files, a vcf file for SNPs, a vcf for SVs and MEI genotypes based on population samples that were used in downstream evolutionary genomic analyses. The data for this paper are all publicly available, with their sources listed in Tables S1 and S2. The annotation files used in this paper, along with the unfiltered vcfs are available at http://zenodo.org/xxxx.

#### Population structure

We used the SNP vcfs to examine population structure. These analyses were performed in ANGSD v0.929 (Korneliussen et al. 2014) using genotype likelihoods in the beagle file as an input to NGSadmix. Population structure inference was based on SNP variants with a minimal quality score of 20 and a minimal mapping quality of 30. The number of genetic clusters (*K*) ranged from 2 to 10, and the maximum iteration of the EM algorithm was set to 2,000. The filtered SNPs across all samples were also used to construct a phylogenetic tree in the FastTree v2.1.11 program with GTR+CAT model (98) with *O. longistaminata* and *O. meridionalis* used as outgroups. The assignment tables from SNP and SVs were compared by flattening the matrix and calculating the Pearson correlation coefficient.

The population structure inference for SV variations were conducted using ADMIXTURE 1.3.0 with a block relaxation algorithm (Alexander et al. 2009). The termination criterion for the algorithm is to stop when the log-likelihood increases by less than 0.0001 between iterations. The binary fileset (.bed) as ADMIXTURE’s input was created from the SV vcf by PLINK 1.9.0 (Purcel et al. 2007).

#### Population genetic statistics

Linkage disequilibrium (LD) decay along physical distance was measured by the squared correlation coefficients (*r*^2^) between all pairs of SNPs, SVs and all variants (SNPs + SVs) within a physical distance of 300 Kb using PopLDdecay (99). Genome-wide genetic diversity was assessed from genotype likelihood in the ANGSD 0.929 (100), based on SNP variants. The −doSaf option was used to calculate the site allele frequency likelihood at all sites, and then the −realSFS was used to obtain a maximum likelihood estimate of the unfolded SFS using the EM algorithm (49). Population genetic statistics, including the number of segregating sites (*S*), Watterson’s θ_w_ and pairwise differences π Tajima’s *D* (Tajima 1989), and Fay and Wu’s *H* (Fay and Wu 2000) were calculated for each population group using the thetaStat program (100). Genetic diversity (π) for SNPs and SVs in each group were compared using vcftools v0.1.15 (101).

The unfolded site frequency spectrum (SFS) was calculated from the allele counts for each position using three *O. longistaminata* and three *O. meridionalis* accessions as outgroups. For japonica, indica and rufipogon, we downsampled to ten samples with the highest coverage and the least missing data to calculate the SFS for each variant type, including sSNPs that were outside outlier windows based on SweeD analyses (see below), nSNPs (Nsyn), and SVs (DEL, DUP, TRA, INV, MEI). For MEIs, we also classified them into 10 families, including four retrotransposon families (*Gypsy, Copia*, LINE, SINE) and six DNA transposon families (CACTA, *Mutator, Helitron, hAT, Harbinger, Mariner*), and calculated the SFS for each family. The number of derived alleles were calculated for each type of variant using *O. longistaminata* and *O. meridionalis* as outgroup. The genetic burden was calculated under additive (2 × homozygous variants + number of heterozygous variants) (63).

#### Selection on SVs and SNPs

SweeD v3.3.2 (64) was used to detect genomic signatures of selection in the indica, japonica and rufipogon samples, based on a sliding window size of 20 Kb. The genes underlying the outlier windows were then annotated based on the MSU annotation (90). Gene Ontology (GO) analyses for these genes were run in agriGO v2.0 (102).

We used SweeD results to define neutral sSNPs, because we assumed that sSNPs outside putative selective sweeps were neutral. The neutral sSNPs were used for calculating the SFS; the sSNP SFS was compared to the SFS of other variant types using the Kolmogorov–Smirnov test in R v3.5.1. For the SFS of individual TEs, we used the unfolded SFS to estimate DFE and α, using polyDFE v.2.0 (103). The results were presented with 95% confidence intervals obtained from the inferred discretized DFEs from 20 bootstrap datasets.

Pairwise genetic differentiation (*F*_ST_) and genetic divergence (*D*_*xy*_) for SNPs and SVs along chromosome between each pair of three groups, indica, japonica and rufipogon, were estimated using VCFtools v0.1.15 (101) and PBScan v1.0 (104) with 20 Kb fixed windows. Genes underlying the *F*_ST_ and *D*_*xy*_ outlier windows were annotated based on the MSU annotation (90). GO analyses were conducted in agriGO v2.0 (102).

To examine gene gain and loss during domestication, we identified private sites in each species, and also identified shared and fixed sites between each pair of the three groups (japonica, indica and rufipogon). The corresponding gene were inferred based on the MSU7 annotation (90). GO analyses were conducted in agriGO v2.0 (102) for each category.

#### TE analyses on the Nipponbare reference

To calculate parameters such as TE insertion time and distance from gene, we focused on the Nipponbare reference and relied on its own TE annotation (46) and gene annotation from MSU annotation (90). Given the TE annotations, a multiple alignment file was generated for each TE family using MAFFT v7.305b with FFT-NS-2 method (105, 106). A consensus sequences of each TE family was extracted from the multiple alignment, and the sequence divergence between each TE copy and the consensus sequence was calculated using EMBOSS 6.5.7.0 (107). The TE insert time for each TE copy were estimated based on the sequence divergence (d_k_) and a substitution rate 6.5 × 10^−9^ substitutions per site per year (108).

## ACKNOWLEDGMENTS

The authors would like to acknowledge the high performance cluster at UCI Irvine. BSG is supported by NSF grants 1741627 and 1655808. JJE is supported by NIH grant R01GM123303. YK was supported by National Natural Science Foundation of China XXXXXXXX. YL was supported by National Natural Science Foundation of China (31771813)

