## Supplementary file for "Evolutionary genomics of structural variation in Asian rice (*Oryza sativa*) and its wild progenitor (*O. rufipogon*)"

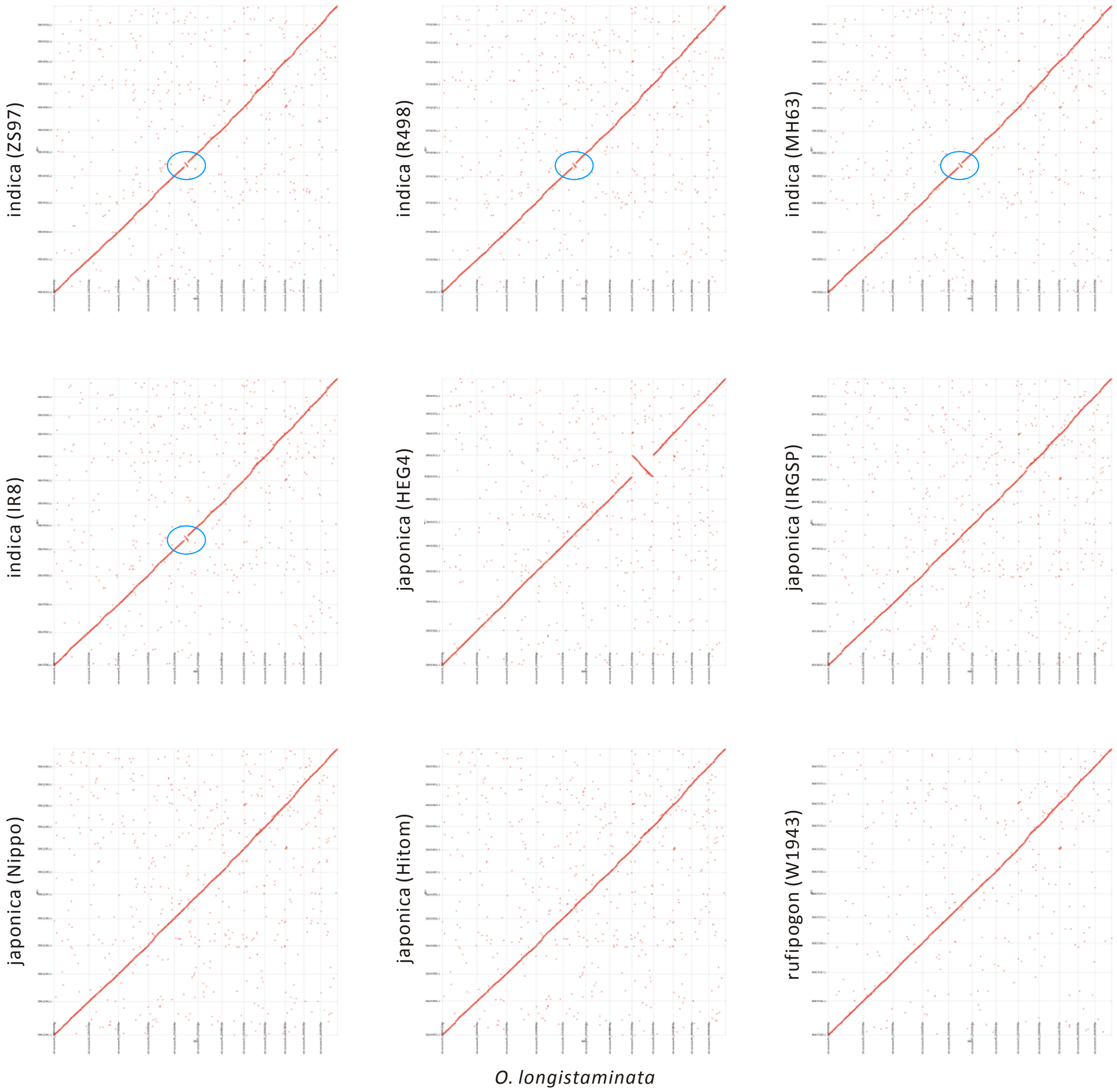

**Fig. S1.** Genome alignment of rufipogon, indica and japonica against *O. longistaminata* performed using MUMmer. A large inversion, marked using a blue circle, was found in indica accessions.

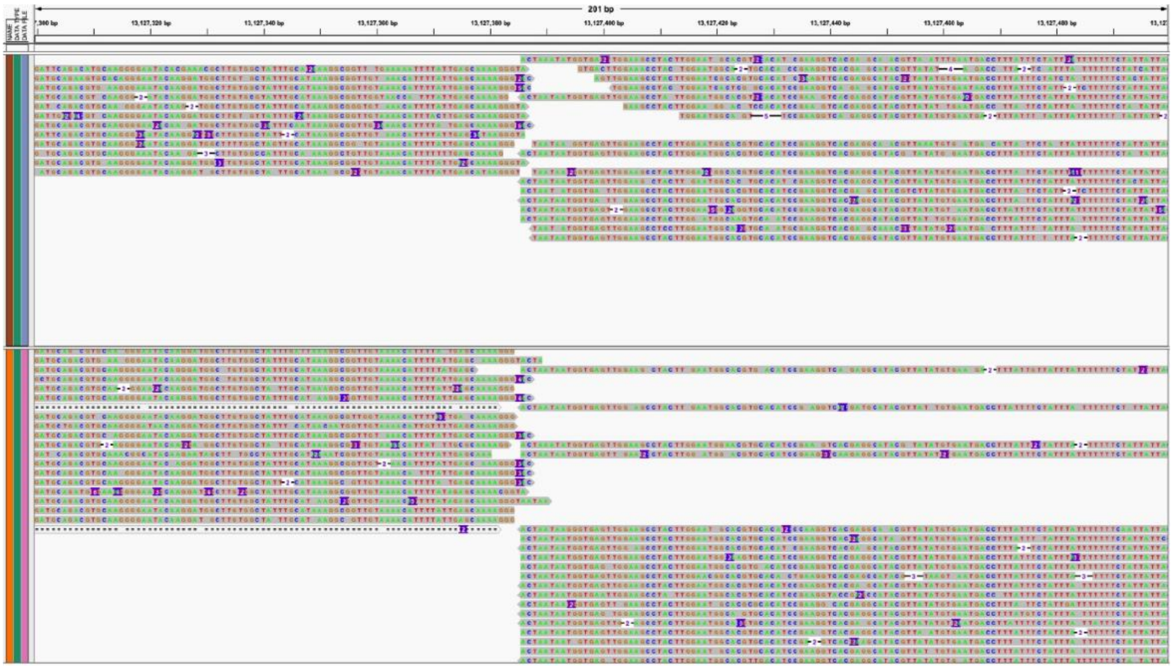

**Fig. S2.** Breakpoint of the inversion in indica identified by genome alignment against *O. longistaminata*.

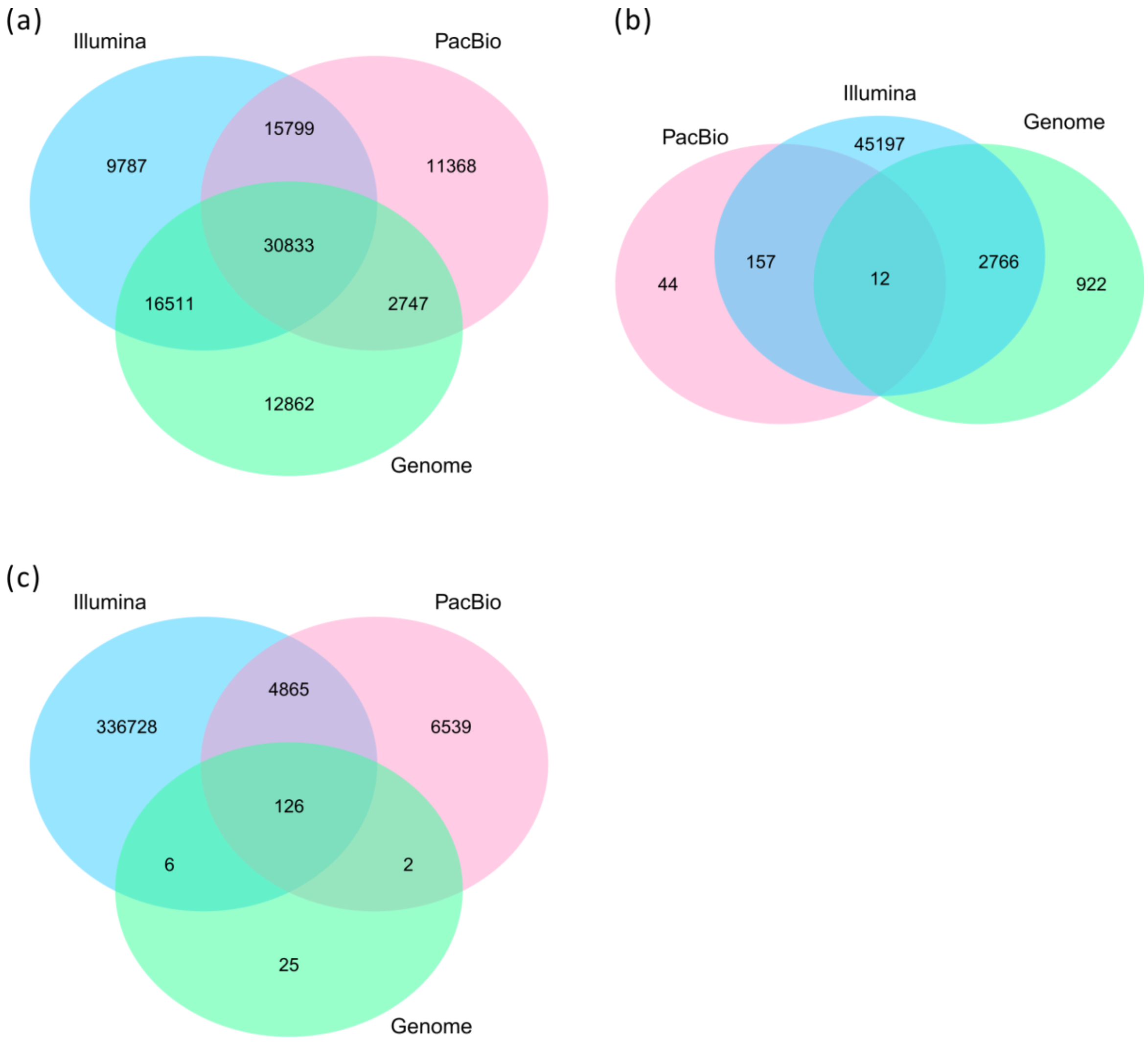

**Fig. S3.** Venn diagrams based on **(a)** DEL, **(b)** DUP, and **(c)** INV detected from Illumina, PacBio, and Genome data. See Figure 1 for the results combined across the three SV categories.

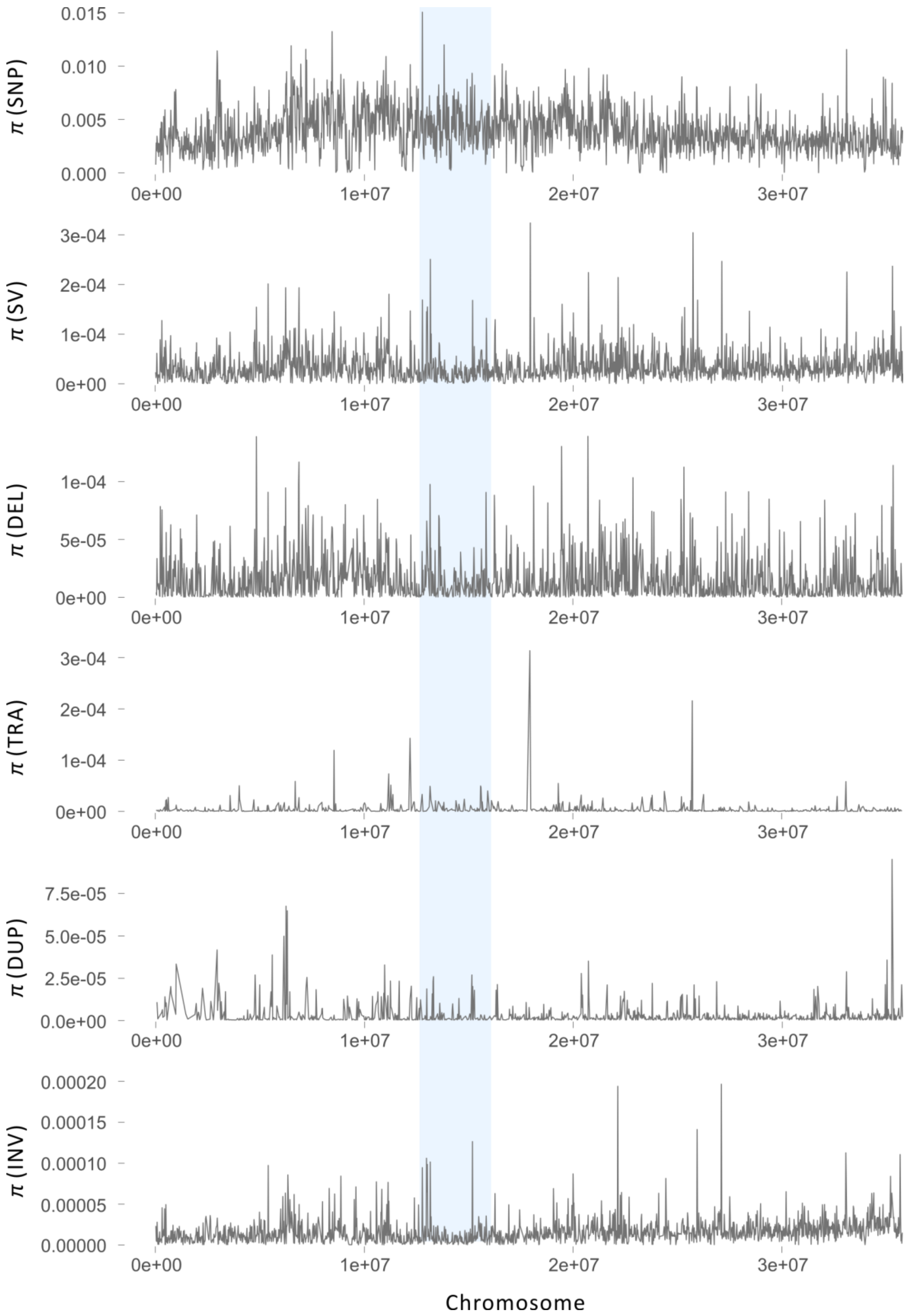

**Fig. S4.** Distribution of genetic diversity ($\pi$) for SNPs and SVs (DEL, DUP, TRA, INV) along chromosome 2 in rufipogon. Light blue bar represents the centromere.

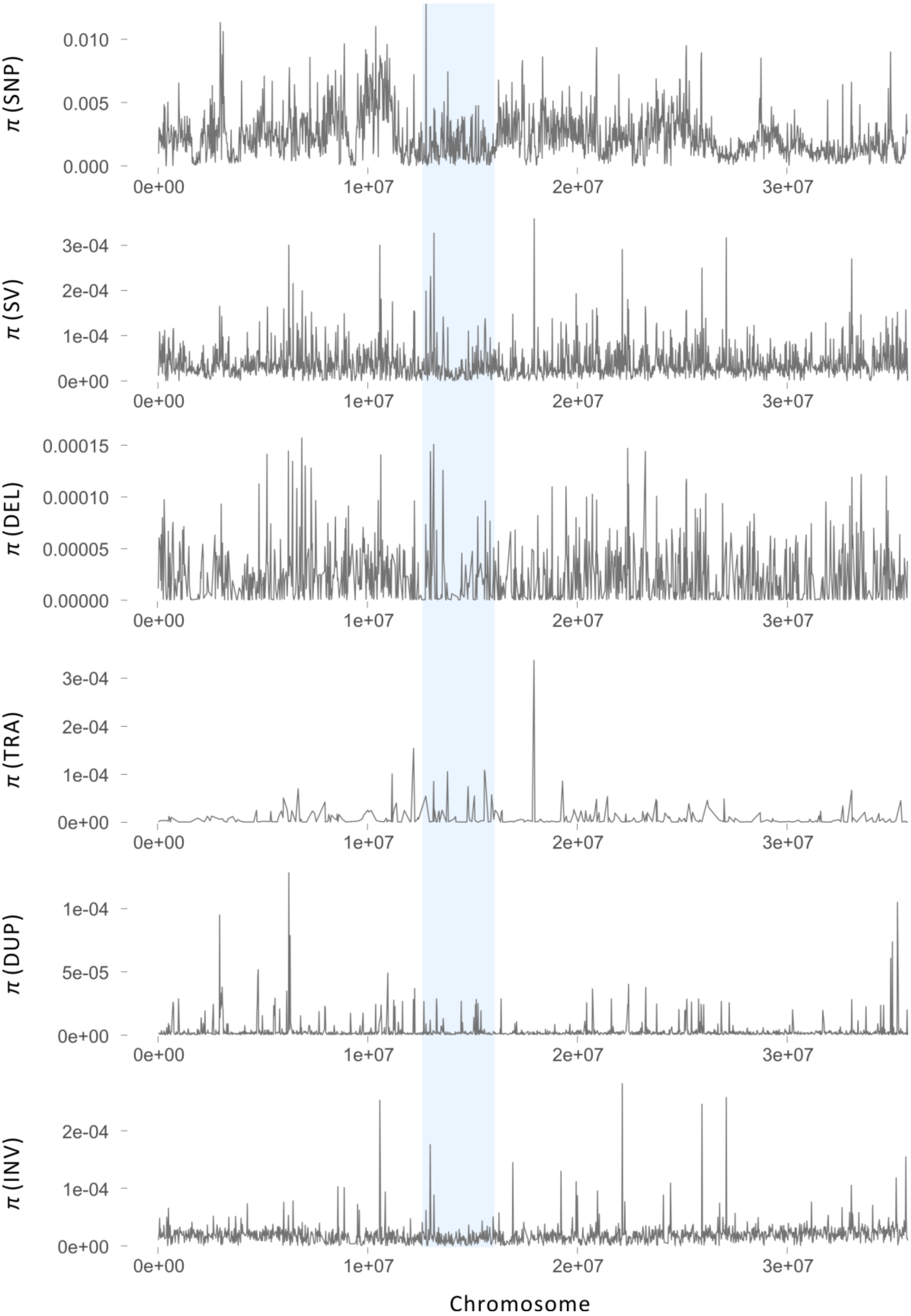

**Fig. S5.** Distribution of genetic diversity ($\pi$) for SNPs and SVs (DEL, DUP, TRA, INV) along chromosome 2 in indica. Light blue bar represent the centromere.

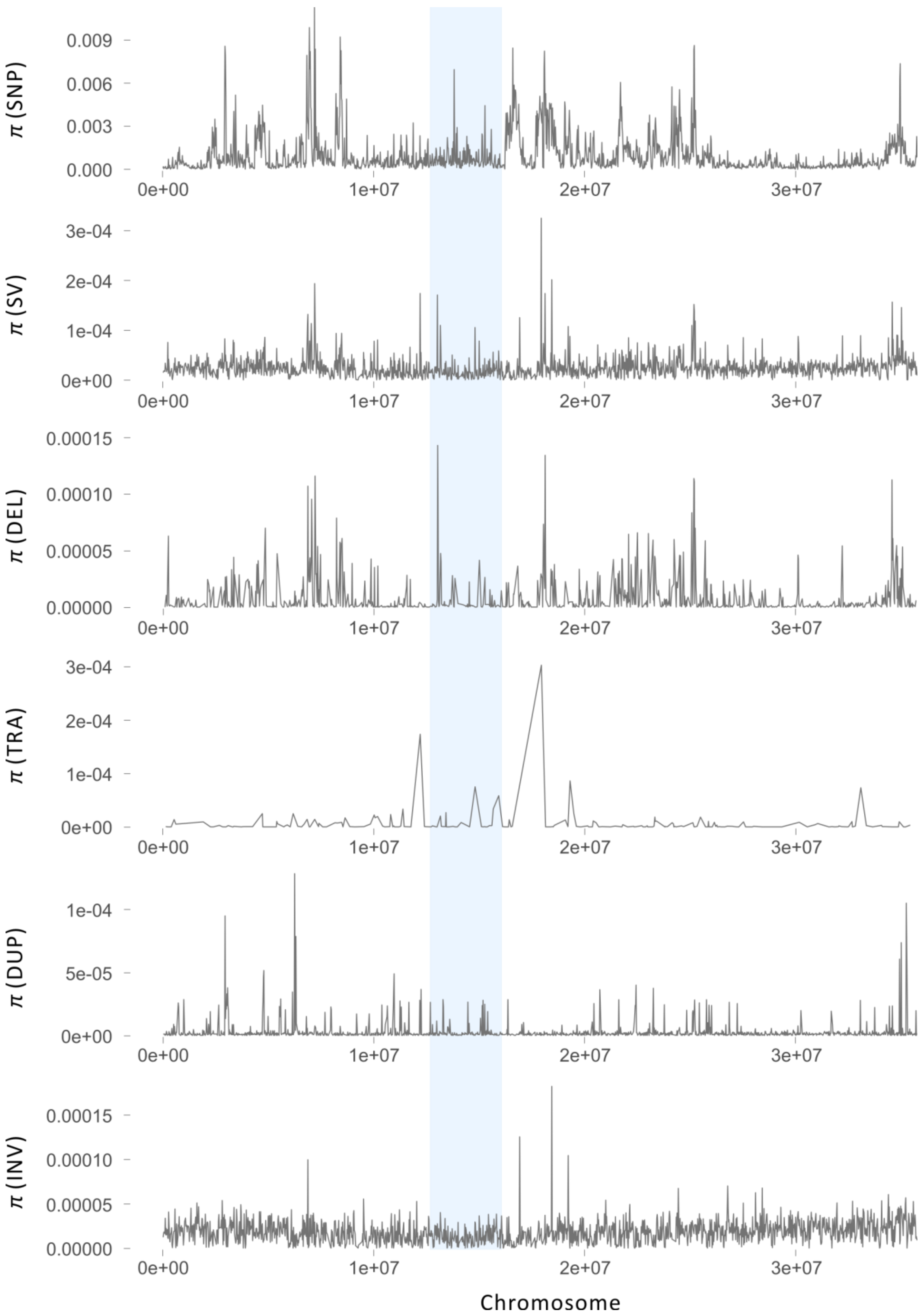

**Fig. S6.** Distribution of genetic diversity ($\pi$) for SNPs and SVs (DEL, DUP, TRA, INV) along chromosome 2 in japonica. Light blue bar represents the centromere.

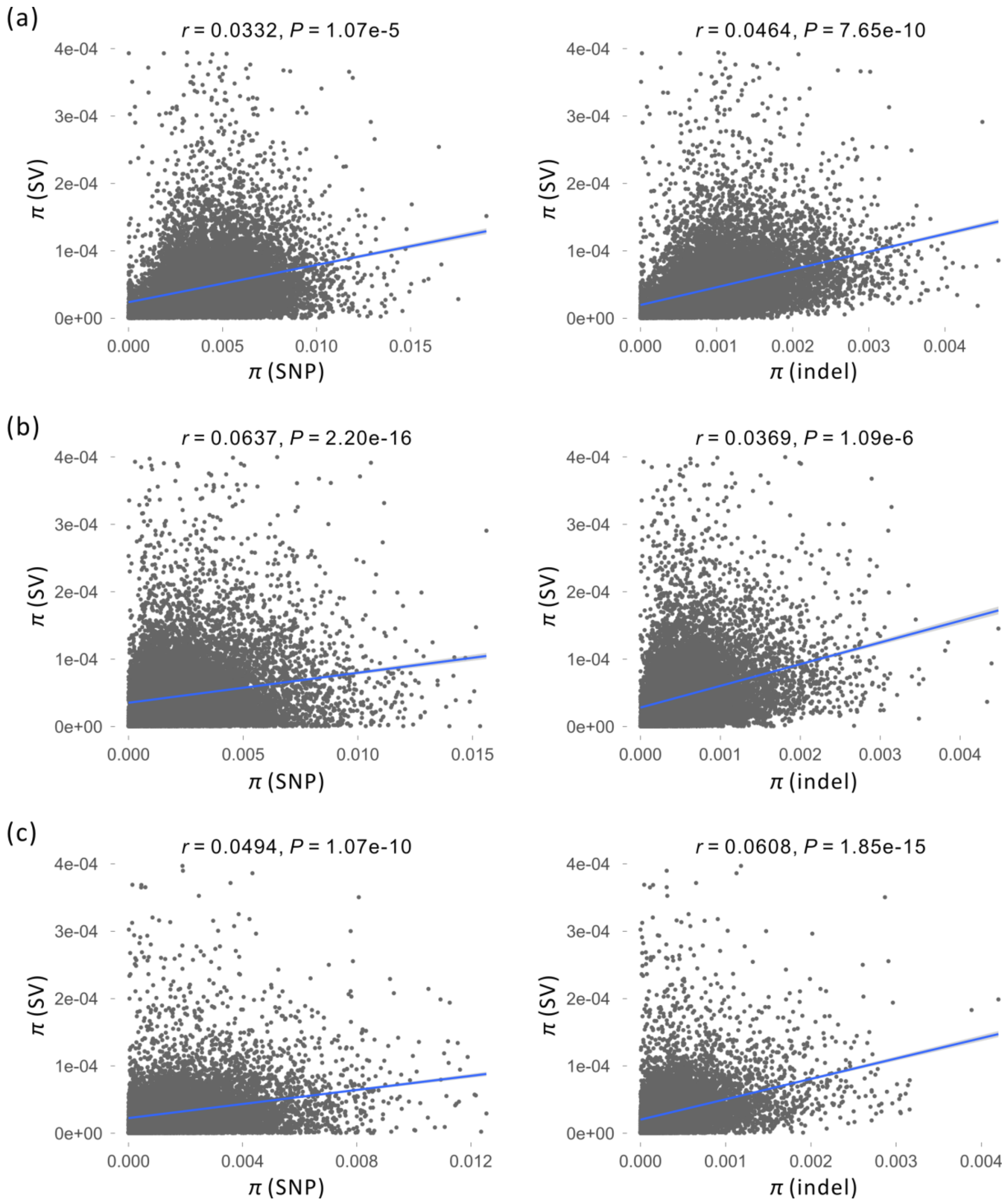

**Fig. S7.** The correlation ($r$) between genetic diversity ($\pi$) of SVs and that of SNPs and indels in rufipogon **(a)**, indica **(b)**, and japonica **(c)**.

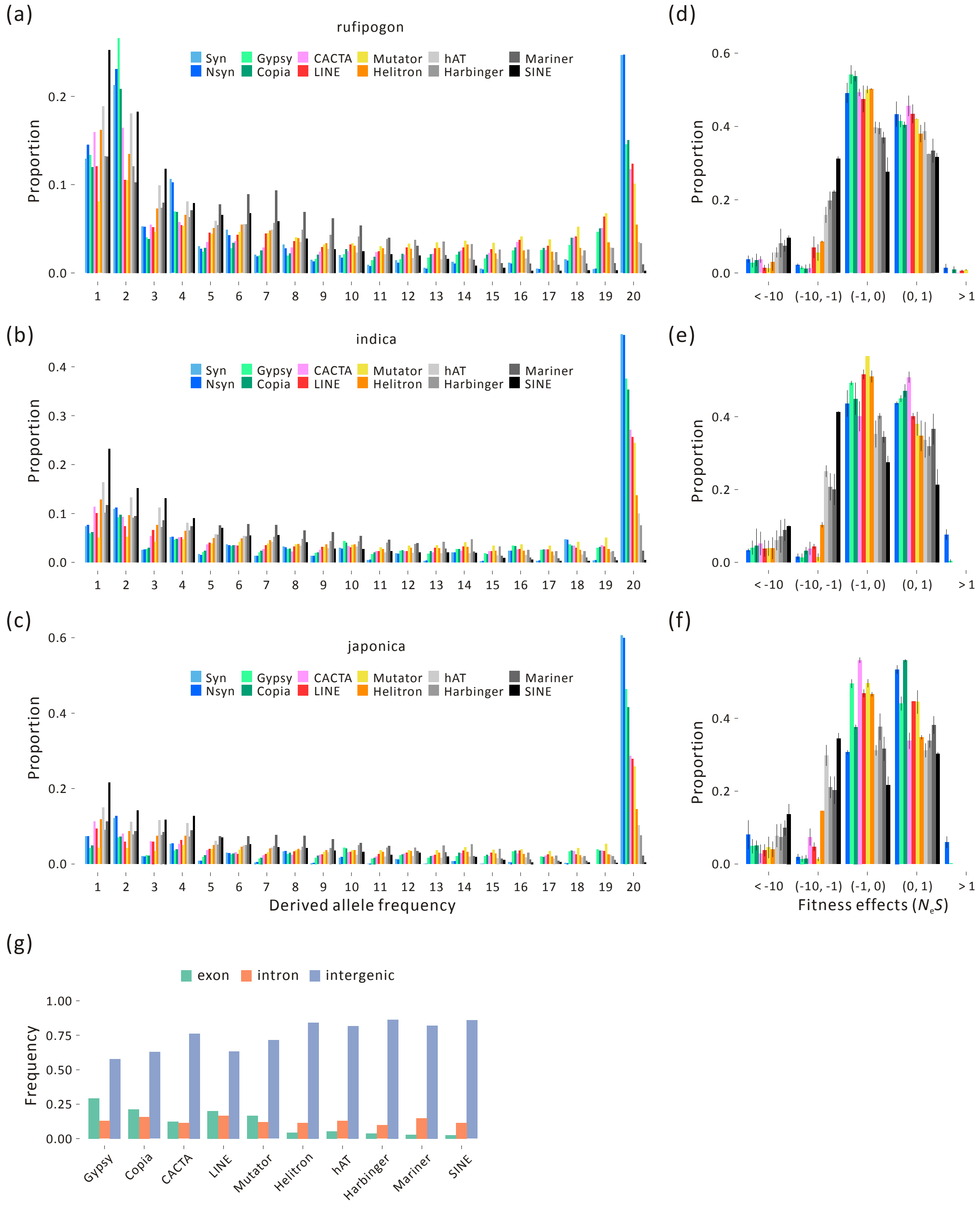

**Fig. S8.** Graphs **(a)**, **(b)** and **(c)** illustrate the SFS for various TE families in rufipogon, indica, japonica, respectively. Graphs **(d)**, **(e)** and **(f)** represent the DFEs estimates for each taxon and each family. **(g)** The frequency of SVs within exons, introns and intergenic regions for TEs of various families within the Nipponbare reference.

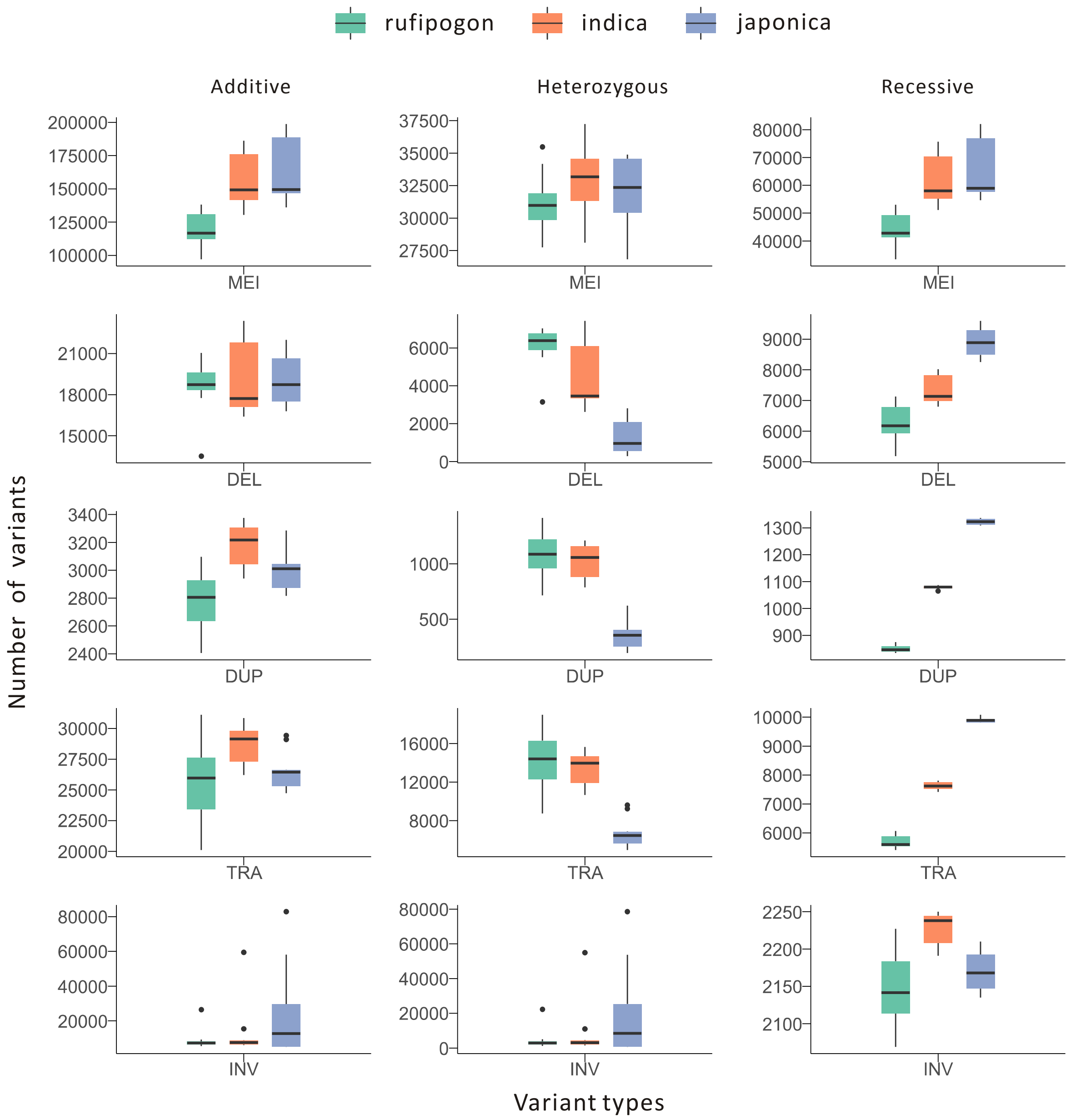

**Fig. S9.** The additive, heterozygous and recessive (homozygous) burden for three taxa and various categories of SV. See Figure 4 of the main text to see the additive, heterozygous and recessive burden for all of SV.

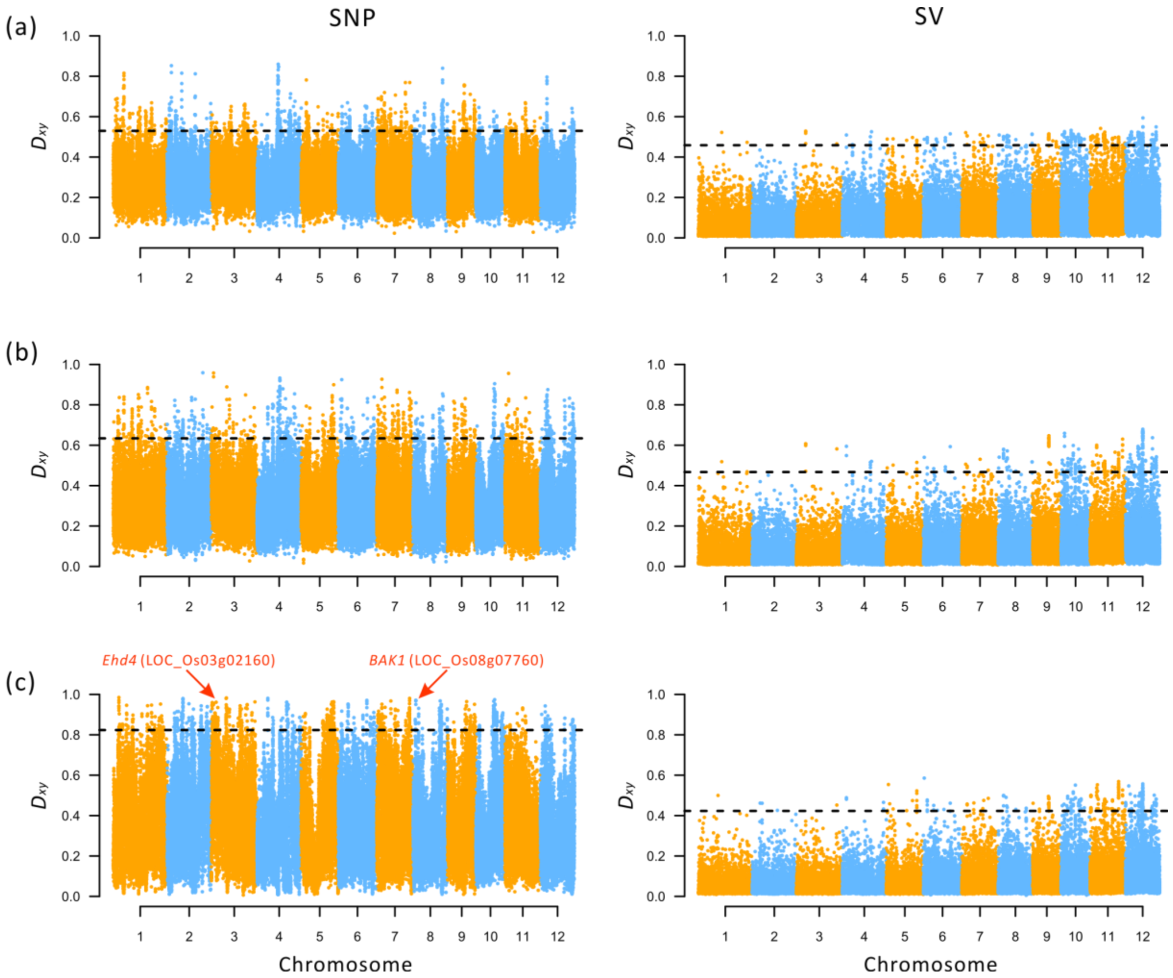

**Fig. S10.** Genomic differentiation ($D_{xy}$) between rufipogon and indica **(a)**, rufipogon and japonica **(b)**, and indica and japonica **(c)** based on SNPs and SVs in 20 kb windows. The dashed line represents the 99% cutoff. The arrows denote a couple of genes of interest.

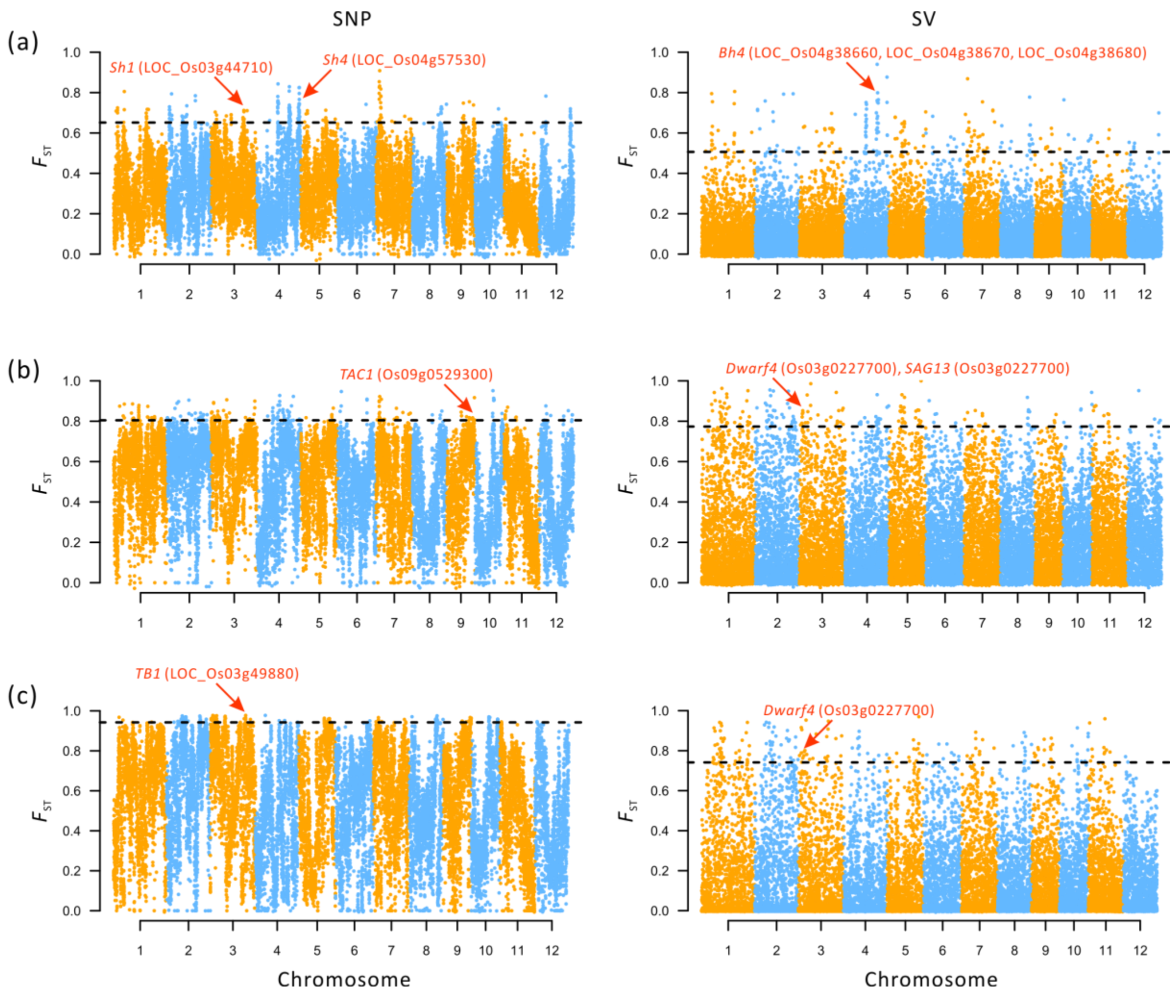

**Fig. S11.** Genomic differentiation ($F_{ST}$) between rufipogon and indica **(a)**, rufipogon and japonica **(b)**, and indica and japonica **(c)** based on SNPs and SVs in 20 kb non-overlapping windows. The dashed line represents the 99% cutoff. Arrows denote genes of interest.

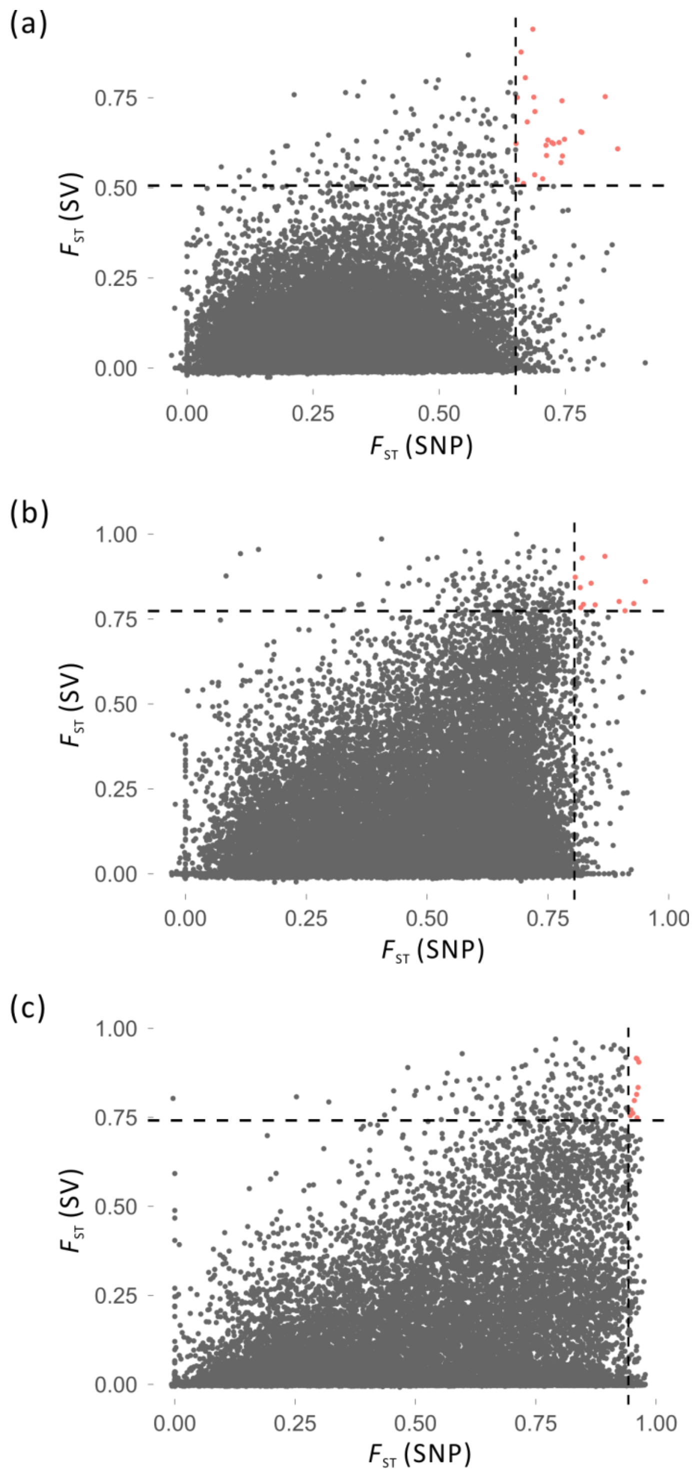

**Fig. S12.** Comparison of $F_{ST}$ between SNPs and SVs. **(a)** $F_{ST}$ is between rufipogon and indica, **(b)** $F_{ST}$ is between rufipogon and japonica, **(c)** $F_{ST}$ is between indica and japonica. The dashed line represents the 99% cutoff. Red dots represent the sites of significant differentiation in both SNP and SV, and there are 26, 12 and 10 sites in **(a)** $F_{ST}$ between rufipogon and indica, **(b)** $F_{ST}$ between rufipogon and japonica, **(c)** $F_{ST}$ between indica and japonica, respectively.

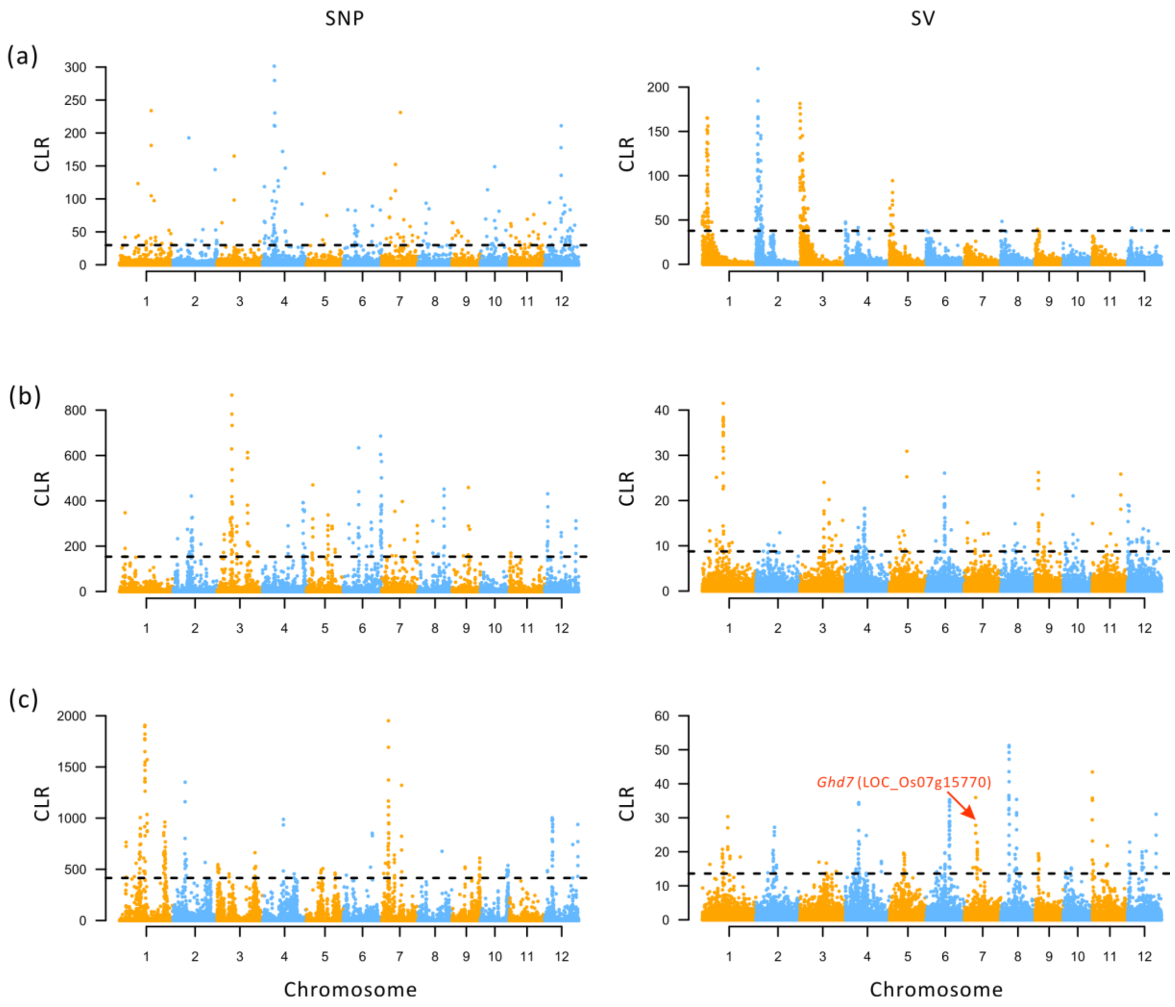

**Fig. S13.** Selective sweeps for rufipogon **(a)**, indica **(b)** and japonica **(c)** detected by composite likelihood ratio (CLR) test based on SNPs and SVs in 20 kb non-overlapping windows. The dashed line represents the 99% cutoff. Red arrow represent a gene *Ghd7* related to tillering.

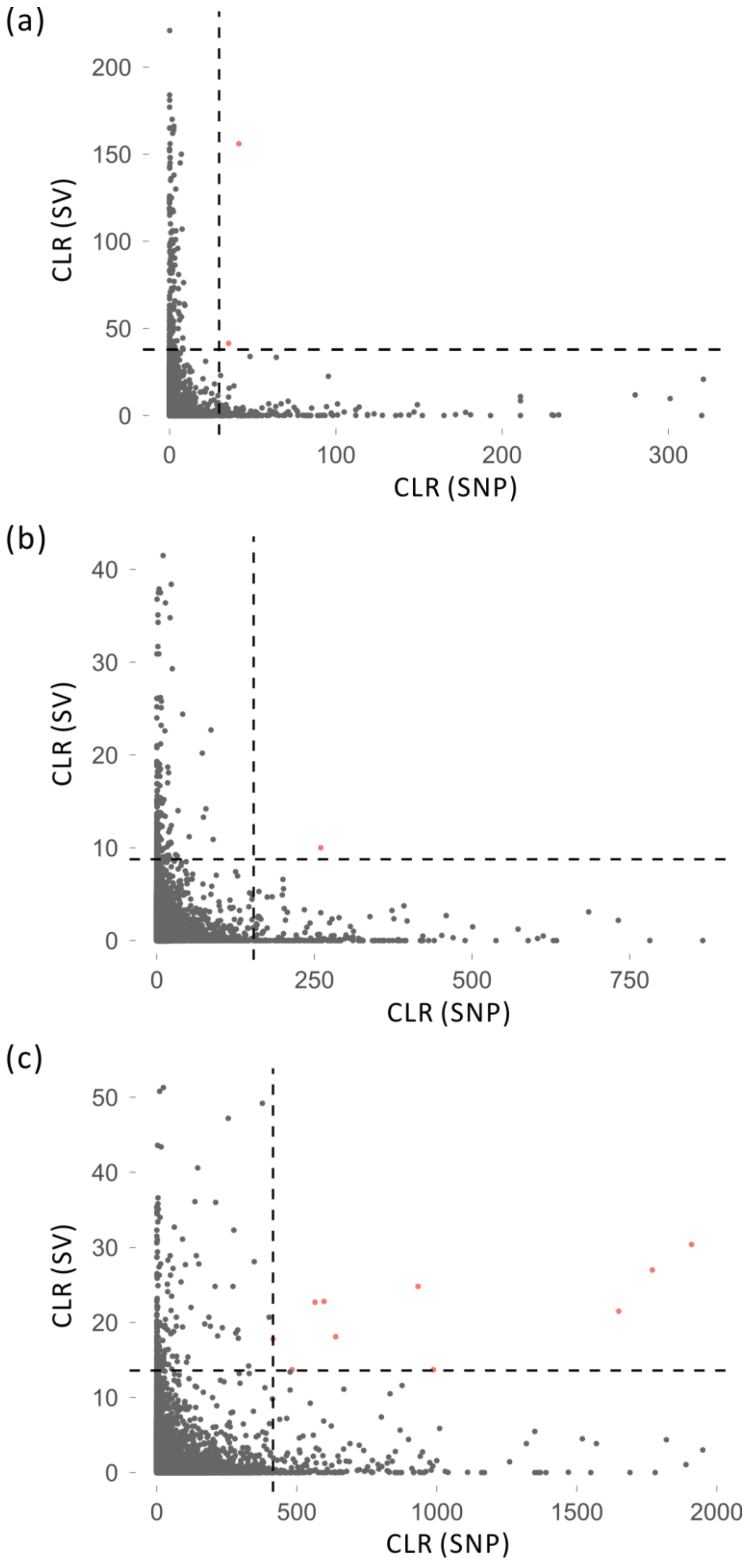

**Fig. S14.** Comparison of CLR between SNPs and SVs for **(a)** rufipogon, **(b)** indica and **(c)** japonica. These graphs reveal little correspondence (red dots) between CLRs based on SNPs (left) and SVs (right) in Figure S13.

**Table S1.** List of the 46 mislabelled accessions.

| | Run accession | Origin | Source[1] | Grouped in this study |
|---|---|---|---|---|
| 1 | ERR2245547 | indica | Zhao et al. 2018 | rufipogon |
| 2 | ERR609561-ERR609580 | indica | Wang et al. 2018 | temperate japonica |
| 3 | ERR617283-ERR617288 | indica | Wang et al. 2018 | rufipogon |
| 4 | ERR619511-ERR619524 | indica | Wang et al. 2018 | aus |
| 5 | ERR621557-ERR621569 | indica | Wang et al. 2018 | rufipogon |
| 6 | ERR622337-ERR622356 | indica | Wang et al. 2018 | aus |
| 7 | ERR625701-ERR625712 | indica | Wang et al. 2018 | rufipogon |
| 8 | SRR3234369 | indica | Zhang et al. 2016a | rufipogon |
| 9 | SRR5011847 | indica | Du et al. 2017 | rufipogon |
| 10 | ERR2241054 | temperate japonica | Zhao et al. 2018 | tropical japonica |
| 11 | ERR2245540 | temperate japonica | Zhao et al. 2018 | tropical japonica |
| 12 | ERR615186-ERR615191 | temperate japonica | Wang et al. 2018 | tropical japonica |
| 13 | ERR615877-ERR615882 | temperate japonica | Wang et al. 2018 | tropical japonica |
| 14 | ERR626638-ERR626643 | temperate japonica | Wang et al. 2018 | indica |
| 15 | ERR629716-ERR629721 | temperate japonica | Wang et al. 2018 | indica |
| 16 | ERR629752-ERR629757 | temperate japonica | Wang et al. 2018 | indica |
| 17 | ERR636841-ERR636846 | temperate japonica | Wang et al. 2018 | temperate japonica |
| 18 | ERR637045-ERR637050 | temperate japonica | Wang et al. 2018 | tropical japonica |
| 19 | ERR607669-ERR607674 | tropical japonica | Wang et al. 2018 | temperate japonica |
| 20 | ERR608057-ERR608062 | tropical japonica | Wang et al. 2018 | temperate japonica |
| 21 | ERR608413-ERR608426 | tropical japonica | Wang et al. 2018 | indica |
| 22 | ERR611071-ERR611090 | tropical japonica | Wang et al. 2018 | indica |

| | | | | |
|---|---|---|---|---|
| 23 | ERR611151-ERR611170 | tropical japonica | Wang et al. 2018 | indica |
| 24 | ERR611517-ERR611536 | tropical japonica | Wang et al. 2018 | indica |
| 25 | ERR613946-ERR613959 | tropical japonica | Wang et al. 2018 | aromatic |
| 26 | ERR616381-ERR616386 | tropical japonica | Wang et al. 2018 | temperate japonica |
| 27 | ERR626504-ERR626509 | tropical japonica | Wang et al. 2018 | indica |
| 28 | ERR629842-ERR629855 | tropical japonica | Wang et al. 2018 | indica |
| 29 | ERR630772-ERR630791 | tropical japonica | Wang et al. 2018 | indica |
| 30 | ERR630970-ERR630989 | tropical japonica | Wang et al. 2018 | indica |
| 31 | ERR636847-ERR636852 | tropical japonica | Wang et al. 2018 | temperate japonica |
| 32 | ERR2240123 | rufipogon | Zhao et al. 2018 | aus |
| 33 | DRR088679 | rufipogon | NCBI | temperate japonica |
| 34 | SRR1016479 | rufipogon | Fawcett et al. 2013 | indica |
| 35 | DRR088678 | rufipogon | NCBI | aus |
| 36 | SRR8324858 | rufipogon | Lou et al. 2015 | indica |
| 37 | SRR8324893 | rufipogon | Lou et al. 2015 | indica |
| 38 | SRR8324917 | rufipogon | Lou et al. 2015 | indica |
| 39 | SRR8324921 | rufipogon | Lou et al. 2015 | indica |
| 40 | SRR8324922 | rufipogon | Lou et al. 2015 | indica |
| 41 | SRR8324923 | rufipogon | Lou et al. 2015 | indica |
| 42 | ERR2245518 | aus | Zhao et al. 2018 | indica |
| 43 | SRR1450138 | nivara | Ammiraju et al. 2006 | indica |
| 44 | SRR1450139 | nivara | Ammiraju et al. 2006 | indica |
| 45 | SRR1450142 | nivara | Ammiraju et al. 2006 | aus |
| 46 | SRR1528449 | nivara | Ammiraju et al. 2006 | aus |

[1]References are provided at the end of the document.

**Table S2.** List of the 358 accessions used in this study.

| Species | Variaty | Depth (X) | Run accession | Source[1] |
|---|---|---|---|---|
| *O. sativa* | indica | 214 | ERR2242621 | Zhao et al. 2018 |
| *O. sativa* | indica | 104 | ERR2245513 | Zhao et al. 2018 |
| *O. sativa* | indica | 149 | ERR2245516 | Zhao et al. 2018 |
| *O. sativa* | indica | 157 | ERR2245525 | Zhao et al. 2018 |
| *O. sativa* | indica | 149 | ERR2242624 | Zhao et al. 2018 |
| *O. sativa* | indica | 177 | ERR2245528 | Zhao et al. 2018 |
| *O. sativa* | indica | 125 | ERR2245530 | Zhao et al. 2018 |
| *O. sativa* | indica | 132 | ERR2245531 | Zhao et al. 2018 |
| *O. sativa* | indica | 166 | ERR2245533 | Zhao et al. 2018 |
| *O. sativa* | indica | 185 | ERR2242626 | Zhao et al. 2018 |
| *O. sativa* | indica | 186 | ERR2245535 | Zhao et al. 2018 |
| *O. sativa* | indica | 173 | ERR2245537 | Zhao et al. 2018 |
| *O. sativa* | indica | 168 | ERR2245538 | Zhao et al. 2018 |
| *O. sativa* | indica | 222 | ERR2245542 | Zhao et al. 2018 |
| *O. sativa* | indica | 117 | ERR2245543 | Zhao et al. 2018 |
| *O. sativa* | indica | 180 | ERR2241060 | Zhao et al. 2018 |
| *O. sativa* | indica | 108 | ERR2245544 | Zhao et al. 2018 |
| *O. sativa* | indica | 37 | ERR606786 | Wang et al. 2018 |
| *O. sativa* | indica | 50 | ERR606887 | Wang et al. 2018 |
| *O. sativa* | indica | 34 | ERR607071 | Wang et al. 2018 |
| *O. sativa* | indica | 35 | ERR607855 | Wang et al. 2018 |
| *O. sativa* | indica | 44 | ERR607931 | Wang et al. 2018 |
| *O. sativa* | indica | 42 | ERR611385 | Wang et al. 2018 |
| *O. sativa* | indica | 44 | ERR611485 | Wang et al. 2018 |
| *O. sativa* | indica | 50 | ERR611723 | Wang et al. 2018 |
| *O. sativa* | indica | 39 | ERR611763 | Wang et al. 2018 |
| *O. sativa* | indica | 46 | ERR611975 | Wang et al. 2018 |
| *O. sativa* | indica | 37 | ERR612509 | Wang et al. 2018 |
| *O. sativa* | indica | 33 | ERR614543 | Wang et al. 2018 |
| *O. sativa* | indica | 39 | ERR614671 | Wang et al. 2018 |

| | | | | |
|---|---|---|---|---|
| *O. sativa* | indica | 30 | ERR626734 | Wang et al. 2018 |
| *O. sativa* | indica | 49 | ERR630642 | Wang et al. 2018 |
| *O. sativa* | indica | 49 | ERR630752 | Wang et al. 2018 |
| *O. sativa* | indica | 41 | ERR630844 | Wang et al. 2018 |
| *O. sativa* | indica | 48 | ERR631080 | Wang et al. 2018 |
| *O. sativa* | indica | 39 | ERR631259 | Wang et al. 2018 |
| *O. sativa* | indica | 50 | ERR631593 | Wang et al. 2018 |
| *O. sativa* | indica | 56 | ERR631613 | Wang et al. 2018 |
| *O. sativa* | indica | 47 | ERR614797 | Wang et al. 2018 |
| *O. sativa* | indica | 48 | ERR630864 | Wang et al. 2018 |
| *O. sativa* | indica | 44 | ERR631679 | Wang et al. 2018 |
| *O. sativa* | indica | 45 | ERR631725 | Wang et al. 2018 |
| *O. sativa* | indica | 46 | ERR631745 | Wang et al. 2018 |
| *O. sativa* | indica | 26 | ERR605587 | Wang et al. 2018 |
| *O. sativa* | indica | 25 | ERR606690 | Wang et al. 2018 |
| *O. sativa* | indica | 29 | ERR607817 | Wang et al. 2018 |
| *O. sativa* | indica | 26 | ERR608075 | Wang et al. 2018 |
| *O. sativa* | indica | 25 | ERR608111 | Wang et al. 2018 |
| *O. sativa* | indica | 24 | ERR608203 | Wang et al. 2018 |
| *O. sativa* | indica | 28 | ERR608257 | Wang et al. 2018 |
| *O. sativa* | indica | 27 | ERR609385 | Wang et al. 2018 |
| *O. sativa* | indica | 28 | ERR609907 | Wang et al. 2018 |
| *O. sativa* | indica | 27 | ERR610367 | Wang et al. 2018 |
| *O. sativa* | indica | 26 | ERR610733 | Wang et al. 2018 |
| *O. sativa* | indica | 29 | ERR611251 | Wang et al. 2018 |
| *O. sativa* | indica | 29 | ERR611359 | Wang et al. 2018 |
| *O. sativa* | indica | 29 | ERR611405 | Wang et al. 2018 |
| *O. sativa* | indica | 29 | ERR611929 | Wang et al. 2018 |
| *O. sativa* | indica | 28 | ERR612299 | Wang et al. 2018 |
| *O. sativa* | indica | 29 | ERR612535 | Wang et al. 2018 |
| *O. sativa* | indica | 26 | ERR612813 | Wang et al. 2018 |
| *O. sativa* | indica | 27 | ERR612915 | Wang et al. 2018 |

| | | | | |
|---|---|---|---|---|
| *O. sativa* | indica | 29 | ERR613011 | Wang et al. 2018 |
| *O. sativa* | indica | 27 | ERR613017 | Wang et al. 2018 |
| *O. sativa* | indica | 27 | ERR613533 | Wang et al. 2018 |
| *O. sativa* | indica | 27 | ERR614088 | Wang et al. 2018 |
| *O. sativa* | indica | 27 | ERR614281 | Wang et al. 2018 |
| *O. sativa* | indica | 24 | ERR615033 | Wang et al. 2018 |
| *O. sativa* | indica | 28 | ERR615780 | Wang et al. 2018 |
| *O. sativa* | indica | 27 | ERR615915 | Wang et al. 2018 |
| *O. sativa* | indica | 28 | ERR615965 | Wang et al. 2018 |
| *O. sativa* | indica | 28 | ERR616060 | Wang et al. 2018 |
| *O. sativa* | indica | 28 | ERR616554 | Wang et al. 2018 |
| *O. sativa* | indica | 28 | ERR616967 | Wang et al. 2018 |
| *O. sativa* | indica | 25 | ERR618181 | Wang et al. 2018 |
| *O. sativa* | indica | 26 | ERR618187 | Wang et al. 2018 |
| *O. sativa* | indica | 27 | ERR619471 | Wang et al. 2018 |
| *O. sativa* | indica | 27 | ERR619612 | Wang et al. 2018 |
| *O. sativa* | indica | 25 | ERR619673 | Wang et al. 2018 |
| *O. sativa* | indica | 28 | ERR619930 | Wang et al. 2018 |
| *O. sativa* | indica | 29 | ERR620191 | Wang et al. 2018 |
| *O. sativa* | indica | 26 | ERR620217 | Wang et al. 2018 |
| *O. sativa* | indica | 29 | ERR620705 | Wang et al. 2018 |
| *O. sativa* | indica | 25 | ERR623190 | Wang et al. 2018 |
| *O. sativa* | indica | 27 | ERR623496 | Wang et al. 2018 |
| *O. sativa* | indica | 28 | ERR625102 | Wang et al. 2018 |
| *O. sativa* | indica | 27 | ERR626441 | Wang et al. 2018 |
| *O. sativa* | indica | 29 | ERR629958 | Wang et al. 2018 |
| *O. sativa* | indica | 25 | ERR630114 | Wang et al. 2018 |
| *O. sativa* | indica | 22 | ERR630464 | Wang et al. 2018 |
| *O. sativa* | indica | 27 | ERR635447 | Wang et al. 2018 |
| *O. sativa* | indica | 27 | ERR661707 | Wang et al. 2018 |
| *O. sativa* | indica | 29 | ERR977887 | Wang et al. 2018 |
| *O. sativa* | Indica | 120 | SRR3234372 | Zhang et al. 2016b |

| | | | | |
|---|---|---|---|---|
| *O. sativa* | indica | 93 | SRR3056468 | Mahesh et al. 2016 |
| *O. sativa* | indica | 20 | SRR2175888 | NCBI |
| *O. sativa* | japonica | 169 | ERR2245515 | Zhao et al. 2018 |
| *O. sativa* | japonica | 109 | ERR2245517 | Zhao et al. 2018 |
| *O. sativa* | japonica | 221 | ERR2245522 | Zhao et al. 2018 |
| *O. sativa* | japonica | 186 | ERR2242623 | Zhao et al. 2018 |
| *O. sativa* | japonica | 34 | ERR606940 | Wang et al. 2018 |
| *O. sativa* | japonica | 44 | ERR611091 | Wang et al. 2018 |
| *O. sativa* | japonica | 36 | ERR612371 | Wang et al. 2018 |
| *O. sativa* | japonica | 77 | ERR617233 | Wang et al. 2018 |
| *O. sativa* | japonica | 33 | ERR622439 | Wang et al. 2018 |
| *O. sativa* | japonica | 35 | ERR630798 | Wang et al. 2018 |
| *O. sativa* | japonica | 27 | ERR606338 | Wang et al. 2018 |
| *O. sativa* | japonica | 29 | ERR606825 | Wang et al. 2018 |
| *O. sativa* | japonica | 27 | ERR606959 | Wang et al. 2018 |
| *O. sativa* | japonica | 27 | ERR613878 | Wang et al. 2018 |
| *O. sativa* | japonica | 27 | ERR617277 | Wang et al. 2018 |
| *O. sativa* | japonica | 29 | ERR619457 | Wang et al. 2018 |
| *O. sativa* | japonica | 29 | ERR622787 | Wang et al. 2018 |
| *O. sativa* | japonica | 28 | ERR624803 | Wang et al. 2018 |
| *O. sativa* | japonica | 28 | ERR625769 | Wang et al. 2018 |
| *O. sativa* | japonica | 27 | ERR625853 | Wang et al. 2018 |
| *O. sativa* | japonica | 23 | ERR605485 | Wang et al. 2018 |
| *O. sativa* | japonica | 20 | ERR605635 | Wang et al. 2018 |
| *O. sativa* | japonica | 22 | ERR605701 | Wang et al. 2018 |
| *O. sativa* | japonica | 22 | ERR605976 | Wang et al. 2018 |
| *O. sativa* | japonica | 21 | ERR607615 | Wang et al. 2018 |
| *O. sativa* | japonica | 20 | ERR607899 | Wang et al. 2018 |
| *O. sativa* | japonica | 23 | ERR607995 | Wang et al. 2018 |
| *O. sativa* | japonica | 19 | ERR608185 | Wang et al. 2018 |
| *O. sativa* | japonica | 25 | ERR608775 | Wang et al. 2018 |
| *O. sativa* | japonica | 26 | ERR609345 | Wang et al. 2018 |

| | | | | |
|---|---|---|---|---|
| *O. sativa* | japonica | 24 | ERR612119 | Wang et al. 2018 |
| *O. sativa* | japonica | 25 | ERR612339 | Wang et al. 2018 |
| *O. sativa* | japonica | 25 | ERR612661 | Wang et al. 2018 |
| *O. sativa* | japonica | 27 | ERR613407 | Wang et al. 2018 |
| *O. sativa* | japonica | 24 | ERR613437 | Wang et al. 2018 |
| *O. sativa* | japonica | 26 | ERR613866 | Wang et al. 2018 |
| *O. sativa* | japonica | 23 | ERR613890 | Wang et al. 2018 |
| *O. sativa* | japonica | 23 | ERR613990 | Wang et al. 2018 |
| *O. sativa* | japonica | 23 | ERR617649 | Wang et al. 2018 |
| *O. sativa* | japonica | 20 | ERR617903 | Wang et al. 2018 |
| *O. sativa* | japonica | 20 | ERR618163 | Wang et al. 2018 |
| *O. sativa* | japonica | 21 | ERR618199 | Wang et al. 2018 |
| *O. sativa* | japonica | 23 | ERR618321 | Wang et al. 2018 |
| *O. sativa* | japonica | 18 | ERR618585 | Wang et al. 2018 |
| *O. sativa* | japonica | 19 | ERR618717 | Wang et al. 2018 |
| *O. sativa* | japonica | 19 | ERR620949 | Wang et al. 2018 |
| *O. sativa* | japonica | 20 | ERR623201 | Wang et al. 2018 |
| *O. sativa* | japonica | 26 | ERR624816 | Wang et al. 2018 |
| *O. sativa* | japonica | 27 | ERR626391 | Wang et al. 2018 |
| *O. sativa* | japonica | 21 | ERR626819 | Wang et al. 2018 |
| *O. sativa* | japonica | 72 | ERR2245519 | Zhao et al. 2018 |
| *O. sativa* | japonica | 141 | ERR2245520 | Zhao et al. 2018 |
| *O. sativa* | japonica | 148 | ERR2245523 | Zhao et al. 2018 |
| *O. sativa* | japonica | 112 | ERR2245524 | Zhao et al. 2018 |
| *O. sativa* | japonica | 167 | ERR2245527 | Zhao et al. 2018 |
| *O. sativa* | japonica | 179 | ERR2242625 | Zhao et al. 2018 |
| *O. sativa* | japonica | 179 | ERR2245529 | Zhao et al. 2018 |
| *O. sativa* | japonica | 166 | ERR2245532 | Zhao et al. 2018 |
| *O. sativa* | japonica | 91 | ERR2245534 | Zhao et al. 2018 |
| *O. sativa* | japonica | 155 | ERR2245536 | Zhao et al. 2018 |
| *O. sativa* | japonica | 145 | ERR2245539 | Zhao et al. 2018 |
| *O. sativa* | japonica | 121 | ERR2245541 | Zhao et al. 2018 |

| | | | | |
|---|---|---|---|---|
| *O. sativa* | japonica | 122 | ERR2241059 | Zhao et al. 2018 |
| *O. sativa* | japonica | 91 | ERR2245545 | Zhao et al. 2018 |
| *O. sativa* | japonica | 122 | ERR2241058 | Zhao et al. 2018 |
| *O. sativa* | japonica | 167 | ERR2241056 | Zhao et al. 2018 |
| *O. sativa* | japonica | 165 | ERR2240127 | Zhao et al. 2018 |
| *O. sativa* | japonica | 139 | ERR2241055 | Zhao et al. 2018 |
| *O. sativa* | japonica | 116 | ERR2245546 | Zhao et al. 2018 |
| *O. sativa* | japonica | 103 | ERR2240128 | Zhao et al. 2018 |
| *O. sativa* | japonica | 31 | ERR607317 | Wang et al. 2018 |
| *O. sativa* | japonica | 51 | ERR611637 | Wang et al. 2018 |
| *O. sativa* | japonica | 31 | ERR624716 | Wang et al. 2018 |
| *O. sativa* | japonica | 36 | ERR622583 | Wang et al. 2018 |
| *O. sativa* | japonica | 31 | ERR624521 | Wang et al. 2018 |
| *O. sativa* | japonica | 37 | ERR624985 | Wang et al. 2018 |
| *O. sativa* | japonica | 33 | ERR625050 | Wang et al. 2018 |
| *O. sativa* | japonica | 45 | ERR630694 | Wang et al. 2018 |
| *O. sativa* | japonica | 36 | ERR630714 | Wang et al. 2018 |
| *O. sativa* | japonica | 30 | ERR978072 | Wang et al. 2018 |
| *O. sativa* | japonica | 21 | ERR605429 | Wang et al. 2018 |
| *O. sativa* | japonica | 19 | ERR607651 | Wang et al. 2018 |
| *O. sativa* | japonica | 26 | ERR608287 | Wang et al. 2018 |
| *O. sativa* | japonica | 24 | ERR608683 | Wang et al. 2018 |
| *O. sativa* | japonica | 21 | ERR608907 | Wang et al. 2018 |
| *O. sativa* | japonica | 20 | ERR609009 | Wang et al. 2018 |
| *O. sativa* | japonica | 29 | ERR609093 | Wang et al. 2018 |
| *O. sativa* | japonica | 21 | ERR613683 | Wang et al. 2018 |
| *O. sativa* | japonica | 28 | ERR620519 | Wang et al. 2018 |
| *O. sativa* | japonica | 20 | ERR621625 | Wang et al. 2018 |
| *O. sativa* | japonica | 21 | ERR621937 | Wang et al. 2018 |
| *O. sativa* | japonica | 28 | ERR622619 | Wang et al. 2018 |
| *O. sativa* | japonica | 25 | ERR622763 | Wang et al. 2018 |
| *O. sativa* | japonica | 23 | ERR622775 | Wang et al. 2018 |

| | | | | |
|---|---|---|---|---|
| *O. sativa* | japonica | 22 | ERR622991 | Wang et al. 2018 |
| *O. sativa* | japonica | 23 | ERR623019 | Wang et al. 2018 |
| *O. sativa* | japonica | 27 | ERR623389 | Wang et al. 2018 |
| *O. sativa* | japonica | 22 | ERR625917 | Wang et al. 2018 |
| *O. sativa* | japonica | 27 | ERR626234 | Wang et al. 2018 |
| *O. sativa* | japonica | 19 | ERR636673 | Wang et al. 2018 |
| *O. sativa* | japonica | 20 | ERR636709 | Wang et al. 2018 |
| *O. sativa* | japonica | 20 | ERR636733 | Wang et al. 2018 |
| *O. sativa* | japonica | 25 | ERR636955 | Wang et al. 2018 |
| *O. sativa* | japonica | 18 | ERR636979 | Wang et al. 2018 |
| *O. sativa* | japonica | 19 | ERR637009 | Wang et al. 2018 |
| *O. sativa* | japonica | 20 | ERR637051 | Wang et al. 2018 |
| *O. rufipogon* | wild & perennial | 141 | ERR2245548 | Zhao et al. 2018 |
| *O. rufipogon* | wild & perennial | 137 | ERR2245549 | Zhao et al. 2018 |
| *O. rufipogon* | wild & perennial | 111 | ERR2245550 | Zhao et al. 2018 |
| *O. rufipogon* | wild & perennial | 120 | ERR2245551 | Zhao et al. 2018 |
| *O. rufipogon* | wild & perennial | 150 | ERR2245552 | Zhao et al. 2018 |
| *O. rufipogon* | wild & perennial | 164 | ERR2245554 | Zhao et al. 2018 |
| *O. rufipogon* | wild & perennial | 89 | ERR2245556 | Zhao et al. 2018 |
| *O. rufipogon* | wild & perennial | 155 | ERR2245557 | Zhao et al. 2018 |
| *O. rufipogon* | wild & perennial | 133 | ERR2240126 | Zhao et al. 2018 |
| *O. rufipogon* | wild & perennial | 137 | ERR2240125 | Zhao et al. 2018 |
| *O. rufipogon* | wild & perennial | 44 | SRR1016473 | Fawcett et al. 2013 |
| *O. rufipogon* | wild & perennial | 32 | SRR1016475 | Fawcett et al. 2013 |
| *O. rufipogon* | wild & perennial | 29 | SRR1016477 | Fawcett et al. 2013 |
| *O. rufipogon* | wild & perennial | 28 | SRR1016481 | Fawcett et al. 2013 |
| *O. rufipogon* | wild & perennial | 25 | SRR1016483 | Fawcett et al. 2013 |
| *O. rufipogon* | wild & perennial | 26 | SRR1016485 | Fawcett et al. 2013 |
| *O. rufipogon* | wild & perennial | 32 | SRR1016487 | Fawcett et al. 2013 |
| *O. rufipogon* | wild & perennial | 45 | SRR1016489 | Fawcett et al. 2013 |
| *O. rufipogon* | wild & perennial | 26 | SRR1016492 | Fawcett et al. 2013 |
| *O. rufipogon* | wild & perennial | 46 | DRR001183 | NCBI |

| | | | | |
|---|---|---|---|---|
| *O. rufipogon* | wild & perennial | 46 | DRR001184 | NCBI |
| *O. rufipogon* | wild & perennial | 51 | DRR001185 | NCBI |
| *O. rufipogon* | wild & perennial | 45 | DRR001186 | NCBI |
| *O. rufipogon* | wild & perennial | 38 | DRR001187 | NCBI |
| *O. rufipogon* | wild & perennial | 41 | DRR001188 | NCBI |
| *O. rufipogon* | wild & perennial | 44 | DRR001189 | NCBI |
| *O. rufipogon* | wild & perennial | 44 | DRR001190 | NCBI |
| *O. rufipogon* | wild & perennial | 36 | DRR088674 | NCBI |
| *O. rufipogon* | wild & perennial | 20 | DRR088675 | NCBI |
| *O. rufipogon* | wild & perennial | 24 | DRR088681 | NCBI |
| *O. rufipogon* | wild & perennial | 21 | DRR088682 | NCBI |
| *O. rufipogon* | wild & perennial | 20 | DRR088684 | NCBI |
| *O. rufipogon* | wild & perennial | 22 | DRR088685 | NCBI |
| *O. rufipogon* | wild & perennial | 20 | DRR088690 | NCBI |
| *O. rufipogon* | wild & perennial | 20 | DRR088691 | NCBI |
| *O. rufipogon* | wild & perennial | 84 | SRR3180882 | NCBI |
| *O. rufipogon* | wild & perennial | 28 | SRR519098 | NCBI |
| *O. rufipogon* | wild & perennial | 38 | SRR519100 | NCBI |
| *O. rufipogon* | wild & perennial | 47 | SRR5536054 | NCBI |
| *O. rufipogon* | wild & perennial | 53 | SRR5536055 | NCBI |
| *O. rufipogon* | wild & perennial | 18 | DRR088676 | NCBI |
| *O. rufipogon* | wild & perennial | 16 | DRR088677 | NCBI |
| *O. rufipogon* | wild & perennial | 19 | DRR088680 | NCBI |
| *O. rufipogon* | wild & perennial | 19 | DRR088683 | NCBI |
| *O. rufipogon* | wild & perennial | 19 | DRR088686 | NCBI |
| *O. rufipogon* | wild & perennial | 15 | DRR088687 | NCBI |
| *O. rufipogon* | wild & perennial | 16 | DRR088688 | NCBI |
| *O. rufipogon* | wild & perennial | 16 | DRR088689 | NCBI |
| *O. rufipogon* | wild & perennial | 19 | DRR088692 | NCBI |
| *O. rufipogon* | wild & perennial | 17 | DRR088693 | NCBI |
| *O. rufipogon* | wild & perennial | 16 | DRR088694 | NCBI |
| *O. rufipogon* | wild & perennial | 19 | SRR8324860 | Lou et al. 2015 |

| | | | | |
|---|---|---|---|---|
| *O. rufipogon* | wild & perennial | 18 | SRR8324919 | Lou et al. 2015 |
| *O. rufipogon* | wild & perennial | 18 | SRR8324916 | Lou et al. 2015 |
| *O. rufipogon* | wild & perennial | 17 | SRR8324913 | Lou et al. 2015 |
| *O. rufipogon* | wild & perennial | 19 | SRR8324911 | Lou et al. 2015 |
| *O. rufipogon* | wild & perennial | 17 | SRR8324908 | Lou et al. 2015 |
| *O. rufipogon* | wild & perennial | 18 | SRR8324907 | Lou et al. 2015 |
| *O. rufipogon* | wild & perennial | 16 | SRR8324888 | Lou et al. 2015 |
| *O. rufipogon* | wild & perennial | 19 | SRR8324880 | Lou et al. 2015 |
| *O. rufipogon* | wild & perennial | 20 | SRR8324857 | Lou et al. 2015 |
| *O. rufipogon* | wild & perennial | 22 | SRR8324859 | Lou et al. 2015 |
| *O. rufipogon* | wild & perennial | 22 | SRR8324861 | Lou et al. 2015 |
| *O. rufipogon* | wild & perennial | 23 | SRR8324862 | Lou et al. 2015 |
| *O. rufipogon* | wild & perennial | 25 | SRR8324863 | Lou et al. 2015 |
| *O. rufipogon* | wild & perennial | 23 | SRR8324864 | Lou et al. 2015 |
| *O. rufipogon* | wild & perennial | 20 | SRR8324877 | Lou et al. 2015 |
| *O. rufipogon* | wild & perennial | 23 | SRR8324878 | Lou et al. 2015 |
| *O. rufipogon* | wild & perennial | 22 | SRR8324879 | Lou et al. 2015 |
| *O. rufipogon* | wild & perennial | 22 | SRR8324881 | Lou et al. 2015 |
| *O. rufipogon* | wild & perennial | 20 | SRR8324882 | Lou et al. 2015 |
| *O. rufipogon* | wild & perennial | 22 | SRR8324883 | Lou et al. 2015 |
| *O. rufipogon* | wild & perennial | 22 | SRR8324884 | Lou et al. 2015 |
| *O. rufipogon* | wild & perennial | 20 | SRR8324885 | Lou et al. 2015 |
| *O. rufipogon* | wild & perennial | 20 | SRR8324886 | Lou et al. 2015 |
| *O. rufipogon* | wild & perennial | 21 | SRR8324887 | Lou et al. 2015 |
| *O. rufipogon* | wild & perennial | 23 | SRR8324889 | Lou et al. 2015 |
| *O. rufipogon* | wild & perennial | 21 | SRR8324890 | Lou et al. 2015 |
| *O. rufipogon* | wild & perennial | 20 | SRR8324891 | Lou et al. 2015 |
| *O. rufipogon* | wild & perennial | 20 | SRR8324892 | Lou et al. 2015 |
| *O. rufipogon* | wild & perennial | 24 | SRR8324894 | Lou et al. 2015 |
| *O. rufipogon* | wild & perennial | 21 | SRR8324895 | Lou et al. 2015 |
| *O. rufipogon* | wild & perennial | 23 | SRR8324896 | Lou et al. 2015 |
| *O. rufipogon* | wild & perennial | 20 | SRR8324909 | Lou et al. 2015 |

| | | | | |
|---|---|---|---|---|
| *O. rufipogon* | wild & perennial | 21 | SRR8324910 | Lou et al. 2015 |
| *O. rufipogon* | wild & perennial | 22 | SRR8324912 | Lou et al. 2015 |
| *O. rufipogon* | wild & perennial | 23 | SRR8324914 | Lou et al. 2015 |
| *O. rufipogon* | wild & perennial | 20 | SRR8324915 | Lou et al. 2015 |
| *O. rufipogon* | wild & perennial | 21 | SRR8324918 | Lou et al. 2015 |
| *O. rufipogon* | wild & perennial | 23 | SRR8324920 | Lou et al. 2015 |
| *O. sativa* | aus | 156 | ERR2245512 | Zhao et al. 2018 |
| *O. sativa* | aus | 333 | ERR2241057 | Zhao et al. 2018 |
| *O. sativa* | aus | 208 | ERR2240124 | Zhao et al. 2018 |
| *O. sativa* | aus | 34 | ERR622357 | Wang et al. 2018 |
| *O. sativa* | aus | 37 | ERR610126 | Wang et al. 2018 |
| *O. sativa* | aus | 41 | ERR622253 | Wang et al. 2018 |
| *O. sativa* | aus | 38 | ERR606520 | Wang et al. 2018 |
| *O. sativa* | aus | 36 | ERR606436 | Wang et al. 2018 |
| *O. sativa* | aus | 28 | ERR625797 | Wang et al. 2018 |
| *O. sativa* | aus | 20 | ERR623168 | Wang et al. 2018 |
| *O. sativa* | aus | 22 | ERR624547 | Wang et al. 2018 |
| *O. sativa* | aus | 34 | ERR635180 | Wang et al. 2018 |
| *O. sativa* | aus | 21 | ERR625368 | Wang et al. 2018 |
| *O. sativa* | aus | 36 | ERR606450 | Wang et al. 2018 |
| *O. sativa* | aus | 33 | ERR605277 | Wang et al. 2018 |
| *O. sativa* | aus | 22 | ERR605509 | Wang et al. 2018 |
| *O. sativa* | aus | 26 | ERR606138 | Wang et al. 2018 |
| *O. sativa* | aus | 24 | ERR606214 | Wang et al. 2018 |
| *O. sativa* | aus | 27 | ERR606226 | Wang et al. 2018 |
| *O. sativa* | aus | 33 | ERR606506 | Wang et al. 2018 |
| *O. sativa* | aus | 31 | ERR609887 | Wang et al. 2018 |
| *O. sativa* | aus | 33 | ERR612451 | Wang et al. 2018 |
| *O. sativa* | aus | 26 | ERR612969 | Wang et al. 2018 |
| *O. sativa* | aus | 22 | ERR618339 | Wang et al. 2018 |
| *O. sativa* | aromatic | 17 | ERR606102 | Wang et al. 2018 |
| *O. sativa* | aromatic | 25 | ERR606560 | Wang et al. 2018 |

| | | | | |
|---|---|---|---|---|
| *O. sativa* | aromatic | 28 | ERR606598 | Wang et al. 2018 |
| *O. sativa* | aromatic | 36 | ERR609633 | Wang et al. 2018 |
| *O. sativa* | aromatic | 30 | ERR610043 | Wang et al. 2018 |
| *O. sativa* | aromatic | 52 | ERR614068 | Wang et al. 2018 |
| *O. sativa* | aromatic | 18 | ERR614201 | Wang et al. 2018 |
| *O. sativa* | aromatic | 18 | ERR616652 | Wang et al. 2018 |
| *O. sativa* | aromatic | 16 | ERR618073 | Wang et al. 2018 |
| *O. sativa* | aromatic | 25 | ERR618133 | Wang et al. 2018 |
| *O. sativa* | aromatic | 21 | ERR625755 | Wang et al. 2018 |
| *O. sativa* | aromatic | 26 | ERR625839 | Wang et al. 2018 |
| *O. sativa* | aromatic | 29 | ERR625867 | Wang et al. 2018 |
| *O. sativa* | aromatic | 26 | ERR626324 | Wang et al. 2018 |
| *O. sativa* | aromatic | 18 | ERR630470 | Wang et al. 2018 |
| *O. sativa* | aromatic | 21 | ERR630476 | Wang et al. 2018 |
| *O. sativa* | aromatic | 20 | ERR630500 | Wang et al. 2018 |
| *O. sativa* | aromatic | 44 | ERR635284 | Wang et al. 2018 |
| *O. nivara* | wild & annual | 29 | SRR1743122 | Huang et al. 2018 |
| *O. nivara* | wild & annual | 106 | SRR7341624 | NCBI |
| *O. nivara* | wild & annual | 68 | SRR1707282 | Zhang et al. 2014 |
| *O. nivara* | wild & annual | 20 | SRR1450140 | Ammiraju et al. 2006 |
| *O. nivara* | wild & annual | 18 | SRR1450141 | Ammiraju et al. 2006 |
| *O. nivara* | wild & annual | 162 | SRR1450143 | Ammiraju et al. 2006 |
| *O. nivara* | wild & annual | 147 | SRR1450144 | Ammiraju et al. 2006 |
| *O. glaberrima* | | 75 | DRR128981 | NCBI |
| *O. glaberrima* | | 155 | SRR1206515 | Zuccolo et al. 2007 |
| *O. glaberrima* | | 155 | SRR1206516 | Zuccolo et al. 2007 |
| *O. glaberrima* | | 145 | SRR1206518 | Zuccolo et al. 2007 |
| *O. glaberrima* | | 159 | SRR1206519 | Zuccolo et al. 2007 |
| *O. glaberrima* | | 49 | SRR1712585 | Zhang et al. 2014 |
| *O. glaberrima* | | 51 | SRR7085035 | Zhang et al. 2014 |
| *O. barthii* | | 166 | SRR1528330 | Zhang et al. 2014 |
| *O. barthii* | | 43 | SRR1712898 | Zhang et al. 2014 |

| | | | |
|---|---|---|---|
| *O. barthii* | 88 | SRR7341625 | NCBI |
| *O. barthii* | 52 | ERR2008866 | Cubry et al. 2018 |
| *O. barthii* | 44 | ERR2008906 | Cubry et al. 2018 |
| *O. barthii* | 48 | ERR2008912 | Cubry et al. 2018 |
| *O. barthii* | 52 | ERR2009093 | Cubry et al. 2018 |
| *O. meridionalis* | 154 | SRR1450058 | Stein et al. 2018 |
| *O. meridionalis* | 157 | SRR1450059 | Stein et al. 2018 |
| *O. meridionalis* | 156 | SRR1450061 | Stein et al. 2018 |

[1]References are provided at the end of the document.

**Table S3.** The number of variants detected by Illumina short reads, PacBio long reads, and genome alignment.

| | Variant type | Illumina | PacBio | Genome alignment | |
|---|---|---|---|---|---|
| | | | | LASTZ | MUMmer |
| rufipogon | Number of sample | 90 | | 1 | 1 |
| | TRA | 73,051 | | | 182 |
| | DUP | 21,090 | | 1,121 | 0 |
| | DEL | 52,946 | | 22,367 | 12,978 |
| | INV | 178,141 | | 37 | 73 |
| | INS | | | 27,460 | 24 |
| | MEI | 280,556 | | 44,832 | 123,896 |
| indica | Number of sample | 96 | 6 | 6 | 6 |
| | TRA | 49,144 | 76,146 | | 1,143 |
| | DUP | 45,404 | 197 | 1,646 | 1 |
| | DEL | 27,188 | 58,621 | 33,551 | 12,788 |
| | INV | 261,158 | 11,081 | 105 | 227 |
| | INS | | 88,336 | 28,284 | 66 |
| | MEI | 280,904 | 269,104 | 66,599 | 221,790 |
| japonica | Number of sample | 106 | 3 | 6 | 6 |
| | TRA | 40,370 | 9,797 | | 221 |
| | DUP | 3,905 | 8 | 1,797 | 1 |
| | DEL | 21,325 | 7,134 | 28,773 | 25,919 |
| | INV | 259,990 | 807 | 67 | 74 |
| | INS | | 23,610 | 35,308 | 49 |
| | MEI | 246,979 | 52,572 | 89,723 | 49,494 |
| All taxa | Number of sample | 358 | 10 | | 15 |
| | TRA | 76,835 | 85,279 | | 1,332 |
| | DUP | 48,132 | 213 | 3,700 | 2 |
| | DEL | 72,930 | 60,747 | 62,953 | 48,153 |

| | | | | |
|---|---|---|---|---|
| INV | 341,752 | 11,532 | 159 | 343 |
| INS | | 103,447 | 70,065 | 113 |
| MEI | 284,741 | 270,708 | | 246,340 |

**Table S4.** The details of assembled genomes used in this study.

| Species | Infraspecific name (Abbr.) | NCBI BioProject | Sequencing technology | Assembly level | Reference |
|---|---|---|---|---|---|
| indica | Zhenshan 97 (ZS97) | PRJNA302542 | PacBio ~120X | Chromosome | Zhang et al. 2016a |
| | Shuhui 498 (R498) | PRJNA318714 | PacBio ~120X | Chromosome | Du et al. 2017 |
| | Minghui 63 (MH63) | PRJNA302543 | Pacbio ~120X | Chromosome | Zhang et al. 2016a |
| | IR8 | PRJNA353946 | PacBio ~70X | Chromosome | Stein et al. 2018 |
| | 93-11 | PRJNA427873 | PacBio ~116X | Chromosome | Zhang et al. 2018 |
| | Tetep | PRJNA482013 | PacBio ~148X | Chromosome | https://www.ncbi.nlm.nih.gov/assembly/GCA_004348155.2 |
| japonica | HEG4 | PRJNA264749 | Illumina ~220X | Chromosome | Lu et al. 2017 |
| | Nipponbare (IRGSP) | PRJDB1747 | Illumina | Chromosome | IRGSP 2005 |
| | Nipponbare (Nippo) | PRJNA427870 | PacBio ~117X | Chromosome | Zhang et al. 2018 |
| | Hitomebore (Hitom) | PRJDA67163 | Illumina ~179X | Chromosome | https://www.ncbi.nlm.nih.gov/assembly/GCA_000321445.1 |
| | Suijing18 | PRJNA396093 | PacBio ~62X | Contig | Nie et al. 2017 |
| | KitaakeX | PRJNA234782 | PacBio ~100X | Scaffold | Jain et al. 2019 |
| rufipogon | W1943 | PRJEB4137 | Illumina ~130X | Scaffold | Stein et al. 2018 |
| *O. glaberrima* | | PRJNA13765 | 454 Titanium | Scaffold | Wang et al. 2014 |
| *O. longistaminata* | | PRJDB6339 | PacBio ~66X | Chromosome | Reuscher et al. 2018 |

**Table S5.** Functional annotation of significantly divergent (top 1% $F_{ST}$) genes between each pair of taxa (rufipogon, indica and japonica).

| Gene ID (MSU7) | Function annotation |
|---|---|
| **Shared genes in indica-rufipogon comparision** | |
| LOC_Os01g01560 | transposon protein, putative, unclassified, expressed |
| LOC_Os01g02740 | retrotransposon protein, putative, unclassified, expressed |
| LOC_Os01g09110 | retrotransposon protein, putative, unclassified |
| LOC_Os01g11190 | expressed protein |
| LOC_Os01g15070 | retrotransposon protein, putative, unclassified |
| LOC_Os01g15260 | dynein light chain type 1 domain containing protein, expressed |
| LOC_Os01g15270 | expressed protein |
| LOC_Os01g15290 | ribosomal L18p/L5e family protein, putative, expressed |
| LOC_Os01g15300 | zinc finger helicase family protein, putative, expressed |
| LOC_Os01g17060 | expressed protein |
| LOC_Os01g18010 | hypothetical protein |
| LOC_Os01g18020 | retrotransposon protein, putative, unclassified |
| LOC_Os01g18250 | subtilase, putative, expressed |
| LOC_Os01g33136 | retrotransposon protein, putative, unclassified, expressed |
| LOC_Os01g40540 | lectin receptor-type protein kinase, putative, expressed |
| LOC_Os01g42890 | expressed protein |
| LOC_Os01g46530 | expressed protein |
| LOC_Os01g50220 | retrotransposon protein, putative, LINE subclass, expressed |
| LOC_Os01g62710 | transposon protein, putative, unclassified |
| LOC_Os01g63040 | expressed protein |
| LOC_Os02g04100 | peptidase, T1 family, putative, expressed |
| LOC_Os02g04110 | expressed protein |
| LOC_Os02g04120 | cytidylyltransferase domain containing protein, expressed |
| LOC_Os02g04130 | DUF1645 domain containing protein, putative, expressed |
| LOC_Os02g16050 | transposon protein, putative, unclassified, expressed |
| LOC_Os02g26180 | transposon protein, putative, CACTA, En/Spm sub-class |

| | |
|---|---|
| LOC_Os02g37230 | expressed protein |
| LOC_Os02g39090 | expressed protein |
| LOC_Os02g42430 | GEM, putative, expressed |
| LOC_Os02g42440 | GEM, putative, expressed |
| LOC_Os02g43800 | retrotransposon protein, putative, Ty3-gypsy subclass, expressed |
| LOC_Os02g48290 | thioredoxin reductase 2, putative, expressed |
| LOC_Os03g27320 | steroid binding protein, putative, expressed |
| LOC_Os03g27340 | expressed protein |
| LOC_Os03g27350 | cysteine protease ATG4, putative, expressed |
| LOC_Os03g27360 | RING-H2 finger protein ATL5H precursor, putative, expressed |
| LOC_Os03g45744 | hypothetical protein |
| LOC_Os03g45750 | DUF260 domain containing protein, putative, expressed |
| LOC_Os03g45760 | expressed protein |
| LOC_Os03g45830 | OsSAUR15 - Auxin-responsive SAUR gene family member |
| LOC_Os03g45850 | OsSAUR16 - Auxin-responsive SAUR gene family member, expressed |
| LOC_Os03g45870 | expressed protein |
| LOC_Os03g45880 | expressed protein |
| LOC_Os03g58080 | BT1 family protein, putative, expressed |
| LOC_Os04g28290 | transposon protein, putative, unclassified, expressed |
| LOC_Os04g28310 | expressed protein |
| LOC_Os04g28480 | retrotransposon protein, putative, unclassified, expressed |
| LOC_Os04g28500 | expressed protein |
| LOC_Os04g28520 | male sterility protein, putative, expressed |
| LOC_Os04g28830 | RALFL34 - Rapid ALkalinization Factor RALF family protein precursor, putative, expressed |
| LOC_Os04g28840 | RALFL35 - Rapid ALkalinization Factor RALF family protein precursor, putative, expressed |
| LOC_Os04g33720 | glycosyl hydrolases, putative, expressed |
| LOC_Os04g40600 | peptide methionine sulfoxide reductase, putative, expressed |
| LOC_Os04g43840 | possible lysine decarboxylase domain containing protein, expressed |
| LOC_Os04g44190 | expressed protein |

| | |
|---|---|
| LOC_Os04g44200 | oxygen-evolving enhancer protein 3, chloroplast precursor, putative, expressed |
| LOC_Os04g44210 | expressed protein |
| LOC_Os04g44230 | cytokinin dehydrogenase precursor, putative, expressed |
| LOC_Os04g44240 | cytokinin-O-glucosyltransferase 3, putative, expressed |
| LOC_Os04g44520 | expressed protein |
| LOC_Os04g56900 | transferase family protein, putative, expressed |
| LOC_Os04g57260 | expressed protein |
| LOC_Os04g57300 | phosphatidylinositol 3- and 4-kinase family protein, putative, expressed |
| LOC_Os04g57310 | thiol-disulphide oxidoreductase DCC, putative, expressed |
| LOC_Os04g57400 | methylthioribose kinase, putative, expressed |
| LOC_Os04g57420 | reticulon domain containing protein, putative, expressed |
| LOC_Os04g57430 | uncharacterized protein At4g06744 precursor, putative, expressed |
| LOC_Os05g01880 | expressed protein |
| LOC_Os05g25940 | expressed protein |
| LOC_Os06g38590 | receptor-like protein kinase precursor, putative, expressed |
| LOC_Os06g38600 | expressed protein |
| LOC_Os07g05470 | AIR9 protein, putative, expressed |
| LOC_Os07g05680 | expressed protein |
| LOC_Os07g07640 | hypothetical protein |
| LOC_Os07g07646 | expressed protein |
| LOC_Os07g07780 | expressed protein |
| LOC_Os07g07790 | LTPL75 - Protease inhibitor/seed storage/LTP family protein precursor, expressed |
| LOC_Os07g07820 | expressed protein |
| LOC_Os07g37270 | MSP domain containing protein, putative, expressed |
| LOC_Os09g22110 | retrotransposon protein, putative, Ty3-gypsy subclass, expressed |
| LOC_Os12g09610 | expressed protein |
| LOC_Os12g09640 | protein phosphatase 2C, putative, expressed |

**Genes in indica-rufipogon_SNP**

| | |
|---|---|
| LOC_Os01g01540 | retrotransposon protein, putative, unclassified, expressed |
| LOC_Os01g01550 | transposon protein, putative, CACTA, En/Spm sub-class, expressed |
| LOC_Os01g01560 | transposon protein, putative, unclassified, expressed |
| LOC_Os01g02740 | retrotransposon protein, putative, unclassified, expressed |
| LOC_Os01g03610 | retrotransposon protein, putative, unclassified, expressed |
| LOC_Os01g03720 | MYB family transcription factor, putative, expressed |
| LOC_Os01g03830 | retrotransposon protein, putative, unclassified, expressed |
| LOC_Os01g03860 | retrotransposon protein, putative, unclassified, expressed |
| LOC_Os01g04110 | mediator of RNA polymerase II transcription subunit 22, putative, expressed |
| LOC_Os01g04250 | expressed protein |
| LOC_Os01g04290 | glycosyl hydrolase family 10 protein, putative, expressed |
| LOC_Os01g05320 | retrotransposon protein, putative, Ty1-copia subclass, expressed |
| LOC_Os01g06460 | uncharacterized Cys-rich domain containing protein, putative, expressed |
| LOC_Os01g06470 | expressed protein |
| LOC_Os01g06490 | OsSCP1 - Putative Serine Carboxypeptidase homologue, expressed |
| LOC_Os01g06500 | PHLOEM 2-LIKE A5, putative, expressed |
| LOC_Os01g06510 | arginyl-tRNA synthetase, putative, expressed |
| LOC_Os01g06520 | verticillium wilt disease resistance protein, putative, expressed |
| LOC_Os01g06530 | retrotransposon protein, putative, unclassified, expressed |
| LOC_Os01g06540 | PHD finger protein, putative, expressed |
| LOC_Os01g06550 | NF-X1-type zinc finger protein, putative, expressed |
| LOC_Os01g06560 | transcription factor HBP-1b, putative, expressed |
| LOC_Os01g06570 | expressed protein |
| LOC_Os01g06580 | fasciclin domain containing protein, expressed |
| LOC_Os01g06590 | zinc finger, C3HC4 type domain containing protein, expressed |
| LOC_Os01g07760 | phospholipase D, putative, expressed |
| LOC_Os01g09110 | retrotransposon protein, putative, unclassified |
| LOC_Os01g09246 | glycine-rich protein, putative, expressed |
| LOC_Os01g10030 | expressed protein |

| | |
|---|---|
| LOC_Os01g10160 | latency associated nuclear antigen, putative, expressed |
| LOC_Os01g10170 | expressed protein |
| LOC_Os01g10260 | expressed protein |
| LOC_Os01g11190 | expressed protein |
| LOC_Os01g11660 | GDSL-like lipase/acylhydrolase, putative, expressed |
| LOC_Os01g11820 | expressed protein |
| LOC_Os01g11840 | xylem cysteine proteinase 2 precursor, putative, expressed |
| LOC_Os01g12170 | retrotransposon protein, putative, unclassified, expressed |
| LOC_Os01g13840 | PE-PGRS family protein, putative, expressed |
| LOC_Os01g14410 | early light-induced protein, chloroplast precursor, putative, expressed |
| LOC_Os01g14420 | RWP-RK domain-containing protein, putative, expressed |
| LOC_Os01g14430 | DNA binding protein, putative, expressed |
| LOC_Os01g15070 | retrotransposon protein, putative, unclassified |
| LOC_Os01g15260 | dynein light chain type 1 domain containing protein, expressed |
| LOC_Os01g15270 | expressed protein |
| LOC_Os01g15290 | ribosomal L18p/L5e family protein, putative, expressed |
| LOC_Os01g15300 | zinc finger helicase family protein, putative, expressed |
| LOC_Os01g15310 | LSM domain containing protein, expressed |
| LOC_Os01g15320 | RALFL9 - Rapid ALkalinization Factor RALF family protein precursor, expressed |
| LOC_Os01g16050 | transposon protein, putative, unclassified, expressed |
| LOC_Os01g16160 | retrotransposon protein, putative, unclassified, expressed |
| LOC_Os01g16190 | 26S proteasome non-ATPase regulatory subunit 14, putative, expressed |
| LOC_Os01g17060 | expressed protein |
| LOC_Os01g18010 | hypothetical protein |
| LOC_Os01g18020 | retrotransposon protein, putative, unclassified |
| LOC_Os01g18250 | subtilase, putative, expressed |
| LOC_Os01g19010 | retrotransposon protein, putative, unclassified, expressed |
| LOC_Os01g19100 | retrotransposon protein, putative, Ty3-gypsy subclass, expressed |
| LOC_Os01g21760 | retrotransposon protein, putative, unclassified |

| | |
|---|---|
| LOC_Os01g25040 | transposon protein, putative, CACTA, En/Spm sub-class, expressed |
| LOC_Os01g25065 | ATP synthase subunit beta, putative, expressed |
| LOC_Os01g25300 | expressed protein |
| LOC_Os01g26290 | expressed protein |
| LOC_Os01g27270 | retrotransposon protein, putative, unclassified, expressed |
| LOC_Os01g29040 | retrotransposon protein, putative, unclassified, expressed |
| LOC_Os01g29507 | retrotransposon, putative, centromere-specific, expressed |
| LOC_Os01g31629 | expressed protein |
| LOC_Os01g32500 | retrotransposon protein, putative, unclassified, expressed |
| LOC_Os01g33136 | retrotransposon protein, putative, unclassified, expressed |
| LOC_Os01g34620 | OsGrx_S15 |
| LOC_Os01g35660 | retrotransposon protein, putative, Ty3-gypsy subclass, expressed |
| LOC_Os01g36460 | MYB family transcription factor, putative, expressed |
| LOC_Os01g37260 | OsFBX14 - F-box domain containing protein, expressed |
| LOC_Os01g40540 | lectin receptor-type protein kinase, putative, expressed |
| LOC_Os01g42890 | expressed protein |
| LOC_Os01g45040 | Leucine rich repeat N-terminal domain containing protein, putative, expressed |
| LOC_Os01g46070 | lactate/malate dehydrogenase, putative, expressed |
| LOC_Os01g46080 | GDSL-like lipase/acylhydrolase, putative, expressed |
| LOC_Os01g46530 | expressed protein |
| LOC_Os01g46570 | CTP synthase, putative, expressed |
| LOC_Os01g46580 | actin-related protein 2/3 complex subunit 2, putative, expressed |
| LOC_Os01g47256 | exosome complex exonuclease RRP40, putative, expressed |
| LOC_Os01g47440 | expressed protein |
| LOC_Os01g47500 | expressed protein |
| LOC_Os01g48000 | S-locus-like receptor protein kinase, putative, expressed |
| LOC_Os01g48525 | expressed protein |
| LOC_Os01g48810 | transcription initiation factor TFIID subunit 11, putative, expressed |
| LOC_Os01g48820 | expressed protein |

| | |
|---|---|
| LOC_Os01g48830 | expressed protein |
| LOC_Os01g50220 | retrotransposon protein, putative, LINE subclass, expressed |
| LOC_Os01g52000 | retrotransposon protein, putative, unclassified, expressed |
| LOC_Os01g53750 | glucan endo-1,3-beta-glucosidase precursor, putative, expressed |
| LOC_Os01g55200 | potassium channel KAT1, putative, expressed |
| LOC_Os01g58020 | ribulose bisphosphate carboxylase large chain precursor, putative, expressed |
| LOC_Os01g62660 | MYB family transcription factor, putative, expressed |
| LOC_Os01g62710 | transposon protein, putative, unclassified |
| LOC_Os01g62960 | hypothetical protein |
| LOC_Os01g63040 | expressed protein |
| LOC_Os01g63050 | expressed protein |
| LOC_Os01g65480 | dnaJ domain containing protein, expressed |
| LOC_Os01g68230 | expressed protein |
| LOC_Os01g68780 | expressed protein |
| LOC_Os01g70790 | SRC2 protein, putative, expressed |
| LOC_Os01g71020 | expressed protein |
| LOC_Os01g73250 | abscisic stress-ripening, putative, expressed |
| LOC_Os02g03840 | expressed protein |
| LOC_Os02g03850 | glycosyltransferase, putative, expressed |
| LOC_Os02g03860 | mitochondrial ATP synthase precursor, putative, expressed |
| LOC_Os02g03910 | OsFBX35 - F-box domain containing protein, expressed |
| LOC_Os02g03950 | zinc finger, C3HC4 type family protein, expressed |
| LOC_Os02g03960 | CPuORF1 - conserved peptide uORF-containing transcript, expressed |
| LOC_Os02g03990 | WD domain, G-beta repeat domain containing protein, expressed |
| LOC_Os02g04010 | cyclin-T1-1, putative, expressed |
| LOC_Os02g04020 | expressed protein |
| LOC_Os02g04030 | phosphatidylinositol transfer, putative, expressed |
| LOC_Os02g04040 | RecF/RecN/SMC N terminal domain containing protein, expressed |
| LOC_Os02g04100 | peptidase, T1 family, putative, expressed |

| | |
|---|---|
| LOC_Os02g04110 | expressed protein |
| LOC_Os02g04120 | cytidylyltransferase domain containing protein, expressed |
| LOC_Os02g04130 | DUF1645 domain containing protein, putative, expressed |
| LOC_Os02g04320 | WD domain, G-beta repeat domain containing protein, expressed |
| LOC_Os02g04325 | expressed protein |
| LOC_Os02g04330 | 5-AMP-activated protein kinase beta-1 subunit-related, putative, expressed |
| LOC_Os02g04340 | NIN-like protein 2, putative, expressed |
| LOC_Os02g04480 | cleavage and polyadenylation specificity factor, putative, expressed |
| LOC_Os02g04490 | histone acetyltransferase HAC1, putative, expressed |
| LOC_Os02g04500 | S-adenosyl-L-methionine-dependent methyltransferase mraW, putative, expressed |
| LOC_Os02g04510 | UOS1, putative, expressed |
| LOC_Os02g05610 | ZOS2-03 - C2H2 zinc finger protein, expressed |
| LOC_Os02g05780 | expressed protein |
| LOC_Os02g06700 | ribosomal protein, putative, expressed |
| LOC_Os02g06730 | expressed protein |
| LOC_Os02g06820 | transposon protein, putative, unclassified, expressed |
| LOC_Os02g06850 | expressed protein |
| LOC_Os02g06860 | OsMADS80 - MADS-box family gene with M-alpha type-box, expressed |
| LOC_Os02g07110 | OsSAUR6 - Auxin-responsive SAUR gene family member, expressed |
| LOC_Os02g07120 | HEAT repeat family protein, putative, expressed |
| LOC_Os02g07200 | protein-O-fucosyltransferase 1, putative, expressed |
| LOC_Os02g07210 | OTU-like cysteine protease family protein, putative, expressed |
| LOC_Os02g07230 | porphobilinogen deaminase, chloroplast precursor, putative, expressed |
| LOC_Os02g07240 | expressed protein |
| LOC_Os02g11665 | transposon protein, putative, unclassified, expressed |
| LOC_Os02g14150 | transposon protein, putative, unclassified, expressed |
| LOC_Os02g15340 | no apical meristem protein, putative, expressed |
| LOC_Os02g16050 | transposon protein, putative, unclassified, expressed |
| LOC_Os02g18400 | hypothetical protein |

| | |
|---|---|
| LOC_Os02g19530 | lectin-like receptor kinase, putative, expressed |
| LOC_Os02g19700 | hypothetical protein |
| LOC_Os02g20270 | expressed protein |
| LOC_Os02g20960 | hypothetical protein |
| LOC_Os02g20970 | C2 domain containing protein, putative, expressed |
| LOC_Os02g21030 | expressed protein |
| LOC_Os02g21040 | aspartic proteinase nepenthesin precursor, putative, expressed |
| LOC_Os02g21220 | expressed protein |
| LOC_Os02g21240 | OsFBX48 - F-box domain containing protein, expressed |
| LOC_Os02g21250 | clathrin adaptor complex small chain domain containing protein, expressed |
| LOC_Os02g26180 | transposon protein, putative, CACTA, En/Spm sub-class |
| LOC_Os02g26310 | expressed protein |
| LOC_Os02g26320 | fasciclin-like arabinogalactan precursor protein, putative, expressed |
| LOC_Os02g26330 | expressed protein |
| LOC_Os02g27110 | expressed protein |
| LOC_Os02g27210 | hypothetical protein |
| LOC_Os02g27870 | expressed protein |
| LOC_Os02g28980 | peptidyl-prolyl isomerase, putative, expressed |
| LOC_Os02g35610 | expressed protein |
| LOC_Os02g37230 | expressed protein |
| LOC_Os02g37862 | 60S ribosomal protein L6, putative, expressed |
| LOC_Os02g37870 | Brix domain containing protein, putative, expressed |
| LOC_Os02g37880 | cyclin G-associated kinase, putative, expressed |
| LOC_Os02g37910 | expressed protein |
| LOC_Os02g37920 | expressed protein |
| LOC_Os02g37930 | hypoxia-responsive family protein, putative, expressed |
| LOC_Os02g37940 | CRP12 - Cysteine-rich family protein precursor, expressed |
| LOC_Os02g39090 | expressed protein |
| LOC_Os02g39910 | B4-BTB1 - Bric-a-Brac, Tramtrack, Broad Complex BTB domain with B4 subfamily conserved sequence, expressed |

| | |
|---|---|
| LOC_Os02g41800 | auxin response factor, putative, expressed |
| LOC_Os02g41860 | aquaporin protein, putative, expressed |
| LOC_Os02g41954 | gibberellin 2-beta-dioxygenase 7, putative, expressed |
| LOC_Os02g42040 | MIF4G domain containing protein, putative, expressed |
| LOC_Os02g42430 | GEM, putative, expressed |
| LOC_Os02g42440 | GEM, putative, expressed |
| LOC_Os02g42585 | AP2 domain containing protein, expressed |
| LOC_Os02g42860 | DEAD-box ATP-dependent RNA helicase, putative, expressed |
| LOC_Os02g42920 | DTA2, putative, expressed |
| LOC_Os02g42950 | YABBY domain containing protein, putative, expressed |
| LOC_Os02g43030 | integral membrane protein, putative, expressed |
| LOC_Os02g43170 | B-box zinc finger family protein, putative, expressed |
| LOC_Os02g43800 | retrotransposon protein, putative, Ty3-gypsy subclass, expressed |
| LOC_Os02g47330 | expressed protein |
| LOC_Os02g47770 | ZF-HD protein dimerisation region containing protein, expressed |
| LOC_Os02g48290 | thioredoxin reductase 2, putative, expressed |
| LOC_Os02g49210 | expressed protein |
| LOC_Os02g55400 | ATPase 8, plasma membrane-type, putative, expressed |
| LOC_Os02g56880 | transcriptional corepressor LEUNIG, putative, expressed |
| LOC_Os02g57570 | MATE efflux family protein, putative, expressed |
| LOC_Os02g57750 | protein binding protein, putative, expressed |
| LOC_Os02g58600 | expressed protein |
| LOC_Os03g02280 | DUF584 domain containing protein, putative, expressed |
| LOC_Os03g06160 | BTBP1 - Bric-a-Brac, Tramtrack, Broad Complex BTB domain with pentapeptide repeats, expressed |
| LOC_Os03g06170 | hsp20/alpha crystallin family protein, putative, expressed |
| LOC_Os03g06180 | expressed protein |
| LOC_Os03g06190 | 3-5 exonuclease family protein, putative, expressed |
| LOC_Os03g06200 | aminotransferase, putative, expressed |
| LOC_Os03g08130 | expressed protein |

| | |
|---|---|
| LOC_Os03g08140 | tubulin--tyrosine ligase-like protein 12, putative, expressed |
| LOC_Os03g08220 | lipoxygenase protein, putative, expressed |
| LOC_Os03g08230 | sodium/calcium exchanger protein, putative, expressed |
| LOC_Os03g08240 | hypothetical protein |
| LOC_Os03g08250 | expressed protein |
| LOC_Os03g08260 | hypothetical protein |
| LOC_Os03g08270 | ataxin-2 C-terminal region family protein, expressed |
| LOC_Os03g10430 | expressed protein |
| LOC_Os03g13660 | expressed protein |
| LOC_Os03g15110 | expressed protein |
| LOC_Os03g18900 | retrotransposon protein, putative, Ty3-gypsy subclass, expressed |
| LOC_Os03g18910 | COBRA-like protein 7 precursor, putative, expressed |
| LOC_Os03g19660 | conserved hypothetical protein |
| LOC_Os03g21140 | RNA-binding zinc finger protein, putative, expressed |
| LOC_Os03g22990 | OsFBX84 - F-box domain containing protein, expressed |
| LOC_Os03g27230 | phospho-2-dehydro-3-deoxyheptonate aldolase, chloroplast precursor, putative, expressed |
| LOC_Os03g27250 | OsFBO14 - F-box and other domain containing protein, expressed |
| LOC_Os03g27260 | 40S ribosomal protein S6, putative, expressed |
| LOC_Os03g27280 | CAMK_CAMK_like |
| LOC_Os03g27320 | steroid binding protein, putative, expressed |
| LOC_Os03g27340 | expressed protein |
| LOC_Os03g27350 | cysteine protease ATG4, putative, expressed |
| LOC_Os03g27360 | RING-H2 finger protein ATL5H precursor, putative, expressed |
| LOC_Os03g27380 | hypothetical protein |
| LOC_Os03g27390 | CPuORF35 - conserved peptide uORF-containing transcript, expressed |
| LOC_Os03g27420 | retrotransposon protein, putative, unclassified, expressed |
| LOC_Os03g27440 | hypothetical protein |
| LOC_Os03g27450 | ADP-ribosylation factor, putative, expressed |
| LOC_Os03g42820 | LIM domain-containing protein, putative, expressed |

| | |
|---|---|
| LOC_Os03g42840 | calcineurin B, putative, expressed |
| LOC_Os03g42900 | KH domain containing protein, putative, expressed |
| LOC_Os03g44380 | 9-cis-epoxycarotenoid dioxygenase 1, chloroplast precursor, putative, expressed |
| LOC_Os03g44690 | expressed protein |
| LOC_Os03g44720 | expressed protein |
| LOC_Os03g45744 | hypothetical protein |
| LOC_Os03g45750 | DUF260 domain containing protein, putative, expressed |
| LOC_Os03g45760 | expressed protein |
| LOC_Os03g45830 | OsSAUR15 - Auxin-responsive SAUR gene family member |
| LOC_Os03g45850 | OsSAUR16 - Auxin-responsive SAUR gene family member, expressed |
| LOC_Os03g45870 | expressed protein |
| LOC_Os03g45880 | expressed protein |
| LOC_Os03g45890 | hypothetical protein |
| LOC_Os03g45900 | expressed protein |
| LOC_Os03g45960 | thaumatin, putative, expressed |
| LOC_Os03g49970 | expressed protein |
| LOC_Os03g51510 | ribosomal protein L6, putative, expressed |
| LOC_Os03g57120 | ferredoxin--NADP reductase, chloroplast precursor, putative, expressed |
| LOC_Os03g58050 | ribosomal protein S13p/S18e, putative, expressed |
| LOC_Os03g58080 | BT1 family protein, putative, expressed |
| LOC_Os03g59225 | expressed protein |
| LOC_Os03g60120 | AP2 domain containing protein, expressed |
| LOC_Os03g63010 | plastid terminal oxidase, putative, expressed |
| LOC_Os03g63210 | expressed protein |
| LOC_Os04g03884 | transposon protein, putative, CACTA, En/Spm sub-class, expressed |
| LOC_Os04g04650 | hypothetical protein |
| LOC_Os04g10890 | expressed protein |
| LOC_Os04g14840 | retrotransposon protein, putative, unclassified, expressed |
| LOC_Os04g18380 | cytochrome P450, putative, expressed |

| | |
|---|---|
| LOC_Os04g19250 | hypothetical protein |
| LOC_Os04g27680 | retrotransposon protein, putative, Ty3-gypsy subclass, expressed |
| LOC_Os04g28290 | transposon protein, putative, unclassified, expressed |
| LOC_Os04g28310 | expressed protein |
| LOC_Os04g28410 | expressed protein |
| LOC_Os04g28480 | retrotransposon protein, putative, unclassified, expressed |
| LOC_Os04g28500 | expressed protein |
| LOC_Os04g28520 | male sterility protein, putative, expressed |
| LOC_Os04g28710 | hypothetical protein |
| LOC_Os04g28720 | RALFL32 - Rapid ALkalinization Factor RALF family protein precursor, putative, expressed |
| LOC_Os04g28790 | hypothetical protein |
| LOC_Os04g28830 | RALFL34 - Rapid ALkalinization Factor RALF family protein precursor, putative, expressed |
| LOC_Os04g28840 | RALFL35 - Rapid ALkalinization Factor RALF family protein precursor, putative, expressed |
| LOC_Os04g28850 | RALFL36 - Rapid ALkalinization Factor RALF family protein precursor, expressed |
| LOC_Os04g28860 | expressed protein |
| LOC_Os04g33720 | glycosyl hydrolases, putative, expressed |
| LOC_Os04g40600 | peptide methionine sulfoxide reductase, putative, expressed |
| LOC_Os04g41430 | retrotransposon protein, putative, unclassified |
| LOC_Os04g42570 | AP2/EREBP transcription factor BABY BOOM, putative, expressed |
| LOC_Os04g43120 | expressed protein |
| LOC_Os04g43140 | DEAD-box ATP-dependent RNA helicase, putative, expressed |
| LOC_Os04g43840 | possible lysine decarboxylase domain containing protein, expressed |
| LOC_Os04g43960 | transposon protein, putative, CACTA, En/Spm sub-class, expressed |
| LOC_Os04g44020 | expressed protein |
| LOC_Os04g44050 | expressed protein |
| LOC_Os04g44070 | expressed protein |
| LOC_Os04g44090 | expressed protein |
| LOC_Os04g44110 | hydrolase, putative, expressed |
| LOC_Os04g44120 | collagen triple helix repeat, putative, expressed |

| | |
|---|---|
| LOC_Os04g44130 | DEF12 - Defensin and Defensin-like DEFL family, expressed |
| LOC_Os04g44190 | expressed protein |
| LOC_Os04g44200 | oxygen-evolving enhancer protein 3, chloroplast precursor, putative, expressed |
| LOC_Os04g44210 | expressed protein |
| LOC_Os04g44224 | brain acid soluble protein 1, putative, expressed |
| LOC_Os04g44230 | cytokinin dehydrogenase precursor, putative, expressed |
| LOC_Os04g44240 | cytokinin-O-glucosyltransferase 3, putative, expressed |
| LOC_Os04g44280 | OsRR5 type-A response regulator, expressed |
| LOC_Os04g44340 | expressed protein |
| LOC_Os04g44520 | expressed protein |
| LOC_Os04g44530 | expressed protein |
| LOC_Os04g44540 | mitochondrial carrier protein, putative, expressed |
| LOC_Os04g44550 | CHCH domain containing protein, expressed |
| LOC_Os04g44560 | expressed protein |
| LOC_Os04g44580 | expressed protein |
| LOC_Os04g44590 | expressed protein |
| LOC_Os04g44600 | CRP1 - Cysteine-rich family protein precursor, expressed |
| LOC_Os04g44610 | white-brown complex homolog protein 11, putative, expressed |
| LOC_Os04g44690 | expressed protein |
| LOC_Os04g44700 | WD-40 repeat family protein, putative, expressed |
| LOC_Os04g44870 | C2 domain containing protein, putative, expressed |
| LOC_Os04g44880 | expressed protein |
| LOC_Os04g44890 | tRNA-dihydrouridine synthase, putative, expressed |
| LOC_Os04g44920 | oxidoreductase, short chain dehydrogenase/reductase family domain containing protein, expressed |
| LOC_Os04g44924 | short-chain dehydrogenase/reductase, putative, expressed |
| LOC_Os04g44940 | hypothetical protein |
| LOC_Os04g45050 | hypothetical protein |
| LOC_Os04g45170 | leucine-rich repeat family protein, putative, expressed |
| LOC_Os04g45180 | EF hand family protein, expressed |

| | |
|---|---|
| LOC_Os04g45190 | MSP domain containing protein, expressed |
| LOC_Os04g45270 | aspartyl protease family protein, putative, expressed |
| LOC_Os04g45280 | histidine triad family protein, putative, expressed |
| LOC_Os04g45290 | glycosyl hydrolases, putative, expressed |
| LOC_Os04g45490 | elongation factor, putative, expressed |
| LOC_Os04g45500 | expressed protein |
| LOC_Os04g45510 | expressed protein |
| LOC_Os04g45600 | NDH-M H PLASTOQUINONE DEHY, putative, expressed |
| LOC_Os04g45620 | expressed protein |
| LOC_Os04g45630 | expressed protein |
| LOC_Os04g45640 | hypothetical protein |
| LOC_Os04g45670 | src homology-3 domain protein 3, putative, expressed |
| LOC_Os04g45700 | COBRA, putative, expressed |
| LOC_Os04g48370 | RNA polymerase IV subunit, putative, expressed |
| LOC_Os04g55560 | AP2 domain containing protein, expressed |
| LOC_Os04g55570 | plant protein of unknown function DUF869 domain containing protein, expressed |
| LOC_Os04g56090 | S-locus-like receptor protein kinase, putative, expressed |
| LOC_Os04g56510 | ABC1 family domain containing protein, putative, expressed |
| LOC_Os04g56520 | alpha-1,4-glucan-protein synthase, putative, expressed |
| LOC_Os04g56530 | STE_MEKK_ste11_MAP3K |
| LOC_Os04g56540 | expressed protein |
| LOC_Os04g56620 | molybdopterin biosynthesis protein CNX1, putative, expressed |
| LOC_Os04g56640 | mediator of RNA polymerase II transcription subunit 7, putative, expressed |
| LOC_Os04g56646 | MEE34, putative, expressed |
| LOC_Os04g56654 | BSD domain-containing protein, putative, expressed |
| LOC_Os04g56670 | S4 domain containing protein, putative, expressed |
| LOC_Os04g56680 | OsSAUR22 - Auxin-responsive SAUR gene family member, expressed |
| LOC_Os04g56810 | hypothetical protein |
| LOC_Os04g56840 | expressed protein |

| | |
|---|---|
| LOC_Os04g56850 | auxin response factor, putative, expressed |
| LOC_Os04g56900 | transferase family protein, putative, expressed |
| LOC_Os04g56939 | expressed protein |
| LOC_Os04g56950 | jasmonate O-methyltransferase, putative, expressed |
| LOC_Os04g56960 | expressed protein |
| LOC_Os04g56970 | tubulin/FtsZ domain containing protein, putative, expressed |
| LOC_Os04g56980 | SWIB/MDM2 domain containing protein, expressed |
| LOC_Os04g56990 | Myb-like DNA-binding domain containing protein, putative, expressed |
| LOC_Os04g56995 | expressed protein |
| LOC_Os04g57010 | zinc finger C-x8-C-x5-C-x3-H type family protein, expressed |
| LOC_Os04g57020 | expressed protein |
| LOC_Os04g57030 | VQ domain containing protein, putative, expressed |
| LOC_Os04g57140 | kinesin motor domain containing protein, putative, expressed |
| LOC_Os04g57150 | tafazzin, putative, expressed |
| LOC_Os04g57154 | expressed protein |
| LOC_Os04g57180 | flavonol synthase/flavanone 3-hydroxylase, putative, expressed |
| LOC_Os04g57190 | ubiquitin carboxyl-terminal hydrolase, family 1, putative, expressed |
| LOC_Os04g57200 | heavy metal transport/detoxification protein, putative, expressed |
| LOC_Os04g57210 | actin-6, putative, expressed |
| LOC_Os04g57220 | ubiquitin-conjugating enzyme, putative, expressed |
| LOC_Os04g57230 | regulatory protein RecX family protein, putative, expressed |
| LOC_Os04g57260 | expressed protein |
| LOC_Os04g57300 | phosphatidylinositol 3- and 4-kinase family protein, putative, expressed |
| LOC_Os04g57310 | thiol-disulphide oxidoreductase DCC, putative, expressed |
| LOC_Os04g57350 | EH domain-containing protein 1, putative, expressed |
| LOC_Os04g57360 | expressed protein |
| LOC_Os04g57370 | acyl-protein thioesterase, putative, expressed |
| LOC_Os04g57380 | acyl-protein thioesterase, putative, expressed |
| LOC_Os04g57400 | methylthioribose kinase, putative, expressed |

| | |
|---|---|
| LOC_Os04g57420 | reticulon domain containing protein, putative, expressed |
| LOC_Os04g57430 | uncharacterized protein At4g06744 precursor, putative, expressed |
| LOC_Os04g57480 | OTU-like cysteine protease family protein, putative, expressed |
| LOC_Os04g57500 | phosphatidate cytidylyltransferase, putative, expressed |
| LOC_Os04g57520 | UBX domain-containing protein, putative, expressed |
| LOC_Os04g57530 | transcription factor, putative, expressed |
| LOC_Os04g57540 | adenylate kinase, putative, expressed |
| LOC_Os04g57550 | amine oxidase, flavin-containing, domain containing protein, expressed |
| LOC_Os04g57560 | amine oxidase, flavin-containing, domain containing protein, expressed |
| LOC_Os04g57950 | dihydroorotate dihydrogenase protein, putative, expressed |
| LOC_Os04g57970 | expressed protein |
| LOC_Os04g58000 | B3 DNA binding domain containing protein, expressed |
| LOC_Os05g01880 | expressed protein |
| LOC_Os05g05830 | bifunctional protein folD, putative, expressed |
| LOC_Os05g05840 | tRNA synthetase class II core domain containing protein, expressed |
| LOC_Os05g05850 | expressed protein |
| LOC_Os05g05860 | retrotransposon protein, putative, unclassified, expressed |
| LOC_Os05g07680 | KIP1, putative, expressed |
| LOC_Os05g07690 | thioredoxin, putative, expressed |
| LOC_Os05g07710 | WD40-like domain containing protein, putative, expressed |
| LOC_Os05g09350 | expressed protein |
| LOC_Os05g09430 | expressed protein |
| LOC_Os05g09460 | hypothetical protein |
| LOC_Os05g25940 | expressed protein |
| LOC_Os05g30790 | CW-type Zinc Finger, putative, expressed |
| LOC_Os05g33830 | zinc finger, C3HC4 type domain containing protein, expressed |
| LOC_Os05g33840 | transketolase, putative, expressed |
| LOC_Os05g33850 | UBA/TS-N domain containing protein, expressed |
| LOC_Os05g33860 | expressed protein |

| | |
|---|---|
| LOC_Os05g33870 | ran GTPase activating protein, putative, expressed |
| LOC_Os05g34040 | monooxygenase, putative, expressed |
| LOC_Os05g34050 | bZIP transcription factor domain containing protein, expressed |
| LOC_Os05g34060 | expressed protein |
| LOC_Os05g34070 | DIP1, putative, expressed |
| LOC_Os05g34100 | expressed protein |
| LOC_Os05g34870 | expressed protein |
| LOC_Os05g34880 | zinc finger, C3HC4 type family protein, expressed |
| LOC_Os05g34890 | expressed protein |
| LOC_Os05g35310 | ankyrin repeat family protein, putative, expressed |
| LOC_Os05g43280 | MATH domain containing protein, expressed |
| LOC_Os05g46920 | expressed protein |
| LOC_Os05g48370 | expressed protein |
| LOC_Os05g48380 | ubiquitin-conjugating enzyme domain containing protein, expressed |
| LOC_Os05g48390 | ubiquitin conjugating enzyme protein, putative, expressed |
| LOC_Os06g10190 | flavin-containing monooxygenase 2, putative, expressed |
| LOC_Os06g12730 | expressed protein |
| LOC_Os06g14220 | STF-1, putative, expressed |
| LOC_Os06g38590 | receptor-like protein kinase precursor, putative, expressed |
| LOC_Os06g38600 | expressed protein |
| LOC_Os06g44220 | OsRCI2-9 - Putative low temperature and salt responsive protein, expressed |
| LOC_Os06g51320 | GASR8 - Gibberellin-regulated GASA/GAST/Snakin family protein precursor, expressed |
| LOC_Os06g51350 | IPP transferase, putative, expressed |
| LOC_Os06g51360 | lysM domain containing protein, putative, expressed |
| LOC_Os07g02690 | DUF538 domain containing protein, putative, expressed |
| LOC_Os07g02710 | expressed protein |
| LOC_Os07g05380 | ATPase BadF/BadG/BcrA/BcrD type, putative, expressed |
| LOC_Os07g05470 | AIR9 protein, putative, expressed |
| LOC_Os07g05480 | photosystem I reaction center subunit, chloroplast precursor, putative, expressed |

| | |
|---|---|
| LOC_Os07g05570 | ERD4 protein, putative, expressed |
| LOC_Os07g05680 | expressed protein |
| LOC_Os07g05930 | expressed protein |
| LOC_Os07g05950 | expressed protein |
| LOC_Os07g05960 | expressed protein |
| LOC_Os07g07480 | HNH endonuclease domain-containing protein, putative, expressed |
| LOC_Os07g07490 | YT521-B, putative, expressed |
| LOC_Os07g07510 | expressed protein |
| LOC_Os07g07540 | SHOOT1 protein, putative, expressed |
| LOC_Os07g07550 | galactose-1-phosphate uridyl transferase, putative, expressed |
| LOC_Os07g07590 | expressed protein |
| LOC_Os07g07610 | RNA polymerase subunit, putative, expressed |
| LOC_Os07g07620 | pentatricopeptide, putative, expressed |
| LOC_Os07g07630 | expressed protein |
| LOC_Os07g07640 | hypothetical protein |
| LOC_Os07g07646 | expressed protein |
| LOC_Os07g07709 | ribosomal protein S13p/S18e, putative, expressed |
| LOC_Os07g07730 | hypothetical protein |
| LOC_Os07g07740 | expressed protein |
| LOC_Os07g07760 | expressed protein |
| LOC_Os07g07780 | expressed protein |
| LOC_Os07g07790 | LTPL75 - Protease inhibitor/seed storage/LTP family protein precursor, expressed |
| LOC_Os07g07820 | expressed protein |
| LOC_Os07g07920 | LTPL100 - Protease inhibitor/seed storage/LTP family protein precursor, expressed |
| LOC_Os07g07930 | LTPL78 - Protease inhibitor/seed storage/LTP family protein precursor, expressed |
| LOC_Os07g07940 | expressed protein |
| LOC_Os07g08510 | expressed protein |
| LOC_Os07g08530 | auxin response factor, putative, expressed |
| LOC_Os07g09675 | expressed protein |

| | |
|---|---|
| LOC_Os07g22890 | expressed protein |
| LOC_Os07g23460 | retrotransposon protein, putative, Ty1-copia subclass, expressed |
| LOC_Os07g26595 | expressed protein |
| LOC_Os07g37270 | MSP domain containing protein, putative, expressed |
| LOC_Os07g39060 | expressed protein |
| LOC_Os07g39070 | 5-formyltetrahydrofolate cyclo-ligase, putative, expressed |
| LOC_Os07g39080 | expressed protein |
| LOC_Os07g39090 | pentatricopeptide repeat domain containing protein, putative, expressed |
| LOC_Os07g39110 | AP2/EREBP transcription factor BABY BOOM, putative, expressed |
| LOC_Os07g45160 | SAC3/GANP family protein, expressed |
| LOC_Os07g45170 | protein phosphatase 2C, putative, expressed |
| LOC_Os07g45430 | expressed protein |
| LOC_Os07g47360 | CW-type Zinc Finger, putative, expressed |
| LOC_Os07g47370 | pentatricopeptide, putative, expressed |
| LOC_Os07g47380 | expressed protein |
| LOC_Os07g47410 | WD40-like domain containing protein, putative, expressed |
| LOC_Os08g34249 | inhibitor I family protein, putative, expressed |
| LOC_Os08g34270 | expressed protein |
| LOC_Os08g37470 | 2-aminoethanethiol dioxygenase, putative, expressed |
| LOC_Os08g37490 | 14-3-3 protein, putative, expressed |
| LOC_Os08g37500 | expressed protein |
| LOC_Os08g37520 | pentatricopeptide, putative, expressed |
| LOC_Os08g37770 | expressed protein |
| LOC_Os08g37780 | NOL1/NOP2/sun family protein, putative, expressed |
| LOC_Os08g38690 | retrotransposon protein, putative, SINE subclass, expressed |
| LOC_Os08g38780 | core histone H2A/H2B/H3/H4, putative, expressed |
| LOC_Os08g44880 | expressed protein |
| LOC_Os09g21520 | DEAD-box ATP-dependent RNA helicase, putative, expressed |
| LOC_Os09g22040 | expressed protein |

| | |
|---|---|
| LOC_Os09g22110 | retrotransposon protein, putative, Ty3-gypsy subclass, expressed |
| LOC_Os09g22530 | expressed protein |
| LOC_Os09g23370 | proline-rich protein 6, putative, expressed |
| LOC_Os09g23390 | hypothetical protein |
| LOC_Os09g23440 | hypothetical protein |
| LOC_Os09g23450 | expressed protein |
| LOC_Os09g23460 | expressed protein |
| LOC_Os09g30280 | glycosyl transferase 8 domain containing protein, putative, expressed |
| LOC_Os09g30290 | expressed protein |
| LOC_Os09g30300 | expressed protein |
| LOC_Os09g34150 | NBS-LRR disease resistance protein, putative, expressed |
| LOC_Os09g37680 | expressed protein |
| LOC_Os09g37700 | proteophosphoglycan ppg4, putative, expressed |
| LOC_Os09g37720 | RNA recognition motif containing protein, expressed |
| LOC_Os09g37730 | expressed protein |
| LOC_Os09g37740 | ribosomal L18p/L5e family protein, putative, expressed |
| LOC_Os09g37750 | hypothetical protein |
| LOC_Os09g37760 | bromodomain domain containing protein, expressed |
| LOC_Os09g38239 | sulfotransferase domain containing protein, expressed |
| LOC_Os10g35120 | expressed protein |
| LOC_Os10g35130 | expressed protein |
| LOC_Os10g35150 | expressed protein |
| LOC_Os11g03840 | D-mannose binding lectin family protein, expressed |
| LOC_Os11g03870 | LTPL31 - Protease inhibitor/seed storage/LTP family protein precursor, expressed |
| LOC_Os11g03900 | glucosamine--fructose-6-phosphate aminotransferase 1, putative, expressed |
| LOC_Os12g01820 | ATCHX, putative, expressed |
| LOC_Os12g09610 | expressed protein |
| LOC_Os12g09640 | protein phosphatase 2C, putative, expressed |
| LOC_Os12g09739 | pollen signalling protein with adenylyl cyclase activity, putative, expressed |

| | |
|---|---|
| LOC_Os12g40290 | expressed protein |
| LOC_Os12g40300 | SKP1, putative, expressed |
| LOC_Os12g40340 | EXS family protein, putative, expressed |
| LOC_Os12g40470 | DC1 domain-containing protein, putative, expressed |
| LOC_Os12g40500 | expressed protein |
| LOC_Os12g40540 | FAD dependent oxidoreductase, putative, expressed |
| LOC_Os12g40550 | peptidase, putative, expressed |
| LOC_Os12g40570 | WRKY94, expressed |
| LOC_Os12g40710 | helix-loop-helix DNA-binding domain containing protein, expressed |

**Genes in indica-rufipogon_SV**

| | |
|---|---|
| LOC_Os01g01270 | retrotransposon protein, putative, unclassified, expressed |
| LOC_Os01g01530 | transposon protein, putative, CACTA, En/Spm sub-class |
| LOC_Os01g01560 | transposon protein, putative, unclassified, expressed |
| LOC_Os01g01760 | expressed protein |
| LOC_Os01g01810 | expressed protein |
| LOC_Os01g02410 | protein kinase domain containing protein, expressed |
| LOC_Os01g02629 | retrotransposon protein, putative, unclassified, expressed |
| LOC_Os01g02740 | retrotransposon protein, putative, unclassified, expressed |
| LOC_Os01g04850 | retrotransposon protein, putative, unclassified, expressed |
| LOC_Os01g05320 | retrotransposon protein, putative, Ty1-copia subclass, expressed |
| LOC_Os01g06530 | retrotransposon protein, putative, unclassified, expressed |
| LOC_Os01g06610 | retrotransposon protein, putative, Ty3-gypsy subclass, expressed |
| LOC_Os01g06740 | ribosome inactivating protein, putative, expressed |
| LOC_Os01g07550 | hypothetical protein |
| LOC_Os01g08970 | SSRP1-like FACT complex subunit, putative, expressed |
| LOC_Os01g09000 | glutaminyl-tRNA synthetase, putative, expressed |
| LOC_Os01g09100 | WRKY10, expressed |
| LOC_Os01g09110 | retrotransposon protein, putative, unclassified |

| | |
|---|---|
| LOC_Os01g09960 | transposon protein, putative, unclassified, expressed |
| LOC_Os01g10030 | expressed protein |
| LOC_Os01g10610 | BES1/BZR1 homolog protein, putative, expressed |
| LOC_Os01g11054 | phosphoenolpyruvate carboxylase, putative, expressed |
| LOC_Os01g11190 | expressed protein |
| LOC_Os01g11660 | GDSL-like lipase/acylhydrolase, putative, expressed |
| LOC_Os01g13300 | B3 DNA binding domain containing protein, expressed |
| LOC_Os01g13620 | retrotransposon protein, putative, unclassified, expressed |
| LOC_Os01g13690 | ligA, putative, expressed |
| LOC_Os01g13720 | expressed protein |
| LOC_Os01g13730 | WD domain, G-beta repeat domain containing protein, expressed |
| LOC_Os01g13810 | expressed protein |
| LOC_Os01g14110 | expressed protein |
| LOC_Os01g14170 | expressed protein |
| LOC_Os01g14200 | retrotransposon protein, putative, unclassified, expressed |
| LOC_Os01g14470 | retrotransposon protein, putative, Ty3-gypsy subclass |
| LOC_Os01g14490 | retrotransposon protein, putative, unclassified, expressed |
| LOC_Os01g14550 | pathogen-related protein, putative, expressed |
| LOC_Os01g14610 | PSF2 - Putative GINS complex subunit, expressed |
| LOC_Os01g14650 | expansin precursor, putative, expressed |
| LOC_Os01g14670 | Cupin domain containing protein, expressed |
| LOC_Os01g14690 | Lg106, putative, expressed |
| LOC_Os01g15070 | retrotransposon protein, putative, unclassified |
| LOC_Os01g15200 | hypothetical protein |
| LOC_Os01g15210 | expressed protein |
| LOC_Os01g15220 | expressed protein |
| LOC_Os01g15229 | expressed protein |
| LOC_Os01g15240 | expressed protein |
| LOC_Os01g15250 | expressed protein |

| | |
|---|---|
| LOC_Os01g15260 | dynein light chain type 1 domain containing protein, expressed |
| LOC_Os01g15270 | expressed protein |
| LOC_Os01g15290 | ribosomal L18p/L5e family protein, putative, expressed |
| LOC_Os01g15300 | zinc finger helicase family protein, putative, expressed |
| LOC_Os01g15350 | zinc finger C-x8-C-x5-C-x3-H type family protein, expressed |
| LOC_Os01g15448 | expressed protein |
| LOC_Os01g15700 | transposon protein, putative, Pong sub-class, expressed |
| LOC_Os01g16180 | MT-A70 domain containing protein, expressed |
| LOC_Os01g17060 | expressed protein |
| LOC_Os01g17130 | retrotransposon protein, putative, Ty3-gypsy subclass, expressed |
| LOC_Os01g18010 | hypothetical protein |
| LOC_Os01g18020 | retrotransposon protein, putative, unclassified |
| LOC_Os01g18230 | FAD-linked oxidoreductase protein, putative, expressed |
| LOC_Os01g18250 | subtilase, putative, expressed |
| LOC_Os01g18980 | transposon protein, putative, unclassified |
| LOC_Os01g19780 | expressed protein |
| LOC_Os01g20060 | expressed protein |
| LOC_Os01g21360 | retrotransposon protein, putative, unclassified |
| LOC_Os01g21550 | expressed protein |
| LOC_Os01g23500 | expressed protein |
| LOC_Os01g23510 | expressed protein |
| LOC_Os01g23940 | expressed protein |
| LOC_Os01g24140 | transposon protein, putative, CACTA, En/Spm sub-class, expressed |
| LOC_Os01g24160 | hypothetical protein |
| LOC_Os01g24800 | retrotransposon protein, putative, unclassified, expressed |
| LOC_Os01g27070 | expressed protein |
| LOC_Os01g27320 | retrotransposon protein, putative, Ty3-gypsy subclass, expressed |
| LOC_Os01g27970 | hypothetical protein |
| LOC_Os01g28000 | retrotransposon protein, putative, Ty3-gypsy subclass, expressed |

| | |
|---|---|
| LOC_Os01g28200 | transposon protein, putative, unclassified, expressed |
| LOC_Os01g28270 | retrotransposon protein, putative, Ty1-copia subclass |
| LOC_Os01g29040 | retrotransposon protein, putative, unclassified, expressed |
| LOC_Os01g29570 | retrotransposon protein, putative, Ty3-gypsy subclass, expressed |
| LOC_Os01g29730 | retrotransposon protein, putative, Ty3-gypsy subclass |
| LOC_Os01g32184 | tetratricopeptide repeat protein 5, putative, expressed |
| LOC_Os01g32530 | transposon protein, putative, CACTA, En/Spm sub-class |
| LOC_Os01g32550 | hypothetical protein |
| LOC_Os01g33136 | retrotransposon protein, putative, unclassified, expressed |
| LOC_Os01g33180 | hypothetical protein |
| LOC_Os01g33740 | retrotransposon protein, putative, unclassified, expressed |
| LOC_Os01g33760 | expressed protein |
| LOC_Os01g33980 | expressed protein |
| LOC_Os01g33990 | transposon protein, putative, CACTA, En/Spm sub-class, expressed |
| LOC_Os01g35980 | expressed protein |
| LOC_Os01g37990 | expressed protein |
| LOC_Os01g38000 | conserved hypothetical protein |
| LOC_Os01g38650 | expressed protein |
| LOC_Os01g39136 | retrotransposon protein, putative, unclassified, expressed |
| LOC_Os01g40410 | transmembrane amino acid transporter protein, putative, expressed |
| LOC_Os01g40540 | lectin receptor-type protein kinase, putative, expressed |
| LOC_Os01g40880 | hypothetical protein |
| LOC_Os01g41030 | ribosomal protein L25, putative, expressed |
| LOC_Os01g41670 | G-patch domain containing protein, putative, expressed |
| LOC_Os01g41960 | retrotransposon protein, putative, unclassified, expressed |
| LOC_Os01g42400 | STE_MEK_ste7_MAP2K |
| LOC_Os01g42890 | expressed protein |
| LOC_Os01g43650 | WRKY11, expressed |
| LOC_Os01g45460 | serine esterase, putative, expressed |

| | |
|---|---|
| LOC_Os01g45470 | expressed protein |
| LOC_Os01g45540 | retrotransposon protein, putative, unclassified, expressed |
| LOC_Os01g45620 | CGMC_MAPKCMGC_2 |
| LOC_Os01g45624 | oleosin, putative, expressed |
| LOC_Os01g45640 | tat pathway signal sequence family protein, expressed |
| LOC_Os01g45650 | OsFBX19 - F-box domain containing protein, expressed |
| LOC_Os01g45659 | expressed protein |
| LOC_Os01g46250 | lipase, putative, expressed |
| LOC_Os01g46340 | chloroplast unusual positioning protein, putative, expressed |
| LOC_Os01g46360 | expressed protein |
| LOC_Os01g46530 | expressed protein |
| LOC_Os01g47300 | retrotransposon protein, putative, unclassified, expressed |
| LOC_Os01g48650 | expressed protein |
| LOC_Os01g50220 | retrotransposon protein, putative, LINE subclass, expressed |
| LOC_Os01g51820 | retrotransposon protein, putative, Ty3-gypsy subclass, expressed |
| LOC_Os01g52130 | sulfate transporter, putative, expressed |
| LOC_Os01g52920 | expressed protein |
| LOC_Os01g57942 | cytochrome b6, putative, expressed |
| LOC_Os01g57945 | cytochrome b6-f complex subunit 4, putative, expressed |
| LOC_Os01g57946 | DNA-directed RNA polymerase subunit alpha, putative, expressed |
| LOC_Os01g57956 | chloroplast 50S ribosomal protein L14, putative, expressed |
| LOC_Os01g58114 | expressed protein |
| LOC_Os01g59420 | expressed protein |
| LOC_Os01g60549 | retrotransposon protein, putative, unclassified, expressed |
| LOC_Os01g60850 | expressed protein |
| LOC_Os01g62290 | DnaK family protein, putative, expressed |
| LOC_Os01g62710 | transposon protein, putative, unclassified |
| LOC_Os01g62960 | hypothetical protein |
| LOC_Os01g63040 | expressed protein |

| | |
|---|---|
| LOC_Os01g63970 | sialyltransferase family domain containing protein, expressed |
| LOC_Os01g66650 | expressed protein |
| LOC_Os01g67410 | AP2/EREBP transcription factor BABY BOOM, putative, expressed |
| LOC_Os01g72140 | glutathione S-transferase, putative, expressed |
| LOC_Os01g73332 | retrotransposon protein, putative, Ty3-gypsy subclass |
| LOC_Os01g73900 | DEAD-box ATP-dependent RNA helicase, putative, expressed |
| LOC_Os01g74200 | protein kinase domain containing protein, expressed |
| LOC_Os02g04040 | RecF/RecN/SMC N terminal domain containing protein, expressed |
| LOC_Os02g04100 | peptidase, T1 family, putative, expressed |
| LOC_Os02g04110 | expressed protein |
| LOC_Os02g04120 | cytidylyltransferase domain containing protein, expressed |
| LOC_Os02g04130 | DUF1645 domain containing protein, putative, expressed |
| LOC_Os02g04880 | expressed protein |
| LOC_Os02g06980 | expressed protein |
| LOC_Os02g06990 | expressed protein |
| LOC_Os02g14420 | hypothetical protein |
| LOC_Os02g14929 | T-complex protein, putative, expressed |
| LOC_Os02g15050 | expressed protein |
| LOC_Os02g15240 | transposon protein, putative, unclassified, expressed |
| LOC_Os02g15380 | transposon protein, putative, Ac/Ds sub-class, expressed |
| LOC_Os02g16050 | transposon protein, putative, unclassified, expressed |
| LOC_Os02g18515 | retrotransposon protein, putative, unclassified, expressed |
| LOC_Os02g18910 | hypothetical protein |
| LOC_Os02g20520 | retrotransposon, putative, centromere-specific |
| LOC_Os02g20820 | retrotransposon protein, putative, unclassified, expressed |
| LOC_Os02g21160 | hypothetical protein |
| LOC_Os02g24920 | expressed protein |
| LOC_Os02g24930 | expressed protein |
| LOC_Os02g26180 | transposon protein, putative, CACTA, En/Spm sub-class |

| | |
|---|---|
| LOC_Os02g26190 | transposon protein, putative, CACTA, En/Spm sub-class, expressed |
| LOC_Os02g26500 | retrotransposon protein, putative, unclassified, expressed |
| LOC_Os02g27910 | expressed protein |
| LOC_Os02g28170 | transferase family protein, putative, expressed |
| LOC_Os02g28980 | peptidyl-prolyl isomerase, putative, expressed |
| LOC_Os02g29970 | expressed protein |
| LOC_Os02g29980 | X8 domain containing protein, expressed |
| LOC_Os02g34430 | serine/threonine-protein kinase, putative, expressed |
| LOC_Os02g35820 | retrotransposon protein, putative, unclassified, expressed |
| LOC_Os02g37230 | expressed protein |
| LOC_Os02g37654 | lecithin:cholesterol acyltransferase, putative, expressed |
| LOC_Os02g38090 | AP2 domain containing protein, expressed |
| LOC_Os02g38100 | hypothetical protein |
| LOC_Os02g39090 | expressed protein |
| LOC_Os02g39780 | retrotransposon protein, putative, unclassified, expressed |
| LOC_Os02g41520 | glycosyl transferase 8 domain containing protein, putative, expressed |
| LOC_Os02g41590 | kinase, pfkB family, putative, expressed |
| LOC_Os02g41630 | phenylalanine ammonia-lyase, putative, expressed |
| LOC_Os02g42430 | GEM, putative, expressed |
| LOC_Os02g42440 | GEM, putative, expressed |
| LOC_Os02g42990 | OsSAUR11 - Auxin-responsive SAUR gene family member, expressed |
| LOC_Os02g43800 | retrotransposon protein, putative, Ty3-gypsy subclass, expressed |
| LOC_Os02g45370 | retrotransposon protein, putative, Ty3-gypsy subclass, expressed |
| LOC_Os02g48290 | thioredoxin reductase 2, putative, expressed |
| LOC_Os02g51010 | membrane associated DUF588 domain containing protein, putative, expressed |
| LOC_Os02g53160 | tyrosine phosphatase family protein, putative, expressed |
| LOC_Os02g57860 | OsFBX71 - F-box domain containing protein, expressed |
| LOC_Os03g06530 | expressed protein |
| LOC_Os03g08170 | protein kinase APK1B, chloroplast precursor, putative, expressed |

| | |
|---|---|
| LOC_Os03g08180 | expressed protein |
| LOC_Os03g08190 | expressed protein |
| LOC_Os03g08200 | DUF567 domain containing protein, putative, expressed |
| LOC_Os03g08210 | hypothetical protein |
| LOC_Os03g08220 | lipoxygenase protein, putative, expressed |
| LOC_Os03g11800 | expressed protein |
| LOC_Os03g12730 | receptor protein kinase CLAVATA1 precursor, putative, expressed |
| LOC_Os03g13920 | transposon protein, putative, CACTA, En/Spm sub-class, expressed |
| LOC_Os03g16860 | DnaK family protein, putative, expressed |
| LOC_Os03g23940 | expressed protein |
| LOC_Os03g23950 | adapitin protein, putative, expressed |
| LOC_Os03g27320 | steroid binding protein, putative, expressed |
| LOC_Os03g27340 | expressed protein |
| LOC_Os03g27350 | cysteine protease ATG4, putative, expressed |
| LOC_Os03g27360 | RING-H2 finger protein ATL5H precursor, putative, expressed |
| LOC_Os03g30460 | vacuolar protein sorting-associated protein 52, putative, expressed |
| LOC_Os03g30470 | CHIT4 - Chitinase family protein precursor, expressed |
| LOC_Os03g30500 | hypothetical protein |
| LOC_Os03g32540 | retrotransposon protein, putative, Ty1-copia subclass, expressed |
| LOC_Os03g34030 | hypothetical protein |
| LOC_Os03g36070 | retrotransposon protein, putative, unclassified |
| LOC_Os03g36110 | expressed protein |
| LOC_Os03g36220 | retrotransposon protein, putative, unclassified, expressed |
| LOC_Os03g42130 | gibberellin 20 oxidase 2, putative, expressed |
| LOC_Os03g42440 | expressed protein |
| LOC_Os03g42490 | expressed protein |
| LOC_Os03g42510 | transposon protein, putative, CACTA, En/Spm sub-class, expressed |
| LOC_Os03g43114 | expressed protein |
| LOC_Os03g45330 | hsp20/alpha crystallin family protein, putative, expressed |

| | |
|---|---|
| LOC_Os03g45340 | hsp20/alpha crystallin family protein, putative, expressed |
| LOC_Os03g45344 | expressed protein |
| LOC_Os03g45350 | expressed protein |
| LOC_Os03g45710 | 2Fe-2S iron-sulfur cluster binding domain containing protein, expressed |
| LOC_Os03g45720 | glyoxalase family protein, putative, expressed |
| LOC_Os03g45730 | DNA-binding protein-like, putative, expressed |
| LOC_Os03g45744 | hypothetical protein |
| LOC_Os03g45750 | DUF260 domain containing protein, putative, expressed |
| LOC_Os03g45760 | expressed protein |
| LOC_Os03g45830 | OsSAUR15 - Auxin-responsive SAUR gene family member |
| LOC_Os03g45850 | OsSAUR16 - Auxin-responsive SAUR gene family member, expressed |
| LOC_Os03g45870 | expressed protein |
| LOC_Os03g45880 | expressed protein |
| LOC_Os03g46240 | hypothetical protein |
| LOC_Os03g46250 | acetyltransferase, GNAT family, putative, expressed |
| LOC_Os03g53010 | acyl-desaturase, chloroplast precursor, putative, expressed |
| LOC_Os03g56740 | expressed protein |
| LOC_Os03g56750 | expressed protein |
| LOC_Os03g58080 | BT1 family protein, putative, expressed |
| LOC_Os03g59030 | UDP-rhamnose rhamnosyltransferase, putative, expressed |
| LOC_Os03g59225 | expressed protein |
| LOC_Os04g09330 | expressed protein |
| LOC_Os04g09420 | hypothetical protein |
| LOC_Os04g14540 | expressed protein |
| LOC_Os04g15900 | retrotransposon protein, putative, Ty3-gypsy subclass, expressed |
| LOC_Os04g16832 | chloroplast 30S ribosomal protein S8, putative, expressed |
| LOC_Os04g16848 | photosystem II reaction center protein H, putative |
| LOC_Os04g23890 | AGC_PVPK_like_kin82y |
| LOC_Os04g25790 | expressed protein |

| | |
|---|---|
| LOC_Os04g28074 | expressed protein |
| LOC_Os04g28234 | Rf1, mitochondrial precursor, putative, expressed |
| LOC_Os04g28280 | BEE 3, putative, expressed |
| LOC_Os04g28290 | transposon protein, putative, unclassified, expressed |
| LOC_Os04g28310 | expressed protein |
| LOC_Os04g28480 | retrotransposon protein, putative, unclassified, expressed |
| LOC_Os04g28500 | expressed protein |
| LOC_Os04g28520 | male sterility protein, putative, expressed |
| LOC_Os04g28805 | expressed protein |
| LOC_Os04g28810 | expressed protein |
| LOC_Os04g28830 | RALFL34 - Rapid ALkalinization Factor RALF family protein precursor, putative, expressed |
| LOC_Os04g28840 | RALFL35 - Rapid ALkalinization Factor RALF family protein precursor, putative, expressed |
| LOC_Os04g29100 | hypothetical protein |
| LOC_Os04g29260 | expressed protein |
| LOC_Os04g29270 | retrotransposon protein, putative, unclassified, expressed |
| LOC_Os04g29280 | expressed protein |
| LOC_Os04g33720 | glycosyl hydrolases, putative, expressed |
| LOC_Os04g38680 | transmembrane amino acid transporter protein, putative, expressed |
| LOC_Os04g38700 | molybdopterin biosynthesis protein CNX3, putative, expressed |
| LOC_Os04g40600 | peptide methionine sulfoxide reductase, putative, expressed |
| LOC_Os04g41080 | zinc finger, C3HC4 type domain containing protein, expressed |
| LOC_Os04g42770 | expressed protein |
| LOC_Os04g43020 | protein kinase, putative, expressed |
| LOC_Os04g43030 | lipase class 3 family protein, putative, expressed |
| LOC_Os04g43324 | G-patch domain containing protein, putative, expressed |
| LOC_Os04g43340 | disease resistance RPP13-like protein 1, putative, expressed |
| LOC_Os04g43350 | expressed protein |
| LOC_Os04g43690 | DUF647 domain containing protein, putative, expressed |
| LOC_Os04g43710 | CAMK_CAMK_like |

| | |
|---|---|
| LOC_Os04g43770 | expressed protein |
| LOC_Os04g43824 | expressed protein |
| LOC_Os04g43840 | possible lysine decarboxylase domain containing protein, expressed |
| LOC_Os04g43880 | expressed protein |
| LOC_Os04g44190 | expressed protein |
| LOC_Os04g44200 | oxygen-evolving enhancer protein 3, chloroplast precursor, putative, expressed |
| LOC_Os04g44210 | expressed protein |
| LOC_Os04g44230 | cytokinin dehydrogenase precursor, putative, expressed |
| LOC_Os04g44240 | cytokinin-O-glucosyltransferase 3, putative, expressed |
| LOC_Os04g44460 | sulfotransferase domain containing protein, expressed |
| LOC_Os04g44470 | KUN1 - Kunitz-type trypsin inhibitor precursor, expressed |
| LOC_Os04g44520 | expressed protein |
| LOC_Os04g45350 | expressed protein |
| LOC_Os04g45560 | hypothetical protein |
| LOC_Os04g45570 | expressed protein |
| LOC_Os04g45740 | hypothetical protein |
| LOC_Os04g53050 | disease resistance RPP13-like protein 1, putative, expressed |
| LOC_Os04g56900 | transferase family protein, putative, expressed |
| LOC_Os04g57260 | expressed protein |
| LOC_Os04g57300 | phosphatidylinositol 3- and 4-kinase family protein, putative, expressed |
| LOC_Os04g57310 | thiol-disulphide oxidoreductase DCC, putative, expressed |
| LOC_Os04g57400 | methylthioribose kinase, putative, expressed |
| LOC_Os04g57420 | reticulon domain containing protein, putative, expressed |
| LOC_Os04g57430 | uncharacterized protein At4g06744 precursor, putative, expressed |
| LOC_Os05g01880 | expressed protein |
| LOC_Os05g02700 | expressed protein |
| LOC_Os05g02710 | retrotransposon protein, putative, unclassified, expressed |
| LOC_Os05g02720 | expressed protein |
| LOC_Os05g08980 | expressed protein |

| | |
|---|---|
| LOC_Os05g09050 | MLO domain containing protein, putative, expressed |
| LOC_Os05g21240 | expressed protein |
| LOC_Os05g22960 | expressed protein |
| LOC_Os05g22970 | glyoxalase family protein, putative, expressed |
| LOC_Os05g22990 | expressed protein |
| LOC_Os05g23000 | expressed protein |
| LOC_Os05g25940 | expressed protein |
| LOC_Os05g28490 | FAD-binding domain-containing protein, putative, expressed |
| LOC_Os05g34330 | retrotransposon protein, putative, unclassified, expressed |
| LOC_Os05g46920 | expressed protein |
| LOC_Os05g48430 | hypothetical protein |
| LOC_Os05g48470 | expressed protein |
| LOC_Os05g51710 | pleckstrin homology domain-containing protein-related taxo, putative, expressed |
| LOC_Os05g51760 | transposon protein, putative, CACTA, En/Spm sub-class, expressed |
| LOC_Os06g01050 | expressed protein |
| LOC_Os06g10660 | lysM domain-containing GPI-anchored protein 1 precursor, putative, expressed |
| LOC_Os06g10670 | aspartic proteinase nepenthesin-1 precursor, putative, expressed |
| LOC_Os06g10680 | expressed protein |
| LOC_Os06g10690 | PHD-finger domain containing protein, putative, expressed |
| LOC_Os06g11650 | phosphate-induced protein 1 conserved region domain containing protein, expressed |
| LOC_Os06g30750 | reticulon domain containing protein, putative, expressed |
| LOC_Os06g30770 | CAF1 family ribonuclease containing protein, putative, expressed |
| LOC_Os06g34700 | expressed protein |
| LOC_Os06g38590 | receptor-like protein kinase precursor, putative, expressed |
| LOC_Os06g38600 | expressed protein |
| LOC_Os06g39708 | photosystem II P680 chlorophyll A apoprotein, putative, expressed |
| LOC_Os06g39712 | OsClp10 - Putative Clp protease homologue, expressed |
| LOC_Os06g39716 | chloroplast 30S ribosomal protein S18, putative, expressed |
| LOC_Os06g39718 | chloroplast 50S ribosomal protein L33, putative, expressed |

| | |
|---|---|
| LOC_Os06g41360 | phosphoribosyl transferase, putative, expressed |
| LOC_Os06g41380 | expressed protein |
| LOC_Os06g41610 | retrotransposon protein, putative, unclassified, expressed |
| LOC_Os06g44510 | hypothetical protein |
| LOC_Os06g44830 | expressed protein |
| LOC_Os06g49920 | expressed protein |
| LOC_Os06g49930 | OsFBX207 - F-box domain containing protein, expressed |
| LOC_Os07g02730 | expressed protein |
| LOC_Os07g02740 | hypothetical protein |
| LOC_Os07g02770 | OsFBL32 - F-box domain and LRR containing protein, expressed |
| LOC_Os07g05470 | AIR9 protein, putative, expressed |
| LOC_Os07g05670 | expressed protein |
| LOC_Os07g05680 | expressed protein |
| LOC_Os07g05690 | expressed protein |
| LOC_Os07g05740 | receptor-like protein kinase 2 precursor, putative, expressed |
| LOC_Os07g05760 | expressed protein |
| LOC_Os07g07640 | hypothetical protein |
| LOC_Os07g07646 | expressed protein |
| LOC_Os07g07780 | expressed protein |
| LOC_Os07g07790 | LTPL75 - Protease inhibitor/seed storage/LTP family protein precursor, expressed |
| LOC_Os07g07820 | expressed protein |
| LOC_Os07g07974 | tesmin/TSO1-like CXC domain containing protein, expressed |
| LOC_Os07g09520 | expressed protein |
| LOC_Os07g09530 | ankyrin repeat family protein, putative, expressed |
| LOC_Os07g11020 | rc - bHLH transcription factor regulating proanthocyanidin production in seeds, expressed |
| LOC_Os07g11330 | RAL2 - Seed allergenic protein RA5/RA14/RA17 precursor, expressed |
| LOC_Os07g14514 | retrotransposon protein, putative, unclassified, expressed |
| LOC_Os07g14540 | vesicle-associated membrane protein, putative, expressed |
| LOC_Os07g14720 | expressed protein |

| | |
|---|---|
| LOC_Os07g18510 | OsFBLD5 - F-box, LRR and FBD domain containing protein, expressed |
| LOC_Os07g20290 | expressed protein |
| LOC_Os07g22024 | zinc finger family protein, putative, expressed |
| LOC_Os07g22730 | AP2 domain containing protein, expressed |
| LOC_Os07g22760 | conserved hypothetical protein |
| LOC_Os07g23100 | expressed protein |
| LOC_Os07g26530 | hypothetical protein |
| LOC_Os07g26990 | expressed protein |
| LOC_Os07g29600 | zinc finger, C3HC4 type, domain containing protein, expressed |
| LOC_Os07g31720 | GTPase activating protein, putative, expressed |
| LOC_Os07g31750 | chalcone synthase, putative, expressed |
| LOC_Os07g31930 | hypothetical protein |
| LOC_Os07g35050 | OsFBX237 - F-box domain containing protein, expressed |
| LOC_Os07g37270 | MSP domain containing protein, putative, expressed |
| LOC_Os07g39120 | expressed protein |
| LOC_Os07g39140 | expressed protein |
| LOC_Os07g39190 | expressed protein |
| LOC_Os07g39200 | hypothetical protein |
| LOC_Os08g02450 | ATCHX, putative, expressed |
| LOC_Os08g02600 | methyltransferase domain containing protein, expressed |
| LOC_Os08g13430 | expressed protein |
| LOC_Os08g13440 | cupin domain containing protein, expressed |
| LOC_Os08g13460 | expressed protein |
| LOC_Os08g14350 | expressed protein |
| LOC_Os08g29040 | protein kinase domain containing protein, expressed |
| LOC_Os08g33050 | MYB family transcription factor, putative, expressed |
| LOC_Os08g37610 | expressed protein |
| LOC_Os08g37630 | elastin precursor, putative, expressed |
| LOC_Os08g37650 | expressed protein |

| | |
|---|---|
| LOC_Os08g37660 | plastocyanin-like domain containing protein, putative, expressed |
| LOC_Os08g37680 | expressed protein |
| LOC_Os08g37830 | expressed protein |
| LOC_Os08g38170 | methyladenine glycosylase, putative, expressed |
| LOC_Os08g38480 | OsFBX300 - F-box domain containing protein, expressed |
| LOC_Os08g38650 | expressed protein |
| LOC_Os08g38670 | expressed protein |
| LOC_Os08g39250 | EDM2, putative, expressed |
| LOC_Os08g39270 | fasciclin domain containing protein, expressed |
| LOC_Os08g40790 | DNA repair ATPase-related, putative, expressed |
| LOC_Os08g42268 | expressed protein |
| LOC_Os08g42320 | expressed protein |
| LOC_Os09g03960 | expressed protein |
| LOC_Os09g08920 | cytochrome P450, putative, expressed |
| LOC_Os09g16850 | A3-BTB1 - Bric-a-Brac, Tramtrack, Broad Complex BTB domain with A3 subfamily conserved sequence, expressed |
| LOC_Os09g21580 | OsFBX324 - F-box domain containing protein, expressed |
| LOC_Os09g21710 | AN1-like zinc finger domain containing protein, expressed |
| LOC_Os09g21720 | expressed protein |
| LOC_Os09g22110 | retrotransposon protein, putative, Ty3-gypsy subclass, expressed |
| LOC_Os09g22440 | expressed protein |
| LOC_Os09g23500 | hypothetical protein |
| LOC_Os09g30170 | F-box protein interaction domain containing protein, expressed |
| LOC_Os10g08018 | expressed protein |
| LOC_Os10g08230 | retrotransposon, putative, centromere-specific, expressed |
| LOC_Os10g40920 | pentatricopeptide, putative, expressed |
| LOC_Os10g40934 | flavonol synthase/flavanone 3-hydroxylase, putative, expressed |
| LOC_Os10g40950 | polyol transporter 5, putative, expressed |
| LOC_Os10g42040 | expressed protein |
| LOC_Os11g03240 | MATE efflux family protein, putative, expressed |

| | |
|---|---|
| LOC_Os11g04130 | nodulin MtN21 family protein, putative, expressed |
| LOC_Os11g11620 | ZmEBE-1 protein, putative, expressed |
| LOC_Os11g22449 | expressed protein |
| LOC_Os11g24170 | CC-NBS-LRR protein, putative, expressed |
| LOC_Os11g39160 | NBS-LRR disease resistance protein, putative, expressed |
| LOC_Os12g09610 | expressed protein |
| LOC_Os12g09640 | protein phosphatase 2C, putative, expressed |
| LOC_Os12g09660 | expressed protein |
| LOC_Os12g20370 | expressed protein |
| LOC_Os12g26220 | expressed protein |
| LOC_Os12g33460 | transposon protein, putative, CACTA, En/Spm sub-class, expressed |
| LOC_Os12g40830 | kinase, pfkB family, putative, expressed |
| LOC_Os12g41050 | hypothetical protein |

**Shared genes in japonica-rufipogon comparision**

| | |
|---|---|
| LOC_Os01g02240 | transposon protein, putative, unclassified, expressed |
| LOC_Os01g05580 | Exo70 exocyst complex subunit family, putative, expressed |
| LOC_Os01g06810 | retrotransposon protein, putative, Ty1-copia subclass, expressed |
| LOC_Os01g08440 | UDP-glucoronosyl and UDP-glucosyl transferase domain containing protein, expressed |
| LOC_Os01g09060 | retrotransposon protein, putative, Ty1-copia subclass |
| LOC_Os01g10770 | expressed protein |
| LOC_Os01g15830 | peroxidase precursor, putative, expressed |
| LOC_Os01g19894 | transposon protein, putative, unclassified, expressed |
| LOC_Os01g27260 | glutathione S-transferase, putative, expressed |
| LOC_Os01g36390 | MCM4 - Putative minichromosome maintenance MCM complex subunit 4, expressed |
| LOC_Os01g37325 | retrotransposon protein, putative, unclassified, expressed |
| LOC_Os01g41430 | UDP-glucoronosyl and UDP-glucosyl transferase, putative, expressed |
| LOC_Os01g53440 | expressed protein |
| LOC_Os01g56120 | RNA recognition motif containing protein, putative, expressed |

| | |
|---|---|
| LOC_Os01g57599 | retrotransposon protein, putative, unclassified, expressed |
| LOC_Os01g63170 | transposon protein, putative, unclassified, expressed |
| LOC_Os02g16030 | harpin-induced protein 1 domain containing protein, expressed |
| LOC_Os02g16040 | ubiquitin-conjugating enzyme, putative, expressed |
| LOC_Os02g43800 | retrotransposon protein, putative, Ty3-gypsy subclass, expressed |
| LOC_Os02g45370 | retrotransposon protein, putative, Ty3-gypsy subclass, expressed |
| LOC_Os03g05560 | zinc finger, C3HC4 type domain containing protein, expressed |
| LOC_Os03g11840 | expressed protein |
| LOC_Os04g15760 | DEFL51 - Defensin and Defensin-like DEFL family, expressed |
| LOC_Os04g21880 | expressed protein |
| LOC_Os04g21890 | disease resistance protein RPM1, putative, expressed |
| LOC_Os04g21900 | expressed protein |
| LOC_Os04g30920 | expressed protein |
| LOC_Os04g42210 | GPI-anchored protein, putative, expressed |
| LOC_Os06g30470 | expressed protein |
| LOC_Os06g39800 | expressed protein |
| LOC_Os07g34990 | retrotransposon protein, putative, unclassified, expressed |
| LOC_Os07g38664 | expressed protein |
| LOC_Os08g12650 | retrotransposon protein, putative, Ty3-gypsy subclass, expressed |
| LOC_Os10g27070 | hypothetical protein |

**Genes in japonica-rufipogon_SNP**

| | |
|---|---|
| LOC_Os01g01560 | transposon protein, putative, unclassified, expressed |
| LOC_Os01g02240 | transposon protein, putative, unclassified, expressed |
| LOC_Os01g02500 | glutamyl-tRNA amidotransferase, putative, expressed |
| LOC_Os01g03830 | retrotransposon protein, putative, unclassified, expressed |
| LOC_Os01g05320 | retrotransposon protein, putative, Ty1-copia subclass, expressed |
| LOC_Os01g05580 | Exo70 exocyst complex subunit family, putative, expressed |
| LOC_Os01g06330 | retrotransposon protein, putative, unclassified, expressed |

| | |
|---|---|
| LOC_Os01g06430 | expressed protein |
| LOC_Os01g06810 | retrotransposon protein, putative, Ty1-copia subclass, expressed |
| LOC_Os01g07450 | MYB family transcription factor, putative, expressed |
| LOC_Os01g08440 | UDP-glucoronosyl and UDP-glucosyl transferase domain containing protein, expressed |
| LOC_Os01g09060 | retrotransposon protein, putative, Ty1-copia subclass |
| LOC_Os01g09110 | retrotransposon protein, putative, unclassified |
| LOC_Os01g09440 | expressed protein |
| LOC_Os01g09450 | OsIAA2 - Auxin-responsive Aux/IAA gene family member, expressed |
| LOC_Os01g09460 | hexokinase, putative, expressed |
| LOC_Os01g09850 | ZOS1-04 - C2H2 zinc finger protein, expressed |
| LOC_Os01g10030 | expressed protein |
| LOC_Os01g10770 | expressed protein |
| LOC_Os01g10850 | peroxidase precursor, putative, expressed |
| LOC_Os01g11200 | transforming protein myb, putative, expressed |
| LOC_Os01g12040 | transposon protein, putative, unclassified, expressed |
| LOC_Os01g12680 | C4-dicarboxylate transporter/malic acid transport protein, expressed |
| LOC_Os01g13690 | ligA, putative, expressed |
| LOC_Os01g13720 | expressed protein |
| LOC_Os01g13730 | WD domain, G-beta repeat domain containing protein, expressed |
| LOC_Os01g14340 | expressed protein |
| LOC_Os01g14360 | expressed protein |
| LOC_Os01g15260 | dynein light chain type 1 domain containing protein, expressed |
| LOC_Os01g15270 | expressed protein |
| LOC_Os01g15290 | ribosomal L18p/L5e family protein, putative, expressed |
| LOC_Os01g15300 | zinc finger helicase family protein, putative, expressed |
| LOC_Os01g15310 | LSM domain containing protein, expressed |
| LOC_Os01g15320 | RALFL9 - Rapid ALkalinization Factor RALF family protein precursor, expressed |
| LOC_Os01g15830 | peroxidase precursor, putative, expressed |
| LOC_Os01g15910 | UTP--glucose-1-phosphate uridylyltransferase, putative, expressed |

| | |
|---|---|
| LOC_Os01g18020 | retrotransposon protein, putative, unclassified |
| LOC_Os01g18250 | subtilase, putative, expressed |
| LOC_Os01g18730 | expressed protein |
| LOC_Os01g19310 | retrotransposon protein, putative, unclassified, expressed |
| LOC_Os01g19710 | transposon protein, putative, CACTA, En/Spm sub-class, expressed |
| LOC_Os01g19720 | transposon protein, putative, CACTA, En/Spm sub-class, expressed |
| LOC_Os01g19894 | transposon protein, putative, unclassified, expressed |
| LOC_Os01g21550 | expressed protein |
| LOC_Os01g21760 | retrotransposon protein, putative, unclassified |
| LOC_Os01g23960 | retrotransposon protein, putative, Ty3-gypsy subclass, expressed |
| LOC_Os01g25040 | transposon protein, putative, CACTA, En/Spm sub-class, expressed |
| LOC_Os01g25100 | glutathione S-transferase, putative, expressed |
| LOC_Os01g25330 | SAC9, putative, expressed |
| LOC_Os01g25340 | expressed protein |
| LOC_Os01g25360 | esterase, putative, expressed |
| LOC_Os01g25370 | SUMO protease, putative, expressed |
| LOC_Os01g27070 | expressed protein |
| LOC_Os01g27150 | cullin, putative, expressed |
| LOC_Os01g27160 | cullin, putative, expressed |
| LOC_Os01g27190 | C2 domain containing protein, putative, expressed |
| LOC_Os01g27230 | 12-oxophytodienoate reductase, putative, expressed |
| LOC_Os01g27260 | glutathione S-transferase, putative, expressed |
| LOC_Os01g27834 | hypothetical protein |
| LOC_Os01g28270 | retrotransposon protein, putative, Ty1-copia subclass |
| LOC_Os01g28540 | stress regulated protein, putative, expressed |
| LOC_Os01g28570 | pentatricopeptide, putative, expressed |
| LOC_Os01g29430 | tetratricopeptide-like helical, putative, expressed |
| LOC_Os01g29450 | retrotransposon protein, putative, unclassified, expressed |
| LOC_Os01g29507 | retrotransposon, putative, centromere-specific, expressed |

| | |
|---|---|
| LOC_Os01g29820 | expressed protein |
| LOC_Os01g31020 | retrotransposon protein, putative, unclassified, expressed |
| LOC_Os01g31430 | retrotransposon protein, putative, Ty3-gypsy subclass, expressed |
| LOC_Os01g32500 | retrotransposon protein, putative, unclassified, expressed |
| LOC_Os01g33030 | Brix domain containing protein, putative, expressed |
| LOC_Os01g34440 | expressed protein |
| LOC_Os01g34510 | hypothetical protein |
| LOC_Os01g36294 | cytochrome P450, putative, expressed |
| LOC_Os01g36330 | expressed protein |
| LOC_Os01g36340 | expressed protein |
| LOC_Os01g36390 | MCM4 - Putative minichromosome maintenance MCM complex subunit 4, expressed |
| LOC_Os01g36490 | hypothetical protein |
| LOC_Os01g36500 | TKL_IRAK_DUF26-lh |
| LOC_Os01g36510 | expressed protein |
| LOC_Os01g37325 | retrotransposon protein, putative, unclassified, expressed |
| LOC_Os01g37750 | glutathione S-transferase, putative, expressed |
| LOC_Os01g38110 | cytochrome P450, putative, expressed |
| LOC_Os01g39136 | retrotransposon protein, putative, unclassified, expressed |
| LOC_Os01g39970 | protein kinase domain containing protein, putative, expressed |
| LOC_Os01g41430 | UDP-glucoronosyl and UDP-glucosyl transferase, putative, expressed |
| LOC_Os01g43760 | cytochrome P450 72A1, putative, expressed |
| LOC_Os01g45040 | Leucine rich repeat N-terminal domain containing protein, putative, expressed |
| LOC_Os01g45060 | polygalacturonase, putative, expressed |
| LOC_Os01g46984 | expressed protein |
| LOC_Os01g47600 | HMG1/2, putative, expressed |
| LOC_Os01g47610 | lipase class 3 family protein, putative, expressed |
| LOC_Os01g47620 | expressed protein |
| LOC_Os01g47630 | expressed protein |
| LOC_Os01g47635 | expressed protein |

| | |
|---|---|
| LOC_Os01g48010 | expressed protein |
| LOC_Os01g48030 | expressed protein |
| LOC_Os01g48040 | S-locus-like receptor protein kinase, putative, expressed |
| LOC_Os01g48060 | auxin response factor, putative, expressed |
| LOC_Os01g48120 | expressed protein |
| LOC_Os01g48380 | PPR repeat domain containing protein, putative, expressed |
| LOC_Os01g48390 | expressed protein |
| LOC_Os01g48400 | hypothetical protein |
| LOC_Os01g48410 | ATROPGEF7/ROPGEF7, putative, expressed |
| LOC_Os01g48420 | peroxiredoxin, putative, expressed |
| LOC_Os01g52000 | retrotransposon protein, putative, unclassified, expressed |
| LOC_Os01g52920 | expressed protein |
| LOC_Os01g53440 | expressed protein |
| LOC_Os01g55710 | DNA-binding protein, putative, expressed |
| LOC_Os01g56120 | RNA recognition motif containing protein, putative, expressed |
| LOC_Os01g56530 | DUF260 domain containing protein, putative, expressed |
| LOC_Os01g56930 | eukaryotic aspartyl protease domain containing protein, expressed |
| LOC_Os01g56940 | flap endonuclease, putative, expressed |
| LOC_Os01g57150 | SR protein related family member, putative, expressed |
| LOC_Os01g57170 | expressed protein |
| LOC_Os01g57450 | universal stress protein domain containing protein, putative, expressed |
| LOC_Os01g57599 | retrotransposon protein, putative, unclassified, expressed |
| LOC_Os01g57954 | translation initiation factor IF-1, chloroplast, putative, expressed |
| LOC_Os01g57956 | chloroplast 50S ribosomal protein L14, putative, expressed |
| LOC_Os01g57958 | chloroplast 50S ribosomal protein L16, putative, expressed |
| LOC_Os01g58870 | domain of unknown function, DUF250 domain containing protein, expressed |
| LOC_Os01g60910 | CAMK_KIN1/SNF1/Nim1_like |
| LOC_Os01g61880 | respiratory burst oxidase, putative, expressed |
| LOC_Os01g63040 | expressed protein |

| | |
|---|---|
| LOC_Os01g63170 | transposon protein, putative, unclassified, expressed |
| LOC_Os01g65380 | patellin protein, putative, expressed |
| LOC_Os01g65400 | DNA polymerase I, putative, expressed |
| LOC_Os01g65410 | serine hydroxymethyltransferase, mitochondrial precursor, putative, expressed |
| LOC_Os01g65900 | chitin-inducible gibberellin-responsive protein, putative, expressed |
| LOC_Os01g65902 | apocytochrome f precursor, putative, expressed |
| LOC_Os01g65920 | F-box/LRR-repeat protein 2, putative, expressed |
| LOC_Os01g65940 | expressed protein |
| LOC_Os01g67100 | expressed protein |
| LOC_Os01g72400 | retrotransposon protein, putative, Ty1-copia subclass, expressed |
| LOC_Os01g72650 | RNA recognition motif containing protein, putative, expressed |
| LOC_Os01g73100 | retrotransposon protein, putative, unclassified, expressed |
| LOC_Os01g73590 | transporter family protein, putative, expressed |
| LOC_Os02g01180 | kinesin motor domain containing protein, putative, expressed |
| LOC_Os02g01190 | POEI25 - Pollen Ole e I allergen and extensin family protein precursor, expressed |
| LOC_Os02g01240 | CPuORF11 - conserved peptide uORF-containing transcript, expressed |
| LOC_Os02g01250 | LSM domain containing protein, expressed |
| LOC_Os02g01270 | START domain containing protein, expressed |
| LOC_Os02g01800 | receptor-like protein kinase precursor, putative, expressed |
| LOC_Os02g01820 | ABC2 homolog 4, putative, expressed |
| LOC_Os02g02150 | ATP-dependent helicase HRQ1, putative, expressed |
| LOC_Os02g02400 | catalase isozyme A, putative, expressed |
| LOC_Os02g03200 | FAD-binding and arabino-lactone oxidase domains containing protein, putative, expressed |
| LOC_Os02g03360 | expressed protein |
| LOC_Os02g03390 | expressed protein |
| LOC_Os02g03400 | microtubule associated protein, putative, expressed |
| LOC_Os02g08160 | OsFBD6 - F-box and FBD domain containing protein, expressed |
| LOC_Os02g08170 | GDU1, putative, expressed |
| LOC_Os02g08180 | protein transport protein SEC61 subunit gamma, putative, expressed |

| | |
|---|---|
| LOC_Os02g08190 | CYPRO4, putative, expressed |
| LOC_Os02g10240 | ZOS2-05 - C2H2 zinc finger protein, expressed |
| LOC_Os02g11000 | agenet domain containing protein, putative, expressed |
| LOC_Os02g11060 | WD domain, G-beta repeat domain containing protein, expressed |
| LOC_Os02g11950 | expressed protein |
| LOC_Os02g11970 | expressed protein |
| LOC_Os02g13800 | HSF-type DNA-binding domain containing protein, expressed |
| LOC_Os02g16030 | harpin-induced protein 1 domain containing protein, expressed |
| LOC_Os02g16040 | ubiquitin-conjugating enzyme, putative, expressed |
| LOC_Os02g16050 | transposon protein, putative, unclassified, expressed |
| LOC_Os02g16490 | expressed protein |
| LOC_Os02g22740 | retrotransposon protein, putative, unclassified, expressed |
| LOC_Os02g25624 | expressed protein |
| LOC_Os02g28530 | expressed protein |
| LOC_Os02g29770 | retrotransposon protein, putative, Ty3-gypsy subclass, expressed |
| LOC_Os02g29774 | expressed protein |
| LOC_Os02g29890 | expressed protein |
| LOC_Os02g29900 | expressed protein |
| LOC_Os02g33550 | harpin-induced protein 1 domain containing protein, expressed |
| LOC_Os02g33560 | expressed protein |
| LOC_Os02g33570 | expressed protein |
| LOC_Os02g34290 | expressed protein |
| LOC_Os02g37880 | cyclin G-associated kinase, putative, expressed |
| LOC_Os02g37910 | expressed protein |
| LOC_Os02g37920 | expressed protein |
| LOC_Os02g37930 | hypoxia-responsive family protein, putative, expressed |
| LOC_Os02g37940 | CRP12 - Cysteine-rich family protein precursor, expressed |
| LOC_Os02g42590 | WD-40 repeat family protein, putative, expressed |
| LOC_Os02g42600 | double-stranded RNA binding motif containing protein, expressed |

| | |
|---|---|
| LOC_Os02g43800 | retrotransposon protein, putative, Ty3-gypsy subclass, expressed |
| LOC_Os02g44720 | expressed protein |
| LOC_Os02g44730 | tetracycline transporter protein, putative, expressed |
| LOC_Os02g44740 | expressed protein |
| LOC_Os02g44750 | expressed protein |
| LOC_Os02g44810 | tRNA pseudouridine synthase family protein, putative, expressed |
| LOC_Os02g44820 | rho-GTPase-activating protein-related, putative, expressed |
| LOC_Os02g44830 | hypothetical protein |
| LOC_Os02g45370 | retrotransposon protein, putative, Ty3-gypsy subclass, expressed |
| LOC_Os02g47490 | expressed protein |
| LOC_Os02g47500 | nodulin, putative, expressed |
| LOC_Os02g47510 | 9-cis-epoxycarotenoid dioxygenase 1, chloroplast precursor, putative, expressed |
| LOC_Os02g47520 | expressed protein |
| LOC_Os02g48200 | lectin-like protein kinase, putative, expressed |
| LOC_Os02g48220 | expressed protein |
| LOC_Os02g48290 | thioredoxin reductase 2, putative, expressed |
| LOC_Os02g53030 | mitogen-activated protein kinase kinase kinase 1, putative, expressed |
| LOC_Os02g53040 | STE_MEKK_ste11_MAP3K |
| LOC_Os02g53050 | tetratricopeptide-like helical, putative, expressed |
| LOC_Os02g53670 | MYB family transcription factor, putative, expressed |
| LOC_Os02g57490 | DUF260 domain containing protein, putative, expressed |
| LOC_Os03g01240 | CCR4-Not complex component, Not1domain containing protein, expressed |
| LOC_Os03g01360 | expressed protein |
| LOC_Os03g01890 | START domain containing protein, expressed |
| LOC_Os03g01900 | 40S ribosomal protein S17, putative, expressed |
| LOC_Os03g01910 | transcription factor BTF3, putative, expressed |
| LOC_Os03g01920 | expressed protein |
| LOC_Os03g01930 | retrotransposon protein, putative, unclassified, expressed |
| LOC_Os03g02100 | valyl-tRNA synthetase, putative, expressed |

| | |
|---|---|
| LOC_Os03g02280 | DUF584 domain containing protein, putative, expressed |
| LOC_Os03g03550 | bZIP family transcription factor, putative, expressed |
| LOC_Os03g03890 | protein kinase family protein, putative, expressed |
| LOC_Os03g03930 | tRNA nucleotidyltransferase/polyA polymerase family protein, putative, expressed |
| LOC_Os03g04500 | tetratricopeptide repeat domain containing protein, expressed |
| LOC_Os03g05340 | retrotransposon protein, putative, unclassified, expressed |
| LOC_Os03g05480 | ZOS3-01 - C2H2 zinc finger protein, expressed |
| LOC_Os03g05490 | ZOS3-02 - C2H2 zinc finger protein, expressed |
| LOC_Os03g05500 | DUF260 domain containing protein, putative, expressed |
| LOC_Os03g05560 | zinc finger, C3HC4 type domain containing protein, expressed |
| LOC_Os03g06090 | high-affinity nickel-transport family protein, putative, expressed |
| LOC_Os03g06120 | PLAC8 family protein, expressed |
| LOC_Os03g07930 | retrotransposon protein, putative, Ty3-gypsy subclass, expressed |
| LOC_Os03g07950 | expressed protein |
| LOC_Os03g10060 | S10/S20 domain containing ribosomal protein, putative, expressed |
| LOC_Os03g10070 | transcription initiation factor IIB, putative, expressed |
| LOC_Os03g10080 | expressed protein |
| LOC_Os03g10110 | cupin domain containing protein, expressed |
| LOC_Os03g10140 | ZOS3-04 - C2H2 zinc finger protein, expressed |
| LOC_Os03g10150 | DUF623 domain containing protein, expressed |
| LOC_Os03g11840 | expressed protein |
| LOC_Os03g12530 | cation efflux family protein, putative, expressed |
| LOC_Os03g12550 | expressed protein |
| LOC_Os03g12790 | MATE efflux family protein, putative, expressed |
| LOC_Os03g13660 | expressed protein |
| LOC_Os03g17634 | PPR repeat domain containing protein, putative, expressed |
| LOC_Os03g18779 | expressed protein |
| LOC_Os03g21000 | thioredoxin, putative, expressed |
| LOC_Os03g21010 | expressed protein |

| | |
|---|---|
| LOC_Os03g21020 | RNA recognition motif containing protein, putative, expressed |
| LOC_Os03g21570 | DNA-binding protein, putative, expressed |
| LOC_Os03g21580 | expressed protein |
| LOC_Os03g21630 | PRC-barrel domain protein, putative, expressed |
| LOC_Os03g21720 | vacuolar-sorting receptor precursor, putative, expressed |
| LOC_Os03g21730 | receptor-like protein kinase precursor, putative, expressed |
| LOC_Os03g24960 | ulp1 protease family, C-terminal catalytic domain containing protein, expressed |
| LOC_Os03g24970 | SWIM zinc finger family protein, putative |
| LOC_Os03g30050 | protein kinase family protein, putative, expressed |
| LOC_Os03g33060 | retrotransposon protein, putative, unclassified, expressed |
| LOC_Os03g34320 | retrotransposon protein, putative, unclassified, expressed |
| LOC_Os03g44380 | 9-cis-epoxycarotenoid dioxygenase 1, chloroplast precursor, putative, expressed |
| LOC_Os03g44500 | serine/threonine protein phosphatase, putative, expressed |
| LOC_Os03g44760 | SWI1, putative, expressed |
| LOC_Os03g45170 | amino acid permease, putative, expressed |
| LOC_Os03g45370 | sodium/calcium exchanger protein, putative, expressed |
| LOC_Os03g45710 | 2Fe-2S iron-sulfur cluster binding domain containing protein, expressed |
| LOC_Os03g46940 | expressed protein |
| LOC_Os03g51610 | Inositol 1, 3, 4-trisphosphate 5/6-kinase, putative, expressed |
| LOC_Os03g51620 | DUF221 domain containing protein, expressed |
| LOC_Os03g51640 | expressed protein |
| LOC_Os03g53270 | stem-specific protein TSJT1, putative, expressed |
| LOC_Os03g53280 | WD domain containing protein, putative, expressed |
| LOC_Os03g55420 | peroxidase precursor, putative, expressed |
| LOC_Os03g57080 | PLA IIIA/PLP7, putative, expressed |
| LOC_Os03g57120 | ferredoxin--NADP reductase, chloroplast precursor, putative, expressed |
| LOC_Os03g58400 | C-5 cytosine-specific DNA methylase, putative, expressed |
| LOC_Os03g58410 | prenylated rab acceptor, putative, expressed |
| LOC_Os03g58420 | WRKY6, expressed |

| | |
|---|---|
| LOC_Os03g58720 | RNA recognition motif containing protein, putative, expressed |
| LOC_Os03g59225 | expressed protein |
| LOC_Os03g59330 | polygalacturonase, putative, expressed |
| LOC_Os03g59760 | RING finger protein 126, putative, expressed |
| LOC_Os03g59770 | EF hand family protein, putative, expressed |
| LOC_Os03g60250 | membrane associated DUF588 domain containing protein, putative, expressed |
| LOC_Os03g60910 | PPR repeat domain containing protein, putative, expressed |
| LOC_Os03g60920 | Methyltransferase small domain containing protein, expressed |
| LOC_Os03g60939 | dolichyl-phosphate beta-glycosyltransferase, putative, expressed |
| LOC_Os03g60950 | expressed protein |
| LOC_Os03g60960 | cytokinin-N-glucosyltransferase 1, putative, expressed |
| LOC_Os03g60976 | expressed protein |
| LOC_Os03g63300 | NLI interacting factor-like phosphatase, putative, expressed |
| LOC_Os04g02660 | retrotransposon protein, putative, unclassified |
| LOC_Os04g04650 | hypothetical protein |
| LOC_Os04g10480 | hypothetical protein |
| LOC_Os04g12390 | transposon protein, putative, CACTA, En/Spm sub-class, expressed |
| LOC_Os04g14840 | retrotransposon protein, putative, unclassified, expressed |
| LOC_Os04g15600 | receptor kinase, putative, expressed |
| LOC_Os04g15670 | expressed protein |
| LOC_Os04g15760 | DEFL51 - Defensin and Defensin-like DEFL family, expressed |
| LOC_Os04g16824 | chloroplast 30S ribosomal protein S3, putative, expressed |
| LOC_Os04g16832 | chloroplast 30S ribosomal protein S8, putative, expressed |
| LOC_Os04g19250 | hypothetical protein |
| LOC_Os04g21880 | expressed protein |
| LOC_Os04g21890 | disease resistance protein RPM1, putative, expressed |
| LOC_Os04g21900 | expressed protein |
| LOC_Os04g22510 | expressed protein |
| LOC_Os04g22800 | expressed protein |

| | |
|---|---|
| LOC_Os04g22820 | expressed protein |
| LOC_Os04g26330 | expressed protein |
| LOC_Os04g28520 | male sterility protein, putative, expressed |
| LOC_Os04g28530 | expressed protein |
| LOC_Os04g28540 | expressed protein |
| LOC_Os04g28630 | hypothetical protein |
| LOC_Os04g28710 | hypothetical protein |
| LOC_Os04g28720 | RALFL32 - Rapid ALkalinization Factor RALF family protein precursor, putative, expressed |
| LOC_Os04g28790 | hypothetical protein |
| LOC_Os04g28850 | RALFL36 - Rapid ALkalinization Factor RALF family protein precursor, expressed |
| LOC_Os04g28860 | expressed protein |
| LOC_Os04g30750 | expressed protein |
| LOC_Os04g30780 | eukaryotic translation initiation factor 3 subunit H, putative, expressed |
| LOC_Os04g30800 | ER lumen protein retaining receptor, putative, expressed |
| LOC_Os04g30810 | OsFBX129 - F-box domain containing protein, expressed |
| LOC_Os04g30880 | TBC domain containing protein, expressed |
| LOC_Os04g30890 | MYB family transcription factor, putative, expressed |
| LOC_Os04g30920 | expressed protein |
| LOC_Os04g34550 | expressed protein |
| LOC_Os04g34800 | hypothetical protein |
| LOC_Os04g34860 | expressed protein |
| LOC_Os04g36030 | AIG1, putative, expressed |
| LOC_Os04g36040 | peptide transporter PTR2, putative, expressed |
| LOC_Os04g40600 | peptide methionine sulfoxide reductase, putative, expressed |
| LOC_Os04g41470 | expressed protein |
| LOC_Os04g41480 | skin secretory protein xP2 precursor, putative, expressed |
| LOC_Os04g41490 | DNA-directed RNA polymerase III subunit RPC1, putative, expressed |
| LOC_Os04g42134 | enhancer of rudimentary protein, putative, expressed |
| LOC_Os04g42140 | eukaryotic initiation factor iso-4F subunit p82-34, putative, expressed |

| | |
|---|---|
| LOC_Os04g42150 | hypothetical protein |
| LOC_Os04g42160 | hypothetical protein |
| LOC_Os04g42174 | expressed protein |
| LOC_Os04g42210 | GPI-anchored protein, putative, expressed |
| LOC_Os04g42320 | AT hook motif family protein, expressed |
| LOC_Os04g42340 | expressed protein |
| LOC_Os04g42350 | heavy metal-associated domain containing protein, expressed |
| LOC_Os04g42360 | expressed protein |
| LOC_Os04g44020 | expressed protein |
| LOC_Os04g44170 | expressed protein |
| LOC_Os04g44230 | cytokinin dehydrogenase precursor, putative, expressed |
| LOC_Os04g44240 | cytokinin-O-glucosyltransferase 3, putative, expressed |
| LOC_Os04g47040 | anthocyanin regulatory Lc protein, putative, expressed |
| LOC_Os04g49260 | heavy metal-associated domain containing protein, expressed |
| LOC_Os04g49270 | tRNA-splicing endonuclease positive effector-related, putative, expressed |
| LOC_Os04g49280 | retrotransposon protein, putative, unclassified, expressed |
| LOC_Os04g49290 | retrotransposon protein, putative, unclassified, expressed |
| LOC_Os04g57550 | amine oxidase, flavin-containing, domain containing protein, expressed |
| LOC_Os04g57560 | amine oxidase, flavin-containing, domain containing protein, expressed |
| LOC_Os04g59120 | auxin-induced protein 5NG4, putative, expressed |
| LOC_Os05g01006 | retrotransposon protein, putative, unclassified, expressed |
| LOC_Os05g01880 | expressed protein |
| LOC_Os05g02435 | expressed protein |
| LOC_Os05g06230 | expressed protein |
| LOC_Os05g06610 | retrotransposon protein, putative, unclassified, expressed |
| LOC_Os05g07680 | KIP1, putative, expressed |
| LOC_Os05g07690 | thioredoxin, putative, expressed |
| LOC_Os05g07710 | WD40-like domain containing protein, putative, expressed |
| LOC_Os05g15490 | retrotransposon protein, putative, Ty3-gypsy subclass, expressed |

| | |
|---|---|
| LOC_Os05g31910 | white-brown complex homolog protein 16, putative, expressed |
| LOC_Os05g35150 | expressed protein |
| LOC_Os05g36320 | expressed protein |
| LOC_Os05g40820 | ribosomal protein L24, putative, expressed |
| LOC_Os05g43070 | expressed protein |
| LOC_Os05g45260 | expressed protein |
| LOC_Os05g45270 | ribosomal L9, putative, expressed |
| LOC_Os05g45280 | protein of unknown function DUF502 domain containing protein, expressed |
| LOC_Os05g45300 | enoyl-CoA hydratase/isomerase family protein, putative, expressed |
| LOC_Os05g45310 | ER lumen protein retaining receptor, putative, expressed |
| LOC_Os05g45320 | ubiquitin domain-containing protein 1, putative, expressed |
| LOC_Os05g49530 | F-box domain containing protein, expressed |
| LOC_Os06g04370 | expressed protein |
| LOC_Os06g05900 | methyltransferase, putative, expressed |
| LOC_Os06g05920 | Rf1, mitochondrial precursor, putative, expressed |
| LOC_Os06g05930 | expressed protein |
| LOC_Os06g05940 | DTA2, putative, expressed |
| LOC_Os06g06090 | CGMC_MAPKCMGC_2_ERK |
| LOC_Os06g09240 | anthocyanidin 3-O-glucosyltransferase, putative, expressed |
| LOC_Os06g14220 | STF-1, putative, expressed |
| LOC_Os06g14560 | expressed protein |
| LOC_Os06g14860 | hypothetical protein |
| LOC_Os06g21099 | expressed protein |
| LOC_Os06g30470 | expressed protein |
| LOC_Os06g35420 | retrotransposon protein, putative, Ty3-gypsy subclass, expressed |
| LOC_Os06g37400 | retrotransposon protein, putative, unclassified, expressed |
| LOC_Os06g39800 | expressed protein |
| LOC_Os06g39830 | expressed protein |
| LOC_Os06g39840 | retrotransposon protein, putative, unclassified, expressed |

| | |
|---|---|
| LOC_Os06g39850 | expressed protein |
| LOC_Os06g39860 | expressed protein |
| LOC_Os06g46540 | cell division cycle protein 23, putative, expressed |
| LOC_Os07g02420 | expressed protein |
| LOC_Os07g02430 | expressed protein |
| LOC_Os07g02450 | receptor-like protein kinase 2 precursor, putative, expressed |
| LOC_Os07g02460 | expressed protein |
| LOC_Os07g05084 | expressed protein |
| LOC_Os07g05120 | expressed protein |
| LOC_Os07g05260 | expressed protein |
| LOC_Os07g05380 | ATPase BadF/BadG/BcrA/BcrD type, putative, expressed |
| LOC_Os07g05410 | expressed protein |
| LOC_Os07g05470 | AIR9 protein, putative, expressed |
| LOC_Os07g05480 | photosystem I reaction center subunit, chloroplast precursor, putative, expressed |
| LOC_Os07g05500 | expressed protein |
| LOC_Os07g05520 | hypothetical protein |
| LOC_Os07g05530 | expressed protein |
| LOC_Os07g05930 | expressed protein |
| LOC_Os07g05950 | expressed protein |
| LOC_Os07g05960 | expressed protein |
| LOC_Os07g06820 | serine/threonine-protein kinase RIO-like, putative, expressed |
| LOC_Os07g08360 | expressed protein |
| LOC_Os07g08380 | expressed protein |
| LOC_Os07g10680 | polygalacturonase, putative, expressed |
| LOC_Os07g10720 | 40S ribosomal protein S15a, putative, expressed |
| LOC_Os07g12240 | EF hand family protein, putative, expressed |
| LOC_Os07g13090 | expressed protein |
| LOC_Os07g20660 | retrotransposon protein, putative, Ty3-gypsy subclass, expressed |
| LOC_Os07g28470 | coiled-coil domain-containing protein 111, putative, expressed |

| | |
|---|---|
| LOC_Os07g34990 | retrotransposon protein, putative, unclassified, expressed |
| LOC_Os07g36610 | CSLF9 - cellulose synthase-like family F; beta1,3;1,4 glucan synthase, expressed |
| LOC_Os07g36630 | CSLF8 - cellulose synthase-like family F; beta1,3;1,4 glucan synthase, expressed |
| LOC_Os07g38664 | expressed protein |
| LOC_Os07g45400 | expressed protein |
| LOC_Os08g12650 | retrotransposon protein, putative, Ty3-gypsy subclass, expressed |
| LOC_Os08g14630 | hypothetical protein |
| LOC_Os08g36390 | ZOS8-07 - C2H2 zinc finger protein, expressed |
| LOC_Os08g36410 | ANTH domain containing protein, expressed |
| LOC_Os08g36750 | NFKB, putative, expressed |
| LOC_Os08g36760 | remorin C-terminal domain containing protein, putative, expressed |
| LOC_Os08g37470 | 2-aminoethanethiol dioxygenase, putative, expressed |
| LOC_Os08g38690 | retrotransposon protein, putative, SINE subclass, expressed |
| LOC_Os08g38780 | core histone H2A/H2B/H3/H4, putative, expressed |
| LOC_Os08g39140 | heat shock protein, putative, expressed |
| LOC_Os08g43370 | 6-phosphogluconolactonase, putative, expressed |
| LOC_Os09g19720 | expressed protein |
| LOC_Os09g19734 | isochorismate synthase 1, chloroplast precursor, putative, expressed |
| LOC_Os09g19750 | expressed protein |
| LOC_Os09g20170 | expressed protein |
| LOC_Os09g20180 | expressed protein |
| LOC_Os09g22110 | retrotransposon protein, putative, Ty3-gypsy subclass, expressed |
| LOC_Os09g29470 | AGAP005770-PA, putative, expressed |
| LOC_Os09g30280 | glycosyl transferase 8 domain containing protein, putative, expressed |
| LOC_Os09g30290 | expressed protein |
| LOC_Os09g30300 | expressed protein |
| LOC_Os09g34150 | NBS-LRR disease resistance protein, putative, expressed |
| LOC_Os09g35960 | TBC domain containing protein, expressed |
| LOC_Os09g35980 | expressed protein |

| | |
|---|---|
| LOC_Os09g36380 | expressed protein |
| LOC_Os09g36440 | transcription initiation factor IIB, putative, expressed |
| LOC_Os09g37990 | expressed protein |
| LOC_Os09g38040 | hydrolase, NUDIX family protein, expressed |
| LOC_Os09g38050 | expressed protein |
| LOC_Os09g38060 | nicotinate-nucleotide pyrophosphorylase, putative, expressed |
| LOC_Os09g39060 | vignain precursor, putative, expressed |
| LOC_Os09g39170 | cysteine proteinase COT44 precursor, putative, expressed |
| LOC_Os09g39720 | 1-aminocyclopropane-1-carboxylate oxidase, putative, expressed |
| LOC_Os09g39730 | Core histone H2A/H2B/H3/H4 domain containing protein, putative, expressed |
| LOC_Os09g39740 | ACT domain containing protein, expressed |
| LOC_Os10g26780 | expressed protein |
| LOC_Os10g27070 | hypothetical protein |
| LOC_Os10g27460 | expressed protein |
| LOC_Os10g28450 | CAF1 family ribonuclease containing protein, expressed |
| LOC_Os10g28570 | expressed protein |
| LOC_Os10g28600 | PPR repeat domain containing protein, putative, expressed |
| LOC_Os10g29300 | expressed protein |
| LOC_Os10g40280 | expressed protein |
| LOC_Os11g02950 | auxin efflux carrier component, putative, expressed |
| LOC_Os11g02964 | FAR1 family protein, expressed |
| LOC_Os11g02990 | conserved hypothetical protein |
| LOC_Os11g03050 | cytidylyltransferase domain containing protein, expressed |
| LOC_Os11g03060 | SNF7 domain containing protein, putative, expressed |
| LOC_Os11g04390 | RNA recognition motif containing protein, expressed |
| LOC_Os11g06290 | expressed protein |
| LOC_Os11g06310 | integral membrane protein, putative, expressed |
| LOC_Os11g17340 | expressed protein |
| LOC_Os11g17640 | expressed protein |

| | |
|---|---|
| LOC_Os11g17650 | expressed protein |
| LOC_Os11g17660 | expressed protein |
| LOC_Os11g17940 | expressed protein |
| LOC_Os11g25900 | hAT dimerisation domain containing protein, putative |
| LOC_Os12g07740 | eukaryotic translation initiation factor 2 subunit gamma, putative, expressed |
| LOC_Os12g09739 | pollen signalling protein with adenylyl cyclase activity, putative, expressed |
| LOC_Os12g29605 | expressed protein |
| LOC_Os12g32470 | expressed protein |
| LOC_Os12g38110 | importin subunit beta, putative, expressed |
| LOC_Os12g38120 | thaumatin family domain containing protein, expressed |
| LOC_Os12g38140 | expressed protein |
| LOC_Os12g39480 | expressed protein |
| LOC_Os12g41550 | expressed protein |
| LOC_Os12g42290 | expressed protein |
| LOC_Os12g42840 | expressed protein |
| LOC_Os12g42860 | 2-aminoethanethiol dioxygenase, putative, expressed |
| LOC_Os12g42876 | 5-methyltetrahydropteroyltriglutamate--homocysteine methyltransferase, putative, expressed |
| LOC_Os12g42900 | lysine-rich arabinogalactan protein 19 precursor, putative, expressed |

**Genes in japonica-rufipogon_SV**

| | |
|---|---|
| LOC_Os01g01090 | retrotransposon protein, putative, unclassified, expressed |
| LOC_Os01g01100 | retrotransposon protein, putative, Ty3-gypsy subclass, expressed |
| LOC_Os01g01530 | transposon protein, putative, CACTA, En/Spm sub-class |
| LOC_Os01g01540 | retrotransposon protein, putative, unclassified, expressed |
| LOC_Os01g01550 | transposon protein, putative, CACTA, En/Spm sub-class, expressed |
| LOC_Os01g01910 | expressed protein |
| LOC_Os01g02240 | transposon protein, putative, unclassified, expressed |
| LOC_Os01g02910 | glycosyltransferase, putative, expressed |
| LOC_Os01g03200 | retrotransposon protein, putative, Ty1-copia subclass, expressed |

| | |
|---|---|
| LOC_Os01g03830 | retrotransposon protein, putative, unclassified, expressed |
| LOC_Os01g03860 | retrotransposon protein, putative, unclassified, expressed |
| LOC_Os01g04230 | inactive receptor kinase At2g26730 precursor, putative, expressed |
| LOC_Os01g04850 | retrotransposon protein, putative, unclassified, expressed |
| LOC_Os01g05240 | retrotransposon protein, putative, unclassified |
| LOC_Os01g05320 | retrotransposon protein, putative, Ty1-copia subclass, expressed |
| LOC_Os01g05580 | Exo70 exocyst complex subunit family, putative, expressed |
| LOC_Os01g06360 | transposon protein, putative, unclassified, expressed |
| LOC_Os01g06430 | expressed protein |
| LOC_Os01g06610 | retrotransposon protein, putative, Ty3-gypsy subclass, expressed |
| LOC_Os01g06620 | expressed protein |
| LOC_Os01g06810 | retrotransposon protein, putative, Ty1-copia subclass, expressed |
| LOC_Os01g08440 | UDP-glucoronosyl and UDP-glucosyl transferase domain containing protein, expressed |
| LOC_Os01g09060 | retrotransposon protein, putative, Ty1-copia subclass |
| LOC_Os01g09110 | retrotransposon protein, putative, unclassified |
| LOC_Os01g10030 | expressed protein |
| LOC_Os01g10195 | expressed protein |
| LOC_Os01g10480 | plastocyanin-like domain containing protein, putative, expressed |
| LOC_Os01g10490 | keratin, type I cytoskeletal 9, putative, expressed |
| LOC_Os01g10504 | OsMADS3 - MADS-box family gene with MIKCc type-box, expressed |
| LOC_Os01g10560 | transposon protein, putative, unclassified |
| LOC_Os01g10770 | expressed protein |
| LOC_Os01g11680 | transposon protein, putative, unclassified, expressed |
| LOC_Os01g11720 | transposable element protein, putative, containing Pfam profile: PF03004, Transposase_24, expressed |
| LOC_Os01g12040 | transposon protein, putative, unclassified, expressed |
| LOC_Os01g12340 | retrotransposon protein, putative, unclassified, expressed |
| LOC_Os01g13300 | B3 DNA binding domain containing protein, expressed |
| LOC_Os01g13620 | retrotransposon protein, putative, unclassified, expressed |
| LOC_Os01g13630 | retrotransposon protein, putative, unclassified, expressed |

| | |
|---|---|
| LOC_Os01g13810 | expressed protein |
| LOC_Os01g14170 | expressed protein |
| LOC_Os01g14200 | retrotransposon protein, putative, unclassified, expressed |
| LOC_Os01g14260 | transposon protein, putative, Ac/Ds sub-class |
| LOC_Os01g14270 | OsFBX4 - F-box domain containing protein, expressed |
| LOC_Os01g15070 | retrotransposon protein, putative, unclassified |
| LOC_Os01g15240 | expressed protein |
| LOC_Os01g15510 | expressed protein |
| LOC_Os01g15830 | peroxidase precursor, putative, expressed |
| LOC_Os01g16050 | transposon protein, putative, unclassified, expressed |
| LOC_Os01g16300 | hypothetical protein |
| LOC_Os01g17130 | retrotransposon protein, putative, Ty3-gypsy subclass, expressed |
| LOC_Os01g18020 | retrotransposon protein, putative, unclassified |
| LOC_Os01g18230 | FAD-linked oxidoreductase protein, putative, expressed |
| LOC_Os01g18250 | subtilase, putative, expressed |
| LOC_Os01g19010 | retrotransposon protein, putative, unclassified, expressed |
| LOC_Os01g19100 | retrotransposon protein, putative, Ty3-gypsy subclass, expressed |
| LOC_Os01g19110 | transposon protein, putative, CACTA, En/Spm sub-class |
| LOC_Os01g19894 | transposon protein, putative, unclassified, expressed |
| LOC_Os01g21120 | AP2 domain containing protein, expressed |
| LOC_Os01g21550 | expressed protein |
| LOC_Os01g22309 | hypothetical protein |
| LOC_Os01g23180 | retrotransposon protein, putative, unclassified |
| LOC_Os01g23239 | retrotransposon protein, putative, Ty3-gypsy subclass, expressed |
| LOC_Os01g24160 | hypothetical protein |
| LOC_Os01g25100 | glutathione S-transferase, putative, expressed |
| LOC_Os01g25600 | tetratricopeptide repeat domain containing protein, expressed |
| LOC_Os01g27070 | expressed protein |
| LOC_Os01g27230 | 12-oxophytodienoate reductase, putative, expressed |

| | |
|---|---|
| LOC_Os01g27260 | glutathione S-transferase, putative, expressed |
| LOC_Os01g27270 | retrotransposon protein, putative, unclassified, expressed |
| LOC_Os01g27320 | retrotransposon protein, putative, Ty3-gypsy subclass, expressed |
| LOC_Os01g27449 | expressed protein |
| LOC_Os01g27880 | sucrose-phosphatase, putative, expressed |
| LOC_Os01g27890 | cytochrome P450, putative, expressed |
| LOC_Os01g28134 | expressed protein |
| LOC_Os01g28200 | transposon protein, putative, unclassified, expressed |
| LOC_Os01g28270 | retrotransposon protein, putative, Ty1-copia subclass |
| LOC_Os01g28300 | OsFBX7 - F-box domain containing protein, expressed |
| LOC_Os01g28320 | retrotransposon, putative, centromere-specific |
| LOC_Os01g28570 | pentatricopeptide, putative, expressed |
| LOC_Os01g28580 | expressed protein |
| LOC_Os01g28590 | expressed protein |
| LOC_Os01g28600 | exo70 exocyst complex subunit family protein, putative, expressed |
| LOC_Os01g28940 | hypothetical protein |
| LOC_Os01g28970 | retrotransposon protein, putative, Ty3-gypsy subclass, expressed |
| LOC_Os01g28980 | proton-dependent oligopeptide transport, putative, expressed |
| LOC_Os01g29040 | retrotransposon protein, putative, unclassified, expressed |
| LOC_Os01g29570 | retrotransposon protein, putative, Ty3-gypsy subclass, expressed |
| LOC_Os01g29610 | expressed protein |
| LOC_Os01g31820 | expressed protein |
| LOC_Os01g32530 | transposon protein, putative, CACTA, En/Spm sub-class |
| LOC_Os01g32550 | hypothetical protein |
| LOC_Os01g33160 | stress responsive A/B Barrel domain containing protein, expressed |
| LOC_Os01g34060 | MYB family transcription factor, putative, expressed |
| LOC_Os01g34510 | hypothetical protein |
| LOC_Os01g34560 | 3-ketoacyl-CoA synthase, putative, expressed |
| LOC_Os01g34620 | OsGrx_S15 |

| | |
|---|---|
| LOC_Os01g35160 | TKL_IRAK_DUF26-lh |
| LOC_Os01g35170 | tRNA methyltransferase, putative, expressed |
| LOC_Os01g35184 | CAMK_KIN1/SNF1/Nim1_like |
| LOC_Os01g36390 | MCM4 - Putative minichromosome maintenance MCM complex subunit 4, expressed |
| LOC_Os01g36860 | DEAD-box ATP-dependent RNA helicase 40, putative, expressed |
| LOC_Os01g37050 | ribonuclease III domain protein, putative, expressed |
| LOC_Os01g37160 | retrotransposon protein, putative, unclassified |
| LOC_Os01g37170 | retrotransposon protein, putative, unclassified, expressed |
| LOC_Os01g37325 | retrotransposon protein, putative, unclassified, expressed |
| LOC_Os01g38400 | retrotransposon protein, putative, Ty3-gypsy subclass, expressed |
| LOC_Os01g39136 | retrotransposon protein, putative, unclassified, expressed |
| LOC_Os01g39340 | expressed protein |
| LOC_Os01g39350 | protein kinase domain containing protein, expressed |
| LOC_Os01g40260 | WRKY77, expressed |
| LOC_Os01g40499 | S-locus lectin protein kinase family protein, putative, expressed |
| LOC_Os01g41030 | ribosomal protein L25, putative, expressed |
| LOC_Os01g41430 | UDP-glucoronosyl and UDP-glucosyl transferase, putative, expressed |
| LOC_Os01g41440 | retrotransposon protein, putative, unclassified |
| LOC_Os01g43280 | hydroquinone glucosyltransferase, putative, expressed |
| LOC_Os01g43790 | transposon protein, putative, unclassified |
| LOC_Os01g43980 | retrotransposon protein, putative, unclassified, expressed |
| LOC_Os01g45000 | expressed protein |
| LOC_Os01g46169 | GDSL-like lipase/acylhydrolase, putative, expressed |
| LOC_Os01g46530 | expressed protein |
| LOC_Os01g47300 | retrotransposon protein, putative, unclassified, expressed |
| LOC_Os01g50220 | retrotransposon protein, putative, LINE subclass, expressed |
| LOC_Os01g53440 | expressed protein |
| LOC_Os01g54060 | expressed protein |
| LOC_Os01g54140 | expressed protein |

| | |
|---|---|
| LOC_Os01g54980 | expressed protein |
| LOC_Os01g56120 | RNA recognition motif containing protein, putative, expressed |
| LOC_Os01g56850 | expressed protein |
| LOC_Os01g57110 | SNF2 family N-terminal domain containing protein, expressed |
| LOC_Os01g57140 | hypothetical protein |
| LOC_Os01g57599 | retrotransposon protein, putative, unclassified, expressed |
| LOC_Os01g58830 | transposon protein, putative, unclassified, expressed |
| LOC_Os01g59230 | retrotransposon protein, putative, unclassified, expressed |
| LOC_Os01g59360 | CAMK_CAMK_like |
| LOC_Os01g59420 | expressed protein |
| LOC_Os01g60080 | monocopper oxidase, putative, expressed |
| LOC_Os01g61920 | Core histone H2A/H2B/H3/H4 domain containing protein, putative, expressed |
| LOC_Os01g62700 | expressed protein |
| LOC_Os01g62710 | transposon protein, putative, unclassified |
| LOC_Os01g62740 | expressed protein |
| LOC_Os01g63170 | transposon protein, putative, unclassified, expressed |
| LOC_Os01g63610 | retrotransposon protein, putative, unclassified, expressed |
| LOC_Os01g63970 | sialyltransferase family domain containing protein, expressed |
| LOC_Os01g64140 | endoglucanase 22 precursor, putative, expressed |
| LOC_Os01g64320 | expressed protein |
| LOC_Os01g64340 | expressed protein |
| LOC_Os01g64370 | expressed protein |
| LOC_Os01g64380 | expressed protein |
| LOC_Os01g66890 | BTBZ1 - Bric-a-Brac, Tramtrack, and Broad Complex BTB domain with TAZ zinc finger and Calmodulin-binding domains, expressed |
| LOC_Os01g66900 | expressed protein |
| LOC_Os01g66910 | expressed protein |
| LOC_Os01g66920 | Ser/Thr protein phosphatase family protein, putative, expressed |
| LOC_Os01g66930 | transposon protein, putative, CACTA, En/Spm sub-class, expressed |
| LOC_Os01g68340 | retrotransposon protein, putative, unclassified, expressed |

| | |
|---|---|
| LOC_Os01g68990 | retrotransposon, putative, centromere-specific, expressed |
| LOC_Os01g71870 | transposon protein, putative, unclassified, expressed |
| LOC_Os01g72200 | expressed protein |
| LOC_Os01g73332 | retrotransposon protein, putative, Ty3-gypsy subclass |
| LOC_Os02g01620 | transposon protein, putative, unclassified, expressed |
| LOC_Os02g03130 | expressed protein |
| LOC_Os02g03140 | inhibitor I family protein, putative, expressed |
| LOC_Os02g04400 | expressed protein |
| LOC_Os02g09270 | cytochrome P450 71A1, putative, expressed |
| LOC_Os02g12140 | retrotransposon protein, putative, Ty3-gypsy subclass, expressed |
| LOC_Os02g14600 | glutelin, putative, expressed |
| LOC_Os02g14610 | expressed protein |
| LOC_Os02g15360 | expressed protein |
| LOC_Os02g16030 | harpin-induced protein 1 domain containing protein, expressed |
| LOC_Os02g16040 | ubiquitin-conjugating enzyme, putative, expressed |
| LOC_Os02g16050 | transposon protein, putative, unclassified, expressed |
| LOC_Os02g16060 | RGA-1, putative, expressed |
| LOC_Os02g16320 | retrotransposon protein, putative, Ty3-gypsy subclass, expressed |
| LOC_Os02g16450 | expressed protein |
| LOC_Os02g16470 | retrotransposon protein, putative, Ty1-copia subclass, expressed |
| LOC_Os02g16500 | expressed protein |
| LOC_Os02g18064 | expressed protein |
| LOC_Os02g18400 | hypothetical protein |
| LOC_Os02g18410 | salt stress root protein RS1, putative, expressed |
| LOC_Os02g18420 | expressed protein |
| LOC_Os02g18515 | retrotransposon protein, putative, unclassified, expressed |
| LOC_Os02g18954 | GDSL-like lipase/acylhydrolase, putative, expressed |
| LOC_Os02g19010 | hypothetical protein |
| LOC_Os02g20040 | loricrin, putative, expressed |

| | |
|---|---|
| LOC_Os02g20210 | disease resistance protein RGA1, putative, expressed |
| LOC_Os02g21320 | expressed protein |
| LOC_Os02g21550 | flavonol synthase/flavanone 3-hydroxylase, putative, expressed |
| LOC_Os02g22200 | expressed protein |
| LOC_Os02g22770 | expressed protein |
| LOC_Os02g22780 | T-complex protein, putative, expressed |
| LOC_Os02g24205 | expressed protein |
| LOC_Os02g24420 | retrotransposon protein, putative, unclassified, expressed |
| LOC_Os02g25310 | zinc knuckle family protein, expressed |
| LOC_Os02g27220 | protein phosphatase 2C, putative, expressed |
| LOC_Os02g27340 | riboflavin biosynthesis protein ribD, putative, expressed |
| LOC_Os02g29450 | expressed protein |
| LOC_Os02g29464 | RecF/RecN/SMC N terminal domain containing protein, expressed |
| LOC_Os02g30760 | expressed protein |
| LOC_Os02g33800 | expressed protein |
| LOC_Os02g35250 | conserved hypothetical protein |
| LOC_Os02g37230 | expressed protein |
| LOC_Os02g38090 | AP2 domain containing protein, expressed |
| LOC_Os02g38100 | hypothetical protein |
| LOC_Os02g43460 | required to maintain repression 1, putative, expressed |
| LOC_Os02g43590 | ELMO/CED-12 family protein, putative, expressed |
| LOC_Os02g43800 | retrotransposon protein, putative, Ty3-gypsy subclass, expressed |
| LOC_Os02g44420 | conserved hypothetical protein |
| LOC_Os02g45280 | cytochrome P450, putative, expressed |
| LOC_Os02g45300 | hypothetical protein |
| LOC_Os02g45370 | retrotransposon protein, putative, Ty3-gypsy subclass, expressed |
| LOC_Os02g47470 | cytochrome P450, putative, expressed |
| LOC_Os02g47680 | retrotransposon protein, putative, Ty3-gypsy subclass |
| LOC_Os02g48260 | retrotransposon protein, putative, unclassified, expressed |

| | |
|---|---|
| LOC_Os02g48270 | DUF260 domain containing protein, putative |
| LOC_Os02g48280 | expressed protein |
| LOC_Os02g48290 | thioredoxin reductase 2, putative, expressed |
| LOC_Os02g48300 | OsFBX60 - F-box domain containing protein, expressed |
| LOC_Os02g49310 | AGC_PVPK_like_kin82y |
| LOC_Os02g49320 | sucrase-related, putative, expressed |
| LOC_Os02g50110 | expressed protein |
| LOC_Os02g50120 | retrotransposon protein, putative, Ty3-gypsy subclass |
| LOC_Os02g51270 | expressed protein |
| LOC_Os02g51970 | phosphate-induced protein 1 conserved region domain containing protein, expressed |
| LOC_Os02g51990 | expressed protein |
| LOC_Os02g52000 | phosphate-induced protein 1 conserved region domain containing protein, expressed |
| LOC_Os02g52650 | chlorophyll A-B binding protein, putative, expressed |
| LOC_Os02g53380 | expressed protein |
| LOC_Os02g53390 | expressed protein |
| LOC_Os02g53400 | thioredoxin, putative, expressed |
| LOC_Os02g53410 | expressed protein |
| LOC_Os02g53420 | DnaK family protein, putative, expressed |
| LOC_Os02g54120 | CCR4-NOT transcription factor, putative, expressed |
| LOC_Os03g01240 | CCR4-Not complex component, Not1domain containing protein, expressed |
| LOC_Os03g02514 | hydrolase, alpha/beta fold family protein, putative, expressed |
| LOC_Os03g02530 | OsRhmbd7 - Putative Rhomboid homologue, expressed |
| LOC_Os03g04169 | ATP phosphoribosyltransferase, putative, expressed |
| LOC_Os03g04630 | cytochrome P450 protein, putative, expressed |
| LOC_Os03g04760 | expressed protein |
| LOC_Os03g04840 | expressed protein |
| LOC_Os03g05370 | expressed protein |
| LOC_Os03g05560 | zinc finger, C3HC4 type domain containing protein, expressed |
| LOC_Os03g06120 | PLAC8 family protein, expressed |

| | |
|---|---|
| LOC_Os03g06139 | ABC-2 type transporter domain containing protein, expressed |
| LOC_Os03g09120 | expressed protein |
| LOC_Os03g09130 | expressed protein |
| LOC_Os03g09140 | ras-related protein, putative, expressed |
| LOC_Os03g09150 | pumilio-family RNA binding repeat domain containing protein, expressed |
| LOC_Os03g11840 | expressed protein |
| LOC_Os03g12510 | non-symbiotic hemoglobin 2, putative, expressed |
| LOC_Os03g12520 | STRUBBELIG-RECEPTOR FAMILY 7 precursor, putative, expressed |
| LOC_Os03g12660 | cytochrome P450, putative, expressed |
| LOC_Os03g13920 | transposon protein, putative, CACTA, En/Spm sub-class, expressed |
| LOC_Os03g16260 | protein kinase, putative, expressed |
| LOC_Os03g20060 | retrotransposon protein, putative, unclassified, expressed |
| LOC_Os03g21914 | ATP binding protein, putative, expressed |
| LOC_Os03g28180 | expressed protein |
| LOC_Os03g36070 | retrotransposon protein, putative, unclassified |
| LOC_Os03g36220 | retrotransposon protein, putative, unclassified, expressed |
| LOC_Os03g37430 | retrotransposon protein, putative, unclassified, expressed |
| LOC_Os03g38520 | expressed protein |
| LOC_Os03g38534 | expressed protein |
| LOC_Os03g38540 | folic acid binding protein, putative, expressed |
| LOC_Os03g44380 | 9-cis-epoxycarotenoid dioxygenase 1, chloroplast precursor, putative, expressed |
| LOC_Os03g50590 | expressed protein |
| LOC_Os03g50600 | expressed protein |
| LOC_Os03g52794 | phosphatidylinositol kinase, putative, expressed |
| LOC_Os03g54930 | VIP4, putative, expressed |
| LOC_Os03g54970 | expressed protein |
| LOC_Os03g57460 | fasciclin domain containing protein, expressed |
| LOC_Os03g57569 | expressed protein |
| LOC_Os03g60030 | transposon protein, putative, CACTA, En/Spm sub-class, expressed |

| | |
|---|---|
| LOC_Os03g61410 | expressed protein |
| LOC_Os03g61419 | expressed protein |
| LOC_Os04g02530 | expressed protein |
| LOC_Os04g09330 | expressed protein |
| LOC_Os04g12250 | retrotransposon protein, putative, Ty3-gypsy subclass, expressed |
| LOC_Os04g15760 | DEFL51 - Defensin and Defensin-like DEFL family, expressed |
| LOC_Os04g20424 | neural Wiskott-Aldrich syndrome protein, putative, expressed |
| LOC_Os04g20430 | expressed protein |
| LOC_Os04g21880 | expressed protein |
| LOC_Os04g21890 | disease resistance protein RPM1, putative, expressed |
| LOC_Os04g21900 | expressed protein |
| LOC_Os04g22210 | SCP-like extracellular protein, expressed |
| LOC_Os04g22220 | SCP-like extracellular protein, expressed |
| LOC_Os04g22230 | SCP-like extracellular protein, expressed |
| LOC_Os04g22440 | expressed protein |
| LOC_Os04g23190 | retrotransposon protein, putative, unclassified, expressed |
| LOC_Os04g23890 | AGC_PVPK_like_kin82y |
| LOC_Os04g25790 | expressed protein |
| LOC_Os04g27010 | hypothetical protein |
| LOC_Os04g27190 | terpene synthase, putative, expressed |
| LOC_Os04g28074 | expressed protein |
| LOC_Os04g28500 | expressed protein |
| LOC_Os04g28760 | expressed protein |
| LOC_Os04g30920 | expressed protein |
| LOC_Os04g41080 | zinc finger, C3HC4 type domain containing protein, expressed |
| LOC_Os04g42110 | cell division control protein 48 homolog B, putative, expressed |
| LOC_Os04g42120 | N-acetyltransferase ESCO2, putative, expressed |
| LOC_Os04g42130 | integral membrane transporter family protein, putative, expressed |
| LOC_Os04g42134 | enhancer of rudimentary protein, putative, expressed |

| | |
|---|---|
| LOC_Os04g42210 | GPI-anchored protein, putative, expressed |
| LOC_Os04g42470 | regulatory subunit, putative, expressed |
| LOC_Os04g42480 | receptor-like protein kinase At3g46290 precursor, putative, expressed |
| LOC_Os04g42770 | expressed protein |
| LOC_Os04g43840 | possible lysine decarboxylase domain containing protein, expressed |
| LOC_Os04g44460 | sulfotransferase domain containing protein, expressed |
| LOC_Os04g44470 | KUN1 - Kunitz-type trypsin inhibitor precursor, expressed |
| LOC_Os04g45560 | hypothetical protein |
| LOC_Os04g45570 | expressed protein |
| LOC_Os04g48160 | IQ calmodulin-binding motif family protein, putative, expressed |
| LOC_Os04g52640 | SHR5-receptor-like kinase, putative, expressed |
| LOC_Os04g55350 | retrotransposon protein, putative, Ty3-gypsy subclass, expressed |
| LOC_Os04g59160 | peroxidase precursor, putative, expressed |
| LOC_Os05g11820 | retrotransposon protein, putative, unclassified, expressed |
| LOC_Os05g16000 | expressed protein |
| LOC_Os05g21240 | expressed protein |
| LOC_Os05g22960 | expressed protein |
| LOC_Os05g22970 | glyoxalase family protein, putative, expressed |
| LOC_Os05g25940 | expressed protein |
| LOC_Os05g28490 | FAD-binding domain-containing protein, putative, expressed |
| LOC_Os05g31520 | expressed protein |
| LOC_Os05g34330 | retrotransposon protein, putative, unclassified, expressed |
| LOC_Os05g36370 | expressed protein |
| LOC_Os05g36380 | expressed protein |
| LOC_Os05g39730 | SAG20, putative, expressed |
| LOC_Os05g40720 | glycosyl transferase, family 8, putative, expressed |
| LOC_Os05g41310 | disease resistance protein RGA2, putative, expressed |
| LOC_Os05g41370 | TKL_IRAK_DUF26-la |
| LOC_Os05g43360 | NADH dehydrogenase flavoprotein 2, mitochondrial precursor, putative, expressed |

| | |
|---|---|
| LOC_Os05g44420 | expressed protein |
| LOC_Os05g44430 | hypothetical protein |
| LOC_Os05g45240 | flavin monooxygenase, putative, expressed |
| LOC_Os05g45260 | expressed protein |
| LOC_Os05g47880 | hypro1, putative, expressed |
| LOC_Os06g05750 | transferase family domain containing protein, expressed |
| LOC_Os06g05800 | RNA recognition motif containing protein, putative, expressed |
| LOC_Os06g05804 | trafficking protein particle complex subunit 4, putative, expressed |
| LOC_Os06g11970 | OsMADS63 - MADS-box family gene with MIKC* type-box, expressed |
| LOC_Os06g13410 | OsSCP33 - Putative Serine Carboxypeptidase homologue, expressed |
| LOC_Os06g30470 | expressed protein |
| LOC_Os06g39800 | expressed protein |
| LOC_Os06g49420 | retrotransposon protein, putative, unclassified, expressed |
| LOC_Os07g01600 | dirigent, putative, expressed |
| LOC_Os07g05740 | receptor-like protein kinase 2 precursor, putative, expressed |
| LOC_Os07g05760 | expressed protein |
| LOC_Os07g15610 | hypothetical protein |
| LOC_Os07g16280 | retrotransposon, putative, centromere-specific, expressed |
| LOC_Os07g20640 | hypothetical protein |
| LOC_Os07g26060 | expressed protein |
| LOC_Os07g34990 | retrotransposon protein, putative, unclassified, expressed |
| LOC_Os07g37890 | protein phosphatase 2C, putative, expressed |
| LOC_Os07g38664 | expressed protein |
| LOC_Os08g12650 | retrotransposon protein, putative, Ty3-gypsy subclass, expressed |
| LOC_Os08g32530 | expressed protein |
| LOC_Os08g35560 | expressed protein |
| LOC_Os08g36240 | expressed protein |
| LOC_Os08g36850 | hypothetical protein |
| LOC_Os08g38170 | methyladenine glycosylase, putative, expressed |

| | |
|---|---|
| LOC_Os09g07930 | retrotransposon protein, putative, unclassified, expressed |
| LOC_Os09g09820 | retrotransposon protein, putative, unclassified, expressed |
| LOC_Os09g21490 | expressed protein |
| LOC_Os09g26330 | hypro1, putative, expressed |
| LOC_Os09g26670 | expressed protein |
| LOC_Os10g27070 | hypothetical protein |
| LOC_Os11g03240 | MATE efflux family protein, putative, expressed |
| LOC_Os11g04130 | nodulin MtN21 family protein, putative, expressed |
| LOC_Os11g04630 | expressed protein |
| LOC_Os11g22449 | expressed protein |
| LOC_Os11g24360 | expressed protein |
| LOC_Os11g24374 | OsSCP55 - Putative Serine Carboxypeptidase homologue, expressed |
| LOC_Os12g29605 | expressed protein |
| LOC_Os12g39690 | hypothetical protein |
| LOC_Os12g41050 | hypothetical protein |
| LOC_Os12g42670 | expressed protein |
| LOC_Os12g42700 | expressed protein |

**Shared genes in indica-japonica comparision**

| | |
|---|---|
| LOC_Os01g01910 | expressed protein |
| LOC_Os01g02240 | transposon protein, putative, unclassified, expressed |
| LOC_Os01g02910 | glycosyltransferase, putative, expressed |
| LOC_Os01g05580 | Exo70 exocyst complex subunit family, putative, expressed |
| LOC_Os01g06360 | transposon protein, putative, unclassified, expressed |
| LOC_Os01g10770 | expressed protein |
| LOC_Os02g16030 | harpin-induced protein 1 domain containing protein, expressed |
| LOC_Os02g16040 | ubiquitin-conjugating enzyme, putative, expressed |
| LOC_Os02g43800 | retrotransposon protein, putative, Ty3-gypsy subclass, expressed |
| LOC_Os02g43960 | expressed protein |

| | |
|---|---|
| LOC_Os02g45370 | retrotransposon protein, putative, Ty3-gypsy subclass, expressed |
| LOC_Os02g46370 | expressed protein |
| LOC_Os02g47470 | cytochrome P450, putative, expressed |
| LOC_Os03g04840 | expressed protein |
| LOC_Os03g05560 | zinc finger, C3HC4 type domain containing protein, expressed |
| LOC_Os04g42210 | GPI-anchored protein, putative, expressed |
| LOC_Os08g12650 | retrotransposon protein, putative, Ty3-gypsy subclass, expressed |
| LOC_Os08g36240 | expressed protein |
| LOC_Os08g36850 | hypothetical protein |
| LOC_Os10g27070 | hypothetical protein |

**Genes in indica-japonica_SNP**

| | |
|---|---|
| LOC_Os01g01360 | peptide transporter PTR2, putative, expressed |
| LOC_Os01g01470 | no apical meristem protein, putative, expressed |
| LOC_Os01g01530 | transposon protein, putative, CACTA, En/Spm sub-class |
| LOC_Os01g01910 | expressed protein |
| LOC_Os01g02240 | transposon protein, putative, unclassified, expressed |
| LOC_Os01g02290 | Ser/Thr receptor-like kinase, putative, expressed |
| LOC_Os01g02910 | glycosyltransferase, putative, expressed |
| LOC_Os01g03860 | retrotransposon protein, putative, unclassified, expressed |
| LOC_Os01g05320 | retrotransposon protein, putative, Ty1-copia subclass, expressed |
| LOC_Os01g05580 | Exo70 exocyst complex subunit family, putative, expressed |
| LOC_Os01g06300 | transposon protein, putative, unclassified, expressed |
| LOC_Os01g06360 | transposon protein, putative, unclassified, expressed |
| LOC_Os01g08890 | expressed protein |
| LOC_Os01g08930 | DEAD-box ATP-dependent RNA helicase, putative, expressed |
| LOC_Os01g08940 | expressed protein |
| LOC_Os01g08950 | expressed protein |
| LOC_Os01g08970 | SSRP1-like FACT complex subunit, putative, expressed |

| | |
|---|---|
| LOC_Os01g09110 | retrotransposon protein, putative, unclassified |
| LOC_Os01g09220 | transposon protein, putative, CACTA, En/Spm sub-class, expressed |
| LOC_Os01g10030 | expressed protein |
| LOC_Os01g10090 | pentatricopeptide, putative, expressed |
| LOC_Os01g10770 | expressed protein |
| LOC_Os01g11200 | transforming protein myb, putative, expressed |
| LOC_Os01g11670 | OsSCP2 - Putative Serine Carboxypeptidase homologue, expressed |
| LOC_Os01g11910 | basic helix-loop-helix, putative, expressed |
| LOC_Os01g12170 | retrotransposon protein, putative, unclassified, expressed |
| LOC_Os01g12680 | C4-dicarboxylate transporter/malic acid transport protein, expressed |
| LOC_Os01g13300 | B3 DNA binding domain containing protein, expressed |
| LOC_Os01g13690 | ligA, putative, expressed |
| LOC_Os01g13720 | expressed protein |
| LOC_Os01g13730 | WD domain, G-beta repeat domain containing protein, expressed |
| LOC_Os01g13840 | PE-PGRS family protein, putative, expressed |
| LOC_Os01g14514 | expressed protein |
| LOC_Os01g14650 | expansin precursor, putative, expressed |
| LOC_Os01g14670 | Cupin domain containing protein, expressed |
| LOC_Os01g15910 | UTP--glucose-1-phosphate uridylyltransferase, putative, expressed |
| LOC_Os01g16030 | ADP-ribosylation factor, putative, expressed |
| LOC_Os01g16370 | NBS-LRR type disease resistance protein, putative, expressed |
| LOC_Os01g18250 | subtilase, putative, expressed |
| LOC_Os01g19010 | retrotransposon protein, putative, unclassified, expressed |
| LOC_Os01g19310 | retrotransposon protein, putative, unclassified, expressed |
| LOC_Os01g19670 | expressed protein |
| LOC_Os01g20850 | expressed protein |
| LOC_Os01g21760 | retrotransposon protein, putative, unclassified |
| LOC_Os01g22080 | expressed protein |
| LOC_Os01g22490 | 40S ribosomal protein S27a, putative, expressed |

| | |
|---|---|
| LOC_Os01g23940 | expressed protein |
| LOC_Os01g24800 | retrotransposon protein, putative, unclassified, expressed |
| LOC_Os01g25510 | alpha-amylase isozyme C, putative, expressed |
| LOC_Os01g28270 | retrotransposon protein, putative, Ty1-copia subclass |
| LOC_Os01g28970 | retrotransposon protein, putative, Ty3-gypsy subclass, expressed |
| LOC_Os01g32530 | transposon protein, putative, CACTA, En/Spm sub-class |
| LOC_Os01g32550 | hypothetical protein |
| LOC_Os01g33030 | Brix domain containing protein, putative, expressed |
| LOC_Os01g34440 | expressed protein |
| LOC_Os01g34620 | OsGrx_S15 |
| LOC_Os01g35870 | hypothetical protein |
| LOC_Os01g37837 | seryl-tRNA synthetase, putative, expressed |
| LOC_Os01g38110 | cytochrome P450, putative, expressed |
| LOC_Os01g38650 | expressed protein |
| LOC_Os01g38690 | nucleic acid binding protein, putative, expressed |
| LOC_Os01g39136 | retrotransposon protein, putative, unclassified, expressed |
| LOC_Os01g39590 | expressed protein |
| LOC_Os01g39860 | 1-aminocyclopropane-1-carboxylate oxidase protein, putative, expressed |
| LOC_Os01g40910 | transposon protein, putative, unclassified |
| LOC_Os01g41145 | retrotransposon protein, putative, unclassified, expressed |
| LOC_Os01g42110 | nodulin MtN3 family protein, putative, expressed |
| LOC_Os01g42740 | ammonium transporter 2, putative, expressed |
| LOC_Os01g42890 | expressed protein |
| LOC_Os01g45550 | auxin efflux carrier component, putative, expressed |
| LOC_Os01g46860 | inorganic phosphate transporter, putative, expressed |
| LOC_Os01g46870 | AP2 domain containing protein, expressed |
| LOC_Os01g48120 | expressed protein |
| LOC_Os01g48650 | expressed protein |
| LOC_Os01g49550 | expressed protein |

| | |
|---|---|
| LOC_Os01g52000 | retrotransposon protein, putative, unclassified, expressed |
| LOC_Os01g52750 | OsSub3 - Putative Subtilisin homologue, expressed |
| LOC_Os01g52920 | expressed protein |
| LOC_Os01g53750 | glucan endo-1,3-beta-glucosidase precursor, putative, expressed |
| LOC_Os01g53880 | OsIAA6 - Auxin-responsive Aux/IAA gene family member, expressed |
| LOC_Os01g53890 | RNA methyltransferase, TrmH family protein, putative, expressed |
| LOC_Os01g53900 | elongation factor, putative, expressed |
| LOC_Os01g55200 | potassium channel KAT1, putative, expressed |
| LOC_Os01g56420 | ctr copper transporter family protein, putative, expressed |
| LOC_Os01g57956 | chloroplast 50S ribosomal protein L14, putative, expressed |
| LOC_Os01g58300 | expressed protein |
| LOC_Os01g58850 | circadian clock coupling factor-related, putative, expressed |
| LOC_Os01g58870 | domain of unknown function, DUF250 domain containing protein, expressed |
| LOC_Os01g59360 | CAMK_CAMK_like |
| LOC_Os01g59760 | bZIP transcription factor, putative, expressed |
| LOC_Os01g60770 | expansin precursor, putative, expressed |
| LOC_Os01g61480 | helix-loop-helix DNA-binding domain containing protein, expressed |
| LOC_Os01g61540 | expressed protein |
| LOC_Os01g61630 | expressed protein |
| LOC_Os01g62310 | homeobox domain containing protein, expressed |
| LOC_Os01g63170 | transposon protein, putative, unclassified, expressed |
| LOC_Os01g64090 | L1P family of ribosomal proteins domain containing protein, expressed |
| LOC_Os01g64640 | histone H3, putative, expressed |
| LOC_Os01g65410 | serine hydroxymethyltransferase, mitochondrial precursor, putative, expressed |
| LOC_Os01g65440 | universal stress protein domain containing protein, putative, expressed |
| LOC_Os01g65640 | expressed protein |
| LOC_Os01g65670 | amino acid transporter, putative, expressed |
| LOC_Os01g65986 | DUF803 domain containing, putative, expressed |
| LOC_Os01g66020 | protein kinase family protein, putative, expressed |

| | |
|---|---|
| LOC_Os01g66810 | expressed protein |
| LOC_Os01g67100 | expressed protein |
| LOC_Os01g67620 | expressed protein |
| LOC_Os01g68070 | PPR repeat containing protein, expressed |
| LOC_Os01g71240 | calcium-transporting ATPase, plasma membrane-type, putative, expressed |
| LOC_Os02g02890 | peptidyl-prolyl cis-trans isomerase, putative, expressed |
| LOC_Os02g06910 | auxin response factor 6, putative, expressed |
| LOC_Os02g06940 | expressed protein |
| LOC_Os02g07130 | expressed protein |
| LOC_Os02g09390 | cytochrome P450, putative, expressed |
| LOC_Os02g09610 | GDSL-like lipase/acylhydrolase, putative, expressed |
| LOC_Os02g09620 | GDSL-like lipase/acylhydrolase, putative, expressed |
| LOC_Os02g10240 | ZOS2-05 - C2H2 zinc finger protein, expressed |
| LOC_Os02g11110 | flavonol-3-O-glycoside-7-O-glucosyltransferase 1, putative, expressed |
| LOC_Os02g11665 | transposon protein, putative, unclassified, expressed |
| LOC_Os02g14140 | cytokinin inducible protein, putative, expressed |
| LOC_Os02g16030 | harpin-induced protein 1 domain containing protein, expressed |
| LOC_Os02g16040 | ubiquitin-conjugating enzyme, putative, expressed |
| LOC_Os02g19770 | eukaryotic translation initiation factor 1A, putative, expressed |
| LOC_Os02g19780 | expressed protein |
| LOC_Os02g19790 | lipoxygenase 4, putative, expressed |
| LOC_Os02g19804 | Zinc finger C-x8-C-x5-C-x3-H type domain containing protein, expressed |
| LOC_Os02g19820 | nodulin MtN3 family protein, putative, expressed |
| LOC_Os02g20270 | expressed protein |
| LOC_Os02g20960 | hypothetical protein |
| LOC_Os02g20970 | C2 domain containing protein, putative, expressed |
| LOC_Os02g21030 | expressed protein |
| LOC_Os02g21040 | aspartic proteinase nepenthesin precursor, putative, expressed |
| LOC_Os02g21080 | AGAP008475-PA, putative, expressed |

| | |
|---|---|
| LOC_Os02g21090 | CPuORF33 - conserved peptide uORF-containing transcript, expressed |
| LOC_Os02g21220 | expressed protein |
| LOC_Os02g21240 | OsFBX48 - F-box domain containing protein, expressed |
| LOC_Os02g21250 | clathrin adaptor complex small chain domain containing protein, expressed |
| LOC_Os02g21430 | AML1, putative, expressed |
| LOC_Os02g21520 | chalcone isomerase 3, putative, expressed |
| LOC_Os02g21530 | expressed protein |
| LOC_Os02g21540 | expressed protein |
| LOC_Os02g21550 | flavonol synthase/flavanone 3-hydroxylase, putative, expressed |
| LOC_Os02g21570 | expressed protein |
| LOC_Os02g21580 | PPR repeat containing protein, expressed |
| LOC_Os02g21680 | expressed protein |
| LOC_Os02g21685 | expressed protein |
| LOC_Os02g22370 | expressed protein |
| LOC_Os02g22374 | expressed protein |
| LOC_Os02g24700 | OsSAUR8 - Auxin-responsive SAUR gene family member, expressed |
| LOC_Os02g24710 | expressed protein |
| LOC_Os02g24720 | LTPL21 - Protease inhibitor/seed storage/LTP family protein precursor, expressed |
| LOC_Os02g24730 | retrotransposon protein, putative, unclassified, expressed |
| LOC_Os02g25830 | expressed protein |
| LOC_Os02g25840 | conserved oligomeric Golgi complex component 4, related, putative, expressed |
| LOC_Os02g25860 | glutelin, putative, expressed |
| LOC_Os02g25900 | pollen signalling protein with adenylyl cyclase activity, putative, expressed |
| LOC_Os02g26310 | expressed protein |
| LOC_Os02g26320 | fasciclin-like arabinogalactan precursor protein, putative, expressed |
| LOC_Os02g26330 | expressed protein |
| LOC_Os02g26349 | transposon protein, putative, CACTA, En/Spm sub-class, expressed |
| LOC_Os02g26370 | wiskott-Aldrich syndrome protein family member 2, putative, expressed |
| LOC_Os02g26700 | cation transport regulator-like protein 1, putative, expressed |

| | |
|---|---|
| LOC_Os02g26710 | cyclo-DOPA 5-O-glucosyltransferase, putative |
| LOC_Os02g26940 | vegetative storage protein, putative, expressed |
| LOC_Os02g26950 | expressed protein |
| LOC_Os02g26960 | expressed protein |
| LOC_Os02g26970 | hypothetical protein |
| LOC_Os02g26984 | expressed protein |
| LOC_Os02g27060 | high mobility group, putative, expressed |
| LOC_Os02g27070 | PPR repeat domain containing protein, putative, expressed |
| LOC_Os02g27110 | expressed protein |
| LOC_Os02g29890 | expressed protein |
| LOC_Os02g29900 | expressed protein |
| LOC_Os02g31210 | expressed protein |
| LOC_Os02g34290 | expressed protein |
| LOC_Os02g35770 | homeobox associated leucine zipper, putative, expressed |
| LOC_Os02g35880 | salt tolerant protein, putative, expressed |
| LOC_Os02g38120 | BTBN3 - Bric-a-Brac, Tramtrack, Broad Complex BTB domain with non-phototropic hypocotyl 3 NPH3 domain, expressed |
| LOC_Os02g38392 | NBS-LRR disease resistance protein, putative, expressed |
| LOC_Os02g38490 | expressed protein |
| LOC_Os02g43800 | retrotransposon protein, putative, Ty3-gypsy subclass, expressed |
| LOC_Os02g43960 | expressed protein |
| LOC_Os02g45090 | expressed protein |
| LOC_Os02g45100 | mitochondrial import inner membrane translocase subunit Tim17, putative, expressed |
| LOC_Os02g45110 | MT-A70 domain containing protein, expressed |
| LOC_Os02g45120 | expressed protein |
| LOC_Os02g45130 | protein kinase, putative, expressed |
| LOC_Os02g45370 | retrotransposon protein, putative, Ty3-gypsy subclass, expressed |
| LOC_Os02g45380 | MATE domain containing protein, expressed |
| LOC_Os02g45410 | expressed protein |
| LOC_Os02g45420 | AP2 domain containing protein, expressed |

| | |
|---|---|
| LOC_Os02g45470 | hypothetical protein |
| LOC_Os02g45480 | nucleic acid binding protein, putative, expressed |
| LOC_Os02g45490 | expressed protein |
| LOC_Os02g45520 | uncharacterized membrane protein, putative, expressed |
| LOC_Os02g45530 | HOTHEAD precursor, putative, expressed |
| LOC_Os02g45540 | synthase, putative, expressed |
| LOC_Os02g45590 | PPR repeat domain containing protein, putative, expressed |
| LOC_Os02g45600 | rhoGAP domain containing protein, expressed |
| LOC_Os02g45630 | expressed protein |
| LOC_Os02g45650 | peptidase, putative, expressed |
| LOC_Os02g45660 | 2-C-methyl-D-erythritol 2,4-cyclodiphosphate synthase, chloroplast precursor, putative, expressed |
| LOC_Os02g45690 | uncharacterized mscS family protein, putative, expressed |
| LOC_Os02g45710 | zinc finger, C3HC4 type domain containing protein, expressed |
| LOC_Os02g46370 | expressed protein |
| LOC_Os02g46710 | expressed protein |
| LOC_Os02g47030 | expressed protein |
| LOC_Os02g47470 | cytochrome P450, putative, expressed |
| LOC_Os02g51040 | expansin precursor, putative, expressed |
| LOC_Os02g52490 | expressed protein |
| LOC_Os02g52500 | expressed protein |
| LOC_Os02g53270 | expressed protein |
| LOC_Os02g53280 | expressed protein |
| LOC_Os02g53290 | expressed protein |
| LOC_Os02g53300 | hypothetical protein |
| LOC_Os02g57570 | MATE efflux family protein, putative, expressed |
| LOC_Os02g57770 | glycosyl hydrolases family 16, putative, expressed |
| LOC_Os03g01360 | expressed protein |
| LOC_Os03g02090 | expressed protein |
| LOC_Os03g02100 | valyl-tRNA synthetase, putative, expressed |

| | |
|---|---|
| LOC_Os03g02110 | WD domain, G-beta repeat domain containing protein, expressed |
| LOC_Os03g02150 | adapitin protein, putative, expressed |
| LOC_Os03g02160 | zinc finger C-x8-C-x5-C-x3-H type family protein, expressed |
| LOC_Os03g02710 | hydroxymethylglutaryl-CoA synthase, putative, expressed |
| LOC_Os03g03550 | bZIP family transcription factor, putative, expressed |
| LOC_Os03g03570 | leucine-rich repeat transmembrane protein kinase, putative, expressed |
| LOC_Os03g03590 | transporter, monovalent cation:proton antiporter-2 family, putative, expressed |
| LOC_Os03g03600 | fasciclin-like arabinogalactan protein, putative, expressed |
| LOC_Os03g03610 | 1,3-beta-glucan synthase component domain containing protein, expressed |
| LOC_Os03g03630 | expressed protein |
| LOC_Os03g03640 | expressed protein |
| LOC_Os03g04430 | protein phosphatase 2C, putative, expressed |
| LOC_Os03g04440 | expressed protein |
| LOC_Os03g04450 | expressed protein |
| LOC_Os03g04760 | expressed protein |
| LOC_Os03g04840 | expressed protein |
| LOC_Os03g05210 | WD domain, G-beta repeat domain containing protein, expressed |
| LOC_Os03g05225 | expressed protein |
| LOC_Os03g05260 | ankyrin repeat domain containing protein, expressed |
| LOC_Os03g05270 | RING finger and CHY zinc finger domain-containing protein 1, putative, expressed |
| LOC_Os03g05340 | retrotransposon protein, putative, unclassified, expressed |
| LOC_Os03g05450 | expressed protein |
| LOC_Os03g05460 | expressed protein |
| LOC_Os03g05470 | tyrosine protein kinase domain containing protein, putative, expressed |
| LOC_Os03g05560 | zinc finger, C3HC4 type domain containing protein, expressed |
| LOC_Os03g05580 | expressed protein |
| LOC_Os03g05600 | hypothetical protein |
| LOC_Os03g05630 | expressed protein |
| LOC_Os03g05650 | expressed protein |

| | |
|---|---|
| LOC_Os03g05660 | appr-1-p processing enzyme family protein, putative, expressed |
| LOC_Os03g05680 | histone demethylase JARID1C, putative, expressed |
| LOC_Os03g05700 | expressed protein |
| LOC_Os03g05710 | acetyltransferase, GNAT family, putative, expressed |
| LOC_Os03g05720 | WD domain, G-beta repeat domain containing protein, expressed |
| LOC_Os03g05730 | cell division control protein 48 homolog E, putative, expressed |
| LOC_Os03g05750 | heavy-metal-associated domain-containing protein, putative, expressed |
| LOC_Os03g05760 | transcription factor Dp, putative, expressed |
| LOC_Os03g05800 | expressed protein |
| LOC_Os03g05806 | pseudouridine synthase family protein, putative, expressed |
| LOC_Os03g05812 | expressed protein |
| LOC_Os03g05820 | kinesin motor domain containing protein, expressed |
| LOC_Os03g05870 | Fe-S metabolism associated domain containing protein, expressed |
| LOC_Os03g05900 | expressed protein |
| LOC_Os03g06090 | high-affinity nickel-transport family protein, putative, expressed |
| LOC_Os03g06120 | PLAC8 family protein, expressed |
| LOC_Os03g06139 | ABC-2 type transporter domain containing protein, expressed |
| LOC_Os03g06160 | BTBP1 - Bric-a-Brac, Tramtrack, Broad Complex BTB domain with pentapeptide repeats, expressed |
| LOC_Os03g06170 | hsp20/alpha crystallin family protein, putative, expressed |
| LOC_Os03g06180 | expressed protein |
| LOC_Os03g06190 | 3-5 exonuclease family protein, putative, expressed |
| LOC_Os03g06200 | aminotransferase, putative, expressed |
| LOC_Os03g06220 | DEAD-box ATP-dependent RNA helicase, putative, expressed |
| LOC_Os03g06240 | YT521-B-like family domain containing protein, expressed |
| LOC_Os03g06340 | XS domain containing protein, expressed |
| LOC_Os03g06350 | DUF623 domain containing protein, expressed |
| LOC_Os03g06360 | late embryogenesis abundant protein D-34, putative, expressed |
| LOC_Os03g06370 | PPR repeat domain containing protein, putative, expressed |
| LOC_Os03g07160 | OsFBK11 - F-box domain and kelch repeat containing protein, expressed |

| | |
|---|---|
| LOC_Os03g08960 | homeobox associated leucine zipper, putative, expressed |
| LOC_Os03g10140 | ZOS3-04 - C2H2 zinc finger protein, expressed |
| LOC_Os03g10160 | pentatricopeptide repeat-containing protein, putative, expressed |
| LOC_Os03g10200 | ethylene-responsive element-binding protein, putative, expressed |
| LOC_Os03g10650 | cyclin, putative, expressed |
| LOC_Os03g10660 | expressed protein |
| LOC_Os03g10990 | retinoblastoma-binding protein-like, putative, expressed |
| LOC_Os03g11020 | PPR repeat containing protein, expressed |
| LOC_Os03g11030 | cyclin, putative, expressed |
| LOC_Os03g11140 | pleckstrin homology domain-containing protein-related taxo, putative, expressed |
| LOC_Os03g11150 | expressed protein |
| LOC_Os03g11160 | cysteine proteinase inhibitor precursor, putative, expressed |
| LOC_Os03g13300 | glutamate decarboxylase, putative, expressed |
| LOC_Os03g13920 | transposon protein, putative, CACTA, En/Spm sub-class, expressed |
| LOC_Os03g20260 | aspartic proteinase nepenthesin-1 precursor, putative, expressed |
| LOC_Os03g21000 | thioredoxin, putative, expressed |
| LOC_Os03g21010 | expressed protein |
| LOC_Os03g21020 | RNA recognition motif containing protein, putative, expressed |
| LOC_Os03g21080 | guanine nucleotide exchange factor, putative, expressed |
| LOC_Os03g21090 | auxin-independent growth promoter protein, putative, expressed |
| LOC_Os03g21140 | RNA-binding zinc finger protein, putative, expressed |
| LOC_Os03g21210 | endoglucanase, putative, expressed |
| LOC_Os03g21220 | expressed protein |
| LOC_Os03g21240 | myb-like DNA-binding domain containing protein, expressed |
| LOC_Os03g21250 | glycosyl transferase, family 8, putative, expressed |
| LOC_Os03g21260 | 2,3-bisphosphoglycerate-independent phosphoglycerate mutase, putative, expressed |
| LOC_Os03g21270 | expressed protein |
| LOC_Os03g21720 | vacuolar-sorting receptor precursor, putative, expressed |
| LOC_Os03g21730 | receptor-like protein kinase precursor, putative, expressed |

| | |
|---|---|
| LOC_Os03g25070 | CAMK_CAMK_like |
| LOC_Os03g27360 | RING-H2 finger protein ATL5H precursor, putative, expressed |
| LOC_Os03g43360 | zinc finger, C3HC4 type domain containing protein, expressed |
| LOC_Os03g43390 | F-box/LRR domain containing protein, putative, expressed |
| LOC_Os03g48000 | CorA-like magnesium transporter protein, putative, expressed |
| LOC_Os03g48020 | expressed protein |
| LOC_Os03g49524 | anthocyanidin 3-O-glucosyltransferase, putative, expressed |
| LOC_Os03g49800 | phosphatidylinositol-4-phosphate 5-kinase, putative, expressed |
| LOC_Os03g49810 | hypothetical protein |
| LOC_Os03g49840 | hypothetical protein |
| LOC_Os03g49880 | TCP family transcription factor, putative, expressed |
| LOC_Os03g49890 | hypothetical protein |
| LOC_Os03g49900 | zinc finger, C3HC4 type domain containing protein, expressed |
| LOC_Os03g49920 | expressed protein |
| LOC_Os03g49970 | expressed protein |
| LOC_Os03g52475 | pentatricopeptide, putative, expressed |
| LOC_Os03g53210 | expressed protein |
| LOC_Os03g56690 | expressed protein |
| LOC_Os03g56700 | expressed protein |
| LOC_Os03g56950 | phytochrome-interacting factor 4, putative, expressed |
| LOC_Os03g60250 | membrane associated DUF588 domain containing protein, putative, expressed |
| LOC_Os03g60939 | dolichyl-phosphate beta-glycosyltransferase, putative, expressed |
| LOC_Os03g60950 | expressed protein |
| LOC_Os03g60960 | cytokinin-N-glucosyltransferase 1, putative, expressed |
| LOC_Os03g60976 | expressed protein |
| LOC_Os04g10890 | expressed protein |
| LOC_Os04g14220 | disease resistance protein RPM1, putative, expressed |
| LOC_Os04g14240 | expressed protein |
| LOC_Os04g15600 | receptor kinase, putative, expressed |

| | |
|---|---|
| LOC_Os04g15670 | expressed protein |
| LOC_Os04g16770 | photosynthetic reaction center protein, putative, expressed |
| LOC_Os04g16772 | chloroplast 30S ribosomal protein S19, putative |
| LOC_Os04g16816 | uncharacterized protein ycf72, putative |
| LOC_Os04g24510 | OsWAK36 - OsWAK receptor-like protein kinase, expressed |
| LOC_Os04g27680 | retrotransposon protein, putative, Ty3-gypsy subclass, expressed |
| LOC_Os04g32470 | expressed protein |
| LOC_Os04g40050 | ribonuclease H2 subunit B, putative, expressed |
| LOC_Os04g40070 | GRAM and C2 domains containing protein, putative, expressed |
| LOC_Os04g40080 | leucine rich repeat containing protein, expressed |
| LOC_Os04g40090 | zinc finger, ZZ type family protein, expressed |
| LOC_Os04g40100 | BTBN11 - Bric-a-Brac, Tramtrack, Broad Complex BTB domain with non-phototropic hypocotyl 3 NPH3 domain, expressed |
| LOC_Os04g40110 | hypothetical protein |
| LOC_Os04g40120 | expressed protein |
| LOC_Os04g40130 | Rf1, mitochondrial precursor, putative, expressed |
| LOC_Os04g40140 | expressed protein |
| LOC_Os04g40600 | peptide methionine sulfoxide reductase, putative, expressed |
| LOC_Os04g42210 | GPI-anchored protein, putative, expressed |
| LOC_Os04g45360 | retrotransposon protein, putative, unclassified, expressed |
| LOC_Os04g46740 | pectinesterase, putative, expressed |
| LOC_Os04g48370 | RNA polymerase IV subunit, putative, expressed |
| LOC_Os04g49510 | CAMK_CAMK_like |
| LOC_Os04g57880 | heat shock protein DnaJ, putative, expressed |
| LOC_Os04g57900 | PPR repeat containing protein, expressed |
| LOC_Os05g01006 | retrotransposon protein, putative, unclassified, expressed |
| LOC_Os05g06230 | expressed protein |
| LOC_Os05g11820 | retrotransposon protein, putative, unclassified, expressed |
| LOC_Os05g12590 | retrotransposon protein, putative, unclassified |
| LOC_Os05g28320 | myb-like DNA-binding domain containing protein, putative, expressed |

| | |
|---|---|
| LOC_Os05g31910 | white-brown complex homolog protein 16, putative, expressed |
| LOC_Os05g33400 | basic 7S globulin precursor, putative, expressed |
| LOC_Os05g34870 | expressed protein |
| LOC_Os05g34880 | zinc finger, C3HC4 type family protein, expressed |
| LOC_Os05g34890 | expressed protein |
| LOC_Os05g34920 | expressed protein |
| LOC_Os05g34930 | hypothetical protein |
| LOC_Os05g34990 | hypothetical protein |
| LOC_Os05g35030 | hypothetical protein |
| LOC_Os05g35080 | expressed protein |
| LOC_Os05g35150 | expressed protein |
| LOC_Os05g35160 | clathrin assembly protein, putative, expressed |
| LOC_Os05g35170 | no apical meristem protein, putative, expressed |
| LOC_Os05g35210 | hypothetical protein |
| LOC_Os05g35420 | expressed protein |
| LOC_Os05g35450 | hypothetical protein |
| LOC_Os05g39230 | low photochemical bleaching 1 protein, putative, expressed |
| LOC_Os05g40820 | ribosomal protein L24, putative, expressed |
| LOC_Os05g40960 | Myb transcription factor, putative, expressed |
| LOC_Os05g40970 | expressed protein |
| LOC_Os05g40980 | zinc finger, C3HC4 type domain containing protein, expressed |
| LOC_Os05g40990 | NADH-cytochrome b5 reductase, putative, expressed |
| LOC_Os05g41000 | endonuclease/exonuclease/phosphatase family domain containing protein, expressed |
| LOC_Os05g41080 | histone H3, putative, expressed |
| LOC_Os05g41090 | CAMK_CAMK_like_CG18020d |
| LOC_Os05g41100 | protein kri1, putative, expressed |
| LOC_Os05g41450 | histone-like transcription factor and archaeal histone, putative, expressed |
| LOC_Os05g41460 | cysteine proteinase inhibitor precursor protein, putative, expressed |
| LOC_Os05g41470 | expressed protein |

| | |
|---|---|
| LOC_Os05g41510 | SH2 motif, putative, expressed |
| LOC_Os05g41530 | ZOS5-11 - C2H2 zinc finger protein, expressed |
| LOC_Os05g41540 | bZIP transcription factor domain containing protein, expressed |
| LOC_Os05g43030 | expressed protein |
| LOC_Os05g44210 | trehalose-6-phosphate synthase, putative, expressed |
| LOC_Os06g35400 | hypothetical protein |
| LOC_Os06g39515 | expressed protein |
| LOC_Os06g46380 | rab geranylgeranyl transferase like protein, putative, expressed |
| LOC_Os06g46390 | LSM domain containing protein, expressed |
| LOC_Os06g46400 | ATCPSF30/CPSF30, putative, expressed |
| LOC_Os07g03260 | CSLC10 - cellulose synthase-like family C, expressed |
| LOC_Os07g05024 | expressed protein |
| LOC_Os07g05030 | expressed protein |
| LOC_Os07g08190 | cyclophilin, putative, expressed |
| LOC_Os07g08200 | methyltransferase domain containing protein, putative, expressed |
| LOC_Os07g08210 | cytidylyltransferase domain containing protein, expressed |
| LOC_Os07g08220 | expressed protein |
| LOC_Os07g08230 | hypothetical protein |
| LOC_Os07g08240 | ZmEBE-1 protein, putative, expressed |
| LOC_Os07g08360 | expressed protein |
| LOC_Os07g08380 | expressed protein |
| LOC_Os07g12240 | EF hand family protein, putative, expressed |
| LOC_Os07g37890 | protein phosphatase 2C, putative, expressed |
| LOC_Os07g42750 | DDT domain containing protein, putative, expressed |
| LOC_Os07g42760 | expressed protein |
| LOC_Os07g44830 | proline-rich family protein, putative, expressed |
| LOC_Os07g44840 | bacterial transferase hexapeptide domain containing protein, expressed |
| LOC_Os07g49060 | retrotransposon protein, putative, unclassified |
| LOC_Os08g04390 | basic helix-loop-helix DND-binding domain containing protein, expressed |

| | |
|---|---|
| LOC_Os08g04400 | pentatricopeptide repeat domain containing protein, putative, expressed |
| LOC_Os08g04420 | tyrosine protein kinase domain containing protein, putative, expressed |
| LOC_Os08g12650 | retrotransposon protein, putative, Ty3-gypsy subclass, expressed |
| LOC_Os08g14630 | hypothetical protein |
| LOC_Os08g30634 | DC1 domain containing protein, expressed |
| LOC_Os08g32840 | bifunctional monodehydroascorbate reductase and carbonic anhydrasenectarin-3 precursor, putative, expressed |
| LOC_Os08g34950 | chaperonin Cpn60/TCP-1, putative, expressed |
| LOC_Os08g34970 | conserved oligomeric Golgi complex component 3, putative, expressed |
| LOC_Os08g35570 | expressed protein |
| LOC_Os08g35590 | expressed protein |
| LOC_Os08g35740 | 12-oxophytodienoate reductase, putative, expressed |
| LOC_Os08g35750 | Cupin domain containing protein, expressed |
| LOC_Os08g35760 | Cupin domain containing protein, expressed |
| LOC_Os08g35770 | expressed protein |
| LOC_Os08g35850 | expressed protein |
| LOC_Os08g35860 | cytokinin dehydrogenase precursor, putative, expressed |
| LOC_Os08g35990 | expressed protein |
| LOC_Os08g36020 | Disease resistance/zinc finger/chromosome condensation-like region protein, putative, expressed |
| LOC_Os08g36030 | plant viral response family protein, putative, expressed |
| LOC_Os08g36060 | expressed protein |
| LOC_Os08g36179 | expressed protein |
| LOC_Os08g36240 | expressed protein |
| LOC_Os08g36390 | ZOS8-07 - C2H2 zinc finger protein, expressed |
| LOC_Os08g36410 | ANTH domain containing protein, expressed |
| LOC_Os08g36440 | HVA22, putative, expressed |
| LOC_Os08g36450 | transcription regulator, putative, expressed |
| LOC_Os08g36520 | peptidase aspartic family protein, putative, expressed |
| LOC_Os08g36560 | emp24/gp25L/p24 family protein, putative, expressed |
| LOC_Os08g36570 | F-box protein, putative, expressed |

| | |
|---|---|
| LOC_Os08g36580 | expressed protein |
| LOC_Os08g36750 | NFKB, putative, expressed |
| LOC_Os08g36760 | remorin C-terminal domain containing protein, putative, expressed |
| LOC_Os08g36850 | hypothetical protein |
| LOC_Os08g37130 | oxidoreductase, short chain dehydrogenase/reductase family domain containing protein, expressed |
| LOC_Os08g37140 | phosphoglycerate mutase, putative, expressed |
| LOC_Os08g37150 | DUF617 domain containing protein, expressed |
| LOC_Os08g37250 | patatin, putative, expressed |
| LOC_Os08g37260 | expressed protein |
| LOC_Os08g37270 | heat shock protein DnaJ, putative, expressed |
| LOC_Os08g37280 | PWWP domain containing protein, expressed |
| LOC_Os08g37300 | expressed protein |
| LOC_Os08g37310 | uncharacterized protein, YigZ family, putative, expressed |
| LOC_Os08g37320 | expressed protein |
| LOC_Os08g37345 | DNA binding protein, putative, expressed |
| LOC_Os08g37350 | ubiquitin carboxyl-terminal hydrolase family protein, expressed |
| LOC_Os08g44350 | histidine-containing phosphotransfer protein, putative, expressed |
| LOC_Os08g44470 | expressed protein |
| LOC_Os08g44480 | 40S ribosomal protein S25, putative, expressed |
| LOC_Os08g44850 | C2 domain containing protein, putative, expressed |
| LOC_Os08g44880 | expressed protein |
| LOC_Os09g04710 | GDSL-like lipase/acylhydrolase, putative, expressed |
| LOC_Os09g24810 | ZF-HD protein dimerisation region containing protein, expressed |
| LOC_Os09g35570 | expressed protein |
| LOC_Os09g37990 | expressed protein |
| LOC_Os09g38040 | hydrolase, NUDIX family protein, expressed |
| LOC_Os09g38050 | expressed protein |
| LOC_Os09g38060 | nicotinate-nucleotide pyrophosphorylase, putative, expressed |
| LOC_Os09g38239 | sulfotransferase domain containing protein, expressed |

| | |
|---|---|
| LOC_Os09g38300 | OsFBK20 - F-box domain and kelch repeat containing protein, expressed |
| LOC_Os09g38310 | pentatricopeptide, putative, expressed |
| LOC_Os09g38320 | phytoene synthase, chloroplast precursor, putative, expressed |
| LOC_Os09g38330 | MORN repeat domain containing protein, expressed |
| LOC_Os09g39410 | male sterility protein, putative, expressed |
| LOC_Os09g39420 | WD domain, G-beta repeat domain containing protein, expressed |
| LOC_Os09g39440 | inosine-uridine preferring nucleoside hydrolase family protein, putative, expressed |
| LOC_Os09g39450 | expressed protein |
| LOC_Os09g39462 | nucleic acid binding protein, putative, expressed |
| LOC_Os09g39474 | expressed protein |
| LOC_Os09g39490 | histone-like transcription factor and archaeal histone, putative, expressed |
| LOC_Os09g39510 | Cupin domain containing protein, expressed |
| LOC_Os09g39520 | Cupin domain containing protein, expressed |
| LOC_Os09g39560 | genetic modifier, putative, expressed |
| LOC_Os09g39570 | beta-amylase, putative, expressed |
| LOC_Os09g39594 | expressed protein |
| LOC_Os09g39620 | protein kinase family protein, putative, expressed |
| LOC_Os09g39640 | protein kinase, putative, expressed |
| LOC_Os09g39650 | protein kinase family protein, putative, expressed |
| LOC_Os09g39660 | ZOS9-21 - C2H2 zinc finger protein, expressed |
| LOC_Os09g39670 | oxidoreductase, short chain dehydrogenase/reductase family domain containing family, expressed |
| LOC_Os09g39680 | transporter, major facilitator family, putative, expressed |
| LOC_Os09g39720 | 1-aminocyclopropane-1-carboxylate oxidase, putative, expressed |
| LOC_Os09g39730 | Core histone H2A/H2B/H3/H4 domain containing protein, putative, expressed |
| LOC_Os09g39740 | ACT domain containing protein, expressed |
| LOC_Os09g39750 | domain of unknown function DUF966 domain containing protein, expressed |
| LOC_Os09g39790 | lipase, putative, expressed |
| LOC_Os09g39970 | expressed protein |
| LOC_Os09g39980 | ubiquitin family protein, putative, expressed |

| | |
|---|---|
| LOC_Os10g26940 | BURP domain containing protein, expressed |
| LOC_Os10g26980 | expressed protein |
| LOC_Os10g27070 | hypothetical protein |
| LOC_Os10g27110 | expressed protein |
| LOC_Os10g27460 | expressed protein |
| LOC_Os10g28450 | CAF1 family ribonuclease containing protein, expressed |
| LOC_Os10g28600 | PPR repeat domain containing protein, putative, expressed |
| LOC_Os10g39434 | expressed protein |
| LOC_Os10g39440 | transporter family protein, putative, expressed |
| LOC_Os10g39450 | RING-H2 finger protein, putative, expressed |
| LOC_Os10g39460 | selenium-binding protein-like, putative, expressed |
| LOC_Os10g39510 | expressed protein |
| LOC_Os10g39520 | MLO domain containing protein, putative, expressed |
| LOC_Os10g39770 | zinc finger, C3HC4 type, domain containing protein, expressed |
| LOC_Os10g39780 | protein phosphatase 2C, putative, expressed |
| LOC_Os10g39790 | CorA-like magnesium transporter protein, putative, expressed |
| LOC_Os10g39800 | expressed protein |
| LOC_Os10g39810 | fatty acid hydroxylase, putative, expressed |
| LOC_Os10g39820 | zinc knuckle domain containing protein, expressed |
| LOC_Os10g39850 | zinc finger, C3HC4 type domain containing protein, expressed |
| LOC_Os10g39860 | expressed protein |
| LOC_Os10g39890 | POEI31 - Pollen Ole e I allergen and extensin family protein precursor, putative, expressed |
| LOC_Os10g39900 | glycosyltransferase family protein 1, putative, expressed |
| LOC_Os10g39910 | expressed protein |
| LOC_Os10g39950 | viral A-type inclusion protein repeat containing protein, expressed |
| LOC_Os11g03390 | FHA domain containing protein, putative, expressed |
| LOC_Os11g03400 | ribosomal protein, putative, expressed |
| LOC_Os11g03410 | exostosin family domain containing protein, expressed |
| LOC_Os11g04160 | dehydrogenase-phosphopantetheinyltransferase, putative, expressed |

| | |
|---|---|
| LOC_Os11g04180 | dual specificity protein phosphatase, putative, expressed |
| LOC_Os12g03940 | expressed protein |

**Genes in indica-japonica_SV**

| | |
|---|---|
| LOC_Os01g01090 | retrotransposon protein, putative, unclassified, expressed |
| LOC_Os01g01100 | retrotransposon protein, putative, Ty3-gypsy subclass, expressed |
| LOC_Os01g01200 | retrotransposon protein, putative, unclassified, expressed |
| LOC_Os01g01530 | transposon protein, putative, CACTA, En/Spm sub-class |
| LOC_Os01g01550 | transposon protein, putative, CACTA, En/Spm sub-class, expressed |
| LOC_Os01g01910 | expressed protein |
| LOC_Os01g02240 | transposon protein, putative, unclassified, expressed |
| LOC_Os01g02910 | glycosyltransferase, putative, expressed |
| LOC_Os01g02990 | expressed protein |
| LOC_Os01g03200 | retrotransposon protein, putative, Ty1-copia subclass, expressed |
| LOC_Os01g03240 | conserved hypothetical protein |
| LOC_Os01g03830 | retrotransposon protein, putative, unclassified, expressed |
| LOC_Os01g03860 | retrotransposon protein, putative, unclassified, expressed |
| LOC_Os01g05320 | retrotransposon protein, putative, Ty1-copia subclass, expressed |
| LOC_Os01g05580 | Exo70 exocyst complex subunit family, putative, expressed |
| LOC_Os01g06360 | transposon protein, putative, unclassified, expressed |
| LOC_Os01g06610 | retrotransposon protein, putative, Ty3-gypsy subclass, expressed |
| LOC_Os01g06810 | retrotransposon protein, putative, Ty1-copia subclass, expressed |
| LOC_Os01g07550 | hypothetical protein |
| LOC_Os01g09060 | retrotransposon protein, putative, Ty1-copia subclass |
| LOC_Os01g09960 | transposon protein, putative, unclassified, expressed |
| LOC_Os01g10195 | expressed protein |
| LOC_Os01g10560 | transposon protein, putative, unclassified |
| LOC_Os01g10770 | expressed protein |
| LOC_Os01g10850 | peroxidase precursor, putative, expressed |

| | |
|---|---|
| LOC_Os01g11054 | phosphoenolpyruvate carboxylase, putative, expressed |
| LOC_Os01g11190 | expressed protein |
| LOC_Os01g11670 | OsSCP2 - Putative Serine Carboxypeptidase homologue, expressed |
| LOC_Os01g11680 | transposon protein, putative, unclassified, expressed |
| LOC_Os01g11720 | transposable element protein, putative, containing Pfam profile: PF03004, Transposase_24, expressed |
| LOC_Os01g13300 | B3 DNA binding domain containing protein, expressed |
| LOC_Os01g13620 | retrotransposon protein, putative, unclassified, expressed |
| LOC_Os01g13630 | retrotransposon protein, putative, unclassified, expressed |
| LOC_Os01g14260 | transposon protein, putative, Ac/Ds sub-class |
| LOC_Os01g14270 | OsFBX4 - F-box domain containing protein, expressed |
| LOC_Os01g14650 | expansin precursor, putative, expressed |
| LOC_Os01g15070 | retrotransposon protein, putative, unclassified |
| LOC_Os01g15080 | hypothetical protein |
| LOC_Os01g15150 | expressed protein |
| LOC_Os01g15420 | retrotransposon protein, putative, Ty3-gypsy subclass, expressed |
| LOC_Os01g16050 | transposon protein, putative, unclassified, expressed |
| LOC_Os01g16300 | hypothetical protein |
| LOC_Os01g18020 | retrotransposon protein, putative, unclassified |
| LOC_Os01g18250 | subtilase, putative, expressed |
| LOC_Os01g19010 | retrotransposon protein, putative, unclassified, expressed |
| LOC_Os01g19100 | retrotransposon protein, putative, Ty3-gypsy subclass, expressed |
| LOC_Os01g19220 | beta-D-xylosidase, putative, expressed |
| LOC_Os01g19230 | expressed protein |
| LOC_Os01g19240 | integral membrane protein DUF6 containing protein, expressed |
| LOC_Os01g19250 | expressed protein |
| LOC_Os01g19894 | transposon protein, putative, unclassified, expressed |
| LOC_Os01g21420 | pre-mRNA-splicing factor SF2, putative, expressed |
| LOC_Os01g22309 | hypothetical protein |
| LOC_Os01g22630 | ECT protein, putative, expressed |

| | |
|---|---|
| LOC_Os01g22780 | GDSL-like lipase/acylhydrolase, putative, expressed |
| LOC_Os01g22930 | expressed protein |
| LOC_Os01g23180 | retrotransposon protein, putative, unclassified |
| LOC_Os01g23239 | retrotransposon protein, putative, Ty3-gypsy subclass, expressed |
| LOC_Os01g24160 | hypothetical protein |
| LOC_Os01g25386 | multidrug resistance-associated protein, putative, expressed |
| LOC_Os01g25600 | tetratricopeptide repeat domain containing protein, expressed |
| LOC_Os01g25920 | expressed protein |
| LOC_Os01g27070 | expressed protein |
| LOC_Os01g27230 | 12-oxophytodienoate reductase, putative, expressed |
| LOC_Os01g27260 | glutathione S-transferase, putative, expressed |
| LOC_Os01g27320 | retrotransposon protein, putative, Ty3-gypsy subclass, expressed |
| LOC_Os01g27510 | expressed protein |
| LOC_Os01g27520 | tRNA synthetases class II domain containing protein, expressed |
| LOC_Os01g27530 | expressed protein |
| LOC_Os01g27880 | sucrose-phosphatase, putative, expressed |
| LOC_Os01g27890 | cytochrome P450, putative, expressed |
| LOC_Os01g28134 | expressed protein |
| LOC_Os01g28270 | retrotransposon protein, putative, Ty1-copia subclass |
| LOC_Os01g28300 | OsFBX7 - F-box domain containing protein, expressed |
| LOC_Os01g28320 | retrotransposon, putative, centromere-specific |
| LOC_Os01g28570 | pentatricopeptide, putative, expressed |
| LOC_Os01g28580 | expressed protein |
| LOC_Os01g28590 | expressed protein |
| LOC_Os01g28600 | exo70 exocyst complex subunit family protein, putative, expressed |
| LOC_Os01g28680 | WD domain, G-beta repeat domain containing protein, expressed |
| LOC_Os01g28690 | nucleoporin autopeptidase domain containing protein, expressed |
| LOC_Os01g28700 | hypothetical protein |
| LOC_Os01g28710 | expressed protein |

| | |
|---|---|
| LOC_Os01g28720 | hypothetical protein |
| LOC_Os01g28730 | protein kinase domain containing protein, expressed |
| LOC_Os01g28830 | expressed protein |
| LOC_Os01g28940 | hypothetical protein |
| LOC_Os01g28970 | retrotransposon protein, putative, Ty3-gypsy subclass, expressed |
| LOC_Os01g28980 | proton-dependent oligopeptide transport, putative, expressed |
| LOC_Os01g29040 | retrotransposon protein, putative, unclassified, expressed |
| LOC_Os01g29570 | retrotransposon protein, putative, Ty3-gypsy subclass, expressed |
| LOC_Os01g29610 | expressed protein |
| LOC_Os01g32270 | hypothetical protein |
| LOC_Os01g32530 | transposon protein, putative, CACTA, En/Spm sub-class |
| LOC_Os01g32550 | hypothetical protein |
| LOC_Os01g33160 | stress responsive A/B Barrel domain containing protein, expressed |
| LOC_Os01g34060 | MYB family transcription factor, putative, expressed |
| LOC_Os01g34510 | hypothetical protein |
| LOC_Os01g37160 | retrotransposon protein, putative, unclassified |
| LOC_Os01g37170 | retrotransposon protein, putative, unclassified, expressed |
| LOC_Os01g38400 | retrotransposon protein, putative, Ty3-gypsy subclass, expressed |
| LOC_Os01g38650 | expressed protein |
| LOC_Os01g39136 | retrotransposon protein, putative, unclassified, expressed |
| LOC_Os01g40120 | expressed protein |
| LOC_Os01g40320 | DNA damage-binding protein 1, putative, expressed |
| LOC_Os01g40499 | S-locus lectin protein kinase family protein, putative, expressed |
| LOC_Os01g40880 | hypothetical protein |
| LOC_Os01g41440 | retrotransposon protein, putative, unclassified |
| LOC_Os01g43590 | HSF-type DNA-binding domain containing protein, expressed |
| LOC_Os01g43790 | transposon protein, putative, unclassified |
| LOC_Os01g43910 | CGMC_MAPKCMGC_2 |
| LOC_Os01g43980 | retrotransposon protein, putative, unclassified, expressed |

| | |
|---|---|
| LOC_Os01g45000 | expressed protein |
| LOC_Os01g45880 | retrotransposon protein, putative, unclassified, expressed |
| LOC_Os01g46169 | GDSL-like lipase/acylhydrolase, putative, expressed |
| LOC_Os01g47300 | retrotransposon protein, putative, unclassified, expressed |
| LOC_Os01g48060 | auxin response factor, putative, expressed |
| LOC_Os01g48640 | expressed protein |
| LOC_Os01g52260 | serine acetyltransferase protein, putative, expressed |
| LOC_Os01g52370 | expressed protein |
| LOC_Os01g52910 | expressed protein |
| LOC_Os01g53210 | metal transporter Nramp3, putative, expressed |
| LOC_Os01g53230 | expressed protein |
| LOC_Os01g53240 | BURP domain containing protein, expressed |
| LOC_Os01g53250 | NADPH reductase, putative, expressed |
| LOC_Os01g53310 | hypothetical protein |
| LOC_Os01g53340 | expressed protein |
| LOC_Os01g54140 | expressed protein |
| LOC_Os01g54980 | expressed protein |
| LOC_Os01g56850 | expressed protein |
| LOC_Os01g57110 | SNF2 family N-terminal domain containing protein, expressed |
| LOC_Os01g57140 | hypothetical protein |
| LOC_Os01g59360 | CAMK_CAMK_like |
| LOC_Os01g60080 | monocopper oxidase, putative, expressed |
| LOC_Os01g61920 | Core histone H2A/H2B/H3/H4 domain containing protein, putative, expressed |
| LOC_Os01g62700 | expressed protein |
| LOC_Os01g62710 | transposon protein, putative, unclassified |
| LOC_Os01g62740 | expressed protein |
| LOC_Os01g63050 | expressed protein |
| LOC_Os01g63970 | sialyltransferase family domain containing protein, expressed |
| LOC_Os01g63980 | ZOS1-17 - C2H2 zinc finger protein, expressed |

| | |
|---|---|
| LOC_Os01g64140 | endoglucanase 22 precursor, putative, expressed |
| LOC_Os01g64200 | transposon protein, putative, unclassified, expressed |
| LOC_Os01g64320 | expressed protein |
| LOC_Os01g64340 | expressed protein |
| LOC_Os01g64370 | expressed protein |
| LOC_Os01g64380 | expressed protein |
| LOC_Os01g65130 | peptide transporter, putative, expressed |
| LOC_Os01g66890 | BTBZ1 - Bric-a-Brac, Tramtrack, and Broad Complex BTB domain with TAZ zinc finger and Calmodulin-binding domains, expressed |
| LOC_Os01g66900 | expressed protein |
| LOC_Os01g66910 | expressed protein |
| LOC_Os01g66920 | Ser/Thr protein phosphatase family protein, putative, expressed |
| LOC_Os01g66930 | transposon protein, putative, CACTA, En/Spm sub-class, expressed |
| LOC_Os01g67620 | expressed protein |
| LOC_Os01g68160 | ZOS1-22 - C2H2 zinc finger protein, expressed |
| LOC_Os01g71870 | transposon protein, putative, unclassified, expressed |
| LOC_Os01g72140 | glutathione S-transferase, putative, expressed |
| LOC_Os01g72200 | expressed protein |
| LOC_Os01g72400 | retrotransposon protein, putative, Ty1-copia subclass, expressed |
| LOC_Os01g72620 | expressed protein |
| LOC_Os01g73820 | expressed protein |
| LOC_Os02g02080 | expressed protein |
| LOC_Os02g04400 | expressed protein |
| LOC_Os02g12140 | retrotransposon protein, putative, Ty3-gypsy subclass, expressed |
| LOC_Os02g14420 | hypothetical protein |
| LOC_Os02g14470 | hypothetical protein |
| LOC_Os02g15300 | expressed protein |
| LOC_Os02g15420 | hypothetical protein |
| LOC_Os02g15630 | retrotransposon protein, putative, unclassified, expressed |
| LOC_Os02g16030 | harpin-induced protein 1 domain containing protein, expressed |

| | |
|---|---|
| LOC_Os02g16040 | ubiquitin-conjugating enzyme, putative, expressed |
| LOC_Os02g16050 | transposon protein, putative, unclassified, expressed |
| LOC_Os02g16320 | retrotransposon protein, putative, Ty3-gypsy subclass, expressed |
| LOC_Os02g16450 | expressed protein |
| LOC_Os02g16470 | retrotransposon protein, putative, Ty1-copia subclass, expressed |
| LOC_Os02g16500 | expressed protein |
| LOC_Os02g17840 | expressed protein |
| LOC_Os02g18064 | expressed protein |
| LOC_Os02g18700 | ABC transporter family protein, putative, expressed |
| LOC_Os02g19370 | retrotransposon protein, putative, unclassified |
| LOC_Os02g20040 | loricrin, putative, expressed |
| LOC_Os02g20210 | disease resistance protein RGA1, putative, expressed |
| LOC_Os02g21320 | expressed protein |
| LOC_Os02g21550 | flavonol synthase/flavanone 3-hydroxylase, putative, expressed |
| LOC_Os02g22200 | expressed protein |
| LOC_Os02g24205 | expressed protein |
| LOC_Os02g24420 | retrotransposon protein, putative, unclassified, expressed |
| LOC_Os02g24920 | expressed protein |
| LOC_Os02g24930 | expressed protein |
| LOC_Os02g25050 | expressed protein |
| LOC_Os02g25310 | zinc knuckle family protein, expressed |
| LOC_Os02g27220 | protein phosphatase 2C, putative, expressed |
| LOC_Os02g27340 | riboflavin biosynthesis protein ribD, putative, expressed |
| LOC_Os02g29450 | expressed protein |
| LOC_Os02g29464 | RecF/RecN/SMC N terminal domain containing protein, expressed |
| LOC_Os02g33720 | RING-H2 finger protein, putative, expressed |
| LOC_Os02g33730 | ubiquinol-cytochrome C reductase hinge protein, putative, expressed |
| LOC_Os02g33800 | expressed protein |
| LOC_Os02g34730 | transposon protein, putative, Mariner sub-class, expressed |

| | |
|---|---|
| LOC_Os02g40140 | transposon protein, putative, unclassified, expressed |
| LOC_Os02g40840 | alcohol oxidase, putative, expressed |
| LOC_Os02g42380 | TCP family transcription factor, putative, expressed |
| LOC_Os02g43460 | required to maintain repression 1, putative, expressed |
| LOC_Os02g43800 | retrotransposon protein, putative, Ty3-gypsy subclass, expressed |
| LOC_Os02g43960 | expressed protein |
| LOC_Os02g44140 | retrotransposon protein, putative, Ty1-copia subclass, expressed |
| LOC_Os02g44420 | conserved hypothetical protein |
| LOC_Os02g45280 | cytochrome P450, putative, expressed |
| LOC_Os02g45370 | retrotransposon protein, putative, Ty3-gypsy subclass, expressed |
| LOC_Os02g46370 | expressed protein |
| LOC_Os02g46600 | RING-H2 finger protein ATL5G, putative, expressed |
| LOC_Os02g47470 | cytochrome P450, putative, expressed |
| LOC_Os02g47680 | retrotransposon protein, putative, Ty3-gypsy subclass |
| LOC_Os02g48260 | retrotransposon protein, putative, unclassified, expressed |
| LOC_Os02g48270 | DUF260 domain containing protein, putative |
| LOC_Os02g48280 | expressed protein |
| LOC_Os02g48290 | thioredoxin reductase 2, putative, expressed |
| LOC_Os02g48300 | OsFBX60 - F-box domain containing protein, expressed |
| LOC_Os02g48940 | expressed protein |
| LOC_Os02g49310 | AGC_PVPK_like_kin82y |
| LOC_Os02g49320 | sucrase-related, putative, expressed |
| LOC_Os02g49510 | amino acid transporter, putative, expressed |
| LOC_Os02g50120 | retrotransposon protein, putative, Ty3-gypsy subclass |
| LOC_Os02g51970 | phosphate-induced protein 1 conserved region domain containing protein, expressed |
| LOC_Os02g51990 | expressed protein |
| LOC_Os02g52000 | phosphate-induced protein 1 conserved region domain containing protein, expressed |
| LOC_Os02g54120 | CCR4-NOT transcription factor, putative, expressed |
| LOC_Os02g54420 | hypro1, putative, expressed |

| | |
|---|---|
| LOC_Os02g54780 | CYPRO4, putative, expressed |
| LOC_Os02g54800 | expressed protein |
| LOC_Os02g54810 | hypothetical protein |
| LOC_Os02g54940 | expressed protein |
| LOC_Os02g54960 | carboxyl-terminal proteinase, putative, expressed |
| LOC_Os03g02514 | hydrolase, alpha/beta fold family protein, putative, expressed |
| LOC_Os03g02530 | OsRhmbd7 - Putative Rhomboid homologue, expressed |
| LOC_Os03g04840 | expressed protein |
| LOC_Os03g05180 | expressed protein |
| LOC_Os03g05370 | expressed protein |
| LOC_Os03g05560 | zinc finger, C3HC4 type domain containing protein, expressed |
| LOC_Os03g05870 | Fe-S metabolism associated domain containing protein, expressed |
| LOC_Os03g06990 | oxidoreductase, putative, expressed |
| LOC_Os03g07160 | OsFBK11 - F-box domain and kelch repeat containing protein, expressed |
| LOC_Os03g08640 | ORC2 - Putative origin recognition complex subunit 2, expressed |
| LOC_Os03g09120 | expressed protein |
| LOC_Os03g09130 | expressed protein |
| LOC_Os03g09140 | ras-related protein, putative, expressed |
| LOC_Os03g09150 | pumilio-family RNA binding repeat domain containing protein, expressed |
| LOC_Os03g10280 | retrotransposon protein, putative, unclassified, expressed |
| LOC_Os03g10290 | expressed protein |
| LOC_Os03g11840 | expressed protein |
| LOC_Os03g12660 | cytochrome P450, putative, expressed |
| LOC_Os03g13920 | transposon protein, putative, CACTA, En/Spm sub-class, expressed |
| LOC_Os03g20060 | retrotransposon protein, putative, unclassified, expressed |
| LOC_Os03g21310 | ulp1 protease family, C-terminal catalytic domain containing protein, expressed |
| LOC_Os03g24830 | uncharacterized protein HI0077, putative, expressed |
| LOC_Os03g28180 | expressed protein |
| LOC_Os03g33420 | retrotransposon protein, putative, Ty3-gypsy subclass, expressed |

| | |
|---|---|
| LOC_Os03g36220 | retrotransposon protein, putative, unclassified, expressed |
| LOC_Os03g38520 | expressed protein |
| LOC_Os03g38534 | expressed protein |
| LOC_Os03g38540 | folic acid binding protein, putative, expressed |
| LOC_Os03g39390 | expressed protein |
| LOC_Os03g40470 | expressed protein |
| LOC_Os03g40860 | expressed protein |
| LOC_Os03g40900 | expressed protein |
| LOC_Os03g40920 | expressed protein |
| LOC_Os03g41570 | transposon protein, putative, unclassified, expressed |
| LOC_Os03g42440 | expressed protein |
| LOC_Os03g42490 | expressed protein |
| LOC_Os03g42510 | transposon protein, putative, CACTA, En/Spm sub-class, expressed |
| LOC_Os03g43114 | expressed protein |
| LOC_Os03g50340 | coatomer alpha subunit, putative, expressed |
| LOC_Os03g59225 | expressed protein |
| LOC_Os03g59400 | expressed protein |
| LOC_Os03g60030 | transposon protein, putative, CACTA, En/Spm sub-class, expressed |
| LOC_Os04g02530 | expressed protein |
| LOC_Os04g07450 | retrotransposon protein, putative, unclassified, expressed |
| LOC_Os04g12620 | expressed protein |
| LOC_Os04g12639 | carboxyl-terminal proteinase, putative, expressed |
| LOC_Os04g20424 | neural Wiskott-Aldrich syndrome protein, putative, expressed |
| LOC_Os04g20430 | expressed protein |
| LOC_Os04g21880 | expressed protein |
| LOC_Os04g21890 | disease resistance protein RPM1, putative, expressed |
| LOC_Os04g21900 | expressed protein |
| LOC_Os04g22210 | SCP-like extracellular protein, expressed |
| LOC_Os04g22220 | SCP-like extracellular protein, expressed |

| | |
|---|---|
| LOC_Os04g22230 | SCP-like extracellular protein, expressed |
| LOC_Os04g22330 | SCP-like extracellular protein, expressed |
| LOC_Os04g22340 | SCP-like extracellular protein, expressed |
| LOC_Os04g22350 | expressed protein |
| LOC_Os04g22380 | short chain dehydrogenase/reductase protein, putative, expressed |
| LOC_Os04g22440 | expressed protein |
| LOC_Os04g28500 | expressed protein |
| LOC_Os04g28760 | expressed protein |
| LOC_Os04g32770 | retrotransposon protein, putative, unclassified, expressed |
| LOC_Os04g40600 | peptide methionine sulfoxide reductase, putative, expressed |
| LOC_Os04g41080 | zinc finger, C3HC4 type domain containing protein, expressed |
| LOC_Os04g41720 | retrotransposon, putative, centromere-specific, expressed |
| LOC_Os04g42110 | cell division control protein 48 homolog B, putative, expressed |
| LOC_Os04g42120 | N-acetyltransferase ESCO2, putative, expressed |
| LOC_Os04g42130 | integral membrane transporter family protein, putative, expressed |
| LOC_Os04g42134 | enhancer of rudimentary protein, putative, expressed |
| LOC_Os04g42210 | GPI-anchored protein, putative, expressed |
| LOC_Os04g48160 | IQ calmodulin-binding motif family protein, putative, expressed |
| LOC_Os04g49290 | retrotransposon protein, putative, unclassified, expressed |
| LOC_Os04g55350 | retrotransposon protein, putative, Ty3-gypsy subclass, expressed |
| LOC_Os04g59160 | peroxidase precursor, putative, expressed |
| LOC_Os05g03810 | trehalose phosphatase, putative, expressed |
| LOC_Os05g03820 | glutamate--cysteine ligase, chloroplast precursor, putative, expressed |
| LOC_Os05g03830 | aminotransferase, putative, expressed |
| LOC_Os05g04750 | OsFBX157 - F-box domain containing protein, expressed |
| LOC_Os05g06180 | expressed protein |
| LOC_Os05g13970 | phosphorylase domain containing protein, putative, expressed |
| LOC_Os05g13980 | hypothetical protein |
| LOC_Os05g25220 | expressed protein |

| | |
|---|---|
| LOC_Os05g25940 | expressed protein |
| LOC_Os05g27720 | transposon protein, putative, CACTA, En/Spm sub-class, expressed |
| LOC_Os05g28490 | FAD-binding domain-containing protein, putative, expressed |
| LOC_Os05g30730 | expressed protein |
| LOC_Os05g30740 | C2 domain containing protein, putative, expressed |
| LOC_Os05g31530 | disease resistance protein RGA4, putative, expressed |
| LOC_Os05g36320 | expressed protein |
| LOC_Os05g36370 | expressed protein |
| LOC_Os05g36380 | expressed protein |
| LOC_Os05g40720 | glycosyl transferase, family 8, putative, expressed |
| LOC_Os05g40840 | hypothetical protein |
| LOC_Os05g40880 | expressed protein |
| LOC_Os05g43360 | NADH dehydrogenase flavoprotein 2, mitochondrial precursor, putative, expressed |
| LOC_Os05g44420 | expressed protein |
| LOC_Os05g44430 | hypothetical protein |
| LOC_Os05g45610 | expressed protein |
| LOC_Os05g47880 | hypro1, putative, expressed |
| LOC_Os05g48180 | expressed protein |
| LOC_Os05g50590 | expressed protein |
| LOC_Os05g51760 | transposon protein, putative, CACTA, En/Spm sub-class, expressed |
| LOC_Os06g05750 | transferase family domain containing protein, expressed |
| LOC_Os06g05800 | RNA recognition motif containing protein, putative, expressed |
| LOC_Os06g05804 | trafficking protein particle complex subunit 4, putative, expressed |
| LOC_Os06g24711 | expressed protein |
| LOC_Os06g28000 | carboxyl-terminal peptidase, putative, expressed |
| LOC_Os06g30470 | expressed protein |
| LOC_Os06g31810 | hypothetical protein |
| LOC_Os06g34370 | transposon protein, putative, Pong sub-class, expressed |
| LOC_Os06g38590 | receptor-like protein kinase precursor, putative, expressed |

| | |
|---|---|
| LOC_Os06g38600 | expressed protein |
| LOC_Os06g39515 | expressed protein |
| LOC_Os06g41480 | vrga1, putative, expressed |
| LOC_Os06g43740 | OsFBX201 - F-box domain containing protein, expressed |
| LOC_Os06g47350 | RNA polymerase I specific transcription initiation factor RRN3 family protein, putative, expressed |
| LOC_Os06g47360 | beta-expansin precursor, putative, expressed |
| LOC_Os06g47440 | expressed protein |
| LOC_Os06g47460 | expressed protein |
| LOC_Os07g08930 | expressed protein |
| LOC_Os07g15610 | hypothetical protein |
| LOC_Os07g16280 | retrotransposon, putative, centromere-specific, expressed |
| LOC_Os07g19560 | TATA-binding protein-associated factor 2N, putative, expressed |
| LOC_Os07g26060 | expressed protein |
| LOC_Os07g31930 | hypothetical protein |
| LOC_Os07g38664 | expressed protein |
| LOC_Os07g39660 | hypothetical protein |
| LOC_Os08g12650 | retrotransposon protein, putative, Ty3-gypsy subclass, expressed |
| LOC_Os08g32530 | expressed protein |
| LOC_Os08g33290 | expressed protein |
| LOC_Os08g35560 | expressed protein |
| LOC_Os08g36240 | expressed protein |
| LOC_Os08g36850 | hypothetical protein |
| LOC_Os08g37030 | gibberellin receptor GID1L2, putative, expressed |
| LOC_Os08g44950 | zinc finger, C3HC4 type domain containing protein, expressed |
| LOC_Os08g44960 | AP2 domain containing protein, expressed |
| LOC_Os08g45180 | kinase, pfkB family, putative, expressed |
| LOC_Os09g07160 | expressed protein |
| LOC_Os09g07930 | retrotransposon protein, putative, unclassified, expressed |
| LOC_Os09g09820 | retrotransposon protein, putative, unclassified, expressed |

| | |
|---|---|
| LOC_Os09g17329 | expressed protein |
| LOC_Os09g21490 | expressed protein |
| LOC_Os09g26040 | expressed protein |
| LOC_Os09g26300 | hypro1, putative, expressed |
| LOC_Os09g26330 | hypro1, putative, expressed |
| LOC_Os09g26670 | expressed protein |
| LOC_Os09g28700 | expressed protein |
| LOC_Os09g28710 | expressed protein |
| LOC_Os09g31250 | transposon protein, putative, unclassified, expressed |
| LOC_Os10g08230 | retrotransposon, putative, centromere-specific, expressed |
| LOC_Os10g22070 | glutathione S-transferase, putative, expressed |
| LOC_Os10g26820 | expressed protein |
| LOC_Os10g27000 | expressed protein |
| LOC_Os10g27070 | hypothetical protein |
| LOC_Os10g38160 | glutathione S-transferase, putative, expressed |
| LOC_Os10g38510 | expressed protein |
| LOC_Os11g03240 | MATE efflux family protein, putative, expressed |
| LOC_Os11g04210 | hydroxymethylglutaryl-CoA lyase, putative, expressed |
| LOC_Os11g04230 | transcription initiation factor, putative, expressed |
| LOC_Os11g04630 | expressed protein |
| LOC_Os11g06990 | expressed protein |
| LOC_Os11g07000 | expressed protein |
| LOC_Os11g07010 | expressed protein |
| LOC_Os11g11620 | ZmEBE-1 protein, putative, expressed |
| LOC_Os11g24360 | expressed protein |
| LOC_Os11g24374 | OsSCP55 - Putative Serine Carboxypeptidase homologue, expressed |
| LOC_Os11g31060 | IQ calmodulin-binding and BAG domain containing protein, putative, expressed |
| LOC_Os11g31070 | hypothetical protein |
| LOC_Os12g07300 | N-acetyltransferase 10, putative, expressed |

| | |
|---|---|
| LOC_Os12g07310 | citrate-binding protein precursor, putative, expressed |
| LOC_Os12g21560 | retrotransposon protein, putative, unclassified, expressed |

**Table S6.** GO analysis for significantly divergent genes (top 1% $F_{ST}$) between each two species, rufipogon, indica and japonica.

| GO term | Ontology | Description | Number in input list | Number in BG/Ref | p-value |
|---|---|---|---|---|---|
| **Shared genes in indica-rufipogon comparision** | | | | | |
| GO:0019430 | Biological Process | removal of superoxide radicals | 1 | 3 | 0.0056 |
| GO:0000305 | Biological Process | response to oxygen radical | 1 | 3 | 0.0056 |
| GO:0000303 | Biological Process | response to superoxide | 1 | 3 | 0.0056 |
| GO:0000302 | Biological Process | response to reactive oxygen species | 1 | 3 | 0.0056 |
| GO:0071451 | Biological Process | cellular response to superoxide | 1 | 3 | 0.0056 |
| GO:0071450 | Biological Process | cellular response to oxygen radical | 1 | 3 | 0.0056 |
| GO:0034614 | Biological Process | cellular response to reactive oxygen species | 1 | 3 | 0.0056 |
| GO:0034599 | Biological Process | cellular response to oxidative stress | 1 | 3 | 0.0056 |
| GO:0070887 | Biological Process | cellular response to chemical stimulus | 1 | 6 | 0.0098 |
| GO:0015979 | Biological Process | photosynthesis | 2 | 115 | 0.012 |
| GO:0009690 | Biological Process | cytokinin metabolic process | 1 | 10 | 0.015 |
| GO:0042445 | Biological Process | hormone metabolic process | 1 | 10 | 0.015 |
| GO:0010817 | Biological Process | regulation of hormone levels | 1 | 10 | 0.015 |
| GO:0034754 | Biological Process | cellular hormone metabolic process | 1 | 10 | 0.015 |
| GO:0009086 | Biological Process | methionine biosynthetic process | 1 | 11 | 0.017 |
| GO:0006800 | Biological Process | oxygen and reactive oxygen species metabolic process | 1 | 12 | 0.018 |
| GO:0006801 | Biological Process | superoxide metabolic process | 1 | 12 | 0.018 |
| GO:0006555 | Biological Process | methionine metabolic process | 1 | 12 | 0.018 |
| GO:0010035 | Biological Process | response to inorganic substance | 1 | 18 | 0.026 |
| GO:0009067 | Biological Process | aspartate family amino acid biosynthetic process | 1 | 26 | 0.037 |
| GO:0000097 | Biological Process | sulfur amino acid biosynthetic process | 1 | 28 | 0.04 |
| GO:0009066 | Biological Process | aspartate family amino acid metabolic process | 1 | 28 | 0.04 |
| GO:0000096 | Biological Process | sulfur amino acid metabolic process | 1 | 31 | 0.044 |
| GO:0046522 | Molecular Function | S-methyl-5-thioribose kinase activity | 1 | 2 | 0.0042 |
| GO:0016667 | Molecular Function | oxidoreductase activity, acting on sulfur group of | 2 | 75 | 0.0053 |

| | | | | | |
|---|---|---|---|---|---|
| | | donors | | | |
| GO:0004791 | Molecular Function | thioredoxin-disulfide reductase activity | 1 | 3 | 0.0056 |
| GO:0016668 | Molecular Function | oxidoreductase activity, acting on sulfur group of donors, NAD or NADP as acceptor | 1 | 9 | 0.014 |
| GO:0016671 | Molecular Function | oxidoreductase activity, acting on sulfur group of donors, disulfide as acceptor | 1 | 9 | 0.014 |
| GO:0019139 | Molecular Function | cytokinin dehydrogenase activity | 1 | 10 | 0.015 |
| GO:0050660 | Molecular Function | FAD binding | 2 | 156 | 0.021 |
| GO:0016645 | Molecular Function | oxidoreductase activity, acting on the CH-NH group of donors | 1 | 16 | 0.024 |
| GO:0004298 | Molecular Function | threonine-type endopeptidase activity | 1 | 24 | 0.035 |
| GO:0070003 | Molecular Function | threonine-type peptidase activity | 1 | 24 | 0.035 |
| GO:0019898 | Cellular Component | extrinsic to membrane | 2 | 22 | 0.00052 |
| GO:0009654 | Cellular Component | oxygen evolving complex | 2 | 26 | 0.00071 |
| GO:0009523 | Cellular Component | photosystem II | 2 | 46 | 0.0021 |
| GO:0044436 | Cellular Component | thylakoid part | 2 | 50 | 0.0024 |
| GO:0043234 | Cellular Component | protein complex | 5 | 706 | 0.003 |
| GO:0009521 | Cellular Component | photosystem | 2 | 72 | 0.0049 |
| GO:0032991 | Cellular Component | macromolecular complex | 6 | 1207 | 0.0064 |
| GO:0034357 | Cellular Component | photosynthetic membrane | 2 | 89 | 0.0073 |
| GO:0009579 | Cellular Component | thylakoid | 2 | 93 | 0.0079 |
| GO:0005875 | Cellular Component | microtubule associated complex | 1 | 7 | 0.011 |
| GO:0019773 | Cellular Component | proteasome core complex, alpha-subunit complex | 1 | 14 | 0.021 |
| GO:0005839 | Cellular Component | proteasome core complex | 1 | 24 | 0.035 |
| GO:0015630 | Cellular Component | microtubule cytoskeleton | 1 | 30 | 0.043 |
| GO:0000502 | Cellular Component | proteasome complex | 1 | 32 | 0.046 |
| **Genes in indica-rufipogon_SNP** | | | | | |
| GO:0010467 | Biological Process | gene expression | 41 | 2424 | 0.00072 |
| GO:0016070 | Biological Process | RNA metabolic process | 23 | 1088 | 0.00077 |

| | | | | | |
|---|---|---|---|---|---|
| GO:0006725 | Biological Process | cellular aromatic compound metabolic process | 7 | 145 | 0.00084 |
| GO:0006351 | Biological Process | transcription, DNA-dependent | 18 | 786 | 0.0012 |
| GO:0006366 | Biological Process | transcription from RNA polymerase II promoter | 4 | 43 | 0.0012 |
| GO:0032774 | Biological Process | RNA biosynthetic process | 18 | 788 | 0.0013 |
| GO:0080090 | Biological Process | regulation of primary metabolic process | 29 | 1626 | 0.0022 |
| GO:0060255 | Biological Process | regulation of macromolecule metabolic process | 29 | 1640 | 0.0025 |
| GO:0044271 | Biological Process | cellular nitrogen compound biosynthetic process | 8 | 231 | 0.0028 |
| GO:0019222 | Biological Process | regulation of metabolic process | 29 | 1664 | 0.003 |
| GO:0006355 | Biological Process | regulation of transcription, DNA-dependent | 16 | 737 | 0.0038 |
| GO:0051252 | Biological Process | regulation of RNA metabolic process | 16 | 740 | 0.0039 |
| GO:0006350 | Biological Process | transcription | 28 | 1629 | 0.0043 |
| GO:0050789 | Biological Process | regulation of biological process | 33 | 2027 | 0.0045 |
| GO:0046483 | Biological Process | heterocycle metabolic process | 9 | 312 | 0.0051 |
| GO:0065007 | Biological Process | biological regulation | 34 | 2125 | 0.0051 |
| GO:0034641 | Biological Process | cellular nitrogen compound metabolic process | 11 | 435 | 0.0053 |
| GO:0019219 | Biological Process | regulation of nucleobase, nucleoside, nucleotide and nucleic acid metabolic process | 26 | 1520 | 0.0063 |
| GO:0051171 | Biological Process | regulation of nitrogen compound metabolic process | 26 | 1520 | 0.0063 |
| GO:0009396 | Biological Process | folic acid and derivative biosynthetic process | 2 | 11 | 0.0072 |
| GO:0006760 | Biological Process | folic acid and derivative metabolic process | 2 | 14 | 0.011 |
| GO:0045449 | Biological Process | regulation of transcription | 25 | 1513 | 0.011 |
| GO:0031323 | Biological Process | regulation of cellular metabolic process | 26 | 1593 | 0.011 |
| GO:0008652 | Biological Process | cellular amino acid biosynthetic process | 5 | 133 | 0.012 |
| GO:0050794 | Biological Process | regulation of cellular process | 30 | 1955 | 0.014 |
| GO:0019438 | Biological Process | aromatic compound biosynthetic process | 3 | 50 | 0.016 |
| GO:0010556 | Biological Process | regulation of macromolecule biosynthetic process | 25 | 1564 | 0.016 |
| GO:0009889 | Biological Process | regulation of biosynthetic process | 25 | 1564 | 0.016 |
| GO:0031326 | Biological Process | regulation of cellular biosynthetic process | 25 | 1564 | 0.016 |
| GO:0009308 | Biological Process | amine metabolic process | 9 | 380 | 0.016 |
| GO:0006221 | Biological Process | pyrimidine nucleotide biosynthetic process | 2 | 18 | 0.017 |

| | | | | | |
|---|---|---|---|---|---|
| GO:0010468 | Biological Process | regulation of gene expression | 25 | 1577 | 0.017 |
| GO:0006220 | Biological Process | pyrimidine nucleotide metabolic process | 2 | 19 | 0.018 |
| GO:0006282 | Biological Process | regulation of DNA repair | 1 | 1 | 0.02 |
| GO:0018065 | Biological Process | protein-cofactor linkage | 1 | 1 | 0.02 |
| GO:0048583 | Biological Process | regulation of response to stimulus | 1 | 1 | 0.02 |
| GO:0080134 | Biological Process | regulation of response to stress | 1 | 1 | 0.02 |
| GO:0080135 | Biological Process | regulation of cellular response to stress | 1 | 1 | 0.02 |
| GO:0018160 | Biological Process | peptidyl-pyrromethane cofactor linkage | 1 | 1 | 0.02 |
| GO:0018198 | Biological Process | peptidyl-cysteine modification | 1 | 1 | 0.02 |
| GO:0043648 | Biological Process | dicarboxylic acid metabolic process | 3 | 55 | 0.02 |
| GO:0009108 | Biological Process | coenzyme biosynthetic process | 3 | 56 | 0.021 |
| GO:0009309 | Biological Process | amine biosynthetic process | 5 | 153 | 0.021 |
| GO:0044106 | Biological Process | cellular amine metabolic process | 8 | 333 | 0.021 |
| GO:0006813 | Biological Process | potassium ion transport | 3 | 57 | 0.022 |
| GO:0043436 | Biological Process | oxoacid metabolic process | 10 | 473 | 0.023 |
| GO:0019752 | Biological Process | carboxylic acid metabolic process | 10 | 473 | 0.023 |
| GO:0006357 | Biological Process | regulation of transcription from RNA polymerase II promoter | 2 | 22 | 0.024 |
| GO:0006082 | Biological Process | organic acid metabolic process | 10 | 474 | 0.024 |
| GO:0006752 | Biological Process | group transfer coenzyme metabolic process | 2 | 23 | 0.025 |
| GO:0042180 | Biological Process | cellular ketone metabolic process | 10 | 480 | 0.026 |
| GO:0051258 | Biological Process | protein polymerization | 2 | 24 | 0.027 |
| GO:0009174 | Biological Process | pyrimidine ribonucleoside monophosphate biosynthetic process | 1 | 2 | 0.03 |
| GO:0009173 | Biological Process | pyrimidine ribonucleoside monophosphate metabolic process | 1 | 2 | 0.03 |
| GO:0006427 | Biological Process | histidyl-tRNA aminoacylation | 1 | 2 | 0.03 |
| GO:0006420 | Biological Process | arginyl-tRNA aminoacylation | 1 | 2 | 0.03 |
| GO:0046049 | Biological Process | UMP metabolic process | 1 | 2 | 0.03 |
| GO:0006222 | Biological Process | UMP biosynthetic process | 1 | 2 | 0.03 |

| | | | | | |
|---|---|---|---|---|---|
| GO:0006399 | Biological Process | tRNA metabolic process | 4 | 114 | 0.03 |
| GO:0006520 | Biological Process | cellular amino acid metabolic process | 7 | 303 | 0.036 |
| GO:0071451 | Biological Process | cellular response to superoxide | 1 | 3 | 0.039 |
| GO:0000305 | Biological Process | response to oxygen radical | 1 | 3 | 0.039 |
| GO:0000303 | Biological Process | response to superoxide | 1 | 3 | 0.039 |
| GO:0000302 | Biological Process | response to reactive oxygen species | 1 | 3 | 0.039 |
| GO:0051052 | Biological Process | regulation of DNA metabolic process | 1 | 3 | 0.039 |
| GO:0007047 | Biological Process | cellular cell wall organization | 1 | 3 | 0.039 |
| GO:0034614 | Biological Process | cellular response to reactive oxygen species | 1 | 3 | 0.039 |
| GO:0034599 | Biological Process | cellular response to oxidative stress | 1 | 3 | 0.039 |
| GO:0019430 | Biological Process | removal of superoxide radicals | 1 | 3 | 0.039 |
| GO:0071450 | Biological Process | cellular response to oxygen radical | 1 | 3 | 0.039 |
| GO:0045229 | Biological Process | external encapsulating structure organization | 1 | 3 | 0.039 |
| GO:0010033 | Biological Process | response to organic substance | 2 | 30 | 0.04 |
| GO:0046417 | Biological Process | chorismate metabolic process | 2 | 30 | 0.04 |
| GO:0009719 | Biological Process | response to endogenous stimulus | 2 | 30 | 0.04 |
| GO:0009073 | Biological Process | aromatic amino acid family biosynthetic process | 2 | 30 | 0.04 |
| GO:0009725 | Biological Process | response to hormone stimulus | 2 | 30 | 0.04 |
| GO:0006412 | Biological Process | translation | 11 | 595 | 0.041 |
| GO:0033554 | Biological Process | cellular response to stress | 4 | 126 | 0.041 |
| GO:0006519 | Biological Process | cellular amino acid and derivative metabolic process | 8 | 391 | 0.047 |
| GO:0051716 | Biological Process | cellular response to stimulus | 4 | 132 | 0.047 |
| GO:0007017 | Biological Process | microtubule-based process | 3 | 79 | 0.049 |
| GO:0009130 | Biological Process | pyrimidine nucleoside monophosphate biosynthetic process | 1 | 4 | 0.049 |
| GO:0009129 | Biological Process | pyrimidine nucleoside monophosphate metabolic process | 1 | 4 | 0.049 |
| GO:0005249 | Molecular Function | voltage-gated potassium channel activity | 3 | 20 | 0.0015 |
| GO:0022843 | Molecular Function | voltage-gated cation channel activity | 3 | 20 | 0.0015 |
| GO:0005261 | Molecular Function | cation channel activity | 3 | 23 | 0.0021 |

| | | | | | |
|---|---|---|---|---|---|
| GO:0005267 | Molecular Function | potassium channel activity | 3 | 23 | 0.0021 |
| GO:0016765 | Molecular Function | transferase activity, transferring alkyl or aryl (other than methyl) groups | 4 | 52 | 0.0023 |
| GO:0003849 | Molecular Function | 3-deoxy-7-phosphoheptulonate synthase activity | 2 | 6 | 0.0027 |
| GO:0022832 | Molecular Function | voltage-gated channel activity | 3 | 38 | 0.0079 |
| GO:0005244 | Molecular Function | voltage-gated ion channel activity | 3 | 38 | 0.0079 |
| GO:0005198 | Molecular Function | structural molecule activity | 11 | 462 | 0.0081 |
| GO:0003702 | Molecular Function | RNA polymerase II transcription factor activity | 3 | 43 | 0.011 |
| GO:0016874 | Molecular Function | ligase activity | 8 | 305 | 0.013 |
| GO:0016455 | Molecular Function | RNA polymerase II transcription mediator activity | 2 | 18 | 0.017 |
| GO:0070569 | Molecular Function | uridylyltransferase activity | 1 | 1 | 0.02 |
| GO:0008108 | Molecular Function | UDP-glucose:hexose-1-phosphate uridylyltransferase activity | 1 | 1 | 0.02 |
| GO:0004418 | Molecular Function | hydroxymethylbilane synthase activity | 1 | 1 | 0.02 |
| GO:0004835 | Molecular Function | tubulin-tyrosine ligase activity | 1 | 1 | 0.02 |
| GO:0050660 | Molecular Function | FAD binding | 5 | 156 | 0.023 |
| GO:0022836 | Molecular Function | gated channel activity | 3 | 59 | 0.024 |
| GO:0008026 | Molecular Function | ATP-dependent helicase activity | 4 | 107 | 0.025 |
| GO:0070035 | Molecular Function | purine NTP-dependent helicase activity | 4 | 107 | 0.025 |
| GO:0016251 | Molecular Function | general RNA polymerase II transcription factor activity | 2 | 23 | 0.025 |
| GO:0016879 | Molecular Function | ligase activity, forming carbon-nitrogen bonds | 6 | 226 | 0.029 |
| GO:0004158 | Molecular Function | dihydroorotate oxidase activity | 1 | 2 | 0.03 |
| GO:0004814 | Molecular Function | arginine-tRNA ligase activity | 1 | 2 | 0.03 |
| GO:0004360 | Molecular Function | glutamine-fructose-6-phosphate transaminase (isomerizing) activity | 1 | 2 | 0.03 |
| GO:0004152 | Molecular Function | dihydroorotate dehydrogenase activity | 1 | 2 | 0.03 |
| GO:0030272 | Molecular Function | 5-formyltetrahydrofolate cyclo-ligase activity | 1 | 2 | 0.03 |
| GO:0046522 | Molecular Function | S-methyl-5-thioribose kinase activity | 1 | 2 | 0.03 |
| GO:0004821 | Molecular Function | histidine-tRNA ligase activity | 1 | 2 | 0.03 |

| | | | | | |
|---|---|---|---|---|---|
| GO:0022838 | Molecular Function | substrate-specific channel activity | 3 | 69 | 0.035 |
| GO:0015267 | Molecular Function | channel activity | 3 | 69 | 0.035 |
| GO:0005216 | Molecular Function | ion channel activity | 3 | 69 | 0.035 |
| GO:0022803 | Molecular Function | passive transmembrane transporter activity | 3 | 69 | 0.035 |
| GO:0004791 | Molecular Function | thioredoxin-disulfide reductase activity | 1 | 3 | 0.039 |
| GO:0004455 | Molecular Function | ketol-acid reductoisomerase activity | 1 | 3 | 0.039 |
| GO:0008466 | Molecular Function | glycogenin glucosyltransferase activity | 1 | 3 | 0.039 |
| GO:0008661 | Molecular Function | 1-deoxy-D-xylulose-5-phosphate synthase activity | 1 | 3 | 0.039 |
| GO:0016882 | Molecular Function | cyclo-ligase activity | 1 | 3 | 0.039 |
| GO:0016667 | Molecular Function | oxidoreductase activity, acting on sulfur group of donors | 3 | 75 | 0.043 |
| GO:0030528 | Molecular Function | transcription regulator activity | 16 | 987 | 0.044 |
| GO:0019843 | Molecular Function | rRNA binding | 2 | 32 | 0.045 |
| GO:0016702 | Molecular Function | oxidoreductase activity, acting on single donors with incorporation of molecular oxygen, incorporation of two atoms of oxygen | 2 | 33 | 0.047 |
| GO:0016813 | Molecular Function | hydrolase activity, acting on carbon-nitrogen (but not peptide) bonds, in linear amidines | 1 | 4 | 0.049 |
| GO:0004324 | Molecular Function | ferredoxin-NADP+ reductase activity | 1 | 4 | 0.049 |
| GO:0017150 | Molecular Function | tRNA dihydrouridine synthase activity | 1 | 4 | 0.049 |
| GO:0003735 | Molecular Function | structural constituent of ribosome | 8 | 396 | 0.05 |
| GO:0008076 | Cellular Component | voltage-gated potassium channel complex | 2 | 5 | 0.002 |
| GO:0034705 | Cellular Component | potassium channel complex | 2 | 5 | 0.002 |
| GO:0034702 | Cellular Component | ion channel complex | 2 | 5 | 0.002 |
| GO:0034703 | Cellular Component | cation channel complex | 2 | 5 | 0.002 |
| GO:0032991 | Cellular Component | macromolecular complex | 22 | 1207 | 0.0058 |
| GO:0031226 | Cellular Component | intrinsic to plasma membrane | 2 | 10 | 0.0061 |
| GO:0005887 | Cellular Component | integral to plasma membrane | 2 | 10 | 0.0061 |
| GO:0005856 | Cellular Component | cytoskeleton | 4 | 71 | 0.0067 |
| GO:0044459 | Cellular Component | plasma membrane part | 3 | 36 | 0.0068 |

| | | | | | |
|---|---|---|---|---|---|
| GO:0005886 | Cellular Component | plasma membrane | 3 | 44 | 0.011 |
| GO:0043234 | Cellular Component | protein complex | 14 | 706 | 0.013 |
| GO:0034357 | Cellular Component | photosynthetic membrane | 4 | 89 | 0.014 |
| GO:0005622 | Cellular Component | intracellular | 55 | 4116 | 0.015 |
| GO:0044436 | Cellular Component | thylakoid part | 3 | 50 | 0.016 |
| GO:0009579 | Cellular Component | thylakoid | 4 | 93 | 0.016 |
| GO:0005654 | Cellular Component | nucleoplasm | 3 | 51 | 0.017 |
| GO:0044451 | Cellular Component | nucleoplasm part | 3 | 51 | 0.017 |
| GO:0016592 | Cellular Component | mediator complex | 2 | 18 | 0.017 |
| GO:0031981 | Cellular Component | nuclear lumen | 3 | 56 | 0.021 |
| GO:0019898 | Cellular Component | extrinsic to membrane | 2 | 22 | 0.024 |
| GO:0005665 | Cellular Component | DNA-directed RNA polymerase II, core complex | 1 | 2 | 0.03 |
| GO:0009654 | Cellular Component | oxygen evolving complex | 2 | 26 | 0.031 |
| GO:0044464 | Cellular Component | cell part | 73 | 5945 | 0.031 |
| GO:0005623 | Cellular Component | cell | 73 | 5945 | 0.031 |
| GO:0005737 | Cellular Component | cytoplasm | 20 | 1263 | 0.032 |
| GO:0070013 | Cellular Component | intracellular organelle lumen | 3 | 70 | 0.036 |
| GO:0043233 | Cellular Component | organelle lumen | 3 | 70 | 0.036 |
| GO:0009521 | Cellular Component | photosystem | 3 | 72 | 0.039 |
| GO:0030054 | Cellular Component | cell junction | 1 | 3 | 0.039 |
| GO:0009505 | Cellular Component | plant-type cell wall | 1 | 3 | 0.039 |
| GO:0015630 | Cellular Component | microtubule cytoskeleton | 2 | 30 | 0.04 |
| GO:0043229 | Cellular Component | intracellular organelle | 38 | 2859 | 0.046 |
| GO:0043226 | Cellular Component | organelle | 38 | 2859 | 0.046 |
| **Genes in indica-rufipogon_SV** | | | | | |
| GO:0022900 | Biological Process | electron transport chain | 4 | 60 | 0.001 |
| GO:0006800 | Biological Process | oxygen and reactive oxygen species metabolic process | 2 | 12 | 0.0041 |
| GO:0006801 | Biological Process | superoxide metabolic process | 2 | 12 | 0.0041 |

| | | | | | |
|---|---|---|---|---|---|
| GO:0045333 | Biological Process | cellular respiration | 3 | 48 | 0.0053 |
| GO:0015980 | Biological Process | energy derivation by oxidation of organic compounds | 3 | 54 | 0.0073 |
| GO:0015979 | Biological Process | photosynthesis | 4 | 115 | 0.0095 |
| GO:0009767 | Biological Process | photosynthetic electron transport chain | 2 | 26 | 0.016 |
| GO:0022904 | Biological Process | respiratory electron transport chain | 2 | 28 | 0.018 |
| GO:0006425 | Biological Process | glutaminyl-tRNA aminoacylation | 1 | 2 | 0.021 |
| GO:0042816 | Biological Process | vitamin B6 metabolic process | 1 | 2 | 0.021 |
| GO:0042819 | Biological Process | vitamin B6 biosynthetic process | 1 | 2 | 0.021 |
| GO:0008615 | Biological Process | pyridoxine biosynthetic process | 1 | 2 | 0.021 |
| GO:0008614 | Biological Process | pyridoxine metabolic process | 1 | 2 | 0.021 |
| GO:0071554 | Biological Process | cell wall organization or biogenesis | 4 | 148 | 0.022 |
| GO:0043174 | Biological Process | nucleoside salvage | 1 | 3 | 0.027 |
| GO:0019430 | Biological Process | removal of superoxide radicals | 1 | 3 | 0.027 |
| GO:0006166 | Biological Process | purine ribonucleoside salvage | 1 | 3 | 0.027 |
| GO:0000305 | Biological Process | response to oxygen radical | 1 | 3 | 0.027 |
| GO:0000303 | Biological Process | response to superoxide | 1 | 3 | 0.027 |
| GO:0000302 | Biological Process | response to reactive oxygen species | 1 | 3 | 0.027 |
| GO:0071451 | Biological Process | cellular response to superoxide | 1 | 3 | 0.027 |
| GO:0071450 | Biological Process | cellular response to oxygen radical | 1 | 3 | 0.027 |
| GO:0034614 | Biological Process | cellular response to reactive oxygen species | 1 | 3 | 0.027 |
| GO:0034599 | Biological Process | cellular response to oxidative stress | 1 | 3 | 0.027 |
| GO:0006091 | Biological Process | generation of precursor metabolites and energy | 5 | 243 | 0.03 |
| GO:0050821 | Biological Process | protein stabilization | 1 | 4 | 0.034 |
| GO:0009775 | Biological Process | photosynthetic electron transport in cytochrome b6/f | 1 | 4 | 0.034 |
| GO:0031647 | Biological Process | regulation of protein stability | 1 | 4 | 0.034 |
| GO:0016998 | Biological Process | cell wall macromolecule catabolic process | 2 | 42 | 0.037 |
| GO:0046129 | Biological Process | purine ribonucleoside biosynthetic process | 1 | 5 | 0.041 |
| GO:0002437 | Biological Process | inflammatory response to antigenic stimulus | 1 | 5 | 0.041 |
| GO:0042455 | Biological Process | ribonucleoside biosynthetic process | 1 | 5 | 0.041 |
| GO:0019724 | Biological Process | B cell mediated immunity | 1 | 5 | 0.041 |

| | | | | | |
|---|---|---|---|---|---|
| GO:0042451 | Biological Process | purine nucleoside biosynthetic process | 1 | 5 | 0.041 |
| GO:0002449 | Biological Process | lymphocyte mediated immunity | 1 | 5 | 0.041 |
| GO:0009163 | Biological Process | nucleoside biosynthetic process | 1 | 5 | 0.041 |
| GO:0006954 | Biological Process | inflammatory response | 1 | 5 | 0.041 |
| GO:0002526 | Biological Process | acute inflammatory response | 1 | 5 | 0.041 |
| GO:0002252 | Biological Process | immune effector process | 1 | 5 | 0.041 |
| GO:0002524 | Biological Process | hypersensitivity | 1 | 5 | 0.041 |
| GO:0016064 | Biological Process | immunoglobulin mediated immune response | 1 | 5 | 0.041 |
| GO:0002250 | Biological Process | adaptive immune response | 1 | 5 | 0.041 |
| GO:0016068 | Biological Process | type I hypersensitivity | 1 | 5 | 0.041 |
| GO:0002460 | Biological Process | adaptive immune response based on somatic recombination of immune receptors built from immunoglobulin superfamily domains | 1 | 5 | 0.041 |
| GO:0002443 | Biological Process | leukocyte mediated immunity | 1 | 5 | 0.041 |
| GO:0002438 | Biological Process | acute inflammatory response to antigenic stimulus | 1 | 5 | 0.041 |
| GO:0044036 | Biological Process | cell wall macromolecule metabolic process | 2 | 45 | 0.042 |
| GO:0044267 | Biological Process | cellular protein metabolic process | 28 | 2877 | 0.044 |
| GO:0019684 | Biological Process | photosynthesis, light reaction | 2 | 47 | 0.045 |
| GO:0070887 | Biological Process | cellular response to chemical stimulus | 1 | 6 | 0.048 |
| GO:0009116 | Biological Process | nucleoside metabolic process | 2 | 49 | 0.049 |
| GO:0008478 | Molecular Function | pyridoxal kinase activity | 1 | 1 | 0.014 |
| GO:0004819 | Molecular Function | glutamine-tRNA ligase activity | 1 | 2 | 0.021 |
| GO:0046522 | Molecular Function | S-methyl-5-thioribose kinase activity | 1 | 2 | 0.021 |
| GO:0004001 | Molecular Function | adenosine kinase activity | 1 | 2 | 0.021 |
| GO:0004791 | Molecular Function | thioredoxin-disulfide reductase activity | 1 | 3 | 0.027 |
| GO:0019206 | Molecular Function | nucleoside kinase activity | 1 | 3 | 0.027 |
| GO:0042301 | Molecular Function | phosphate binding | 1 | 4 | 0.034 |
| GO:0008097 | Molecular Function | 5S rRNA binding | 1 | 4 | 0.034 |
| GO:0043168 | Molecular Function | anion binding | 1 | 4 | 0.034 |
| GO:0004462 | Molecular Function | lactoylglutathione lyase activity | 1 | 4 | 0.034 |

| | | | | | |
|---|---|---|---|---|---|
| GO:0008373 | Molecular Function | sialyltransferase activity | 1 | 5 | 0.041 |
| GO:0005198 | Molecular Function | structural molecule activity | 7 | 462 | 0.045 |
| GO:0004749 | Molecular Function | ribose phosphate diphosphokinase activity | 1 | 6 | 0.048 |
| GO:0008725 | Molecular Function | DNA-3-methyladenine glycosylase I activity | 1 | 6 | 0.048 |
| GO:0034357 | Cellular Component | photosynthetic membrane | 5 | 89 | 0.0005 |
| GO:0009579 | Cellular Component | thylakoid | 5 | 93 | 0.00061 |
| GO:0009521 | Cellular Component | photosystem | 4 | 72 | 0.0019 |
| GO:0009523 | Cellular Component | photosystem II | 3 | 46 | 0.0048 |
| GO:0044436 | Cellular Component | thylakoid part | 3 | 50 | 0.0059 |
| GO:0019898 | Cellular Component | extrinsic to membrane | 2 | 22 | 0.012 |
| GO:0009654 | Cellular Component | oxygen evolving complex | 2 | 26 | 0.016 |
| GO:0032991 | Cellular Component | macromolecular complex | 14 | 1207 | 0.045 |
| GO:0030173 | Cellular Component | integral to Golgi membrane | 1 | 6 | 0.048 |
| GO:0031228 | Cellular Component | intrinsic to Golgi membrane | 1 | 6 | 0.048 |
| GO:0012511 | Cellular Component | monolayer-surrounded lipid storage body | 1 | 6 | 0.048 |
| GO:0005811 | Cellular Component | lipid particle | 1 | 6 | 0.048 |
| **Share genes in japonica-rufipogon comparision** | | | | | |
| GO:0006270 | Biological Process | DNA replication initiation | 1 | 8 | 0.006 |
| GO:0006261 | Biological Process | DNA-dependent DNA replication | 1 | 17 | 0.012 |
| GO:0006887 | Biological Process | exocytosis | 1 | 64 | 0.042 |
| GO:0032940 | Biological Process | secretion by cell | 1 | 66 | 0.044 |
| GO:0046903 | Biological Process | secretion | 1 | 66 | 0.044 |
| GO:0046914 | Molecular Function | transition metal ion binding | 7 | 4875 | 0.028 |
| GO:0008270 | Molecular Function | zinc ion binding | 6 | 3939 | 0.035 |
| GO:0016758 | Molecular Function | transferase activity, transferring hexosyl groups | 2 | 489 | 0.041 |
| GO:0005938 | Cellular Component | cell cortex | 1 | 56 | 0.037 |
| GO:0000145 | Cellular Component | exocyst | 1 | 56 | 0.037 |
| GO:0044448 | Cellular Component | cell cortex part | 1 | 56 | 0.037 |

**Genes in japonica-rufipogon_SNP**

| | | | | | |
|---|---|---|---|---|---|
| GO:0006412 | Biological Process | translation | 14 | 595 | 0.0016 |
| GO:0051651 | Biological Process | maintenance of location in cell | 2 | 5 | 0.0017 |
| GO:0045185 | Biological Process | maintenance of protein location | 2 | 5 | 0.0017 |
| GO:0006621 | Biological Process | protein retention in ER lumen | 2 | 5 | 0.0017 |
| GO:0051235 | Biological Process | maintenance of location | 2 | 5 | 0.0017 |
| GO:0032507 | Biological Process | maintenance of protein location in cell | 2 | 5 | 0.0017 |
| GO:0006733 | Biological Process | oxidoreduction coenzyme metabolic process | 3 | 34 | 0.0047 |
| GO:0006413 | Biological Process | translational initiation | 3 | 36 | 0.0055 |
| GO:0065008 | Biological Process | regulation of biological quality | 6 | 176 | 0.0068 |
| GO:0006352 | Biological Process | transcription initiation | 3 | 46 | 0.01 |
| GO:0006461 | Biological Process | protein complex assembly | 4 | 94 | 0.013 |
| GO:0070271 | Biological Process | protein complex biogenesis | 4 | 94 | 0.013 |
| GO:0032313 | Biological Process | regulation of Rab GTPase activity | 2 | 19 | 0.016 |
| GO:0032483 | Biological Process | regulation of Rab protein signal transduction | 2 | 19 | 0.016 |
| GO:0032482 | Biological Process | Rab protein signal transduction | 2 | 19 | 0.016 |
| GO:0065003 | Biological Process | macromolecular complex assembly | 5 | 169 | 0.022 |
| GO:0022607 | Biological Process | cellular component assembly | 5 | 172 | 0.024 |
| GO:0042221 | Biological Process | response to chemical stimulus | 7 | 301 | 0.024 |
| GO:0006979 | Biological Process | response to oxidative stress | 5 | 174 | 0.025 |
| GO:0046496 | Biological Process | nicotinamide nucleotide metabolic process | 2 | 25 | 0.025 |
| GO:0009820 | Biological Process | alkaloid metabolic process | 2 | 25 | 0.025 |
| GO:0006769 | Biological Process | nicotinamide metabolic process | 2 | 25 | 0.025 |
| GO:0044267 | Biological Process | cellular protein metabolic process | 37 | 2877 | 0.025 |
| GO:0019362 | Biological Process | pyridine nucleotide metabolic process | 2 | 27 | 0.029 |
| GO:0043603 | Biological Process | cellular amide metabolic process | 2 | 28 | 0.031 |
| GO:0050896 | Biological Process | response to stimulus | 16 | 1026 | 0.031 |
| GO:0033554 | Biological Process | cellular response to stress | 4 | 126 | 0.032 |
| GO:0043933 | Biological Process | macromolecular complex subunit organization | 5 | 190 | 0.034 |
| GO:0006950 | Biological Process | response to stress | 14 | 873 | 0.035 |

| | | | | | |
|---|---|---|---|---|---|
| GO:0007088 | Biological Process | regulation of mitosis | 1 | 3 | 0.036 |
| GO:0042450 | Biological Process | arginine biosynthetic process via ornithine | 1 | 3 | 0.036 |
| GO:0051783 | Biological Process | regulation of nuclear division | 1 | 3 | 0.036 |
| GO:0000305 | Biological Process | response to oxygen radical | 1 | 3 | 0.036 |
| GO:0006591 | Biological Process | ornithine metabolic process | 1 | 3 | 0.036 |
| GO:0000303 | Biological Process | response to superoxide | 1 | 3 | 0.036 |
| GO:0000302 | Biological Process | response to reactive oxygen species | 1 | 3 | 0.036 |
| GO:0071450 | Biological Process | cellular response to oxygen radical | 1 | 3 | 0.036 |
| GO:0034614 | Biological Process | cellular response to reactive oxygen species | 1 | 3 | 0.036 |
| GO:0034599 | Biological Process | cellular response to oxidative stress | 1 | 3 | 0.036 |
| GO:0071451 | Biological Process | cellular response to superoxide | 1 | 3 | 0.036 |
| GO:0019430 | Biological Process | removal of superoxide radicals | 1 | 3 | 0.036 |
| GO:0030071 | Biological Process | regulation of mitotic metaphase/anaphase transition | 1 | 3 | 0.036 |
| GO:0007091 | Biological Process | mitotic metaphase/anaphase transition | 1 | 3 | 0.036 |
| GO:0051716 | Biological Process | cellular response to stimulus | 4 | 132 | 0.037 |
| GO:0010564 | Biological Process | regulation of cell cycle process | 1 | 4 | 0.045 |
| GO:0051726 | Biological Process | regulation of cell cycle | 1 | 4 | 0.045 |
| GO:0007346 | Biological Process | regulation of mitotic cell cycle | 1 | 4 | 0.045 |
| GO:0032318 | Biological Process | regulation of Ras GTPase activity | 2 | 35 | 0.045 |
| GO:0030243 | Biological Process | cellulose metabolic process | 2 | 36 | 0.047 |
| GO:0030244 | Biological Process | cellulose biosynthetic process | 2 | 36 | 0.047 |
| GO:0055114 | Biological Process | oxidation reduction | 19 | 1351 | 0.048 |
| GO:0032259 | Biological Process | methylation | 2 | 37 | 0.05 |
| GO:0046923 | Molecular Function | ER retention sequence binding | 2 | 5 | 0.0017 |
| GO:0005048 | Molecular Function | signal sequence binding | 2 | 6 | 0.0023 |
| GO:0042277 | Molecular Function | peptide binding | 2 | 6 | 0.0023 |
| GO:0005083 | Molecular Function | small GTPase regulator activity | 4 | 62 | 0.0032 |
| GO:0030695 | Molecular Function | GTPase regulator activity | 4 | 63 | 0.0034 |
| GO:0003735 | Molecular Function | structural constituent of ribosome | 10 | 396 | 0.0045 |
| GO:0060589 | Molecular Function | nucleoside-triphosphatase regulator activity | 4 | 70 | 0.0048 |

| | | | | | |
|---|---|---|---|---|---|
| GO:0003743 | Molecular Function | translation initiation factor activity | 3 | 38 | 0.0063 |
| GO:0031625 | Molecular Function | ubiquitin protein ligase binding | 2 | 13 | 0.0081 |
| GO:0047800 | Molecular Function | cysteamine dioxygenase activity | 2 | 13 | 0.0081 |
| GO:0005089 | Molecular Function | Rho guanyl-nucleotide exchange factor activity | 2 | 13 | 0.0081 |
| GO:0005088 | Molecular Function | Ras guanyl-nucleotide exchange factor activity | 2 | 13 | 0.0081 |
| GO:0019899 | Molecular Function | enzyme binding | 2 | 14 | 0.0092 |
| GO:0004650 | Molecular Function | polygalacturonase activity | 3 | 45 | 0.0097 |
| GO:0005198 | Molecular Function | structural molecule activity | 10 | 462 | 0.012 |
| GO:0016209 | Molecular Function | antioxidant activity | 6 | 203 | 0.013 |
| GO:0005099 | Molecular Function | Ras GTPase activator activity | 2 | 19 | 0.016 |
| GO:0005097 | Molecular Function | Rab GTPase activator activity | 2 | 19 | 0.016 |
| GO:0005085 | Molecular Function | guanyl-nucleotide exchange factor activity | 2 | 19 | 0.016 |
| GO:0050660 | Molecular Function | FAD binding | 5 | 156 | 0.017 |
| GO:0004514 | Molecular Function | nicotinate-nucleotide diphosphorylase (carboxylating) activity | 1 | 1 | 0.018 |
| GO:0050486 | Molecular Function | intramolecular transferase activity, transferring hydroxy groups | 1 | 1 | 0.018 |
| GO:0008909 | Molecular Function | isochorismate synthase activity | 1 | 1 | 0.018 |
| GO:0000166 | Molecular Function | nucleotide binding | 48 | 3874 | 0.02 |
| GO:0001882 | Molecular Function | nucleoside binding | 41 | 3257 | 0.026 |
| GO:0042085 | Molecular Function | 5-methyltetrahydropteroyltri-L-glutamate-dependent methyltransferase activity | 1 | 2 | 0.027 |
| GO:0003871 | Molecular Function | 5-methyltetrahydropteroyltriglutamate-homocysteine S-methyltransferase activity | 1 | 2 | 0.027 |
| GO:0008425 | Molecular Function | 2-polyprenyl-6-methoxy-1,4-benzoquinone methyltransferase activity | 1 | 2 | 0.027 |
| GO:0008169 | Molecular Function | C-methyltransferase activity | 1 | 2 | 0.027 |
| GO:0030580 | Molecular Function | quinone cofactor methyltransferase activity | 1 | 2 | 0.027 |
| GO:0004601 | Molecular Function | peroxidase activity | 5 | 181 | 0.029 |
| GO:0016684 | Molecular Function | oxidoreductase activity, acting on peroxide as acceptor | 5 | 181 | 0.029 |

| | | | | | |
|---|---|---|---|---|---|
| GO:0008135 | Molecular Function | translation factor activity, nucleic acid binding | 3 | 75 | 0.035 |
| GO:0003896 | Molecular Function | DNA primase activity | 1 | 3 | 0.036 |
| GO:0004791 | Molecular Function | thioredoxin-disulfide reductase activity | 1 | 3 | 0.036 |
| GO:0004056 | Molecular Function | argininosuccinate lyase activity | 1 | 3 | 0.036 |
| GO:0001883 | Molecular Function | purine nucleoside binding | 40 | 3247 | 0.037 |
| GO:0030554 | Molecular Function | adenyl nucleotide binding | 40 | 3247 | 0.037 |
| GO:0017076 | Molecular Function | purine nucleotide binding | 42 | 3456 | 0.039 |
| GO:0016760 | Molecular Function | cellulose synthase (UDP-forming) activity | 2 | 33 | 0.041 |
| GO:0016759 | Molecular Function | cellulose synthase activity | 2 | 33 | 0.041 |
| GO:0016702 | Molecular Function | oxidoreductase activity, acting on single donors with incorporation of molecular oxygen, incorporation of two atoms of oxygen | 2 | 33 | 0.041 |
| GO:0004096 | Molecular Function | catalase activity | 1 | 4 | 0.045 |
| GO:0004324 | Molecular Function | ferredoxin-NADP+ reductase activity | 1 | 4 | 0.045 |
| GO:0016842 | Molecular Function | amidine-lyase activity | 1 | 4 | 0.045 |
| GO:0017057 | Molecular Function | 6-phosphogluconolactonase activity | 1 | 4 | 0.045 |
| GO:0005096 | Molecular Function | GTPase activator activity | 2 | 35 | 0.045 |
| GO:0051213 | Molecular Function | dioxygenase activity | 2 | 37 | 0.05 |
| GO:0005840 | Cellular Component | ribosome | 10 | 397 | 0.0046 |
| GO:0031461 | Cellular Component | cullin-RING ubiquitin ligase complex | 2 | 13 | 0.0081 |
| GO:0030529 | Cellular Component | ribonucleoprotein complex | 10 | 438 | 0.0087 |
| GO:0032991 | Cellular Component | macromolecular complex | 20 | 1207 | 0.0095 |
| GO:0042651 | Cellular Component | thylakoid membrane | 2 | 23 | 0.022 |
| GO:0005673 | Cellular Component | transcription factor TFIIE complex | 1 | 2 | 0.027 |
| GO:0044464 | Cellular Component | cell part | 68 | 5945 | 0.029 |
| GO:0005623 | Cellular Component | cell | 68 | 5945 | 0.029 |
| GO:0000152 | Cellular Component | nuclear ubiquitin ligase complex | 1 | 3 | 0.036 |
| GO:0005680 | Cellular Component | anaphase-promoting complex | 1 | 3 | 0.036 |
| GO:0033279 | Cellular Component | ribosomal subunit | 3 | 82 | 0.044 |

**Genes in japonica-rufipogon_SV**

| | | | | | |
|---|---|---|---|---|---|
| GO:0043413 | Biological Process | macromolecule glycosylation | 5 | 60 | 0.0071 |
| GO:0006486 | Biological Process | protein amino acid glycosylation | 5 | 60 | 0.0071 |
| GO:0070085 | Biological Process | glycosylation | 5 | 60 | 0.0071 |
| GO:0009100 | Biological Process | glycoprotein metabolic process | 5 | 60 | 0.0071 |
| GO:0009101 | Biological Process | glycoprotein biosynthetic process | 5 | 60 | 0.0071 |
| GO:0006520 | Biological Process | cellular amino acid metabolic process | 12 | 303 | 0.015 |
| GO:0016070 | Biological Process | RNA metabolic process | 31 | 1088 | 0.018 |
| GO:0006563 | Biological Process | L-serine metabolic process | 3 | 28 | 0.019 |
| GO:0006519 | Biological Process | cellular amino acid and derivative metabolic process | 14 | 391 | 0.02 |
| GO:0016138 | Biological Process | glycoside biosynthetic process | 3 | 31 | 0.025 |
| GO:0046351 | Biological Process | disaccharide biosynthetic process | 3 | 31 | 0.025 |
| GO:0006901 | Biological Process | vesicle coating | 2 | 12 | 0.027 |
| GO:0006900 | Biological Process | membrane budding | 2 | 12 | 0.027 |
| GO:0048268 | Biological Process | clathrin coat assembly | 2 | 12 | 0.027 |
| GO:0006355 | Biological Process | regulation of transcription, DNA-dependent | 22 | 737 | 0.027 |
| GO:0051252 | Biological Process | regulation of RNA metabolic process | 22 | 740 | 0.028 |
| GO:0044106 | Biological Process | cellular amine metabolic process | 12 | 333 | 0.028 |
| GO:0009312 | Biological Process | oligosaccharide biosynthetic process | 3 | 33 | 0.029 |
| GO:0016192 | Biological Process | vesicle-mediated transport | 8 | 186 | 0.029 |
| GO:0016050 | Biological Process | vesicle organization | 2 | 13 | 0.031 |
| GO:0071669 | Biological Process | plant-type cell wall organization or biogenesis | 3 | 34 | 0.031 |
| GO:0009664 | Biological Process | plant-type cell wall organization | 3 | 34 | 0.031 |
| GO:0034622 | Biological Process | cellular macromolecular complex assembly | 6 | 121 | 0.031 |
| GO:0009308 | Biological Process | amine metabolic process | 13 | 380 | 0.033 |
| GO:0051179 | Biological Process | localization | 42 | 1651 | 0.034 |
| GO:0045449 | Biological Process | regulation of transcription | 39 | 1513 | 0.034 |
| GO:0019219 | Biological Process | regulation of nucleobase, nucleoside, nucleotide and nucleic acid metabolic process | 39 | 1520 | 0.036 |
| GO:0051171 | Biological Process | regulation of nitrogen compound metabolic process | 39 | 1520 | 0.036 |

| | | | | | |
|---|---|---|---|---|---|
| GO:0044262 | Biological Process | cellular carbohydrate metabolic process | 13 | 385 | 0.036 |
| GO:0009075 | Biological Process | histidine family amino acid metabolic process | 2 | 15 | 0.039 |
| GO:0006547 | Biological Process | histidine metabolic process | 2 | 15 | 0.039 |
| GO:0009069 | Biological Process | serine family amino acid metabolic process | 3 | 39 | 0.043 |
| GO:0061024 | Biological Process | membrane organization | 3 | 39 | 0.043 |
| GO:0016044 | Biological Process | cellular membrane organization | 3 | 39 | 0.043 |
| GO:0051234 | Biological Process | establishment of localization | 41 | 1638 | 0.044 |
| GO:0006810 | Biological Process | transport | 41 | 1638 | 0.044 |
| GO:0006525 | Biological Process | arginine metabolic process | 2 | 17 | 0.048 |
| GO:0009064 | Biological Process | glutamine family amino acid metabolic process | 3 | 41 | 0.048 |
| GO:0006351 | Biological Process | transcription, DNA-dependent | 22 | 786 | 0.048 |
| GO:0031497 | Biological Process | chromatin assembly | 4 | 70 | 0.049 |
| GO:0006334 | Biological Process | nucleosome assembly | 4 | 70 | 0.049 |
| GO:0065004 | Biological Process | protein-DNA complex assembly | 4 | 70 | 0.049 |
| GO:0034728 | Biological Process | nucleosome organization | 4 | 70 | 0.049 |
| GO:0032774 | Biological Process | RNA biosynthetic process | 22 | 788 | 0.049 |
| GO:0045735 | Molecular Function | nutrient reservoir activity | 7 | 77 | 0.00096 |
| GO:0030145 | Molecular Function | manganese ion binding | 6 | 58 | 0.0012 |
| GO:0016757 | Molecular Function | transferase activity, transferring glycosyl groups | 21 | 595 | 0.0059 |
| GO:0008378 | Molecular Function | galactosyltransferase activity | 4 | 43 | 0.011 |
| GO:0000166 | Molecular Function | nucleotide binding | 90 | 3874 | 0.022 |
| GO:0046873 | Molecular Function | metal ion transmembrane transporter activity | 6 | 112 | 0.023 |
| GO:0005515 | Molecular Function | protein binding | 60 | 2460 | 0.026 |
| GO:0005545 | Molecular Function | phosphatidylinositol binding | 2 | 12 | 0.027 |
| GO:0030276 | Molecular Function | clathrin binding | 2 | 12 | 0.027 |
| GO:0010181 | Molecular Function | FMN binding | 3 | 35 | 0.033 |
| GO:0003879 | Molecular Function | ATP phosphoribosyltransferase activity | 1 | 1 | 0.037 |
| GO:0051020 | Molecular Function | GTPase binding | 1 | 1 | 0.037 |
| GO:0004514 | Molecular Function | nicotinate-nucleotide diphosphorylase (carboxylating) activity | 1 | 1 | 0.037 |

| | | | | | |
|---|---|---|---|---|---|
| GO:0008685 | Molecular Function | 2-C-methyl-D-erythritol 2,4-cyclodiphosphate synthase activity | 1 | 1 | 0.037 |
| GO:0003825 | Molecular Function | alpha,alpha-trehalose-phosphate synthase (UDP-forming) activity | 1 | 1 | 0.037 |
| GO:0005576 | Cellular Component | extracellular region | 14 | 251 | 0.00047 |
| GO:0005801 | Cellular Component | cis-Golgi network | 2 | 4 | 0.0049 |
| GO:0048046 | Cellular Component | apoplast | 6 | 95 | 0.011 |
| GO:0044431 | Cellular Component | Golgi apparatus part | 4 | 50 | 0.018 |
| GO:0005794 | Cellular Component | Golgi apparatus | 4 | 56 | 0.025 |
| GO:0030118 | Cellular Component | clathrin coat | 3 | 32 | 0.027 |
| GO:0048475 | Cellular Component | coated membrane | 4 | 59 | 0.03 |
| GO:0030117 | Cellular Component | membrane coat | 4 | 59 | 0.03 |
| GO:0000786 | Cellular Component | nucleosome | 4 | 63 | 0.036 |
| GO:0032993 | Cellular Component | protein-DNA complex | 4 | 63 | 0.036 |
| GO:0030870 | Cellular Component | Mre11 complex | 1 | 1 | 0.037 |

**Shared genes in indica-japonica comparision**

| | | | | | |
|---|---|---|---|---|---|
| GO:0006887 | Biological Process | exocytosis | 1 | 64 | 0.027 |
| GO:0032940 | Biological Process | secretion by cell | 1 | 66 | 0.027 |
| GO:0046903 | Biological Process | secretion | 1 | 66 | 0.027 |
| GO:0051246 | Biological Process | regulation of protein metabolic process | 1 | 106 | 0.044 |
| GO:0070001 | Molecular Function | aspartic-type peptidase activity | 2 | 607 | 0.025 |
| GO:0004190 | Molecular Function | aspartic-type endopeptidase activity | 2 | 607 | 0.025 |
| GO:0046914 | Molecular Function | transition metal ion binding | 5 | 4875 | 0.035 |
| GO:0004175 | Molecular Function | endopeptidase activity | 2 | 856 | 0.047 |
| GO:0005938 | Cellular Component | cell cortex | 1 | 56 | 0.023 |
| GO:0000145 | Cellular Component | exocyst | 1 | 56 | 0.023 |
| GO:0044448 | Cellular Component | cell cortex part | 1 | 56 | 0.023 |

**Genes in indica-japonica_SNP**

| | | | | | |
|---|---|---|---|---|---|
| GO:0051179 | Biological Process | localization | 31 | 1651 | 0.0027 |
| GO:0030001 | Biological Process | metal ion transport | 8 | 214 | 0.003 |
| GO:0051234 | Biological Process | establishment of localization | 30 | 1638 | 0.0045 |
| GO:0006810 | Biological Process | transport | 30 | 1638 | 0.0045 |
| GO:0045449 | Biological Process | regulation of transcription | 28 | 1513 | 0.0053 |
| GO:0019219 | Biological Process | regulation of nucleobase, nucleoside, nucleotide and nucleic acid metabolic process | 28 | 1520 | 0.0056 |
| GO:0051171 | Biological Process | regulation of nitrogen compound metabolic process | 28 | 1520 | 0.0056 |
| GO:0065007 | Biological Process | biological regulation | 36 | 2125 | 0.0065 |
| GO:0080090 | Biological Process | regulation of primary metabolic process | 29 | 1626 | 0.0074 |
| GO:0071669 | Biological Process | plant-type cell wall organization or biogenesis | 3 | 34 | 0.0075 |
| GO:0009664 | Biological Process | plant-type cell wall organization | 3 | 34 | 0.0075 |
| GO:0009889 | Biological Process | regulation of biosynthetic process | 28 | 1564 | 0.0081 |
| GO:0010556 | Biological Process | regulation of macromolecule biosynthetic process | 28 | 1564 | 0.0081 |
| GO:0031326 | Biological Process | regulation of cellular biosynthetic process | 28 | 1564 | 0.0081 |
| GO:0060255 | Biological Process | regulation of macromolecule metabolic process | 29 | 1640 | 0.0083 |
| GO:0010468 | Biological Process | regulation of gene expression | 28 | 1577 | 0.009 |
| GO:0050789 | Biological Process | regulation of biological process | 34 | 2027 | 0.0094 |
| GO:0050794 | Biological Process | regulation of cellular process | 33 | 1955 | 0.0096 |
| GO:0006901 | Biological Process | vesicle coating | 2 | 12 | 0.0098 |
| GO:0006900 | Biological Process | membrane budding | 2 | 12 | 0.0098 |
| GO:0048268 | Biological Process | clathrin coat assembly | 2 | 12 | 0.0098 |
| GO:0019222 | Biological Process | regulation of metabolic process | 29 | 1664 | 0.01 |
| GO:0031323 | Biological Process | regulation of cellular metabolic process | 28 | 1593 | 0.01 |
| GO:0010467 | Biological Process | gene expression | 39 | 2424 | 0.01 |
| GO:0016050 | Biological Process | vesicle organization | 2 | 13 | 0.011 |
| GO:0055085 | Biological Process | transmembrane transport | 15 | 701 | 0.012 |
| GO:0034622 | Biological Process | cellular macromolecular complex assembly | 5 | 121 | 0.012 |
| GO:0006350 | Biological Process | transcription | 28 | 1629 | 0.013 |
| GO:0071555 | Biological Process | cell wall organization | 4 | 83 | 0.015 |

| | | | | | |
|---|---|---|---|---|---|
| GO:0006812 | Biological Process | cation transport | 9 | 348 | 0.016 |
| GO:0046488 | Biological Process | phosphatidylinositol metabolic process | 2 | 18 | 0.02 |
| GO:0034621 | Biological Process | cellular macromolecular complex subunit organization | 5 | 142 | 0.022 |
| GO:0016070 | Biological Process | RNA metabolic process | 19 | 1088 | 0.033 |
| GO:0006811 | Biological Process | ion transport | 9 | 408 | 0.039 |
| GO:0065003 | Biological Process | macromolecular complex assembly | 5 | 169 | 0.042 |
| GO:0006563 | Biological Process | L-serine metabolic process | 2 | 28 | 0.042 |
| GO:0042450 | Biological Process | arginine biosynthetic process via ornithine | 1 | 3 | 0.043 |
| GO:0006591 | Biological Process | ornithine metabolic process | 1 | 3 | 0.043 |
| GO:0006434 | Biological Process | seryl-tRNA aminoacylation | 1 | 3 | 0.043 |
| GO:0022607 | Biological Process | cellular component assembly | 5 | 172 | 0.044 |
| GO:0034728 | Biological Process | nucleosome organization | 3 | 70 | 0.045 |
| GO:0065004 | Biological Process | protein-DNA complex assembly | 3 | 70 | 0.045 |
| GO:0031497 | Biological Process | chromatin assembly | 3 | 70 | 0.045 |
| GO:0006334 | Biological Process | nucleosome assembly | 3 | 70 | 0.045 |
| GO:0006323 | Biological Process | DNA packaging | 3 | 71 | 0.047 |
| GO:0030145 | Molecular Function | manganese ion binding | 6 | 58 | 6.80E-05 |
| GO:0045735 | Molecular Function | nutrient reservoir activity | 6 | 77 | 0.00029 |
| GO:0046873 | Molecular Function | metal ion transmembrane transporter activity | 6 | 112 | 0.0018 |
| GO:0005545 | Molecular Function | phosphatidylinositol binding | 2 | 12 | 0.0098 |
| GO:0030276 | Molecular Function | clathrin binding | 2 | 12 | 0.0098 |
| GO:0005515 | Molecular Function | protein binding | 39 | 2460 | 0.013 |
| GO:0008324 | Molecular Function | cation transmembrane transporter activity | 8 | 280 | 0.014 |
| GO:0016307 | Molecular Function | phosphatidylinositol phosphate kinase activity | 2 | 18 | 0.02 |
| GO:0051020 | Molecular Function | GTPase binding | 1 | 1 | 0.022 |
| GO:0008685 | Molecular Function | 2-C-methyl-D-erythritol 2,4-cyclodiphosphate synthase activity | 1 | 1 | 0.022 |
| GO:0004514 | Molecular Function | nicotinate-nucleotide diphosphorylase (carboxylating) activity | 1 | 1 | 0.022 |

| | | | | | |
|---|---|---|---|---|---|
| GO:0003825 | Molecular Function | alpha,alpha-trehalose-phosphate synthase (UDP-forming) activity | 1 | 1 | 0.022 |
| GO:0001727 | Molecular Function | lipid kinase activity | 2 | 20 | 0.023 |
| GO:0005249 | Molecular Function | voltage-gated potassium channel activity | 2 | 20 | 0.023 |
| GO:0035091 | Molecular Function | phosphoinositide binding | 2 | 20 | 0.023 |
| GO:0022843 | Molecular Function | voltage-gated cation channel activity | 2 | 20 | 0.023 |
| GO:0005267 | Molecular Function | potassium channel activity | 2 | 23 | 0.03 |
| GO:0005261 | Molecular Function | cation channel activity | 2 | 23 | 0.03 |
| GO:0016853 | Molecular Function | isomerase activity | 5 | 161 | 0.035 |
| GO:0004828 | Molecular Function | serine-tRNA ligase activity | 1 | 3 | 0.043 |
| GO:0016849 | Molecular Function | phosphorus-oxygen lyase activity | 1 | 3 | 0.043 |
| GO:0008897 | Molecular Function | holo-[acyl-carrier-protein] synthase activity | 1 | 3 | 0.043 |
| GO:0004421 | Molecular Function | hydroxymethylglutaryl-CoA synthase activity | 1 | 3 | 0.043 |
| GO:0004056 | Molecular Function | argininosuccinate lyase activity | 1 | 3 | 0.043 |
| GO:0004619 | Molecular Function | phosphoglycerate mutase activity | 1 | 3 | 0.043 |
| GO:0004437 | Molecular Function | inositol or phosphatidylinositol phosphatase activity | 2 | 29 | 0.044 |
| GO:0016859 | Molecular Function | cis-trans isomerase activity | 2 | 29 | 0.044 |
| GO:0003755 | Molecular Function | peptidyl-prolyl cis-trans isomerase activity | 2 | 29 | 0.044 |
| GO:0030528 | Molecular Function | transcription regulator activity | 17 | 987 | 0.047 |
| GO:0004428 | Molecular Function | inositol or phosphatidylinositol kinase activity | 2 | 31 | 0.05 |
| GO:0048046 | Cellular Component | apoplast | 6 | 95 | 0.00082 |
| GO:0005576 | Cellular Component | extracellular region | 9 | 251 | 0.0022 |
| GO:0044464 | Cellular Component | cell part | 84 | 5945 | 0.0056 |
| GO:0005623 | Cellular Component | cell | 84 | 5945 | 0.0056 |
| GO:0030118 | Cellular Component | clathrin coat | 3 | 32 | 0.0064 |
| GO:0016020 | Cellular Component | membrane | 33 | 2115 | 0.027 |
| GO:0032991 | Cellular Component | macromolecular complex | 21 | 1207 | 0.027 |
| GO:0048475 | Cellular Component | coated membrane | 3 | 59 | 0.03 |
| GO:0030117 | Cellular Component | membrane coat | 3 | 59 | 0.03 |
| GO:0000786 | Cellular Component | nucleosome | 3 | 63 | 0.035 |

| | | | | | |
|---|---|---|---|---|---|
| GO:0032993 | Cellular Component | protein-DNA complex | 3 | 63 | 0.035 |

**Genes in indica-japonica_SV**

| | | | | | |
|---|---|---|---|---|---|
| GO:0043413 | Biological Process | macromolecule glycosylation | 5 | 60 | 4.00E-05 |
| GO:0009100 | Biological Process | glycoprotein metabolic process | 5 | 60 | 4.00E-05 |
| GO:0006486 | Biological Process | protein amino acid glycosylation | 5 | 60 | 4.00E-05 |
| GO:0009101 | Biological Process | glycoprotein biosynthetic process | 5 | 60 | 4.00E-05 |
| GO:0070085 | Biological Process | glycosylation | 5 | 60 | 4.00E-05 |
| GO:0044262 | Biological Process | cellular carbohydrate metabolic process | 8 | 385 | 0.0023 |
| GO:0055114 | Biological Process | oxidation reduction | 15 | 1351 | 0.014 |
| GO:0016138 | Biological Process | glycoside biosynthetic process | 2 | 31 | 0.016 |
| GO:0046351 | Biological Process | disaccharide biosynthetic process | 2 | 31 | 0.016 |
| GO:0006421 | Biological Process | asparaginyl-tRNA aminoacylation | 1 | 2 | 0.017 |
| GO:0009312 | Biological Process | oligosaccharide biosynthetic process | 2 | 33 | 0.018 |
| GO:0071451 | Biological Process | cellular response to superoxide | 1 | 3 | 0.023 |
| GO:0019430 | Biological Process | removal of superoxide radicals | 1 | 3 | 0.023 |
| GO:0000305 | Biological Process | response to oxygen radical | 1 | 3 | 0.023 |
| GO:0000303 | Biological Process | response to superoxide | 1 | 3 | 0.023 |
| GO:0000302 | Biological Process | response to reactive oxygen species | 1 | 3 | 0.023 |
| GO:0071450 | Biological Process | cellular response to oxygen radical | 1 | 3 | 0.023 |
| GO:0034614 | Biological Process | cellular response to reactive oxygen species | 1 | 3 | 0.023 |
| GO:0005986 | Biological Process | sucrose biosynthetic process | 1 | 3 | 0.023 |
| GO:0034599 | Biological Process | cellular response to oxidative stress | 1 | 3 | 0.023 |
| GO:0016192 | Biological Process | vesicle-mediated transport | 4 | 186 | 0.026 |
| GO:0005975 | Biological Process | carbohydrate metabolic process | 10 | 833 | 0.027 |
| GO:0005984 | Biological Process | disaccharide metabolic process | 2 | 44 | 0.03 |
| GO:0016137 | Biological Process | glycoside metabolic process | 2 | 44 | 0.03 |
| GO:0009311 | Biological Process | oligosaccharide metabolic process | 2 | 47 | 0.034 |
| GO:0051649 | Biological Process | establishment of localization in cell | 5 | 299 | 0.034 |
| GO:0042221 | Biological Process | response to chemical stimulus | 5 | 301 | 0.035 |

| | | | | | |
|---|---|---|---|---|---|
| GO:0006422 | Biological Process | aspartyl-tRNA aminoacylation | 1 | 5 | 0.035 |
| GO:0006750 | Biological Process | glutathione biosynthetic process | 1 | 5 | 0.035 |
| GO:0045333 | Biological Process | cellular respiration | 2 | 48 | 0.035 |
| GO:0044272 | Biological Process | sulfur compound biosynthetic process | 2 | 48 | 0.035 |
| GO:0051641 | Biological Process | cellular localization | 5 | 309 | 0.038 |
| GO:0070887 | Biological Process | cellular response to chemical stimulus | 1 | 6 | 0.04 |
| GO:0006122 | Biological Process | mitochondrial electron transport, ubiquinol to cytochrome c | 1 | 6 | 0.04 |
| GO:0042775 | Biological Process | mitochondrial ATP synthesis coupled electron transport | 1 | 6 | 0.04 |
| GO:0006014 | Biological Process | D-ribose metabolic process | 1 | 6 | 0.04 |
| GO:0033554 | Biological Process | cellular response to stress | 3 | 126 | 0.04 |
| GO:0050896 | Biological Process | response to stimulus | 11 | 1026 | 0.042 |
| GO:0015980 | Biological Process | energy derivation by oxidation of organic compounds | 2 | 54 | 0.043 |
| GO:0051716 | Biological Process | cellular response to stimulus | 3 | 132 | 0.045 |
| GO:0006749 | Biological Process | glutathione metabolic process | 1 | 7 | 0.046 |
| GO:0008378 | Molecular Function | galactosyltransferase activity | 4 | 43 | 0.00017 |
| GO:0016491 | Molecular Function | oxidoreductase activity | 19 | 1672 | 0.0047 |
| GO:0004816 | Molecular Function | asparagine-tRNA ligase activity | 1 | 2 | 0.017 |
| GO:0004357 | Molecular Function | glutamate-cysteine ligase activity | 1 | 2 | 0.017 |
| GO:0010181 | Molecular Function | FMN binding | 2 | 35 | 0.02 |
| GO:0004791 | Molecular Function | thioredoxin-disulfide reductase activity | 1 | 3 | 0.023 |
| GO:0005542 | Molecular Function | folic acid binding | 1 | 3 | 0.023 |
| GO:0050307 | Molecular Function | sucrose-phosphatase activity | 1 | 3 | 0.023 |
| GO:0016757 | Molecular Function | transferase activity, transferring glycosyl groups | 8 | 595 | 0.026 |
| GO:0004815 | Molecular Function | aspartate-tRNA ligase activity | 1 | 5 | 0.035 |
| GO:0008373 | Molecular Function | sialyltransferase activity | 1 | 5 | 0.035 |
| GO:0009001 | Molecular Function | serine O-acetyltransferase activity | 1 | 6 | 0.04 |
| GO:0016412 | Molecular Function | serine O-acyltransferase activity | 1 | 6 | 0.04 |
| GO:0004747 | Molecular Function | ribokinase activity | 1 | 6 | 0.04 |

| | | | | | |
|---|---|---|---|---|---|
| GO:0008964 | Molecular Function | phosphoenolpyruvate carboxylase activity | 1 | 7 | 0.046 |
| GO:0016413 | Molecular Function | O-acetyltransferase activity | 1 | 7 | 0.046 |
| GO:0044431 | Cellular Component | Golgi apparatus part | 3 | 50 | 0.0037 |
| GO:0005576 | Cellular Component | extracellular region | 6 | 251 | 0.0042 |
| GO:0005794 | Cellular Component | Golgi apparatus | 3 | 56 | 0.005 |
| GO:0030870 | Cellular Component | Mre11 complex | 1 | 1 | 0.012 |
| GO:0000139 | Cellular Component | Golgi membrane | 2 | 29 | 0.014 |
| GO:0012505 | Cellular Component | endomembrane system | 3 | 104 | 0.025 |
| GO:0005801 | Cellular Component | cis-Golgi network | 1 | 4 | 0.029 |
| GO:0030173 | Cellular Component | integral to Golgi membrane | 1 | 6 | 0.04 |
| GO:0031228 | Cellular Component | intrinsic to Golgi membrane | 1 | 6 | 0.04 |
| GO:0000808 | Cellular Component | origin recognition complex | 1 | 6 | 0.04 |
| GO:0000145 | Cellular Component | exocyst | 2 | 56 | 0.046 |
| GO:0044448 | Cellular Component | cell cortex part | 2 | 56 | 0.046 |
| GO:0005938 | Cellular Component | cell cortex | 2 | 56 | 0.046 |

**Table S7.** List of genes within the windows that were top 1% by CLR, as identified by SweeD, for both SNPs and SVs.

| | Gene ID (MSU7) | Function annotation |
|---|---|---|
| **Shared genes in rufipogon_SNP-SV (two overlapping windows)** | | |
| 1 | LOC_Os01g08890 | expressed protein |
| 2 | LOC_Os01g08930 | DEAD-box ATP-dependent RNA helicase, putative, expressed |
| 3 | LOC_Os01g29040 | retrotransposon protein, putative, unclassified, expressed |
| **Shared genes in indica_SNP-SV comparison (only one overlapping window)** | | |
| 1 | LOC_Os01g72400 | retrotransposon protein, putative, Ty1-copia subclass, expressed |
| 2 | LOC_Os02g12140 | retrotransposon protein, putative, Ty3-gypsy subclass, expressed |
| 3 | LOC_Os02g25290 | retrotransposon protein, putative, unclassified, expressed |
| **Shared genes in japonica_SNP-SV (ten overlapping windows)** | | |
| 1 | LOC_Os01g01100 | retrotransposon protein, putative, Ty3-gypsy subclass, expressed |
| 2 | LOC_Os01g02629 | retrotransposon protein, putative, unclassified, expressed |
| 3 | LOC_Os01g09110 | retrotransposon protein, putative, unclassified |
| 4 | LOC_Os01g18020 | retrotransposon protein, putative, unclassified |
| 5 | LOC_Os01g27070 | expressed protein |
| 6 | LOC_Os01g36740 | expressed protein |
| 7 | LOC_Os01g36760 | expressed protein |
| 8 | LOC_Os01g36790 | TKL_IRAK_DUF26-lf 2 - DUF26 kinases have homology to DUF26 containing loci, expressed |
| 9 | LOC_Os01g40120 | expressed protein |
| 10 | LOC_Os01g52910 | expressed protein |
| 11 | LOC_Os01g58049 | photosystem I assembly protein ycf4, putative, expressed |
| 12 | LOC_Os01g63040 | expressed protein |
| 13 | LOC_Os04g28520 | male sterility protein, putative, expressed |
| 14 | LOC_Os04g28530 | expressed protein |
| 15 | LOC_Os04g28540 | expressed protein |
| 16 | LOC_Os07g17460 | OsFBL36 - F-box domain and LRR containing protein, expressed |
| 17 | LOC_Os07g17560 | expressed protein |

**Table S8.** Functional annotation of genes from potentially selective regions (top 1% CLR) identified by SweeD.

| Gene ID (MSU7) | Function annotation |
|---|---|
| **Genes in indica_SNP CLR** | |
| LOC_Os01g01200 | retrotransposon protein, putative, unclassified, expressed |
| LOC_Os01g01430 | No apical meristem protein, putative, expressed |
| LOC_Os01g01450 | stress responsive protein, putative, expressed |
| LOC_Os01g01470 | no apical meristem protein, putative, expressed |
| LOC_Os01g02240 | transposon protein, putative, unclassified, expressed |
| LOC_Os01g02860 | expressed protein |
| LOC_Os01g02910 | glycosyltransferase, putative, expressed |
| LOC_Os01g03240 | conserved hypothetical protein |
| LOC_Os01g03720 | MYB family transcription factor, putative, expressed |
| LOC_Os01g03770 | hypothetical protein |
| LOC_Os01g03830 | retrotransposon protein, putative, unclassified, expressed |
| LOC_Os01g03860 | retrotransposon protein, putative, unclassified, expressed |
| LOC_Os01g04000 | hypothetical protein |
| LOC_Os01g04730 | ribosomal protein L24, putative, expressed |
| LOC_Os01g05320 | retrotransposon protein, putative, Ty1-copia subclass, expressed |
| LOC_Os01g06240 | protein kinase, putative, expressed |
| LOC_Os01g06610 | retrotransposon protein, putative, Ty3-gypsy subclass, expressed |
| LOC_Os01g07970 | expressed protein |
| LOC_Os01g08020 | boron transporter protein, putative, expressed |
| LOC_Os01g08850 | expressed protein |
| LOC_Os01g08860 | hsp20/alpha crystallin family protein, putative, expressed |
| LOC_Os01g08870 | expressed protein |
| LOC_Os01g08880 | expressed protein |
| LOC_Os01g08890 | expressed protein |
| LOC_Os01g08930 | DEAD-box ATP-dependent RNA helicase, putative, expressed |
| LOC_Os01g08940 | expressed protein |
| LOC_Os01g08950 | expressed protein |

| | |
|---|---|
| LOC_Os01g08970 | SSRP1-like FACT complex subunit, putative, expressed |
| LOC_Os01g08980 | expressed protein |
| LOC_Os01g09110 | retrotransposon protein, putative, unclassified |
| LOC_Os01g09220 | transposon protein, putative, CACTA, En/Spm sub-class, expressed |
| LOC_Os01g10030 | expressed protein |
| LOC_Os01g11670 | OsSCP2 - Putative Serine Carboxypeptidase homologue, expressed |
| LOC_Os01g11820 | expressed protein |
| LOC_Os01g11970 | expressed protein |
| LOC_Os01g13050 | long cell-linked locus protein, putative, expressed |
| LOC_Os01g13470 | KH domain containing protein, putative, expressed |
| LOC_Os01g14340 | expressed protein |
| LOC_Os01g15420 | retrotransposon protein, putative, Ty3-gypsy subclass, expressed |
| LOC_Os01g16050 | transposon protein, putative, unclassified, expressed |
| LOC_Os01g20850 | expressed protein |
| LOC_Os01g21420 | pre-mRNA-splicing factor SF2, putative, expressed |
| LOC_Os01g21760 | retrotransposon protein, putative, unclassified |
| LOC_Os01g23180 | retrotransposon protein, putative, unclassified |
| LOC_Os01g24030 | AMP-binding enzyme, putative, expressed |
| LOC_Os01g25250 | expressed protein |
| LOC_Os01g25510 | alpha-amylase isozyme C, putative, expressed |
| LOC_Os01g29570 | retrotransposon protein, putative, Ty3-gypsy subclass, expressed |
| LOC_Os01g33740 | retrotransposon protein, putative, unclassified, expressed |
| LOC_Os01g34510 | hypothetical protein |
| LOC_Os01g34620 | OsGrx_S15 |
| LOC_Os01g35230 | 1-aminocyclopropane-1-carboxylate oxidase homolog 1, putative, expressed |
| LOC_Os01g35870 | hypothetical protein |
| LOC_Os01g37040 | retrotransposon protein, putative, Ty1-copia subclass, expressed |
| LOC_Os01g37310 | SWIM zinc finger family protein, putative, expressed |
| LOC_Os01g37320 | expressed protein |
| LOC_Os01g38670 | transporter family protein, putative, expressed |

| | |
|---|---|
| LOC_Os01g38750 | expressed protein |
| LOC_Os01g39530 | retrotransposon, putative, centromere-specific |
| LOC_Os01g40080 | hypothetical protein |
| LOC_Os01g40910 | transposon protein, putative, unclassified |
| LOC_Os01g42600 | hypothetical protein |
| LOC_Os01g42890 | expressed protein |
| LOC_Os01g43290 | expressed protein |
| LOC_Os01g48120 | expressed protein |
| LOC_Os01g49065 | expressed protein |
| LOC_Os01g52920 | expressed protein |
| LOC_Os01g54620 | CESA4 - cellulose synthase, expressed |
| LOC_Os01g55710 | DNA-binding protein, putative, expressed |
| LOC_Os01g57280 | rp1, putative, expressed |
| LOC_Os01g60150 | expressed protein |
| LOC_Os01g60600 | WRKY108, expressed |
| LOC_Os01g61310 | DUF640 domain containing protein, putative, expressed |
| LOC_Os01g61350 | glutaredoxin, putative, expressed |
| LOC_Os01g61760 | myosin heavy chain-related, putative, expressed |
| LOC_Os01g62290 | DnaK family protein, putative, expressed |
| LOC_Os01g62710 | transposon protein, putative, unclassified |
| LOC_Os01g62770 | expressed protein |
| LOC_Os01g64090 | L1P family of ribosomal proteins domain containing protein, expressed |
| LOC_Os01g64640 | histone H3, putative, expressed |
| LOC_Os01g65640 | expressed protein |
| LOC_Os01g65670 | amino acid transporter, putative, expressed |
| LOC_Os01g66020 | protein kinase family protein, putative, expressed |
| LOC_Os01g67620 | expressed protein |
| LOC_Os01g71660 | retrotransposon protein, putative, unclassified, expressed |
| LOC_Os02g01800 | receptor-like protein kinase precursor, putative, expressed |
| LOC_Os02g07110 | OsSAUR6 - Auxin-responsive SAUR gene family member, expressed |

| | |
|---|---|
| LOC_Os02g07120 | HEAT repeat family protein, putative, expressed |
| LOC_Os02g08100 | AMP-binding domain containing protein, expressed |
| LOC_Os02g12140 | retrotransposon protein, putative, Ty3-gypsy subclass, expressed |
| LOC_Os02g12630 | retrotransposon protein, putative, Ty3-gypsy subclass, expressed |
| LOC_Os02g12650 | puromycin-sensitive aminopeptidase, putative, expressed |
| LOC_Os02g16050 | transposon protein, putative, unclassified, expressed |
| LOC_Os02g18064 | expressed protein |
| LOC_Os02g18350 | expressed protein |
| LOC_Os02g19250 | transposon protein, putative, CACTA, En/Spm sub-class, expressed |
| LOC_Os02g19370 | retrotransposon protein, putative, unclassified |
| LOC_Os02g19850 | hAT dimerisation domain-containing protein, putative, expressed |
| LOC_Os02g19860 | pre-mRNA-splicing factor ATP-dependent RNA helicase, putative, expressed |
| LOC_Os02g21680 | expressed protein |
| LOC_Os02g21685 | expressed protein |
| LOC_Os02g22120 | expressed protein |
| LOC_Os02g25290 | retrotransposon protein, putative, unclassified, expressed |
| LOC_Os02g25310 | zinc knuckle family protein, expressed |
| LOC_Os02g25580 | Sec1 family transport protein, putative, expressed |
| LOC_Os02g25610 | expressed protein |
| LOC_Os02g25620 | expressed protein |
| LOC_Os02g25790 | retrotransposon protein, putative, Ty1-copia subclass, expressed |
| LOC_Os02g25900 | pollen signalling protein with adenylyl cyclase activity, putative, expressed |
| LOC_Os02g25960 | ankyrin repeat family protein, putative, expressed |
| LOC_Os02g26130 | hypothetical protein |
| LOC_Os02g26140 | microtubule-binding protein TANGLED1, putative, expressed |
| LOC_Os02g26150 | hypothetical protein |
| LOC_Os02g26160 | receptor lectin kinase like protein, putative, expressed |
| LOC_Os02g26170 | expressed protein |
| LOC_Os02g26320 | fasciclin-like arabinogalactan precursor protein, putative, expressed |
| LOC_Os02g26330 | expressed protein |

| | |
|---|---|
| LOC_Os02g26349 | transposon protein, putative, CACTA, En/Spm sub-class, expressed |
| LOC_Os02g26940 | vegetative storage protein, putative, expressed |
| LOC_Os02g26950 | expressed protein |
| LOC_Os02g26960 | expressed protein |
| LOC_Os02g26970 | hypothetical protein |
| LOC_Os02g26984 | expressed protein |
| LOC_Os02g27060 | high mobility group, putative, expressed |
| LOC_Os02g27070 | PPR repeat domain containing protein, putative, expressed |
| LOC_Os02g27110 | expressed protein |
| LOC_Os02g28920 | transposon protein, putative, unclassified, expressed |
| LOC_Os02g30050 | ribosomal protein L29, putative, expressed |
| LOC_Os02g32380 | expressed protein |
| LOC_Os02g33680 | U-box domain containing protein, expressed |
| LOC_Os02g37654 | lecithin:cholesterol acyltransferase, putative, expressed |
| LOC_Os02g42310 | OsSCP8 - Putative Serine Carboxypeptidase homologue, expressed |
| LOC_Os02g43800 | retrotransposon protein, putative, Ty3-gypsy subclass, expressed |
| LOC_Os02g43960 | expressed protein |
| LOC_Os02g46370 | expressed protein |
| LOC_Os02g48184 | expressed protein |
| LOC_Os02g48660 | 60S ribosomal protein L31, putative, expressed |
| LOC_Os02g51500 | membrane-anchored ubiquitin-fold protein, putative, expressed |
| LOC_Os02g53260 | acetyltransferase, GNAT family, putative, expressed |
| LOC_Os02g54520 | MYB family transcription factor, putative, expressed |
| LOC_Os02g54600 | STE_MEK_ste7_MAP2K |
| LOC_Os02g57530 | ethylene receptor, putative, expressed |
| LOC_Os02g58490 | PINHEAD, putative, expressed |
| LOC_Os02g58554 | expressed protein |
| LOC_Os03g02100 | valyl-tRNA synthetase, putative, expressed |
| LOC_Os03g03890 | protein kinase family protein, putative, expressed |
| LOC_Os03g10240 | DUF677 domain containing protein, putative, expressed |

| | |
|---|---|
| LOC_Os03g10250 | expressed protein |
| LOC_Os03g10260 | expressed protein |
| LOC_Os03g11140 | pleckstrin homology domain-containing protein-related taxo, putative, expressed |
| LOC_Os03g11150 | expressed protein |
| LOC_Os03g11160 | cysteine proteinase inhibitor precursor, putative, expressed |
| LOC_Os03g11170 | cysteine proteinase inhibitor precursor, putative, expressed |
| LOC_Os03g11180 | cysteine proteinase inhibitor 6 precursor, putative, expressed |
| LOC_Os03g11190 | expressed protein |
| LOC_Os03g11200 | pre-mRNA-processing factor 39, putative, expressed |
| LOC_Os03g11210 | DUF538 domain containing protein, putative, expressed |
| LOC_Os03g12730 | receptor protein kinase CLAVATA1 precursor, putative, expressed |
| LOC_Os03g13300 | glutamate decarboxylase, putative, expressed |
| LOC_Os03g13325 | ompA/MotB, putative, expressed |
| LOC_Os03g18940 | exonuclease, putative, expressed |
| LOC_Os03g18950 | zinc finger C-x8-C-x5-C-x3-H type family protein, expressed |
| LOC_Os03g18960 | calmodulin binding protein, putative, expressed |
| LOC_Os03g18970 | protein phosphotase protein, putative, expressed |
| LOC_Os03g18980 | kinesin motor domain containing protein, expressed |
| LOC_Os03g18990 | expressed protein |
| LOC_Os03g19000 | peroxisomal biogenesis factor 11, putative, expressed |
| LOC_Os03g19059 | anaphase-promoting complex subunit 11, putative, expressed |
| LOC_Os03g19080 | expressed protein |
| LOC_Os03g19170 | GCRP7 - Glycine and cysteine rich family protein precursor, expressed |
| LOC_Os03g19180 | GCRP8 - Glycine and cysteine rich family protein precursor, putative, expressed |
| LOC_Os03g19190 | TOPBP1C - Similar to DNA replication protein TOPBP1 from, expressed |
| LOC_Os03g19230 | expressed protein |
| LOC_Os03g20420 | alpha-N-arabinofuranosidase A, putative, expressed |
| LOC_Os03g20500 | OsFBX82 - F-box domain containing protein, expressed |
| LOC_Os03g20520 | endoplasmic reticulum-Golgi intermediate compartment protein 3, putative, expressed |
| LOC_Os03g20750 | outer mitochondrial membrane porin, putative, expressed |

| | |
|---|---|
| LOC_Os03g20760 | LTPL66 - Protease inhibitor/seed storage/LTP family protein precursor, expressed |
| LOC_Os03g20770 | expressed protein |
| LOC_Os03g20830 | hypothetical protein |
| LOC_Os03g20840 | disease resistance RPP13-like protein 1, putative, expressed |
| LOC_Os03g20850 | exostosin family domain containing protein, expressed |
| LOC_Os03g20860 | expressed protein |
| LOC_Os03g20870 | zinc finger, C3HC4 type domain containing protein, expressed |
| LOC_Os03g20880 | pyruvate kinase, putative, expressed |
| LOC_Os03g21000 | thioredoxin, putative, expressed |
| LOC_Os03g21010 | expressed protein |
| LOC_Os03g21020 | RNA recognition motif containing protein, putative, expressed |
| LOC_Os03g21070 | expressed protein |
| LOC_Os03g21080 | guanine nucleotide exchange factor, putative, expressed |
| LOC_Os03g21120 | nucleoside-triphosphatase, putative, expressed |
| LOC_Os03g21140 | RNA-binding zinc finger protein, putative, expressed |
| LOC_Os03g21210 | endoglucanase, putative, expressed |
| LOC_Os03g21220 | expressed protein |
| LOC_Os03g21240 | myb-like DNA-binding domain containing protein, expressed |
| LOC_Os03g21250 | glycosyl transferase, family 8, putative, expressed |
| LOC_Os03g21260 | 2,3-bisphosphoglycerate-independent phosphoglycerate mutase, putative, expressed |
| LOC_Os03g21270 | expressed protein |
| LOC_Os03g21310 | ulp1 protease family, C-terminal catalytic domain containing protein, expressed |
| LOC_Os03g21330 | transposon protein, putative, unclassified, expressed |
| LOC_Os03g21480 | HAD superfamily phosphatase, putative, expressed |
| LOC_Os03g21490 | ABC transporter, ATP-binding protein, putative, expressed |
| LOC_Os03g21500 | expressed protein |
| LOC_Os03g21510 | inactive receptor kinase At2g26730 precursor, putative, expressed |
| LOC_Os03g21570 | DNA-binding protein, putative, expressed |
| LOC_Os03g21580 | expressed protein |
| LOC_Os03g21680 | phospholipid-transporting ATPase, putative, expressed |

| | |
|---|---|
| LOC_Os03g21690 | lung seven transmembrane domain containing protein, putative, expressed |
| LOC_Os03g21720 | vacuolar-sorting receptor precursor, putative, expressed |
| LOC_Os03g21730 | receptor-like protein kinase precursor, putative, expressed |
| LOC_Os03g24650 | terpene synthase family, metal binding domain containing protein, expressed |
| LOC_Os03g25070 | CAMK_CAMK_like |
| LOC_Os03g27230 | phospho-2-dehydro-3-deoxyheptonate aldolase, chloroplast precursor, putative, expressed |
| LOC_Os03g27250 | OsFBO14 - F-box and other domain containing protein, expressed |
| LOC_Os03g27260 | 40S ribosomal protein S6, putative, expressed |
| LOC_Os03g27280 | CAMK_CAMK_like |
| LOC_Os03g27290 | cytochrome c oxidase subunit, putative, expressed |
| LOC_Os03g27320 | steroid binding protein, putative, expressed |
| LOC_Os03g27340 | expressed protein |
| LOC_Os03g27350 | cysteine protease ATG4, putative, expressed |
| LOC_Os03g27590 | OsSCP18 - Putative Serine Carboxypeptidase homologue, expressed |
| LOC_Os03g30000 | glycosyl transferase 8 domain containing protein, putative, expressed |
| LOC_Os03g36490 | retrotransposon protein, putative, Ty3-gypsy subclass, expressed |
| LOC_Os03g36500 | expressed protein |
| LOC_Os03g36520 | retrotransposon, putative, centromere-specific, expressed |
| LOC_Os03g39150 | protein kinase domain containing protein, expressed |
| LOC_Os03g42235 | expressed protein |
| LOC_Os03g43360 | zinc finger, C3HC4 type domain containing protein, expressed |
| LOC_Os03g43390 | F-box/LRR domain containing protein, putative, expressed |
| LOC_Os03g43420 | single-stranded DNA-binding protein, putative, expressed |
| LOC_Os03g43430 | mttA/Hcf106 family protein, putative, expressed |
| LOC_Os03g43440 | CAMK_KIN1/SNF1/Nim1_like |
| LOC_Os03g43460 | hypothetical protein |
| LOC_Os03g43464 | hypothetical protein |
| LOC_Os03g43470 | pentatricopeptide, putative, expressed |
| LOC_Os03g43480 | AGG2, putative, expressed |
| LOC_Os03g43500 | expressed protein |

| | |
|---|---|
| LOC_Os03g43510 | expressed protein |
| LOC_Os03g43524 | expressed protein |
| LOC_Os03g43570 | UBX domain-containing protein 1, putative, expressed |
| LOC_Os03g43580 | IQ calmodulin-binding motif family protein, putative, expressed |
| LOC_Os03g43590 | LSTK-1-like kinase, putative, expressed |
| LOC_Os03g43620 | hypothetical protein |
| LOC_Os03g56690 | expressed protein |
| LOC_Os03g56700 | expressed protein |
| LOC_Os03g62600 | ras-related protein, putative, expressed |
| LOC_Os04g16270 | nucleic acid binding protein, putative, expressed |
| LOC_Os04g16820 | DNA-directed RNA polymerase subunit beta, putative, expressed |
| LOC_Os04g24110 | anthocyanin 3-O-beta-glucosyltransferase, putative, expressed |
| LOC_Os04g25784 | expressed protein |
| LOC_Os04g28074 | expressed protein |
| LOC_Os04g31260 | hypothetical protein |
| LOC_Os04g34120 | expressed protein |
| LOC_Os04g34150 | expressed protein |
| LOC_Os04g34160 | expressed protein |
| LOC_Os04g34180 | expressed protein |
| LOC_Os04g39530 | transposon protein, putative, CACTA, En/Spm sub-class, expressed |
| LOC_Os04g40600 | peptide methionine sulfoxide reductase, putative, expressed |
| LOC_Os04g41720 | retrotransposon, putative, centromere-specific, expressed |
| LOC_Os04g46540 | expressed protein |
| LOC_Os04g55050 | amidase, putative, expressed |
| LOC_Os04g55060 | 3-oxoacyl-synthase III, chloroplast precursor, putative, expressed |
| LOC_Os04g55070 | gibberellin 20 oxidase 2, putative, expressed |
| LOC_Os04g55080 | cyclic nucleotide-gated ion channel, putative, expressed |
| LOC_Os04g55090 | PPR repeat domain containing protein, putative, expressed |
| LOC_Os04g55100 | expressed protein |
| LOC_Os04g55110 | expressed protein |

| | |
|---|---|
| LOC_Os04g55120 | jp18, putative, expressed |
| LOC_Os04g55180 | hydrolase, alpha/beta fold family domain containing protein, expressed |
| LOC_Os04g55190 | SPOC domain containing protein, putative, expressed |
| LOC_Os04g55200 | coatomer subunit epsilon, putative, expressed |
| LOC_Os04g55210 | chloride channel protein, putative, expressed |
| LOC_Os04g55220 | C2 domain containing protein, expressed |
| LOC_Os04g55230 | tetratricopeptide repeat domain containing protein, putative, expressed |
| LOC_Os04g55240 | VQ domain containing protein, putative, expressed |
| LOC_Os04g55510 | zinc finger, C3HC4 type domain containing protein, expressed |
| LOC_Os04g56620 | molybdopterin biosynthesis protein CNX1, putative, expressed |
| LOC_Os04g56640 | mediator of RNA polymerase II transcription subunit 7, putative, expressed |
| LOC_Os04g56646 | MEE34, putative, expressed |
| LOC_Os04g56654 | BSD domain-containing protein, putative, expressed |
| LOC_Os04g56670 | S4 domain containing protein, putative, expressed |
| LOC_Os04g56680 | OsSAUR22 - Auxin-responsive SAUR gene family member, expressed |
| LOC_Os04g56700 | naringenin,2-oxoglutarate 3-dioxygenase, putative, expressed |
| LOC_Os04g56710 | GTP cyclohydrolase I 1, putative, expressed |
| LOC_Os04g57550 | amine oxidase, flavin-containing, domain containing protein, expressed |
| LOC_Os04g57560 | amine oxidase, flavin-containing, domain containing protein, expressed |
| LOC_Os04g57600 | zinc finger C-x8-C-x5-C-x3-H type family protein, expressed |
| LOC_Os05g03240 | hypothetical protein |
| LOC_Os05g08380 | myosin heavy chain-related, putative, expressed |
| LOC_Os05g08390 | expressed protein |
| LOC_Os05g08410 | ATP10 protein, expressed |
| LOC_Os05g08414 | expressed protein |
| LOC_Os05g09120 | hypothetical protein |
| LOC_Os05g09160 | expressed protein |
| LOC_Os05g09200 | expressed protein |
| LOC_Os05g12460 | expressed protein |
| LOC_Os05g22960 | expressed protein |

| | |
|---|---|
| LOC_Os05g23450 | expressed protein |
| LOC_Os05g29880 | 3-hydroxyacyl-CoA dehydrogenase, putative, expressed |
| LOC_Os05g29930 | late embryogenesis abundant protein, putative, expressed |
| LOC_Os05g30130 | chaperone protein dnaJ 49, putative, expressed |
| LOC_Os05g30160 | expressed protein |
| LOC_Os05g30180 | hypothetical protein |
| LOC_Os05g30190 | phytochrome A-associated F-box protein, putative, expressed |
| LOC_Os05g30200 | expressed protein |
| LOC_Os05g30210 | expressed protein |
| LOC_Os05g30240 | PPR repeat domain containing protein, putative, expressed |
| LOC_Os05g30250 | Os5bglu19 - beta-glucosidase homologue, similar to G |
| LOC_Os05g30470 | collagen alpha-1, putative, expressed |
| LOC_Os05g34820 | expressed protein |
| LOC_Os05g34920 | expressed protein |
| LOC_Os05g34930 | hypothetical protein |
| LOC_Os05g37810 | expressed protein |
| LOC_Os06g03520 | DUF581 domain containing protein, expressed |
| LOC_Os06g06850 | resistance protein LR10, putative, expressed |
| LOC_Os06g08910 | glutamate receptor 2 |
| LOC_Os06g09150 | transposon protein, putative, CACTA, En/Spm sub-class, expressed |
| LOC_Os06g09160 | expressed protein |
| LOC_Os06g09170 | zinc ion binding protein, putative, expressed |
| LOC_Os06g14860 | hypothetical protein |
| LOC_Os06g15930 | retrotransposon protein, putative, unclassified, expressed |
| LOC_Os06g20960 | SAM dependent carboxyl methyltransferase, putative, expressed |
| LOC_Os06g21099 | expressed protein |
| LOC_Os06g21210 | glycine rich protein family protein, putative, expressed |
| LOC_Os06g21470 | expressed protein |
| LOC_Os06g21530 | expressed protein |
| LOC_Os06g21540 | inhibitor I family protein, putative, expressed |

| | |
|---|---|
| LOC_Os06g21550 | thioredoxin domain-containing protein 17, putative, expressed |
| LOC_Os06g21560 | CDK5RAP3, putative, expressed |
| LOC_Os06g21570 | Os6bglu24 - beta-glucosidase homologue, similar to G |
| LOC_Os06g21580 | expressed protein |
| LOC_Os06g21590 | chlorophyll A-B binding protein, putative, expressed |
| LOC_Os06g21600 | selenoprotein O, putative, expressed |
| LOC_Os06g21770 | hypothetical protein |
| LOC_Os06g22390 | expressed protein |
| LOC_Os06g22394 | expressed protein |
| LOC_Os06g28660 | retrotransposon protein, putative, unclassified, expressed |
| LOC_Os06g31800 | THION2 - Plant thionin family protein precursor, expressed |
| LOC_Os06g31810 | hypothetical protein |
| LOC_Os06g38115 | expressed protein |
| LOC_Os06g38590 | receptor-like protein kinase precursor, putative, expressed |
| LOC_Os06g38660 | expressed protein |
| LOC_Os06g38710 | expressed protein |
| LOC_Os06g38720 | expressed protein |
| LOC_Os06g38740 | expressed protein |
| LOC_Os06g44460 | D-3-phosphoglycerate dehydrogenase, chloroplast precursor, putative, expressed |
| LOC_Os06g49920 | expressed protein |
| LOC_Os06g49930 | OsFBX207 - F-box domain containing protein, expressed |
| LOC_Os06g49980 | expressed protein |
| LOC_Os06g50050 | AAA-type ATPase family protein, putative, expressed |
| LOC_Os06g50070 | expressed protein |
| LOC_Os06g50090 | expressed protein |
| LOC_Os06g50100 | tyrosine protein kinase domain containing protein, putative, expressed |
| LOC_Os06g50110 | expressed protein |
| LOC_Os06g50120 | novel plant SNARE 11, putative, expressed |
| LOC_Os06g50130 | expressed protein |
| LOC_Os06g50140 | endoglucanase, putative, expressed |

| | |
|---|---|
| LOC_Os06g50146 | calcium-dependent protein kinase CPK1 adapter protein 2, putative, expressed |
| LOC_Os06g50170 | BRE, putative, expressed |
| LOC_Os06g50180 | DUF803 domain containing, putative, expressed |
| LOC_Os06g50330 | senescence-associated protein, putative, expressed |
| LOC_Os06g50350 | expressed protein |
| LOC_Os06g50360 | pseudouridine synthase family protein, putative, expressed |
| LOC_Os06g51084 | 1,4-alpha-glucan-branching enzyme, chloroplast precursor, putative, expressed |
| LOC_Os06g51120 | expressed protein |
| LOC_Os06g51130 | spotted leaf 11, putative, expressed |
| LOC_Os06g51140 | ZOS6-11 - C2H2 zinc finger protein, expressed |
| LOC_Os06g51150 | catalase isozyme B, putative, expressed |
| LOC_Os06g51170 | serine/threonine-protein kinase Cx32, chloroplast precursor, putative, expressed |
| LOC_Os06g51230 | expressed protein |
| LOC_Os06g51240 | expressed protein |
| LOC_Os06g51250 | EF hand family protein, putative, expressed |
| LOC_Os06g51260 | MYB family transcription factor, putative, expressed |
| LOC_Os06g51280 | acetolactate synthase 1, chloroplast precursor, putative, expressed |
| LOC_Os06g51290 | phytoene synthase, chloroplast precursor, putative, expressed |
| LOC_Os06g51320 | GASR8 - Gibberellin-regulated GASA/GAST/Snakin family protein precursor, expressed |
| LOC_Os06g51350 | IPP transferase, putative, expressed |
| LOC_Os06g51360 | lysM domain containing protein, putative, expressed |
| LOC_Os06g51450 | PHD-finger domain containing protein, expressed |
| LOC_Os06g51460 | white-brown complex homolog protein, putative, expressed |
| LOC_Os06g51470 | expressed protein |
| LOC_Os06g51480 | expressed protein |
| LOC_Os06g51490 | PHD-finger domain containing protein, putative, expressed |
| LOC_Os06g51500 | expressed protein |
| LOC_Os06g51510 | expressed protein |
| LOC_Os06g51520 | lysine ketoglutarate reductase trans-splicing related 1, putative, expressed |
| LOC_Os07g03170 | expressed protein |

| | |
|---|---|
| LOC_Os07g03840 | lectin-like receptor kinase, putative, expressed |
| LOC_Os07g10210 | expressed protein |
| LOC_Os07g13090 | expressed protein |
| LOC_Os07g14750 | 60S acidic ribosomal protein, putative, expressed |
| LOC_Os07g14760 | expressed protein |
| LOC_Os07g19560 | TATA-binding protein-associated factor 2N, putative, expressed |
| LOC_Os07g24930 | retrotransposon protein, putative, Ty3-gypsy subclass, expressed |
| LOC_Os07g29040 | hypothetical protein |
| LOC_Os07g29050 | expressed protein |
| LOC_Os07g44220 | expressed protein |
| LOC_Os07g44830 | proline-rich family protein, putative, expressed |
| LOC_Os07g48602 | EMB1011, putative, expressed |
| LOC_Os07g48610 | hydrolase, alpha/beta fold family domain containing protein, expressed |
| LOC_Os07g48620 | expressed protein |
| LOC_Os07g48690 | DUF630/DUF632 domains containing protein, putative, expressed |
| LOC_Os07g48700 | expressed protein |
| LOC_Os07g48710 | VQ domain containing protein, putative, expressed |
| LOC_Os07g48760 | CAMK_KIN1/SNF1/Nim1_like |
| LOC_Os07g48770 | serine hydrolase domain containing protein, expressed |
| LOC_Os08g16359 | expressed protein |
| LOC_Os08g18930 | expressed protein |
| LOC_Os08g24900 | retrotransposon protein, putative, unclassified, expressed |
| LOC_Os08g26210 | expressed protein |
| LOC_Os08g26460 | expressed protein |
| LOC_Os08g34249 | inhibitor I family protein, putative, expressed |
| LOC_Os08g34610 | hypothetical protein |
| LOC_Os08g34620 | expressed protein |
| LOC_Os08g34640 | receptor-like protein kinase precursor, putative, expressed |
| LOC_Os08g34700 | GDU1, putative, expressed |
| LOC_Os08g34710 | expressed protein |

| | |
|---|---|
| LOC_Os08g34730 | expressed protein |
| LOC_Os08g34740 | SGT1 protein, putative, expressed |
| LOC_Os08g34750 | serine esterase family protein, putative, expressed |
| LOC_Os08g34760 | expressed protein |
| LOC_Os08g34780 | adenylyl cyclase-associated protein, putative, expressed |
| LOC_Os08g35110 | OsSAUR33 - Auxin-responsive SAUR gene family member, expressed |
| LOC_Os08g35130 | expressed protein |
| LOC_Os08g39174 | retrotransposon protein, putative, Ty1-copia subclass, expressed |
| LOC_Os09g04710 | GDSL-like lipase/acylhydrolase, putative, expressed |
| LOC_Os09g09820 | retrotransposon protein, putative, unclassified, expressed |
| LOC_Os09g21620 | OsFBX325 - F-box domain containing protein, expressed |
| LOC_Os09g22040 | expressed protein |
| LOC_Os09g37990 | expressed protein |
| LOC_Os11g03240 | MATE efflux family protein, putative, expressed |
| LOC_Os11g03430 | CDC45B - Putative DNA replication initiation protein, expressed |
| LOC_Os11g03460 | Mad3/BUB1 homology region 1 domain containing protein, expressed |
| LOC_Os11g04130 | nodulin MtN21 family protein, putative, expressed |
| LOC_Os11g04770 | EF hand family protein, putative, expressed |
| LOC_Os11g04950 | splicing factor, putative, expressed |
| LOC_Os11g04954 | DNA repair protein Rad51, putative, expressed |
| LOC_Os11g04980 | acyl-CoA synthetase protein, putative, expressed |
| LOC_Os11g05260 | phosphoglycerate mutase, putative, expressed |
| LOC_Os11g05340 | transposon protein, putative, unclassified, expressed |
| LOC_Os11g22890 | expressed protein |
| LOC_Os11g47410 | retrotransposon protein, putative, unclassified, expressed |
| LOC_Os12g04630 | expressed protein |
| LOC_Os12g04770 | expressed protein |
| LOC_Os12g04790 | expressed protein |
| LOC_Os12g04970 | retrotransposon protein, putative, unclassified, expressed |
| LOC_Os12g05270 | zinc finger, C3HC4 type domain containing protein, expressed |

| | |
|---|---|
| LOC_Os12g05310 | retrotransposon protein, putative, unclassified, expressed |
| LOC_Os12g05320 | expressed protein |
| LOC_Os12g05330 | protein of unknown function containing protein, expressed |
| LOC_Os12g05340 | expressed protein |
| LOC_Os12g05410 | G10 protein, putative, expressed |
| LOC_Os12g05470 | plastocyanin-like domain containing protein, putative, expressed |
| LOC_Os12g23550 | expressed protein |
| LOC_Os12g40940 | expressed protein |
| LOC_Os12g40980 | expressed protein |
| LOC_Os12g41050 | hypothetical protein |

**Genes in indica_SV CLR**

| | |
|---|---|
| LOC_Os01g01090 | retrotransposon protein, putative, unclassified, expressed |
| LOC_Os01g01100 | retrotransposon protein, putative, Ty3-gypsy subclass, expressed |
| LOC_Os01g02629 | retrotransposon protein, putative, unclassified, expressed |
| LOC_Os01g02990 | expressed protein |
| LOC_Os01g03610 | retrotransposon protein, putative, unclassified, expressed |
| LOC_Os01g03830 | retrotransposon protein, putative, unclassified, expressed |
| LOC_Os01g04790 | retrotransposon protein, putative, unclassified |
| LOC_Os01g05320 | retrotransposon protein, putative, Ty1-copia subclass, expressed |
| LOC_Os01g06680 | retrotransposon protein, putative, unclassified, expressed |
| LOC_Os01g09110 | retrotransposon protein, putative, unclassified |
| LOC_Os01g10170 | expressed protein |
| LOC_Os01g10970 | auxin-induced protein 5NG4, putative, expressed |
| LOC_Os01g11270 | cytochrome P450, putative, expressed |
| LOC_Os01g11720 | transposable element protein, putative, containing Pfam profile: PF03004, Transposase_24, expressed |
| LOC_Os01g12710 | oxidoreductase, short chain dehydrogenase/reductase family domain containing protein, expressed |
| LOC_Os01g13229 | cyclin-A1, putative, expressed |
| LOC_Os01g13250 | sarcoma antigen NY-SAR-91, putative, expressed |
| LOC_Os01g13300 | B3 DNA binding domain containing protein, expressed |

| | |
|---|---|
| LOC_Os01g13440 | pollen allergen Cyn d 23, putative, expressed |
| LOC_Os01g14730 | retrotransposon protein, putative, unclassified |
| LOC_Os01g15080 | hypothetical protein |
| LOC_Os01g15190 | retrotransposon protein, putative, unclassified, expressed |
| LOC_Os01g17010 | phospholipid-transporting ATPase, putative, expressed |
| LOC_Os01g17130 | retrotransposon protein, putative, Ty3-gypsy subclass, expressed |
| LOC_Os01g18020 | retrotransposon protein, putative, unclassified |
| LOC_Os01g19010 | retrotransposon protein, putative, unclassified, expressed |
| LOC_Os01g19640 | retrotransposon protein, putative, Ty3-gypsy subclass |
| LOC_Os01g19970 | MYB family transcription factor, putative, expressed |
| LOC_Os01g19980 | hypothetical protein |
| LOC_Os01g19990 | expressed protein |
| LOC_Os01g20000 | expressed protein |
| LOC_Os01g20010 | expressed protein |
| LOC_Os01g20020 | hypothetical protein |
| LOC_Os01g20030 | expressed protein |
| LOC_Os01g20060 | expressed protein |
| LOC_Os01g20100 | expressed protein |
| LOC_Os01g21360 | retrotransposon protein, putative, unclassified |
| LOC_Os01g21490 | transposon protein, putative, CACTA, En/Spm sub-class |
| LOC_Os01g21550 | expressed protein |
| LOC_Os01g22050 | retrotransposon protein, putative, unclassified, expressed |
| LOC_Os01g22309 | hypothetical protein |
| LOC_Os01g23239 | retrotransposon protein, putative, Ty3-gypsy subclass, expressed |
| LOC_Os01g23320 | retrotransposon, putative, centromere-specific, expressed |
| LOC_Os01g23340 | expressed protein |
| LOC_Os01g23480 | expressed protein |
| LOC_Os01g23940 | expressed protein |
| LOC_Os01g24400 | retrotransposon protein, putative, unclassified, expressed |
| LOC_Os01g25020 | expressed protein |

| | |
|---|---|
| LOC_Os01g25030 | dirigent, putative |
| LOC_Os01g25040 | transposon protein, putative, CACTA, En/Spm sub-class, expressed |
| LOC_Os01g25050 | expressed protein |
| LOC_Os01g25060 | transposon protein, putative, CACTA, En/Spm sub-class, expressed |
| LOC_Os01g25065 | ATP synthase subunit beta, putative, expressed |
| LOC_Os01g25070 | expressed protein |
| LOC_Os01g25080 | expressed protein |
| LOC_Os01g25940 | expressed protein |
| LOC_Os01g27070 | expressed protein |
| LOC_Os01g28270 | retrotransposon protein, putative, Ty1-copia subclass |
| LOC_Os01g28650 | retrotransposon, putative, centromere-specific, expressed |
| LOC_Os01g28882 | retrotransposon protein, putative, unclassified, expressed |
| LOC_Os01g28930 | expressed protein |
| LOC_Os01g28940 | hypothetical protein |
| LOC_Os01g28970 | retrotransposon protein, putative, Ty3-gypsy subclass, expressed |
| LOC_Os01g28980 | proton-dependent oligopeptide transport, putative, expressed |
| LOC_Os01g28989 | expressed protein |
| LOC_Os01g29040 | retrotransposon protein, putative, unclassified, expressed |
| LOC_Os01g29507 | retrotransposon, putative, centromere-specific, expressed |
| LOC_Os01g29790 | retrotransposon protein, putative, unclassified, expressed |
| LOC_Os01g29890 | expressed protein |
| LOC_Os01g30970 | PMR5, putative, expressed |
| LOC_Os01g31050 | expressed protein |
| LOC_Os01g31160 | retrotransposon protein, putative, Ty3-gypsy subclass, expressed |
| LOC_Os01g31170 | expressed protein |
| LOC_Os01g31310 | expressed protein |
| LOC_Os01g32530 | transposon protein, putative, CACTA, En/Spm sub-class |
| LOC_Os01g32550 | hypothetical protein |
| LOC_Os01g32610 | expressed protein |
| LOC_Os01g33142 | hypothetical protein |

| | |
|---|---|
| LOC_Os01g33180 | hypothetical protein |
| LOC_Os01g33470 | retrotransposon protein, putative, unclassified, expressed |
| LOC_Os01g33740 | retrotransposon protein, putative, unclassified, expressed |
| LOC_Os01g34510 | hypothetical protein |
| LOC_Os01g35610 | expressed protein |
| LOC_Os01g35789 | expressed protein |
| LOC_Os01g36410 | expressed protein |
| LOC_Os01g37340 | retrotransposon protein, putative, unclassified, expressed |
| LOC_Os01g38120 | plant protein of unknown function domain containing protein, expressed |
| LOC_Os01g39136 | retrotransposon protein, putative, unclassified, expressed |
| LOC_Os01g39980 | hypothetical protein |
| LOC_Os01g40140 | expressed protein |
| LOC_Os01g40910 | transposon protein, putative, unclassified |
| LOC_Os01g41100 | retrotransposon protein, putative, unclassified, expressed |
| LOC_Os01g42610 | terpene synthase, putative, expressed |
| LOC_Os01g47300 | retrotransposon protein, putative, unclassified, expressed |
| LOC_Os01g48830 | expressed protein |
| LOC_Os01g49154 | expressed protein |
| LOC_Os01g52920 | expressed protein |
| LOC_Os01g56840 | DUF domain containing protein, expressed |
| LOC_Os01g61460 | leaf senescence related protein, putative, expressed |
| LOC_Os01g63910 | AT hook motif family protein, expressed |
| LOC_Os01g65130 | peptide transporter, putative, expressed |
| LOC_Os01g71390 | retrotransposon protein, putative, unclassified, expressed |
| LOC_Os01g71850 | transposon protein, putative, unclassified, expressed |
| LOC_Os01g72400 | retrotransposon protein, putative, Ty1-copia subclass, expressed |
| LOC_Os01g73332 | retrotransposon protein, putative, Ty3-gypsy subclass |
| LOC_Os01g73440 | expressed protein |
| LOC_Os02g04725 | dolichol phosphate-mannose biosynthesis regulatory protein, putative, expressed |
| LOC_Os02g06640 | ubiquitin family protein, putative, expressed |

| | |
|---|---|
| LOC_Os02g09580 | OsFBX39 - F-box domain containing protein, expressed |
| LOC_Os02g09860 | retrotransposon protein, putative, Ty3-gypsy subclass |
| LOC_Os02g11110 | flavonol-3-O-glycoside-7-O-glucosyltransferase 1, putative, expressed |
| LOC_Os02g12140 | retrotransposon protein, putative, Ty3-gypsy subclass, expressed |
| LOC_Os02g16220 | expressed protein |
| LOC_Os02g17020 | hypothetical protein |
| LOC_Os02g18000 | disease resistance protein RGA2, putative, expressed |
| LOC_Os02g18064 | expressed protein |
| LOC_Os02g18115 | expressed protein |
| LOC_Os02g18130 | expressed protein |
| LOC_Os02g18150 | expressed protein |
| LOC_Os02g18160 | expressed protein |
| LOC_Os02g18784 | expressed protein |
| LOC_Os02g20520 | retrotransposon, putative, centromere-specific |
| LOC_Os02g22460 | retrotransposon protein, putative, Ty3-gypsy subclass, expressed |
| LOC_Os02g24090 | retrotransposon protein, putative, unclassified, expressed |
| LOC_Os02g25010 | hypothetical protein |
| LOC_Os02g25290 | retrotransposon protein, putative, unclassified, expressed |
| LOC_Os02g25990 | retrotransposon, putative, centromere-specific |
| LOC_Os02g26190 | transposon protein, putative, CACTA, En/Spm sub-class, expressed |
| LOC_Os02g27910 | expressed protein |
| LOC_Os02g29570 | retrotransposon protein, putative, Ty3-gypsy subclass, expressed |
| LOC_Os02g30760 | expressed protein |
| LOC_Os02g31845 | expressed protein |
| LOC_Os02g32690 | pleiotropic drug resistance protein 15, putative, expressed |
| LOC_Os02g36830 | cytokinin-O-glucosyltransferase 2, putative, expressed |
| LOC_Os02g36840 | cytokinin-O-glucosyltransferase 2, putative, expressed |
| LOC_Os02g39910 | B4-BTB1 - Bric-a-Brac, Tramtrack, Broad Complex BTB domain with B4 subfamily conserved sequence, expressed |
| LOC_Os02g41961 | expressed protein |
| LOC_Os02g42180 | retrotransposon protein, putative, unclassified, expressed |

| | |
|---|---|
| LOC_Os02g43800 | retrotransposon protein, putative, Ty3-gypsy subclass, expressed |
| LOC_Os02g44550 | NADP-dependent malic enzyme, putative, expressed |
| LOC_Os02g45370 | retrotransposon protein, putative, Ty3-gypsy subclass, expressed |
| LOC_Os02g47680 | retrotransposon protein, putative, Ty3-gypsy subclass |
| LOC_Os02g48184 | expressed protein |
| LOC_Os02g50120 | retrotransposon protein, putative, Ty3-gypsy subclass |
| LOC_Os02g50610 | retrotransposon protein, putative, Ty3-gypsy subclass, expressed |
| LOC_Os03g03840 | expressed protein |
| LOC_Os03g05220 | expressed protein |
| LOC_Os03g06610 | expressed protein |
| LOC_Os03g07500 | hypothetical protein |
| LOC_Os03g21010 | expressed protein |
| LOC_Os03g28180 | expressed protein |
| LOC_Os03g28270 | Leucine Rich Repeat family protein, expressed |
| LOC_Os03g31500 | expressed protein |
| LOC_Os03g32940 | retrotransposon protein, putative, unclassified, expressed |
| LOC_Os03g33384 | hypothetical protein |
| LOC_Os03g35849 | expressed protein |
| LOC_Os03g41010 | hypothetical protein |
| LOC_Os03g41612 | ribosomal protein L25, putative, expressed |
| LOC_Os03g41640 | GRF zinc finger family protein, expressed |
| LOC_Os03g41675 | expressed protein |
| LOC_Os03g41690 | expressed protein |
| LOC_Os03g41780 | retrotransposon, putative, centromere-specific, expressed |
| LOC_Os03g41800 | transposon protein, putative, unclassified, expressed |
| LOC_Os03g41820 | expressed protein |
| LOC_Os03g41870 | nucleic acid binding protein, putative |
| LOC_Os03g44910 | EBNA-1, putative, expressed |
| LOC_Os03g44920 | OsFBX98 - F-box domain containing protein, expressed |
| LOC_Os03g48364 | retrotransposon protein, putative, unclassified |

| | |
|---|---|
| LOC_Os03g55860 | retrotransposon protein, putative, unclassified, expressed |
| LOC_Os03g57750 | expressed protein |
| LOC_Os03g57760 | 25 |
| LOC_Os03g57770 | expressed protein |
| LOC_Os03g57780 | protein kinase, putative, expressed |
| LOC_Os03g58790 | ATPase, putative, expressed |
| LOC_Os03g59770 | EF hand family protein, putative, expressed |
| LOC_Os03g59840 | expressed protein |
| LOC_Os03g59940 | expressed protein |
| LOC_Os03g59950 | receptor-like kinase, putative, expressed |
| LOC_Os03g64014 | expressed protein |
| LOC_Os04g03120 | retrotransposon protein, putative, unclassified, expressed |
| LOC_Os04g03220 | expressed protein |
| LOC_Os04g04650 | hypothetical protein |
| LOC_Os04g12390 | transposon protein, putative, CACTA, En/Spm sub-class, expressed |
| LOC_Os04g12430 | expressed protein |
| LOC_Os04g12460 | Leucine Rich Repeat family protein, expressed |
| LOC_Os04g13080 | retrotransposon, putative, centromere-specific, expressed |
| LOC_Os04g14680 | OsAPx3 - Peroxisomal Ascorbate Peroxidase encoding gene 5,8, expressed |
| LOC_Os04g14690 | flavin-containing monooxygenase family protein, putative, expressed |
| LOC_Os04g14730 | expressed protein |
| LOC_Os04g18380 | cytochrome P450, putative, expressed |
| LOC_Os04g19250 | hypothetical protein |
| LOC_Os04g22110 | retrotransposon protein, putative, unclassified, expressed |
| LOC_Os04g24950 | hypothetical protein |
| LOC_Os04g25060 | cysteine-rich receptor-like protein kinase, putative, expressed |
| LOC_Os04g25100 | transposon protein, putative, unclassified, expressed |
| LOC_Os04g25130 | retrotransposon protein, putative, LINE subclass, expressed |
| LOC_Os04g25150 | pollen allergen, putative, expressed |
| LOC_Os04g25180 | hypothetical protein |

| | |
|---|---|
| LOC_Os04g25230 | hypothetical protein |
| LOC_Os04g25270 | hypothetical protein |
| LOC_Os04g25290 | expressed protein |
| LOC_Os04g25350 | expressed protein |
| LOC_Os04g25360 | tRNA methyltransferase complex GCD14 subunit, putative, expressed |
| LOC_Os04g25810 | retrotransposon protein, putative, unclassified |
| LOC_Os04g26330 | expressed protein |
| LOC_Os04g26380 | aspartic proteinase-like protein 2 precursor, putative, expressed |
| LOC_Os04g26400 | expressed protein |
| LOC_Os04g26440 | hypothetical protein |
| LOC_Os04g26450 | expressed protein |
| LOC_Os04g26520 | hypothetical protein |
| LOC_Os04g26590 | expressed protein |
| LOC_Os04g28450 | hypothetical protein |
| LOC_Os04g28710 | hypothetical protein |
| LOC_Os04g28720 | RALFL32 - Rapid ALkalinization Factor RALF family protein precursor, putative, expressed |
| LOC_Os04g33910 | expressed protein |
| LOC_Os04g40600 | peptide methionine sulfoxide reductase, putative, expressed |
| LOC_Os04g44610 | white-brown complex homolog protein 11, putative, expressed |
| LOC_Os05g03810 | trehalose phosphatase, putative, expressed |
| LOC_Os05g13530 | expressed protein |
| LOC_Os05g13540 | hypothetical protein |
| LOC_Os05g15200 | zinc knuckle family protein, expressed |
| LOC_Os05g19110 | expressed protein |
| LOC_Os05g19120 | hypothetical protein |
| LOC_Os05g19130 | hypothetical protein |
| LOC_Os05g24450 | hypothetical protein |
| LOC_Os05g25940 | expressed protein |
| LOC_Os05g27980 | DUF260 domain containing protein, putative, expressed |
| LOC_Os05g32020 | expressed protein |

| | |
|---|---|
| LOC_Os05g39630 | expressed protein |
| LOC_Os06g09470 | transposon protein, putative, CACTA, En/Spm sub-class |
| LOC_Os06g11670 | expressed protein |
| LOC_Os06g13910 | expressed protein |
| LOC_Os06g25250 | endoribonuclease Dicer, putative, expressed |
| LOC_Os06g25294 | 60S ribosomal protein L18a-1, putative, expressed |
| LOC_Os06g25470 | expressed protein |
| LOC_Os06g25510 | expressed protein |
| LOC_Os06g25540 | expressed protein |
| LOC_Os06g26080 | expressed protein |
| LOC_Os06g26120 | hypothetical protein |
| LOC_Os06g34530 | zinc finger, C3HC4 type domain containing protein, expressed |
| LOC_Os06g34950 | expressed protein |
| LOC_Os06g41210 | hypothetical protein |
| LOC_Os06g43070 | expressed protein |
| LOC_Os07g05260 | expressed protein |
| LOC_Os07g05290 | expressed protein |
| LOC_Os07g15110 | retrotransposon protein, putative, unclassified, expressed |
| LOC_Os07g15610 | hypothetical protein |
| LOC_Os07g19340 | retrotransposon protein, putative, unclassified, expressed |
| LOC_Os08g05040 | expressed protein |
| LOC_Os08g05440 | NB-ARC domain containing protein, expressed |
| LOC_Os08g16750 | hypothetical protein |
| LOC_Os08g26210 | expressed protein |
| LOC_Os08g31970 | NHL repeat-containing protein, putative, expressed |
| LOC_Os08g31990 | expressed protein |
| LOC_Os08g33090 | inactive receptor kinase At2g26730 precursor, putative, expressed |
| LOC_Os09g04470 | retrotransposon protein, putative, Ty3-gypsy subclass |
| LOC_Os09g04490 | inactive receptor kinase At1g27190 precursor, putative, expressed |
| LOC_Os09g04500 | expressed protein |

| | |
|---|---|
| LOC_Os09g06310 | hypothetical protein |
| LOC_Os09g06340 | retrotransposon protein, putative, unclassified |
| LOC_Os09g11020 | NBS-LRR disease resistance protein, putative, expressed |
| LOC_Os09g12970 | plant protein of unknown function domain containing protein, expressed |
| LOC_Os09g16720 | expressed protein |
| LOC_Os09g20684 | SMP-30/Gluconolaconase/LRE-like region containing protein, expressed |
| LOC_Os09g22160 | expressed protein |
| LOC_Os09g24190 | expressed protein |
| LOC_Os10g04180 | NB-ARC domain containing protein, expressed |
| LOC_Os10g04210 | expressed protein |
| LOC_Os10g14814 | ATFMN/FHY, putative, expressed |
| LOC_Os10g15230 | retrotransposon protein, putative, Ty3-gypsy subclass |
| LOC_Os10g19912 | expressed protein |
| LOC_Os10g19914 | hypothetical protein |
| LOC_Os10g19919 | retrotransposon protein, putative, Ty3-gypsy subclass, expressed |
| LOC_Os10g19925 | expressed protein |
| LOC_Os11g01480 | MYB family transcription factor, putative, expressed |
| LOC_Os11g01490 | outer membrane protein, OMP85 family, putative, expressed |
| LOC_Os11g01530 | ferritin-1, chloroplast precursor, putative, expressed |
| LOC_Os11g01540 | expressed protein |
| LOC_Os11g01570 | PMR5, putative, expressed |
| LOC_Os11g02630 | expressed protein |
| LOC_Os11g03200 | EMB2748, putative, expressed |
| LOC_Os11g03210 | glutathione S-transferase, N-terminal domain containing protein |
| LOC_Os11g03220 | RNA binding protein, putative, expressed |
| LOC_Os11g03230 | nucleoside-triphosphatase, putative, expressed |
| LOC_Os11g04670 | 2-isopropylmalate synthase B, putative, expressed |
| LOC_Os11g04680 | zinc finger, C3HC4 type domain containing protein, expressed |
| LOC_Os11g04690 | zinc finger, C3HC4 type domain containing protein, expressed |
| LOC_Os11g04710 | cytochrome P450, putative, expressed |

| | |
|---|---|
| LOC_Os11g33430 | plant protein of unknown function domain containing protein, expressed |
| LOC_Os11g39420 | jacalin-like lectin domain containing protein, expressed |
| LOC_Os11g39490 | jacalin-like lectin domain containing protein, expressed |
| LOC_Os12g07370 | csAtPR5, putative, expressed |
| LOC_Os12g07380 | expressed protein |
| LOC_Os12g07390 | expressed protein |
| LOC_Os12g13460 | SET domain-containing protein, putative, expressed |
| LOC_Os12g13470 | hypothetical protein |
| LOC_Os12g19530 | ATP-binding region, ATPase-like domain-containing protein, putative, expressed |
| LOC_Os12g20260 | expressed protein |
| LOC_Os12g22760 | expressed protein |
| LOC_Os12g24320 | ATPase 3, putative, expressed |
| LOC_Os12g34930 | expressed protein |
| LOC_Os12g39560 | expressed protein |

**Genes in japonica_SNP CLR**

| | |
|---|---|
| LOC_Os01g01090 | retrotransposon protein, putative, unclassified, expressed |
| LOC_Os01g01100 | retrotransposon protein, putative, Ty3-gypsy subclass, expressed |
| LOC_Os01g01530 | transposon protein, putative, CACTA, En/Spm sub-class |
| LOC_Os01g01540 | retrotransposon protein, putative, unclassified, expressed |
| LOC_Os01g01550 | transposon protein, putative, CACTA, En/Spm sub-class, expressed |
| LOC_Os01g01570 | kinesin heavy chain, putative, expressed |
| LOC_Os01g01810 | expressed protein |
| LOC_Os01g02310 | expressed protein |
| LOC_Os01g02380 | retrotransposon protein, putative, Ty3-gypsy subclass, expressed |
| LOC_Os01g02629 | retrotransposon protein, putative, unclassified, expressed |
| LOC_Os01g02990 | expressed protein |
| LOC_Os01g03610 | retrotransposon protein, putative, unclassified, expressed |
| LOC_Os01g05320 | retrotransposon protein, putative, Ty1-copia subclass, expressed |
| LOC_Os01g06610 | retrotransposon protein, putative, Ty3-gypsy subclass, expressed |

| | |
|---|---|
| LOC_Os01g06810 | retrotransposon protein, putative, Ty1-copia subclass, expressed |
| LOC_Os01g07570 | carboxyl-terminal proteinase, putative, expressed |
| LOC_Os01g07940 | AGC_PVPK_like_kin82y |
| LOC_Os01g08380 | transferase family protein, putative, expressed |
| LOC_Os01g08490 | conserved hypothetical protein |
| LOC_Os01g09060 | retrotransposon protein, putative, Ty1-copia subclass |
| LOC_Os01g09110 | retrotransposon protein, putative, unclassified |
| LOC_Os01g10030 | expressed protein |
| LOC_Os01g10130 | RNA-dependent RNA polymerase, putative, expressed |
| LOC_Os01g10180 | avr9/Cf-9 rapidly elicited protein 146, putative, expressed |
| LOC_Os01g10195 | expressed protein |
| LOC_Os01g10290 | autophagy-related protein 3, putative, expressed |
| LOC_Os01g10370 | AP2 domain containing protein, expressed |
| LOC_Os01g12150 | expressed protein |
| LOC_Os01g12340 | retrotransposon protein, putative, unclassified, expressed |
| LOC_Os01g14160 | retrotransposon, putative, centromere-specific, expressed |
| LOC_Os01g18020 | retrotransposon protein, putative, unclassified |
| LOC_Os01g18390 | OsRCI2-1 - Putative low temperature and salt responsive protein, expressed |
| LOC_Os01g18980 | transposon protein, putative, unclassified |
| LOC_Os01g19010 | retrotransposon protein, putative, unclassified, expressed |
| LOC_Os01g19529 | pentatricopeptide, putative, expressed |
| LOC_Os01g19894 | transposon protein, putative, unclassified, expressed |
| LOC_Os01g20140 | latency associated nuclear antigen, putative, expressed |
| LOC_Os01g22309 | hypothetical protein |
| LOC_Os01g23850 | transposon protein, putative, CACTA, En/Spm sub-class, expressed |
| LOC_Os01g23940 | expressed protein |
| LOC_Os01g27070 | expressed protein |
| LOC_Os01g27230 | 12-oxophytodienoate reductase, putative, expressed |
| LOC_Os01g27970 | hypothetical protein |
| LOC_Os01g28020 | retrotransposon protein, putative, Ty3-gypsy subclass |

| | |
|---|---|
| LOC_Os01g28270 | retrotransposon protein, putative, Ty1-copia subclass |
| LOC_Os01g28580 | expressed protein |
| LOC_Os01g28590 | expressed protein |
| LOC_Os01g28600 | exo70 exocyst complex subunit family protein, putative, expressed |
| LOC_Os01g28650 | retrotransposon, putative, centromere-specific, expressed |
| LOC_Os01g28680 | WD domain, G-beta repeat domain containing protein, expressed |
| LOC_Os01g29040 | retrotransposon protein, putative, unclassified, expressed |
| LOC_Os01g29409 | tRNA methyltransferase, putative, expressed |
| LOC_Os01g29430 | tetratricopeptide-like helical, putative, expressed |
| LOC_Os01g29450 | retrotransposon protein, putative, unclassified, expressed |
| LOC_Os01g31270 | annexin, putative, expressed |
| LOC_Os01g31280 | expressed protein |
| LOC_Os01g31820 | expressed protein |
| LOC_Os01g32500 | retrotransposon protein, putative, unclassified, expressed |
| LOC_Os01g32530 | transposon protein, putative, CACTA, En/Spm sub-class |
| LOC_Os01g35230 | 1-aminocyclopropane-1-carboxylate oxidase homolog 1, putative, expressed |
| LOC_Os01g35240 | hypothetical protein |
| LOC_Os01g35254 | transposon protein, putative, CACTA, En/Spm sub-class, expressed |
| LOC_Os01g36520 | expressed protein |
| LOC_Os01g36530 | transposon protein, putative, CACTA, En/Spm sub-class, expressed |
| LOC_Os01g36540 | pentatricopeptide, putative, expressed |
| LOC_Os01g36550 | protein kinase family protein, putative, expressed |
| LOC_Os01g36560 | auxin-induced protein 5NG4, putative, expressed |
| LOC_Os01g36570 | glycine-rich protein 2, putative |
| LOC_Os01g36600 | PPR repeat domain containing protein, putative, expressed |
| LOC_Os01g36610 | expressed protein |
| LOC_Os01g36620 | pectate lyase family protein, expressed |
| LOC_Os01g36640 | disease resistance protein RPM1, putative, expressed |
| LOC_Os01g36670 | expressed protein |
| LOC_Os01g36690 | expressed protein |

| | |
|---|---|
| LOC_Os01g36720 | transporter, major facilitator family, putative, expressed |
| LOC_Os01g36740 | expressed protein |
| LOC_Os01g36760 | expressed protein |
| LOC_Os01g36790 | TKL_IRAK_DUF26-lf |
| LOC_Os01g36810 | expressed protein |
| LOC_Os01g36820 | expressed protein |
| LOC_Os01g36830 | polygalacturonase, putative, expressed |
| LOC_Os01g36840 | ATP-binding region, ATPase-like domain containing protein, expressed |
| LOC_Os01g36860 | DEAD-box ATP-dependent RNA helicase 40, putative, expressed |
| LOC_Os01g36870 | phox domain-containing protein, putative, expressed |
| LOC_Os01g36880 | hypothetical protein |
| LOC_Os01g36890 | DEAD-box ATP-dependent RNA helicase, putative, expressed |
| LOC_Os01g36930 | ubiquitin carboxyl-terminal hydrolase 6, putative, expressed |
| LOC_Os01g36940 | OsFBX13 - F-box domain containing protein, expressed |
| LOC_Os01g36950 | N-rich protein, putative, expressed |
| LOC_Os01g36970 | expressed protein |
| LOC_Os01g36984 | expressed protein |
| LOC_Os01g37000 | carboxyl-terminal peptidase, putative, expressed |
| LOC_Os01g37100 | RWP-RK domain-containing protein, putative, expressed |
| LOC_Os01g37325 | retrotransposon protein, putative, unclassified, expressed |
| LOC_Os01g39030 | expressed protein |
| LOC_Os01g39136 | retrotransposon protein, putative, unclassified, expressed |
| LOC_Os01g39380 | SCD1, putative, expressed |
| LOC_Os01g39399 | expressed protein |
| LOC_Os01g39430 | anthocyanin regulatory protein, putative |
| LOC_Os01g39510 | hypothetical protein |
| LOC_Os01g39970 | protein kinase domain containing protein, putative, expressed |
| LOC_Os01g39980 | hypothetical protein |
| LOC_Os01g40120 | expressed protein |
| LOC_Os01g42810 | leaf senescence related protein, putative, expressed |

| | |
|---|---|
| LOC_Os01g43895 | expressed protein |
| LOC_Os01g44010 | retrotransposon protein, putative, unclassified, expressed |
| LOC_Os01g44069 | glycerol-3-phosphate acyltransferase, putative, expressed |
| LOC_Os01g46554 | transposon protein, putative, CACTA, En/Spm sub-class, expressed |
| LOC_Os01g46984 | expressed protein |
| LOC_Os01g48600 | ubiquitin carboxyl-terminal hydrolase family protein, putative, expressed |
| LOC_Os01g51400 | Leucine Rich Repeat family protein, expressed |
| LOC_Os01g51650 | retrotransposon protein, putative, unclassified, expressed |
| LOC_Os01g52910 | expressed protein |
| LOC_Os01g53450 | aminotransferase, classes I and II, domain containing protein, expressed |
| LOC_Os01g56120 | RNA recognition motif containing protein, putative, expressed |
| LOC_Os01g57942 | cytochrome b6, putative, expressed |
| LOC_Os01g57945 | cytochrome b6-f complex subunit 4, putative, expressed |
| LOC_Os01g57946 | DNA-directed RNA polymerase subunit alpha, putative, expressed |
| LOC_Os01g58049 | photosystem I assembly protein ycf4, putative, expressed |
| LOC_Os01g59250 | expressed protein |
| LOC_Os01g60870 | expressed protein |
| LOC_Os01g61350 | glutaredoxin, putative, expressed |
| LOC_Os01g61520 | expressed protein |
| LOC_Os01g61590 | CAMK_CAMK_like |
| LOC_Os01g61620 | protein kinase family protein, putative, expressed |
| LOC_Os01g61630 | expressed protein |
| LOC_Os01g61640 | NLI interacting factor-like phosphatase, putative, expressed |
| LOC_Os01g61670 | ureidoglycolate hydrolase, putative, expressed |
| LOC_Os01g61680 | expressed protein |
| LOC_Os01g61690 | OsSCP5 - Putative Serine Carboxypeptidase homologue, expressed |
| LOC_Os01g61700 | ATP-dependent Clp protease adaptor protein ClpS containing protein, expressed |
| LOC_Os01g61710 | coatomer subunit delta, putative, expressed |
| LOC_Os01g61720 | IQ calmodulin-binding motif domain containing protein, expressed |
| LOC_Os01g61730 | expressed protein |

| | |
|---|---|
| LOC_Os01g61750 | hypothetical protein |
| LOC_Os01g61780 | vacuolar ATP synthase 98 kDa subunit, putative, expressed |
| LOC_Os01g61800 | ZmEBE-1 protein, putative, expressed |
| LOC_Os01g61810 | histone-like transcription factor and archaeal histone, putative, expressed |
| LOC_Os01g61814 | 40S ribosomal protein S23, putative, expressed |
| LOC_Os01g61830 | zinc finger C-x8-C-x5-C-x3-H type family protein, expressed |
| LOC_Os01g62180 | expressed protein |
| LOC_Os01g63040 | expressed protein |
| LOC_Os01g63050 | expressed protein |
| LOC_Os01g63150 | retrotransposon protein, putative, Ty3-gypsy subclass, expressed |
| LOC_Os01g63160 | MYB family transcription factor, putative, expressed |
| LOC_Os01g63170 | transposon protein, putative, unclassified, expressed |
| LOC_Os01g63180 | laccase-6 precursor, putative, expressed |
| LOC_Os01g63190 | laccase precursor protein, putative, expressed |
| LOC_Os01g63200 | laccase precursor protein, putative, expressed |
| LOC_Os01g63210 | SOUL heme-binding protein, putative, expressed |
| LOC_Os01g63220 | kinase, pfkB family, putative, expressed |
| LOC_Os01g63230 | growth regulator related protein, putative, expressed |
| LOC_Os01g63240 | reticulon domain containing protein, putative, expressed |
| LOC_Os01g63610 | retrotransposon protein, putative, unclassified, expressed |
| LOC_Os01g63670 | hypothetical protein |
| LOC_Os01g63680 | MYB family transcription factor, putative, expressed |
| LOC_Os01g63690 | hs1, putative, expressed |
| LOC_Os01g63710 | CDC6 - Putative DNA replication initiation protein, expressed |
| LOC_Os01g63720 | hypothetical protein |
| LOC_Os01g63730 | hypothetical protein |
| LOC_Os01g63770 | transmembrane amino acid transporter protein, putative, expressed |
| LOC_Os01g63780 | nsp1-like C-terminal region family protein, expressed |
| LOC_Os01g63790 | prolipoprotein diacylglyceryl transferase, putative, expressed |
| LOC_Os01g63800 | pleckstrin homology domain-containing protein, putative, expressed |

| | |
|---|---|
| LOC_Os01g63810 | starch binding domain containing protein, putative, expressed |
| LOC_Os01g63820 | double-stranded RNA binding motif containing protein, expressed |
| LOC_Os01g63830 | dehydrodolichyl diphosphate synthase, putative, expressed |
| LOC_Os01g63900 | WD domain, G-beta repeat domain containing protein, expressed |
| LOC_Os01g63910 | AT hook motif family protein, expressed |
| LOC_Os01g63920 | hypothetical protein |
| LOC_Os01g63930 | cytochrome P450, putative, expressed |
| LOC_Os01g67790 | F-box protein interaction domain containing protein, expressed |
| LOC_Os01g71250 | expressed protein |
| LOC_Os01g71870 | transposon protein, putative, unclassified, expressed |
| LOC_Os02g02400 | catalase isozyme A, putative, expressed |
| LOC_Os02g03870 | periplasmic beta-glucosidase precursor, putative, expressed |
| LOC_Os02g04870 | retrotransposon protein, putative, Ty1-copia subclass, expressed |
| LOC_Os02g05040 | cyclin-related protein, putative, expressed |
| LOC_Os02g05210 | retrotransposon protein, putative, unclassified, expressed |
| LOC_Os02g05640 | homeobox associated leucine zipper, putative, expressed |
| LOC_Os02g12300 | pectate lyase precursor, putative, expressed |
| LOC_Os02g14600 | glutelin, putative, expressed |
| LOC_Os02g15070 | glutelin, putative, expressed |
| LOC_Os02g16940 | OsSub13 - Putative Subtilisin homologue, expressed |
| LOC_Os02g16995 | expressed protein |
| LOC_Os02g17000 | OsSub14 - Putative Subtilisin homologue, expressed |
| LOC_Os02g17020 | hypothetical protein |
| LOC_Os02g17170 | expressed protein |
| LOC_Os02g17180 | OsFBX42 - F-box domain containing protein, expressed |
| LOC_Os02g18064 | expressed protein |
| LOC_Os02g18150 | expressed protein |
| LOC_Os02g18380 | 60S ribosomal protein L27-3, putative, expressed |
| LOC_Os02g18390 | expressed protein |
| LOC_Os02g18400 | hypothetical protein |

| | |
|---|---|
| LOC_Os02g18410 | salt stress root protein RS1, putative, expressed |
| LOC_Os02g18420 | expressed protein |
| LOC_Os02g29180 | expressed protein |
| LOC_Os02g30760 | expressed protein |
| LOC_Os02g31280 | expressed protein |
| LOC_Os02g36414 | transporter family protein, putative, expressed |
| LOC_Os02g36720 | expressed protein |
| LOC_Os02g36910 | transposon protein, putative, unclassified |
| LOC_Os02g37109 | vegetative cell wall protein gp1 precursor, putative, expressed |
| LOC_Os02g40330 | retrotransposon protein, putative, Ty3-gypsy subclass, expressed |
| LOC_Os02g40710 | ammonium transporter protein, putative, expressed |
| LOC_Os02g42560 | expressed protein |
| LOC_Os02g43290 | protein kinase domain containing protein, expressed |
| LOC_Os02g43460 | required to maintain repression 1, putative, expressed |
| LOC_Os02g43800 | retrotransposon protein, putative, Ty3-gypsy subclass, expressed |
| LOC_Os02g46560 | helix-loop-helix DNA-binding protein, putative, expressed |
| LOC_Os02g47280 | growth-regulating factor, putative, expressed |
| LOC_Os02g53750 | tyrosine protein kinase domain containing protein, putative, expressed |
| LOC_Os02g56010 | anthocyanidin 3-O-glucosyltransferase, putative, expressed |
| LOC_Os02g57620 | Citrate transporter protein, putative, expressed |
| LOC_Os03g02010 | DNA methyltransferase protein, putative, expressed |
| LOC_Os03g02030 | folylpolyglutamate synthase, mitochondrial precursor, putative, expressed |
| LOC_Os03g02040 | remorin, putative, expressed |
| LOC_Os03g02050 | LTPL151 - Protease inhibitor/seed storage/LTP family protein precursor, expressed |
| LOC_Os03g02060 | hypothetical protein |
| LOC_Os03g02070 | heavy-metal-associated domain-containing protein, putative, expressed |
| LOC_Os03g02080 | expressed protein |
| LOC_Os03g02090 | expressed protein |
| LOC_Os03g02110 | WD domain, G-beta repeat domain containing protein, expressed |
| LOC_Os03g02150 | adapitin protein, putative, expressed |

| | |
|---|---|
| LOC_Os03g02160 | zinc finger C-x8-C-x5-C-x3-H type family protein, expressed |
| LOC_Os03g02170 | transmembrane and coiled-coil domain-containing protein 4, putative, expressed |
| LOC_Os03g02180 | cytochrome P450, putative, expressed |
| LOC_Os03g02200 | thymidine kinase, putative, expressed |
| LOC_Os03g02980 | CAMK_CAMK_like_ULKh_APGy |
| LOC_Os03g03010 | expressed protein |
| LOC_Os03g03890 | protein kinase family protein, putative, expressed |
| LOC_Os03g06960 | vesicle-associated membrane protein, putative, expressed |
| LOC_Os03g15250 | lectin-like receptor kinase, putative, expressed |
| LOC_Os03g15550 | phosphatidylinositol-4-phosphate 5-kinase, putative, expressed |
| LOC_Os03g17490 | expressed protein |
| LOC_Os03g17500 | expressed protein |
| LOC_Os03g17510 | pentatricopeptide, putative, expressed |
| LOC_Os03g17520 | CDP-alcohol phosphatidyltransferase, putative, expressed |
| LOC_Os03g17530 | expressed protein |
| LOC_Os03g17540 | TBC domain containing protein, expressed |
| LOC_Os03g18779 | expressed protein |
| LOC_Os03g19436 | expressed protein |
| LOC_Os03g19660 | conserved hypothetical protein |
| LOC_Os03g24430 | cytokinin-O-glucosyltransferase 3, putative, expressed |
| LOC_Os03g26329 | retrotransposon protein, putative, unclassified, expressed |
| LOC_Os03g27590 | OsSCP18 - Putative Serine Carboxypeptidase homologue, expressed |
| LOC_Os03g38410 | expressed protein |
| LOC_Os03g41110 | ZOS3-14 - C2H2 zinc finger protein, expressed |
| LOC_Os03g45710 | 2Fe-2S iron-sulfur cluster binding domain containing protein, expressed |
| LOC_Os03g47149 | expressed protein |
| LOC_Os03g51980 | expressed protein |
| LOC_Os03g51984 | differentiation protein, putative, expressed |
| LOC_Os03g51990 | ACT domain containing protein, expressed |
| LOC_Os03g52000 | BSD domain containing protein, expressed |

| | |
|---|---|
| LOC_Os03g52794 | phosphatidylinositol kinase, putative, expressed |
| LOC_Os03g53010 | acyl-desaturase, chloroplast precursor, putative, expressed |
| LOC_Os03g53020 | helix-loop-helix DNA-binding domain containing protein, expressed |
| LOC_Os03g53030 | hypothetical protein |
| LOC_Os03g53080 | zinc finger, C3HC4 type domain containing protein, expressed |
| LOC_Os03g53090 | LRR-kinase protein, putative, expressed |
| LOC_Os03g53100 | response regulator receiver domain containing protein, expressed |
| LOC_Os03g53110 | CorA-like magnesium transporter protein, putative, expressed |
| LOC_Os03g53140 | protein transport protein Sec23B, putative, expressed |
| LOC_Os03g56820 | fatty acid hydroxylase, putative, expressed |
| LOC_Os03g57569 | expressed protein |
| LOC_Os03g58830 | BEE 1, putative, expressed |
| LOC_Os03g58920 | galactosyltransferase family protein, putative, expressed |
| LOC_Os03g59050 | DEAD-box ATP-dependent RNA helicase, putative, expressed |
| LOC_Os03g59210 | expressed protein |
| LOC_Os03g59220 | cytochrome b-c1 complex subunit 7, putative, expressed |
| LOC_Os03g59225 | expressed protein |
| LOC_Os03g59300 | expressed protein |
| LOC_Os03g59310 | ribosomal protein, putative, expressed |
| LOC_Os03g59330 | polygalacturonase, putative, expressed |
| LOC_Os03g62560 | expressed protein |
| LOC_Os03g63300 | NLI interacting factor-like phosphatase, putative, expressed |
| LOC_Os04g04340 | retrotransposon protein, putative, unclassified, expressed |
| LOC_Os04g07630 | hypothetical protein |
| LOC_Os04g10150 | expressed protein |
| LOC_Os04g11330 | retrotransposon protein, putative, unclassified, expressed |
| LOC_Os04g16848 | photosystem II reaction center protein H, putative |
| LOC_Os04g27320 | transposon protein, putative, unclassified |
| LOC_Os04g28520 | male sterility protein, putative, expressed |
| LOC_Os04g28530 | expressed protein |

| | |
|---|---|
| LOC_Os04g28540 | expressed protein |
| LOC_Os04g28980 | expressed protein |
| LOC_Os04g32650 | tRNA synthetase class II core domain containing protein, expressed |
| LOC_Os04g32660 | cytochrome b-c1 complex subunit Rieske, mitochondrial precursor, putative, expressed |
| LOC_Os04g32670 | exostosin family domain containing protein, expressed |
| LOC_Os04g40180 | expressed protein |
| LOC_Os04g40600 | peptide methionine sulfoxide reductase, putative, expressed |
| LOC_Os04g42889 | expressed protein |
| LOC_Os04g42920 | dehydrogenase, putative, expressed |
| LOC_Os04g42930 | OsGrx_C2 |
| LOC_Os04g42940 | expressed protein |
| LOC_Os04g43080 | hypothetical protein |
| LOC_Os04g46910 | actin-depolymerizing factor, putative, expressed |
| LOC_Os04g51590 | retrotransposon protein, putative, unclassified, expressed |
| LOC_Os04g55350 | retrotransposon protein, putative, Ty3-gypsy subclass, expressed |
| LOC_Os04g55960 | NADPH reductase, putative, expressed |
| LOC_Os05g11790 | CAMK_KIN1/SNF1/Nim1_like |
| LOC_Os05g16360 | expressed protein |
| LOC_Os05g19420 | retrotransposon protein, putative, Ty3-gypsy subclass |
| LOC_Os05g23250 | Leucine Rich Repeat family protein, expressed |
| LOC_Os05g23260 | OsSigP4 - Putative Type I Signal Peptidase homologue; employs a putative Ser/His catalytic dyad, expressed |
| LOC_Os05g28490 | FAD-binding domain-containing protein, putative, expressed |
| LOC_Os05g49530 | F-box domain containing protein, expressed |
| LOC_Os06g05240 | zinc carboxypeptidase family protein, putative, expressed |
| LOC_Os06g15850 | transposon protein, putative, CACTA, En/Spm sub-class, expressed |
| LOC_Os06g37224 | cytochrome P450, putative, expressed |
| LOC_Os06g37310 | retrotransposon protein, putative, unclassified, expressed |
| LOC_Os06g39708 | photosystem II P680 chlorophyll A apoprotein, putative, expressed |
| LOC_Os06g39712 | OsClp10 - Putative Clp protease homologue, expressed |
| LOC_Os06g39716 | chloroplast 30S ribosomal protein S18, putative, expressed |

| | |
|---|---|
| LOC_Os06g44030 | WD domain, G-beta repeat domain containing protein, expressed |
| LOC_Os06g49420 | retrotransposon protein, putative, unclassified, expressed |
| LOC_Os07g01690 | expressed protein |
| LOC_Os07g03900 | lectin-like receptor kinase, putative, expressed |
| LOC_Os07g03980 | expressed protein |
| LOC_Os07g03990 | leucine-rich repeat family protein, putative, expressed |
| LOC_Os07g06620 | YABBY domain containing protein, putative, expressed |
| LOC_Os07g09570 | hypothetical protein |
| LOC_Os07g09580 | aminoacyl-tRNA synthetase, putative, expressed |
| LOC_Os07g09610 | CAMK_KIN1/SNF1/Nim1_like_AMPKh |
| LOC_Os07g09700 | expressed protein |
| LOC_Os07g09710 | OsFBX220 - F-box domain containing protein, expressed |
| LOC_Os07g09950 | OsFBX222 - F-box domain containing protein, expressed |
| LOC_Os07g10160 | glucosyltransferase, putative, expressed |
| LOC_Os07g10254 | expressed protein |
| LOC_Os07g10256 | mitosis protein dim1, putative, expressed |
| LOC_Os07g10310 | expressed protein |
| LOC_Os07g10320 | expressed protein |
| LOC_Os07g10340 | expressed protein |
| LOC_Os07g10350 | S1 RNA binding domain containing protein, expressed |
| LOC_Os07g10390 | cleavage and polyadenylation specificity factor, putative, expressed |
| LOC_Os07g10400 | pentatricopeptide, putative, expressed |
| LOC_Os07g10410 | AP2 domain containing protein, expressed |
| LOC_Os07g10420 | expressed protein |
| LOC_Os07g10440 | expressed protein |
| LOC_Os07g10460 | 5-nucleotidase surE, putative, expressed |
| LOC_Os07g10490 | zeta-carotene desaturase, chloroplast/chromoplast precursor, putative, expressed |
| LOC_Os07g10495 | expressed protein |
| LOC_Os07g10510 | expressed protein |
| LOC_Os07g10535 | expressed protein |

| | |
|---|---|
| LOC_Os07g10540 | peptidase, M24 family protein, putative, expressed |
| LOC_Os07g10560 | expressed protein |
| LOC_Os07g10650 | hypothetical protein |
| LOC_Os07g10680 | polygalacturonase, putative, expressed |
| LOC_Os07g10720 | 40S ribosomal protein S15a, putative, expressed |
| LOC_Os07g10780 | remorin, putative, expressed |
| LOC_Os07g11020 | rc - bHLH transcription factor regulating proanthocyanidin production in seeds, expressed |
| LOC_Os07g17460 | OsFBL36 - F-box domain and LRR containing protein, expressed |
| LOC_Os07g17560 | expressed protein |
| LOC_Os07g28090 | ABC transporter, ATP-binding protein, putative, expressed |
| LOC_Os07g28110 | cytochrome P450, putative, expressed |
| LOC_Os07g28150 | hAT dimerisation domain containing protein, putative, expressed |
| LOC_Os07g29760 | cysteine proteinase A494 precursor, putative, expressed |
| LOC_Os07g35050 | OsFBX237 - F-box domain containing protein, expressed |
| LOC_Os08g03410 | glutelin, putative, expressed |
| LOC_Os08g15306 | chloroplast 50S ribosomal protein L20, putative, expressed |
| LOC_Os08g18079 | expressed protein |
| LOC_Os08g32570 | expressed protein |
| LOC_Os08g32600 | STE_MEKK_ste11_MAP3K |
| LOC_Os08g42910 | peptidase, M24 family protein, putative, expressed |
| LOC_Os09g17329 | expressed protein |
| LOC_Os09g38470 | expressed protein |
| LOC_Os09g38580 | cyclic nucleotide-gated ion channel, putative, expressed |
| LOC_Os09g38660 | expressed protein |
| LOC_Os09g38670 | thioredoxin, putative, expressed |
| LOC_Os09g38680 | expressed protein |
| LOC_Os09g38690 | transporter family protein, putative, expressed |
| LOC_Os09g38700 | STRUBBELIG-RECEPTOR FAMILY 5 precursor, putative, expressed |
| LOC_Os09g39020 | N-rich protein, putative, expressed |
| LOC_Os09g39050 | OsFBX342 - F-box domain containing protein, expressed |

| | |
|---|---|
| LOC_Os09g39060 | vignain precursor, putative, expressed |
| LOC_Os10g27460 | expressed protein |
| LOC_Os10g39489 | retrotransposon protein, putative, unclassified, expressed |
| LOC_Os10g39500 | expressed protein |
| LOC_Os10g39970 | harpin-induced protein 1 domain containing protein, expressed |
| LOC_Os10g40000 | retinol dehydrogenase 12, putative, expressed |
| LOC_Os10g41030 | agenet domain containing protein, putative, expressed |
| LOC_Os10g41040 | ternary complex factor MIP1, putative, expressed |
| LOC_Os10g41120 | expressed protein |
| LOC_Os10g41160 | expressed protein |
| LOC_Os10g41170 | dehydrogenase, putative, expressed |
| LOC_Os10g41370 | WD repeat-containing protein 8, putative, expressed |
| LOC_Os10g41380 | expressed protein |
| LOC_Os10g41390 | protein kinase domain containing protein, expressed |
| LOC_Os10g41410 | nucleoside diphosphate kinase, putative, expressed |
| LOC_Os10g41510 | calcineurin B, putative, expressed |
| LOC_Os10g41520 | casein kinase II subunit beta-4, putative, expressed |
| LOC_Os10g41534 | ICE-like protease p20 domain containing protein, expressed |
| LOC_Os11g04130 | nodulin MtN21 family protein, putative, expressed |
| LOC_Os11g04950 | splicing factor, putative, expressed |
| LOC_Os11g04954 | DNA repair protein Rad51, putative, expressed |
| LOC_Os11g04980 | acyl-CoA synthetase protein, putative, expressed |
| LOC_Os12g04970 | retrotransposon protein, putative, unclassified, expressed |
| LOC_Os12g10930 | NL0E, putative, expressed |
| LOC_Os12g11040 | expressed protein |
| LOC_Os12g11060 | hypothetical protein |
| LOC_Os12g11500 | resistance protein SlVe1 precursor, putative, expressed |
| LOC_Os12g11610 | expressed protein |
| LOC_Os12g11660 | RALFL45 - Rapid ALkalinization Factor RALF family protein precursor, expressed |
| LOC_Os12g11690 | expressed protein |

| | |
|---|---|
| LOC_Os12g11790 | expressed protein |
| LOC_Os12g11840 | hypothetical protein |
| LOC_Os12g31290 | expressed protein |
| LOC_Os12g37110 | OsFBD16 - F-box and FBD domain containing protein, expressed |
| LOC_Os12g43034 | retrotransposon protein, putative, unclassified, expressed |
| LOC_Os12g43310 | hypothetical protein |
| LOC_Os12g43330 | expressed protein |
| LOC_Os12g43380 | thaumatin, putative, expressed |
| LOC_Os12g43400 | expressed protein |
| LOC_Os12g43510 | expressed protein |

**Genes in japonica_SV CLR**

| | |
|---|---|
| LOC_Os01g01090 | retrotransposon protein, putative, unclassified, expressed |
| LOC_Os01g01100 | retrotransposon protein, putative, Ty3-gypsy subclass, expressed |
| LOC_Os01g01530 | transposon protein, putative, CACTA, En/Spm sub-class |
| LOC_Os01g01540 | retrotransposon protein, putative, unclassified, expressed |
| LOC_Os01g01550 | transposon protein, putative, CACTA, En/Spm sub-class, expressed |
| LOC_Os01g02629 | retrotransposon protein, putative, unclassified, expressed |
| LOC_Os01g03610 | retrotransposon protein, putative, unclassified, expressed |
| LOC_Os01g03720 | MYB family transcription factor, putative, expressed |
| LOC_Os01g04510 | expressed protein |
| LOC_Os01g04790 | retrotransposon protein, putative, unclassified |
| LOC_Os01g05320 | retrotransposon protein, putative, Ty1-copia subclass, expressed |
| LOC_Os01g07550 | hypothetical protein |
| LOC_Os01g09060 | retrotransposon protein, putative, Ty1-copia subclass |
| LOC_Os01g09110 | retrotransposon protein, putative, unclassified |
| LOC_Os01g11270 | cytochrome P450, putative, expressed |
| LOC_Os01g12440 | AP2 domain containing protein, expressed |
| LOC_Os01g12610 | expressed protein |
| LOC_Os01g13620 | retrotransposon protein, putative, unclassified, expressed |

| | |
|---|---|
| LOC_Os01g14720 | transcription regulator, putative, expressed |
| LOC_Os01g14730 | retrotransposon protein, putative, unclassified |
| LOC_Os01g15150 | expressed protein |
| LOC_Os01g17130 | retrotransposon protein, putative, Ty3-gypsy subclass, expressed |
| LOC_Os01g18020 | retrotransposon protein, putative, unclassified |
| LOC_Os01g19010 | retrotransposon protein, putative, unclassified, expressed |
| LOC_Os01g19710 | transposon protein, putative, CACTA, En/Spm sub-class, expressed |
| LOC_Os01g20060 | expressed protein |
| LOC_Os01g21210 | retrotransposon protein, putative, unclassified, expressed |
| LOC_Os01g21360 | retrotransposon protein, putative, unclassified |
| LOC_Os01g21550 | expressed protein |
| LOC_Os01g22050 | retrotransposon protein, putative, unclassified, expressed |
| LOC_Os01g22080 | expressed protein |
| LOC_Os01g22830 | retrotransposon protein, putative, Ty3-gypsy subclass, expressed |
| LOC_Os01g22840 | retrotransposon protein, putative, Ty3-gypsy subclass, expressed |
| LOC_Os01g23180 | retrotransposon protein, putative, unclassified |
| LOC_Os01g23239 | retrotransposon protein, putative, Ty3-gypsy subclass, expressed |
| LOC_Os01g23340 | expressed protein |
| LOC_Os01g23480 | expressed protein |
| LOC_Os01g24860 | retrotransposon protein, putative, unclassified, expressed |
| LOC_Os01g25050 | expressed protein |
| LOC_Os01g27070 | expressed protein |
| LOC_Os01g27320 | retrotransposon protein, putative, Ty3-gypsy subclass, expressed |
| LOC_Os01g27410 | hypothetical protein |
| LOC_Os01g27449 | expressed protein |
| LOC_Os01g27970 | hypothetical protein |
| LOC_Os01g28020 | retrotransposon protein, putative, Ty3-gypsy subclass |
| LOC_Os01g28270 | retrotransposon protein, putative, Ty1-copia subclass |
| LOC_Os01g28660 | expressed protein |
| LOC_Os01g28850 | expressed protein |

| | |
|---|---|
| LOC_Os01g28970 | retrotransposon protein, putative, Ty3-gypsy subclass, expressed |
| LOC_Os01g28980 | proton-dependent oligopeptide transport, putative, expressed |
| LOC_Os01g28989 | expressed protein |
| LOC_Os01g29030 | retrotransposon protein, putative, unclassified, expressed |
| LOC_Os01g29040 | retrotransposon protein, putative, unclassified, expressed |
| LOC_Os01g29104 | expressed protein |
| LOC_Os01g29200 | transposon protein, putative, unclassified |
| LOC_Os01g29730 | retrotransposon protein, putative, Ty3-gypsy subclass |
| LOC_Os01g30970 | PMR5, putative, expressed |
| LOC_Os01g31440 | retrotransposon protein, putative, unclassified, expressed |
| LOC_Os01g31450 | retrotransposon protein, putative, unclassified, expressed |
| LOC_Os01g31460 | retrotransposon protein, putative, unclassified, expressed |
| LOC_Os01g32530 | transposon protein, putative, CACTA, En/Spm sub-class |
| LOC_Os01g32550 | hypothetical protein |
| LOC_Os01g33142 | hypothetical protein |
| LOC_Os01g33580 | retrotransposon, putative, centromere-specific |
| LOC_Os01g34510 | hypothetical protein |
| LOC_Os01g34560 | 3-ketoacyl-CoA synthase, putative, expressed |
| LOC_Os01g34910 | expressed protein |
| LOC_Os01g35610 | expressed protein |
| LOC_Os01g35810 | expressed protein |
| LOC_Os01g36120 | expressed protein |
| LOC_Os01g36410 | expressed protein |
| LOC_Os01g36720 | transporter, major facilitator family, putative, expressed |
| LOC_Os01g36740 | expressed protein |
| LOC_Os01g36760 | expressed protein |
| LOC_Os01g36790 | TKL_IRAK_DUF26-lf |
| LOC_Os01g39136 | retrotransposon protein, putative, unclassified, expressed |
| LOC_Os01g39350 | protein kinase domain containing protein, expressed |
| LOC_Os01g39490 | expressed protein |

| | |
|---|---|
| LOC_Os01g39510 | hypothetical protein |
| LOC_Os01g39520 | retrotransposon, putative, centromere-specific, expressed |
| LOC_Os01g39530 | retrotransposon, putative, centromere-specific |
| LOC_Os01g39560 | anthocyanin regulatory protein, putative, expressed |
| LOC_Os01g39570 | retrotransposon protein, putative, Ty3-gypsy subclass, expressed |
| LOC_Os01g39980 | hypothetical protein |
| LOC_Os01g40120 | expressed protein |
| LOC_Os01g44010 | retrotransposon protein, putative, unclassified, expressed |
| LOC_Os01g46169 | GDSL-like lipase/acylhydrolase, putative, expressed |
| LOC_Os01g48650 | expressed protein |
| LOC_Os01g48830 | expressed protein |
| LOC_Os01g52910 | expressed protein |
| LOC_Os01g53200 | flavin monooxygenase, putative, expressed |
| LOC_Os01g53210 | metal transporter Nramp3, putative, expressed |
| LOC_Os01g58024 | NADPH-dependent oxidoreductase, putative, expressed |
| LOC_Os01g58049 | photosystem I assembly protein ycf4, putative, expressed |
| LOC_Os01g58950 | cytochrome P450, putative, expressed |
| LOC_Os01g58970 | cytochrome P450, putative, expressed |
| LOC_Os01g59360 | CAMK_CAMK_like |
| LOC_Os01g60560 | retrotransposon protein, putative, unclassified, expressed |
| LOC_Os01g61300 | expressed protein |
| LOC_Os01g61460 | leaf senescence related protein, putative, expressed |
| LOC_Os01g62520 | retrotransposon protein, putative, unclassified |
| LOC_Os01g63040 | expressed protein |
| LOC_Os01g64510 | retrotransposon protein, putative, unclassified, expressed |
| LOC_Os01g65000 | ammonium transporter protein, putative, expressed |
| LOC_Os01g65130 | peptide transporter, putative, expressed |
| LOC_Os01g71850 | transposon protein, putative, unclassified, expressed |
| LOC_Os01g71940 | retrotransposon protein, putative, unclassified |
| LOC_Os02g03750 | polygalacturonase, putative, expressed |

| | |
|---|---|
| LOC_Os02g04360 | expressed protein |
| LOC_Os02g06310 | expressed protein |
| LOC_Os02g15360 | expressed protein |
| LOC_Os02g17760 | cytochrome P450, putative, expressed |
| LOC_Os02g18115 | expressed protein |
| LOC_Os02g19540 | OsFBX47 - F-box domain containing protein, expressed |
| LOC_Os02g20520 | retrotransposon, putative, centromere-specific |
| LOC_Os02g24190 | cyclin-T1-2, putative, expressed |
| LOC_Os02g24560 | retrotransposon protein, putative, unclassified, expressed |
| LOC_Os02g24576 | chloroplast 30S ribosomal protein S7, putative, expressed |
| LOC_Os02g24586 | uncharacterized protein ycf68, putative, expressed |
| LOC_Os02g24830 | expressed protein |
| LOC_Os02g25755 | expressed protein |
| LOC_Os02g25780 | expressed protein |
| LOC_Os02g25800 | retrotransposon protein, putative, unclassified, expressed |
| LOC_Os02g25960 | ankyrin repeat family protein, putative, expressed |
| LOC_Os02g25990 | retrotransposon, putative, centromere-specific |
| LOC_Os02g26349 | transposon protein, putative, CACTA, En/Spm sub-class, expressed |
| LOC_Os02g27210 | hypothetical protein |
| LOC_Os02g27370 | expressed protein |
| LOC_Os02g28170 | transferase family protein, putative, expressed |
| LOC_Os02g28465 | expressed protein |
| LOC_Os02g29570 | retrotransposon protein, putative, Ty3-gypsy subclass, expressed |
| LOC_Os02g29700 | retrotransposon, putative, centromere-specific, expressed |
| LOC_Os02g30760 | expressed protein |
| LOC_Os02g32910 | retrotransposon protein, putative, unclassified, expressed |
| LOC_Os02g39530 | expressed protein |
| LOC_Os02g40330 | retrotransposon protein, putative, Ty3-gypsy subclass, expressed |
| LOC_Os02g43800 | retrotransposon protein, putative, Ty3-gypsy subclass, expressed |
| LOC_Os02g47680 | retrotransposon protein, putative, Ty3-gypsy subclass |

| | |
|---|---|
| LOC_Os02g49170 | retrotransposon protein, putative, unclassified, expressed |
| LOC_Os02g50120 | retrotransposon protein, putative, Ty3-gypsy subclass |
| LOC_Os02g55770 | expressed protein |
| LOC_Os03g13920 | transposon protein, putative, CACTA, En/Spm sub-class, expressed |
| LOC_Os03g18070 | omega-3 fatty acid desaturase, chloroplast precursor, putative, expressed |
| LOC_Os03g19660 | conserved hypothetical protein |
| LOC_Os03g21170 | expressed protein |
| LOC_Os03g21914 | ATP binding protein, putative, expressed |
| LOC_Os03g22190 | expressed protein |
| LOC_Os03g25450 | h/ACA ribonucleoprotein complex subunit 4, putative, expressed |
| LOC_Os03g25640 | OsFBX91 - F-box domain containing protein, expressed |
| LOC_Os03g25700 | retrotransposon protein, putative, Ty3-gypsy subclass, expressed |
| LOC_Os03g26044 | CSLA5 - cellulose synthase-like family A; mannan synthase, expressed |
| LOC_Os03g26280 | expressed protein |
| LOC_Os03g26329 | retrotransposon protein, putative, unclassified, expressed |
| LOC_Os03g34070 | retrotransposon protein, putative, unclassified, expressed |
| LOC_Os03g37430 | retrotransposon protein, putative, unclassified, expressed |
| LOC_Os03g37760 | expressed protein |
| LOC_Os03g37770 | expressed protein |
| LOC_Os03g37780 | expressed protein |
| LOC_Os03g37810 | expressed protein |
| LOC_Os03g41780 | retrotransposon, putative, centromere-specific, expressed |
| LOC_Os03g41800 | transposon protein, putative, unclassified, expressed |
| LOC_Os03g41810 | hypothetical protein |
| LOC_Os03g41830 | retrotransposon protein, putative, unclassified, expressed |
| LOC_Os03g44370 | transposon protein, putative, CACTA, En/Spm sub-class, expressed |
| LOC_Os03g60710 | protein kinase domain containing protein, expressed |
| LOC_Os04g04600 | retrotransposon protein, putative, Ty3-gypsy subclass |
| LOC_Os04g12370 | expressed protein |
| LOC_Os04g14890 | retrotransposon protein, putative, Ty3-gypsy subclass, expressed |

| | |
|---|---|
| LOC_Os04g15760 | DEFL51 - Defensin and Defensin-like DEFL family, expressed |
| LOC_Os04g16732 | NADPH-dependent oxidoreductase, putative, expressed |
| LOC_Os04g16742 | NADPH-dependent oxidoreductase, putative, expressed |
| LOC_Os04g16792 | uncharacterized protein ycf76, putative |
| LOC_Os04g16794 | hypothetical protein |
| LOC_Os04g19740 | transketolase, chloroplast precursor, putative, expressed |
| LOC_Os04g19750 | OsFBL11 - F-box domain and LRR containing protein, expressed |
| LOC_Os04g19810 | OsFBL12 - F-box domain and LRR containing protein, expressed |
| LOC_Os04g21780 | expressed protein |
| LOC_Os04g22110 | retrotransposon protein, putative, unclassified, expressed |
| LOC_Os04g22290 | expressed protein |
| LOC_Os04g24540 | expressed protein |
| LOC_Os04g28520 | male sterility protein, putative, expressed |
| LOC_Os04g28530 | expressed protein |
| LOC_Os04g28540 | expressed protein |
| LOC_Os04g41720 | retrotransposon, putative, centromere-specific, expressed |
| LOC_Os04g48710 | expressed protein |
| LOC_Os05g04630 | retrotransposon protein, putative, SINE subclass, expressed |
| LOC_Os05g08170 | retrotransposon protein, putative, Ty3-gypsy subclass, expressed |
| LOC_Os05g20710 | retrotransposon protein, putative, Ty3-gypsy subclass, expressed |
| LOC_Os05g30540 | retrotransposon protein, putative, unclassified |
| LOC_Os05g32020 | expressed protein |
| LOC_Os05g32090 | expressed protein |
| LOC_Os05g39630 | expressed protein |
| LOC_Os06g11670 | expressed protein |
| LOC_Os06g20140 | aspartic proteinase nepenthesin-1 precursor, putative, expressed |
| LOC_Os06g20200 | gibberellin receptor GID1L2, putative, expressed |
| LOC_Os06g23770 | expressed protein |
| LOC_Os06g26080 | expressed protein |
| LOC_Os06g26120 | hypothetical protein |

| | |
|---|---|
| LOC_Os06g31280 | THION1 - Plant thionin family protein precursor, putative, expressed |
| LOC_Os06g31800 | THION2 - Plant thionin family protein precursor, expressed |
| LOC_Os06g31810 | hypothetical protein |
| LOC_Os06g32290 | THION10 - Plant thionin family protein precursor |
| LOC_Os06g32320 | THION11 - Plant thionin family protein precursor |
| LOC_Os06g32480 | expressed protein |
| LOC_Os06g32500 | retrotransposon protein, putative, Ty3-gypsy subclass, expressed |
| LOC_Os06g32550 | THION14 - Plant thionin family protein precursor, putative, expressed |
| LOC_Os06g32650 | expressed protein |
| LOC_Os06g32710 | expressed protein |
| LOC_Os06g32720 | zinc finger C-x8-C-x5-C-x3-H type family protein, expressed |
| LOC_Os06g32730 | eukaryotic initiation factor iso-4F subunit p82-34, putative, expressed |
| LOC_Os06g40415 | expressed protein |
| LOC_Os06g46438 | NADPH-dependent oxidoreductase, putative |
| LOC_Os07g02030 | thiamine pyrophosphate enzyme, N-terminal TPP binding domain containing protein, expressed |
| LOC_Os07g05930 | expressed protein |
| LOC_Os07g08040 | expressed protein |
| LOC_Os07g15610 | hypothetical protein |
| LOC_Os07g15770 | CCT motif family protein, expressed |
| LOC_Os07g15820 | expressed protein |
| LOC_Os07g17390 | PsbP, putative, expressed |
| LOC_Os07g17460 | OsFBL36 - F-box domain and LRR containing protein, expressed |
| LOC_Os07g17550 | hypothetical protein |
| LOC_Os07g17560 | expressed protein |
| LOC_Os07g17770 | tryptophanyl-tRNA synthetase, cytoplasmic, putative, expressed |
| LOC_Os07g20530 | hypothetical protein |
| LOC_Os07g20880 | retrotransposon protein, putative, unclassified, expressed |
| LOC_Os07g23485 | expressed protein |
| LOC_Os07g23900 | OsFBX234 - F-box domain containing protein, expressed |
| LOC_Os07g27250 | expressed protein |

| | |
|---|---|
| LOC_Os08g11200 | retrotransposon protein, putative, unclassified, expressed |
| LOC_Os08g11220 | expressed protein |
| LOC_Os08g11599 | expressed protein |
| LOC_Os08g12160 | expressed protein |
| LOC_Os08g15530 | expressed protein |
| LOC_Os08g19550 | expressed protein |
| LOC_Os08g22354 | polyadenylate-binding protein, putative, expressed |
| LOC_Os08g22389 | hypothetical protein |
| LOC_Os08g26210 | expressed protein |
| LOC_Os09g06310 | hypothetical protein |
| LOC_Os09g06930 | hypothetical protein |
| LOC_Os10g04180 | NB-ARC domain containing protein, expressed |
| LOC_Os10g04210 | expressed protein |
| LOC_Os10g19919 | retrotransposon protein, putative, Ty3-gypsy subclass, expressed |
| LOC_Os10g21398 | NADPH-dependent oxidoreductase, putative, expressed |
| LOC_Os10g26700 | YGL010w, putative, expressed |
| LOC_Os10g34370 | expressed protein |
| LOC_Os10g37680 | expressed protein |
| LOC_Os11g01154 | dehydrogenase, putative, expressed |
| LOC_Os11g01170 | tonneau 1b, putative, expressed |
| LOC_Os11g01180 | ulp1 protease family protein, putative, expressed |
| LOC_Os11g01200 | senescence-induced receptor-like serine/threonine-protein kinase precursor, putative, expressed |
| LOC_Os11g01210 | HCF152, putative, expressed |
| LOC_Os11g01400 | expressed protein |
| LOC_Os11g01490 | outer membrane protein, OMP85 family, putative, expressed |
| LOC_Os11g01530 | ferritin-1, chloroplast precursor, putative, expressed |
| LOC_Os11g02770 | expressed protein |
| LOC_Os11g02774 | histidine triad family protein, putative, expressed |
| LOC_Os11g02820 | CRP10 - Cysteine-rich family protein precursor, expressed |
| LOC_Os11g02850 | expressed protein |

| | |
|---|---|
| LOC_Os11g03230 | nucleoside-triphosphatase, putative, expressed |
| LOC_Os11g04160 | dehydrogenase-phosphopantetheinyltransferase, putative, expressed |
| LOC_Os11g04210 | hydroxymethylglutaryl-CoA lyase, putative, expressed |
| LOC_Os11g04230 | transcription initiation factor, putative, expressed |
| LOC_Os11g04270 | expressed protein |
| LOC_Os11g04280 | zinc finger, C3HC4 type domain containing protein |
| LOC_Os11g15720 | expressed protein |
| LOC_Os11g25490 | expressed protein |
| LOC_Os11g28184 | expressed protein |
| LOC_Os11g33360 | hypothetical protein |
| LOC_Os11g45720 | pectinesterase, putative, expressed |
| LOC_Os12g01180 | WRKY57, expressed |
| LOC_Os12g03940 | expressed protein |
| LOC_Os12g15390 | expressed protein |
| LOC_Os12g19440 | hypothetical protein |
| LOC_Os12g20260 | expressed protein |
| LOC_Os12g22920 | expressed protein |
| LOC_Os12g37630 | domain of unknown function family protein, expressed |
| LOC_Os12g37740 | resistance protein LR10, putative, expressed |

**Genes in rufipogon_SNP CLR**

| | |
|---|---|
| LOC_Os01g01530 | transposon protein, putative, CACTA, En/Spm sub-class |
| LOC_Os01g01540 | retrotransposon protein, putative, unclassified, expressed |
| LOC_Os01g01550 | transposon protein, putative, CACTA, En/Spm sub-class, expressed |
| LOC_Os01g02240 | transposon protein, putative, unclassified, expressed |
| LOC_Os01g02610 | protein kinase domain containing protein, expressed |
| LOC_Os01g02629 | retrotransposon protein, putative, unclassified, expressed |
| LOC_Os01g03452 | transposon protein, putative, unclassified, expressed |
| LOC_Os01g03720 | MYB family transcription factor, putative, expressed |
| LOC_Os01g03830 | retrotransposon protein, putative, unclassified, expressed |

| | |
|---|---|
| LOC_Os01g03880 | retrotransposon protein, putative, Ty3-gypsy subclass |
| LOC_Os01g04850 | retrotransposon protein, putative, unclassified, expressed |
| LOC_Os01g05320 | retrotransposon protein, putative, Ty1-copia subclass, expressed |
| LOC_Os01g06530 | retrotransposon protein, putative, unclassified, expressed |
| LOC_Os01g06540 | PHD finger protein, putative, expressed |
| LOC_Os01g06800 | expressed protein |
| LOC_Os01g08450 | ras-related protein, putative, expressed |
| LOC_Os01g08890 | expressed protein |
| LOC_Os01g08930 | DEAD-box ATP-dependent RNA helicase, putative, expressed |
| LOC_Os01g12170 | retrotransposon protein, putative, unclassified, expressed |
| LOC_Os01g14260 | transposon protein, putative, Ac/Ds sub-class |
| LOC_Os01g14450 | retrotransposon protein, putative, Ty3-gypsy subclass, expressed |
| LOC_Os01g14790 | expressed protein |
| LOC_Os01g14990 | GCRP6 - Glycine and cysteine rich family protein precursor, putative, expressed |
| LOC_Os01g15080 | hypothetical protein |
| LOC_Os01g15420 | retrotransposon protein, putative, Ty3-gypsy subclass, expressed |
| LOC_Os01g16050 | transposon protein, putative, unclassified, expressed |
| LOC_Os01g17970 | retrotransposon protein, putative, Ty1-copia subclass |
| LOC_Os01g18000 | retrotransposon protein, putative, Ty3-gypsy subclass, expressed |
| LOC_Os01g18140 | expressed protein |
| LOC_Os01g19010 | retrotransposon protein, putative, unclassified, expressed |
| LOC_Os01g19710 | transposon protein, putative, CACTA, En/Spm sub-class, expressed |
| LOC_Os01g19894 | transposon protein, putative, unclassified, expressed |
| LOC_Os01g20060 | expressed protein |
| LOC_Os01g20130 | transposon protein, putative, CACTA, En/Spm sub-class, expressed |
| LOC_Os01g20140 | latency associated nuclear antigen, putative, expressed |
| LOC_Os01g21490 | transposon protein, putative, CACTA, En/Spm sub-class |
| LOC_Os01g22300 | retrotransposon protein, putative, Ty3-gypsy subclass |
| LOC_Os01g22309 | hypothetical protein |
| LOC_Os01g22880 | expressed protein |

| | |
|---|---|
| LOC_Os01g23070 | expressed protein |
| LOC_Os01g23180 | retrotransposon protein, putative, unclassified |
| LOC_Os01g23420 | transposon protein, putative, unclassified, expressed |
| LOC_Os01g24130 | transposon protein, putative, CACTA, En/Spm sub-class, expressed |
| LOC_Os01g24360 | retrotransposon protein, putative, unclassified |
| LOC_Os01g25065 | ATP synthase subunit beta, putative, expressed |
| LOC_Os01g27070 | expressed protein |
| LOC_Os01g27230 | 12-oxophytodienoate reductase, putative, expressed |
| LOC_Os01g27330 | expressed protein |
| LOC_Os01g27670 | transposon protein, putative, CACTA, En/Spm sub-class |
| LOC_Os01g27970 | hypothetical protein |
| LOC_Os01g28020 | retrotransposon protein, putative, Ty3-gypsy subclass |
| LOC_Os01g28270 | retrotransposon protein, putative, Ty1-copia subclass |
| LOC_Os01g29040 | retrotransposon protein, putative, unclassified, expressed |
| LOC_Os01g29104 | expressed protein |
| LOC_Os01g32530 | transposon protein, putative, CACTA, En/Spm sub-class |
| LOC_Os01g32550 | hypothetical protein |
| LOC_Os01g32610 | expressed protein |
| LOC_Os01g33150 | retrotransposon protein, putative, Ty3-gypsy subclass |
| LOC_Os01g33940 | expressed protein |
| LOC_Os01g34130 | retrotransposon protein, putative, Ty3-gypsy subclass |
| LOC_Os01g34510 | hypothetical protein |
| LOC_Os01g35660 | retrotransposon protein, putative, Ty3-gypsy subclass, expressed |
| LOC_Os01g35789 | expressed protein |
| LOC_Os01g35890 | retrotransposon protein, putative, Ty1-copia subclass |
| LOC_Os01g36370 | transposon protein, putative, unclassified, expressed |
| LOC_Os01g39136 | retrotransposon protein, putative, unclassified, expressed |
| LOC_Os01g40120 | expressed protein |
| LOC_Os01g40910 | transposon protein, putative, unclassified |
| LOC_Os01g41430 | UDP-glucoronosyl and UDP-glucosyl transferase, putative, expressed |

| | |
|---|---|
| LOC_Os01g41480 | retrotransposon protein, putative, Ty3-gypsy subclass, expressed |
| LOC_Os01g41810 | cytochrome P450 72A1, putative, expressed |
| LOC_Os01g42400 | STE_MEK_ste7_MAP2K |
| LOC_Os01g44010 | retrotransposon protein, putative, unclassified, expressed |
| LOC_Os01g44970 | polygalacturonase, putative, expressed |
| LOC_Os01g44980 | peptide deformylase, putative, expressed |
| LOC_Os01g45010 | expressed protein |
| LOC_Os01g45020 | F-box protein interaction domain containing protein, expressed |
| LOC_Os01g45150 | expressed protein |
| LOC_Os01g45160 | expressed protein |
| LOC_Os01g46400 | expressed protein |
| LOC_Os01g46410 | leaf senescence related protein, putative, expressed |
| LOC_Os01g48640 | expressed protein |
| LOC_Os01g48680 | two pore calcium channel protein 1, putative, expressed |
| LOC_Os01g49420 | retrotransposon protein, putative, unclassified, expressed |
| LOC_Os01g50520 | cytochrome P450 monooxygenase CYP711A12, putative, expressed |
| LOC_Os01g50560 | expressed protein |
| LOC_Os01g50600 | expressed protein |
| LOC_Os01g52920 | expressed protein |
| LOC_Os01g54600 | WRKY13, expressed |
| LOC_Os01g57942 | cytochrome b6, putative, expressed |
| LOC_Os01g57945 | cytochrome b6-f complex subunit 4, putative, expressed |
| LOC_Os01g57946 | DNA-directed RNA polymerase subunit alpha, putative, expressed |
| LOC_Os01g57954 | translation initiation factor IF-1, chloroplast, putative, expressed |
| LOC_Os01g57956 | chloroplast 50S ribosomal protein L14, putative, expressed |
| LOC_Os01g57964 | photosystem I P700 chlorophyll a apoprotein A1, putative, expressed |
| LOC_Os01g57966 | photosystem I assembly protein ycf3, putative, expressed |
| LOC_Os01g58020 | ribulose bisphosphate carboxylase large chain precursor, putative, expressed |
| LOC_Os01g58022 | NADPH-dependent oxidoreductase, putative, expressed |
| LOC_Os01g58024 | NADPH-dependent oxidoreductase, putative, expressed |

| | |
|---|---|
| LOC_Os01g58035 | expressed protein |
| LOC_Os01g58038 | photosystem II reaction center protein Z, putative, expressed |
| LOC_Os01g58039 | uncharacterized protein ycf70, putative, expressed |
| LOC_Os01g58049 | photosystem I assembly protein ycf4, putative, expressed |
| LOC_Os01g58444 | expressed protein |
| LOC_Os01g58500 | expressed protein |
| LOC_Os01g58510 | NB-ARC domain containing protein, expressed |
| LOC_Os01g62960 | hypothetical protein |
| LOC_Os01g64880 | expressed protein |
| LOC_Os01g65902 | apocytochrome f precursor, putative, expressed |
| LOC_Os01g65970 | transcription factor HBP-1b, putative, expressed |
| LOC_Os01g68750 | adaptin, putative, expressed |
| LOC_Os01g68760 | WW domain containing protein, expressed |
| LOC_Os01g69890 | expressed protein |
| LOC_Os01g71540 | expressed protein |
| LOC_Os01g71624 | expressed protein |
| LOC_Os01g71930 | glycosyl hydrolases family 17 protein, expressed |
| LOC_Os01g71960 | endonuclease, putative, expressed |
| LOC_Os01g72120 | glutathione S-transferase, putative, expressed |
| LOC_Os01g73100 | retrotransposon protein, putative, unclassified, expressed |
| LOC_Os02g01765 | expressed protein |
| LOC_Os02g05210 | retrotransposon protein, putative, unclassified, expressed |
| LOC_Os02g12654 | expressed protein |
| LOC_Os02g13930 | expressed protein |
| LOC_Os02g16050 | transposon protein, putative, unclassified, expressed |
| LOC_Os02g16930 | hypothetical protein |
| LOC_Os02g17450 | retrotransposon protein, putative, unclassified, expressed |
| LOC_Os02g17760 | cytochrome P450, putative, expressed |
| LOC_Os02g20729 | expressed protein |
| LOC_Os02g20940 | CHCH domain containing protein, expressed |

| | |
|---|---|
| LOC_Os02g21570 | expressed protein |
| LOC_Os02g21580 | PPR repeat containing protein, expressed |
| LOC_Os02g24576 | chloroplast 30S ribosomal protein S7, putative, expressed |
| LOC_Os02g24586 | uncharacterized protein ycf68, putative, expressed |
| LOC_Os02g24612 | ribosomal protein S2, putative, expressed |
| LOC_Os02g24614 | DNA-directed RNA polymerase subunit beta, putative, expressed |
| LOC_Os02g24632 | photosystem II 44 kDa reaction center protein, putative, expressed |
| LOC_Os02g28050 | hypothetical protein |
| LOC_Os02g28530 | expressed protein |
| LOC_Os02g28920 | transposon protein, putative, unclassified, expressed |
| LOC_Os02g29570 | retrotransposon protein, putative, Ty3-gypsy subclass, expressed |
| LOC_Os02g29910 | hypothetical protein |
| LOC_Os02g30280 | expressed protein |
| LOC_Os02g30450 | harpin-induced protein 1 domain containing protein, expressed |
| LOC_Os02g30460 | uncharacterized protein MJ0304, putative, expressed |
| LOC_Os02g30480 | retrotransposon protein, putative, unclassified, expressed |
| LOC_Os02g36250 | expressed protein |
| LOC_Os02g40870 | phosphatidylinositol N-acetylglucosaminyltransferase subunit C, putative, expressed |
| LOC_Os02g41450 | no apical meristem protein, putative, expressed |
| LOC_Os02g42030 | hypothetical protein |
| LOC_Os02g43800 | retrotransposon protein, putative, Ty3-gypsy subclass, expressed |
| LOC_Os02g45370 | retrotransposon protein, putative, Ty3-gypsy subclass, expressed |
| LOC_Os02g48560 | fatty acid desaturase, putative, expressed |
| LOC_Os02g48840 | dual specificity protein phosphatase, putative, expressed |
| LOC_Os02g49332 | CSLE2 - cellulose synthase-like family E, expressed |
| LOC_Os02g50480 | histidine kinase, putative, expressed |
| LOC_Os02g53690 | growth regulating factor protein, putative, expressed |
| LOC_Os02g55840 | expressed protein |
| LOC_Os02g56030 | expressed protein |
| LOC_Os02g56080 | retrotransposon protein, putative, Ty3-gypsy subclass, expressed |

| | |
|---|---|
| LOC_Os02g56090 | expressed protein |
| LOC_Os02g56460 | dehydrogenase, putative, expressed |
| LOC_Os02g56740 | translation initiation factor eIF-2B subunit epsilon, putative, expressed |
| LOC_Os02g57570 | MATE efflux family protein, putative, expressed |
| LOC_Os02g57620 | Citrate transporter protein, putative, expressed |
| LOC_Os02g57640 | KH domain containing protein, putative, expressed |
| LOC_Os02g57650 | No apical meristem protein, putative, expressed |
| LOC_Os02g58110 | expressed protein |
| LOC_Os02g58120 | oxidoreductase/ transition metal ion binding protein, putative, expressed |
| LOC_Os02g58160 | expressed protein |
| LOC_Os02g58230 | C2 domain containing protein, putative, expressed |
| LOC_Os02g58250 | expressed protein |
| LOC_Os02g58260 | metallo-beta-lactamase family protein, putative, expressed |
| LOC_Os02g58410 | expressed protein |
| LOC_Os02g58440 | zinc finger C-x8-C-x5-C-x3-H type family protein, expressed |
| LOC_Os03g05750 | heavy-metal-associated domain-containing protein, putative, expressed |
| LOC_Os03g06610 | expressed protein |
| LOC_Os03g09830 | expressed protein |
| LOC_Os03g13920 | transposon protein, putative, CACTA, En/Spm sub-class, expressed |
| LOC_Os03g23940 | expressed protein |
| LOC_Os03g23950 | adapitin protein, putative, expressed |
| LOC_Os03g24960 | ulp1 protease family, C-terminal catalytic domain containing protein, expressed |
| LOC_Os03g24970 | SWIM zinc finger family protein, putative |
| LOC_Os03g28920 | expressed protein |
| LOC_Os03g29200 | retrotransposon protein, putative, unclassified, expressed |
| LOC_Os03g32190 | transposon protein, putative, unclassified, expressed |
| LOC_Os03g32526 | tRNA-splicing endonuclease positive effector-related, putative, expressed |
| LOC_Os03g36070 | retrotransposon protein, putative, unclassified |
| LOC_Os03g40180 | 60S ribosomal protein L15, putative, expressed |
| LOC_Os03g42180 | hypothetical protein |

| | |
|---|---|
| LOC_Os03g45710 | 2Fe-2S iron-sulfur cluster binding domain containing protein, expressed |
| LOC_Os03g46560 | RPD1, putative, expressed |
| LOC_Os03g48260 | expressed protein |
| LOC_Os03g48471 | ATP synthase subunit alpha, putative, expressed |
| LOC_Os03g59760 | RING finger protein 126, putative, expressed |
| LOC_Os03g59770 | EF hand family protein, putative, expressed |
| LOC_Os04g02090 | retrotransposon protein, putative, unclassified, expressed |
| LOC_Os04g02970 | subtilisin-like protease precursor, putative, expressed |
| LOC_Os04g03310 | expressed protein |
| LOC_Os04g03320 | jacalin-like lectin domain containing protein, expressed |
| LOC_Os04g03579 | protein kinase, putative, expressed |
| LOC_Os04g04010 | expressed protein |
| LOC_Os04g04020 | protein transport protein Sec24-like, putative, expressed |
| LOC_Os04g04030 | expressed protein |
| LOC_Os04g09350 | expressed protein |
| LOC_Os04g10890 | expressed protein |
| LOC_Os04g13430 | expressed protein |
| LOC_Os04g13470 | expressed protein |
| LOC_Os04g14290 | hypothetical protein |
| LOC_Os04g16742 | NADPH-dependent oxidoreductase, putative, expressed |
| LOC_Os04g16745 | chloroplast 30S ribosomal protein S14, putative, expressed |
| LOC_Os04g16748 | ATP synthase B chain, putative, expressed |
| LOC_Os04g16782 | chloroplast 50S ribosomal protein L23, putative |
| LOC_Os04g16792 | uncharacterized protein ycf76, putative |
| LOC_Os04g16794 | hypothetical protein |
| LOC_Os04g16848 | photosystem II reaction center protein H, putative |
| LOC_Os04g17050 | glutaredoxin family protein, expressed |
| LOC_Os04g17064 | expressed protein |
| LOC_Os04g17090 | retrotransposon protein, putative, unclassified, expressed |
| LOC_Os04g17490 | expressed protein |

| | |
|---|---|
| LOC_Os04g18380 | cytochrome P450, putative, expressed |
| LOC_Os04g19740 | transketolase, chloroplast precursor, putative, expressed |
| LOC_Os04g20424 | neural Wiskott-Aldrich syndrome protein, putative, expressed |
| LOC_Os04g20430 | expressed protein |
| LOC_Os04g20800 | pumilio domain-containing protein, putative, expressed |
| LOC_Os04g21880 | expressed protein |
| LOC_Os04g21890 | disease resistance protein RPM1, putative, expressed |
| LOC_Os04g21900 | expressed protein |
| LOC_Os04g22380 | short chain dehydrogenase/reductase protein, putative, expressed |
| LOC_Os04g25440 | cytokinin-O-glucosyltransferase 2, putative, expressed |
| LOC_Os04g27700 | expressed protein |
| LOC_Os04g28500 | expressed protein |
| LOC_Os04g30920 | expressed protein |
| LOC_Os04g30980 | expressed protein |
| LOC_Os04g31000 | methyltransferase domain containing protein, expressed |
| LOC_Os04g33340 | expressed protein |
| LOC_Os04g40600 | peptide methionine sulfoxide reductase, putative, expressed |
| LOC_Os04g53496 | NBS-LRR disease resistance protein, putative, expressed |
| LOC_Os05g01390 | retrotransposon protein, putative, unclassified |
| LOC_Os05g10270 | expressed protein |
| LOC_Os05g14150 | polygalacturonase precursor, putative, expressed |
| LOC_Os05g22940 | acetyl-CoA carboxylase, putative, expressed |
| LOC_Os05g28090 | expressed protein |
| LOC_Os05g31630 | transposon protein, putative, unclassified, expressed |
| LOC_Os05g46390 | OsFBX175 - F-box domain containing protein, expressed |
| LOC_Os05g49990 | expressed protein |
| LOC_Os05g50110 | oleosin, putative, expressed |
| LOC_Os06g05600 | OsFBDUF31 - F-box and DUF domain containing protein, expressed |
| LOC_Os06g07340 | hypothetical protein |
| LOC_Os06g07350 | RNA-binding motif protein, putative, expressed |

| | |
|---|---|
| LOC_Os06g16070 | receptor-like protein kinase 2 precursor, putative, expressed |
| LOC_Os06g16110 | expressed protein |
| LOC_Os06g16150 | expressed protein |
| LOC_Os06g16550 | hypothetical protein |
| LOC_Os06g16820 | expressed protein |
| LOC_Os06g18930 | cadmium tolerance factor, putative, expressed |
| LOC_Os06g20370 | microtubule associated protein, putative, expressed |
| LOC_Os06g20550 | expressed protein |
| LOC_Os06g30470 | expressed protein |
| LOC_Os06g36970 | transposon protein, putative, unclassified, expressed |
| LOC_Os06g38590 | receptor-like protein kinase precursor, putative, expressed |
| LOC_Os06g38710 | expressed protein |
| LOC_Os06g38720 | expressed protein |
| LOC_Os06g38740 | expressed protein |
| LOC_Os06g39708 | photosystem II P680 chlorophyll A apoprotein, putative, expressed |
| LOC_Os06g39712 | OsClp10 - Putative Clp protease homologue, expressed |
| LOC_Os06g39716 | chloroplast 30S ribosomal protein S18, putative, expressed |
| LOC_Os06g39728 | cytochrome b559 subunit alpha, putative |
| LOC_Os06g39748 | expressed protein |
| LOC_Os06g45360 | peptidase, M24 family protein, putative, expressed |
| LOC_Os06g49505 | arogenate dehydrogenase 1, chloroplast precursor, putative, expressed |
| LOC_Os06g49510 | zinc knuckle family protein, putative, expressed |
| LOC_Os06g49530 | OsFBX206 - F-box domain containing protein, expressed |
| LOC_Os06g49620 | expressed protein |
| LOC_Os07g07790 | LTPL75 - Protease inhibitor/seed storage/LTP family protein precursor, expressed |
| LOC_Os07g07870 | LTPL77 - Protease inhibitor/seed storage/LTP family protein precursor, putative, expressed |
| LOC_Os07g07880 | trp repressor/replication initiator, putative, expressed |
| LOC_Os07g07890 | expressed protein |
| LOC_Os07g07900 | expressed protein |
| LOC_Os07g09490 | ATPase, putative, expressed |

| | |
|---|---|
| LOC_Os07g11630 | LTPL163 - Protease inhibitor/seed storage/LTP family protein precursor, expressed |
| LOC_Os07g12510 | AP2 domain containing protein, expressed |
| LOC_Os07g12940 | expressed protein |
| LOC_Os07g13030 | expressed protein |
| LOC_Os07g13040 | expressed protein |
| LOC_Os07g13050 | hypothetical protein |
| LOC_Os07g18930 | expressed protein |
| LOC_Os07g19020 | conserved hypothetical protein |
| LOC_Os07g19030 | tic22-like family domain containing protein, expressed |
| LOC_Os07g23460 | retrotransposon protein, putative, Ty1-copia subclass, expressed |
| LOC_Os07g26740 | 60S ribosomal protein L44, putative, expressed |
| LOC_Os07g26750 | expressed protein |
| LOC_Os07g26770 | circumsporozoite protein precursor, putative |
| LOC_Os07g30410 | TKL_IRAK_DUF26-la |
| LOC_Os07g30494 | hypothetical protein |
| LOC_Os07g38930 | glucan endo-1,3-beta-glucosidase precursor, putative, expressed |
| LOC_Os07g41930 | DC1 domain-containing protein, putative, expressed |
| LOC_Os07g41980 | C1-like domain containing protein, expressed |
| LOC_Os07g42030 | DC1 domain-containing protein, putative, expressed |
| LOC_Os07g42080 | expressed protein |
| LOC_Os08g10730 | expressed protein |
| LOC_Os08g11599 | expressed protein |
| LOC_Os08g12960 | speckle-type POZ protein-like, putative, expressed |
| LOC_Os08g13040 | hypothetical protein |
| LOC_Os08g15306 | chloroplast 50S ribosomal protein L20, putative, expressed |
| LOC_Os08g15530 | expressed protein |
| LOC_Os08g15540 | expressed protein |
| LOC_Os08g15545 | DEFL55 - Defensin and Defensin-like DEFL family, expressed |
| LOC_Os08g15560 | hypothetical protein |
| LOC_Os08g17050 | expressed protein |

| | |
|---|---|
| LOC_Os08g26210 | expressed protein |
| LOC_Os08g44880 | expressed protein |
| LOC_Os09g08910 | ATP synthase, putative, expressed |
| LOC_Os09g12580 | expressed protein |
| LOC_Os09g14870 | expressed protein |
| LOC_Os09g21520 | DEAD-box ATP-dependent RNA helicase, putative, expressed |
| LOC_Os09g23210 | retrotransposon protein, putative, unclassified |
| LOC_Os09g24810 | ZF-HD protein dimerisation region containing protein, expressed |
| LOC_Os10g09360 | SAM dependent carboxyl methyltransferase family protein, putative, expressed |
| LOC_Os10g10244 | alanyl-tRNA synthetase, putative, expressed |
| LOC_Os10g10260 | expressed protein |
| LOC_Os10g18870 | dirigent, putative, expressed |
| LOC_Os10g21230 | ATP synthase C chain, putative, expressed |
| LOC_Os10g21236 | chloroplast ATP synthase a chain precursor, putative, expressed |
| LOC_Os10g22039 | mitochondrial import inner membrane translocase subunit Tim17, putative, expressed |
| LOC_Os10g24070 | expressed protein |
| LOC_Os10g25110 | pumilio-family RNA binding repeat containing protein, expressed |
| LOC_Os10g28450 | CAF1 family ribonuclease containing protein, expressed |
| LOC_Os10g33040 | receptor-like protein kinase precursor, putative, expressed |
| LOC_Os11g02379 | LTPL6 - Protease inhibitor/seed storage/LTP family protein precursor, expressed |
| LOC_Os11g03380 | expressed protein |
| LOC_Os11g03390 | FHA domain containing protein, putative, expressed |
| LOC_Os11g03400 | ribosomal protein, putative, expressed |
| LOC_Os11g03410 | exostosin family domain containing protein, expressed |
| LOC_Os11g04020 | major facilitator superfamily antiporter, putative, expressed |
| LOC_Os11g28470 | expressed protein |
| LOC_Os11g30060 | NBS-LRR type disease resistance protein Hom-B, putative, expressed |
| LOC_Os11g38270 | hypothetical protein |
| LOC_Os11g39770 | expressed protein |
| LOC_Os11g39840 | expressed protein |

| | |
|---|---|
| LOC_Os11g39850 | expressed protein |
| LOC_Os11g48060 | laccase-22 precursor, putative, expressed |
| LOC_Os11g48064 | hypothetical protein |
| LOC_Os12g07730 | SWIRM domain containing protein, expressed |
| LOC_Os12g07740 | eukaryotic translation initiation factor 2 subunit gamma, putative, expressed |
| LOC_Os12g10810 | expressed protein |
| LOC_Os12g14630 | retrotransposon protein, putative, LINE subclass, expressed |
| LOC_Os12g14910 | hypothetical protein |
| LOC_Os12g23300 | expressed protein |
| LOC_Os12g23310 | expressed protein |
| LOC_Os12g23330 | expressed protein |
| LOC_Os12g23550 | expressed protein |
| LOC_Os12g28720 | hypothetical protein |
| LOC_Os12g30950 | F-box protein interaction domain containing protein, expressed |
| LOC_Os12g31090 | hypothetical protein |
| LOC_Os12g33270 | expressed protein |
| LOC_Os12g33952 | expressed protein |
| LOC_Os12g34062 | mitochondrial ribosomal protein S12, putative, expressed |
| LOC_Os12g34094 | NADH-ubiquinone oxidoreductase chain 4, putative, expressed |
| LOC_Os12g34100 | expressed protein |
| LOC_Os12g34108 | ATP synthase protein 9, mitochondrial, putative, expressed |
| LOC_Os12g36680 | expressed protein |
| LOC_Os12g39530 | hypothetical protein |

**Genes in rufipogon_SV CLR**

| | |
|---|---|
| LOC_Os01g01010 | TBC domain containing protein, expressed |
| LOC_Os01g01019 | expressed protein |
| LOC_Os01g01030 | monocopper oxidase, putative, expressed |
| LOC_Os01g01040 | expressed protein |
| LOC_Os01g01060 | 40S ribosomal protein S5, putative, expressed |

| | |
|---|---|
| LOC_Os01g01070 | expressed protein |
| LOC_Os01g01080 | decarboxylase, putative, expressed |
| LOC_Os01g01090 | retrotransposon protein, putative, unclassified, expressed |
| LOC_Os01g01280 | expressed protein |
| LOC_Os01g01290 | histone-like transcription factor and archaeal histone, putative, expressed |
| LOC_Os01g01295 | expressed protein |
| LOC_Os01g01302 | shikimate kinase, putative, expressed |
| LOC_Os01g01307 | translocon-associated protein beta domain containing protein, expressed |
| LOC_Os01g01320 | retrotransposon protein, putative, Ty1-copia subclass, expressed |
| LOC_Os01g01340 | light-induced protein 1-like, putative, expressed |
| LOC_Os01g01350 | SNF7 domain containing protein, putative, expressed |
| LOC_Os01g01530 | transposon protein, putative, CACTA, En/Spm sub-class |
| LOC_Os01g01910 | expressed protein |
| LOC_Os01g01990 | expressed protein |
| LOC_Os01g02000 | phosphate transporter 1, putative, expressed |
| LOC_Os01g02020 | acetyl-CoA acetyltransferase, cytosolic, putative, expressed |
| LOC_Os01g02040 | retrotransposon protein, putative, unclassified, expressed |
| LOC_Os01g02050 | phosphoenolpyruvate carboxylase, putative, expressed |
| LOC_Os01g03720 | MYB family transcription factor, putative, expressed |
| LOC_Os01g04160 | expressed protein |
| LOC_Os01g04170 | expressed protein |
| LOC_Os01g04190 | transporter family protein, putative, expressed |
| LOC_Os01g04200 | metallo-beta-lactamase family protein, putative, expressed |
| LOC_Os01g04210 | DNA binding protein, putative, expressed |
| LOC_Os01g04230 | inactive receptor kinase At2g26730 precursor, putative, expressed |
| LOC_Os01g05180 | expressed protein |
| LOC_Os01g05240 | retrotransposon protein, putative, unclassified |
| LOC_Os01g05320 | retrotransposon protein, putative, Ty1-copia subclass, expressed |
| LOC_Os01g06160 | retrotransposon protein, putative, Ty3-gypsy subclass, expressed |
| LOC_Os01g06740 | ribosome inactivating protein, putative, expressed |

| | |
|---|---|
| LOC_Os01g06810 | retrotransposon protein, putative, Ty1-copia subclass, expressed |
| LOC_Os01g07180 | expressed protein |
| LOC_Os01g07376 | thioredoxin, putative, expressed |
| LOC_Os01g07382 | expressed protein |
| LOC_Os01g07390 | zinc finger, C3HC4 type family protein, expressed |
| LOC_Os01g07410 | growth regulator related protein, putative, expressed |
| LOC_Os01g07420 | hydrolase, alpha/beta fold family protein, putative, expressed |
| LOC_Os01g07450 | MYB family transcription factor, putative, expressed |
| LOC_Os01g07460 | conserved hypothetical protein |
| LOC_Os01g07480 | DUF260 domain containing protein, putative, expressed |
| LOC_Os01g07490 | RST1, putative, expressed |
| LOC_Os01g07500 | alliin lyase precursor, putative, expressed |
| LOC_Os01g07510 | conserved hypothetical protein |
| LOC_Os01g07550 | hypothetical protein |
| LOC_Os01g07590 | universal stress protein domain containing protein, putative, expressed |
| LOC_Os01g07600 | expressed protein |
| LOC_Os01g07610 | PPR repeat domain containing protein, putative, expressed |
| LOC_Os01g07620 | expressed protein |
| LOC_Os01g07630 | BRASSINOSTEROID INSENSITIVE 1-associated receptor kinase 1 precursor, putative, expressed |
| LOC_Os01g07640 | ankyrin, putative, expressed |
| LOC_Os01g07650 | expressed protein |
| LOC_Os01g07660 | expressed protein |
| LOC_Os01g07670 | expressed protein |
| LOC_Os01g07700 | expressed protein |
| LOC_Os01g07710 | expressed protein |
| LOC_Os01g07720 | dolichyl-P-Man Man-PP-dolichyl mannosyltransferase, putative, expressed |
| LOC_Os01g07730 | phosphate/phosphoenolpyruvate translocator-related protein, putative, expressed |
| LOC_Os01g07740 | DEAD-box ATP-dependent RNA helicase 14, putative, expressed |
| LOC_Os01g07750 | expressed protein |
| LOC_Os01g07760 | phospholipase D, putative, expressed |

| | |
|---|---|
| LOC_Os01g07770 | peroxidase precursor, putative, expressed |
| LOC_Os01g07780 | embryo-specific 3, putative, expressed |
| LOC_Os01g07790 | polygalacturonase, putative, expressed |
| LOC_Os01g07810 | protein of unknown function domain containing protein, expressed |
| LOC_Os01g07850 | glyoxalase family protein, putative, expressed |
| LOC_Os01g07860 | retrotransposon protein, putative, unclassified, expressed |
| LOC_Os01g08190 | transcriptional corepressor LEUNIG, putative, expressed |
| LOC_Os01g08200 | ubiquitin carboxyl-terminal hydrolase 14, putative, expressed |
| LOC_Os01g08220 | gibberellin 3-beta-dioxygenase 2-2, putative, expressed |
| LOC_Os01g08240 | hypothetical protein |
| LOC_Os01g08250 | retrotransposon protein, putative, unclassified, expressed |
| LOC_Os01g08260 | pleiotropic drug resistance protein 4, putative, expressed |
| LOC_Os01g08310 | serine/threonine protein phosphatase 2A 59 kDa regulatory subunit Bgamma isoform, putative, expressed |
| LOC_Os01g08320 | OsIAA1 - Auxin-responsive Aux/IAA gene family member, expressed |
| LOC_Os01g08330 | aspartic proteinase nepenthesin-1 precursor, putative, expressed |
| LOC_Os01g08340 | zinc finger family protein, putative, expressed |
| LOC_Os01g08350 | ATP synthase subunit alpha, mitochondrial, putative, expressed |
| LOC_Os01g08360 | hypothetical protein |
| LOC_Os01g08370 | symbiosis-related disease resistance protein, putative, expressed |
| LOC_Os01g08380 | transferase family protein, putative, expressed |
| LOC_Os01g08390 | hypothetical protein |
| LOC_Os01g08400 | syntaxin, putative, expressed |
| LOC_Os01g08410 | early fruit mRNA, putative, expressed |
| LOC_Os01g08420 | EMB2410, putative, expressed |
| LOC_Os01g08430 | expressed protein |
| LOC_Os01g08440 | UDP-glucoronosyl and UDP-glucosyl transferase domain containing protein, expressed |
| LOC_Os01g08450 | ras-related protein, putative, expressed |
| LOC_Os01g08540 | DNA mismatch repair protein MSH6-2, putative, expressed |
| LOC_Os01g08550 | aminoacyl-tRNA synthetase, putative, expressed |
| LOC_Os01g08710 | WRKY102, expressed |

| | |
|---|---|
| LOC_Os01g08740 | expressed protein |
| LOC_Os01g08750 | expressed protein |
| LOC_Os01g08780 | endonuclease/exonuclease/phosphatase family domain containing protein, expressed |
| LOC_Os01g08790 | histone-like transcription factor and archaeal histone, putative, expressed |
| LOC_Os01g08800 | cytochrome P450, putative, expressed |
| LOC_Os01g08810 | cytochrome P450, putative, expressed |
| LOC_Os01g08814 | expressed protein |
| LOC_Os01g08820 | PHD-finger domain containing protein, putative, expressed |
| LOC_Os01g08830 | OsFBDUF1 - F-box and DUF domain containing protein, expressed |
| LOC_Os01g08840 | expressed protein |
| LOC_Os01g08850 | expressed protein |
| LOC_Os01g08860 | hsp20/alpha crystallin family protein, putative, expressed |
| LOC_Os01g08870 | expressed protein |
| LOC_Os01g08880 | expressed protein |
| LOC_Os01g08890 | expressed protein |
| LOC_Os01g08930 | DEAD-box ATP-dependent RNA helicase, putative, expressed |
| LOC_Os01g08940 | expressed protein |
| LOC_Os01g08950 | expressed protein |
| LOC_Os01g08970 | SSRP1-like FACT complex subunit, putative, expressed |
| LOC_Os01g08980 | expressed protein |
| LOC_Os01g08990 | expressed protein |
| LOC_Os01g09000 | glutaminyl-tRNA synthetase, putative, expressed |
| LOC_Os01g09060 | retrotransposon protein, putative, Ty1-copia subclass |
| LOC_Os01g09080 | WRKY107, expressed |
| LOC_Os01g09110 | retrotransposon protein, putative, unclassified |
| LOC_Os01g09150 | expressed protein |
| LOC_Os01g09200 | AGC_AGC_other_NDRh_TRCd |
| LOC_Os01g09220 | transposon protein, putative, CACTA, En/Spm sub-class, expressed |
| LOC_Os01g09500 | expressed protein |
| LOC_Os01g09510 | 60S acidic ribosomal protein, putative, expressed |

| | |
|---|---|
| LOC_Os01g09540 | HAD superfamily phosphatase, putative, expressed |
| LOC_Os01g09550 | no apical meristem protein, putative, expressed |
| LOC_Os01g09560 | mitochondrial-processing peptidase subunit alpha, mitochondrial precursor, putative, expressed |
| LOC_Os01g09570 | 6-phosphofructokinase, putative, expressed |
| LOC_Os01g09960 | transposon protein, putative, unclassified, expressed |
| LOC_Os01g10030 | expressed protein |
| LOC_Os01g10560 | transposon protein, putative, unclassified |
| LOC_Os01g11820 | expressed protein |
| LOC_Os01g12040 | transposon protein, putative, unclassified, expressed |
| LOC_Os01g13050 | long cell-linked locus protein, putative, expressed |
| LOC_Os01g13500 | expressed protein |
| LOC_Os01g13510 | expressed protein |
| LOC_Os01g15420 | retrotransposon protein, putative, Ty3-gypsy subclass, expressed |
| LOC_Os01g16730 | ulp1 protease family, C-terminal catalytic domain containing protein, expressed |
| LOC_Os01g16740 | transposon protein, putative, unclassified, expressed |
| LOC_Os01g18050 | tubulin/FtsZ domain containing protein, putative, expressed |
| LOC_Os01g18320 | protoporphyrinogen oxidase, chloroplast precursor, putative, expressed |
| LOC_Os01g19910 | expressed protein |
| LOC_Os01g24030 | AMP-binding enzyme, putative, expressed |
| LOC_Os01g28680 | WD domain, G-beta repeat domain containing protein, expressed |
| LOC_Os01g29040 | retrotransposon protein, putative, unclassified, expressed |
| LOC_Os01g29680 | retrotransposon protein, putative, Ty1-copia subclass, expressed |
| LOC_Os01g32490 | hypothetical protein |
| LOC_Os01g34560 | 3-ketoacyl-CoA synthase, putative, expressed |
| LOC_Os01g34620 | OsGrx_S15 |
| LOC_Os01g37170 | retrotransposon protein, putative, unclassified, expressed |
| LOC_Os01g37590 | peptide transporter PTR2, putative, expressed |
| LOC_Os01g38100 | expressed protein |
| LOC_Os01g38680 | transporter family protein, putative, expressed |
| LOC_Os01g41145 | retrotransposon protein, putative, unclassified, expressed |

| | |
|---|---|
| LOC_Os01g46860 | inorganic phosphate transporter, putative, expressed |
| LOC_Os01g52020 | subtilase, putative, expressed |
| LOC_Os01g52750 | OsSub3 - Putative Subtilisin homologue, expressed |
| LOC_Os01g58230 | expressed protein |
| LOC_Os01g60020 | NAC domain transcription factor, putative, expressed |
| LOC_Os01g60549 | retrotransposon protein, putative, unclassified, expressed |
| LOC_Os01g61520 | expressed protein |
| LOC_Os01g62090 | expressed protein |
| LOC_Os01g62120 | expressed protein |
| LOC_Os01g66010 | amino acid transporter, putative, expressed |
| LOC_Os01g68230 | expressed protein |
| LOC_Os01g69904 | expressed protein |
| LOC_Os01g71150 | expressed protein |
| LOC_Os01g72650 | RNA recognition motif containing protein, putative, expressed |
| LOC_Os01g72700 | expressed protein |
| LOC_Os01g73530 | ABC transporter, ATP-binding protein, putative, expressed |
| LOC_Os01g74450 | aquaporin protein, putative, expressed |
| LOC_Os02g01170 | HECT-domain domain containing protein, expressed |
| LOC_Os02g01180 | kinesin motor domain containing protein, putative, expressed |
| LOC_Os02g01190 | POEI25 - Pollen Ole e I allergen and extensin family protein precursor, expressed |
| LOC_Os02g01200 | glutaredoxin, putative, expressed |
| LOC_Os02g01220 | rhodanese-like domain containing protein, putative, expressed |
| LOC_Os02g01240 | CPuORF11 - conserved peptide uORF-containing transcript, expressed |
| LOC_Os02g01250 | LSM domain containing protein, expressed |
| LOC_Os02g01270 | START domain containing protein, expressed |
| LOC_Os02g01280 | T-complex protein, putative, expressed |
| LOC_Os02g01290 | expressed protein |
| LOC_Os02g01300 | invertase/pectin methylesterase inhibitor family protein, putative, expressed |
| LOC_Os02g01310 | invertase/pectin methylesterase inhibitor family protein, putative, expressed |
| LOC_Os02g02700 | AMP-binding enzyme, putative, expressed |

| | |
|---|---|
| LOC_Os02g03250 | expressed protein |
| LOC_Os02g03260 | 3-isopropylmalate dehydratase large subunit 2, putative, expressed |
| LOC_Os02g03270 | AT hook motif domain containing protein, expressed |
| LOC_Os02g03280 | transmembrane BAX inhibitor motif-containing protein, putative, expressed |
| LOC_Os02g03294 | cyclin, putative, expressed |
| LOC_Os02g03310 | KH domain containing protein, putative, expressed |
| LOC_Os02g03320 | plant neutral invertase domain containing protein, expressed |
| LOC_Os02g03330 | expressed protein |
| LOC_Os02g03340 | RNA polymerase II transcription factor B subunit 4, putative, expressed |
| LOC_Os02g03360 | expressed protein |
| LOC_Os02g03410 | CAMK_CAMK_like |
| LOC_Os02g03420 | pentatricopeptide repeat-containing protein, putative, expressed |
| LOC_Os02g03430 | expressed protein |
| LOC_Os02g03440 | LSM domain containing protein, expressed |
| LOC_Os02g03450 | vesicle transport v-SNARE protein, putative, expressed |
| LOC_Os02g03460 | phosphatidate cytidylyltransferase, putative, expressed |
| LOC_Os02g03470 | tetratricopeptide repeat containing protein, putative, expressed |
| LOC_Os02g03480 | THION24 - Plant thionin family protein precursor, expressed |
| LOC_Os02g03490 | major ampullate spidroin 3, putative, expressed |
| LOC_Os02g03530 | PPR repeat domain containing protein, putative, expressed |
| LOC_Os02g03540 | phosphoribosyl transferase, putative, expressed |
| LOC_Os02g03550 | glycosyl hydrolases family 16, putative, expressed |
| LOC_Os02g03560 | expressed protein |
| LOC_Os02g03570 | hsp20/alpha crystallin family protein, putative, expressed |
| LOC_Os02g03575 | expressed protein |
| LOC_Os02g03580 | transcription factor, putative, expressed |
| LOC_Os02g03590 | expressed protein |
| LOC_Os02g03600 | dnaJ domain containing protein, expressed |
| LOC_Os02g03620 | seven in absentia protein family domain containing protein, expressed |
| LOC_Os02g03660 | OsFBDUF12 - F-box and DUF domain containing protein, expressed |

| | |
|---|---|
| LOC_Os02g03670 | expressed protein |
| LOC_Os02g03680 | expressed protein |
| LOC_Os02g03690 | beta-amylase, putative, expressed |
| LOC_Os02g03700 | expressed protein |
| LOC_Os02g03710 | UP-9A, putative, expressed |
| LOC_Os02g03720 | expressed protein |
| LOC_Os02g03730 | SWIB/MDM2 domain containing protein, expressed |
| LOC_Os02g03740 | pyridoxal biosynthesis protein PDX2, putative, expressed |
| LOC_Os02g03760 | expressed protein |
| LOC_Os02g03780 | hypothetical protein |
| LOC_Os02g03790 | expressed protein |
| LOC_Os02g03810 | OsFBX34 - F-box domain containing protein, expressed |
| LOC_Os02g03820 | expressed protein |
| LOC_Os02g03830 | expressed protein |
| LOC_Os02g03840 | expressed protein |
| LOC_Os02g03850 | glycosyltransferase, putative, expressed |
| LOC_Os02g03860 | mitochondrial ATP synthase precursor, putative, expressed |
| LOC_Os02g03870 | periplasmic beta-glucosidase precursor, putative, expressed |
| LOC_Os02g03880 | expressed protein |
| LOC_Os02g03890 | nuclear transport factor 2, putative, expressed |
| LOC_Os02g03900 | metal transporter Nramp6, putative, expressed |
| LOC_Os02g03910 | OsFBX35 - F-box domain containing protein, expressed |
| LOC_Os02g03940 | expressed protein |
| LOC_Os02g03950 | zinc finger, C3HC4 type family protein, expressed |
| LOC_Os02g03960 | CPuORF1 - conserved peptide uORF-containing transcript, expressed |
| LOC_Os02g03970 | PPR repeat domain containing protein, putative, expressed |
| LOC_Os02g03990 | WD domain, G-beta repeat domain containing protein, expressed |
| LOC_Os02g04010 | cyclin-T1-1, putative, expressed |
| LOC_Os02g04020 | expressed protein |
| LOC_Os02g04030 | phosphatidylinositol transfer, putative, expressed |

| | |
|---|---|
| LOC_Os02g04040 | RecF/RecN/SMC N terminal domain containing protein, expressed |
| LOC_Os02g04090 | retrotransposon protein, putative, unclassified, expressed |
| LOC_Os02g04100 | peptidase, T1 family, putative, expressed |
| LOC_Os02g04110 | expressed protein |
| LOC_Os02g04120 | cytidylyltransferase domain containing protein, expressed |
| LOC_Os02g04130 | DUF1645 domain containing protein, putative, expressed |
| LOC_Os02g04160 | transcription elongation factor 1, putative, expressed |
| LOC_Os02g04170 | L-aspartate oxidase 1, putative, expressed |
| LOC_Os02g04190 | THION23 - Plant thionin family protein precursor, expressed |
| LOC_Os02g04210 | expressed protein |
| LOC_Os02g04220 | expressed protein |
| LOC_Os02g04230 | CGMC_MAPKCMGC_2 |
| LOC_Os02g04240 | wee1-like protein kinase, putative, expressed |
| LOC_Os02g04250 | glycosyltransferase, putative, expressed |
| LOC_Os02g04260 | expressed protein |
| LOC_Os02g04270 | DNA polymerase V, putative, expressed |
| LOC_Os02g04580 | retrotransposon protein, putative, unclassified |
| LOC_Os02g04900 | expressed protein |
| LOC_Os02g05070 | expressed protein |
| LOC_Os02g05240 | expressed protein |
| LOC_Os02g05780 | expressed protein |
| LOC_Os02g06360 | uncharacterized protein CPn_0526/CP_0226/CPj0526/CpB0547, putative, expressed |
| LOC_Os02g06380 | plant-specific domain TIGR01627 family protein, expressed |
| LOC_Os02g06390 | expressed protein |
| LOC_Os02g06400 | growth regulator related protein, putative, expressed |
| LOC_Os02g06730 | expressed protein |
| LOC_Os02g06750 | retrotransposon protein, putative, unclassified, expressed |
| LOC_Os02g06754 | expressed protein |
| LOC_Os02g06760 | expressed protein |
| LOC_Os02g06770 | expressed protein |

| | |
|---|---|
| LOC_Os02g06779 | expressed protein |
| LOC_Os02g06790 | expressed protein |
| LOC_Os02g06820 | transposon protein, putative, unclassified, expressed |
| LOC_Os02g06850 | expressed protein |
| LOC_Os02g06860 | OsMADS80 - MADS-box family gene with M-alpha type-box, expressed |
| LOC_Os02g07360 | pentatricopeptide containing protein, putative, expressed |
| LOC_Os02g07410 | glycine cleavage system H protein, putative, expressed |
| LOC_Os02g07420 | splicing factor, arginine/serine-rich 12, putative, expressed |
| LOC_Os02g07430 | OsMADS29 - MADS-box family gene with MIKCc type-box, expressed |
| LOC_Os02g07440 | DEFL78 - Defensin and Defensin-like DEFL family, expressed |
| LOC_Os02g07450 | core histone H2A/H2B/H3/H4, putative, expressed |
| LOC_Os02g07460 | expressed protein |
| LOC_Os02g07480 | transglycosylase SLT domain containing protein, expressed |
| LOC_Os02g07490 | glyceraldehyde-3-phosphate dehydrogenase, putative, expressed |
| LOC_Os02g07495 | expressed protein |
| LOC_Os02g07790 | serine/threonine-protein kinase HT1, putative, expressed |
| LOC_Os02g07800 | transcription factor, putative, expressed |
| LOC_Os02g07810 | inactive receptor kinase At2g26730 precursor, putative, expressed |
| LOC_Os02g07820 | DUF581 domain containing protein, expressed |
| LOC_Os02g07840 | bZIP transcription factor domain containing protein, expressed |
| LOC_Os02g07850 | DUF630/DUF632 domains containing protein, putative, expressed |
| LOC_Os02g08160 | OsFBD6 - F-box and FBD domain containing protein, expressed |
| LOC_Os02g08170 | GDU1, putative, expressed |
| LOC_Os02g08180 | protein transport protein SEC61 subunit gamma, putative, expressed |
| LOC_Os02g08190 | CYPRO4, putative, expressed |
| LOC_Os02g08200 | zinc finger, C3HC4 type domain containing protein, expressed |
| LOC_Os02g08210 | expressed protein |
| LOC_Os02g08220 | expressed protein |
| LOC_Os02g08230 | WAX2, putative, expressed |
| LOC_Os02g08240 | STE_PAK_Ste20_STLK |

| | |
|---|---|
| LOC_Os02g08260 | HD domain containing protein 2, putative, expressed |
| LOC_Os02g08270 | class I glutamine amidotransferase, putative, expressed |
| LOC_Os02g08300 | RAD23 DNA repair protein, putative, expressed |
| LOC_Os02g08310 | tubby, putative, expressed |
| LOC_Os02g08360 | RNA recognition motif, putative, expressed |
| LOC_Os02g08370 | ubiquitin carboxyl-terminal hydrolase, family 1, putative, expressed |
| LOC_Os02g08380 | CR084 protein, putative, expressed |
| LOC_Os02g08540 | expressed protein |
| LOC_Os02g08544 | homeobox protein knotted-1, putative, expressed |
| LOC_Os02g09050 | expressed protein |
| LOC_Os02g09060 | BRCA1-associated protein, putative, expressed |
| LOC_Os02g09070 | zinc-binding protein, putative, expressed |
| LOC_Os02g09080 | expressed protein |
| LOC_Os02g09090 | expressed protein |
| LOC_Os02g09100 | isoleucyl-tRNA synthetase, cytoplasmic, putative, expressed |
| LOC_Os02g10310 | fumarylacetoacetase, putative, expressed |
| LOC_Os02g10350 | MLO domain containing protein, putative, expressed |
| LOC_Os02g10360 | MONOCULM 1, putative, expressed |
| LOC_Os02g10380 | Regulator of chromosome condensation domain containing protein, expressed |
| LOC_Os02g10390 | chlorophyll A-B binding protein, putative, expressed |
| LOC_Os02g10400 | expressed protein |
| LOC_Os02g10410 | expressed protein |
| LOC_Os02g10420 | expressed protein |
| LOC_Os02g12654 | expressed protein |
| LOC_Os02g13510 | receptor-like protein kinase 5 precursor, putative, expressed |
| LOC_Os02g16050 | transposon protein, putative, unclassified, expressed |
| LOC_Os02g18910 | hypothetical protein |
| LOC_Os02g33900 | transposon protein, putative, unclassified, expressed |
| LOC_Os02g35740 | expressed protein |
| LOC_Os02g43800 | retrotransposon protein, putative, Ty3-gypsy subclass, expressed |

| | |
|---|---|
| LOC_Os02g44704 | LYR motif containing protein, putative, expressed |
| LOC_Os02g45300 | hypothetical protein |
| LOC_Os02g49460 | auxin-independent growth promoter protein, putative, expressed |
| LOC_Os02g52660 | expressed protein |
| LOC_Os02g53000 | lysM domain-containing GPI-anchored protein precursor, putative, expressed |
| LOC_Os02g53310 | expressed protein |
| LOC_Os03g01100 | endonuclease, putative, expressed |
| LOC_Os03g01110 | RNA methyltransferase, TrmH family protein, putative, expressed |
| LOC_Os03g01120 | E1-E2 ATPase domain containing protein, expressed |
| LOC_Os03g01130 | ethylene-responsive element-binding protein, putative, expressed |
| LOC_Os03g01150 | palmitoyl-protein thioesterase 1 precursor, putative, expressed |
| LOC_Os03g01190 | dehydrogenase, putative, expressed |
| LOC_Os03g01216 | phosphatidylserine decarboxylase, putative, expressed |
| LOC_Os03g01222 | formyl transferase, putative, expressed |
| LOC_Os03g01240 | CCR4-Not complex component, Not1domain containing protein, expressed |
| LOC_Os03g01280 | expressed protein |
| LOC_Os03g01320 | LTPL116 - Protease inhibitor/seed storage/LTP family protein precursor, expressed |
| LOC_Os03g01330 | sodium/potassium/calcium exchanger 1, putative, expressed |
| LOC_Os03g01360 | expressed protein |
| LOC_Os03g01420 | expressed protein |
| LOC_Os03g01430 | expressed protein |
| LOC_Os03g01436 | expressed protein |
| LOC_Os03g01450 | expressed protein |
| LOC_Os03g01500 | hypothetical protein |
| LOC_Os03g01520 | expressed protein |
| LOC_Os03g01540 | DNA binding protein, putative, expressed |
| LOC_Os03g01550 | expressed protein |
| LOC_Os03g01560 | expressed protein |
| LOC_Os03g01580 | expressed protein |
| LOC_Os03g01590 | PAIR1, putative, expressed |

| | |
|---|---|
| LOC_Os03g01610 | expansin precursor, putative, expressed |
| LOC_Os03g02380 | major facilitator superfamily domain-containing protein 5, putative, expressed |
| LOC_Os03g02390 | mitochondrial import inner membrane translocase subunit Tim17, putative, expressed |
| LOC_Os03g02400 | plastocyanin-like domain containing protein, putative, expressed |
| LOC_Os03g03980 | coiled-coil domain-containing protein 124, putative, expressed |
| LOC_Os03g03990 | signal recognition particle 43 kDa protein, chloroplast precursor, putative, expressed |
| LOC_Os03g04000 | AMP-binding domain containing protein, expressed |
| LOC_Os03g04010 | hypothetical protein |
| LOC_Os03g04080 | expressed protein |
| LOC_Os03g04100 | expressed protein |
| LOC_Os03g04110 | lysM domain-containing GPI-anchored protein precursor, putative, expressed |
| LOC_Os03g04169 | ATP phosphoribosyltransferase, putative, expressed |
| LOC_Os03g04190 | cytochrome P450, putative, expressed |
| LOC_Os03g04210 | IQ calmodulin-binding motif family protein, putative, expressed |
| LOC_Os03g04220 | glutathione S-transferase, putative, expressed |
| LOC_Os03g04270 | OsFBL10 - F-box domain and LRR containing protein, expressed |
| LOC_Os03g04300 | ankyrin repeat domain containing protein, expressed |
| LOC_Os03g04310 | BHLH transcription factor, putative, expressed |
| LOC_Os03g04320 | expressed protein |
| LOC_Os03g04330 | expressed protein |
| LOC_Os03g04340 | C-methyltransferase, putative, expressed |
| LOC_Os03g04350 | expressed protein |
| LOC_Os03g04370 | protein-methionine-S-oxide reductase, putative, expressed |
| LOC_Os03g04400 | heat shock protein DnaJ, putative, expressed |
| LOC_Os03g04410 | aconitate hydratase protein, putative, expressed |
| LOC_Os03g04420 | expressed protein |
| LOC_Os03g04430 | protein phosphatase 2C, putative, expressed |
| LOC_Os03g04440 | expressed protein |
| LOC_Os03g04450 | expressed protein |
| LOC_Os03g04470 | expressed protein |

| | |
|---|---|
| LOC_Os03g04480 | CorA-like magnesium transporter protein, putative, expressed |
| LOC_Os03g04490 | cyclin-dependent kinase inhibitor, putative, expressed |
| LOC_Os03g04500 | tetratricopeptide repeat domain containing protein, expressed |
| LOC_Os03g04530 | cytochrome P450, putative, expressed |
| LOC_Os03g04540 | expressed protein |
| LOC_Os03g04545 | expressed protein |
| LOC_Os03g04550 | expressed protein |
| LOC_Os03g05120 | expressed protein |
| LOC_Os03g05130 | expressed protein |
| LOC_Os03g05150 | hypothetical protein |
| LOC_Os03g05160 | GATA zinc finger domain containing protein, expressed |
| LOC_Os03g05170 | expressed protein |
| LOC_Os03g05180 | expressed protein |
| LOC_Os03g05200 | DENN domain containing protein, expressed |
| LOC_Os03g05210 | WD domain, G-beta repeat domain containing protein, expressed |
| LOC_Os03g05270 | RING finger and CHY zinc finger domain-containing protein 1, putative, expressed |
| LOC_Os03g05280 | ras-related protein, putative, expressed |
| LOC_Os03g05300 | cyclin-dependent kinases regulatory subunit 1, putative, expressed |
| LOC_Os03g05310 | pheophorbide a oxygenase, chloroplast precursor, putative, expressed |
| LOC_Os03g05320 | expressed protein |
| LOC_Os03g05330 | HEAT repeat family protein, putative, expressed |
| LOC_Os03g05334 | expressed protein |
| LOC_Os03g05340 | retrotransposon protein, putative, unclassified, expressed |
| LOC_Os03g06240 | YT521-B-like family domain containing protein, expressed |
| LOC_Os03g06340 | XS domain containing protein, expressed |
| LOC_Os03g06350 | DUF623 domain containing protein, expressed |
| LOC_Os03g06360 | late embryogenesis abundant protein D-34, putative, expressed |
| LOC_Os03g06370 | PPR repeat domain containing protein, putative, expressed |
| LOC_Os03g06390 | expressed protein |
| LOC_Os03g06410 | protein kinase domain containing protein, expressed |

| | |
|---|---|
| LOC_Os03g06420 | expressed protein |
| LOC_Os03g06430 | expressed protein |
| LOC_Os03g06440 | RNA helicase SDE3, putative, expressed |
| LOC_Os03g06490 | expressed protein |
| LOC_Os03g06510 | KIP1, putative, expressed |
| LOC_Os03g07160 | OsFBK11 - F-box domain and kelch repeat containing protein, expressed |
| LOC_Os03g07170 | lactose permease-related, putative, expressed |
| LOC_Os03g07190 | expressed protein |
| LOC_Os03g07220 | PPR repeat domain containing protein, putative, expressed |
| LOC_Os03g07226 | thioredoxin, putative, expressed |
| LOC_Os03g07234 | pentatricopeptide, putative, expressed |
| LOC_Os03g07250 | cytochrome P450, putative, expressed |
| LOC_Os03g07260 | tetratricopeptide repeat domain containing protein, expressed |
| LOC_Os03g07290 | structural constituent of ribosome, putative, expressed |
| LOC_Os03g07300 | ribulose-phosphate 3-epimerase, chloroplast precursor, putative, expressed |
| LOC_Os03g07330 | pentatricopeptide, putative, expressed |
| LOC_Os03g07340 | ribosomal L18p/L5e family protein, putative, expressed |
| LOC_Os03g08380 | ABC transporter, ATP-binding protein, putative, expressed |
| LOC_Os03g08390 | expressed protein |
| LOC_Os03g10440 | glycosyl hydrolase family 10 protein, putative, expressed |
| LOC_Os03g10500 | adenylyl-sulfate kinase, putative, expressed |
| LOC_Os03g10510 | outer mitochondrial membrane porin, putative, expressed |
| LOC_Os03g10520 | radical SAM enzyme, putative, expressed |
| LOC_Os03g10530 | expressed protein |
| LOC_Os03g10820 | expressed protein |
| LOC_Os03g10830 | hypothetical protein |
| LOC_Os03g10840 | expressed protein |
| LOC_Os03g11600 | YABBY domain containing protein, putative, expressed |
| LOC_Os03g11640 | hypothetical protein |
| LOC_Os03g11650 | ATMAP70 protein, putative, expressed |

| | |
|---|---|
| LOC_Os03g11660 | cysteine synthase, putative, expressed |
| LOC_Os03g11670 | pentatricopeptide repeat domain containing protein, putative, expressed |
| LOC_Os03g11790 | OsFBX79 - F-box domain containing protein, expressed |
| LOC_Os03g11890 | potyvirus VPg interacting protein, putative, expressed |
| LOC_Os03g11910 | DnaK family protein, putative, expressed |
| LOC_Os03g11930 | expressed protein |
| LOC_Os03g11950 | SEC14 cytosolic factor family protein, putative, expressed |
| LOC_Os03g12610 | DNA polymerase I, putative, expressed |
| LOC_Os03g12620 | glycosyl hydrolases family 17, putative, expressed |
| LOC_Os03g36320 | expressed protein |
| LOC_Os03g42235 | expressed protein |
| LOC_Os03g52475 | pentatricopeptide, putative, expressed |
| LOC_Os03g56100 | expressed protein |
| LOC_Os03g60250 | membrane associated DUF588 domain containing protein, putative, expressed |
| LOC_Os04g25784 | expressed protein |
| LOC_Os04g38990 | expressed protein |
| LOC_Os04g44020 | expressed protein |
| LOC_Os04g44690 | expressed protein |
| LOC_Os05g01710 | transcription initiation factor IIA gamma chain, putative, expressed |
| LOC_Os05g01730 | drought induced 19 protein, putative, expressed |
| LOC_Os05g01750 | TruB family pseudouridylate synthase, putative, expressed |
| LOC_Os05g03050 | DNA repair protein Rad51, putative, expressed |
| LOC_Os05g03060 | expressed protein |
| LOC_Os05g03070 | transporter, putative, expressed |
| LOC_Os05g03100 | HECT-domain domain containing protein, expressed |
| LOC_Os05g05120 | hypothetical protein |
| LOC_Os05g05130 | GA20663-PA, putative, expressed |
| LOC_Os05g05140 | expressed protein |
| LOC_Os05g05150 | SLD5 - Putative GINS complex subunit, expressed |
| LOC_Os05g05160 | CGMC_MAPKCGMC_2 |

| | |
|---|---|
| LOC_Os05g05170 | jacalin-like lectin domain containing protein, expressed |
| LOC_Os05g05180 | OsGH3 |
| LOC_Os05g05200 | O-acyltransferase, putative, expressed |
| LOC_Os05g05220 | glioma tumor suppressor candidate region gene 2 protein, putative, expressed |
| LOC_Os05g05230 | expressed protein |
| LOC_Os05g05240 | serine/threonine protein phosphatase, putative, expressed |
| LOC_Os05g05760 | expressed protein |
| LOC_Os05g05790 | double-stranded RNA binding motif containing protein, expressed |
| LOC_Os05g05810 | DEAD-box ATP-dependent RNA helicase, putative, expressed |
| LOC_Os05g05820 | pro-resilin precursor, putative, expressed |
| LOC_Os05g05830 | bifunctional protein folD, putative, expressed |
| LOC_Os05g05840 | tRNA synthetase class II core domain containing protein, expressed |
| LOC_Os05g05850 | expressed protein |
| LOC_Os05g05860 | retrotransposon protein, putative, unclassified, expressed |
| LOC_Os05g10290 | expressed protein |
| LOC_Os06g27510 | hypothetical protein |
| LOC_Os06g39515 | expressed protein |
| LOC_Os06g51180 | lysine ketoglutarate reductase trans-splicing related 1, putative, expressed |
| LOC_Os07g10990 | hypothetical protein |
| LOC_Os07g25880 | hypothetical protein |
| LOC_Os07g42560 | expressed protein |
| LOC_Os08g01040 | zinc finger, C3HC4 type domain containing protein, expressed |
| LOC_Os08g02350 | expressed protein |
| LOC_Os08g20020 | octicosapeptide/Phox/Bem1p, putative, expressed |
| LOC_Os09g16580 | amidase family protein, putative, expressed |
| LOC_Os10g36984 | expressed protein |
| LOC_Os11g38840 | expressed protein |
| LOC_Os11g39160 | NBS-LRR disease resistance protein, putative, expressed |
| LOC_Os12g07160 | plant protein of unknown function DUF869 domain containing protein, expressed |
| LOC_Os12g19304 | Fe-S metabolism associated domain containing protein, expressed |

**Table S9.** GO analysis for genes from potentially selective regions (top 1% CLR) identified by SweeD.

| GO term | Ontology | Description | Number in input list | Number in BG/Ref | p-value |
|---|---|---|---|---|---|
| **Genes in indica_SNP CLR** | | | | | |
| GO:0006468 | Biological Process | protein amino acid phosphorylation | 24 | 1570 | 0.0026 |
| GO:0044267 | Biological Process | cellular protein metabolic process | 37 | 2877 | 0.0037 |
| GO:0043687 | Biological Process | post-translational protein modification | 26 | 1815 | 0.0041 |
| GO:0043412 | Biological Process | macromolecule modification | 27 | 1979 | 0.0064 |
| GO:0016310 | Biological Process | phosphorylation | 24 | 1695 | 0.0066 |
| GO:0006464 | Biological Process | protein modification process | 26 | 1929 | 0.0087 |
| GO:0006820 | Biological Process | anion transport | 3 | 55 | 0.012 |
| GO:0006796 | Biological Process | phosphate metabolic process | 24 | 1795 | 0.013 |
| GO:0006793 | Biological Process | phosphorus metabolic process | 24 | 1795 | 0.013 |
| GO:0009611 | Biological Process | response to wounding | 2 | 21 | 0.015 |
| GO:0033615 | Biological Process | mitochondrial proton-transporting ATP synthase complex assembly | 1 | 1 | 0.016 |
| GO:0009605 | Biological Process | response to external stimulus | 2 | 25 | 0.02 |
| GO:0007131 | Biological Process | reciprocal meiotic recombination | 1 | 2 | 0.024 |
| GO:0070071 | Biological Process | proton-transporting two-sector ATPase complex assembly | 1 | 2 | 0.024 |
| GO:0009306 | Biological Process | protein secretion | 1 | 2 | 0.024 |
| GO:0006890 | Biological Process | retrograde vesicle-mediated transport, Golgi to ER | 1 | 2 | 0.024 |
| GO:0070272 | Biological Process | proton-transporting ATP synthase complex biogenesis | 1 | 2 | 0.024 |
| GO:0007127 | Biological Process | meiosis I | 1 | 2 | 0.024 |
| GO:0043461 | Biological Process | proton-transporting ATP synthase complex assembly | 1 | 2 | 0.024 |
| GO:0009755 | Biological Process | hormone-mediated signaling pathway | 1 | 3 | 0.032 |
| GO:0051327 | Biological Process | M phase of meiotic cell cycle | 1 | 3 | 0.032 |
| GO:0007126 | Biological Process | meiosis | 1 | 3 | 0.032 |
| GO:0051321 | Biological Process | meiotic cell cycle | 1 | 3 | 0.032 |
| GO:0032870 | Biological Process | cellular response to hormone stimulus | 1 | 3 | 0.032 |

| | | | | | |
|---|---|---|---|---|---|
| GO:0071369 | Biological Process | cellular response to ethylene stimulus | 1 | 3 | 0.032 |
| GO:0009723 | Biological Process | response to ethylene stimulus | 1 | 3 | 0.032 |
| GO:0071495 | Biological Process | cellular response to endogenous stimulus | 1 | 3 | 0.032 |
| GO:0071310 | Biological Process | cellular response to organic substance | 1 | 3 | 0.032 |
| GO:0009873 | Biological Process | ethylene mediated signaling pathway | 1 | 3 | 0.032 |
| GO:0006438 | Biological Process | valyl-tRNA aminoacylation | 1 | 5 | 0.048 |
| GO:0007165 | Biological Process | signal transduction | 5 | 240 | 0.05 |
| GO:0022836 | Molecular Function | gated channel activity | 4 | 59 | 0.0017 |
| GO:0004672 | Molecular Function | protein kinase activity | 24 | 1575 | 0.0027 |
| GO:0015267 | Molecular Function | channel activity | 4 | 69 | 0.003 |
| GO:0005216 | Molecular Function | ion channel activity | 4 | 69 | 0.003 |
| GO:0022838 | Molecular Function | substrate-specific channel activity | 4 | 69 | 0.003 |
| GO:0022803 | Molecular Function | passive transmembrane transporter activity | 4 | 69 | 0.003 |
| GO:0004866 | Molecular Function | endopeptidase inhibitor activity | 4 | 72 | 0.0034 |
| GO:0030414 | Molecular Function | peptidase inhibitor activity | 4 | 72 | 0.0034 |
| GO:0005524 | Molecular Function | ATP binding | 39 | 3085 | 0.0039 |
| GO:0032559 | Molecular Function | adenyl ribonucleotide binding | 39 | 3087 | 0.0039 |
| GO:0032555 | Molecular Function | purine ribonucleotide binding | 41 | 3292 | 0.0039 |
| GO:0032553 | Molecular Function | ribonucleotide binding | 41 | 3292 | 0.0039 |
| GO:0016773 | Molecular Function | phosphotransferase activity, alcohol group as acceptor | 25 | 1726 | 0.0042 |
| GO:0022832 | Molecular Function | voltage-gated channel activity | 3 | 38 | 0.0045 |
| GO:0005244 | Molecular Function | voltage-gated ion channel activity | 3 | 38 | 0.0045 |
| GO:0004869 | Molecular Function | cysteine-type endopeptidase inhibitor activity | 2 | 11 | 0.0048 |
| GO:0016301 | Molecular Function | kinase activity | 25 | 1750 | 0.005 |
| GO:0004674 | Molecular Function | protein serine/threonine kinase activity | 22 | 1478 | 0.0053 |
| GO:0000166 | Molecular Function | nucleotide binding | 46 | 3874 | 0.0054 |
| GO:0001882 | Molecular Function | nucleoside binding | 40 | 3257 | 0.0056 |
| GO:0008509 | Molecular Function | anion transmembrane transporter activity | 3 | 44 | 0.0066 |
| GO:0017076 | Molecular Function | purine nucleotide binding | 41 | 3456 | 0.0089 |
| GO:0001883 | Molecular Function | purine nucleoside binding | 39 | 3247 | 0.0089 |

| | | | | | |
|---|---|---|---|---|---|
| GO:0030554 | Molecular Function | adenyl nucleotide binding | 39 | 3247 | 0.0089 |
| GO:0005253 | Molecular Function | anion channel activity | 2 | 18 | 0.011 |
| GO:0008308 | Molecular Function | voltage-gated anion channel activity | 2 | 18 | 0.011 |
| GO:0070008 | Molecular Function | serine-type exopeptidase activity | 3 | 61 | 0.015 |
| GO:0004185 | Molecular Function | serine-type carboxypeptidase activity | 3 | 61 | 0.015 |
| GO:0004315 | Molecular Function | 3-oxoacyl-[acyl-carrier-protein] synthase activity | 1 | 1 | 0.016 |
| GO:0051020 | Molecular Function | GTPase binding | 1 | 1 | 0.016 |
| GO:0004180 | Molecular Function | carboxypeptidase activity | 3 | 63 | 0.017 |
| GO:0016667 | Molecular Function | oxidoreductase activity, acting on sulfur group of donors | 3 | 75 | 0.026 |
| GO:0000287 | Molecular Function | magnesium ion binding | 4 | 134 | 0.026 |
| GO:0008238 | Molecular Function | exopeptidase activity | 3 | 82 | 0.032 |
| GO:0015380 | Molecular Function | anion exchanger activity | 1 | 3 | 0.032 |
| GO:0005452 | Molecular Function | inorganic anion exchanger activity | 1 | 3 | 0.032 |
| GO:0015301 | Molecular Function | anion:anion antiporter activity | 1 | 3 | 0.032 |
| GO:0004617 | Molecular Function | phosphoglycerate dehydrogenase activity | 1 | 3 | 0.032 |
| GO:0004619 | Molecular Function | phosphoglycerate mutase activity | 1 | 3 | 0.032 |
| GO:0005198 | Molecular Function | structural molecule activity | 8 | 462 | 0.039 |
| GO:0004096 | Molecular Function | catalase activity | 1 | 4 | 0.04 |
| GO:0004857 | Molecular Function | enzyme inhibitor activity | 4 | 157 | 0.043 |
| GO:0015075 | Molecular Function | ion transmembrane transporter activity | 7 | 389 | 0.043 |
| GO:0004351 | Molecular Function | glutamate decarboxylase activity | 1 | 5 | 0.048 |
| GO:0004832 | Molecular Function | valine-tRNA ligase activity | 1 | 5 | 0.048 |
| GO:0030660 | Cellular Component | Golgi-associated vesicle membrane | 2 | 15 | 0.0082 |
| GO:0005798 | Cellular Component | Golgi-associated vesicle | 2 | 15 | 0.0082 |
| GO:0030662 | Cellular Component | coated vesicle membrane | 2 | 25 | 0.02 |
| GO:0012506 | Cellular Component | vesicle membrane | 2 | 25 | 0.02 |
| GO:0030120 | Cellular Component | vesicle coat | 2 | 25 | 0.02 |
| GO:0030135 | Cellular Component | coated vesicle | 2 | 25 | 0.02 |
| GO:0030659 | Cellular Component | cytoplasmic vesicle membrane | 2 | 25 | 0.02 |

| | | | | | |
|---|---|---|---|---|---|
| GO:0016023 | Cellular Component | cytoplasmic membrane-bounded vesicle | 2 | 26 | 0.022 |
| GO:0031988 | Cellular Component | membrane-bounded vesicle | 2 | 26 | 0.022 |
| GO:0009986 | Cellular Component | cell surface | 1 | 2 | 0.024 |
| GO:0000139 | Cellular Component | Golgi membrane | 2 | 29 | 0.026 |
| GO:0044433 | Cellular Component | cytoplasmic vesicle part | 2 | 33 | 0.033 |
| GO:0031982 | Cellular Component | vesicle | 2 | 34 | 0.034 |
| GO:0031410 | Cellular Component | cytoplasmic vesicle | 2 | 34 | 0.034 |
| GO:0031090 | Cellular Component | organelle membrane | 5 | 228 | 0.042 |
| GO:0005905 | Cellular Component | coated pit | 1 | 5 | 0.048 |
| GO:0030132 | Cellular Component | clathrin coat of coated pit | 1 | 5 | 0.048 |

**Genes in indica_SV CLR**

| | | | | | |
|---|---|---|---|---|---|
| GO:0030488 | Biological Process | tRNA methylation | 1 | 3 | 0.014 |
| GO:0055072 | Biological Process | iron ion homeostasis | 1 | 4 | 0.017 |
| GO:0009098 | Biological Process | leucine biosynthetic process | 1 | 4 | 0.017 |
| GO:0006879 | Biological Process | cellular iron ion homeostasis | 1 | 4 | 0.017 |
| GO:0006551 | Biological Process | leucine metabolic process | 1 | 4 | 0.017 |
| GO:0030005 | Biological Process | cellular di-, tri-valent inorganic cation homeostasis | 1 | 5 | 0.021 |
| GO:0055066 | Biological Process | di-, tri-valent inorganic cation homeostasis | 1 | 5 | 0.021 |
| GO:0006826 | Biological Process | iron ion transport | 1 | 7 | 0.028 |
| GO:0006873 | Biological Process | cellular ion homeostasis | 1 | 7 | 0.028 |
| GO:0030003 | Biological Process | cellular cation homeostasis | 1 | 7 | 0.028 |
| GO:0055082 | Biological Process | cellular chemical homeostasis | 1 | 7 | 0.028 |
| GO:0006771 | Biological Process | riboflavin metabolic process | 1 | 9 | 0.035 |
| GO:0001510 | Biological Process | RNA methylation | 1 | 9 | 0.035 |
| GO:0042727 | Biological Process | riboflavin and derivative biosynthetic process | 1 | 9 | 0.035 |
| GO:0042726 | Biological Process | riboflavin and derivative metabolic process | 1 | 9 | 0.035 |
| GO:0009231 | Biological Process | riboflavin biosynthetic process | 1 | 9 | 0.035 |
| GO:0015914 | Biological Process | phospholipid transport | 1 | 10 | 0.038 |
| GO:0006400 | Biological Process | tRNA modification | 1 | 11 | 0.041 |

| | | | | | |
|---|---|---|---|---|---|
| GO:0050801 | Biological Process | ion homeostasis | 1 | 13 | 0.048 |
| GO:0048878 | Biological Process | chemical homeostasis | 1 | 13 | 0.048 |
| GO:0055080 | Biological Process | cation homeostasis | 1 | 13 | 0.048 |
| GO:0009082 | Biological Process | branched chain family amino acid biosynthetic process | 1 | 13 | 0.048 |
| GO:0003852 | Molecular Function | 2-isopropylmalate synthase activity | 1 | 2 | 0.011 |
| GO:0008531 | Molecular Function | riboflavin kinase activity | 1 | 2 | 0.011 |
| GO:0046872 | Molecular Function | metal ion binding | 29 | 5476 | 0.011 |
| GO:0008199 | Molecular Function | ferric iron binding | 1 | 3 | 0.014 |
| GO:0016429 | Molecular Function | tRNA (adenine-N1-)-methyltransferase activity | 1 | 3 | 0.014 |
| GO:0016426 | Molecular Function | tRNA (adenine)-methyltransferase activity | 1 | 3 | 0.014 |
| GO:0046914 | Molecular Function | transition metal ion binding | 26 | 4875 | 0.016 |
| GO:0008097 | Molecular Function | 5S rRNA binding | 1 | 4 | 0.017 |
| GO:0043169 | Molecular Function | cation binding | 29 | 5696 | 0.019 |
| GO:0043167 | Molecular Function | ion binding | 29 | 5700 | 0.019 |
| GO:0015662 | Molecular Function | ATPase activity, coupled to transmembrane movement of ions, phosphorylative mechanism | 2 | 67 | 0.025 |
| GO:0004470 | Molecular Function | malic enzyme activity | 1 | 7 | 0.028 |
| GO:0008270 | Molecular Function | zinc ion binding | 21 | 3939 | 0.032 |
| GO:0008553 | Molecular Function | hydrogen-exporting ATPase activity, phosphorylative mechanism | 1 | 9 | 0.035 |
| GO:0016671 | Molecular Function | oxidoreductase activity, acting on sulfur group of donors, disulfide as acceptor | 1 | 9 | 0.035 |
| GO:0016887 | Molecular Function | ATPase activity | 4 | 355 | 0.038 |
| GO:0004012 | Molecular Function | phospholipid-translocating ATPase activity | 1 | 10 | 0.038 |
| GO:0008175 | Molecular Function | tRNA methyltransferase activity | 1 | 10 | 0.038 |
| GO:0005548 | Molecular Function | phospholipid transporter activity | 1 | 10 | 0.038 |
| GO:0015247 | Molecular Function | aminophospholipid transporter activity | 1 | 10 | 0.038 |
| GO:0017111 | Molecular Function | nucleoside-triphosphatase activity | 6 | 726 | 0.044 |
| GO:0046912 | Molecular Function | transferase activity, transferring acyl groups, acyl groups converted into alkyl on transfer | 1 | 12 | 0.045 |

| | | | | | |
|---|---|---|---|---|---|
| GO:0042625 | Molecular Function | ATPase activity, coupled to transmembrane movement of ions | 2 | 94 | 0.045 |
| GO:0004525 | Molecular Function | ribonuclease III activity | 1 | 13 | 0.048 |
| GO:0016462 | Molecular Function | pyrophosphatase activity | 6 | 743 | 0.049 |
| GO:0031227 | Cellular Component | intrinsic to endoplasmic reticulum membrane | 2 | 23 | 0.0035 |
| GO:0031300 | Cellular Component | intrinsic to organelle membrane | 2 | 37 | 0.0083 |
| GO:0042175 | Cellular Component | nuclear envelope-endoplasmic reticulum network | 2 | 47 | 0.013 |
| GO:0005789 | Cellular Component | endoplasmic reticulum membrane | 2 | 47 | 0.013 |
| GO:0044432 | Cellular Component | endoplasmic reticulum part | 2 | 49 | 0.014 |
| GO:0071212 | Cellular Component | subsynaptic reticulum | 2 | 49 | 0.014 |
| GO:0030176 | Cellular Component | integral to endoplasmic reticulum membrane | 1 | 3 | 0.014 |
| GO:0008076 | Cellular Component | voltage-gated potassium channel complex | 1 | 5 | 0.021 |
| GO:0034702 | Cellular Component | ion channel complex | 1 | 5 | 0.021 |
| GO:0034703 | Cellular Component | cation channel complex | 1 | 5 | 0.021 |
| GO:0034705 | Cellular Component | potassium channel complex | 1 | 5 | 0.021 |
| GO:0031226 | Cellular Component | intrinsic to plasma membrane | 1 | 10 | 0.038 |
| GO:0005887 | Cellular Component | integral to plasma membrane | 1 | 10 | 0.038 |
| **Genes in japonica_SNP CLR** | | | | | |
| GO:0022900 | Biological Process | electron transport chain | 5 | 60 | 0.00017 |
| GO:0022904 | Biological Process | respiratory electron transport chain | 3 | 28 | 0.0019 |
| GO:0055114 | Biological Process | oxidation reduction | 21 | 1351 | 0.0033 |
| GO:0019538 | Biological Process | protein metabolic process | 50 | 4333 | 0.0046 |
| GO:0016310 | Biological Process | phosphorylation | 24 | 1695 | 0.0055 |
| GO:0045333 | Biological Process | cellular respiration | 3 | 48 | 0.0079 |
| GO:0006468 | Biological Process | protein amino acid phosphorylation | 22 | 1570 | 0.0087 |
| GO:0006793 | Biological Process | phosphorus metabolic process | 24 | 1795 | 0.011 |
| GO:0006796 | Biological Process | phosphate metabolic process | 24 | 1795 | 0.011 |
| GO:0015980 | Biological Process | energy derivation by oxidation of organic compounds | 3 | 54 | 0.011 |
| GO:0000279 | Biological Process | M phase | 2 | 19 | 0.012 |

| | | | | | |
|---|---|---|---|---|---|
| GO:0022403 | Biological Process | cell cycle phase | 2 | 20 | 0.013 |
| GO:0009808 | Biological Process | lignin metabolic process | 2 | 21 | 0.014 |
| GO:0046274 | Biological Process | lignin catabolic process | 2 | 21 | 0.014 |
| GO:0046271 | Biological Process | phenylpropanoid catabolic process | 2 | 21 | 0.014 |
| GO:0042219 | Biological Process | cellular amino acid derivative catabolic process | 2 | 21 | 0.014 |
| GO:0006091 | Biological Process | generation of precursor metabolites and energy | 6 | 243 | 0.015 |
| GO:0019748 | Biological Process | secondary metabolic process | 3 | 64 | 0.017 |
| GO:0009698 | Biological Process | phenylpropanoid metabolic process | 2 | 24 | 0.018 |
| GO:0016192 | Biological Process | vesicle-mediated transport | 5 | 186 | 0.019 |
| GO:0022402 | Biological Process | cell cycle process | 2 | 25 | 0.02 |
| GO:0009767 | Biological Process | photosynthetic electron transport chain | 2 | 26 | 0.021 |
| GO:0007131 | Biological Process | reciprocal meiotic recombination | 1 | 2 | 0.024 |
| GO:0007127 | Biological Process | meiosis I | 1 | 2 | 0.024 |
| GO:0044267 | Biological Process | cellular protein metabolic process | 33 | 2877 | 0.024 |
| GO:0007049 | Biological Process | cell cycle | 2 | 30 | 0.027 |
| GO:0019439 | Biological Process | aromatic compound catabolic process | 2 | 31 | 0.029 |
| GO:0043412 | Biological Process | macromolecule modification | 24 | 1979 | 0.03 |
| GO:0006426 | Biological Process | glycyl-tRNA aminoacylation | 1 | 3 | 0.032 |
| GO:0051327 | Biological Process | M phase of meiotic cell cycle | 1 | 3 | 0.032 |
| GO:0007126 | Biological Process | meiosis | 1 | 3 | 0.032 |
| GO:0051321 | Biological Process | meiotic cell cycle | 1 | 3 | 0.032 |
| GO:0043687 | Biological Process | post-translational protein modification | 22 | 1815 | 0.037 |
| GO:0006464 | Biological Process | protein modification process | 23 | 1929 | 0.039 |
| GO:0050821 | Biological Process | protein stabilization | 1 | 4 | 0.039 |
| GO:0009775 | Biological Process | photosynthetic electron transport in cytochrome b6/f | 1 | 4 | 0.039 |
| GO:0006102 | Biological Process | isocitrate metabolic process | 1 | 4 | 0.039 |
| GO:0031647 | Biological Process | regulation of protein stability | 1 | 4 | 0.039 |
| GO:0044255 | Biological Process | cellular lipid metabolic process | 5 | 229 | 0.04 |
| GO:0046039 | Biological Process | GTP metabolic process | 1 | 5 | 0.047 |
| GO:0046036 | Biological Process | CTP metabolic process | 1 | 5 | 0.047 |

| | | | | | |
|---|---|---|---|---|---|
| GO:0009209 | Biological Process | pyrimidine ribonucleoside triphosphate biosynthetic process | 1 | 5 | 0.047 |
| GO:0009208 | Biological Process | pyrimidine ribonucleoside triphosphate metabolic process | 1 | 5 | 0.047 |
| GO:0009148 | Biological Process | pyrimidine nucleoside triphosphate biosynthetic process | 1 | 5 | 0.047 |
| GO:0006183 | Biological Process | GTP biosynthetic process | 1 | 5 | 0.047 |
| GO:0006228 | Biological Process | UTP biosynthetic process | 1 | 5 | 0.047 |
| GO:0006241 | Biological Process | CTP biosynthetic process | 1 | 5 | 0.047 |
| GO:0046051 | Biological Process | UTP metabolic process | 1 | 5 | 0.047 |
| GO:0016679 | Molecular Function | oxidoreductase activity, acting on diphenols and related substances as donors | 4 | 36 | 0.00029 |
| GO:0016681 | Molecular Function | oxidoreductase activity, acting on diphenols and related substances as donors, cytochrome as acceptor | 2 | 9 | 0.0033 |
| GO:0008121 | Molecular Function | ubiquinol-cytochrome-c reductase activity | 2 | 9 | 0.0033 |
| GO:0016301 | Molecular Function | kinase activity | 25 | 1750 | 0.0041 |
| GO:0004674 | Molecular Function | protein serine/threonine kinase activity | 22 | 1478 | 0.0044 |
| GO:0016491 | Molecular Function | oxidoreductase activity | 24 | 1672 | 0.0046 |
| GO:0008238 | Molecular Function | exopeptidase activity | 4 | 82 | 0.005 |
| GO:0005524 | Molecular Function | ATP binding | 38 | 3085 | 0.0052 |
| GO:0032559 | Molecular Function | adenyl ribonucleotide binding | 38 | 3087 | 0.0052 |
| GO:0004650 | Molecular Function | polygalacturonase activity | 3 | 45 | 0.0067 |
| GO:0016773 | Molecular Function | phosphotransferase activity, alcohol group as acceptor | 24 | 1726 | 0.0068 |
| GO:0008235 | Molecular Function | metalloexopeptidase activity | 2 | 14 | 0.0071 |
| GO:0004672 | Molecular Function | protein kinase activity | 22 | 1575 | 0.009 |
| GO:0001883 | Molecular Function | purine nucleoside binding | 38 | 3247 | 0.012 |
| GO:0030554 | Molecular Function | adenyl nucleotide binding | 38 | 3247 | 0.012 |
| GO:0001882 | Molecular Function | nucleoside binding | 38 | 3257 | 0.012 |
| GO:0032555 | Molecular Function | purine ribonucleotide binding | 38 | 3292 | 0.014 |
| GO:0032553 | Molecular Function | ribonucleotide binding | 38 | 3292 | 0.014 |

| | | | | | |
|---|---|---|---|---|---|
| GO:0008471 | Molecular Function | laccase activity | 2 | 21 | 0.014 |
| GO:0051537 | Molecular Function | 2 iron, 2 sulfur cluster binding | 2 | 22 | 0.016 |
| GO:0004180 | Molecular Function | carboxypeptidase activity | 3 | 63 | 0.016 |
| GO:0019136 | Molecular Function | deoxynucleoside kinase activity | 1 | 1 | 0.016 |
| GO:0004797 | Molecular Function | thymidine kinase activity | 1 | 1 | 0.016 |
| GO:0017169 | Molecular Function | CDP-alcohol phosphatidyltransferase activity | 1 | 1 | 0.016 |
| GO:0016719 | Molecular Function | carotene 7,8-desaturase activity | 1 | 1 | 0.016 |
| GO:0008444 | Molecular Function | CDP-diacylglycerol-glycerol-3-phosphate 3-phosphatidyltransferase activity | 1 | 1 | 0.016 |
| GO:0019205 | Molecular Function | nucleobase, nucleoside, nucleotide kinase activity | 2 | 25 | 0.02 |
| GO:0016682 | Molecular Function | oxidoreductase activity, acting on diphenols and related substances as donors, oxygen as acceptor | 2 | 27 | 0.022 |
| GO:0022892 | Molecular Function | substrate-specific transporter activity | 10 | 593 | 0.024 |
| GO:0004181 | Molecular Function | metallocarboxypeptidase activity | 1 | 2 | 0.024 |
| GO:0017056 | Molecular Function | structural constituent of nuclear pore | 1 | 2 | 0.024 |
| GO:0016667 | Molecular Function | oxidoreductase activity, acting on sulfur group of donors | 3 | 75 | 0.025 |
| GO:0017171 | Molecular Function | serine hydrolase activity | 5 | 201 | 0.025 |
| GO:0008236 | Molecular Function | serine-type peptidase activity | 5 | 201 | 0.025 |
| GO:0000166 | Molecular Function | nucleotide binding | 42 | 3874 | 0.026 |
| GO:0022891 | Molecular Function | substrate-specific transmembrane transporter activity | 9 | 517 | 0.026 |
| GO:0045735 | Molecular Function | nutrient reservoir activity | 3 | 77 | 0.026 |
| GO:0017076 | Molecular Function | purine nucleotide binding | 38 | 3456 | 0.028 |
| GO:0004326 | Molecular Function | tetrahydrofolylpolyglutamate synthase activity | 1 | 3 | 0.032 |
| GO:0004820 | Molecular Function | glycine-tRNA ligase activity | 1 | 3 | 0.032 |
| GO:0019206 | Molecular Function | nucleoside kinase activity | 1 | 3 | 0.032 |
| GO:0010181 | Molecular Function | FMN binding | 2 | 35 | 0.035 |
| GO:0005543 | Molecular Function | phospholipid binding | 2 | 36 | 0.037 |
| GO:0022857 | Molecular Function | transmembrane transporter activity | 10 | 643 | 0.038 |
| GO:0004096 | Molecular Function | catalase activity | 1 | 4 | 0.039 |

| | | | | | |
|---|---|---|---|---|---|
| GO:0004450 | Molecular Function | isocitrate dehydrogenase (NADP+) activity | 1 | 4 | 0.039 |
| GO:0043168 | Molecular Function | anion binding | 1 | 4 | 0.039 |
| GO:0042301 | Molecular Function | phosphate binding | 1 | 4 | 0.039 |
| GO:0030570 | Molecular Function | pectate lyase activity | 1 | 4 | 0.039 |
| GO:0016837 | Molecular Function | carbon-oxygen lyase activity, acting on polysaccharides | 1 | 4 | 0.039 |
| GO:0015078 | Molecular Function | hydrogen ion transmembrane transporter activity | 3 | 95 | 0.044 |
| GO:0004550 | Molecular Function | nucleoside diphosphate kinase activity | 1 | 5 | 0.047 |
| GO:0004386 | Molecular Function | helicase activity | 4 | 165 | 0.047 |
| GO:0009579 | Cellular Component | thylakoid | 4 | 93 | 0.0077 |
| GO:0030119 | Cellular Component | AP-type membrane coat adaptor complex | 2 | 16 | 0.0089 |
| GO:0030131 | Cellular Component | clathrin adaptor complex | 2 | 16 | 0.0089 |
| GO:0048475 | Cellular Component | coated membrane | 3 | 59 | 0.013 |
| GO:0030117 | Cellular Component | membrane coat | 3 | 59 | 0.013 |
| GO:0005956 | Cellular Component | protein kinase CK2 complex | 1 | 2 | 0.024 |
| GO:0030118 | Cellular Component | clathrin coat | 2 | 32 | 0.03 |
| GO:0044425 | Cellular Component | membrane part | 14 | 1007 | 0.035 |
| GO:0016020 | Cellular Component | membrane | 25 | 2115 | 0.035 |
| GO:0032991 | Cellular Component | macromolecular complex | 16 | 1207 | 0.037 |
| GO:0034357 | Cellular Component | photosynthetic membrane | 3 | 89 | 0.038 |
| **Genes in japonica_SV CLR** | | | | | |
| GO:0042773 | Biological Process | ATP synthesis coupled electron transport | 2 | 19 | 0.0026 |
| GO:0055114 | Biological Process | oxidation reduction | 12 | 1351 | 0.0038 |
| GO:0022904 | Biological Process | respiratory electron transport chain | 2 | 28 | 0.0053 |
| GO:0045333 | Biological Process | cellular respiration | 2 | 48 | 0.014 |
| GO:0006436 | Biological Process | tryptophanyl-tRNA aminoacylation | 1 | 3 | 0.014 |
| GO:0015980 | Biological Process | energy derivation by oxidation of organic compounds | 2 | 54 | 0.018 |
| GO:0055072 | Biological Process | iron ion homeostasis | 1 | 4 | 0.018 |
| GO:0006879 | Biological Process | cellular iron ion homeostasis | 1 | 4 | 0.018 |

| | | | | | |
|---|---|---|---|---|---|
| GO:0022900 | Biological Process | electron transport chain | 2 | 60 | 0.022 |
| GO:0030005 | Biological Process | cellular di-, tri-valent inorganic cation homeostasis | 1 | 5 | 0.022 |
| GO:0055066 | Biological Process | di-, tri-valent inorganic cation homeostasis | 1 | 5 | 0.022 |
| GO:0006952 | Biological Process | defense response | 5 | 452 | 0.026 |
| GO:0006826 | Biological Process | iron ion transport | 1 | 7 | 0.029 |
| GO:0006873 | Biological Process | cellular ion homeostasis | 1 | 7 | 0.029 |
| GO:0030003 | Biological Process | cellular cation homeostasis | 1 | 7 | 0.029 |
| GO:0055082 | Biological Process | cellular chemical homeostasis | 1 | 7 | 0.029 |
| GO:0006508 | Biological Process | proteolysis | 11 | 1579 | 0.029 |
| GO:0006119 | Biological Process | oxidative phosphorylation | 2 | 88 | 0.043 |
| GO:0048878 | Biological Process | chemical homeostasis | 1 | 13 | 0.05 |
| GO:0050801 | Biological Process | ion homeostasis | 1 | 13 | 0.05 |
| GO:0055080 | Biological Process | cation homeostasis | 1 | 13 | 0.05 |
| GO:0016651 | Molecular Function | oxidoreductase activity, acting on NADH or NADPH | 5 | 64 | 5.30E-06 |
| GO:0003954 | Molecular Function | NADH dehydrogenase activity | 3 | 30 | 0.00024 |
| GO:0050136 | Molecular Function | NADH dehydrogenase (quinone) activity | 3 | 30 | 0.00024 |
| GO:0008137 | Molecular Function | NADH dehydrogenase (ubiquinone) activity | 3 | 30 | 0.00024 |
| GO:0016655 | Molecular Function | oxidoreductase activity, acting on NADH or NADPH, quinone or similar compound as acceptor | 3 | 34 | 0.00033 |
| GO:0016491 | Molecular Function | oxidoreductase activity | 13 | 1672 | 0.0078 |
| GO:0004802 | Molecular Function | transketolase activity | 1 | 2 | 0.011 |
| GO:0004830 | Molecular Function | tryptophan-tRNA ligase activity | 1 | 3 | 0.014 |
| GO:0008897 | Molecular Function | holo-[acyl-carrier-protein] synthase activity | 1 | 3 | 0.014 |
| GO:0008199 | Molecular Function | ferric iron binding | 1 | 3 | 0.014 |
| GO:0003824 | Molecular Function | catalytic activity | 57 | 12724 | 0.016 |
| GO:0070011 | Molecular Function | peptidase activity, acting on L-amino acid peptides | 11 | 1470 | 0.018 |
| GO:0008233 | Molecular Function | peptidase activity | 11 | 1489 | 0.02 |
| GO:0070001 | Molecular Function | aspartic-type peptidase activity | 6 | 607 | 0.025 |
| GO:0004190 | Molecular Function | aspartic-type endopeptidase activity | 6 | 607 | 0.025 |
| GO:0004497 | Molecular Function | monooxygenase activity | 5 | 458 | 0.027 |

| | | | | | |
|---|---|---|---|---|---|
| GO:0016780 | Molecular Function | phosphotransferase activity, for other substituted phosphate groups | 1 | 9 | 0.036 |
| GO:0016747 | Molecular Function | transferase activity, transferring acyl groups other than amino-acyl groups | 4 | 348 | 0.04 |
| GO:0016744 | Molecular Function | transferase activity, transferring aldehyde or ketonic groups | 1 | 11 | 0.043 |
| GO:0016746 | Molecular Function | transferase activity, transferring acyl groups | 4 | 367 | 0.047 |
| GO:0009579 | Cellular Component | thylakoid | 2 | 93 | 0.047 |

**Genes in rufipogon_SNP CLR**

| | | | | | |
|---|---|---|---|---|---|
| GO:0006091 | Biological Process | generation of precursor metabolites and energy | 16 | 243 | 2.50E-11 |
| GO:0022900 | Biological Process | electron transport chain | 8 | 60 | 1.80E-08 |
| GO:0006119 | Biological Process | oxidative phosphorylation | 9 | 88 | 1.90E-08 |
| GO:0015986 | Biological Process | ATP synthesis coupled proton transport | 7 | 69 | 8.10E-07 |
| GO:0015985 | Biological Process | energy coupled proton transport, down electrochemical gradient | 7 | 69 | 8.10E-07 |
| GO:0034220 | Biological Process | ion transmembrane transport | 7 | 69 | 8.10E-07 |
| GO:0015979 | Biological Process | photosynthesis | 8 | 115 | 1.90E-06 |
| GO:0015992 | Biological Process | proton transport | 7 | 80 | 2.00E-06 |
| GO:0006818 | Biological Process | hydrogen transport | 7 | 80 | 2.00E-06 |
| GO:0006754 | Biological Process | ATP biosynthetic process | 8 | 127 | 3.80E-06 |
| GO:0046034 | Biological Process | ATP metabolic process | 8 | 129 | 4.20E-06 |
| GO:0009142 | Biological Process | nucleoside triphosphate biosynthetic process | 8 | 132 | 5.00E-06 |
| GO:0009145 | Biological Process | purine nucleoside triphosphate biosynthetic process | 8 | 132 | 5.00E-06 |
| GO:0009206 | Biological Process | purine ribonucleoside triphosphate biosynthetic process | 8 | 132 | 5.00E-06 |
| GO:0009201 | Biological Process | ribonucleoside triphosphate biosynthetic process | 8 | 132 | 5.00E-06 |
| GO:0009199 | Biological Process | ribonucleoside triphosphate metabolic process | 8 | 134 | 5.50E-06 |
| GO:0009205 | Biological Process | purine ribonucleoside triphosphate metabolic process | 8 | 134 | 5.50E-06 |
| GO:0009144 | Biological Process | purine nucleoside triphosphate metabolic process | 8 | 134 | 5.50E-06 |

| | | | | | |
|---|---|---|---|---|---|
| GO:0009141 | Biological Process | nucleoside triphosphate metabolic process | 8 | 135 | 5.80E-06 |
| GO:0009152 | Biological Process | purine ribonucleotide biosynthetic process | 8 | 145 | 9.50E-06 |
| GO:0009150 | Biological Process | purine ribonucleotide metabolic process | 8 | 147 | 1.00E-05 |
| GO:0009260 | Biological Process | ribonucleotide biosynthetic process | 8 | 149 | 1.20E-05 |
| GO:0006164 | Biological Process | purine nucleotide biosynthetic process | 8 | 150 | 1.20E-05 |
| GO:0009259 | Biological Process | ribonucleotide metabolic process | 8 | 151 | 1.30E-05 |
| GO:0006163 | Biological Process | purine nucleotide metabolic process | 8 | 154 | 1.40E-05 |
| GO:0019684 | Biological Process | photosynthesis, light reaction | 5 | 47 | 2.60E-05 |
| GO:0009165 | Biological Process | nucleotide biosynthetic process | 8 | 179 | 4.10E-05 |
| GO:0009767 | Biological Process | photosynthetic electron transport chain | 4 | 26 | 4.70E-05 |
| GO:0022904 | Biological Process | respiratory electron transport chain | 4 | 28 | 6.10E-05 |
| GO:0006412 | Biological Process | translation | 14 | 595 | 6.70E-05 |
| GO:0015672 | Biological Process | monovalent inorganic cation transport | 7 | 155 | 0.00012 |
| GO:0006753 | Biological Process | nucleoside phosphate metabolic process | 8 | 218 | 0.00015 |
| GO:0009117 | Biological Process | nucleotide metabolic process | 8 | 218 | 0.00015 |
| GO:0055086 | Biological Process | nucleobase, nucleoside and nucleotide metabolic process | 8 | 244 | 0.00032 |
| GO:0045333 | Biological Process | cellular respiration | 4 | 48 | 0.00042 |
| GO:0006811 | Biological Process | ion transport | 10 | 408 | 0.00055 |
| GO:0015980 | Biological Process | energy derivation by oxidation of organic compounds | 4 | 54 | 0.00063 |
| GO:0046483 | Biological Process | heterocycle metabolic process | 8 | 312 | 0.0015 |
| GO:0055114 | Biological Process | oxidation reduction | 19 | 1351 | 0.0022 |
| GO:0023034 | Biological Process | intracellular signaling pathway | 3 | 36 | 0.0023 |
| GO:0006812 | Biological Process | cation transport | 8 | 348 | 0.0029 |
| GO:0006810 | Biological Process | transport | 20 | 1638 | 0.008 |
| GO:0051234 | Biological Process | establishment of localization | 20 | 1638 | 0.008 |
| GO:0042773 | Biological Process | ATP synthesis coupled electron transport | 2 | 19 | 0.0087 |
| GO:0051179 | Biological Process | localization | 20 | 1651 | 0.0087 |
| GO:0055085 | Biological Process | transmembrane transport | 11 | 701 | 0.0089 |
| GO:0023033 | Biological Process | signaling pathway | 3 | 61 | 0.0092 |

| | | | | | |
|---|---|---|---|---|---|
| GO:0010467 | Biological Process | gene expression | 26 | 2424 | 0.013 |
| GO:0016197 | Biological Process | endosome transport | 1 | 1 | 0.013 |
| GO:0042147 | Biological Process | retrograde transport, endosome to Golgi | 1 | 1 | 0.013 |
| GO:0046907 | Biological Process | intracellular transport | 5 | 233 | 0.023 |
| GO:0033036 | Biological Process | macromolecule localization | 7 | 411 | 0.023 |
| GO:0006571 | Biological Process | tyrosine biosynthetic process | 1 | 3 | 0.027 |
| GO:0006570 | Biological Process | tyrosine metabolic process | 1 | 4 | 0.033 |
| GO:0050821 | Biological Process | protein stabilization | 1 | 4 | 0.033 |
| GO:0009775 | Biological Process | photosynthetic electron transport in cytochrome b6/f | 1 | 4 | 0.033 |
| GO:0031647 | Biological Process | regulation of protein stability | 1 | 4 | 0.033 |
| GO:0006419 | Biological Process | alanyl-tRNA aminoacylation | 1 | 4 | 0.033 |
| GO:0042549 | Biological Process | photosystem II stabilization | 1 | 5 | 0.04 |
| GO:0042548 | Biological Process | regulation of photosynthesis, light reaction | 1 | 5 | 0.04 |
| GO:0010109 | Biological Process | regulation of photosynthesis | 1 | 5 | 0.04 |
| GO:0043467 | Biological Process | regulation of generation of precursor metabolites and energy | 1 | 5 | 0.04 |
| GO:0016192 | Biological Process | vesicle-mediated transport | 4 | 186 | 0.04 |
| GO:0006842 | Biological Process | tricarboxylic acid transport | 1 | 6 | 0.046 |
| GO:0015746 | Biological Process | citrate transport | 1 | 6 | 0.046 |
| GO:0016655 | Molecular Function | oxidoreductase activity, acting on NADH or NADPH, quinone or similar compound as acceptor | 5 | 34 | 6.20E-06 |
| GO:0015078 | Molecular Function | hydrogen ion transmembrane transporter activity | 6 | 95 | 6.30E-05 |
| GO:0050136 | Molecular Function | NADH dehydrogenase (quinone) activity | 4 | 30 | 7.80E-05 |
| GO:0008137 | Molecular Function | NADH dehydrogenase (ubiquinone) activity | 4 | 30 | 7.80E-05 |
| GO:0003954 | Molecular Function | NADH dehydrogenase activity | 4 | 30 | 7.80E-05 |
| GO:0003735 | Molecular Function | structural constituent of ribosome | 11 | 396 | 0.0001 |
| GO:0016651 | Molecular Function | oxidoreductase activity, acting on NADH or NADPH | 5 | 64 | 0.0001 |
| GO:0015077 | Molecular Function | monovalent inorganic cation transmembrane transporter activity | 6 | 126 | 0.00027 |
| GO:0005198 | Molecular Function | structural molecule activity | 11 | 462 | 0.00037 |

| | | | | | |
|---|---|---|---|---|---|
| GO:0022890 | Molecular Function | inorganic cation transmembrane transporter activity | 6 | 180 | 0.0017 |
| GO:0016168 | Molecular Function | chlorophyll binding | 2 | 8 | 0.002 |
| GO:0022892 | Molecular Function | substrate-specific transporter activity | 11 | 593 | 0.0027 |
| GO:0022857 | Molecular Function | transmembrane transporter activity | 11 | 643 | 0.0048 |
| GO:0015075 | Molecular Function | ion transmembrane transporter activity | 8 | 389 | 0.0056 |
| GO:0045156 | Molecular Function | electron transporter, transferring electrons within the cyclic electron transport pathway of photosynthesis activity | 2 | 15 | 0.0057 |
| GO:0009055 | Molecular Function | electron carrier activity | 10 | 599 | 0.0083 |
| GO:0022891 | Molecular Function | substrate-specific transmembrane transporter activity | 9 | 517 | 0.0094 |
| GO:0005215 | Molecular Function | transporter activity | 13 | 941 | 0.012 |
| GO:0008324 | Molecular Function | cation transmembrane transporter activity | 6 | 280 | 0.013 |
| GO:0016820 | Molecular Function | hydrolase activity, acting on acid anhydrides, catalyzing transmembrane movement of substances | 4 | 147 | 0.019 |
| GO:0017176 | Molecular Function | phosphatidylinositol N-acetylglucosaminyltransferase activity | 1 | 2 | 0.02 |
| GO:0016885 | Molecular Function | ligase activity, forming carbon-carbon bonds | 1 | 2 | 0.02 |
| GO:0003989 | Molecular Function | acetyl-CoA carboxylase activity | 1 | 2 | 0.02 |
| GO:0004802 | Molecular Function | transketolase activity | 1 | 2 | 0.02 |
| GO:0016421 | Molecular Function | CoA carboxylase activity | 1 | 2 | 0.02 |
| GO:0046933 | Molecular Function | hydrogen ion transporting ATP synthase activity, rotational mechanism | 2 | 32 | 0.022 |
| GO:0046961 | Molecular Function | proton-transporting ATPase activity, rotational mechanism | 2 | 35 | 0.026 |
| GO:0004665 | Molecular Function | prephenate dehydrogenase (NADP+) activity | 1 | 3 | 0.027 |
| GO:0042586 | Molecular Function | peptide deformylase activity | 1 | 3 | 0.027 |
| GO:0009374 | Molecular Function | biotin binding | 1 | 3 | 0.027 |
| GO:0042625 | Molecular Function | ATPase activity, coupled to transmembrane movement of ions | 3 | 94 | 0.028 |
| GO:0004813 | Molecular Function | alanine-tRNA ligase activity | 1 | 4 | 0.033 |
| GO:0042301 | Molecular Function | phosphate binding | 1 | 4 | 0.033 |

| | | | | | |
|---|---|---|---|---|---|
| GO:0043168 | Molecular Function | anion binding | 1 | 4 | 0.033 |
| GO:0016491 | Molecular Function | oxidoreductase activity | 18 | 1672 | 0.036 |
| GO:0016817 | Molecular Function | hydrolase activity, acting on acid anhydrides | 10 | 764 | 0.037 |
| GO:0004650 | Molecular Function | polygalacturonase activity | 2 | 45 | 0.04 |
| GO:0042623 | Molecular Function | ATPase activity, coupled | 5 | 275 | 0.041 |
| GO:0019829 | Molecular Function | cation-transporting ATPase activity | 2 | 47 | 0.043 |
| GO:0015137 | Molecular Function | citrate transmembrane transporter activity | 1 | 6 | 0.046 |
| GO:0015142 | Molecular Function | tricarboxylic acid transmembrane transporter activity | 1 | 6 | 0.046 |
| GO:0009579 | Cellular Component | thylakoid | 9 | 93 | 3.00E-08 |
| GO:0032991 | Cellular Component | macromolecular complex | 27 | 1207 | 5.80E-08 |
| GO:0034357 | Cellular Component | photosynthetic membrane | 8 | 89 | 3.10E-07 |
| GO:0045259 | Cellular Component | proton-transporting ATP synthase complex | 6 | 44 | 1.00E-06 |
| GO:0016469 | Cellular Component | proton-transporting two-sector ATPase complex | 7 | 73 | 1.20E-06 |
| GO:0044425 | Cellular Component | membrane part | 21 | 1007 | 5.90E-06 |
| GO:0009521 | Cellular Component | photosystem | 6 | 72 | 1.50E-05 |
| GO:0045263 | Cellular Component | proton-transporting ATP synthase complex, coupling factor F(o) | 4 | 20 | 1.90E-05 |
| GO:0043234 | Cellular Component | protein complex | 16 | 706 | 3.10E-05 |
| GO:0033177 | Cellular Component | proton-transporting two-sector ATPase complex, proton-transporting domain | 4 | 30 | 7.80E-05 |
| GO:0016020 | Cellular Component | membrane | 30 | 2115 | 9.20E-05 |
| GO:0005840 | Cellular Component | ribosome | 11 | 397 | 0.0001 |
| GO:0044444 | Cellular Component | cytoplasmic part | 18 | 966 | 0.00012 |
| GO:0030529 | Cellular Component | ribonucleoprotein complex | 11 | 438 | 0.00024 |
| GO:0005737 | Cellular Component | cytoplasm | 19 | 1263 | 0.001 |
| GO:0044424 | Cellular Component | intracellular part | 38 | 3529 | 0.0023 |
| GO:0033279 | Cellular Component | ribosomal subunit | 4 | 82 | 0.0027 |
| GO:0033178 | Cellular Component | proton-transporting two-sector ATPase complex, catalytic domain | 3 | 39 | 0.0028 |
| GO:0009539 | Cellular Component | photosystem II reaction center | 2 | 11 | 0.0033 |

| | | | | | |
|---|---|---|---|---|---|
| GO:0009523 | Cellular Component | photosystem II | 3 | 46 | 0.0044 |
| GO:0044436 | Cellular Component | thylakoid part | 3 | 50 | 0.0055 |
| GO:0015935 | Cellular Component | small ribosomal subunit | 3 | 51 | 0.0058 |
| GO:0048475 | Cellular Component | coated membrane | 3 | 59 | 0.0084 |
| GO:0030117 | Cellular Component | membrane coat | 3 | 59 | 0.0084 |
| GO:0042651 | Cellular Component | thylakoid membrane | 2 | 23 | 0.012 |
| GO:0045261 | Cellular Component | proton-transporting ATP synthase complex, catalytic core F(1) | 2 | 24 | 0.013 |
| GO:0005622 | Cellular Component | intracellular | 39 | 4116 | 0.017 |
| GO:0030075 | Cellular Component | plasma membrane-derived thylakoid | 1 | 3 | 0.027 |
| GO:0030096 | Cellular Component | plasma membrane-derived thylakoid photosystem II | 1 | 3 | 0.027 |
| GO:0009536 | Cellular Component | plastid | 2 | 37 | 0.028 |
| GO:0031361 | Cellular Component | integral to thylakoid membrane | 1 | 5 | 0.04 |
| GO:0012511 | Cellular Component | monolayer-surrounded lipid storage body | 1 | 6 | 0.046 |
| GO:0005811 | Cellular Component | lipid particle | 1 | 6 | 0.046 |

**Genes in rufipogon_SV CLR**

| | | | | | |
|---|---|---|---|---|---|
| GO:0034641 | Biological Process | cellular nitrogen compound metabolic process | 13 | 435 | 0.00038 |
| GO:0044281 | Biological Process | small molecule metabolic process | 22 | 1034 | 0.0005 |
| GO:0043436 | Biological Process | oxoacid metabolic process | 13 | 473 | 0.00082 |
| GO:0019752 | Biological Process | carboxylic acid metabolic process | 13 | 473 | 0.00082 |
| GO:0006082 | Biological Process | organic acid metabolic process | 13 | 474 | 0.00083 |
| GO:0042180 | Biological Process | cellular ketone metabolic process | 13 | 480 | 0.00093 |
| GO:0044106 | Biological Process | cellular amine metabolic process | 10 | 333 | 0.0017 |
| GO:0006520 | Biological Process | cellular amino acid metabolic process | 9 | 303 | 0.0031 |
| GO:0009056 | Biological Process | catabolic process | 11 | 433 | 0.0037 |
| GO:0019674 | Biological Process | NAD metabolic process | 2 | 8 | 0.0038 |
| GO:0019359 | Biological Process | nicotinamide nucleotide biosynthetic process | 2 | 8 | 0.0038 |
| GO:0009435 | Biological Process | NAD biosynthetic process | 2 | 8 | 0.0038 |
| GO:0009308 | Biological Process | amine metabolic process | 10 | 380 | 0.0043 |

| | | | | | |
|---|---|---|---|---|---|
| GO:0006519 | Biological Process | cellular amino acid and derivative metabolic process | 10 | 391 | 0.0052 |
| GO:0019363 | Biological Process | pyridine nucleotide biosynthetic process | 2 | 10 | 0.0056 |
| GO:0008104 | Biological Process | protein localization | 8 | 307 | 0.011 |
| GO:0044283 | Biological Process | small molecule biosynthetic process | 8 | 320 | 0.014 |
| GO:0043632 | Biological Process | modification-dependent macromolecule catabolic process | 5 | 143 | 0.014 |
| GO:0006511 | Biological Process | ubiquitin-dependent protein catabolic process | 5 | 143 | 0.014 |
| GO:0019941 | Biological Process | modification-dependent protein catabolic process | 5 | 143 | 0.014 |
| GO:0051186 | Biological Process | cofactor metabolic process | 5 | 143 | 0.014 |
| GO:0009161 | Biological Process | ribonucleoside monophosphate metabolic process | 2 | 17 | 0.014 |
| GO:0009156 | Biological Process | ribonucleoside monophosphate biosynthetic process | 2 | 17 | 0.014 |
| GO:0051188 | Biological Process | cofactor biosynthetic process | 4 | 94 | 0.014 |
| GO:0051179 | Biological Process | localization | 25 | 1651 | 0.017 |
| GO:0044257 | Biological Process | cellular protein catabolic process | 5 | 153 | 0.018 |
| GO:0051603 | Biological Process | proteolysis involved in cellular protein catabolic process | 5 | 153 | 0.018 |
| GO:0044248 | Biological Process | cellular catabolic process | 8 | 337 | 0.018 |
| GO:0006732 | Biological Process | coenzyme metabolic process | 4 | 101 | 0.018 |
| GO:0009108 | Biological Process | coenzyme biosynthetic process | 3 | 56 | 0.019 |
| GO:0060548 | Biological Process | negative regulation of cell death | 1 | 1 | 0.019 |
| GO:0043067 | Biological Process | regulation of programmed cell death | 1 | 1 | 0.019 |
| GO:0043066 | Biological Process | negative regulation of apoptosis | 1 | 1 | 0.019 |
| GO:0043069 | Biological Process | negative regulation of programmed cell death | 1 | 1 | 0.019 |
| GO:0042981 | Biological Process | regulation of apoptosis | 1 | 1 | 0.019 |
| GO:0010941 | Biological Process | regulation of cell death | 1 | 1 | 0.019 |
| GO:0016070 | Biological Process | RNA metabolic process | 18 | 1088 | 0.019 |
| GO:0033036 | Biological Process | macromolecule localization | 9 | 411 | 0.02 |
| GO:0009123 | Biological Process | nucleoside monophosphate metabolic process | 2 | 21 | 0.02 |
| GO:0009124 | Biological Process | nucleoside monophosphate biosynthetic process | 2 | 21 | 0.02 |
| GO:0006753 | Biological Process | nucleoside phosphate metabolic process | 6 | 218 | 0.021 |

| | | | | | |
|---|---|---|---|---|---|
| GO:0009117 | Biological Process | nucleotide metabolic process | 6 | 218 | 0.021 |
| GO:0046394 | Biological Process | carboxylic acid biosynthetic process | 6 | 222 | 0.022 |
| GO:0016053 | Biological Process | organic acid biosynthetic process | 6 | 222 | 0.022 |
| GO:0045184 | Biological Process | establishment of protein localization | 7 | 294 | 0.025 |
| GO:0015031 | Biological Process | protein transport | 7 | 294 | 0.025 |
| GO:0009820 | Biological Process | alkaloid metabolic process | 2 | 25 | 0.027 |
| GO:0046496 | Biological Process | nicotinamide nucleotide metabolic process | 2 | 25 | 0.027 |
| GO:0006769 | Biological Process | nicotinamide metabolic process | 2 | 25 | 0.027 |
| GO:0051234 | Biological Process | establishment of localization | 24 | 1638 | 0.027 |
| GO:0006810 | Biological Process | transport | 24 | 1638 | 0.027 |
| GO:0006418 | Biological Process | tRNA aminoacylation for protein translation | 3 | 66 | 0.028 |
| GO:0043038 | Biological Process | amino acid activation | 3 | 66 | 0.028 |
| GO:0043039 | Biological Process | tRNA aminoacylation | 3 | 66 | 0.028 |
| GO:0006427 | Biological Process | histidyl-tRNA aminoacylation | 1 | 2 | 0.028 |
| GO:0006425 | Biological Process | glutaminyl-tRNA aminoacylation | 1 | 2 | 0.028 |
| GO:0019362 | Biological Process | pyridine nucleotide metabolic process | 2 | 27 | 0.031 |
| GO:0009165 | Biological Process | nucleotide biosynthetic process | 5 | 179 | 0.032 |
| GO:0055086 | Biological Process | nucleobase, nucleoside and nucleotide metabolic process | 6 | 244 | 0.033 |
| GO:0043603 | Biological Process | cellular amide metabolic process | 2 | 28 | 0.033 |
| GO:0030163 | Biological Process | protein catabolic process | 5 | 184 | 0.035 |
| GO:0007049 | Biological Process | cell cycle | 2 | 30 | 0.037 |
| GO:0050794 | Biological Process | regulation of cellular process | 27 | 1955 | 0.037 |
| GO:0009057 | Biological Process | macromolecule catabolic process | 7 | 322 | 0.038 |
| GO:0044265 | Biological Process | cellular macromolecule catabolic process | 6 | 254 | 0.038 |
| GO:0006644 | Biological Process | phospholipid metabolic process | 3 | 77 | 0.041 |
| GO:0045454 | Biological Process | cell redox homeostasis | 4 | 132 | 0.041 |
| GO:0010467 | Biological Process | gene expression | 32 | 2424 | 0.041 |
| GO:0009110 | Biological Process | vitamin biosynthetic process | 2 | 32 | 0.041 |
| GO:0006767 | Biological Process | water-soluble vitamin metabolic process | 2 | 32 | 0.041 |

| | | | | | |
|---|---|---|---|---|---|
| GO:0006766 | Biological Process | vitamin metabolic process | 2 | 32 | 0.041 |
| GO:0042364 | Biological Process | water-soluble vitamin biosynthetic process | 2 | 32 | 0.041 |
| GO:0008652 | Biological Process | cellular amino acid biosynthetic process | 4 | 133 | 0.042 |
| GO:0006733 | Biological Process | oxidoreduction coenzyme metabolic process | 2 | 34 | 0.046 |
| GO:0010498 | Biological Process | proteasomal protein catabolic process | 1 | 4 | 0.047 |
| GO:0043161 | Biological Process | proteasomal ubiquitin-dependent protein catabolic process | 1 | 4 | 0.047 |
| GO:0034660 | Biological Process | ncRNA metabolic process | 4 | 139 | 0.048 |
| GO:0006886 | Biological Process | intracellular protein transport | 5 | 202 | 0.048 |
| GO:0019725 | Biological Process | cellular homeostasis | 4 | 140 | 0.049 |
| GO:0043565 | Molecular Function | sequence-specific DNA binding | 12 | 453 | 0.0017 |
| GO:0016538 | Molecular Function | cyclin-dependent protein kinase regulator activity | 2 | 6 | 0.0024 |
| GO:0019887 | Molecular Function | protein kinase regulator activity | 2 | 9 | 0.0047 |
| GO:0019207 | Molecular Function | kinase regulator activity | 2 | 9 | 0.0047 |
| GO:0042623 | Molecular Function | ATPase activity, coupled | 8 | 275 | 0.0059 |
| GO:0016887 | Molecular Function | ATPase activity | 9 | 355 | 0.0083 |
| GO:0017111 | Molecular Function | nucleoside-triphosphatase activity | 14 | 726 | 0.012 |
| GO:0004707 | Molecular Function | MAP kinase activity | 2 | 16 | 0.012 |
| GO:0004702 | Molecular Function | receptor signaling protein serine/threonine kinase activity | 2 | 16 | 0.012 |
| GO:0005057 | Molecular Function | receptor signaling protein activity | 2 | 16 | 0.012 |
| GO:0016462 | Molecular Function | pyrophosphatase activity | 14 | 743 | 0.014 |
| GO:0051536 | Molecular Function | iron-sulfur cluster binding | 4 | 95 | 0.015 |
| GO:0051540 | Molecular Function | metal cluster binding | 4 | 95 | 0.015 |
| GO:0016818 | Molecular Function | hydrolase activity, acting on acid anhydrides, in phosphorus-containing anhydrides | 14 | 759 | 0.016 |
| GO:0016817 | Molecular Function | hydrolase activity, acting on acid anhydrides | 14 | 764 | 0.017 |
| GO:0015922 | Molecular Function | aspartate oxidase activity | 1 | 1 | 0.019 |
| GO:0003879 | Molecular Function | ATP phosphoribosyltransferase activity | 1 | 1 | 0.019 |
| GO:0004334 | Molecular Function | fumarylacetoacetase activity | 1 | 1 | 0.019 |

| | | | | | |
|---|---|---|---|---|---|
| GO:0016823 | Molecular Function | hydrolase activity, acting on acid carbon-carbon bonds, in ketonic substances | 1 | 1 | 0.019 |
| GO:0016822 | Molecular Function | hydrolase activity, acting on acid carbon-carbon bonds | 1 | 1 | 0.019 |
| GO:0008864 | Molecular Function | formyltetrahydrofolate deformylase activity | 1 | 1 | 0.019 |
| GO:0008987 | Molecular Function | quinolinate synthetase A activity | 1 | 1 | 0.019 |
| GO:0008734 | Molecular Function | L-aspartate oxidase activity | 1 | 1 | 0.019 |
| GO:0016790 | Molecular Function | thiolester hydrolase activity | 3 | 57 | 0.019 |
| GO:0005215 | Molecular Function | transporter activity | 16 | 941 | 0.021 |
| GO:0008173 | Molecular Function | RNA methyltransferase activity | 2 | 24 | 0.025 |
| GO:0015399 | Molecular Function | primary active transmembrane transporter activity | 5 | 170 | 0.026 |
| GO:0015405 | Molecular Function | P-P-bond-hydrolysis-driven transmembrane transporter activity | 5 | 170 | 0.026 |
| GO:0016875 | Molecular Function | ligase activity, forming carbon-oxygen bonds | 3 | 66 | 0.028 |
| GO:0004812 | Molecular Function | aminoacyl-tRNA ligase activity | 3 | 66 | 0.028 |
| GO:0016876 | Molecular Function | ligase activity, forming aminoacyl-tRNA and related compounds | 3 | 66 | 0.028 |
| GO:0004819 | Molecular Function | glutamine-tRNA ligase activity | 1 | 2 | 0.028 |
| GO:0004821 | Molecular Function | histidine-tRNA ligase activity | 1 | 2 | 0.028 |
| GO:0004609 | Molecular Function | phosphatidylserine decarboxylase activity | 1 | 2 | 0.028 |
| GO:0008474 | Molecular Function | palmitoyl-(protein) hydrolase activity | 1 | 2 | 0.028 |
| GO:0070818 | Molecular Function | protoporphyrinogen oxidase activity | 1 | 2 | 0.028 |
| GO:0004729 | Molecular Function | oxygen-dependent protoporphyrinogen oxidase activity | 1 | 2 | 0.028 |
| GO:0004750 | Molecular Function | ribulose-phosphate 3-epimerase activity | 1 | 2 | 0.028 |
| GO:0016874 | Molecular Function | ligase activity | 7 | 305 | 0.03 |
| GO:0016831 | Molecular Function | carboxy-lyase activity | 3 | 68 | 0.03 |
| GO:0043566 | Molecular Function | structure-specific DNA binding | 2 | 28 | 0.033 |
| GO:0004455 | Molecular Function | ketol-acid reductoisomerase activity | 1 | 3 | 0.037 |
| GO:0030234 | Molecular Function | enzyme regulator activity | 6 | 255 | 0.039 |
| GO:0051539 | Molecular Function | 4 iron, 4 sulfur cluster binding | 2 | 31 | 0.039 |
| GO:0004822 | Molecular Function | isoleucine-tRNA ligase activity | 1 | 4 | 0.047 |

| | | | | | |
|---|---|---|---|---|---|
| GO:0044464 | Cellular Component | cell part | 79 | 5945 | 0.00081 |
| GO:0005623 | Cellular Component | cell | 79 | 5945 | 0.00081 |
| GO:0016591 | Cellular Component | DNA-directed RNA polymerase II, holoenzyme | 2 | 15 | 0.011 |
| GO:0005622 | Cellular Component | intracellular | 53 | 4116 | 0.014 |
| GO:0000441 | Cellular Component | SSL2-core TFIIH complex | 1 | 1 | 0.019 |
| GO:0032806 | Cellular Component | carboxy-terminal domain protein kinase complex | 1 | 1 | 0.019 |
| GO:0000439 | Cellular Component | core TFIIH complex | 1 | 1 | 0.019 |
| GO:0031224 | Cellular Component | intrinsic to membrane | 14 | 775 | 0.019 |
| GO:0005665 | Cellular Component | DNA-directed RNA polymerase II, core complex | 1 | 2 | 0.028 |
| GO:0016021 | Cellular Component | integral to membrane | 13 | 752 | 0.032 |
| GO:0005783 | Cellular Component | endoplasmic reticulum | 4 | 123 | 0.033 |
| GO:0016020 | Cellular Component | membrane | 29 | 2115 | 0.034 |
| GO:0005737 | Cellular Component | cytoplasm | 19 | 1263 | 0.037 |
| GO:0005960 | Cellular Component | glycine cleavage complex | 1 | 3 | 0.037 |
| GO:0005672 | Cellular Component | transcription factor TFIIA complex | 1 | 3 | 0.037 |

**Table S10.** Functional annotation of fixed genes between pairs of taxa (rufipogon, indica and japonica) and private genes underlying SVs in each subspecies.

| Gene ID (MSU7) | Function annotation |
|---|---|
| **Fixed genes (SVs) in japonica-rufipogon comparison** | |
| LOC_Os01g14160 | retrotransposon, putative, centromere-specific, expressed |
| **Fixed genes (SVs) in indica-japonica comparison** | |
| LOC_Os01g01540 | retrotransposon protein, putative, unclassified, expressed |
| LOC_Os01g01550 | transposon protein, putative, CACTA, En/Spm sub-class, expressed |
| LOC_Os01g07430 | myb-like DNA-binding domain containing protein, putative, expressed |
| LOC_Os01g10160 | latency associated nuclear antigen, putative, expressed |
| LOC_Os01g10170 | expressed protein |
| LOC_Os01g49154 | expressed protein |
| LOC_Os01g49750 | expressed protein |
| LOC_Os02g04725 | dolichol phosphate-mannose biosynthesis regulatory protein, putative, expressed |
| LOC_Os02g16540 | WRKY39, expressed |
| LOC_Os02g51330 | MAG2, putative, expressed |
| LOC_Os03g01340 | transposon protein, putative, unclassified |
| LOC_Os03g21380 | OsCML27 - Calmodulin-related calcium sensor protein, expressed |
| LOC_Os03g59930 | retrotransposon, putative, centromere-specific, expressed |
| LOC_Os04g01874 | protein kinase, putative, expressed |
| LOC_Os04g53930 | organic cation transporter protein, putative, expressed |
| LOC_Os05g50850 | expressed protein |
| LOC_Os06g22370 | expressed protein |
| LOC_Os08g04340 | plastocyanin-like domain containing protein, putative, expressed |
| LOC_Os08g04400 | pentatricopeptide repeat domain containing protein, putative, expressed |
| LOC_Os08g04420 | tyrosine protein kinase domain containing protein, putative, expressed |
| LOC_Os08g04430 | expressed protein |
| LOC_Os08g04440 | RNA recognition motif containing protein, putative, expressed |
| LOC_Os08g04450 | DAG protein, chloroplast precursor, putative, expressed |

| | |
|---|---|
| LOC_Os08g04460 | NADPH-dependent FMN reductase domain containing protein, expressed |
| LOC_Os08g04470 | U box protein 8, putative, expressed |
| LOC_Os08g04490 | expressed protein |
| LOC_Os08g04500 | terpene synthase, putative, expressed |
| LOC_Os08g04540 | decarboxylase, putative, expressed |
| LOC_Os08g04550 | expressed protein |
| LOC_Os08g04570 | expressed protein |
| LOC_Os08g04580 | csAtPR5, putative, expressed |
| LOC_Os08g04710 | expressed protein |
| LOC_Os08g04760 | retrotransposon protein, putative, unclassified, expressed |
| LOC_Os08g04770 | AMP-binding domain containing protein, expressed |
| LOC_Os08g04780 | amine oxidase, putative, expressed |
| LOC_Os08g04790 | xylosyltransferase, putative, expressed |
| LOC_Os08g04800 | triacylglycerol lipase like protein, putative, expressed |
| LOC_Os08g04810 | serine esterase family protein, putative, expressed |
| LOC_Os08g04840 | MYB family transcription factor, putative, expressed |
| LOC_Os08g04900 | hypothetical protein |
| LOC_Os08g04910 | expressed protein |
| LOC_Os08g05040 | expressed protein |
| LOC_Os08g05090 | expressed protein |
| LOC_Os08g05100 | expressed protein |
| LOC_Os08g05110 | expressed protein |
| LOC_Os08g05160 | expressed protein |
| LOC_Os08g05280 | oxidoreductase family, NAD-binding Rossmann fold containing protein, expressed |
| LOC_Os08g05310 | expressed protein |
| LOC_Os08g05340 | expressed protein |
| LOC_Os08g05370 | conserved hypothetical protein |
| LOC_Os08g05480 | OsFBX261 - F-box domain containing protein, expressed |
| LOC_Os08g05490 | tRNA synthetase, putative, expressed |

| | |
|---|---|
| LOC_Os08g05510 | MYB family transcription factor, putative, expressed |

**Private genes (DUPs) in indica against rufipogon**

| | |
|---|---|
| LOC_Os01g01010 | TBC domain containing protein, expressed |
| LOC_Os01g01040 | expressed protein |
| LOC_Os01g01060 | 40S ribosomal protein S5, putative, expressed |
| LOC_Os01g01070 | expressed protein |
| LOC_Os01g01080 | decarboxylase, putative, expressed |
| LOC_Os01g01090 | retrotransposon protein, putative, unclassified, expressed |
| LOC_Os01g01100 | retrotransposon protein, putative, Ty3-gypsy subclass, expressed |
| LOC_Os01g01110 | expressed protein |
| LOC_Os01g01115 | pentatricopeptide, putative, expressed |
| LOC_Os01g01120 | enolase-phosphatase E1, putative, expressed |
| LOC_Os01g01140 | expressed protein |
| LOC_Os01g01150 | RNA recognition motif, putative, expressed |
| LOC_Os01g01160 | heat shock protein DnaJ, putative, expressed |
| LOC_Os01g01210 | retrotransposon protein, putative, unclassified |
| LOC_Os01g01240 | retrotransposon protein, putative, Ty3-gypsy subclass |
| LOC_Os01g01270 | retrotransposon protein, putative, unclassified, expressed |
| LOC_Os01g01290 | histone-like transcription factor and archaeal histone, putative, expressed |
| LOC_Os01g01295 | expressed protein |
| LOC_Os01g01302 | shikimate kinase, putative, expressed |
| LOC_Os01g01307 | translocon-associated protein beta domain containing protein, expressed |
| LOC_Os01g01320 | retrotransposon protein, putative, Ty1-copia subclass, expressed |
| LOC_Os01g01350 | SNF7 domain containing protein, putative, expressed |
| LOC_Os01g01360 | peptide transporter PTR2, putative, expressed |
| LOC_Os01g01369 | 3-beta-hydroxysteroid-Delta-isomerase, putative, expressed |
| LOC_Os01g01380 | expressed protein |
| LOC_Os01g01390 | expressed protein |

| | |
|---|---|
| LOC_Os01g01400 | expressed protein |
| LOC_Os01g01430 | No apical meristem protein, putative, expressed |
| LOC_Os01g01450 | stress responsive protein, putative, expressed |
| LOC_Os01g01460 | retrotransposon protein, putative, Ty3-gypsy subclass |
| LOC_Os01g01470 | no apical meristem protein, putative, expressed |
| LOC_Os01g01484 | light-mediated development protein DET1, putative, expressed |
| LOC_Os01g01500 | expressed protein |
| LOC_Os01g01510 | sas10/Utp3 family protein, expressed |
| LOC_Os01g01520 | transferase family protein, putative, expressed |
| LOC_Os01g01530 | transposon protein, putative, CACTA, En/Spm sub-class |
| LOC_Os01g01540 | retrotransposon protein, putative, unclassified, expressed |
| LOC_Os01g01550 | transposon protein, putative, CACTA, En/Spm sub-class, expressed |
| LOC_Os01g01560 | transposon protein, putative, unclassified, expressed |
| LOC_Os01g01580 | expressed protein |
| LOC_Os01g01600 | ethylene-responsive protein related, putative, expressed |
| LOC_Os01g01610 | IscA-like iron-sulfur assembly protein, mitochondrial precursor, putative, expressed |
| LOC_Os01g01620 | kinase, pfkB family, putative, expressed |
| LOC_Os01g01640 | Rer1 protein, putative, expressed |
| LOC_Os01g01650 | isoflavone reductase homolog IRL, putative, expressed |
| LOC_Os01g01670 | expressed protein |
| LOC_Os01g01680 | expressed protein |
| LOC_Os01g01689 | phosphatidylinositol 3- and 4-kinase family protein, expressed |
| LOC_Os01g01700 | anaphase-promoting complex subunit 11, putative, expressed |
| LOC_Os01g01710 | 1-deoxy-D-xylulose 5-phosphate reductoisomerase, chloroplast precursor, putative, expressed |
| LOC_Os01g01740 | protein kinase domain containing protein, expressed |
| LOC_Os01g01770 | ZOS1-01 - C2H2 zinc finger protein, expressed |
| LOC_Os01g01780 | exostosin family protein, putative, expressed |
| LOC_Os01g01790 | expressed protein |
| LOC_Os01g01800 | expressed protein |

| | |
|---|---|
| LOC_Os01g01810 | expressed protein |
| LOC_Os01g01840 | helix-loop-helix DNA-binding domain containing protein, expressed |
| LOC_Os01g01860 | transposon protein, putative, unclassified |
| LOC_Os01g01870 | helix-loop-helix DNA-binding domain containing protein, expressed |
| LOC_Os01g01900 | retrotransposon protein, putative, unclassified, expressed |
| LOC_Os01g01920 | HD domain containing protein 2, putative, expressed |
| LOC_Os01g01925 | expressed protein |
| LOC_Os01g01930 | expressed protein |
| LOC_Os01g01940 | expressed protein |
| LOC_Os01g01950 | expressed protein |
| LOC_Os01g01960 | transcriptional repressor, putative, expressed |
| LOC_Os01g02040 | retrotransposon protein, putative, unclassified, expressed |
| LOC_Os01g02050 | phosphoenolpyruvate carboxylase, putative, expressed |
| LOC_Os01g02060 | TOO MANY MOUTHS precursor, putative, expressed |
| LOC_Os01g02070 | invertase/pectin methylesterase inhibitor family protein, putative, expressed |
| LOC_Os01g02090 | expressed protein |
| LOC_Os01g02110 | helix-loop-helix DNA-binding domain containing protein, expressed |
| LOC_Os01g02120 | osMFT2 MFT-Like2 homologous to Mother of FT and TFL1 gene; contains Pfam profile PF01161: Phosphatidylethanolamine-binding protein, expressed |
| LOC_Os01g02150 | sulfated surface glycoprotein 185 precursor, putative, expressed |
| LOC_Os01g02170 | expressed protein |
| LOC_Os01g02180 | expressed protein |
| LOC_Os01g02190 | aquaporin protein, putative, expressed |
| LOC_Os01g02200 | armadillo/beta-catenin repeat family protein, putative, expressed |
| LOC_Os01g02240 | transposon protein, putative, unclassified, expressed |
| LOC_Os01g02380 | retrotransposon protein, putative, Ty3-gypsy subclass, expressed |
| LOC_Os01g02550 | protein kinase domain containing protein, expressed |
| LOC_Os01g02590 | receptor-like kinase, putative, expressed |
| LOC_Os01g02610 | protein kinase domain containing protein, expressed |
| LOC_Os01g02870 | auxin-induced protein 5NG4, putative, expressed |

| | |
|---|---|
| LOC_Os01g02880 | fructose-bisphospate aldolase isozyme, putative, expressed |
| LOC_Os01g02890 | phosphatidylserine synthase, putative, expressed |
| LOC_Os01g02930 | glycosyltransferase protein, putative, expressed |
| LOC_Os01g02990 | expressed protein |
| LOC_Os01g03020 | tRNA synthetase class I, putative, expressed |
| LOC_Os01g03030 | alanyl-tRNA synthetase, putative, expressed |
| LOC_Os01g03040 | expressed protein |
| LOC_Os01g03050 | fruit protein PKIWI502, putative, expressed |
| LOC_Os01g03060 | splicing factor, putative, expressed |
| LOC_Os01g03080 | expressed protein |
| LOC_Os01g03090 | methyltransferase, putative, expressed |
| LOC_Os01g03144 | white-brown complex homolog protein 7, putative, expressed |
| LOC_Os01g03160 | xylosyltransferase, putative, expressed |
| LOC_Os01g03170 | seven in absentia protein family protein, expressed |
| LOC_Os01g03210 | expressed protein |
| LOC_Os01g03280 | hypothetical protein |
| LOC_Os01g03410 | ubiquitin, putative, expressed |
| LOC_Os01g03429 | expressed protein |
| LOC_Os01g03490 | heavy-metal-associated domain-containing protein, putative, expressed |
| LOC_Os01g03500 | expressed protein |
| LOC_Os01g03510 | WD domain, G-beta repeat domain containing protein, expressed |
| LOC_Os01g03530 | multicopper oxidase domain containing protein, expressed |
| LOC_Os01g03570 | transcription factor X1, putative, expressed |
| LOC_Os01g03610 | retrotransposon protein, putative, unclassified, expressed |
| LOC_Os01g03660 | MYB family transcription factor, putative, expressed |
| LOC_Os01g03690 | TKL_IRAK_DUF26-lg |
| LOC_Os01g03710 | mannose-6-phosphate isomerase, putative, expressed |
| LOC_Os01g03720 | MYB family transcription factor, putative, expressed |
| LOC_Os01g03730 | nuclease PA3, putative, expressed |

| | |
|---|---|
| LOC_Os01g03740 | nuclease PA3, putative, expressed |
| LOC_Os01g03750 | expressed protein |
| LOC_Os01g03760 | expressed protein |
| LOC_Os01g03800 | transposon protein, putative, Mutator sub-class, expressed |
| LOC_Os01g03830 | retrotransposon protein, putative, unclassified, expressed |
| LOC_Os01g03860 | retrotransposon protein, putative, unclassified, expressed |
| LOC_Os01g03880 | retrotransposon protein, putative, Ty3-gypsy subclass |
| LOC_Os01g03900 | expressed protein |
| LOC_Os01g03914 | cation efflux family protein, putative, expressed |
| LOC_Os01g04110 | mediator of RNA polymerase II transcription subunit 22, putative, expressed |
| LOC_Os01g04230 | inactive receptor kinase At2g26730 precursor, putative, expressed |
| LOC_Os01g04450 | OsWAK2 - OsWAK receptor-like cytoplasmic kinase OsWAK-RLCK, expressed |
| LOC_Os01g04500 | expressed protein |
| LOC_Os01g04510 | expressed protein |
| LOC_Os01g04540 | serine/threonine-protein kinase At1g18390 precursor, putative, expressed |
| LOC_Os01g04680 | retrotransposon protein, putative, unclassified, expressed |
| LOC_Os01g04690 | expressed protein |
| LOC_Os01g04699 | expressed protein |
| LOC_Os01g04710 | aspartic proteinase, putative, expressed |
| LOC_Os01g04720 | leucine-rich repeat protein-related, putative, expressed |
| LOC_Os01g04750 | B3 DNA binding domain containing protein, expressed |
| LOC_Os01g04814 | vacuolar protein sorting-associating protein 4B, putative, expressed |
| LOC_Os01g04830 | expressed protein |
| LOC_Os01g04950 | peptide transporter PTR2, putative, expressed |
| LOC_Os01g05130 | expressed protein |
| LOC_Os01g05320 | retrotransposon protein, putative, Ty1-copia subclass, expressed |
| LOC_Os01g05610 | Core histone H2A/H2B/H3/H4 domain containing protein, putative, expressed |
| LOC_Os01g06280 | TKL_IRAK_CrRLK1L-1 4 - The CrRLK1L-1 subfamily has homology to the CrRLK1L homolog, expressed |
| LOC_Os01g06360 | transposon protein, putative, unclassified, expressed |

| | |
|---|---|
| LOC_Os01g06540 | PHD finger protein, putative, expressed |
| LOC_Os01g06580 | fasciclin domain containing protein, expressed |
| LOC_Os01g06610 | retrotransposon protein, putative, Ty3-gypsy subclass, expressed |
| LOC_Os01g06740 | ribosome inactivating protein, putative, expressed |
| LOC_Os01g06800 | expressed protein |
| LOC_Os01g06810 | retrotransposon protein, putative, Ty1-copia subclass, expressed |
| LOC_Os01g07470 | hypothetical protein |
| LOC_Os01g07510 | conserved hypothetical protein |
| LOC_Os01g07550 | hypothetical protein |
| LOC_Os01g07630 | BRASSINOSTEROID INSENSITIVE 1-associated receptor kinase 1 precursor, putative, expressed |
| LOC_Os01g07640 | ankyrin, putative, expressed |
| LOC_Os01g07650 | expressed protein |
| LOC_Os01g07660 | expressed protein |
| LOC_Os01g07670 | expressed protein |
| LOC_Os01g07700 | expressed protein |
| LOC_Os01g07710 | expressed protein |
| LOC_Os01g07720 | dolichyl-P-Man Man-PP-dolichyl mannosyltransferase, putative, expressed |
| LOC_Os01g07730 | phosphate/phosphoenolpyruvate translocator-related protein, putative, expressed |
| LOC_Os01g07740 | DEAD-box ATP-dependent RNA helicase 14, putative, expressed |
| LOC_Os01g07750 | expressed protein |
| LOC_Os01g07760 | phospholipase D, putative, expressed |
| LOC_Os01g07770 | peroxidase precursor, putative, expressed |
| LOC_Os01g07780 | embryo-specific 3, putative, expressed |
| LOC_Os01g07790 | polygalacturonase, putative, expressed |
| LOC_Os01g07810 | protein of unknown function domain containing protein, expressed |
| LOC_Os01g07850 | glyoxalase family protein, putative, expressed |
| LOC_Os01g07870 | ABC transporter family protein, putative, expressed |
| LOC_Os01g07890 | expressed protein |
| LOC_Os01g07900 | expressed protein |

| | |
|---|---|
| LOC_Os01g07910 | NADH-cytochrome b5 reductase, putative, expressed |
| LOC_Os01g08250 | retrotransposon protein, putative, unclassified, expressed |
| LOC_Os01g08440 | UDP-glucoronosyl and UDP-glucosyl transferase domain containing protein, expressed |
| LOC_Os01g08450 | ras-related protein, putative, expressed |
| LOC_Os01g08580 | frigida, putative, expressed |
| LOC_Os01g08610 | hAT dimerisation domain-containing protein, putative, expressed |
| LOC_Os01g08740 | expressed protein |
| LOC_Os01g09060 | retrotransposon protein, putative, Ty1-copia subclass |
| LOC_Os01g09110 | retrotransposon protein, putative, unclassified |
| LOC_Os01g09220 | transposon protein, putative, CACTA, En/Spm sub-class, expressed |
| LOC_Os01g09240 | retrotransposon protein, putative, LINE subclass |
| LOC_Os01g09440 | expressed protein |
| LOC_Os01g09450 | OsIAA2 - Auxin-responsive Aux/IAA gene family member, expressed |
| LOC_Os01g09460 | hexokinase, putative, expressed |
| LOC_Os01g09470 | IQ calmodulin-binding motif family protein, expressed |
| LOC_Os01g09500 | expressed protein |
| LOC_Os01g09510 | 60S acidic ribosomal protein, putative, expressed |
| LOC_Os01g09540 | HAD superfamily phosphatase, putative, expressed |
| LOC_Os01g09550 | no apical meristem protein, putative, expressed |
| LOC_Os01g09570 | 6-phosphofructokinase, putative, expressed |
| LOC_Os01g09720 | dof zinc finger domain containing protein, putative, expressed |
| LOC_Os01g09850 | ZOS1-04 - C2H2 zinc finger protein, expressed |
| LOC_Os01g09960 | transposon protein, putative, unclassified, expressed |
| LOC_Os01g09970 | expressed protein |
| LOC_Os01g10030 | expressed protein |
| LOC_Os01g10160 | latency associated nuclear antigen, putative, expressed |
| LOC_Os01g10370 | AP2 domain containing protein, expressed |
| LOC_Os01g10420 | expressed protein |
| LOC_Os01g10560 | transposon protein, putative, unclassified |

| | |
|---|---|
| LOC_Os01g10850 | peroxidase precursor, putative, expressed |
| LOC_Os01g10870 | CAMK_KIN1/SNF1/Nim1_like |
| LOC_Os01g10940 | S-adenosylmethionine synthetase 2, putative, expressed |
| LOC_Os01g10970 | auxin-induced protein 5NG4, putative, expressed |
| LOC_Os01g10980 | auxin-induced protein 5NG4, putative, expressed |
| LOC_Os01g11190 | expressed protein |
| LOC_Os01g11200 | transforming protein myb, putative, expressed |
| LOC_Os01g11320 | transposon protein, putative, Pong sub-class |
| LOC_Os01g11330 | retrotransposon protein, putative, unclassified, expressed |
| LOC_Os01g11360 | expressed protein |
| LOC_Os01g11650 | GDSL-like lipase/acylhydrolase, putative, expressed |
| LOC_Os01g11670 | OsSCP2 - Putative Serine Carboxypeptidase homologue, expressed |
| LOC_Os01g11680 | transposon protein, putative, unclassified, expressed |
| LOC_Os01g11690 | expressed protein |
| LOC_Os01g11820 | expressed protein |
| LOC_Os01g11830 | oryzain alpha chain precursor, putative, expressed |
| LOC_Os01g11840 | xylem cysteine proteinase 2 precursor, putative, expressed |
| LOC_Os01g11850 | expressed protein |
| LOC_Os01g12030 | endoglucanase precursor, putative, expressed |
| LOC_Os01g12040 | transposon protein, putative, unclassified, expressed |
| LOC_Os01g12070 | endoglucanase precursor, putative, expressed |
| LOC_Os01g12160 | OsGH3 3 - Probable indole-3-acetic acid-amido synthetase, expressed |
| LOC_Os01g12330 | expressed protein |
| LOC_Os01g12390 | inactive receptor kinase At2g26730 precursor, putative, expressed |
| LOC_Os01g12440 | AP2 domain containing protein, expressed |
| LOC_Os01g12860 | MYB family transcription factor, putative, expressed |
| LOC_Os01g13050 | long cell-linked locus protein, putative, expressed |
| LOC_Os01g13300 | B3 DNA binding domain containing protein, expressed |
| LOC_Os01g13370 | seven in absentia protein family domain containing protein, expressed |

| | |
|---|---|
| LOC_Os01g13620 | retrotransposon protein, putative, unclassified, expressed |
| LOC_Os01g13630 | retrotransposon protein, putative, unclassified, expressed |
| LOC_Os01g13840 | PE-PGRS family protein, putative, expressed |
| LOC_Os01g14020 | expressed protein |
| LOC_Os01g14030 | DUF260 domain containing protein, putative, expressed |
| LOC_Os01g14040 | D-tyrosyl-tRNA, putative, expressed |
| LOC_Os01g14360 | expressed protein |
| LOC_Os01g14410 | early light-induced protein, chloroplast precursor, putative, expressed |
| LOC_Os01g14580 | dehydrogenase, putative, expressed |
| LOC_Os01g14650 | expansin precursor, putative, expressed |
| LOC_Os01g14690 | Lg106, putative, expressed |
| LOC_Os01g14720 | transcription regulator, putative, expressed |
| LOC_Os01g14890 | expressed protein |
| LOC_Os01g14926 | protein kinase domain containing protein, expressed |
| LOC_Os01g15070 | retrotransposon protein, putative, unclassified |
| LOC_Os01g15080 | hypothetical protein |
| LOC_Os01g15150 | expressed protein |
| LOC_Os01g15190 | retrotransposon protein, putative, unclassified, expressed |
| LOC_Os01g15240 | expressed protein |
| LOC_Os01g15280 | transposon protein, putative, unclassified, expressed |
| LOC_Os01g15320 | RALFL9 - Rapid ALkalinization Factor RALF family protein precursor, expressed |
| LOC_Os01g15370 | retrotransposon protein, putative, Ty3-gypsy subclass |
| LOC_Os01g15420 | retrotransposon protein, putative, Ty3-gypsy subclass, expressed |
| LOC_Os01g15740 | expressed protein |
| LOC_Os01g16050 | transposon protein, putative, unclassified, expressed |
| LOC_Os01g16120 | zinc finger, C3HC4 type domain containing protein, expressed |
| LOC_Os01g16170 | PQ loop repeat domain containing protein, expressed |
| LOC_Os01g16240 | OsCam1-3 - Calmodulin, expressed |
| LOC_Os01g16370 | NBS-LRR type disease resistance protein, putative, expressed |

| | |
|---|---|
| LOC_Os01g16520 | glutamyl-tRNA synthetase, cytoplasmic, putative, expressed |
| LOC_Os01g16750 | flavin monooxygenase, putative, expressed |
| LOC_Os01g17060 | expressed protein |
| LOC_Os01g17130 | retrotransposon protein, putative, Ty3-gypsy subclass, expressed |
| LOC_Os01g17150 | plastid-specific 50S ribosomal protein 5, chloroplast precursor, putative, expressed |
| LOC_Os01g17160 | OsSubt1 - Putative Subtilisin homologue, expressed |
| LOC_Os01g17214 | major facilitator superfamily antiporter, putative, expressed |
| LOC_Os01g17279 | exonuclease, putative, expressed |
| LOC_Os01g17390 | OsFBX5 - F-box domain containing protein, expressed |
| LOC_Os01g18010 | hypothetical protein |
| LOC_Os01g18240 | MYB family transcription factor, putative, expressed |
| LOC_Os01g18250 | subtilase, putative, expressed |
| LOC_Os01g18310 | expressed protein |
| LOC_Os01g18584 | WRKY9, expressed |
| LOC_Os01g18620 | transferase family protein, putative, expressed |
| LOC_Os01g18630 | aspartic proteinase oryzasin-1 precursor, putative, expressed |
| LOC_Os01g18660 | aminotransferase, putative, expressed |
| LOC_Os01g18670 | MDR-like ABC transporter, putative, expressed |
| LOC_Os01g18730 | expressed protein |
| LOC_Os01g18800 | CAMK_KIN1/SNF1/Nim1_like 9 - CAMK includes calcium/calmodulin depedent protein kinases, expressed |
| LOC_Os01g18910 | peroxidase precursor, putative, expressed |
| LOC_Os01g18920 | hAT dimerisation domain-containing protein, putative |
| LOC_Os01g18930 | peroxidase precursor, putative, expressed |
| LOC_Os01g18970 | peroxidase precursor, putative, expressed |
| LOC_Os01g19010 | retrotransposon protein, putative, unclassified, expressed |
| LOC_Os01g19050 | transposon protein, putative, unclassified, expressed |
| LOC_Os01g19100 | retrotransposon protein, putative, Ty3-gypsy subclass, expressed |
| LOC_Os01g19130 | protein phosphatase 2C, putative, expressed |
| LOC_Os01g19310 | retrotransposon protein, putative, unclassified, expressed |

| | |
|---|---|
| LOC_Os01g19410 | expressed protein |
| LOC_Os01g19500 | expressed protein |
| LOC_Os01g19529 | pentatricopeptide, putative, expressed |
| LOC_Os01g19640 | retrotransposon protein, putative, Ty3-gypsy subclass |
| LOC_Os01g19710 | transposon protein, putative, CACTA, En/Spm sub-class, expressed |
| LOC_Os01g19720 | transposon protein, putative, CACTA, En/Spm sub-class, expressed |
| LOC_Os01g19740 | calvin cycle protein CP12, putative, expressed |
| LOC_Os01g19760 | expressed protein |
| LOC_Os01g19780 | expressed protein |
| LOC_Os01g19894 | transposon protein, putative, unclassified, expressed |
| LOC_Os01g19910 | expressed protein |
| LOC_Os01g20040 | retrotransposon protein, putative, unclassified, expressed |
| LOC_Os01g20060 | expressed protein |
| LOC_Os01g20150 | transposon protein, putative, CACTA, En/Spm sub-class, expressed |
| LOC_Os01g20860 | phospholipase D |
| LOC_Os01g20930 | RING-H2 finger protein, putative, expressed |
| LOC_Os01g20980 | pectinesterase, putative, expressed |
| LOC_Os01g21010 | expressed protein |
| LOC_Os01g21050 | pentatricopeptide, putative, expressed |
| LOC_Os01g21210 | retrotransposon protein, putative, unclassified, expressed |
| LOC_Os01g21320 | NAD dependent epimerase/dehydratase family domain containing protein, expressed |
| LOC_Os01g21360 | retrotransposon protein, putative, unclassified |
| LOC_Os01g21420 | pre-mRNA-splicing factor SF2, putative, expressed |
| LOC_Os01g21760 | retrotransposon protein, putative, unclassified |
| LOC_Os01g21900 | expressed protein |
| LOC_Os01g21960 | serine/threonine-protein kinase, putative, expressed |
| LOC_Os01g21970 | protein kinase, putative, expressed |
| LOC_Os01g21980 | expressed protein |
| LOC_Os01g21990 | CRS2-associated factor 2, chloroplast precursor, putative, expressed |

| | |
|---|---|
| LOC_Os01g22030 | expressed protein |
| LOC_Os01g22050 | retrotransposon protein, putative, unclassified, expressed |
| LOC_Os01g22090 | expressed protein |
| LOC_Os01g22140 | transposon protein, putative, CACTA, En/Spm sub-class, expressed |
| LOC_Os01g22230 | peroxidase precursor, putative, expressed |
| LOC_Os01g22309 | hypothetical protein |
| LOC_Os01g22320 | transposon protein, putative, unclassified |
| LOC_Os01g22490 | 40S ribosomal protein S27a, putative, expressed |
| LOC_Os01g22500 | craniofacial development protein 1, putative, expressed |
| LOC_Os01g22560 | glycerol-3-phosphate acyltransferase 1, putative, expressed |
| LOC_Os01g22640 | GDSL-like lipase/acylhydrolase, putative, expressed |
| LOC_Os01g22650 | expressed protein |
| LOC_Os01g22720 | expressed protein |
| LOC_Os01g22780 | GDSL-like lipase/acylhydrolase, putative, expressed |
| LOC_Os01g22830 | retrotransposon protein, putative, Ty3-gypsy subclass, expressed |
| LOC_Os01g22840 | retrotransposon protein, putative, Ty3-gypsy subclass, expressed |
| LOC_Os01g22880 | expressed protein |
| LOC_Os01g22910 | retrotransposon protein, putative, unclassified, expressed |
| LOC_Os01g22954 | serine carboxypeptidase, putative, expressed |
| LOC_Os01g23239 | retrotransposon protein, putative, Ty3-gypsy subclass, expressed |
| LOC_Os01g23420 | transposon protein, putative, unclassified, expressed |
| LOC_Os01g23650 | expressed protein |
| LOC_Os01g24030 | AMP-binding enzyme, putative, expressed |
| LOC_Os01g24070 | GATA zinc finger domain containing protein, expressed |
| LOC_Os01g24080 | expressed protein |
| LOC_Os01g24130 | transposon protein, putative, CACTA, En/Spm sub-class, expressed |
| LOC_Os01g24160 | hypothetical protein |
| LOC_Os01g24360 | retrotransposon protein, putative, unclassified |
| LOC_Os01g24450 | expressed protein |

| | |
|---|---|
| LOC_Os01g24470 | PHS1, putative, expressed |
| LOC_Os01g24700 | expressed protein |
| LOC_Os01g24760 | expressed protein |
| LOC_Os01g24880 | zinc finger, C3HC4 type domain containing protein, expressed |
| LOC_Os01g24890 | transposon protein, putative, CACTA, En/Spm sub-class, expressed |
| LOC_Os01g24910 | transposon protein, putative, CACTA, En/Spm sub-class, expressed |
| LOC_Os01g24920 | poly synthetase 2-B, putative, expressed |
| LOC_Os01g24970 | expressed protein |
| LOC_Os01g24980 | naringenin,2-oxoglutarate 3-dioxygenase, putative, expressed |
| LOC_Os01g25080 | expressed protein |
| LOC_Os01g25150 | jacalin-like lectin domain containing protein |
| LOC_Os01g25160 | jacalin-like lectin domain containing protein, expressed |
| LOC_Os01g25300 | expressed protein |
| LOC_Os01g25330 | SAC9, putative, expressed |
| LOC_Os01g25510 | alpha-amylase isozyme C, putative, expressed |
| LOC_Os01g25960 | retrotransposon protein, putative, Ty1-copia subclass, expressed |
| LOC_Os01g25990 | expressed protein |
| LOC_Os01g26120 | membrane associated DUF588 domain containing protein, putative, expressed |
| LOC_Os01g27070 | expressed protein |
| LOC_Os01g27230 | 12-oxophytodienoate reductase, putative, expressed |
| LOC_Os01g27270 | retrotransposon protein, putative, unclassified, expressed |
| LOC_Os01g27320 | retrotransposon protein, putative, Ty3-gypsy subclass, expressed |
| LOC_Os01g27430 | retrotransposon protein, putative, unclassified, expressed |
| LOC_Os01g27700 | laccase precursor protein, putative, expressed |
| LOC_Os01g27720 | E3 ubiquitin-protein ligase MGRN1, putative, expressed |
| LOC_Os01g27834 | hypothetical protein |
| LOC_Os01g27880 | sucrose-phosphatase, putative, expressed |
| LOC_Os01g27940 | cyclin-L1-1, putative, expressed |
| LOC_Os01g27970 | hypothetical protein |

| | |
|---|---|
| LOC_Os01g28010 | retrotransposon protein, putative, Ty3-gypsy subclass |
| LOC_Os01g28020 | retrotransposon protein, putative, Ty3-gypsy subclass |
| LOC_Os01g28200 | transposon protein, putative, unclassified, expressed |
| LOC_Os01g28270 | retrotransposon protein, putative, Ty1-copia subclass |
| LOC_Os01g28660 | expressed protein |
| LOC_Os01g28680 | WD domain, G-beta repeat domain containing protein, expressed |
| LOC_Os01g28720 | hypothetical protein |
| LOC_Os01g28882 | retrotransposon protein, putative, unclassified, expressed |
| LOC_Os01g28980 | proton-dependent oligopeptide transport, putative, expressed |
| LOC_Os01g29030 | retrotransposon protein, putative, unclassified, expressed |
| LOC_Os01g29040 | retrotransposon protein, putative, unclassified, expressed |
| LOC_Os01g29050 | retrotransposon protein, putative, unclassified, expressed |
| LOC_Os01g29290 | retrotransposon protein, putative, Ty3-gypsy subclass, expressed |
| LOC_Os01g29550 | expressed protein |
| LOC_Os01g29570 | retrotransposon protein, putative, Ty3-gypsy subclass, expressed |
| LOC_Os01g29680 | retrotransposon protein, putative, Ty1-copia subclass, expressed |
| LOC_Os01g29760 | retrotransposon protein, putative, unclassified, expressed |
| LOC_Os01g29820 | expressed protein |
| LOC_Os01g29830 | expressed protein |
| LOC_Os01g29870 | multidrug resistance protein 9, putative, expressed |
| LOC_Os01g31020 | retrotransposon protein, putative, unclassified, expressed |
| LOC_Os01g31050 | expressed protein |
| LOC_Os01g31140 | hypothetical protein |
| LOC_Os01g31230 | retrotransposon, putative, centromere-specific, expressed |
| LOC_Os01g31280 | expressed protein |
| LOC_Os01g31310 | expressed protein |
| LOC_Os01g31450 | retrotransposon protein, putative, unclassified, expressed |
| LOC_Os01g31560 | expressed protein |
| LOC_Os01g31629 | expressed protein |

| | |
|---|---|
| LOC_Os01g31750 | transposon protein, putative, unclassified, expressed |
| LOC_Os01g31800 | Core histone H2A/H2B/H3/H4 domain containing protein, putative, expressed |
| LOC_Os01g31870 | natural resistance-associated macrophage protein, putative, expressed |
| LOC_Os01g32000 | retrotransposon protein, putative, unclassified, expressed |
| LOC_Os01g32080 | thiamine pyrophosphate enzyme, C-terminal TPP binding domain containing protein, expressed |
| LOC_Os01g32120 | OsCML11 - Calmodulin-related calcium sensor protein, expressed |
| LOC_Os01g32150 | expressed protein |
| LOC_Os01g32200 | hypothetical protein |
| LOC_Os01g32300 | expressed protein |
| LOC_Os01g32310 | expressed protein |
| LOC_Os01g32364 | Os1bglu1 - beta-mannosidase/glucosidase homologue, expressed |
| LOC_Os01g32470 | retrotransposon protein, putative, unclassified |
| LOC_Os01g32500 | retrotransposon protein, putative, unclassified, expressed |
| LOC_Os01g32530 | transposon protein, putative, CACTA, En/Spm sub-class |
| LOC_Os01g32550 | hypothetical protein |
| LOC_Os01g32610 | expressed protein |
| LOC_Os01g32650 | expressed protein |
| LOC_Os01g32810 | expressed protein |
| LOC_Os01g33090 | expressed protein |
| LOC_Os01g33136 | retrotransposon protein, putative, unclassified, expressed |
| LOC_Os01g33142 | hypothetical protein |
| LOC_Os01g33180 | hypothetical protein |
| LOC_Os01g33470 | retrotransposon protein, putative, unclassified, expressed |
| LOC_Os01g33615 | expressed protein |
| LOC_Os01g33740 | retrotransposon protein, putative, unclassified, expressed |
| LOC_Os01g33762 | expressed protein |
| LOC_Os01g33960 | transposon protein, putative, CACTA, En/Spm sub-class, expressed |
| LOC_Os01g33970 | transposon protein, putative, CACTA, En/Spm sub-class, expressed |
| LOC_Os01g33980 | expressed protein |

| | |
|---|---|
| LOC_Os01g34030 | retrotransposon protein, putative, unclassified, expressed |
| LOC_Os01g34130 | retrotransposon protein, putative, Ty3-gypsy subclass |
| LOC_Os01g34460 | expressed protein |
| LOC_Os01g34470 | tetratricopeptide repeat containing protein, putative, expressed |
| LOC_Os01g34480 | NAD dependent epimerase/dehydratase family protein, putative, expressed |
| LOC_Os01g34510 | hypothetical protein |
| LOC_Os01g34520 | expressed protein |
| LOC_Os01g34560 | 3-ketoacyl-CoA synthase, putative, expressed |
| LOC_Os01g34620 | OsGrx_S15 |
| LOC_Os01g34710 | transposon protein, putative, unclassified, expressed |
| LOC_Os01g34760 | retrotransposon protein, putative, Ty3-gypsy subclass, expressed |
| LOC_Os01g34910 | expressed protein |
| LOC_Os01g35050 | early-responsive to dehydration protein-related, putative, expressed |
| LOC_Os01g35660 | retrotransposon protein, putative, Ty3-gypsy subclass, expressed |
| LOC_Os01g35880 | expressed protein |
| LOC_Os01g36120 | expressed protein |
| LOC_Os01g36340 | expressed protein |
| LOC_Os01g37280 | expressed protein |
| LOC_Os01g37600 | agenet domain containing protein, expressed |
| LOC_Os01g37630 | esterase, putative, expressed |
| LOC_Os01g37670 | OsFBX15 - F-box domain containing protein, expressed |
| LOC_Os01g37690 | sodium/calcium exchanger protein, putative, expressed |
| LOC_Os01g37700 | expressed protein |
| LOC_Os01g37825 | M16 domain containing zinc peptidase, putative, expressed |
| LOC_Os01g37837 | seryl-tRNA synthetase, putative, expressed |
| LOC_Os01g37910 | vacuolar-processing enzyme precursor, putative, expressed |
| LOC_Os01g37990 | expressed protein |
| LOC_Os01g38100 | expressed protein |
| LOC_Os01g38110 | cytochrome P450, putative, expressed |

| | |
|---|---|
| LOC_Os01g38310 | expressed protein |
| LOC_Os01g38400 | retrotransposon protein, putative, Ty3-gypsy subclass, expressed |
| LOC_Os01g38520 | expressed protein |
| LOC_Os01g38530 | ELF3 protein, putative, expressed |
| LOC_Os01g38620 | 3-5 exonuclease/ nucleic acid binding protein, putative, expressed |
| LOC_Os01g38940 | expressed protein |
| LOC_Os01g39100 | zinc finger C-x8-C-x5-C-x3-H type family protein, expressed |
| LOC_Os01g39136 | retrotransposon protein, putative, unclassified, expressed |
| LOC_Os01g39280 | expressed protein |
| LOC_Os01g39350 | protein kinase domain containing protein, expressed |
| LOC_Os01g39380 | SCD1, putative, expressed |
| LOC_Os01g39700 | expressed protein |
| LOC_Os01g39810 | Alg9-like mannosyltransferase protein, putative, expressed |
| LOC_Os01g39860 | 1-aminocyclopropane-1-carboxylate oxidase protein, putative, expressed |
| LOC_Os01g39870 | expressed protein |
| LOC_Os01g39980 | hypothetical protein |
| LOC_Os01g40000 | expressed protein |
| LOC_Os01g40180 | expressed protein |
| LOC_Os01g40499 | S-locus lectin protein kinase family protein, putative, expressed |
| LOC_Os01g40540 | lectin receptor-type protein kinase, putative, expressed |
| LOC_Os01g40910 | transposon protein, putative, unclassified |
| LOC_Os01g41030 | ribosomal protein L25, putative, expressed |
| LOC_Os01g41100 | retrotransposon protein, putative, unclassified, expressed |
| LOC_Os01g41110 | expressed protein |
| LOC_Os01g41145 | retrotransposon protein, putative, unclassified, expressed |
| LOC_Os01g41250 | OsFBX17 - F-box domain containing protein, expressed |
| LOC_Os01g41610 | mitochondrial ATP synthase g subunit family protein, putative, expressed |
| LOC_Os01g41890 | MLA1, putative, expressed |
| LOC_Os01g41960 | retrotransposon protein, putative, unclassified, expressed |

| | |
|---|---|
| LOC_Os01g42140 | expressed protein |
| LOC_Os01g42320 | aspartic proteinase nepenthesin-2 precursor, putative, expressed |
| LOC_Os01g42400 | STE_MEK_ste7_MAP2K |
| LOC_Os01g42610 | terpene synthase, putative, expressed |
| LOC_Os01g42890 | expressed protein |
| LOC_Os01g42990 | PPR repeat domain containing protein, putative, expressed |
| LOC_Os01g43290 | expressed protein |
| LOC_Os01g44010 | retrotransposon protein, putative, unclassified, expressed |
| LOC_Os01g44090 | X8 domain containing protein, expressed |
| LOC_Os01g44200 | retrotransposon protein, putative, unclassified, expressed |
| LOC_Os01g45040 | Leucine rich repeat N-terminal domain containing protein, putative, expressed |
| LOC_Os01g45480 | retrotransposon protein, putative, unclassified, expressed |
| LOC_Os01g45540 | retrotransposon protein, putative, unclassified, expressed |
| LOC_Os01g45550 | auxin efflux carrier component, putative, expressed |
| LOC_Os01g45670 | expressed protein |
| LOC_Os01g45820 | expressed protein |
| LOC_Os01g45880 | retrotransposon protein, putative, unclassified, expressed |
| LOC_Os01g46169 | GDSL-like lipase/acylhydrolase, putative, expressed |
| LOC_Os01g46200 | expressed protein |
| LOC_Os01g46290 | lipase, putative, expressed |
| LOC_Os01g46300 | hypothetical protein |
| LOC_Os01g46400 | expressed protein |
| LOC_Os01g46430 | glycosyl transferase, group 1 domain containing protein, expressed |
| LOC_Os01g46530 | expressed protein |
| LOC_Os01g46580 | actin-related protein 2/3 complex subunit 2, putative, expressed |
| LOC_Os01g46720 | expressed protein |
| LOC_Os01g46860 | inorganic phosphate transporter, putative, expressed |
| LOC_Os01g47300 | retrotransposon protein, putative, unclassified, expressed |
| LOC_Os01g47680 | 60S ribosomal protein L18a-1, putative, expressed |

| | |
|---|---|
| LOC_Os01g47710 | homeobox domain containing protein, expressed |
| LOC_Os01g47730 | ras-related protein, putative, expressed |
| LOC_Os01g47770 | CS domain containing protein, putative, expressed |
| LOC_Os01g47870 | retrotransposon protein, putative, unclassified, expressed |
| LOC_Os01g48000 | S-locus-like receptor protein kinase, putative, expressed |
| LOC_Os01g48440 | glycosyltransferase family 43 protein, putative, expressed |
| LOC_Os01g48525 | expressed protein |
| LOC_Os01g48580 | ubiquitin-conjugating enzyme E2, putative, expressed |
| LOC_Os01g48650 | expressed protein |
| LOC_Os01g48720 | RNA polymerase III subunit RPC82 family protein, putative, expressed |
| LOC_Os01g48830 | expressed protein |
| LOC_Os01g48900 | dynamin-related protein 1A, putative, expressed |
| LOC_Os01g48930 | splicing factor-related, putative, expressed |
| LOC_Os01g49010 | ORC4 - Putative origin recognition complex subunit 4, expressed |
| LOC_Os01g49026 | transmembrane protein 93, putative, expressed |
| LOC_Os01g49030 | expressed protein |
| LOC_Os01g49065 | expressed protein |
| LOC_Os01g49090 | retrotransposon protein, putative, unclassified, expressed |
| LOC_Os01g49154 | expressed protein |
| LOC_Os01g49420 | retrotransposon protein, putative, unclassified, expressed |
| LOC_Os01g49750 | expressed protein |
| LOC_Os01g50020 | expressed protein |
| LOC_Os01g50680 | OsSub2 - Putative Subtilisin homologue, expressed |
| LOC_Os01g50690 | WD domain, G-beta repeat domain containing protein, expressed |
| LOC_Os01g50710 | hypothetical protein |
| LOC_Os01g50780 | expressed protein |
| LOC_Os01g51330 | expressed protein |
| LOC_Os01g51450 | NAP domain containing protein, putative, expressed |
| LOC_Os01g51650 | retrotransposon protein, putative, unclassified, expressed |

| | |
|---|---|
| LOC_Os01g51735 | expressed protein |
| LOC_Os01g52020 | subtilase, putative, expressed |
| LOC_Os01g52070 | potassium channel AKT1, putative, expressed |
| LOC_Os01g52140 | expressed protein |
| LOC_Os01g52214 | ubiquinone oxidoreductase, putative, expressed |
| LOC_Os01g52250 | starch synthase, putative, expressed |
| LOC_Os01g52450 | hexokinase, putative, expressed |
| LOC_Os01g52750 | OsSub3 - Putative Subtilisin homologue, expressed |
| LOC_Os01g52910 | expressed protein |
| LOC_Os01g52920 | expressed protein |
| LOC_Os01g53040 | WRKY14, expressed |
| LOC_Os01g53294 | respiratory burst oxidase protein B, putative, expressed |
| LOC_Os01g53350 | anthocyanidin 5,3-O-glucosyltransferase, putative, expressed |
| LOC_Os01g53520 | reticulon domain containing protein, putative, expressed |
| LOC_Os01g53720 | expressed protein |
| LOC_Os01g53740 | hypothetical protein |
| LOC_Os01g53750 | glucan endo-1,3-beta-glucosidase precursor, putative, expressed |
| LOC_Os01g53760 | expressed protein |
| LOC_Os01g53900 | elongation factor, putative, expressed |
| LOC_Os01g54060 | expressed protein |
| LOC_Os01g54240 | nucleoporin, putative, expressed |
| LOC_Os01g54300 | OsMan02 - Endo-Beta-Mannanase, expressed |
| LOC_Os01g55140 | expressed protein |
| LOC_Os01g55200 | potassium channel KAT1, putative, expressed |
| LOC_Os01g55330 | embryonic protein DC-8, putative, expressed |
| LOC_Os01g55795 | expressed protein |
| LOC_Os01g56060 | hypothetical protein |
| LOC_Os01g56280 | transposon protein, putative, unclassified |
| LOC_Os01g56580 | CK1_CaseinKinase_1a 3 - CK1 includes the casein kinase 1 kinases, expressed |

| | |
|---|---|
| LOC_Os01g57220 | secretory carrier-associated membrane protein, putative, expressed |
| LOC_Os01g57350 | diacylglycerol kinase, putative, expressed |
| LOC_Os01g57530 | expressed protein |
| LOC_Os01g57599 | retrotransposon protein, putative, unclassified, expressed |
| LOC_Os01g57954 | translation initiation factor IF-1, chloroplast, putative, expressed |
| LOC_Os01g57962 | photosystem I P700 chlorophyll a apoprotein A2, putative, expressed |
| LOC_Os01g57964 | photosystem I P700 chlorophyll a apoprotein A1, putative, expressed |
| LOC_Os01g58022 | NADPH-dependent oxidoreductase, putative, expressed |
| LOC_Os01g58024 | NADPH-dependent oxidoreductase, putative, expressed |
| LOC_Os01g58160 | expressed protein |
| LOC_Os01g58230 | expressed protein |
| LOC_Os01g58300 | expressed protein |
| LOC_Os01g58330 | expressed protein |
| LOC_Os01g58870 | domain of unknown function, DUF250 domain containing protein, expressed |
| LOC_Os01g58950 | cytochrome P450, putative, expressed |
| LOC_Os01g58970 | cytochrome P450, putative, expressed |
| LOC_Os01g59100 | cytokinin-N-glucosyltransferase 1, putative, expressed |
| LOC_Os01g59110 | indole-3-acetate beta-glucosyltransferase, putative, expressed |
| LOC_Os01g59360 | CAMK_CAMK_like |
| LOC_Os01g59420 | expressed protein |
| LOC_Os01g59530 | OsCML1 - Calmodulin-related calcium sensor protein, expressed |
| LOC_Os01g60020 | NAC domain transcription factor, putative, expressed |
| LOC_Os01g60140 | ATCHX15, putative, expressed |
| LOC_Os01g60150 | expressed protein |
| LOC_Os01g60549 | retrotransposon protein, putative, unclassified, expressed |
| LOC_Os01g60560 | retrotransposon protein, putative, unclassified, expressed |
| LOC_Os01g60760 | expressed protein |
| LOC_Os01g60770 | expansin precursor, putative, expressed |
| LOC_Os01g60850 | expressed protein |

| | |
|---|---|
| LOC_Os01g60860 | spotted leaf 11, putative, expressed |
| LOC_Os01g61020 | curculin-like lectin family protein, putative, expressed |
| LOC_Os01g61030 | VHS and GAT domain containing protein, expressed |
| LOC_Os01g61060 | SAG20, putative, expressed |
| LOC_Os01g61160 | laccase precursor protein, putative, expressed |
| LOC_Os01g61170 | prenylated rab acceptor, putative, expressed |
| LOC_Os01g61380 | lactate/malate dehydrogenase, putative, expressed |
| LOC_Os01g61460 | leaf senescence related protein, putative, expressed |
| LOC_Os01g61880 | respiratory burst oxidase, putative, expressed |
| LOC_Os01g61940 | white-brown complex homolog protein, putative, expressed |
| LOC_Os01g62290 | DnaK family protein, putative, expressed |
| LOC_Os01g62310 | homeobox domain containing protein, expressed |
| LOC_Os01g62370 | ENTH domain containing protein, expressed |
| LOC_Os01g62710 | transposon protein, putative, unclassified |
| LOC_Os01g62840 | mannose-1-phosphate guanyltransferase, putative, expressed |
| LOC_Os01g63040 | expressed protein |
| LOC_Os01g63170 | transposon protein, putative, unclassified, expressed |
| LOC_Os01g63180 | laccase-6 precursor, putative, expressed |
| LOC_Os01g63770 | transmembrane amino acid transporter protein, putative, expressed |
| LOC_Os01g63900 | WD domain, G-beta repeat domain containing protein, expressed |
| LOC_Os01g63980 | ZOS1-17 - C2H2 zinc finger protein, expressed |
| LOC_Os01g64170 | glycosyl hydrolases family 17, putative, expressed |
| LOC_Os01g64330 | retrotransposon protein, putative, unclassified, expressed |
| LOC_Os01g64520 | uricase, putative, expressed |
| LOC_Os01g64630 | actin, putative, expressed |
| LOC_Os01g64640 | histone H3, putative, expressed |
| LOC_Os01g64700 | OsFBT3 - F-box and tubby domain containing protein, expressed |
| LOC_Os01g64790 | AP2 domain containing protein, expressed |
| LOC_Os01g64860 | OsSub11 - Putative Subtilisin homologue, expressed |

| | |
|---|---|
| LOC_Os01g64870 | expressed protein |
| LOC_Os01g64880 | expressed protein |
| LOC_Os01g65130 | peptide transporter, putative, expressed |
| LOC_Os01g65370 | MYB family transcription factor, putative, expressed |
| LOC_Os01g65410 | serine hydroxymethyltransferase, mitochondrial precursor, putative, expressed |
| LOC_Os01g65470 | conserved hypothetical protein |
| LOC_Os01g65490 | DNA binding protein, putative, expressed |
| LOC_Os01g65530 | RNA recognition motif containing protein, putative, expressed |
| LOC_Os01g65640 | expressed protein |
| LOC_Os01g65660 | amino acid transporter, putative, expressed |
| LOC_Os01g65670 | amino acid transporter, putative, expressed |
| LOC_Os01g65680 | 4,5-DOPA dioxygenase extradiol, putative, expressed |
| LOC_Os01g65750 | transposon protein, putative, unclassified, expressed |
| LOC_Os01g65850 | CHD3-type chromatin-remodeling factor PICKLE, putative, expressed |
| LOC_Os01g65860 | ANTH domain containing protein, expressed |
| LOC_Os01g65902 | apocytochrome f precursor, putative, expressed |
| LOC_Os01g65920 | F-box/LRR-repeat protein 2, putative, expressed |
| LOC_Os01g66010 | amino acid transporter, putative, expressed |
| LOC_Os01g66020 | protein kinase family protein, putative, expressed |
| LOC_Os01g66100 | gibberellin 20 oxidase 2, putative, expressed |
| LOC_Os01g66110 | methyltransferase, putative, expressed |
| LOC_Os01g66360 | 2-C-methyl-D-erythritol 4-phosphate cytidylyltransferase, putative, expressed |
| LOC_Os01g66590 | DUF260 domain containing protein, putative, expressed |
| LOC_Os01g66700 | beta-hexosaminidase precursor, putative, expressed |
| LOC_Os01g66710 | polygalacturonase, putative, expressed |
| LOC_Os01g67240 | formin-like protein 1 precursor, putative, expressed |
| LOC_Os01g67620 | expressed protein |
| LOC_Os01g67760 | expressed protein |
| LOC_Os01g67850 | zinc finger, RING-type, putative, expressed |

| | |
|---|---|
| LOC_Os01g67860 | fructose-bisphospate aldolase isozyme, putative, expressed |
| LOC_Os01g67940 | retrotransposon, putative, centromere-specific |
| LOC_Os01g68070 | PPR repeat containing protein, expressed |
| LOC_Os01g68260 | lysine ketoglutarate reductase trans-splicing related 1, putative, expressed |
| LOC_Os01g68340 | retrotransposon protein, putative, unclassified, expressed |
| LOC_Os01g68860 | zinc finger C-x8-C-x5-C-x3-H type family protein, expressed |
| LOC_Os01g68990 | retrotransposon, putative, centromere-specific, expressed |
| LOC_Os01g69230 | exo70 exocyst complex subunit domain containing protein, expressed |
| LOC_Os01g69900 | CBS domain-containing protein, putative, expressed |
| LOC_Os01g69930 | dnaJ domain containing protein, expressed |
| LOC_Os01g69940 | OsFBX32 - F-box domain containing protein, expressed |
| LOC_Os01g70060 | protein of unknown function, DUF618 domain containing protein, expressed |
| LOC_Os01g70130 | protein kinase domain containing protein, expressed |
| LOC_Os01g70310 | inducer of CBF expression 2, putative, expressed |
| LOC_Os01g70450 | expressed protein |
| LOC_Os01g70540 | retrotransposon protein, putative, unclassified, expressed |
| LOC_Os01g70550 | heparan-alpha-glucosaminide N-acetyltransferase, putative, expressed |
| LOC_Os01g70880 | histone-like transcription factor and archaeal histone, putative, expressed |
| LOC_Os01g71020 | expressed protein |
| LOC_Os01g71150 | expressed protein |
| LOC_Os01g71170 | classical arabinogalactan protein 26 precursor, putative, expressed |
| LOC_Os01g71390 | retrotransposon protein, putative, unclassified, expressed |
| LOC_Os01g71490 | expressed protein |
| LOC_Os01g71540 | expressed protein |
| LOC_Os01g71620 | expressed protein |
| LOC_Os01g71640 | expressed protein |
| LOC_Os01g71840 | retrotransposon protein, putative, unclassified, expressed |
| LOC_Os01g72130 | glutathione S-transferase, putative, expressed |
| LOC_Os01g72200 | expressed protein |

| | |
|---|---|
| LOC_Os01g72260 | cytochrome P450, putative, expressed |
| LOC_Os01g72834 | RNA recognition motif containing protein, putative, expressed |
| LOC_Os01g73250 | abscisic stress-ripening, putative, expressed |
| LOC_Os01g73310 | actin, putative, expressed |
| LOC_Os01g73332 | retrotransposon protein, putative, Ty3-gypsy subclass |
| LOC_Os01g73390 | expressed protein |
| LOC_Os01g73410 | FYVE zinc finger domain containing protein, expressed |
| LOC_Os01g73514 | 5-oxoprolinase, putative, expressed |
| LOC_Os01g73530 | ABC transporter, ATP-binding protein, putative, expressed |
| LOC_Os01g73590 | transporter family protein, putative, expressed |
| LOC_Os01g73620 | expressed protein |
| LOC_Os01g73700 | RCLEA2 - Root cap and Late embryogenesis related family protein precursor, putative, expressed |
| LOC_Os01g73900 | DEAD-box ATP-dependent RNA helicase, putative, expressed |
| LOC_Os01g73910 | peptidase, putative, expressed |
| LOC_Os01g74030 | pumilio-family RNA binding protein, putative, expressed |
| LOC_Os01g74200 | protein kinase domain containing protein, expressed |
| LOC_Os01g74350 | soluble inorganic pyrophosphatase, putative, expressed |
| LOC_Os01g74450 | aquaporin protein, putative, expressed |
| LOC_Os01g74550 | transposon protein, putative, unclassified, expressed |
| LOC_Os01g74590 | MYB family transcription factor, putative, expressed |
| LOC_Os01g74610 | zinc finger, C3HC4 type domain containing protein, expressed |
| LOC_Os02g01020 | expressed protein |
| LOC_Os02g01030 | tetratricopeptide repeat domain containing protein, expressed |
| LOC_Os02g01060 | harpin-induced protein 1 domain containing protein, expressed |
| LOC_Os02g01070 | FG-GAP repeat-containing protein, putative, expressed |
| LOC_Os02g01180 | kinesin motor domain containing protein, putative, expressed |
| LOC_Os02g01190 | POEI25 - Pollen Ole e I allergen and extensin family protein precursor, expressed |
| LOC_Os02g01220 | rhodanese-like domain containing protein, putative, expressed |
| LOC_Os02g01250 | LSM domain containing protein, expressed |

| | |
|---|---|
| LOC_Os02g01270 | START domain containing protein, expressed |
| LOC_Os02g01280 | T-complex protein, putative, expressed |
| LOC_Os02g01290 | expressed protein |
| LOC_Os02g01300 | invertase/pectin methylesterase inhibitor family protein, putative, expressed |
| LOC_Os02g01310 | invertase/pectin methylesterase inhibitor family protein, putative, expressed |
| LOC_Os02g01326 | auxin-induced protein 5NG4, putative, expressed |
| LOC_Os02g01332 | ribosomal protein L6, putative, expressed |
| LOC_Os02g01340 | ferredoxin--NADP reductase, chloroplast precursor, putative, expressed |
| LOC_Os02g01355 | MADS-box transcription factor, putative, expressed |
| LOC_Os02g01380 | expressed protein |
| LOC_Os02g01500 | 2-oxo acid dehydrogenases acyltransferase domain containing protein, expressed |
| LOC_Os02g01510 | lactate/malate dehydrogenase, putative, expressed |
| LOC_Os02g01610 | pentatricopeptide, putative, expressed |
| LOC_Os02g01700 | RNA recognition motif containing protein, putative, expressed |
| LOC_Os02g01770 | expressed protein |
| LOC_Os02g01940 | transcription factor jumonji, putative, expressed |
| LOC_Os02g01960 | tetratricopeptide repeat domain containing protein, expressed |
| LOC_Os02g02050 | SNF2 family N-terminal domain containing protein, expressed |
| LOC_Os02g02090 | peptidyl-prolyl cis-trans isomerase, putative, expressed |
| LOC_Os02g02130 | ribosomal protein L37, putative, expressed |
| LOC_Os02g02140 | receptor protein kinase CLAVATA1 precursor, putative, expressed |
| LOC_Os02g02210 | aminotransferase, putative, expressed |
| LOC_Os02g02230 | cytochrome P450, putative, expressed |
| LOC_Os02g02290 | SNF2 family N-terminal domain containing protein, expressed |
| LOC_Os02g02310 | auxin-induced protein 5NG4, putative, expressed |
| LOC_Os02g02340 | glycerol-3-phosphate acyltransferase, putative, expressed |
| LOC_Os02g02424 | ZOS2-02 - C2H2 zinc finger protein, expressed |
| LOC_Os02g02470 | oxidoreductase, short chain dehydrogenase/reductase family domain containing protein, expressed |
| LOC_Os02g02490 | phytosulfokine receptor precursor, putative, expressed |

| | |
|---|---|
| LOC_Os02g02524 | canopy homolog 2 precursor, putative, expressed |
| LOC_Os02g02580 | expressed protein |
| LOC_Os02g02590 | PPR repeat domain containing protein, putative, expressed |
| LOC_Os02g02690 | expressed protein |
| LOC_Os02g02710 | zinc carboxypeptidase family protein, putative, expressed |
| LOC_Os02g02720 | SNARE domain containing protein, putative, expressed |
| LOC_Os02g02750 | CDP-alcohol phosphatidyltransferase, putative, expressed |
| LOC_Os02g02770 | PPR repeat domain containing protein, putative, expressed |
| LOC_Os02g02780 | protein kinase family protein, putative, expressed |
| LOC_Os02g02790 | C2 domain containing protein, expressed |
| LOC_Os02g02820 | helix-loop-helix DNA-binding domain containing protein, expressed |
| LOC_Os02g02850 | bifunctional protein folD, putative, expressed |
| LOC_Os02g02860 | glutamyl-tRNA synthetase, putative, expressed |
| LOC_Os02g02880 | expressed protein |
| LOC_Os02g02900 | expressed protein |
| LOC_Os02g02910 | expressed protein |
| LOC_Os02g02920 | HVA22, putative, expressed |
| LOC_Os02g02940 | succinate dehydrogenase subunit 3, putative, expressed |
| LOC_Os02g02950 | pentatricopeptide, putative, expressed |
| LOC_Os02g02960 | histone deacetylase complex subunit SAP18, putative, expressed |
| LOC_Os02g02970 | expressed protein |
| LOC_Os02g02980 | MATE domain containing protein, expressed |
| LOC_Os02g03000 | division protein, putative, expressed |
| LOC_Os02g03010 | expressed protein |
| LOC_Os02g03020 | EF hand family protein, putative, expressed |
| LOC_Os02g03050 | dimethyladenosine transferase, putative, expressed |
| LOC_Os02g03070 | SSE1, putative, expressed |
| LOC_Os02g03100 | oxidoreductase, aldo/keto reductase family protein, putative, expressed |
| LOC_Os02g03110 | CDP-alcohol phosphatidyltransferase, putative, expressed |

| | |
|---|---|
| LOC_Os02g03130 | expressed protein |
| LOC_Os02g03200 | FAD-binding and arabino-lactone oxidase domains containing protein, putative, expressed |
| LOC_Os02g03230 | membrane related protein CP5, putative, expressed |
| LOC_Os02g03250 | expressed protein |
| LOC_Os02g03260 | 3-isopropylmalate dehydratase large subunit 2, putative, expressed |
| LOC_Os02g03270 | AT hook motif domain containing protein, expressed |
| LOC_Os02g03280 | transmembrane BAX inhibitor motif-containing protein, putative, expressed |
| LOC_Os02g03294 | cyclin, putative, expressed |
| LOC_Os02g03310 | KH domain containing protein, putative, expressed |
| LOC_Os02g03360 | expressed protein |
| LOC_Os02g03410 | CAMK_CAMK_like 12 - CAMK includes calcium/calmodulin depedent protein kinases, expressed |
| LOC_Os02g03430 | expressed protein |
| LOC_Os02g03440 | LSM domain containing protein, expressed |
| LOC_Os02g03450 | vesicle transport v-SNARE protein, putative, expressed |
| LOC_Os02g03480 | THION24 - Plant thionin family protein precursor, expressed |
| LOC_Os02g03490 | major ampullate spidroin 3, putative, expressed |
| LOC_Os02g03640 | expressed protein |
| LOC_Os02g03650 | expressed protein |
| LOC_Os02g03750 | polygalacturonase, putative, expressed |
| LOC_Os02g04390 | expressed protein |
| LOC_Os02g04810 | auxin response factor 5, putative, expressed |
| LOC_Os02g04950 | splicing factor 3B subunit 1, putative, expressed |
| LOC_Os02g05060 | OsSAUR5 - Auxin-responsive SAUR gene family member, expressed |
| LOC_Os02g05244 | expressed protein |
| LOC_Os02g05280 | hypothetical protein |
| LOC_Os02g05300 | expressed protein |
| LOC_Os02g05340 | proteasome/cyclosome repeat containing protein, expressed |
| LOC_Os02g05470 | CCT motif family protein, expressed |
| LOC_Os02g05510 | GATA transcription factor 25, putative, expressed |

| | |
|---|---|
| LOC_Os02g05520 | expressed protein |
| LOC_Os02g05610 | ZOS2-03 - C2H2 zinc finger protein, expressed |
| LOC_Os02g05630 | protein phosphatase 2C, putative, expressed |
| LOC_Os02g05640 | homeobox associated leucine zipper, putative, expressed |
| LOC_Os02g05650 | expressed protein |
| LOC_Os02g05670 | expressed protein |
| LOC_Os02g05692 | zinc finger, C3HC4 type domain containing protein, expressed |
| LOC_Os02g05700 | OsFBO8 - F-box and other domain containing protein, expressed |
| LOC_Os02g05720 | pentatricopeptide repeat protein PPR986-12, putative, expressed |
| LOC_Os02g05730 | protein kinase domain containing protein, expressed |
| LOC_Os02g05790 | expressed protein |
| LOC_Os02g05800 | expressed protein |
| LOC_Os02g05810 | expressed protein |
| LOC_Os02g05820 | protein kinase domain containing protein, expressed |
| LOC_Os02g05830 | ribulose bisphosphate carboxylase small chain, chloroplast precursor, putative, expressed |
| LOC_Os02g05840 | VIL2 protein, putative, expressed |
| LOC_Os02g05870 | expressed protein |
| LOC_Os02g05880 | RNA polymerase Rpb1, domain 2 family protein, expressed |
| LOC_Os02g06100 | expressed protein |
| LOC_Os02g06540 | transporter family protein, putative, expressed |
| LOC_Os02g06930 | protein kinase, putative, expressed |
| LOC_Os02g06940 | expressed protein |
| LOC_Os02g06950 | no apical meristem protein, putative, expressed |
| LOC_Os02g06970 | expressed protein |
| LOC_Os02g07120 | HEAT repeat family protein, putative, expressed |
| LOC_Os02g07130 | expressed protein |
| LOC_Os02g07180 | expressed protein |
| LOC_Os02g07190 | expressed protein |
| LOC_Os02g07210 | OTU-like cysteine protease family protein, putative, expressed |

| | |
|---|---|
| LOC_Os02g07300 | hypothetical protein |
| LOC_Os02g07320 | expressed protein |
| LOC_Os02g07350 | inositol-1-monophosphatase, putative, expressed |
| LOC_Os02g07360 | pentatricopeptide containing protein, putative, expressed |
| LOC_Os02g07370 | retrotransposon protein, putative, unclassified, expressed |
| LOC_Os02g07410 | glycine cleavage system H protein, putative, expressed |
| LOC_Os02g07430 | OsMADS29 - MADS-box family gene with MIKCc type-box, expressed |
| LOC_Os02g07440 | DEFL78 - Defensin and Defensin-like DEFL family, expressed |
| LOC_Os02g07480 | transglycosylase SLT domain containing protein, expressed |
| LOC_Os02g07650 | zinc-binding protein, putative, expressed |
| LOC_Os02g07654 | expressed protein |
| LOC_Os02g07690 | VQ domain containing protein, putative, expressed |
| LOC_Os02g07720 | cytidylyltransferase domain containing protein, expressed |
| LOC_Os02g07730 | haloacid dehalogenase-like hydrolase domain-containing protein 3, putative, expressed |
| LOC_Os02g07740 | RNA binding protein, putative, expressed |
| LOC_Os02g07770 | Myb transcription factor, putative, expressed |
| LOC_Os02g07800 | transcription factor, putative, expressed |
| LOC_Os02g07820 | DUF581 domain containing protein, expressed |
| LOC_Os02g07850 | DUF630/DUF632 domains containing protein, putative, expressed |
| LOC_Os02g07920 | expressed protein |
| LOC_Os02g07930 | B-box zinc finger family protein, putative, expressed |
| LOC_Os02g07970 | expressed protein |
| LOC_Os02g07980 | expressed protein |
| LOC_Os02g07990 | hypothetical protein |
| LOC_Os02g08000 | expressed protein |
| LOC_Os02g08020 | hypothetical protein |
| LOC_Os02g08070 | OsSPL5 - SBP-box gene family member, expressed |
| LOC_Os02g08080 | expressed protein |
| LOC_Os02g08100 | AMP-binding domain containing protein, expressed |

| | |
|---|---|
| LOC_Os02g08120 | calmodulin binding protein, putative, expressed |
| LOC_Os02g08130 | FGGY family of carbohydrate kinases, putative, expressed |
| LOC_Os02g08150 | CCT/B-box zinc finger protein, putative, expressed |
| LOC_Os02g08220 | expressed protein |
| LOC_Os02g08230 | WAX2, putative, expressed |
| LOC_Os02g08300 | RAD23 DNA repair protein, putative, expressed |
| LOC_Os02g08320 | haemolysin-III, putative, expressed |
| LOC_Os02g08330 | gp176, putative, expressed |
| LOC_Os02g08350 | expressed protein |
| LOC_Os02g08360 | RNA recognition motif, putative, expressed |
| LOC_Os02g08370 | ubiquitin carboxyl-terminal hydrolase, family 1, putative, expressed |
| LOC_Os02g08380 | CR084 protein, putative, expressed |
| LOC_Os02g08410 | chorismate mutase, chloroplast precursor, putative, expressed |
| LOC_Os02g08420 | cinnamoyl CoA reductase, putative, expressed |
| LOC_Os02g09130 | palmitoyltransferase TIP1, putative, expressed |
| LOC_Os02g09490 | dehydrogenase, putative, expressed |
| LOC_Os02g09740 | STRUBBELIG-RECEPTOR FAMILY 7 precursor, putative, expressed |
| LOC_Os02g09930 | CSLA1 - cellulose synthase-like family A; mannan synthase, expressed |
| LOC_Os02g09960 | LYK8, putative, expressed |
| LOC_Os02g10140 | bZIP transcription factor domain containing protein, expressed |
| LOC_Os02g10180 | dnaJ homolog subfamily C member, putative, expressed |
| LOC_Os02g10200 | zinc finger A20 and AN1 domain-containing stress-associated protein, putative, expressed |
| LOC_Os02g10210 | expressed protein |
| LOC_Os02g10880 | UDP-glucoronosyl and UDP-glucosyl transferase domain containing protein, expressed |
| LOC_Os02g11050 | 26S protease regulatory subunit, putative, expressed |
| LOC_Os02g11665 | transposon protein, putative, unclassified, expressed |
| LOC_Os02g12140 | retrotransposon protein, putative, Ty3-gypsy subclass, expressed |
| LOC_Os02g12240 | expressed protein |
| LOC_Os02g12300 | pectate lyase precursor, putative, expressed |

| | |
|---|---|
| LOC_Os02g12654 | expressed protein |
| LOC_Os02g13020 | expressed protein |
| LOC_Os02g13440 | expressed protein |
| LOC_Os02g13510 | receptor-like protein kinase 5 precursor, putative, expressed |
| LOC_Os02g13780 | receptor-like protein kinase 2 precursor, putative, expressed |
| LOC_Os02g13790 | expressed protein |
| LOC_Os02g13870 | aquaporin protein, putative, expressed |
| LOC_Os02g14140 | cytokinin inducible protein, putative, expressed |
| LOC_Os02g14250 | transposon protein, putative, unclassified, expressed |
| LOC_Os02g14310 | expressed protein |
| LOC_Os02g14580 | retrotransposon protein, putative, unclassified, expressed |
| LOC_Os02g14610 | expressed protein |
| LOC_Os02g14650 | hypothetical protein |
| LOC_Os02g14800 | expressed protein |
| LOC_Os02g14850 | expressed protein |
| LOC_Os02g15310 | RNA recognition motif containing protein, expressed |
| LOC_Os02g15360 | expressed protein |
| LOC_Os02g15550 | spermidine synthase, putative, expressed |
| LOC_Os02g15690 | polygalacturonase, putative, expressed |
| LOC_Os02g15740 | expressed protein |
| LOC_Os02g15750 | expressed protein |
| LOC_Os02g15760 | helix-loop-helix DNA-binding domain containing protein, expressed |
| LOC_Os02g15770 | expressed protein |
| LOC_Os02g15800 | expressed protein |
| LOC_Os02g15810 | HMG1/2, putative, expressed |
| LOC_Os02g15820 | extra-large G-protein-related, putative, expressed |
| LOC_Os02g16050 | transposon protein, putative, unclassified, expressed |
| LOC_Os02g16160 | expressed protein |
| LOC_Os02g16310 | retrotransposon, putative, centromere-specific, expressed |

| | |
|---|---|
| LOC_Os02g16320 | retrotransposon protein, putative, Ty3-gypsy subclass, expressed |
| LOC_Os02g16490 | expressed protein |
| LOC_Os02g16780 | expansin precursor, putative, expressed |
| LOC_Os02g17220 | transposon protein, putative, unclassified, expressed |
| LOC_Os02g17760 | cytochrome P450, putative, expressed |
| LOC_Os02g17840 | expressed protein |
| LOC_Os02g18064 | expressed protein |
| LOC_Os02g18150 | expressed protein |
| LOC_Os02g18350 | expressed protein |
| LOC_Os02g18515 | retrotransposon protein, putative, unclassified, expressed |
| LOC_Os02g18700 | ABC transporter family protein, putative, expressed |
| LOC_Os02g18780 | expressed protein |
| LOC_Os02g19030 | hypothetical protein |
| LOC_Os02g19530 | lectin-like receptor kinase, putative, expressed |
| LOC_Os02g19550 | lectin-like receptor kinase, putative, expressed |
| LOC_Os02g19720 | retrotransposon protein, putative, unclassified, expressed |
| LOC_Os02g19900 | expressed protein |
| LOC_Os02g20130 | DEFL72 - Defensin and Defensin-like DEFL family |
| LOC_Os02g20210 | disease resistance protein RGA1, putative, expressed |
| LOC_Os02g20290 | expressed protein |
| LOC_Os02g20330 | expressed protein |
| LOC_Os02g20360 | tyrosine aminotransferase, putative, expressed |
| LOC_Os02g20420 | retrotransposon protein, putative, LINE subclass, expressed |
| LOC_Os02g20490 | expressed protein |
| LOC_Os02g20520 | retrotransposon, putative, centromere-specific |
| LOC_Os02g20540 | fasciclin domain containing protein, expressed |
| LOC_Os02g20590 | MBTB1 - Bric-a-Brac, Tramtrack, Broad Complex BTB domain with Meprin and TRAF Homology MATH domain, expressed |
| LOC_Os02g20700 | expressed protein |
| LOC_Os02g20820 | retrotransposon protein, putative, unclassified, expressed |

| | |
|---|---|
| LOC_Os02g20844 | expressed protein |
| LOC_Os02g20930 | apoptosis inhibitor 5, putative, expressed |
| LOC_Os02g20934 | expressed protein |
| LOC_Os02g20940 | CHCH domain containing protein, expressed |
| LOC_Os02g21980 | retrotransposon protein, putative, unclassified, expressed |
| LOC_Os02g22160 | DNA binding protein, putative, expressed |
| LOC_Os02g22374 | expressed protein |
| LOC_Os02g22740 | retrotransposon protein, putative, unclassified, expressed |
| LOC_Os02g22800 | expressed protein |
| LOC_Os02g24090 | retrotransposon protein, putative, unclassified, expressed |
| LOC_Os02g24390 | expressed protein |
| LOC_Os02g24420 | retrotransposon protein, putative, unclassified, expressed |
| LOC_Os02g24576 | chloroplast 30S ribosomal protein S7, putative, expressed |
| LOC_Os02g25755 | expressed protein |
| LOC_Os02g25780 | expressed protein |
| LOC_Os02g25860 | glutelin, putative, expressed |
| LOC_Os02g25870 | elongation factor Tu, putative, expressed |
| LOC_Os02g25960 | ankyrin repeat family protein, putative, expressed |
| LOC_Os02g25990 | retrotransposon, putative, centromere-specific |
| LOC_Os02g26130 | hypothetical protein |
| LOC_Os02g26210 | flowering promoting factor-like 1, putative, expressed |
| LOC_Os02g26320 | fasciclin-like arabinogalactan precursor protein, putative, expressed |
| LOC_Os02g26370 | wiskott-Aldrich syndrome protein family member 2, putative, expressed |
| LOC_Os02g26390 | expressed protein |
| LOC_Os02g26400 | nuclease, EndA/NucM family protein, expressed |
| LOC_Os02g26560 | expressed protein |
| LOC_Os02g26580 | expressed protein |
| LOC_Os02g26590 | hypothetical protein |
| LOC_Os02g26600 | ATP binding, related, putative, expressed |

| | |
|---|---|
| LOC_Os02g26670 | F-box/Kelch-repeat protein, putative, expressed |
| LOC_Os02g26700 | cation transport regulator-like protein 1, putative, expressed |
| LOC_Os02g27360 | aspartic proteinase-like protein 2 precursor, putative, expressed |
| LOC_Os02g27470 | importin-7, putative, expressed |
| LOC_Os02g27500 | Leucine Rich Repeat family protein, expressed |
| LOC_Os02g27910 | expressed protein |
| LOC_Os02g28050 | hypothetical protein |
| LOC_Os02g28120 | retrotransposon, putative, centromere-specific |
| LOC_Os02g28140 | expressed protein |
| LOC_Os02g28200 | transferase family protein, putative, expressed |
| LOC_Os02g28210 | expressed protein |
| LOC_Os02g28260 | transposon protein, putative, CACTA, En/Spm sub-class |
| LOC_Os02g28290 | hypothetical protein |
| LOC_Os02g28410 | transferase family protein, putative, expressed |
| LOC_Os02g28870 | U-box domain-containing protein, putative, expressed |
| LOC_Os02g28900 | cytokinin-O-glucosyltransferase 2, putative, expressed |
| LOC_Os02g28920 | transposon protein, putative, unclassified, expressed |
| LOC_Os02g28980 | peptidyl-prolyl isomerase, putative, expressed |
| LOC_Os02g28990 | expressed protein |
| LOC_Os02g29000 | Cupin domain containing protein, expressed |
| LOC_Os02g29010 | retrotransposon protein, putative, unclassified, expressed |
| LOC_Os02g29230 | expressed protein |
| LOC_Os02g29570 | retrotransposon protein, putative, Ty3-gypsy subclass, expressed |
| LOC_Os02g29730 | retrotransposon protein, putative, Ty3-gypsy subclass, expressed |
| LOC_Os02g29740 | retrotransposon protein, putative, Ty3-gypsy subclass, expressed |
| LOC_Os02g29910 | hypothetical protein |
| LOC_Os02g30180 | ZOS2-07 - C2H2 zinc finger protein, expressed |
| LOC_Os02g30200 | PE-PGRS family protein, putative, expressed |
| LOC_Os02g30280 | expressed protein |

| | |
|---|---|
| LOC_Os02g30310 | ThiF family domain containing protein, putative, expressed |
| LOC_Os02g30760 | expressed protein |
| LOC_Os02g30780 | LGC1, putative, expressed |
| LOC_Os02g30790 | SAR DNA-binding protein-like, putative, expressed |
| LOC_Os02g30800 | POLE1 - Putative DNA polymerase epsilon catalytic subunit, expressed |
| LOC_Os02g30900 | protein kinase domain containing protein, expressed |
| LOC_Os02g30974 | expressed protein |
| LOC_Os02g31280 | expressed protein |
| LOC_Os02g31290 | AML1, putative, expressed |
| LOC_Os02g31300 | UBX domain containing protein, expressed |
| LOC_Os02g32060 | hydrolase, NUDIX family, domain containing protein, expressed |
| LOC_Os02g32160 | copine, putative, expressed |
| LOC_Os02g32180 | expressed protein |
| LOC_Os02g32200 | thioesterase family protein, putative, expressed |
| LOC_Os02g32210 | expressed protein |
| LOC_Os02g32280 | expressed protein |
| LOC_Os02g32350 | TUDOR protein with multiple SNc domains, putative, expressed |
| LOC_Os02g32504 | heparan-alpha-glucosaminide N-acetyltransferase, putative, expressed |
| LOC_Os02g32660 | 1,4-alpha-glucan-branching enzyme, chloroplast precursor, putative, expressed |
| LOC_Os02g32920 | expressed protein |
| LOC_Os02g32960 | expressed protein |
| LOC_Os02g33230 | nucleoside-diphosphate-sugar epimerase, putative, expressed |
| LOC_Os02g33460 | transposon protein, putative, Mutator sub-class, expressed |
| LOC_Os02g33570 | expressed protein |
| LOC_Os02g33580 | digalactosyldiacylglycerol synthase, chloroplast precursor, putative, expressed |
| LOC_Os02g33610 | metallo-beta-lactamase, putative, expressed |
| LOC_Os02g33670 | expressed protein |
| LOC_Os02g33720 | RING-H2 finger protein, putative, expressed |
| LOC_Os02g33900 | transposon protein, putative, unclassified, expressed |

| | |
|---|---|
| LOC_Os02g34030 | retrotransposon protein, putative, unclassified, expressed |
| LOC_Os02g34060 | expressed protein |
| LOC_Os02g34140 | expressed protein |
| LOC_Os02g34150 | hypothetical protein |
| LOC_Os02g34290 | expressed protein |
| LOC_Os02g34310 | expressed protein |
| LOC_Os02g34510 | vacuolar ATP synthase, putative, expressed |
| LOC_Os02g34690 | transmembrane 9 superfamily member, putative, expressed |
| LOC_Os02g34710 | expressed protein |
| LOC_Os02g34800 | leucine-rich repeat receptor protein kinase EXS precursor, putative, expressed |
| LOC_Os02g34980 | hypothetical protein |
| LOC_Os02g34990 | ACT domain containing protein, putative, expressed |
| LOC_Os02g35450 | DNA repair protein Rad51, putative, expressed |
| LOC_Os02g35500 | NAD binding domain of 6-phosphogluconate dehydrogenase containing protein, expressed |
| LOC_Os02g35560 | OsFBX53 - F-box domain containing protein, expressed |
| LOC_Os02g35650 | late embryogenesis abundant protein, putative, expressed |
| LOC_Os02g35660 | PTF1, putative, expressed |
| LOC_Os02g35740 | expressed protein |
| LOC_Os02g35860 | expressed protein |
| LOC_Os02g35870 | galactosyltransferase, putative, expressed |
| LOC_Os02g35920 | expressed protein |
| LOC_Os02g35940 | gibberellin receptor GID1L2, putative, expressed |
| LOC_Os02g35950 | RNA recognition motif containing protein, putative, expressed |
| LOC_Os02g36000 | zinc finger protein, putative, expressed |
| LOC_Os02g36020 | transposon protein, putative, CACTA, En/Spm sub-class, expressed |
| LOC_Os02g36080 | transposon protein, putative, unclassified, expressed |
| LOC_Os02g36200 | hypothetical protein |
| LOC_Os02g36390 | nucleotide-sugar transporter family protein, putative, expressed |
| LOC_Os02g36910 | transposon protein, putative, unclassified |

| | |
|---|---|
| LOC_Os02g37990 | histamine oxidase, putative |
| LOC_Os02g38090 | AP2 domain containing protein, expressed |
| LOC_Os02g38130 | no apical meristem protein, putative, expressed |
| LOC_Os02g38490 | expressed protein |
| LOC_Os02g38508 | expressed protein |
| LOC_Os02g38800 | 26S proteasome non-ATPase regulatory subunit 6, putative, expressed |
| LOC_Os02g39450 | retrotransposon protein, putative, unclassified, expressed |
| LOC_Os02g39470 | cyclin, N-terminal domain containing protein, expressed |
| LOC_Os02g39790 | CPuORF9 - conserved peptide uORF-containing transcript, expressed |
| LOC_Os02g39910 | B4-BTB1 - Bric-a-Brac, Tramtrack, Broad Complex BTB domain with B4 subfamily conserved sequence, expressed |
| LOC_Os02g40010 | phosphoribosyl transferase, putative, expressed |
| LOC_Os02g40330 | retrotransposon protein, putative, Ty3-gypsy subclass, expressed |
| LOC_Os02g41480 | OsWAK12 - OsWAK receptor-like cytoplasmic kinase OsWAK-RLCK, expressed |
| LOC_Os02g41630 | phenylalanine ammonia-lyase, putative, expressed |
| LOC_Os02g41860 | aquaporin protein, putative, expressed |
| LOC_Os02g41954 | gibberellin 2-beta-dioxygenase 7, putative, expressed |
| LOC_Os02g42030 | hypothetical protein |
| LOC_Os02g42060 | response regulator receiver domain containing protein, expressed |
| LOC_Os02g42110 | wall-associated receptor kinase-like 22 precursor, putative, expressed |
| LOC_Os02g42520 | dehydrogenase, putative, expressed |
| LOC_Os02g42585 | AP2 domain containing protein, expressed |
| LOC_Os02g42970 | NAC domain containing protein, putative, expressed |
| LOC_Os02g43030 | integral membrane protein, putative, expressed |
| LOC_Os02g43090 | myristoyl-acyl carrier protein thioesterase, chloroplast precursor, putative, expressed |
| LOC_Os02g43560 | WRKY34, expressed |
| LOC_Os02g43590 | ELMO/CED-12 family protein, putative, expressed |
| LOC_Os02g43610 | expressed protein |
| LOC_Os02g43800 | retrotransposon protein, putative, Ty3-gypsy subclass, expressed |
| LOC_Os02g44290 | phosphatase, putative, expressed |

| | |
|---|---|
| LOC_Os02g44300 | MSP domain containing protein, expressed |
| LOC_Os02g44310 | LTPL112 - Protease inhibitor/seed storage/LTP family protein precursor, expressed |
| LOC_Os02g44470 | actin-depolymerizing factor, putative, expressed |
| LOC_Os02g44500 | glutathione peroxidase, putative, expressed |
| LOC_Os02g44870 | dehydrin, putative, expressed |
| LOC_Os02g44910 | transmembrane protein, putative, expressed |
| LOC_Os02g44920 | protein kinase domain containing protein, expressed |
| LOC_Os02g45300 | hypothetical protein |
| LOC_Os02g45310 | dehydration response related protein, putative, expressed |
| LOC_Os02g45370 | retrotransposon protein, putative, Ty3-gypsy subclass, expressed |
| LOC_Os02g45590 | PPR repeat domain containing protein, putative, expressed |
| LOC_Os02g46210 | DUF581 domain containing protein, expressed |
| LOC_Os02g46260 | OsSCP9 - Putative Serine Carboxypeptidase homologue, expressed |
| LOC_Os02g46390 | expressed protein |
| LOC_Os02g46400 | expressed protein |
| LOC_Os02g46440 | expressed protein |
| LOC_Os02g46450 | helicase conserved C-terminal domain containing protein, expressed |
| LOC_Os02g46473 | expressed protein |
| LOC_Os02g46780 | MYB family transcription factor, putative, expressed |
| LOC_Os02g47220 | CPuORF20 - conserved peptide uORF-containing transcript, expressed |
| LOC_Os02g47280 | growth-regulating factor, putative, expressed |
| LOC_Os02g47330 | expressed protein |
| LOC_Os02g47990 | retrotransposon protein, putative, unclassified, expressed |
| LOC_Os02g48020 | expressed protein |
| LOC_Os02g48080 | cysteine-rich receptor-like protein kinase 7 precursor, putative, expressed |
| LOC_Os02g48360 | pyrophosphate--fructose 6-phosphate 1-phosphotransferase subunit alpha, putative, expressed |
| LOC_Os02g48390 | phosphoribosyl transferase, putative, expressed |
| LOC_Os02g48560 | fatty acid desaturase, putative, expressed |
| LOC_Os02g48610 | mitochondrial import inner membrane translocase subunit Tim17, putative, expressed |

| | |
|---|---|
| LOC_Os02g48630 | expressed protein |
| LOC_Os02g48640 | expressed protein |
| LOC_Os02g48850 | plant-specific domain TIGR01615 family protein, expressed |
| LOC_Os02g49060 | amino acid transporter, putative, expressed |
| LOC_Os02g49170 | retrotransposon protein, putative, unclassified, expressed |
| LOC_Os02g49220 | hypothetical protein |
| LOC_Os02g49332 | CSLE2 - cellulose synthase-like family E, expressed |
| LOC_Os02g49410 | histone-like transcription factor and archaeal histone, putative, expressed |
| LOC_Os02g49460 | auxin-independent growth promoter protein, putative, expressed |
| LOC_Os02g49580 | expressed protein |
| LOC_Os02g49590 | ASC1, putative, expressed |
| LOC_Os02g49600 | BRASSINOSTEROID INSENSITIVE 1-associated receptor kinase 1 precursor, putative, expressed |
| LOC_Os02g49650 | leaf senescence related protein, putative, expressed |
| LOC_Os02g49690 | ACT domain-containing protein, putative, expressed |
| LOC_Os02g49750 | expressed protein |
| LOC_Os02g49820 | RNA recognition motif family protein, expressed |
| LOC_Os02g49850 | plastocyanin-like domain containing protein, putative, expressed |
| LOC_Os02g49950 | spotted leaf 11, putative, expressed |
| LOC_Os02g50174 | caleosin related protein, putative, expressed |
| LOC_Os02g50210 | erwinia induced protein 2, putative, expressed |
| LOC_Os02g50610 | retrotransposon protein, putative, Ty3-gypsy subclass, expressed |
| LOC_Os02g50670 | expressed protein |
| LOC_Os02g50970 | protein kinase domain containing protein, expressed |
| LOC_Os02g51040 | expansin precursor, putative, expressed |
| LOC_Os02g51070 | starch synthase, putative, expressed |
| LOC_Os02g51110 | aquaporin protein, putative, expressed |
| LOC_Os02g51130 | glycosyl transferase 8 domain containing protein, putative, expressed |
| LOC_Os02g51280 | TCP-domain protein, putative, expressed |
| LOC_Os02g51310 | TCP family transcription factor, putative, expressed |

| | |
|---|---|
| LOC_Os02g51320 | helix-loop-helix DNA-binding domain containing protein, expressed |
| LOC_Os02g51330 | MAG2, putative, expressed |
| LOC_Os02g51350 | OsFBK10 - F-box domain and kelch repeat containing protein, expressed |
| LOC_Os02g51360 | expressed protein |
| LOC_Os02g51490 | methyltransferase, putative, expressed |
| LOC_Os02g51540 | eukaryotic aspartyl protease domain containing protein, expressed |
| LOC_Os02g51660 | trpH, putative, expressed |
| LOC_Os02g51670 | ethylene-responsive transcription factor, putative, expressed |
| LOC_Os02g51680 | uncharacterized glycosyl hydrolase Rv2006/MT2062, putative, expressed |
| LOC_Os02g51700 | expressed protein |
| LOC_Os02g51710 | wound/stress protein, putative, expressed |
| LOC_Os02g51720 | expressed protein |
| LOC_Os02g51970 | phosphate-induced protein 1 conserved region domain containing protein, expressed |
| LOC_Os02g52170 | expressed protein |
| LOC_Os02g52850 | receptor-like protein kinase like protein, putative, expressed |
| LOC_Os02g52900 | glutaredoxin 2, putative, expressed |
| LOC_Os02g53020 | transposon protein, putative, CACTA, En/Spm sub-class, expressed |
| LOC_Os02g53260 | acetyltransferase, GNAT family, putative, expressed |
| LOC_Os02g53440 | expressed protein |
| LOC_Os02g53600 | DEFL75 - Defensin and Defensin-like DEFL family, expressed |
| LOC_Os02g53690 | growth regulating factor protein, putative, expressed |
| LOC_Os02g53700 | DENN domain containing protein, expressed |
| LOC_Os02g53710 | expressed protein |
| LOC_Os02g53860 | OsSub22 - Putative Subtilisin homologue, expressed |
| LOC_Os02g53960 | CXE carboxylesterase, putative, expressed |
| LOC_Os02g53990 | cyclin-related protein, putative, expressed |
| LOC_Os02g54520 | MYB family transcription factor, putative, expressed |
| LOC_Os02g54860 | ankyrin repeat-rich protein, putative, expressed |
| LOC_Os02g55250 | BHLH transcription factor, putative, expressed |

| | |
|---|---|
| LOC_Os02g55380 | AP2 domain containing protein, expressed |
| LOC_Os02g56190 | expressed protein |
| LOC_Os02g56260 | expressed protein |
| LOC_Os02g56530 | ankyrin repeat domain containing protein, expressed |
| LOC_Os02g56710 | hypothetical protein |
| LOC_Os02g56970 | uncharacterized protein yqjG, putative, expressed |
| LOC_Os02g57190 | Myosin head domain containing protein, expressed |
| LOC_Os02g57620 | Citrate transporter protein, putative, expressed |
| LOC_Os02g57770 | glycosyl hydrolases family 16, putative, expressed |
| LOC_Os02g57854 | vacuolar ATP synthase subunit F, putative, expressed |
| LOC_Os02g58020 | ABC transporter, ATP-binding protein, putative, expressed |
| LOC_Os02g58030 | leucine-rich repeat-containing protein 40, putative, expressed |
| LOC_Os02g58050 | hypothetical protein |
| LOC_Os02g58120 | oxidoreductase/ transition metal ion binding protein, putative, expressed |
| LOC_Os02g58300 | expressed protein |
| LOC_Os02g58310 | expressed protein |
| LOC_Os02g58330 | expressed protein |
| LOC_Os02g58440 | zinc finger C-x8-C-x5-C-x3-H type family protein, expressed |
| LOC_Os02g58460 | beta-catenin-like protein 1, putative, expressed |
| LOC_Os02g58470 | expressed protein |
| LOC_Os02g58554 | expressed protein |
| LOC_Os02g58600 | expressed protein |
| LOC_Os03g01170 | transporter family protein, putative, expressed |
| LOC_Os03g01240 | CCR4-Not complex component, Not1domain containing protein, expressed |
| LOC_Os03g01270 | expansin precursor, putative, expressed |
| LOC_Os03g01450 | expressed protein |
| LOC_Os03g01560 | expressed protein |
| LOC_Os03g01580 | expressed protein |
| LOC_Os03g01590 | PAIR1, putative, expressed |

| | |
|---|---|
| LOC_Os03g01610 | expansin precursor, putative, expressed |
| LOC_Os03g01720 | protein binding protein, putative, expressed |
| LOC_Os03g01740 | expressed protein |
| LOC_Os03g01750 | dual specificity protein phosphatase, putative, expressed |
| LOC_Os03g01760 | exostosin family domain containing protein, expressed |
| LOC_Os03g01790 | zinc finger, C3HC4 type domain containing protein, expressed |
| LOC_Os03g01820 | fatty acid hydroxylase, putative, expressed |
| LOC_Os03g02010 | DNA methyltransferase protein, putative, expressed |
| LOC_Os03g02100 | valyl-tRNA synthetase, putative, expressed |
| LOC_Os03g02260 | DnaK family protein, putative, expressed |
| LOC_Os03g02460 | retinol dehydrogenase, putative, expressed |
| LOC_Os03g02514 | hydrolase, alpha/beta fold family protein, putative, expressed |
| LOC_Os03g02530 | OsRhmbd7 - Putative Rhomboid homologue, expressed |
| LOC_Os03g02900 | B3 DNA binding domain containing protein, expressed |
| LOC_Os03g03034 | flavonol synthase/flavanone 3-hydroxylase, putative, expressed |
| LOC_Os03g03180 | uncharacterized Cys-rich domain containing protein, putative, expressed |
| LOC_Os03g03400 | diacylglycerol kinase, putative, expressed |
| LOC_Os03g03410 | serine/threonine-protein kinase, putative, expressed |
| LOC_Os03g03430 | CBS domain containing protein, expressed |
| LOC_Os03g03440 | expressed protein |
| LOC_Os03g03450 | anthranilate phosphoribosyltransferase, chloroplast precursor, putative, expressed |
| LOC_Os03g03460 | erythronate-4-phosphate dehydrogenase, putative, expressed |
| LOC_Os03g03470 | expressed protein |
| LOC_Os03g03500 | heavy metal-associated domain containing protein, expressed |
| LOC_Os03g03510 | CAMK_KIN1/SNF1/Nim1_like 15 - CAMK includes calcium/calmodulin depedent protein kinases, expressed |
| LOC_Os03g03530 | expressed protein |
| LOC_Os03g03550 | bZIP family transcription factor, putative, expressed |
| LOC_Os03g03610 | 1,3-beta-glucan synthase component domain containing protein, expressed |
| LOC_Os03g03690 | glycosylphosphatidylinositol anchor biosynthesis protein 11, putative, expressed |

| | |
|---|---|
| LOC_Os03g03710 | ribonuclease protein, putative, expressed |
| LOC_Os03g03760 | MYB family transcription factor, putative, expressed |
| LOC_Os03g03850 | GATA zinc finger domain containing protein, expressed |
| LOC_Os03g04169 | ATP phosphoribosyltransferase, putative, expressed |
| LOC_Os03g04190 | cytochrome P450, putative, expressed |
| LOC_Os03g04195 | expressed protein |
| LOC_Os03g04270 | OsFBL10 - F-box domain and LRR containing protein, expressed |
| LOC_Os03g04300 | ankyrin repeat domain containing protein, expressed |
| LOC_Os03g04340 | C-methyltransferase, putative, expressed |
| LOC_Os03g04350 | expressed protein |
| LOC_Os03g04370 | protein-methionine-S-oxide reductase, putative, expressed |
| LOC_Os03g04400 | heat shock protein DnaJ, putative, expressed |
| LOC_Os03g04410 | aconitate hydratase protein, putative, expressed |
| LOC_Os03g04450 | expressed protein |
| LOC_Os03g04470 | expressed protein |
| LOC_Os03g04480 | CorA-like magnesium transporter protein, putative, expressed |
| LOC_Os03g04530 | cytochrome P450, putative, expressed |
| LOC_Os03g04540 | expressed protein |
| LOC_Os03g04545 | expressed protein |
| LOC_Os03g04550 | expressed protein |
| LOC_Os03g04580 | expressed protein |
| LOC_Os03g04590 | ribosomal protein, putative, expressed |
| LOC_Os03g04600 | expressed protein |
| LOC_Os03g04610 | expressed protein |
| LOC_Os03g04620 | CCT motif family protein, expressed |
| LOC_Os03g04630 | cytochrome P450 protein, putative, expressed |
| LOC_Os03g04690 | expressed protein |
| LOC_Os03g04710 | expressed protein |
| LOC_Os03g04750 | 60S ribosomal protein L21-2, putative, expressed |

| | |
|---|---|
| LOC_Os03g04760 | expressed protein |
| LOC_Os03g04770 | beta-amylase, putative, expressed |
| LOC_Os03g04780 | RNA recognition motif containing protein, putative, expressed |
| LOC_Os03g04830 | expressed protein |
| LOC_Os03g04890 | zinc finger family protein, putative, expressed |
| LOC_Os03g04930 | expressed protein |
| LOC_Os03g04970 | T-complex protein, putative, expressed |
| LOC_Os03g04980 | PHD-finger domain containing protein, putative, expressed |
| LOC_Os03g04990 | expressed protein |
| LOC_Os03g05020 | PIR, putative, expressed |
| LOC_Os03g05060 | exostosin family domain containing protein, expressed |
| LOC_Os03g05120 | expressed protein |
| LOC_Os03g05130 | expressed protein |
| LOC_Os03g05180 | expressed protein |
| LOC_Os03g05260 | ankyrin repeat domain containing protein, expressed |
| LOC_Os03g05270 | RING finger and CHY zinc finger domain-containing protein 1, putative, expressed |
| LOC_Os03g05300 | cyclin-dependent kinases regulatory subunit 1, putative, expressed |
| LOC_Os03g05310 | pheophorbide a oxygenase, chloroplast precursor, putative, expressed |
| LOC_Os03g05320 | expressed protein |
| LOC_Os03g05330 | HEAT repeat family protein, putative, expressed |
| LOC_Os03g05340 | retrotransposon protein, putative, unclassified, expressed |
| LOC_Os03g05370 | expressed protein |
| LOC_Os03g05410 | expressed protein |
| LOC_Os03g05450 | expressed protein |
| LOC_Os03g05530 | nodulin, putative, expressed |
| LOC_Os03g05540 | tetratricopeptide repeat containing protein, putative, expressed |
| LOC_Os03g05580 | expressed protein |
| LOC_Os03g05630 | expressed protein |
| LOC_Os03g05660 | appr-1-p processing enzyme family protein, putative, expressed |

| | |
|---|---|
| LOC_Os03g05680 | histone demethylase JARID1C, putative, expressed |
| LOC_Os03g05720 | WD domain, G-beta repeat domain containing protein, expressed |
| LOC_Os03g05730 | cell division control protein 48 homolog E, putative, expressed |
| LOC_Os03g05750 | heavy-metal-associated domain-containing protein, putative, expressed |
| LOC_Os03g05760 | transcription factor Dp, putative, expressed |
| LOC_Os03g05780 | 4-coumarate--CoA ligase-like 7, putative, expressed |
| LOC_Os03g05800 | expressed protein |
| LOC_Os03g05890 | expressed protein |
| LOC_Os03g06040 | expansin precursor, putative, expressed |
| LOC_Os03g06070 | la domain containing protein, putative, expressed |
| LOC_Os03g06120 | PLAC8 family protein, expressed |
| LOC_Os03g06139 | ABC-2 type transporter domain containing protein, expressed |
| LOC_Os03g06160 | BTBP1 - Bric-a-Brac, Tramtrack, Broad Complex BTB domain with pentapeptide repeats, expressed |
| LOC_Os03g06170 | hsp20/alpha crystallin family protein, putative, expressed |
| LOC_Os03g06180 | expressed protein |
| LOC_Os03g06340 | XS domain containing protein, expressed |
| LOC_Os03g06370 | PPR repeat domain containing protein, putative, expressed |
| LOC_Os03g06440 | RNA helicase SDE3, putative, expressed |
| LOC_Os03g06530 | expressed protein |
| LOC_Os03g06570 | IQ calmodulin-binding motif family protein, putative, expressed |
| LOC_Os03g06600 | expressed protein |
| LOC_Os03g06610 | expressed protein |
| LOC_Os03g06670 | Core histone H2A/H2B/H3/H4 domain containing protein, putative, expressed |
| LOC_Os03g06710 | PPR repeat domain containing protein, putative, expressed |
| LOC_Os03g06720 | expressed protein |
| LOC_Os03g06890 | DUF593 domain containing protein, expressed |
| LOC_Os03g06900 | DNA topoisomerase 3 protein, putative, expressed |
| LOC_Os03g06910 | PPR repeat containing protein, expressed |
| LOC_Os03g06920 | DRD1, putative, expressed |

| | |
|---|---|
| LOC_Os03g06930 | homeodomain protein, putative, expressed |
| LOC_Os03g06950 | ubiquitin carboxyl-terminal hydrolase domain containing protein, expressed |
| LOC_Os03g06960 | vesicle-associated membrane protein, putative, expressed |
| LOC_Os03g06970 | tetratricopeptide repeat domain containing protein, expressed |
| LOC_Os03g06980 | nucleic acid binding protein, putative, expressed |
| LOC_Os03g06990 | oxidoreductase, putative, expressed |
| LOC_Os03g07100 | LTPL82 - Protease inhibitor/seed storage/LTP family protein precursor, expressed |
| LOC_Os03g07110 | calmodulin-binding protein, putative, expressed |
| LOC_Os03g07130 | RING finger protein 13, putative, expressed |
| LOC_Os03g07140 | male sterility protein, putative, expressed |
| LOC_Os03g07160 | OsFBK11 - F-box domain and kelch repeat containing protein, expressed |
| LOC_Os03g07220 | PPR repeat domain containing protein, putative, expressed |
| LOC_Os03g07226 | thioredoxin, putative, expressed |
| LOC_Os03g07250 | cytochrome P450, putative, expressed |
| LOC_Os03g07260 | tetratricopeptide repeat domain containing protein, expressed |
| LOC_Os03g07290 | structural constituent of ribosome, putative, expressed |
| LOC_Os03g07300 | ribulose-phosphate 3-epimerase, chloroplast precursor, putative, expressed |
| LOC_Os03g07350 | CSLA4 - cellulose synthase-like family A; mannan synthase, expressed |
| LOC_Os03g07370 | HNH endonuclease domain-containing protein, putative, expressed |
| LOC_Os03g07480 | sucrose transporter, putativ, expressed |
| LOC_Os03g07530 | OsFBK12 - F-box domain and kelch repeat containing protein, expressed |
| LOC_Os03g07540 | bHLH family protein, putative, expressed |
| LOC_Os03g07560 | hypothetical protein |
| LOC_Os03g07570 | aminotransferase, putative, expressed |
| LOC_Os03g07580 | OsNucAP1 - Putative Nucleoporin Autopeptidase homologue, expressed |
| LOC_Os03g07590 | expressed protein |
| LOC_Os03g07700 | expressed protein |
| LOC_Os03g07710 | hypothetical protein |
| LOC_Os03g07720 | expressed protein |

| | |
|---|---|
| LOC_Os03g07730 | expressed protein |
| LOC_Os03g07750 | expressed protein |
| LOC_Os03g07790 | zinc finger, C3HC4 type domain containing protein, expressed |
| LOC_Os03g07800 | RNA recognition motif containing protein, putative, expressed |
| LOC_Os03g07820 | exostosin family protein, putative, expressed |
| LOC_Os03g07830 | AP2 domain containing protein, expressed |
| LOC_Os03g07840 | adenylosuccinate synthetase, chloroplast precursor, putative, expressed |
| LOC_Os03g07850 | tRNA methyltransferase, putative, expressed |
| LOC_Os03g07960 | expressed protein |
| LOC_Os03g07970 | expressed protein |
| LOC_Os03g07990 | acetyltransferase, GNAT family, putative, expressed |
| LOC_Os03g08070 | copper-transporting ATPase PAA1, putative, expressed |
| LOC_Os03g08080 | YGGT family protein, putative, expressed |
| LOC_Os03g08130 | expressed protein |
| LOC_Os03g08140 | tubulin--tyrosine ligase-like protein 12, putative, expressed |
| LOC_Os03g08170 | protein kinase APK1B, chloroplast precursor, putative, expressed |
| LOC_Os03g08300 | glycosyl transferase, group 1 domain containing protein, expressed |
| LOC_Os03g08680 | pentatricopeptide, putative, expressed |
| LOC_Os03g08720 | transferase family protein, putative, expressed |
| LOC_Os03g09070 | leucine rich repeat domain containing protein, putative, expressed |
| LOC_Os03g09140 | ras-related protein, putative, expressed |
| LOC_Os03g09250 | inositol-3-phosphate synthase, putative, expressed |
| LOC_Os03g09840 | C2 domain containing protein, putative, expressed |
| LOC_Os03g09860 | acetyltransferase, GNAT family, putative, expressed |
| LOC_Os03g09880 | AIR12, putative, expressed |
| LOC_Os03g09910 | aminotransferase, classes I and II, domain containing protein, expressed |
| LOC_Os03g09970 | sulfate transporter, putative, expressed |
| LOC_Os03g10040 | OsFBX77 - F-box domain containing protein, expressed |
| LOC_Os03g10110 | cupin domain containing protein, expressed |

| | |
|---|---|
| LOC_Os03g10220 | methyladenine glycosylase, putative, expressed |
| LOC_Os03g10280 | retrotransposon protein, putative, unclassified, expressed |
| LOC_Os03g10290 | expressed protein |
| LOC_Os03g10320 | expressed protein |
| LOC_Os03g10370 | ADP-ribosylation factor, putative, expressed |
| LOC_Os03g10380 | hypothetical protein |
| LOC_Os03g10390 | expressed protein |
| LOC_Os03g10430 | expressed protein |
| LOC_Os03g10440 | glycosyl hydrolase family 10 protein, putative, expressed |
| LOC_Os03g10460 | expressed protein |
| LOC_Os03g10590 | splicing factor, putative, expressed |
| LOC_Os03g10710 | expressed protein |
| LOC_Os03g10730 | expressed protein |
| LOC_Os03g10740 | expressed protein |
| LOC_Os03g10750 | CUE domain containing protein, expressed |
| LOC_Os03g10770 | helix-loop-helix DNA-binding domain containing protein, expressed |
| LOC_Os03g10790 | cytochrome b5-like Heme/Steroid binding domain containing protein, expressed |
| LOC_Os03g10800 | BTBN4 - Bric-a-Brac, Tramtrack, Broad Complex BTB domain with non-phototropic hypocotyl 3 NPH3 domain, expressed |
| LOC_Os03g10810 | HLS, putative, expressed |
| LOC_Os03g10820 | expressed protein |
| LOC_Os03g11110 | zinc finger family protein, putative, expressed |
| LOC_Os03g11120 | methionyl-tRNA synthetase, putative, expressed |
| LOC_Os03g11220 | keratin, type I cytoskeletal 9, putative, expressed |
| LOC_Os03g11230 | expressed protein |
| LOC_Os03g11310 | PPR repeat domain containing protein, putative, expressed |
| LOC_Os03g11330 | glycosyl transferase 8 domain containing protein, putative, expressed |
| LOC_Os03g11335 | expressed protein |
| LOC_Os03g11410 | mitochondrial-processing peptidase subunit, mitochondrial precursor, putative, expressed |
| LOC_Os03g11490 | expressed protein |

| | |
|---|---|
| LOC_Os03g12730 | receptor protein kinase CLAVATA1 precursor, putative, expressed |
| LOC_Os03g12790 | MATE efflux family protein, putative, expressed |
| LOC_Os03g13070 | thaumatin family domain containing protein, expressed |
| LOC_Os03g13300 | glutamate decarboxylase, putative, expressed |
| LOC_Os03g13320 | retrotransposon protein, putative, unclassified, expressed |
| LOC_Os03g13560 | hydroxyproline-rich glycoprotein family protein, putative, expressed |
| LOC_Os03g13690 | expressed protein |
| LOC_Os03g13720 | expressed protein |
| LOC_Os03g13790 | myb/SANT domain protein, putative, expressed |
| LOC_Os03g13800 | ribosomal protein L7Ae, putative, expressed |
| LOC_Os03g13810 | expressed protein |
| LOC_Os03g13870 | expressed protein |
| LOC_Os03g14030 | thaumatin, putative, expressed |
| LOC_Os03g14040 | expressed protein |
| LOC_Os03g14090 | armadillo/beta-catenin repeat family protein, putative, expressed |
| LOC_Os03g14140 | POEI16 - Pollen Ole e I allergen and extensin family protein precursor, expressed |
| LOC_Os03g14250 | plant protein of unknown function DUF869 domain containing protein, expressed |
| LOC_Os03g14254 | expressed protein |
| LOC_Os03g14258 | expressed protein |
| LOC_Os03g14290 | expressed protein |
| LOC_Os03g14400 | cytochrome P450, putative, expressed |
| LOC_Os03g14430 | expressed protein |
| LOC_Os03g14450 | enolase, putative, expressed |
| LOC_Os03g14500 | expressed protein |
| LOC_Os03g14510 | immature colon carcinoma transcript 1 protein precursor, putative, expressed |
| LOC_Os03g14530 | S10/S20 domain containing ribosomal protein, putative, expressed |
| LOC_Os03g14540 | UDP-glucuronate 4-epimerase, putative, expressed |
| LOC_Os03g14580 | expressed protein |
| LOC_Os03g14590 | calcium-binding EF hand family protein, putative, expressed |

| | |
|---|---|
| LOC_Os03g14610 | HVA22, putative, expressed |
| LOC_Os03g14615 | LTPL117 - Protease inhibitor/seed storage/LTP family protein precursor, expressed |
| LOC_Os03g14630 | LTPL106 - Protease inhibitor/seed storage/LTP family protein precursor, expressed |
| LOC_Os03g14690 | vacuolar ATP synthase 98 kDa subunit, putative, expressed |
| LOC_Os03g14700 | C2 domain containing protein, expressed |
| LOC_Os03g14710 | protein kinase, putative, expressed |
| LOC_Os03g14800 | serine palmitoyltransferase 1, putative, expressed |
| LOC_Os03g14910 | expressed protein |
| LOC_Os03g14980 | lissencephaly type-1-like homology motif, putative, expressed |
| LOC_Os03g15110 | expressed protein |
| LOC_Os03g15130 | expressed protein |
| LOC_Os03g15250 | lectin-like receptor kinase, putative, expressed |
| LOC_Os03g15260 | expressed protein |
| LOC_Os03g16430 | RNA polymerase sigma factor, putative, expressed |
| LOC_Os03g16760 | protein phosphatase 2C, putative, expressed |
| LOC_Os03g17210 | expressed protein |
| LOC_Os03g17240 | ankyrin repeat-containing protein, putative, expressed |
| LOC_Os03g17260 | expressed protein |
| LOC_Os03g17270 | expressed protein |
| LOC_Os03g17330 | expressed protein |
| LOC_Os03g17340 | expressed protein |
| LOC_Os03g17359 | expressed protein |
| LOC_Os03g17380 | expressed protein |
| LOC_Os03g17410 | expressed protein |
| LOC_Os03g17432 | pre-mRNA-splicing factor ATP-dependent RNA helicase DHX16, putative, expressed |
| LOC_Os03g17450 | nucleotide binding protein, putative, expressed |
| LOC_Os03g17490 | expressed protein |
| LOC_Os03g17500 | expressed protein |
| LOC_Os03g17510 | pentatricopeptide, putative, expressed |

| | |
|---|---|
| LOC_Os03g17520 | CDP-alcohol phosphatidyltransferase, putative, expressed |
| LOC_Os03g17540 | TBC domain containing protein, expressed |
| LOC_Os03g17580 | ribosomal protein L10, putative, expressed |
| LOC_Os03g17590 | cat eye syndrome critical region protein 5 precursor, putative, expressed |
| LOC_Os03g17634 | PPR repeat domain containing protein, putative, expressed |
| LOC_Os03g17720 | expressed protein |
| LOC_Os03g17780 | WD domain, G-beta repeat domain containing protein, expressed |
| LOC_Os03g17790 | OsRCI2-5 - Putative low temperature and salt responsive protein, expressed |
| LOC_Os03g17800 | phosphatidylinositol transfer protein alpha isoform, putative, expressed |
| LOC_Os03g17840 | expressed protein |
| LOC_Os03g17860 | OsPDIL5-1 protein disulfide isomerase PDIL5-1, expressed |
| LOC_Os03g17870 | metallothionein, putative, expressed |
| LOC_Os03g17880 | expressed protein |
| LOC_Os03g17900 | hydrolase, alpha/beta fold family protein, putative, expressed |
| LOC_Os03g17920 | RNA pseudouridine synthase, putative, expressed |
| LOC_Os03g17980 | CAMK_KIN1/SNF1/Nim1_like_AMPKh 2 - CAMK includes calcium/calmodulin depedent protein kinases, expressed |
| LOC_Os03g17990 | Yip1 domain containing protein, expressed |
| LOC_Os03g18000 | phosphoinositide-specific phospholipase C, putative, expressed |
| LOC_Os03g18030 | leucoanthocyanidin dioxygenase, putative, expressed |
| LOC_Os03g18050 | OsSAUR13 - Auxin-responsive SAUR gene family member, expressed |
| LOC_Os03g18070 | omega-3 fatty acid desaturase, chloroplast precursor, putative, expressed |
| LOC_Os03g18110 | proteins of unknown function domain containing protein, putative, expressed |
| LOC_Os03g18140 | expressed protein |
| LOC_Os03g18170 | STE_MEKK_ste11_MAP3K 13 - STE kinases include homologs to sterile 7, sterile 11 and sterile 20 from yeast, expressed |
| LOC_Os03g18230 | expressed protein |
| LOC_Os03g18250 | expressed protein |
| LOC_Os03g18330 | transcription factor like protein, putative, expressed |
| LOC_Os03g18400 | expressed protein |

| | |
|---|---|
| LOC_Os03g18470 | expressed protein |
| LOC_Os03g18490 | RPGR, putative, expressed |
| LOC_Os03g18630 | receptor-like kinase RHG1, putative, expressed |
| LOC_Os03g18720 | RNA recognition motif containing protein, putative, expressed |
| LOC_Os03g18729 | expressed protein |
| LOC_Os03g18790 | SAG20, putative, expressed |
| LOC_Os03g18820 | glycosyltransferase, putative, expressed |
| LOC_Os03g18940 | exonuclease, putative, expressed |
| LOC_Os03g18950 | zinc finger C-x8-C-x5-C-x3-H type family protein, expressed |
| LOC_Os03g18960 | calmodulin binding protein, putative, expressed |
| LOC_Os03g18970 | protein phosphotase protein, putative, expressed |
| LOC_Os03g19020 | PHD-finger family protein, expressed |
| LOC_Os03g19059 | anaphase-promoting complex subunit 11, putative, expressed |
| LOC_Os03g19080 | expressed protein |
| LOC_Os03g19190 | TOPBP1C - Similar to DNA replication protein TOPBP1 from, expressed |
| LOC_Os03g19380 | calvin cycle protein CP12, putative, expressed |
| LOC_Os03g19660 | conserved hypothetical protein |
| LOC_Os03g19840 | cysteine-rich repeat secretory protein 55 precursor, putative, expressed |
| LOC_Os03g19860 | expressed protein |
| LOC_Os03g19960 | ATP-dependent RNA helicase, putative, expressed |
| LOC_Os03g20060 | retrotransposon protein, putative, unclassified, expressed |
| LOC_Os03g20090 | MYB family transcription factor, putative, expressed |
| LOC_Os03g20230 | aspartic proteinase nepenthesin-1 precursor, putative, expressed |
| LOC_Os03g20600 | expressed protein |
| LOC_Os03g21140 | RNA-binding zinc finger protein, putative, expressed |
| LOC_Os03g21380 | OsCML27 - Calmodulin-related calcium sensor protein, expressed |
| LOC_Os03g21664 | expressed protein |
| LOC_Os03g21730 | receptor-like protein kinase precursor, putative, expressed |
| LOC_Os03g21780 | pentatricopeptide repeat domain containing protein, putative, expressed |

| | |
|---|---|
| LOC_Os03g21830 | appr-1-p processing enzyme family protein, putative, expressed |
| LOC_Os03g21840 | reticulon domain containing protein, putative, expressed |
| LOC_Os03g21880 | THION28 - Plant thionin family protein precursor, expressed |
| LOC_Os03g21950 | fumarate hydratase, mitochondrial precursor, putative, expressed |
| LOC_Os03g21970 | basic helix-loop-helix domain containing protein, expressed |
| LOC_Os03g21980 | tRNA pseudouridine synthase family protein, putative, expressed |
| LOC_Os03g21990 | WD domain, G-beta repeat domain containing protein, expressed |
| LOC_Os03g21999 | expressed protein |
| LOC_Os03g22360 | expressed protein |
| LOC_Os03g22370 | ultraviolet-B-repressible protein, putative, expressed |
| LOC_Os03g22470 | desiccation-related protein PCC13-62 precursor, putative, expressed |
| LOC_Os03g22655 | tubby-like F-box protein 3, putative, expressed |
| LOC_Os03g22690 | nucleoporin interacting component, putative, expressed |
| LOC_Os03g22760 | hypothetical protein |
| LOC_Os03g22770 | CCT/B-box zinc finger protein, putative, expressed |
| LOC_Os03g22780 | DVR, putative, expressed |
| LOC_Os03g22800 | OsFBT5 - F-box and tubby domain containing protein, expressed |
| LOC_Os03g22820 | expressed protein |
| LOC_Os03g22830 | zinc finger, C3HC4 type domain containing protein, expressed |
| LOC_Os03g22950 | acyl carrier protein, putative, expressed |
| LOC_Os03g22990 | OsFBX84 - F-box domain containing protein, expressed |
| LOC_Os03g23020 | expressed protein |
| LOC_Os03g23040 | expressed protein |
| LOC_Os03g23060 | transposon protein, putative, unclassified, expressed |
| LOC_Os03g23940 | expressed protein |
| LOC_Os03g23950 | adapitin protein, putative, expressed |
| LOC_Os03g23970 | diphthine synthase, putative, expressed |
| LOC_Os03g23980 | NAD dependent epimerase/dehydratase family protein, putative, expressed |
| LOC_Os03g24130 | CXXC1 - Cysteine-rich protein with CXXC and CXXXC motifs precursor, putative, expressed |

| | |
|---|---|
| LOC_Os03g24160 | phosphatidylinositol-4-phosphate 5-kinase, putative, expressed |
| LOC_Os03g24170 | phosphatidylinositol-4-phosphate 5-kinase, putative, expressed |
| LOC_Os03g24184 | TRAF-type zinc finger domain-containing protein 1, putative, expressed |
| LOC_Os03g24200 | OsFBX85 - F-box domain containing protein, expressed |
| LOC_Os03g24339 | PWWP domain containing protein, expressed |
| LOC_Os03g24380 | glutathione peroxidase domain containing protein, expressed |
| LOC_Os03g24950 | expressed protein |
| LOC_Os03g24960 | ulp1 protease family, C-terminal catalytic domain containing protein, expressed |
| LOC_Os03g24970 | SWIM zinc finger family protein, putative |
| LOC_Os03g25070 | CAMK_CAMK_like 18 - CAMK includes calcium/calmodulin depedent protein kinases, expressed |
| LOC_Os03g25130 | OB-fold nucleic acid binding domain containing protein, putative, expressed |
| LOC_Os03g25140 | peptidyl-prolyl isomerase PASTICCINO1, putative, expressed |
| LOC_Os03g25270 | pyrrolidone-carboxylate peptidase, putative, expressed |
| LOC_Os03g25480 | cytochrome P450, putative, expressed |
| LOC_Os03g25640 | OsFBX91 - F-box domain containing protein, expressed |
| LOC_Os03g25700 | retrotransposon protein, putative, Ty3-gypsy subclass, expressed |
| LOC_Os03g25750 | SNARE associated Golgi protein, putative, expressed |
| LOC_Os03g25760 | calmodulin-binding protein, putative, expressed |
| LOC_Os03g25984 | hypothetical protein |
| LOC_Os03g26090 | GPI-anchored wall transfer protein 1, putative, expressed |
| LOC_Os03g26329 | retrotransposon protein, putative, unclassified, expressed |
| LOC_Os03g26440 | tRNA pseudouridine synthase family protein, putative, expressed |
| LOC_Os03g26500 | expressed protein |
| LOC_Os03g26910 | uncharacterized glycosyl hydrolase Rv2006/MT2062, putative, expressed |
| LOC_Os03g26990 | VQ domain containing protein, putative, expressed |
| LOC_Os03g27370 | phospholipase D, putative, expressed |
| LOC_Os03g27610 | patatin, putative, expressed |
| LOC_Os03g27800 | paramyosin, putative, expressed |
| LOC_Os03g29330 | expressed protein |

| | |
|---|---|
| LOC_Os03g29710 | hypothetical protein |
| LOC_Os03g30250 | COBRA-like protein precursor, putative, expressed |
| LOC_Os03g30260 | COBRA-like protein precursor, putative, expressed |
| LOC_Os03g30290 | expressed protein |
| LOC_Os03g30810 | retrotransposon protein, putative, unclassified, expressed |
| LOC_Os03g30950 | acyl-desaturase, chloroplast precursor, putative, expressed |
| LOC_Os03g31594 | jmjC domain containing protein, expressed |
| LOC_Os03g31690 | GCN5-related N-acetyltransferase, putative, expressed |
| LOC_Os03g32526 | tRNA-splicing endonuclease positive effector-related, putative, expressed |
| LOC_Os03g33070 | transposon protein, putative, CACTA, En/Spm sub-class, expressed |
| LOC_Os03g34030 | hypothetical protein |
| LOC_Os03g35490 | retrotransposon, putative, centromere-specific, expressed |
| LOC_Os03g35600 | serine/threonine-protein kinase receptor precursor, putative, expressed |
| LOC_Os03g36830 | expressed protein |
| LOC_Os03g37170 | expressed protein |
| LOC_Os03g37960 | acyl CoA binding protein, putative, expressed |
| LOC_Os03g37970 | ribosomal protein L13, putative, expressed |
| LOC_Os03g38000 | 40S ribosomal protein S3-1, putative, expressed |
| LOC_Os03g38010 | nuf2 family protein, expressed |
| LOC_Os03g38804 | expressed protein |
| LOC_Os03g39150 | protein kinase domain containing protein, expressed |
| LOC_Os03g39660 | expressed protein |
| LOC_Os03g40180 | 60S ribosomal protein L15, putative, expressed |
| LOC_Os03g40710 | ZOS3-13 - C2H2 zinc finger protein, expressed |
| LOC_Os03g40930 | expressed protein |
| LOC_Os03g40950 | expressed protein |
| LOC_Os03g41300 | expressed protein |
| LOC_Os03g41339 | expressed protein |
| LOC_Os03g41380 | expressed protein |

| | |
|---|---|
| LOC_Os03g41419 | serpin domain containing protein, putative, expressed |
| LOC_Os03g41530 | single-stranded DNA-binding protein, putative, expressed |
| LOC_Os03g41570 | transposon protein, putative, unclassified, expressed |
| LOC_Os03g41580 | transposon protein, putative, CACTA, En/Spm sub-class |
| LOC_Os03g41675 | expressed protein |
| LOC_Os03g41740 | retrotransposon protein, putative, unclassified, expressed |
| LOC_Os03g41870 | nucleic acid binding protein, putative |
| LOC_Os03g42010 | DNA-directed polymerase, putative, expressed |
| LOC_Os03g42220 | T-complex protein, putative, expressed |
| LOC_Os03g42235 | expressed protein |
| LOC_Os03g42400 | expressed protein |
| LOC_Os03g42420 | B3 DNA binding domain containing protein, expressed |
| LOC_Os03g42440 | expressed protein |
| LOC_Os03g42820 | LIM domain-containing protein, putative, expressed |
| LOC_Os03g42930 | expressed protein |
| LOC_Os03g43340 | expressed protein |
| LOC_Os03g44230 | transmembrane amino acid transporter protein, putative, expressed |
| LOC_Os03g44310 | initiation factor 2 subunit family domain containing protein, expressed |
| LOC_Os03g44380 | 9-cis-epoxycarotenoid dioxygenase 1, chloroplast precursor, putative, expressed |
| LOC_Os03g44440 | cyclic nucleotide-gated ion channel, putative, expressed |
| LOC_Os03g45230 | expressed protein |
| LOC_Os03g45700 | retrotransposon protein, putative, unclassified |
| LOC_Os03g45710 | 2Fe-2S iron-sulfur cluster binding domain containing protein, expressed |
| LOC_Os03g46940 | expressed protein |
| LOC_Os03g47016 | homeobox protein knotted-1-like 10, putative, expressed |
| LOC_Os03g47140 | growth regulating factor protein, putative, expressed |
| LOC_Os03g47480 | hypothetical protein |
| LOC_Os03g48480 | thioesterase family protein, putative, expressed |
| LOC_Os03g48626 | expressed protein |

| | |
|---|---|
| LOC_Os03g48750 | Cupin domain containing protein, expressed |
| LOC_Os03g48870 | expressed protein |
| LOC_Os03g48880 | receptor protein kinase TMK1 precursor, putative, expressed |
| LOC_Os03g49350 | lipozygenase protein, putative, expressed |
| LOC_Os03g49464 | tetratricopeptide-like helical, putative, expressed |
| LOC_Os03g49480 | elongation of fatty acids protein 2, putative, expressed |
| LOC_Os03g49510 | phosphatidylinositol-4-phosphate 5-kinase, putative, expressed |
| LOC_Os03g49790 | pattern formation protein EMB30, putative, expressed |
| LOC_Os03g49860 | expressed protein |
| LOC_Os03g50190 | expressed protein |
| LOC_Os03g52290 | DUF617 domain containing protein, expressed |
| LOC_Os03g52340 | endoplasmic oxidoreductin-1 precursor, putative, expressed |
| LOC_Os03g52475 | pentatricopeptide, putative, expressed |
| LOC_Os03g52560 | glycosyl hydrolases, putative, expressed |
| LOC_Os03g52640 | expressed protein |
| LOC_Os03g54130 | cysteine protease 1 precursor, putative, expressed |
| LOC_Os03g54790 | ABC transporter, ATP-binding protein, putative, expressed |
| LOC_Os03g55010 | UDP-glucoronosyl and UDP-glucosyl transferase domain containing protein, expressed |
| LOC_Os03g55140 | transducin family protein, putative, expressed |
| LOC_Os03g55210 | TKL_IRAK_CrRLK1L-1 |
| LOC_Os03g55360 | heat shock protein DnaJ, putative, expressed |
| LOC_Os03g55380 | oxidoreductase, putative, expressed |
| LOC_Os03g55600 | CAMK_CAMK_like |
| LOC_Os03g55940 | expressed protein |
| LOC_Os03g55950 | H-BTB1 - Bric-a-Brac, Tramtrack, Broad Complex BTB domain with H family conserved sequence, expressed |
| LOC_Os03g56030 | expressed protein |
| LOC_Os03g56100 | expressed protein |
| LOC_Os03g56170 | expressed protein |
| LOC_Os03g56180 | legume lectins beta domain containing protein, expressed |

| | |
|---|---|
| LOC_Os03g56260 | kinesin motor domain containing protein, putative, expressed |
| LOC_Os03g56270 | receptor protein kinase CLAVATA1 precursor, putative, expressed |
| LOC_Os03g56380 | methyltransferase, putative, expressed |
| LOC_Os03g56390 | tRNA wybutosine-synthesizing protein 1, putative, expressed |
| LOC_Os03g56930 | app1, putative, expressed |
| LOC_Os03g56950 | phytochrome-interacting factor 4, putative, expressed |
| LOC_Os03g57020 | expressed protein |
| LOC_Os03g57050 | expressed protein |
| LOC_Os03g57060 | expressed protein |
| LOC_Os03g57130 | expressed protein |
| LOC_Os03g57160 | zinc ion binding protein, putative, expressed |
| LOC_Os03g57270 | ubiquitin carboxyl-terminal hydrolase, family 1, putative, expressed |
| LOC_Os03g57360 | expressed protein |
| LOC_Os03g57390 | hypothetical protein |
| LOC_Os03g57510 | CAMK_CAMK_like |
| LOC_Os03g57854 | MBTB3 - Bric-a-Brac, Tramtrack, Broad Complex BTB domain with Meprin and TRAF Homology MATH domain, expressed |
| LOC_Os03g58110 | uncharacterized protein At4g06744 precursor, putative, expressed |
| LOC_Os03g59930 | retrotransposon, putative, centromere-specific, expressed |
| LOC_Os03g60500 | expressed protein |
| LOC_Os03g60740 | expressed protein |
| LOC_Os03g61290 | ATCHX, putative, expressed |
| LOC_Os03g62070 | hydrolase, putative, expressed |
| LOC_Os03g62230 | ZOS3-24 - C2H2 zinc finger protein, expressed |
| LOC_Os03g62820 | expressed protein |
| LOC_Os03g63999 | expressed protein |
| LOC_Os03g64190 | la domain containing protein, putative, expressed |
| LOC_Os03g64219 | OTU-like cysteine protease family protein, putative, expressed |
| LOC_Os03g64230 | expressed protein |
| LOC_Os03g64320 | expressed protein |

| | |
|---|---|
| LOC_Os03g64340 | heavy metal-associated domain containing protein, expressed |
| LOC_Os03g64350 | rhodanese-like domain containing protein, putative, expressed |
| LOC_Os03g64360 | OsFBX113 - F-box domain containing protein, expressed |
| LOC_Os03g64370 | pentatricopeptide, putative, expressed |
| LOC_Os04g01130 | chromatin modification-related protein EAF3, putative, expressed |
| LOC_Os04g01140 | cytochrome P450 93A2, putative, expressed |
| LOC_Os04g01160 | zinc finger family protein, putative, expressed |
| LOC_Os04g01170 | expressed protein |
| LOC_Os04g01180 | retrotransposon protein, putative, LINE subclass, expressed |
| LOC_Os04g01380 | retrotransposon protein, putative, Ty3-gypsy subclass, expressed |
| LOC_Os04g01660 | erythronate-4-phosphate dehydrogenase domain containing protein, expressed |
| LOC_Os04g01780 | uncharacterized ACR, COG1399 family protein, expressed |
| LOC_Os04g01990 | pentatricopeptide domain containing protein, putative, expressed |
| LOC_Os04g02100 | expressed protein |
| LOC_Os04g02580 | expressed protein |
| LOC_Os04g02640 | 3-ketoacyl-CoA synthase 6, putative, expressed |
| LOC_Os04g02660 | retrotransposon protein, putative, unclassified |
| LOC_Os04g02730 | tRNA-dihydrouridine synthase 3-like, putative, expressed |
| LOC_Os04g02860 | disease resistance protein RPM1, putative, expressed |
| LOC_Os04g02900 | dehydrogenase E1 component domain containing protein, expressed |
| LOC_Os04g02920 | leucine-rich repeat family protein, putative, expressed |
| LOC_Os04g03370 | cysteine-rich receptor-like protein kinase 41 precursor, putative, expressed |
| LOC_Os04g03380 | legume lectins beta domain containing protein, putative, expressed |
| LOC_Os04g03579 | protein kinase, putative, expressed |
| LOC_Os04g03796 | OsSub37 - Putative Subtilisin homologue, expressed |
| LOC_Os04g03860 | expressed protein |
| LOC_Os04g03884 | transposon protein, putative, CACTA, En/Spm sub-class, expressed |
| LOC_Os04g03910 | expressed protein |
| LOC_Os04g04000 | ECT protein, putative, expressed |

| | |
|---|---|
| LOC_Os04g04240 | sterol 3-beta-glucosyltransferase, putative, expressed |
| LOC_Os04g04310 | hypothetical protein |
| LOC_Os04g04320 | expressed protein |
| LOC_Os04g04650 | hypothetical protein |
| LOC_Os04g04660 | expressed protein |
| LOC_Os04g04680 | expressed protein |
| LOC_Os04g04750 | peroxidase precursor, putative, expressed |
| LOC_Os04g04970 | expressed protein |
| LOC_Os04g05010 | CBS domain containing membrane protein, putative, expressed |
| LOC_Os04g05030 | serine-rich 25 kDa antigen protein, putative, expressed |
| LOC_Os04g05070 | expressed protein |
| LOC_Os04g06770 | argonaute, putative, expressed |
| LOC_Os04g06790 | 2-aminoethanethiol dioxygenase, putative, expressed |
| LOC_Os04g07450 | retrotransposon protein, putative, unclassified, expressed |
| LOC_Os04g08180 | expressed protein |
| LOC_Os04g08310 | expressed protein |
| LOC_Os04g08320 | YIF1B, putative, expressed |
| LOC_Os04g08330 | expressed protein |
| LOC_Os04g08350 | cysteine synthase, chloroplast/chromoplast precursor, putative, expressed |
| LOC_Os04g08400 | retrotransposon protein, putative, unclassified, expressed |
| LOC_Os04g09380 | retrotransposon protein, putative, Ty1-copia subclass, expressed |
| LOC_Os04g09420 | hypothetical protein |
| LOC_Os04g09920 | cytochrome P450, putative, expressed |
| LOC_Os04g10150 | expressed protein |
| LOC_Os04g10180 | expressed protein |
| LOC_Os04g10214 | expressed protein |
| LOC_Os04g10219 | expressed protein |
| LOC_Os04g10230 | expressed protein |
| LOC_Os04g10240 | nicotiana lesion-inducing like, putative, expressed |

| | |
|---|---|
| LOC_Os04g10260 | basic region leucine zipper domain containing protein, expressed |
| LOC_Os04g10290 | expressed protein |
| LOC_Os04g10380 | glycine-rich protein, putative, expressed |
| LOC_Os04g10410 | amidase, putative, expressed |
| LOC_Os04g11165 | gamma-thionin family domain containing protein, expressed |
| LOC_Os04g11195 | gamma-thionin family domain containing protein, expressed |
| LOC_Os04g11270 | expressed protein |
| LOC_Os04g11390 | expressed protein |
| LOC_Os04g11820 | white-brown complex homolog protein, putative, expressed |
| LOC_Os04g11880 | WD domain, G-beta repeat domain containing protein, expressed |
| LOC_Os04g11890 | OsFBX121 - F-box domain containing protein, expressed |
| LOC_Os04g12190 | hAT dimerisation domain-containing protein, putative, expressed |
| LOC_Os04g12370 | expressed protein |
| LOC_Os04g12499 | amino acid transporter protein, putative, expressed |
| LOC_Os04g12540 | receptor-like protein kinase, putative, expressed |
| LOC_Os04g13130 | hypothetical protein |
| LOC_Os04g13160 | OsFBD9 - F-box and FBD domain containing protein, expressed |
| LOC_Os04g13190 | expressed protein |
| LOC_Os04g13200 | expressed protein |
| LOC_Os04g13220 | ABC transporter family protein, putative, expressed |
| LOC_Os04g13470 | expressed protein |
| LOC_Os04g13500 | retrotransposon protein, putative, unclassified, expressed |
| LOC_Os04g14110 | TRAF-type zinc finger family protein, expressed |
| LOC_Os04g14130 | pentatricopeptide, putative, expressed |
| LOC_Os04g14350 | expressed protein |
| LOC_Os04g14450 | pentatricopeptide, putative, expressed |
| LOC_Os04g14510 | expressed protein |
| LOC_Os04g14654 | Sec1 family transport protein, putative, expressed |
| LOC_Os04g14740 | expressed protein |

| | |
|---|---|
| LOC_Os04g14790 | FAD dependent oxidoreductase domain containing protein, expressed |
| LOC_Os04g15000 | expressed protein |
| LOC_Os04g15600 | receptor kinase, putative, expressed |
| LOC_Os04g15630 | xa21, putative, expressed |
| LOC_Os04g15790 | eukaryotic aspartyl protease domain containing protein, expressed |
| LOC_Os04g15800 | expressed protein |
| LOC_Os04g16160 | expressed protein |
| LOC_Os04g16270 | nucleic acid binding protein, putative, expressed |
| LOC_Os04g16742 | NADPH-dependent oxidoreductase, putative, expressed |
| LOC_Os04g16772 | chloroplast 30S ribosomal protein S19, putative |
| LOC_Os04g16792 | uncharacterized protein ycf76, putative |
| LOC_Os04g16794 | hypothetical protein |
| LOC_Os04g17080 | transposon protein, putative, unclassified |
| LOC_Os04g17150 | hypothetical protein |
| LOC_Os04g18010 | cleavage and polyadenylation specificity factor subunit 1, putative, expressed |
| LOC_Os04g19470 | transposon protein, putative, CACTA, En/Spm sub-class |
| LOC_Os04g19500 | retrotransposon protein, putative, unclassified, expressed |
| LOC_Os04g19580 | retrotransposon protein, putative, unclassified, expressed |
| LOC_Os04g19630 | expressed protein |
| LOC_Os04g19670 | retrotransposon protein, putative, Ty3-gypsy subclass, expressed |
| LOC_Os04g19750 | OsFBL11 - F-box domain and LRR containing protein, expressed |
| LOC_Os04g19810 | OsFBL12 - F-box domain and LRR containing protein, expressed |
| LOC_Os04g19920 | OsFBL13 - F-box domain and LRR containing protein, expressed |
| LOC_Os04g20164 | amine oxidase precursor, putative, expressed |
| LOC_Os04g20230 | neighbor of COX4, putative, expressed |
| LOC_Os04g20424 | neural Wiskott-Aldrich syndrome protein, putative, expressed |
| LOC_Os04g20890 | retrotransposon protein, putative, unclassified, expressed |
| LOC_Os04g20960 | expressed protein |
| LOC_Os04g21020 | expressed protein |

| | |
|---|---|
| LOC_Os04g21130 | F-box protein PP2-B1, putative, expressed |
| LOC_Os04g21160 | triacylglycerol lipase 1 precursor, putative, expressed |
| LOC_Os04g21320 | membrane associated DUF588 domain containing protein, putative, expressed |
| LOC_Os04g21340 | expressed protein |
| LOC_Os04g21360 | expressed protein |
| LOC_Os04g21370 | expressed protein |
| LOC_Os04g22820 | expressed protein |
| LOC_Os04g23180 | cation efflux family protein, putative, expressed |
| LOC_Os04g23190 | retrotransposon protein, putative, unclassified, expressed |
| LOC_Os04g23200 | expressed protein |
| LOC_Os04g23210 | expressed protein |
| LOC_Os04g23440 | helix-loop-helix DNA-binding domain containing protein, expressed |
| LOC_Os04g24190 | growth-regulating factor 11, putative, expressed |
| LOC_Os04g25280 | retrotransposon protein, putative, unclassified, expressed |
| LOC_Os04g25790 | expressed protein |
| LOC_Os04g26470 | hypothetical protein |
| LOC_Os04g26870 | oxidoreductase, aldo/keto reductase family protein, putative, expressed |
| LOC_Os04g28630 | hypothetical protein |
| LOC_Os04g29100 | hypothetical protein |
| LOC_Os04g30410 | expressed protein |
| LOC_Os04g30570 | transferase family protein, putative, expressed |
| LOC_Os04g31210 | bile acid sodium symporter family protein, putative, expressed |
| LOC_Os04g31230 | expressed protein |
| LOC_Os04g31260 | hypothetical protein |
| LOC_Os04g31320 | SWIB/MDM2 domain containing protein, expressed |
| LOC_Os04g31390 | vacuolar protein sorting-associated protein 11, putative, expressed |
| LOC_Os04g31580 | OsFBX131 - F-box domain containing protein, expressed |
| LOC_Os04g31610 | OsFBO3 - F-box and other domain containing protein, expressed |
| LOC_Os04g31620 | retrotransposon protein, putative, unclassified, expressed |

| | |
|---|---|
| LOC_Os04g32000 | expressed protein |
| LOC_Os04g32690 | expressed protein |
| LOC_Os04g33280 | expressed protein |
| LOC_Os04g33370 | cytochrome P450, putative, expressed |
| LOC_Os04g33380 | expressed protein |
| LOC_Os04g33520 | RCLEA6 - Root cap and Late embryogenesis related family protein precursor, expressed |
| LOC_Os04g33530 | Ser/Thr protein phosphatase family protein, putative, expressed |
| LOC_Os04g33550 | expressed protein |
| LOC_Os04g33590 | hydrolase, alpha/beta fold family protein, putative, expressed |
| LOC_Os04g33610 | expressed protein |
| LOC_Os04g33690 | sec23/Sec24 trunk domain containing protein, expressed |
| LOC_Os04g33720 | glycosyl hydrolases, putative, expressed |
| LOC_Os04g33760 | DUF584 domain containing protein, putative, expressed |
| LOC_Os04g33810 | RNA recognition motif containing protein, putative, expressed |
| LOC_Os04g33880 | HVA22, putative, expressed |
| LOC_Os04g33885 | expressed protein |
| LOC_Os04g33890 | expressed protein |
| LOC_Os04g33920 | LTPL102 - Protease inhibitor/seed storage/LTP family protein precursor, expressed |
| LOC_Os04g34360 | serine/threonine-protein kinase receptor precursor, putative, expressed |
| LOC_Os04g34699 | expressed protein |
| LOC_Os04g34720 | PHD-finger domain containing protein, putative, expressed |
| LOC_Os04g35620 | expressed protein |
| LOC_Os04g36040 | peptide transporter PTR2, putative, expressed |
| LOC_Os04g37430 | lipoxygenase protein, putative, expressed |
| LOC_Os04g38060 | retrotransposon protein, putative, unclassified, expressed |
| LOC_Os04g38610 | RAN guanine nucleotide release factor, putative, expressed |
| LOC_Os04g38885 | expressed protein |
| LOC_Os04g38990 | expressed protein |
| LOC_Os04g39530 | transposon protein, putative, CACTA, En/Spm sub-class, expressed |

| | |
|---|---|
| LOC_Os04g40190 | circumsporozoite protein, putative, expressed |
| LOC_Os04g40600 | peptide methionine sulfoxide reductase, putative, expressed |
| LOC_Os04g40950 | glyceraldehyde-3-phosphate dehydrogenase, putative, expressed |
| LOC_Os04g41010 | expressed protein |
| LOC_Os04g41980 | ATOZI1, putative, expressed |
| LOC_Os04g41990 | autophagy-related protein 10, putative, expressed |
| LOC_Os04g42030 | expressed protein |
| LOC_Os04g42100 | retrotransposon protein, putative, unclassified, expressed |
| LOC_Os04g42130 | integral membrane transporter family protein, putative, expressed |
| LOC_Os04g42134 | enhancer of rudimentary protein, putative, expressed |
| LOC_Os04g42140 | eukaryotic initiation factor iso-4F subunit p82-34, putative, expressed |
| LOC_Os04g42150 | hypothetical protein |
| LOC_Os04g42320 | AT hook motif family protein, expressed |
| LOC_Os04g42340 | expressed protein |
| LOC_Os04g42350 | heavy metal-associated domain containing protein, expressed |
| LOC_Os04g42360 | expressed protein |
| LOC_Os04g42370 | expressed protein |
| LOC_Os04g43770 | expressed protein |
| LOC_Os04g43840 | possible lysine decarboxylase domain containing protein, expressed |
| LOC_Os04g43880 | expressed protein |
| LOC_Os04g44020 | expressed protein |
| LOC_Os04g44050 | expressed protein |
| LOC_Os04g44070 | expressed protein |
| LOC_Os04g44090 | expressed protein |
| LOC_Os04g44110 | hydrolase, putative, expressed |
| LOC_Os04g44120 | collagen triple helix repeat, putative, expressed |
| LOC_Os04g44130 | DEF12 - Defensin and Defensin-like DEFL family, expressed |
| LOC_Os04g44160 | hypothetical protein |
| LOC_Os04g44600 | CRP1 - Cysteine-rich family protein precursor, expressed |

| | |
|---|---|
| LOC_Os04g44610 | white-brown complex homolog protein 11, putative, expressed |
| LOC_Os04g44700 | WD-40 repeat family protein, putative, expressed |
| LOC_Os04g44710 | double-stranded RNA binding motif containing protein, expressed |
| LOC_Os04g44730 | lanC-like protein 2, putative, expressed |
| LOC_Os04g44740 | glycosyltransferase sugar-binding region containing DXD motif, putative, expressed |
| LOC_Os04g44880 | expressed protein |
| LOC_Os04g44890 | tRNA-dihydrouridine synthase, putative, expressed |
| LOC_Os04g44920 | oxidoreductase, short chain dehydrogenase/reductase family domain containing protein, expressed |
| LOC_Os04g44924 | short-chain dehydrogenase/reductase, putative, expressed |
| LOC_Os04g44940 | hypothetical protein |
| LOC_Os04g45070 | remorin, putative, expressed |
| LOC_Os04g45080 | expressed protein |
| LOC_Os04g45110 | serpin domain containing protein, putative, expressed |
| LOC_Os04g45360 | retrotransposon protein, putative, unclassified, expressed |
| LOC_Os04g45490 | elongation factor, putative, expressed |
| LOC_Os04g45500 | expressed protein |
| LOC_Os04g45510 | expressed protein |
| LOC_Os04g45590 | glyoxalase family protein, putative, expressed |
| LOC_Os04g45730 | protein kinase domain containing protein, expressed |
| LOC_Os04g45800 | diacylglycerol kinase, putative, expressed |
| LOC_Os04g45821 | expressed protein |
| LOC_Os04g45870 | expressed protein |
| LOC_Os04g45910 | placental protein 11 precursor, putative, expressed |
| LOC_Os04g46030 | expressed protein |
| LOC_Os04g46040 | hypothetical protein |
| LOC_Os04g46120 | plastocyanin-like domain containing protein, putative, expressed |
| LOC_Os04g46430 | expressed protein |
| LOC_Os04g47420 | transmembrane amino acid transporter protein, putative, expressed |
| LOC_Os04g47555 | expressed protein |

| | |
|---|---|
| LOC_Os04g47610 | hypothetical protein |
| LOC_Os04g49160 | zinc finger, C3HC4 type domain containing protein, expressed |
| LOC_Os04g49990 | AT hook motif domain containing protein, expressed |
| LOC_Os04g51440 | villin protein, putative, expressed |
| LOC_Os04g51794 | DNA binding protein, putative, expressed |
| LOC_Os04g51910 | proteasome/cyclosome repeat containing protein, expressed |
| LOC_Os04g51980 | transferase family domain containing protein, expressed |
| LOC_Os04g52020 | PHD-finger domain containing protein, putative, expressed |
| LOC_Os04g52140 | protein kinase domain containing protein, expressed |
| LOC_Os04g52361 | ribosomal protein S17, putative, expressed |
| LOC_Os04g52690 | AAA family ATPase, putative, expressed |
| LOC_Os04g52810 | no apical meristem protein, putative, expressed |
| LOC_Os04g52830 | OsFBK15 - F-box domain and kelch repeat containing protein, expressed |
| LOC_Os04g53030 | NBS-LRR disease resistance protein, putative, expressed |
| LOC_Os04g53200 | exonuclease, putative, expressed |
| LOC_Os04g53400 | MBTB6 - Bric-a-Brac, Tramtrack, Broad Complex BTB domain with Meprin and TRAF Homology MATH domain, expressed |
| LOC_Os04g53496 | NBS-LRR disease resistance protein, putative, expressed |
| LOC_Os04g53540 | homeobox and START domains containing protein, putative, expressed |
| LOC_Os04g53720 | SNF2 family N-terminal domain containing protein, expressed |
| LOC_Os04g54180 | serine/threonine-protein kinase receptor precursor, putative, expressed |
| LOC_Os04g54200 | diacylglycerol kinase, putative, expressed |
| LOC_Os04g54474 | transcription factor, putative, expressed |
| LOC_Os04g54820 | programmed cell death 2 C-terminal domain-containing protein, putative, expressed |
| LOC_Os04g54860 | glutaredoxin, putative, expressed |
| LOC_Os04g54960 | expressed protein |
| LOC_Os04g55060 | 3-oxoacyl-synthase III, chloroplast precursor, putative, expressed |
| LOC_Os04g55180 | hydrolase, alpha/beta fold family domain containing protein, expressed |
| LOC_Os04g55360 | ubiquitin carboxyl-terminal hydrolase domain containing protein, expressed |
| LOC_Os04g55370 | plant protein of unknown function domain containing protein, expressed |

| | |
|---|---|
| LOC_Os04g55510 | zinc finger, C3HC4 type domain containing protein, expressed |
| LOC_Os04g55550 | expressed protein |
| LOC_Os04g55570 | plant protein of unknown function DUF869 domain containing protein, expressed |
| LOC_Os04g55879 | retrotransposon protein, putative, unclassified, expressed |
| LOC_Os04g55950 | expressed protein |
| LOC_Os04g55970 | AP2-like ethylene-responsive transcription factor AINTEGUMENTA, putative, expressed |
| LOC_Os04g56030 | glycine-rich cell wall structural protein precursor, putative, expressed |
| LOC_Os04g56050 | expressed protein |
| LOC_Os04g56480 | pelota, putative, expressed |
| LOC_Os04g56520 | alpha-1,4-glucan-protein synthase, putative, expressed |
| LOC_Os04g56530 | STE_MEKK_ste11_MAP3K 1 - STE kinases include homologs to sterile 7, sterile 11 and sterile 20 from yeast, expressed |
| LOC_Os04g56610 | expressed protein |
| LOC_Os04g56720 | RCC2, putative, expressed |
| LOC_Os04g56730 | repressor of RNA polymerase III transcription MAF1, putative, expressed |
| LOC_Os04g56770 | expressed protein |
| LOC_Os04g56790 | TLD family protein, putative, expressed |
| LOC_Os04g56990 | Myb-like DNA-binding domain containing protein, putative, expressed |
| LOC_Os04g57480 | OTU-like cysteine protease family protein, putative, expressed |
| LOC_Os04g59540 | phosphatidylinositol-4-phosphate 5-Kinase, putative, expressed |
| LOC_Os04g59550 | phosphate/phosphate translocator, putative, expressed |
| LOC_Os04g59600 | pentatricopeptide, putative, expressed |
| LOC_Os05g01030 | phospholipid-transporting ATPase, putative, expressed |
| LOC_Os05g01050 | DNA-binding protein-related, putative, expressed |
| LOC_Os05g01080 | expressed protein |
| LOC_Os05g01090 | pex14, putative, expressed |
| LOC_Os05g01100 | expressed protein |
| LOC_Os05g01110 | ribosomal protein L28 protein, putative, expressed |
| LOC_Os05g01120 | cytochrome P450, putative, expressed |
| LOC_Os05g01230 | zinc finger, C3HC4 type domain containing protein, expressed |

| | |
|---|---|
| LOC_Os05g01240 | AML1, putative, expressed |
| LOC_Os05g01256 | helix-loop-helix DNA-binding domain containing protein, expressed |
| LOC_Os05g01270 | peptidyl-prolyl cis-trans isomerase, putative, expressed |
| LOC_Os05g01360 | expressed protein |
| LOC_Os05g01470 | methionine S-methyltransferase, putative, expressed |
| LOC_Os05g01500 | tubulin-specific chaperone E, putative, expressed |
| LOC_Os05g01530 | expressed protein |
| LOC_Os05g01540 | serine/arginine repetitive matrix protein 1, putative, expressed |
| LOC_Os05g01590 | heat shock protein DnaJ, putative, expressed |
| LOC_Os05g01620 | OsFBX155 - F-box domain containing protein, expressed |
| LOC_Os05g01730 | drought induced 19 protein, putative, expressed |
| LOC_Os05g01750 | TruB family pseudouridylate synthase, putative, expressed |
| LOC_Os05g01780 | STE_PAK_Ste20++TranslationKinase_Slob_Wnk |
| LOC_Os05g01790 | expressed protein |
| LOC_Os05g01820 | cytochrome b5-like Heme/Steroid binding domain containing protein, expressed |
| LOC_Os05g01830 | expressed protein |
| LOC_Os05g01840 | hypothetical protein |
| LOC_Os05g01890 | expressed protein |
| LOC_Os05g01920 | inhibitor I family protein, putative, expressed |
| LOC_Os05g01990 | DEAD-box ATP-dependent RNA helicase, putative, expressed |
| LOC_Os05g02010 | expressed protein |
| LOC_Os05g02030 | OB-fold nucleic acid binding domain containing protein, putative, expressed |
| LOC_Os05g02060 | mitochondrial import inner membrane translocase subunit Tim17, putative, expressed |
| LOC_Os05g02100 | expressed protein |
| LOC_Os05g02200 | cysteine-rich repeat secretory protein 55 precursor, putative, expressed |
| LOC_Os05g02210 | pentatricopeptide, putative, expressed |
| LOC_Os05g02220 | ACT domain containing protein, expressed |
| LOC_Os05g02230 | expressed protein |
| LOC_Os05g02360 | hypothetical protein |

| | |
|---|---|
| LOC_Os05g02700 | expressed protein |
| LOC_Os05g02940 | calcium-transporting ATPase 2, endoplasmic reticulum-type, putative, expressed |
| LOC_Os05g03820 | glutamate--cysteine ligase, chloroplast precursor, putative, expressed |
| LOC_Os05g04100 | expressed protein |
| LOC_Os05g04160 | pentatricopeptide containing protein, putative, expressed |
| LOC_Os05g04170 | AMP-binding enzyme, putative, expressed |
| LOC_Os05g04215 | expressed protein |
| LOC_Os05g04220 | nitrogen regulatory protein P-II, putative, expressed |
| LOC_Os05g04250 | expressed protein |
| LOC_Os05g04260 | retrotransposon protein, putative, unclassified, expressed |
| LOC_Os05g04370 | expressed protein |
| LOC_Os05g04530 | IF, putative, expressed |
| LOC_Os05g04540 | expressed protein |
| LOC_Os05g04680 | expressed protein |
| LOC_Os05g04740 | helix-loop-helix DNA-binding domain containing protein, expressed |
| LOC_Os05g04760 | expressed protein |
| LOC_Os05g04840 | expressed protein |
| LOC_Os05g04860 | agenet domain containing protein, putative, expressed |
| LOC_Os05g04870 | oxidoreductase, short chain dehydrogenase/reductase family protein, putative, expressed |
| LOC_Os05g04890 | expressed protein |
| LOC_Os05g04950 | protein binding protein, putative, expressed |
| LOC_Os05g04990 | adenosylmethionine decarboxylase, putative, expressed |
| LOC_Os05g05020 | expressed protein |
| LOC_Os05g05040 | expressed protein |
| LOC_Os05g05100 | expressed protein |
| LOC_Os05g05110 | expressed protein |
| LOC_Os05g05140 | expressed protein |
| LOC_Os05g05150 | SLD5 - Putative GINS complex subunit, expressed |
| LOC_Os05g05160 | CGMC_MAPKCGMC_2 1 - CGMC includes CDA, MAPK, GSK3, and CLKC kinases, expressed |

| | |
|---|---|
| LOC_Os05g05170 | jacalin-like lectin domain containing protein, expressed |
| LOC_Os05g05200 | O-acyltransferase, putative, expressed |
| LOC_Os05g05530 | expressed protein |
| LOC_Os05g05860 | retrotransposon protein, putative, unclassified, expressed |
| LOC_Os05g06230 | expressed protein |
| LOC_Os05g06490 | RNA polymerase Rpb7, N-terminal domain containing protein, expressed |
| LOC_Os05g06830 | hypothetical protein |
| LOC_Os05g06840 | POLE2A - Putative DNA polymerase epsilon complex subunit, expressed |
| LOC_Os05g06890 | relA-SpoT like protein RSH4, putative, expressed |
| LOC_Os05g06900 | hypothetical protein |
| LOC_Os05g06990 | TKL_IRAK_CrRLK1L-1 10 - The CrRLK1L-1 subfamily has homology to the CrRLK1L homolog, expressed |
| LOC_Os05g07000 | splicing factor, arginine/serine-rich 7, putative, expressed |
| LOC_Os05g07010 | myb-like DNA-binding domain containing protein, expressed |
| LOC_Os05g07050 | pre-mRNA-processing-splicing factor 8, putative, expressed |
| LOC_Os05g07070 | zinc finger, C3HC4 type domain containing protein, expressed |
| LOC_Os05g07090 | acyl-coenzyme A dehydrogenase, mitochondrial precursor, putative, expressed |
| LOC_Os05g07960 | F-box protein interaction domain containing protein, expressed |
| LOC_Os05g08340 | expressed protein |
| LOC_Os05g08600 | splicing factor, putative, expressed |
| LOC_Os05g09708 | expressed protein |
| LOC_Os05g10270 | expressed protein |
| LOC_Os05g10730 | ABC transporter, ATP-binding protein, putative, expressed |
| LOC_Os05g10810 | O-acyltransferase, putative, expressed |
| LOC_Os05g10940 | metal cation transporter, putative, expressed |
| LOC_Os05g11820 | retrotransposon protein, putative, unclassified, expressed |
| LOC_Os05g11930 | expressed protein |
| LOC_Os05g12180 | chalcone synthase, putative, expressed |
| LOC_Os05g12380 | expressed protein |
| LOC_Os05g12400 | BURP domain containing protein, expressed |

| | |
|---|---|
| LOC_Os05g12420 | retrotransposon protein, putative, Ty3-gypsy subclass |
| LOC_Os05g12474 | expressed protein |
| LOC_Os05g13480 | S-adenosylmethionine decarboxylase proenzyme, putative, expressed |
| LOC_Os05g13530 | expressed protein |
| LOC_Os05g13760 | expressed protein |
| LOC_Os05g13900 | retrotransposon protein, putative, unclassified, expressed |
| LOC_Os05g13970 | phosphorylase domain containing protein, putative, expressed |
| LOC_Os05g13980 | hypothetical protein |
| LOC_Os05g14280 | RNA-directed DNA polymerase, putative, expressed |
| LOC_Os05g14360 | expressed protein |
| LOC_Os05g14370 | WRKY82, expressed |
| LOC_Os05g14390 | expressed protein |
| LOC_Os05g14540 | hypothetical protein |
| LOC_Os05g14600 | retrotransposon protein, putative, unclassified, expressed |
| LOC_Os05g14820 | transmembrane amino acid transporter protein, putative, expressed |
| LOC_Os05g16000 | expressed protein |
| LOC_Os05g16140 | expressed protein |
| LOC_Os05g16170 | expressed protein |
| LOC_Os05g16315 | translation initiation factor IF-2, putative, expressed |
| LOC_Os05g16360 | expressed protein |
| LOC_Os05g16410 | retrotransposon, putative, centromere-specific, expressed |
| LOC_Os05g16430 | SHR5-receptor-like kinase, putative, expressed |
| LOC_Os05g16460 | expressed protein |
| LOC_Os05g16630 | thioredoxin domain-containing protein 9, putative, expressed |
| LOC_Os05g16660 | WD domain, G-beta repeat domain containing protein, expressed |
| LOC_Os05g18670 | C4-dicarboxylate transporter/malic acid transport protein domain containing protein, expressed |
| LOC_Os05g18770 | Sad1 / UNC-like C-terminal domain containing protein, putative, expressed |
| LOC_Os05g18800 | expressed protein |
| LOC_Os05g18860 | STIGL2 - Stigma-specific Stig1 family protein precursor, putative, expressed |

| | |
|---|---|
| LOC_Os05g18870 | hypothetical protein |
| LOC_Os05g18950 | pentatricopeptide, putative, expressed |
| LOC_Os05g18960 | expressed protein |
| LOC_Os05g18970 | expressed protein |
| LOC_Os05g19000 | expressed protein |
| LOC_Os05g19970 | ZOS5-06 - C2H2 zinc finger protein, expressed |
| LOC_Os05g20070 | retrotransposon protein, putative, unclassified |
| LOC_Os05g20914 | expressed protein |
| LOC_Os05g21240 | expressed protein |
| LOC_Os05g22420 | retrotransposon protein, putative, unclassified, expressed |
| LOC_Os05g23250 | Leucine Rich Repeat family protein, expressed |
| LOC_Os05g24160 | double-stranded RNA binding motif containing protein, expressed |
| LOC_Os05g24990 | transposon protein, putative, unclassified, expressed |
| LOC_Os05g25310 | acyl-CoA synthetase protein, putative, expressed |
| LOC_Os05g25400 | RNA binding protein, putative, expressed |
| LOC_Os05g25940 | expressed protein |
| LOC_Os05g26210 | ATOFP17/OFP17, putative, expressed |
| LOC_Os05g26240 | PROLM1 - Prolamin precursor, expressed |
| LOC_Os05g26630 | expressed protein |
| LOC_Os05g26680 | retrotransposon protein, putative, unclassified, expressed |
| LOC_Os05g26840 | permease domain containing protein, putative, expressed |
| LOC_Os05g27980 | DUF260 domain containing protein, putative, expressed |
| LOC_Os05g28490 | FAD-binding domain-containing protein, putative, expressed |
| LOC_Os05g28520 | MRH1, putative, expressed |
| LOC_Os05g28780 | GCRP10 - Glycine and cysteine rich family protein precursor, expressed |
| LOC_Os05g29000 | expressed protein |
| LOC_Os05g29010 | POLA4 - Putative DNA polymerase alpha complex subunit, expressed |
| LOC_Os05g29080 | expressed protein |
| LOC_Os05g29676 | RING-H2 finger protein, putative, expressed |

| | |
|---|---|
| LOC_Os05g29790 | pectinesterase, putative, expressed |
| LOC_Os05g30030 | expressed protein |
| LOC_Os05g30180 | hypothetical protein |
| LOC_Os05g30240 | PPR repeat domain containing protein, putative, expressed |
| LOC_Os05g30250 | Os5bglu19 - beta-glucosidase homologue, similar to G |
| LOC_Os05g30340 | expressed protein |
| LOC_Os05g31010 | hypothetical protein |
| LOC_Os05g32090 | expressed protein |
| LOC_Os05g33190 | endo-1,3;1,4-beta-D-glucanase precursor, putative, expressed |
| LOC_Os05g33230 | cobalt ion transporter, putative, expressed |
| LOC_Os05g33260 | 1-5-phosphoribosyl-5-5-phosphoribosylaminomethylideneaminoimidazole-4-carboxamide isomerase, chloroplast precursor, putative, expressed |
| LOC_Os05g33280 | thylakoid lumen protein, putative, expressed |
| LOC_Os05g33300 | tat pathway signal sequence family protein, expressed |
| LOC_Os05g33360 | transmembrane BAX inhibitor motif-containing protein, putative, expressed |
| LOC_Os05g33400 | basic 7S globulin precursor, putative, expressed |
| LOC_Os05g33514 | expressed protein |
| LOC_Os05g33730 | gibberellin receptor GID1L2, putative, expressed |
| LOC_Os05g33810 | OsSPL9 - SBP-box gene family member, expressed |
| LOC_Os05g33850 | UBA/TS-N domain containing protein, expressed |
| LOC_Os05g33860 | expressed protein |
| LOC_Os05g34290 | glutathione gamma-glutamylcysteinyltransferase 1, putative, expressed |
| LOC_Os05g34460 | OsDegp7 - Putative Deg protease homologue, expressed |
| LOC_Os05g34470 | expressed protein |
| LOC_Os05g34480 | expressed protein |
| LOC_Os05g34670 | KH domain containing protein, putative, expressed |
| LOC_Os05g35140 | nodulin MtN3 family protein, putative, expressed |
| LOC_Os05g35490 | hypothetical protein |
| LOC_Os05g36140 | expressed protein |
| LOC_Os05g36320 | expressed protein |

| | |
|---|---|
| LOC_Os05g36940 | retrotransposon protein, putative, unclassified, expressed |
| LOC_Os05g38350 | glycerol-3-phosphate acyltransferase 8, putative, expressed |
| LOC_Os05g39140 | expressed protein |
| LOC_Os05g40300 | expressed protein |
| LOC_Os05g40310 | expressed protein |
| LOC_Os05g40320 | PPR repeat domain containing protein, putative, expressed |
| LOC_Os05g40700 | transmembrane protein 56, putative, expressed |
| LOC_Os05g40820 | ribosomal protein L24, putative, expressed |
| LOC_Os05g41200 | OsCML9 - Calmodulin-related calcium sensor protein, expressed |
| LOC_Os05g41300 | expressed protein |
| LOC_Os05g41440 | cytochrome P450, putative, expressed |
| LOC_Os05g43530 | CSLC7 - cellulose synthase-like family C, expressed |
| LOC_Os05g44200 | GDSL-like lipase/acylhydrolase, putative, expressed |
| LOC_Os05g44570 | histidine-containing phosphotransfer protein, putative, expressed |
| LOC_Os05g45070 | harpin-induced protein 1 domain containing protein, expressed |
| LOC_Os05g45400 | expressed protein |
| LOC_Os05g45460 | POEI52 - Pollen Ole e I allergen and extensin family protein precursor, expressed |
| LOC_Os05g45470 | hypothetical protein |
| LOC_Os05g45520 | expressed protein |
| LOC_Os05g45580 | expressed protein |
| LOC_Os05g45760 | EMB1796, putative, expressed |
| LOC_Os05g45770 | Divergent PAP2 family domain containing protein, expressed |
| LOC_Os05g46070 | hAT dimerisation domain containing protein |
| LOC_Os05g46170 | expressed protein |
| LOC_Os05g50850 | expressed protein |
| LOC_Os05g51530 | vacuolar ATP synthase subunit C, putative, expressed |
| LOC_Os05g51640 | adhesion regulating molecule family, putative, expressed |
| LOC_Os05g51670 | NAD dependent epimerase/dehydratase family protein, putative, expressed |
| LOC_Os05g51750 | aspartyl protease family protein, putative, expressed |

| | |
|---|---|
| LOC_Os06g01140 | expressed protein |
| LOC_Os06g01200 | zinc finger, C3HC4 type domain containing protein, expressed |
| LOC_Os06g01250 | cytochrome P450, putative, expressed |
| LOC_Os06g01290 | expressed protein |
| LOC_Os06g01304 | spotted leaf 11, putative, expressed |
| LOC_Os06g01340 | CCT/B-box zinc finger protein, putative, expressed |
| LOC_Os06g01350 | transferase family protein, putative, expressed |
| LOC_Os06g01370 | DNA-binding protein-related, putative, expressed |
| LOC_Os06g01400 | glycyl-tRNA synthetase 2, chloroplast/mitochondrial precursor, putative, expressed |
| LOC_Os06g01410 | glutamine cyclotransferase precursor, putative, expressed |
| LOC_Os06g01440 | integral membrane protein, putative, expressed |
| LOC_Os06g01500 | IQ calmodulin-binding motif family protein, putative, expressed |
| LOC_Os06g01600 | expressed protein |
| LOC_Os06g01680 | expressed protein |
| LOC_Os06g01690 | septum site-determining protein minD, putative, expressed |
| LOC_Os06g01700 | CWC15 homolog A, putative, expressed |
| LOC_Os06g01780 | ethylene-responsive element-binding protein, putative, expressed |
| LOC_Os06g01840 | nodulin, putative, expressed |
| LOC_Os06g01890 | MADS-box transcription factor, putative, expressed |
| LOC_Os06g01950 | phosphoglycerate mutase, putative, expressed |
| LOC_Os06g01966 | auxin-induced protein 5NG4, putative, expressed |
| LOC_Os06g01980 | AAA family ATPase, putative, expressed |
| LOC_Os06g02000 | adenylate kinase, putative, expressed |
| LOC_Os06g02019 | cytochrome P450, putative, expressed |
| LOC_Os06g02040 | late embryogenesis abundant group 1, putative, expressed |
| LOC_Os06g02054 | OsFBX180 - F-box domain containing protein, expressed |
| LOC_Os06g02090 | expressed protein |
| LOC_Os06g02100 | OsFBX181 - F-box domain containing protein, expressed |
| LOC_Os06g02110 | OsFBX182 - F-box domain containing protein, expressed |

| | |
|---|---|
| LOC_Os06g02120 | PPR repeat domain containing protein, putative, expressed |
| LOC_Os06g02130 | G-protein alpha subunit, putative, expressed |
| LOC_Os06g02144 | 6-phosphogluconate dehydrogenase, decarboxylating, putative, expressed |
| LOC_Os06g02160 | AMP-binding enzyme family protein, expressed |
| LOC_Os06g02180 | CSLD2 - cellulose synthase-like family D, expressed |
| LOC_Os06g02190 | expressed protein |
| LOC_Os06g02200 | pentatricopeptide, putative, expressed |
| LOC_Os06g02210 | bark storage protein A precursor, putative, expressed |
| LOC_Os06g02240 | RNA recognition motif containing protein, putative, expressed |
| LOC_Os06g02370 | expressed protein |
| LOC_Os06g02380 | T-complex protein, putative, expressed |
| LOC_Os06g02390 | ADP-ribosylation factor 1, putative, expressed |
| LOC_Os06g02450 | expressed protein |
| LOC_Os06g02470 | ATOZI1, putative, expressed |
| LOC_Os06g02500 | superoxide dismutase, chloroplast, putative, expressed |
| LOC_Os06g02520 | interacting protein of DMI3, putative, expressed |
| LOC_Os06g02530 | expressed protein |
| LOC_Os06g02540 | expressed protein |
| LOC_Os06g02610 | folic acid binding protein, putative, expressed |
| LOC_Os06g02620 | chaperone protein dnaJ, putative, expressed |
| LOC_Os06g03090 | expressed protein |
| LOC_Os06g03486 | expressed protein |
| LOC_Os06g03520 | DUF581 domain containing protein, expressed |
| LOC_Os06g03530 | pentatricopeptide, putative, expressed |
| LOC_Os06g03570 | pentatricopeptide, putative, expressed |
| LOC_Os06g03600 | transcriptional corepressor SEUSS, putative, expressed |
| LOC_Os06g03620 | expressed protein |
| LOC_Os06g03640 | BAG domain containing protein, expressed |
| LOC_Os06g03650 | expressed protein |

| | |
|---|---|
| LOC_Os06g03690 | RNA recognition motif containing protein, putative, expressed |
| LOC_Os06g03720 | ribonucleoside-diphosphate reductase small chain, putative, expressed |
| LOC_Os06g03780 | NUC153 domain containing protein, expressed |
| LOC_Os06g03810 | expressed protein |
| LOC_Os06g03820 | expressed protein |
| LOC_Os06g03840 | H-BTB4 - Bric-a-Brac, Tramtrack, Broad Complex BTB domain with H family conserved sequence, expressed |
| LOC_Os06g03850 | impaired sucrose induction 1, putative, expressed |
| LOC_Os06g03910 | hydrolase, NUDIX family, domain containing protein, expressed |
| LOC_Os06g03940 | spastin, putative, expressed |
| LOC_Os06g03960 | expressed protein |
| LOC_Os06g03970 | receptor-like protein kinase 5 precursor, putative, expressed |
| LOC_Os06g03980 | expressed protein |
| LOC_Os06g04040 | WD domain, G-beta repeat domain containing protein, expressed |
| LOC_Os06g04070 | pyridoxal-dependent decarboxylase protein, putative, expressed |
| LOC_Os06g04190 | rad1, putative, expressed |
| LOC_Os06g04195 | expressed protein |
| LOC_Os06g04210 | expressed protein |
| LOC_Os06g04250 | phosphate-induced protein 1 conserved region domain containing protein, expressed |
| LOC_Os06g04370 | expressed protein |
| LOC_Os06g04380 | aminomethyltransferase, putative, expressed |
| LOC_Os06g04440 | expressed protein |
| LOC_Os06g04450 | Sec1 family transport protein, putative, expressed |
| LOC_Os06g04500 | cornichon protein, putative, expressed |
| LOC_Os06g04530 | OsClp9 - Putative Clp protease homologue, expressed |
| LOC_Os06g04540 | DNA binding protein, putative, expressed |
| LOC_Os06g04550 | expressed protein |
| LOC_Os06g04560 | kinesin motor domain containing protein, expressed |
| LOC_Os06g04600 | Brix domain containing protein, putative, expressed |
| LOC_Os06g04610 | ribosome-binding factor A, chloroplast precursor, putative, expressed |

| | |
|---|---|
| LOC_Os06g04620 | GDP-mannose 4,6 dehydratase 2, putative, expressed |
| LOC_Os06g04640 | bromodomain domain containing protein, expressed |
| LOC_Os06g04810 | leucine rich repeat protein, putative, expressed |
| LOC_Os06g04830 | leucine rich repeat protein, putative, expressed |
| LOC_Os06g04860 | hypothetical protein |
| LOC_Os06g04870 | homeobox associated leucine zipper, putative, expressed |
| LOC_Os06g04880 | serine threonine kinase, putative, expressed |
| LOC_Os06g04910 | hydrolase, NUDIX family protein, expressed |
| LOC_Os06g04920 | zinc finger family protein, putative, expressed |
| LOC_Os06g05080 | cytochrome c oxidase subunit 5B, mitochondrial precursor, putative, expressed |
| LOC_Os06g05090 | methyltransferase domain containing protein, expressed |
| LOC_Os06g05110 | superoxide dismutase, chloroplast, putative, expressed |
| LOC_Os06g05140 | pentatricopeptide, putative, expressed |
| LOC_Os06g05390 | expressed protein |
| LOC_Os06g05400 | IscA-like iron-sulfur assembly protein, mitochondrial precursor, putative, expressed |
| LOC_Os06g05520 | OsMKK1 - putative MAPKK based on amino acid sequence homology, expressed |
| LOC_Os06g05530 | RNA recognition motif containing protein, putative, expressed |
| LOC_Os06g05690 | cysteine synthase, chloroplast/chromoplast precursor, putative, expressed |
| LOC_Os06g05710 | pollen-specific protein, putative, expressed |
| LOC_Os06g05720 | expressed protein |
| LOC_Os06g05740 | expressed protein |
| LOC_Os06g05760 | ubiquitin family protein, putative, expressed |
| LOC_Os06g05770 | expressed protein |
| LOC_Os06g05804 | trafficking protein particle complex subunit 4, putative, expressed |
| LOC_Os06g05820 | OsLonP2 - Putative Lon protease homologue, expressed |
| LOC_Os06g05860 | 6-phosphofructokinase, putative, expressed |
| LOC_Os06g05870 | dual specificity protein phosphatase, putative, expressed |
| LOC_Os06g05900 | methyltransferase, putative, expressed |
| LOC_Os06g05920 | Rf1, mitochondrial precursor, putative, expressed |

| | |
|---|---|
| LOC_Os06g05930 | expressed protein |
| LOC_Os06g05940 | DTA2, putative, expressed |
| LOC_Os06g05960 | expressed protein |
| LOC_Os06g06080 | serine esterase family protein, putative, expressed |
| LOC_Os06g06090 | CGMC_MAPKCMGC_2_ERK 12 - CGMC includes CDA, MAPK, GSK3, and CLKC kinases, expressed |
| LOC_Os06g06115 | expressed protein |
| LOC_Os06g06120 | expressed protein |
| LOC_Os06g06130 | glutamate receptor, putative, expressed |
| LOC_Os06g06160 | IQ calmodulin-binding motif domain containing protein, expressed |
| LOC_Os06g06170 | expressed protein |
| LOC_Os06g06180 | transferase family protein, putative, expressed |
| LOC_Os06g06190 | uncharacterized protein ycf45, putative, expressed |
| LOC_Os06g06210 | expressed protein |
| LOC_Os06g06280 | galactosyltransferase family protein, putative, expressed |
| LOC_Os06g06350 | AMP-binding enzyme, putative, expressed |
| LOC_Os06g06430 | expressed protein |
| LOC_Os06g06530 | proline-rich cell wall protein-like, putative, expressed |
| LOC_Os06g06570 | expressed protein |
| LOC_Os06g06580 | expressed protein |
| LOC_Os06g06690 | hypothetical protein |
| LOC_Os06g06900 | helix-loop-helix DNA-binding domain containing protein, expressed |
| LOC_Os06g07350 | RNA-binding motif protein, putative, expressed |
| LOC_Os06g08120 | plant protein of unknown function domain containing protein, expressed |
| LOC_Os06g08290 | MYB family transcription factor, putative, expressed |
| LOC_Os06g08300 | FAD dependent oxidoreductase domain containing protein, expressed |
| LOC_Os06g08360 | ethylene-responsive element-binding protein, putative, expressed |
| LOC_Os06g08370 | hypothetical protein |
| LOC_Os06g08440 | two-component response regulator, putative, expressed |
| LOC_Os06g08470 | expressed protein |

| | |
|---|---|
| LOC_Os06g08480 | CHD3-type chromatin-remodeling factor PICKLE, putative, expressed |
| LOC_Os06g08490 | X0001, putative, expressed |
| LOC_Os06g08520 | transposon protein, putative, unclassified, expressed |
| LOC_Os06g08650 | pentatricopeptide, putative, expressed |
| LOC_Os06g08670 | glutathione peroxidase, putative, expressed |
| LOC_Os06g08690 | leucine-rich repeat receptor protein kinase EXS precursor, putative, expressed |
| LOC_Os06g08730 | expressed protein |
| LOC_Os06g08760 | conserved hypothetical protein |
| LOC_Os06g08870 | hypothetical protein |
| LOC_Os06g09717 | ethylene-responsive transcription factor, putative, expressed |
| LOC_Os06g11050 | cyclin-dependent kinase inhibitor, putative, expressed |
| LOC_Os06g11380 | geminivirus Rep-interacting motor protein, putative, expressed |
| LOC_Os06g11410 | cyclin, putative, expressed |
| LOC_Os06g11430 | GRAM domain containing protein, expressed |
| LOC_Os06g11620 | RNA recognition motif containing protein, putative, expressed |
| LOC_Os06g11630 | OsFBX192 - F-box domain containing protein, expressed |
| LOC_Os06g11740 | OsAPRL2 adenosine 5'-phosphosulfate reductase-like OsAPRL2, expressed |
| LOC_Os06g11780 | MYB family transcription factor, putative, expressed |
| LOC_Os06g11790 | TLD family protein, putative, expressed |
| LOC_Os06g11850 | expressed protein |
| LOC_Os06g11960 | 2,3-bisphosphoglycerate-dependent phosphoglycerate mutase, putative, expressed |
| LOC_Os06g11970 | OsMADS63 - MADS-box family gene with MIKC* type-box, expressed |
| LOC_Os06g11980 | DUF581 domain containing protein, expressed |
| LOC_Os06g12030 | glutaredoxin domain containing protein, putative, expressed |
| LOC_Os06g12040 | mTERF family protein, expressed |
| LOC_Os06g12110 | mTERF family protein, expressed |
| LOC_Os06g12129 | expressed protein |
| LOC_Os06g12140 | expressed protein |
| LOC_Os06g12150 | shikimate kinase, putative, expressed |

| | |
|---|---|
| LOC_Os06g12180 | jacalin-like lectin domain containing protein, expressed |
| LOC_Os06g12220 | HVA22, putative, expressed |
| LOC_Os06g12260 | N-rich protein, putative, expressed |
| LOC_Os06g12270 | hypothetical protein |
| LOC_Os06g12400 | Homeobox domain containing protein, expressed |
| LOC_Os06g12455 | expressed protein |
| LOC_Os06g12800 | expressed protein |
| LOC_Os06g12876 | expressed protein |
| LOC_Os06g14530 | expressed protein |
| LOC_Os06g15620 | GASR7 - Gibberellin-regulated GASA/GAST/Snakin family protein precursor, expressed |
| LOC_Os06g17470 | expressed protein |
| LOC_Os06g17830 | oxidoreductase, 2OG-Fe oxygenase family protein, putative, expressed |
| LOC_Os06g17840 | expressed protein |
| LOC_Os06g18840 | retrotransposon protein, putative, LINE subclass, expressed |
| LOC_Os06g19960 | aconitate hydratase protein, putative, expressed |
| LOC_Os06g20370 | microtubule associated protein, putative, expressed |
| LOC_Os06g20410 | BAH domain containing protein, expressed |
| LOC_Os06g20420 | expressed protein |
| LOC_Os06g20430 | BPI/LBP family protein At3g20270 precursor, putative, expressed |
| LOC_Os06g20550 | expressed protein |
| LOC_Os06g20820 | expressed protein |
| LOC_Os06g20860 | paramyosin, putative, expressed |
| LOC_Os06g20870 | pentatricopeptide repeat protein PPR1106-17, putative, expressed |
| LOC_Os06g20900 | expressed protein |
| LOC_Os06g20920 | SAM dependent carboxyl methyltransferase, putative, expressed |
| LOC_Os06g20960 | SAM dependent carboxyl methyltransferase, putative, expressed |
| LOC_Os06g21110 | expressed protein |
| LOC_Os06g22300 | transposon protein, putative, Ac/Ds sub-class, expressed |
| LOC_Os06g22670 | tesmin/TSO1-like CXC domain containing protein, expressed |

| | |
|---|---|
| LOC_Os06g23780 | glutaryl-CoA dehydrogenase, mitochondrial precursor, putative, expressed |
| LOC_Os06g24020 | retrotransposon protein, putative, unclassified |
| LOC_Os06g24850 | OsIAA22 - Auxin-responsive Aux/IAA gene family member, expressed |
| LOC_Os06g24900 | hypothetical protein |
| LOC_Os06g25590 | expressed protein |
| LOC_Os06g25660 | transposon protein, putative, unclassified, expressed |
| LOC_Os06g25670 | expressed protein |
| LOC_Os06g27840 | retrotransposon protein, putative, LINE subclass, expressed |
| LOC_Os06g28740 | expressed protein |
| LOC_Os06g31210 | methionyl-tRNA synthetase, putative, expressed |
| LOC_Os06g31270 | polygalacturonase, putative, expressed |
| LOC_Os06g31810 | hypothetical protein |
| LOC_Os06g33220 | expressed protein |
| LOC_Os06g33230 | transposon protein, putative, Mariner sub-class, expressed |
| LOC_Os06g33320 | extra-large G-protein-related, putative, expressed |
| LOC_Os06g33710 | spermidine synthase, putative, expressed |
| LOC_Os06g33970 | VQ domain containing protein, putative, expressed |
| LOC_Os06g34450 | zinc finger, C3HC4 type domain containing protein, expressed |
| LOC_Os06g34570 | retrotransposon protein, putative, unclassified, expressed |
| LOC_Os06g34700 | expressed protein |
| LOC_Os06g34710 | expressed protein |
| LOC_Os06g35400 | hypothetical protein |
| LOC_Os06g35560 | reticuline oxidase-like protein precursor, putative, expressed |
| LOC_Os06g35830 | expressed protein |
| LOC_Os06g36010 | plastocyanin-like domain containing protein, putative, expressed |
| LOC_Os06g36050 | expressed protein |
| LOC_Os06g36090 | ABC-2 type transporter, putative, expressed |
| LOC_Os06g36220 | OsFBX198 - F-box domain containing protein, expressed |
| LOC_Os06g36230 | expressed protein |

| | |
|---|---|
| LOC_Os06g36490 | ethylene-responsive element-binding protein, putative, expressed |
| LOC_Os06g36590 | transporter, monovalent cation:proton antiporter-2 family, putative, expressed |
| LOC_Os06g36810 | expressed protein |
| LOC_Os06g36970 | transposon protein, putative, unclassified, expressed |
| LOC_Os06g36990 | phospholipid-transporting ATPase 4, putative, expressed |
| LOC_Os06g37000 | RNA recognition motif containing protein, expressed |
| LOC_Os06g39400 | retrotransposon protein, putative, unclassified, expressed |
| LOC_Os06g39515 | expressed protein |
| LOC_Os06g39610 | expressed protein |
| LOC_Os06g39660 | FAD binding domain containing protein, expressed |
| LOC_Os06g39748 | expressed protein |
| LOC_Os06g39900 | agenet domain containing protein, putative, expressed |
| LOC_Os06g39906 | homeobox domain containing protein, expressed |
| LOC_Os06g39970 | CESA11 - cellulose synthase, expressed |
| LOC_Os06g40020 | DEAD-box ATP-dependent RNA helicase 52A, putative, expressed |
| LOC_Os06g40050 | mitochondrial carrier protein, putative, expressed |
| LOC_Os06g40460 | expressed protein |
| LOC_Os06g40600 | elongation factor, putative, expressed |
| LOC_Os06g40670 | expressed protein |
| LOC_Os06g40680 | expressed protein |
| LOC_Os06g40740 | expressed protein |
| LOC_Os06g41030 | DUF1680 domain containing protein, putative, expressed |
| LOC_Os06g42790 | hydrolase, NUDIX family, domain containing protein, expressed |
| LOC_Os06g42890 | DNA binding protein, putative, expressed |
| LOC_Os06g43690 | WD repeat-containing protein, putative, expressed |
| LOC_Os06g43700 | transmembrane amino acid transporter protein, putative, expressed |
| LOC_Os06g43870 | expressed protein |
| LOC_Os06g43890 | expressed protein |
| LOC_Os06g43900 | ribosomal protein L35, putative, expressed |

| | |
|---|---|
| LOC_Os06g43990 | expressed protein |
| LOC_Os06g44140 | transmembrane 9 superfamily member, putative, expressed |
| LOC_Os06g44160 | heat shock protein DnaJ, putative, expressed |
| LOC_Os06g44830 | expressed protein |
| LOC_Os06g45440 | haloacid dehalogenase-like hydrolase domain-containing protein 3, putative, expressed |
| LOC_Os06g45470 | expressed protein |
| LOC_Os06g45480 | hydrolase, putative, expressed |
| LOC_Os06g45860 | molybdenum cofactor sulfurase, putative, expressed |
| LOC_Os06g48680 | expressed protein |
| LOC_Os06g49505 | arogenate dehydrogenase 1, chloroplast precursor, putative, expressed |
| LOC_Os06g49510 | zinc knuckle family protein, putative, expressed |
| LOC_Os06g49620 | expressed protein |
| LOC_Os06g49640 | uncharacterized UPF0114 domain containing protein, expressed |
| LOC_Os06g49650 | harpin-induced protein 1 domain containing protein, expressed |
| LOC_Os06g49770 | LTPL86 - Protease inhibitor/seed storage/LTP family protein precursor, expressed |
| LOC_Os06g49840 | OsMADS16 - MADS-box family gene with MIKCc type-box, expressed |
| LOC_Os06g50300 | heat shock protein, putative, expressed |
| LOC_Os06g50370 | zinc finger, C3HC4 type, domain containing protein, expressed |
| LOC_Os06g50380 | protein phosphatase 2C, putative, expressed |
| LOC_Os06g50430 | expressed protein |
| LOC_Os06g50890 | RNA recognition motif containing protein, expressed |
| LOC_Os06g50910 | Phosphatidylinositol kinase and FAT containing domain protein, putative, expressed |
| LOC_Os06g51084 | 1,4-alpha-glucan-branching enzyme, chloroplast precursor, putative, expressed |
| LOC_Os06g51180 | lysine ketoglutarate reductase trans-splicing related 1, putative, expressed |
| LOC_Os06g51210 | zinc finger family protein, putative, expressed |
| LOC_Os06g51280 | acetolactate synthase 1, chloroplast precursor, putative, expressed |
| LOC_Os06g51290 | phytoene synthase, chloroplast precursor, putative, expressed |
| LOC_Os06g51320 | GASR8 - Gibberellin-regulated GASA/GAST/Snakin family protein precursor, expressed |
| LOC_Os06g51380 | ROOT HAIRLESS 1, putative, expressed |

| | |
|---|---|
| LOC_Os06g51390 | expressed protein |
| LOC_Os06g51400 | expressed protein |
| LOC_Os06g51520 | lysine ketoglutarate reductase trans-splicing related 1, putative, expressed |
| LOC_Os07g01060 | expressed protein |
| LOC_Os07g01120 | expressed protein |
| LOC_Os07g01130 | pentatricopeptide containing protein, putative, expressed |
| LOC_Os07g01160 | expressed protein |
| LOC_Os07g02070 | retrotransposon protein, putative, Ty3-gypsy subclass |
| LOC_Os07g02490 | retrotransposon protein, putative, unclassified, expressed |
| LOC_Os07g02760 | OsFBD13 - F-box and FBD domain containing protein, expressed |
| LOC_Os07g03120 | expressed protein |
| LOC_Os07g03160 | WD domain, G-beta repeat domain containing protein, expressed |
| LOC_Os07g03210 | expressed protein |
| LOC_Os07g03220 | bZIP transcription factor domain containing protein, expressed |
| LOC_Os07g03260 | CSLC10 - cellulose synthase-like family C, expressed |
| LOC_Os07g04020 | expressed protein |
| LOC_Os07g04150 | stress-induced protein, putative, expressed |
| LOC_Os07g04170 | OsMADS90 - MADS-box family gene with M-beta type-box, expressed |
| LOC_Os07g04200 | bacterial transferase hexapeptide domain containing protein, expressed |
| LOC_Os07g04210 | Ser/Thr protein phosphatase family protein, putative, expressed |
| LOC_Os07g04220 | wound and phytochrome signaling involved receptor like kinase, putative, expressed |
| LOC_Os07g04230 | mTERF domain containing protein, expressed |
| LOC_Os07g04530 | GYF domain containing protein, putative, expressed |
| LOC_Os07g04540 | expressed protein |
| LOC_Os07g04550 | pleckstrin homology domain-containing protein, putative, expressed |
| LOC_Os07g04570 | RNA recognition motif containing protein, putative, expressed |
| LOC_Os07g04580 | zinc finger C-x8-C-x5-C-x3-H type family protein, expressed |
| LOC_Os07g04780 | expressed protein |
| LOC_Os07g04790 | OsFBX215 - F-box domain containing protein, expressed |

| | |
|---|---|
| LOC_Os07g04800 | hypothetical protein |
| LOC_Os07g04840 | PsbP, putative, expressed |
| LOC_Os07g04850 | pre-mRNA-processing factor 31, putative, expressed |
| LOC_Os07g04880 | retrotransposon protein, putative, unclassified |
| LOC_Os07g04910 | reticulon domain containing protein, putative, expressed |
| LOC_Os07g04930 | retrotransposon protein, putative, unclassified, expressed |
| LOC_Os07g04980 | expressed protein |
| LOC_Os07g04990 | oxidoreductase, aldo/keto reductase family protein, putative, expressed |
| LOC_Os07g05024 | expressed protein |
| LOC_Os07g05030 | expressed protein |
| LOC_Os07g05050 | DEAD-box ATP-dependent RNA helicase, putative, expressed |
| LOC_Os07g05060 | expressed protein |
| LOC_Os07g05084 | expressed protein |
| LOC_Os07g05100 | expressed protein |
| LOC_Os07g05120 | expressed protein |
| LOC_Os07g05145 | expressed protein |
| LOC_Os07g05150 | Skp1 family, dimerisation domain containing protein, expressed |
| LOC_Os07g05160 | SKP1-like protein 1B, putative, expressed |
| LOC_Os07g05190 | leucine-rich repeat family protein, putative, expressed |
| LOC_Os07g05360 | photosystem II 10 kDa polypeptide, chloroplast precursor, putative, expressed |
| LOC_Os07g05370 | TKL_IRAK_CrRLK1L-1 |
| LOC_Os07g05450 | sulfotransferase domain containing protein, expressed |
| LOC_Os07g05470 | AIR9 protein, putative, expressed |
| LOC_Os07g05560 | expressed protein |
| LOC_Os07g05570 | ERD4 protein, putative, expressed |
| LOC_Os07g05580 | ribosomal protein L7Ae, putative, expressed |
| LOC_Os07g05600 | retrotransposon protein, putative, unclassified, expressed |
| LOC_Os07g05650 | MAGE-8 antigen-related, putative, expressed |
| LOC_Os07g05670 | expressed protein |

| | |
|---|---|
| LOC_Os07g05930 | expressed protein |
| LOC_Os07g07440 | versicolorin reductase, putative, expressed |
| LOC_Os07g07460 | hypothetical protein |
| LOC_Os07g07470 | transketolase, putative, expressed |
| LOC_Os07g07480 | HNH endonuclease domain-containing protein, putative, expressed |
| LOC_Os07g07540 | SHOOT1 protein, putative, expressed |
| LOC_Os07g07550 | galactose-1-phosphate uridyl transferase, putative, expressed |
| LOC_Os07g07590 | expressed protein |
| LOC_Os07g07610 | RNA polymerase subunit, putative, expressed |
| LOC_Os07g07620 | pentatricopeptide, putative, expressed |
| LOC_Os07g07630 | expressed protein |
| LOC_Os07g07640 | hypothetical protein |
| LOC_Os07g07690 | PHD-finger domain containing protein, expressed |
| LOC_Os07g07800 | PHA synthase, putative, expressed |
| LOC_Os07g07820 | expressed protein |
| LOC_Os07g07880 | trp repressor/replication initiator, putative, expressed |
| LOC_Os07g07890 | expressed protein |
| LOC_Os07g07900 | expressed protein |
| LOC_Os07g07920 | LTPL100 - Protease inhibitor/seed storage/LTP family protein precursor, expressed |
| LOC_Os07g07930 | LTPL78 - Protease inhibitor/seed storage/LTP family protein precursor, expressed |
| LOC_Os07g08040 | expressed protein |
| LOC_Os07g08750 | STE_PAK_Ste20_Slob_Wnk |
| LOC_Os07g11980 | expressed protein |
| LOC_Os07g12240 | EF hand family protein, putative, expressed |
| LOC_Os07g12530 | ABC1 family domain containing protein, putative, expressed |
| LOC_Os07g12630 | transcription elongation factor protein, putative, expressed |
| LOC_Os07g12640 | glycerol-3-phosphate dehydrogenase, putative, expressed |
| LOC_Os07g12660 | expressed protein |
| LOC_Os07g12670 | expressed protein |

| | |
|---|---|
| LOC_Os07g12700 | expressed protein |
| LOC_Os07g12710 | GIY-YIG catalytic domain containing protein, putative, expressed |
| LOC_Os07g12790 | expressed protein |
| LOC_Os07g12800 | expressed protein |
| LOC_Os07g12810 | expressed protein |
| LOC_Os07g12890 | metal cation transporter, putative, expressed |
| LOC_Os07g13040 | expressed protein |
| LOC_Os07g14350 | methyltransferase, putative, expressed |
| LOC_Os07g14514 | retrotransposon protein, putative, unclassified, expressed |
| LOC_Os07g14540 | vesicle-associated membrane protein, putative, expressed |
| LOC_Os07g14580 | mRNA-decapping enzyme, putative, expressed |
| LOC_Os07g14590 | hydrolase, putative, expressed |
| LOC_Os07g15110 | retrotransposon protein, putative, unclassified, expressed |
| LOC_Os07g17470 | hypothetical protein |
| LOC_Os07g18110 | expressed protein |
| LOC_Os07g18120 | aldehyde oxidase, putative, expressed |
| LOC_Os07g18170 | retrotransposon protein, putative, unclassified |
| LOC_Os07g18250 | expressed protein |
| LOC_Os07g18510 | OsFBLD5 - F-box, LRR and FBD domain containing protein, expressed |
| LOC_Os07g19320 | stripe rust resistance protein Yr10, putative, expressed |
| LOC_Os07g22340 | hypothetical protein |
| LOC_Os07g22350 | glucose-6-phosphate 1-dehydrogenase, chloroplast precursor, putative, expressed |
| LOC_Os07g22730 | AP2 domain containing protein, expressed |
| LOC_Os07g22930 | starch synthase, putative, expressed |
| LOC_Os07g23100 | expressed protein |
| LOC_Os07g23130 | expressed protein |
| LOC_Os07g23140 | hypothetical protein |
| LOC_Os07g23150 | transferase family protein, putative, expressed |
| LOC_Os07g23190 | lysine ketoglutarate reductase trans-splicing related 1, putative, expressed |

| | |
|---|---|
| LOC_Os07g23200 | F-actin-capping protein subunit alpha, putative, expressed |
| LOC_Os07g25420 | 26S proteasome non-ATPase regulatory subunit 8, putative, expressed |
| LOC_Os07g25450 | histone-lysine N-methyltransferase, putative, expressed |
| LOC_Os07g25490 | expressed protein |
| LOC_Os07g25510 | expressed protein |
| LOC_Os07g25520 | expressed protein |
| LOC_Os07g25530 | expressed protein |
| LOC_Os07g25670 | expressed protein |
| LOC_Os07g25680 | protein kinase domain containing protein, expressed |
| LOC_Os07g25710 | myb-like DNA-binding domain containing protein, expressed |
| LOC_Os07g25800 | OsFBDUF37 - F-box and DUF domain containing protein, expressed |
| LOC_Os07g26440 | expressed protein |
| LOC_Os07g26460 | GLTP domain containing protein, putative, expressed |
| LOC_Os07g26490 | ankyrin repeat domain containing protein, expressed |
| LOC_Os07g26570 | expressed protein |
| LOC_Os07g26610 | phosphoglucomutase, putative, expressed |
| LOC_Os07g26740 | 60S ribosomal protein L44, putative, expressed |
| LOC_Os07g27880 | O-methyltransferase, putative, expressed |
| LOC_Os07g30020 | anthranilate phosphoribosyltransferase, putative, expressed |
| LOC_Os07g30160 | trehalose phosphatase, putative, expressed |
| LOC_Os07g30250 | RFT1, putative, expressed |
| LOC_Os07g30280 | csAtPR5, putative, expressed |
| LOC_Os07g30790 | ubiquinol-cytochrome C chaperone family protein, putative, expressed |
| LOC_Os07g30980 | uvrD/REP helicase family protein, putative, expressed |
| LOC_Os07g30990 | ubiquitin family protein, putative, expressed |
| LOC_Os07g31070 | expressed protein |
| LOC_Os07g31340 | regulator of nonsense transcripts 1, putative, expressed |
| LOC_Os07g31350 | expressed protein |
| LOC_Os07g31450 | CHR4/MI-2-LIKE, putative, expressed |

| | |
|---|---|
| LOC_Os07g31460 | peptide-Nasparagine amidase, putative, expressed |
| LOC_Os07g31770 | chalcone synthase, putative, expressed |
| LOC_Os07g31930 | hypothetical protein |
| LOC_Os07g32380 | protein phosphatase 2C, putative, expressed |
| LOC_Os07g32460 | src homology-3 domain protein 3, putative, expressed |
| LOC_Os07g33330 | RNA recognition motif containing protein, putative, expressed |
| LOC_Os07g34510 | retrotransposon protein, putative, unclassified, expressed |
| LOC_Os07g34520 | isocitrate lyase, putative, expressed |
| LOC_Os07g34560 | expressed protein |
| LOC_Os07g34570 | FAD dependent oxidoreductase domain containing protein, expressed |
| LOC_Os07g34690 | expressed protein |
| LOC_Os07g34710 | peroxidase precursor, putative, expressed |
| LOC_Os07g35150 | hypothetical protein |
| LOC_Os07g35580 | TKL_IRAK_DUF26-lc 20 - DUF26 kinases have homology to DUF26 containing loci, expressed |
| LOC_Os07g35870 | basic helix-loop-helix DND-binding domain containing protein, expressed |
| LOC_Os07g36170 | chitin-inducible gibberellin-responsive protein, putative, expressed |
| LOC_Os07g36180 | pentatricopeptide, putative, expressed |
| LOC_Os07g36190 | hydrolase, NUDIX family, domain containing protein, expressed |
| LOC_Os07g36200 | eukaryotic translation initiation factor-related, putative, expressed |
| LOC_Os07g36390 | PPR repeat containing protein, expressed |
| LOC_Os07g36410 | expressed protein |
| LOC_Os07g36590 | serine/threonine-protein kinase receptor precursor, putative, expressed |
| LOC_Os07g36630 | CSLF8 - cellulose synthase-like family F; beta1,3;1,4 glucan synthase, expressed |
| LOC_Os07g36830 | OsFBX250 - F-box domain containing protein, expressed |
| LOC_Os07g37890 | protein phosphatase 2C, putative, expressed |
| LOC_Os07g40070 | expressed protein |
| LOC_Os07g42250 | strictosidine synthase, putative, expressed |
| LOC_Os07g42580 | expressed protein |
| LOC_Os07g42626 | STRUBBELIG-RECEPTOR FAMILY 6 precursor, putative, expressed |

| | |
|---|---|
| LOC_Os07g43660 | expressed protein |
| LOC_Os07g45750 | retrotransposon protein, putative, unclassified, expressed |
| LOC_Os07g46220 | SNARE associated Golgi protein, putative, expressed |
| LOC_Os07g46350 | OsSCP40 - Putative Serine Carboxypeptidase homologue, expressed |
| LOC_Os07g46830 | sex determination protein tasselseed-2, putative, expressed |
| LOC_Os07g48750 | alpha-N-arabinofuranosidase, putative, expressed |
| LOC_Os07g49020 | expressed protein |
| LOC_Os07g49030 | PHD-finger family protein, expressed |
| LOC_Os07g49050 | expressed protein |
| LOC_Os07g49090 | WD-40 repeat family protein, putative, expressed |
| LOC_Os08g01040 | zinc finger, C3HC4 type domain containing protein, expressed |
| LOC_Os08g01090 | B3 DNA binding domain containing protein, expressed |
| LOC_Os08g01100 | HMG1/2, putative, expressed |
| LOC_Os08g01170 | acetyltransferase, GNAT family, putative, expressed |
| LOC_Os08g01210 | harpin-induced protein 1 domain containing protein, expressed |
| LOC_Os08g01230 | hypothetical protein |
| LOC_Os08g01240 | expressed protein |
| LOC_Os08g01270 | protein kinase family protein, putative, expressed |
| LOC_Os08g01274 | expressed protein |
| LOC_Os08g01290 | OsFBT12 - F-box and tubby domain containing protein, expressed |
| LOC_Os08g01310 | phospholipase C, putative, expressed |
| LOC_Os08g01320 | C3-BTB1 - Bric-a-Brac, Tramtrack, Broad Complex BTB domain with C3 subfamily conserved sequence, expressed |
| LOC_Os08g01330 | no apical meristem protein, putative, expressed |
| LOC_Os08g01350 | transmembrane protein 18, putative, expressed |
| LOC_Os08g01360 | RING finger protein 13, putative, expressed |
| LOC_Os08g01370 | expressed protein |
| LOC_Os08g01380 | 2Fe-2S iron-sulfur cluster binding domain containing protein, expressed |
| LOC_Os08g01390 | phosphatidylinositol-4-phosphate 5-Kinase, putative, expressed |
| LOC_Os08g01430 | OsFBX260 - F-box domain containing protein, expressed |

| | |
|---|---|
| LOC_Os08g01460 | expressed protein |
| LOC_Os08g01550 | expressed protein |
| LOC_Os08g01600 | polygalacturonase, putative, expressed |
| LOC_Os08g01620 | shwachman-Bodian-Diamond syndrome protein, putative, expressed |
| LOC_Os08g01630 | DNA repair metallo-beta-lactamase, putative, expressed |
| LOC_Os08g01660 | mago nashi, putative, expressed |
| LOC_Os08g01680 | WD domain, G-beta repeat domain containing protein, expressed |
| LOC_Os08g01700 | basic helix-loop-helix family protein, putative, expressed |
| LOC_Os08g01710 | GLTP domain containing protein, putative, expressed |
| LOC_Os08g01740 | VHS and GAT domain containing protein, expressed |
| LOC_Os08g01760 | dehydrogenase, putative, expressed |
| LOC_Os08g01770 | adenylate kinase, putative, expressed |
| LOC_Os08g01900 | armadillo/beta-catenin repeat family protein, putative, expressed |
| LOC_Os08g01910 | expressed protein |
| LOC_Os08g01930 | KH domain-containing protein, putative, expressed |
| LOC_Os08g02340 | 60S acidic ribosomal protein, putative, expressed |
| LOC_Os08g02350 | expressed protein |
| LOC_Os08g02440 | expressed protein |
| LOC_Os08g02530 | OsSAUR32 - Auxin-responsive SAUR gene family member, expressed |
| LOC_Os08g02600 | methyltransferase domain containing protein, expressed |
| LOC_Os08g02630 | photosystem II core complex proteins psbY, chloroplast precursor, putative, expressed |
| LOC_Os08g02640 | succinate dehydrogenase iron-sulfur subunit 2,mitochondrial precursor, putative, expressed |
| LOC_Os08g02850 | zinc finger family protein, putative, expressed |
| LOC_Os08g02880 | CXXXC11 - Cysteine-rich protein with paired CXXXC motifs precursor, expressed |
| LOC_Os08g02909 | expressed protein |
| LOC_Os08g02950 | expressed protein |
| LOC_Os08g02960 | expressed protein |
| LOC_Os08g02970 | expressed protein |
| LOC_Os08g02990 | retrotransposon protein, putative, unclassified, expressed |

| | |
|---|---|
| LOC_Os08g03360 | ATBET9, putative, expressed |
| LOC_Os08g03420 | kelch repeat protein, putative, expressed |
| LOC_Os08g03430 | extracellular ligand-gated ion channel, putative, expressed |
| LOC_Os08g03550 | expressed protein |
| LOC_Os08g03560 | chloroplast unusual positioning protein, putative, expressed |
| LOC_Os08g03600 | expressed protein |
| LOC_Os08g03670 | galactosyltransferase family protein, putative, expressed |
| LOC_Os08g03676 | pentatricopeptide, putative, expressed |
| LOC_Os08g03690 | LTPL24 - Protease inhibitor/seed storage/LTP family protein precursor, expressed |
| LOC_Os08g04090 | expressed protein |
| LOC_Os08g04110 | transporter, putative, expressed |
| LOC_Os08g04120 | expressed protein |
| LOC_Os08g04130 | copine-6, putative, expressed |
| LOC_Os08g04160 | expressed protein |
| LOC_Os08g04170 | zinc finger C-x8-C-x5-C-x3-H type family protein, expressed |
| LOC_Os08g04220 | expressed protein |
| LOC_Os08g04310 | plastocyanin-like domain containing protein, putative, expressed |
| LOC_Os08g04340 | plastocyanin-like domain containing protein, putative, expressed |
| LOC_Os08g04390 | basic helix-loop-helix DND-binding domain containing protein, expressed |
| LOC_Os08g04400 | pentatricopeptide repeat domain containing protein, putative, expressed |
| LOC_Os08g04580 | csAtPR5, putative, expressed |
| LOC_Os08g04630 | external NADH-ubiquinone oxidoreductase 1, mitochondrial precursor, putative, expressed |
| LOC_Os08g04770 | AMP-binding domain containing protein, expressed |
| LOC_Os08g04780 | amine oxidase, putative, expressed |
| LOC_Os08g04800 | triacylglycerol lipase like protein, putative, expressed |
| LOC_Os08g04900 | hypothetical protein |
| LOC_Os08g04910 | expressed protein |
| LOC_Os08g04920 | OsDegp11 - Putative Deg protease homologue, expressed |
| LOC_Os08g05440 | NB-ARC domain containing protein, expressed |

| | |
|---|---|
| LOC_Os08g06490 | pentatricopeptide, putative, expressed |
| LOC_Os08g06520 | expressed protein |
| LOC_Os08g06530 | rubredoxin family protein, putative, expressed |
| LOC_Os08g06590 | expressed protein |
| LOC_Os08g07030 | hypothetical protein |
| LOC_Os08g07060 | CRR6, putative, expressed |
| LOC_Os08g07760 | BRASSINOSTEROID INSENSITIVE 1-associated receptor kinase 1 precursor, putative, expressed |
| LOC_Os08g07774 | disease resistance protein RPM1, putative, expressed |
| LOC_Os08g07840 | PolI-like DNA polymerase, putative, expressed |
| LOC_Os08g07880 | phosphopantothenate--cysteine ligase, putative, expressed |
| LOC_Os08g07950 | NB-ARC domain containing protein, expressed |
| LOC_Os08g08000 | DNA binding protein, putative, expressed |
| LOC_Os08g08020 | expressed protein |
| LOC_Os08g08060 | vacuolar protein sorting-associated protein 18, putative, expressed |
| LOC_Os08g08080 | BUD13, putative, expressed |
| LOC_Os08g08084 | expressed protein |
| LOC_Os08g08090 | wound induced protein, putative, expressed |
| LOC_Os08g08120 | B-box zinc finger family protein, putative, expressed |
| LOC_Os08g08205 | expressed protein |
| LOC_Os08g08210 | SET domain containing protein, expressed |
| LOC_Os08g08250 | expressed protein |
| LOC_Os08g08830 | expressed protein |
| LOC_Os08g08840 | phosphate/phosphate translocator, putative, expressed |
| LOC_Os08g09110 | NB-ARC domain containing protein, expressed |
| LOC_Os08g09120 | expressed protein |
| LOC_Os08g09130 | conserved hypothetical protein |
| LOC_Os08g09170 | expressed protein |
| LOC_Os08g09180 | postsynaptic protein CRIPT, putative, expressed |
| LOC_Os08g09210 | phosphoribosylamine--glycine ligase, putative, expressed |

| | |
|---|---|
| LOC_Os08g09270 | pentatricopeptide, putative, expressed |
| LOC_Os08g09280 | hypothetical protein |
| LOC_Os08g09340 | bromodomain containing protein, expressed |
| LOC_Os08g09350 | gar2, putative, expressed |
| LOC_Os08g09690 | nuclear transcription factor Y subunit, putative, expressed |
| LOC_Os08g09940 | GTP-binding protein, putative, expressed |
| LOC_Os08g10000 | expressed protein |
| LOC_Os08g10030 | KH domain-containing protein, putative, expressed |
| LOC_Os08g10070 | protein kinase, putative, expressed |
| LOC_Os08g12160 | expressed protein |
| LOC_Os08g12650 | retrotransposon protein, putative, Ty3-gypsy subclass, expressed |
| LOC_Os08g12670 | expressed protein |
| LOC_Os08g12740 | disease resistance protein RGA3, putative, expressed |
| LOC_Os08g12750 | serine/threonine-protein kinase HT1, putative, expressed |
| LOC_Os08g12760 | YT521-B, putative, expressed |
| LOC_Os08g12780 | chloroplast envelope membrane protein, putative, expressed |
| LOC_Os08g12890 | DNA binding protein, putative, expressed |
| LOC_Os08g12990 | retrotransposon protein, putative, unclassified, expressed |
| LOC_Os08g13060 | MBTB22 - Bric-a-Brac, Tramtrack, Broad Complex BTB domain with Meprin and TRAF Homology MATH domain, expressed |
| LOC_Os08g13270 | expressed protein |
| LOC_Os08g13310 | expressed protein |
| LOC_Os08g13320 | expressed protein |
| LOC_Os08g13990 | pentatricopeptide, putative, expressed |
| LOC_Os08g14195 | expressed protein |
| LOC_Os08g14420 | expressed protein |
| LOC_Os08g14440 | uridylyltransferase-related, putative, expressed |
| LOC_Os08g14450 | RNA polymerase sigma factor, putative, expressed |
| LOC_Os08g14630 | hypothetical protein |
| LOC_Os08g15080 | proline rich protein 3, putative, expressed |

| | |
|---|---|
| LOC_Os08g15230 | heat shock protein-related, putative, expressed |
| LOC_Os08g15410 | expressed protein |
| LOC_Os08g15480 | conserved hypothetical protein |
| LOC_Os08g15505 | expressed protein |
| LOC_Os08g15545 | DEFL55 - Defensin and Defensin-like DEFL family, expressed |
| LOC_Os08g17784 | purple acid phosphatase, putative, expressed |
| LOC_Os08g17820 | retrotransposon protein, putative, unclassified, expressed |
| LOC_Os08g18000 | OsFBX283 - F-box domain containing protein, expressed |
| LOC_Os08g18110 | alpha-soluble NSF attachment protein, putative, expressed |
| LOC_Os08g18910 | expressed protein |
| LOC_Os08g18970 | pectate lyase family protein, expressed |
| LOC_Os08g19140 | adenylate kinase family protein, putative, expressed |
| LOC_Os08g19170 | G-patch domain containing protein, expressed |
| LOC_Os08g19320 | aberrant root formation protein 4, putative, expressed |
| LOC_Os08g19330 | expressed protein |
| LOC_Os08g19420 | O-methyltransferase, putative, expressed |
| LOC_Os08g19520 | retrotransposon protein, putative, unclassified, expressed |
| LOC_Os08g20200 | male sterility protein, putative, expressed |
| LOC_Os08g22149 | CBS domain containing membrane protein, putative, expressed |
| LOC_Os08g23150 | indole-3-glycerol phosphate synthase, chloroplast precursor, putative, expressed |
| LOC_Os08g23720 | glucan endo-1,3-beta-glucosidase precursor, putative, expressed |
| LOC_Os08g24400 | SWP, putative, expressed |
| LOC_Os08g24460 | hypothetical protein |
| LOC_Os08g24470 | expressed protein |
| LOC_Os08g24480 | HAT family dimerisation domain containing protein, identical, putative, expressed |
| LOC_Os08g27400 | transposon protein, putative, CACTA, En/Spm sub-class |
| LOC_Os08g28730 | NAD dependent epimerase/dehydratase family protein, putative, expressed |
| LOC_Os08g28880 | patatin, putative, expressed |
| LOC_Os08g28920 | hypothetical protein |

| | |
|---|---|
| LOC_Os08g28970 | expressed protein |
| LOC_Os08g29110 | thioredoxin, putative, expressed |
| LOC_Os08g29124 | expressed protein |
| LOC_Os08g29220 | expressed protein |
| LOC_Os08g29400 | OsFBX290 - F-box domain containing protein, expressed |
| LOC_Os08g29854 | RGH1A, putative, expressed |
| LOC_Os08g30020 | membrane protein, putative, expressed |
| LOC_Os08g30070 | expressed protein |
| LOC_Os08g30634 | DC1 domain containing protein, expressed |
| LOC_Os08g30780 | ABC transporter, ATP-binding protein, putative, expressed |
| LOC_Os08g30790 | hypersensitive-induced response protein, putative, expressed |
| LOC_Os08g30810 | puromycin-sensitive aminopeptidase, putative, expressed |
| LOC_Os08g31040 | expressed protein |
| LOC_Os08g31070 | expressed protein |
| LOC_Os08g31080 | DUF260 domain containing protein, putative, expressed |
| LOC_Os08g33590 | basic helix-loop-helix, putative, expressed |
| LOC_Os08g33750 | myb-like DNA-binding domain containing protein, expressed |
| LOC_Os08g33790 | hypothetical protein |
| LOC_Os08g33920 | 60S ribosomal protein L34, putative, expressed |
| LOC_Os08g34010 | ZF-HD protein dimerisation region containing protein, expressed |
| LOC_Os08g34050 | phosphofructokinase, putative, expressed |
| LOC_Os08g34060 | DUF1336 domain containing protein, expressed |
| LOC_Os08g34130 | mitochondrial glycoprotein, putative, expressed |
| LOC_Os08g34249 | inhibitor I family protein, putative, expressed |
| LOC_Os08g34410 | expressed protein |
| LOC_Os08g34500 | expressed protein |
| LOC_Os08g35490 | ctr copper transporter family protein, putative, expressed |
| LOC_Os08g35650 | expressed protein |
| LOC_Os08g36150 | activator of 90 kDa heat shock protein ATPase homolog, putative, expressed |

| | |
|---|---|
| LOC_Os08g36450 | transcription regulator, putative, expressed |
| LOC_Os08g36460 | expressed protein |
| LOC_Os08g36570 | F-box protein, putative, expressed |
| LOC_Os08g36680 | bifunctional monodehydroascorbate reductase and carbonic anhydrasenectarin-3 precursor, putative, expressed |
| LOC_Os08g36994 | protein transport protein, putative, expressed |
| LOC_Os08g37090 | expressed protein |
| LOC_Os08g37104 | expressed protein |
| LOC_Os08g37270 | heat shock protein DnaJ, putative, expressed |
| LOC_Os08g37280 | PWWP domain containing protein, expressed |
| LOC_Os08g37350 | ubiquitin carboxyl-terminal hydrolase family protein, expressed |
| LOC_Os08g37390 | cyclin, putative, expressed |
| LOC_Os08g37490 | 14-3-3 protein, putative, expressed |
| LOC_Os08g37605 | riboflavin biosynthesis protein ribAB, chloroplast precursor, putative, expressed |
| LOC_Os08g37760 | zinc finger, C3HC4 type domain containing protein, expressed |
| LOC_Os08g37810 | transcription factor like protein, putative, expressed |
| LOC_Os08g38020 | bZIP transcription factor domain containing protein, expressed |
| LOC_Os08g38430 | retrotransposon protein, putative, unclassified, expressed |
| LOC_Os08g39174 | retrotransposon protein, putative, Ty1-copia subclass, expressed |
| LOC_Os08g39550 | polygalacturonase inhibitor 2 precursor, putative, expressed |
| LOC_Os08g40750 | expressed protein |
| LOC_Os08g40790 | DNA repair ATPase-related, putative, expressed |
| LOC_Os08g41290 | AIR12, putative, expressed |
| LOC_Os08g41300 | 60S ribosomal protein L32, putative, expressed |
| LOC_Os08g41310 | hypothetical protein |
| LOC_Os08g41489 | expressed protein |
| LOC_Os08g41500 | ubiquitin-associated protein, putative, expressed |
| LOC_Os08g41790 | phosphoribulokinase/Uridine kinase family protein, expressed |
| LOC_Os08g41880 | nucleotide pyrophosphatase/phosphodiesterase, putative, expressed |
| LOC_Os08g42000 | nuclear transport factor, putative, expressed |

| | |
|---|---|
| LOC_Os08g42010 | nodulin, putative, expressed |
| LOC_Os08g42100 | ACT domain containing protein, expressed |
| LOC_Os08g42268 | expressed protein |
| LOC_Os08g42320 | expressed protein |
| LOC_Os08g42420 | expressed protein |
| LOC_Os08g42550 | AP2 domain containing protein, expressed |
| LOC_Os08g42720 | solute carrier family 35 member F1, putative, expressed |
| LOC_Os08g43460 | expressed protein |
| LOC_Os08g44010 | WD domain, G-beta repeat domain containing protein, expressed |
| LOC_Os08g44150 | transmembrane 9 superfamily member, putative, expressed |
| LOC_Os08g44160 | expressed protein |
| LOC_Os08g44170 | transposon protein, putative, unclassified, expressed |
| LOC_Os08g44220 | GLTP domain containing protein, putative, expressed |
| LOC_Os08g44280 | protein-L-isoaspartate O-methyltransferase, putative, expressed |
| LOC_Os08g44370 | inosine-uridine preferring nucleoside hydrolase family protein, putative, expressed |
| LOC_Os08g44380 | L1P family of ribosomal proteins domain containing protein, expressed |
| LOC_Os08g44400 | glutathione S-transferase, N-terminal domain containing protein, expressed |
| LOC_Os08g44410 | harpin-induced protein 1 domain containing protein, expressed |
| LOC_Os08g44540 | expressed protein |
| LOC_Os08g44560 | expressed protein |
| LOC_Os08g45070 | expressed protein |
| LOC_Os09g01140 | expressed protein |
| LOC_Os09g01290 | hhH-GPD superfamily base excision DNA repair protein, putative, expressed |
| LOC_Os09g01310 | expressed protein |
| LOC_Os09g01330 | expressed protein |
| LOC_Os09g01360 | expressed protein |
| LOC_Os09g01370 | transposon protein, putative, unclassified, expressed |
| LOC_Os09g01620 | RNA pseudouridine synthase, putative, expressed |
| LOC_Os09g01650 | peptidyl-prolyl cis-trans isomerase, FKBP-type, putative, expressed |

| | |
|---|---|
| LOC_Os09g01660 | expressed protein |
| LOC_Os09g01690 | expressed protein |
| LOC_Os09g02130 | expressed protein |
| LOC_Os09g02140 | expressed protein |
| LOC_Os09g02214 | Citrate transporter protein, putative, expressed |
| LOC_Os09g02260 | PPR repeat containing protein, putative, expressed |
| LOC_Os09g02350 | expressed protein |
| LOC_Os09g02360 | cyclin, putative, expressed |
| LOC_Os09g02380 | expressed protein |
| LOC_Os09g02410 | S-domain receptor-like protein kinase, putative, expressed |
| LOC_Os09g02430 | expressed protein |
| LOC_Os09g02550 | expressed protein |
| LOC_Os09g02640 | expressed protein |
| LOC_Os09g02670 | expressed protein |
| LOC_Os09g02729 | phospholipase C, putative, expressed |
| LOC_Os09g02770 | translation initiation factor IF-2, putative, expressed |
| LOC_Os09g02790 | zinc-binding protein, putative, expressed |
| LOC_Os09g03990 | expressed protein |
| LOC_Os09g04160 | expressed protein |
| LOC_Os09g04300 | expressed protein |
| LOC_Os09g04310 | expressed protein |
| LOC_Os09g04440 | DNA-binding protein, putative, expressed |
| LOC_Os09g04470 | retrotransposon protein, putative, Ty3-gypsy subclass |
| LOC_Os09g08150 | expressed protein |
| LOC_Os09g08890 | mitochondrial 60S ribosomal protein L5, putative, expressed |
| LOC_Os09g08990 | cytochrome P450, putative, expressed |
| LOC_Os09g09320 | nuclear transport factor, putative, expressed |
| LOC_Os09g09350 | expressed protein |
| LOC_Os09g09360 | esterase/lipase/thioesterase, putative, expressed |

| | |
|---|---|
| LOC_Os09g09370 | BTBN18 - Bric-a-Brac, Tramtrack, Broad Complex BTB domain with non-phototropic hypocotyl 3 NPH3 domain, expressed |
| LOC_Os09g09470 | WD domain, G-beta repeat domain containing protein, expressed |
| LOC_Os09g09480 | SNF7 domain containing protein, putative, expressed |
| LOC_Os09g09530 | expressed protein |
| LOC_Os09g09930 | heavy metal transport/detoxification protein, putative, expressed |
| LOC_Os09g09970 | retrotransposon protein, putative, unclassified, expressed |
| LOC_Os09g10560 | expressed protein |
| LOC_Os09g10600 | enoyl-acyl-carrier-protein reductase NADH, chloroplast precursor, expressed |
| LOC_Os09g10630 | expressed protein |
| LOC_Os09g10640 | retrotransposon protein, putative, unclassified, expressed |
| LOC_Os09g10650 | phosphatidylinositol-4-phosphate 5-Kinase family protein, putative, expressed |
| LOC_Os09g10710 | R-interacting factor 1, putative, expressed |
| LOC_Os09g10740 | mitochondrial import inner membrane translocase subunit Tim17, putative, expressed |
| LOC_Os09g10820 | oxidoreductase domain protein, putative, expressed |
| LOC_Os09g10850 | meiotic coiled-coil protein 7, putative, expressed |
| LOC_Os09g10930 | gamma response I protein, putative, expressed |
| LOC_Os09g10980 | ZOS9-02 - C2H2 zinc finger protein, expressed |
| LOC_Os09g10990 | transposon protein, putative, CACTA, En/Spm sub-class, expressed |
| LOC_Os09g11000 | expressed protein |
| LOC_Os09g11010 | retrotransposon protein, putative, unclassified, expressed |
| LOC_Os09g13610 | PFT1, putative, expressed |
| LOC_Os09g13810 | expressed protein |
| LOC_Os09g15330 | transporter family protein, putative, expressed |
| LOC_Os09g15389 | expressed protein |
| LOC_Os09g15480 | Ser/Thr-rich protein T10 in DGCR region, putative, expressed |
| LOC_Os09g15530 | expressed protein |
| LOC_Os09g15580 | expressed protein |
| LOC_Os09g15590 | OsFBX316 - F-box domain containing protein, expressed |
| LOC_Os09g17610 | RING finger protein, putative, expressed |

| | |
|---|---|
| LOC_Os09g17750 | oxidoreductase, short chain dehydrogenase/reductase family domain containing protein, expressed |
| LOC_Os09g17790 | expressed protein |
| LOC_Os09g17880 | senescence-induced receptor-like serine/threonine-protein kinase precursor, putative, expressed |
| LOC_Os09g19320 | expressed protein |
| LOC_Os09g19734 | isochorismate synthase 1, chloroplast precursor, putative, expressed |
| LOC_Os09g19750 | expressed protein |
| LOC_Os09g19880 | expressed protein |
| LOC_Os09g19970 | expressed protein |
| LOC_Os09g19980 | GTP binding protein, putative, expressed |
| LOC_Os09g20040 | resistance-like protein I2GA-SH194-2, putative, expressed |
| LOC_Os09g20340 | DUF630/DUF632 domains containing protein, putative, expressed |
| LOC_Os09g20360 | expressed protein |
| LOC_Os09g20400 | expressed protein |
| LOC_Os09g24200 | RAD23 DNA repair protein, putative, expressed |
| LOC_Os09g24210 | expressed protein |
| LOC_Os09g24440 | expressed protein |
| LOC_Os09g24530 | ribulose-1,5 bisphosphate carboxylase/oxygenase large subunit N-methyltransferase, chloroplast precursor, putative, expressed |
| LOC_Os09g24540 | peptidyl-prolyl cis-trans isomerase, putative, expressed |
| LOC_Os09g24570 | growth regulator related protein, putative, expressed |
| LOC_Os09g24640 | pentatricopeptide, putative, expressed |
| LOC_Os09g24680 | pentatricopeptide, putative, expressed |
| LOC_Os09g24700 | expressed protein |
| LOC_Os09g24820 | ZF-HD protein dimerisation region containing protein, expressed |
| LOC_Os09g26060 | transposon protein, putative, unclassified, expressed |
| LOC_Os09g26470 | zinc finger, C3HC4 type family protein, expressed |
| LOC_Os09g26680 | transposon protein, putative, Ac/Ds sub-class, expressed |
| LOC_Os09g26690 | transposon protein, putative, unclassified, expressed |
| LOC_Os09g26730 | chaperonin, putative, expressed |
| LOC_Os09g26830 | lung seven transmembrane domain containing protein, putative, expressed |

| | |
|---|---|
| LOC_Os09g26880 | aldehyde dehydrogenase, putative, expressed |
| LOC_Os09g27970 | expressed protein |
| LOC_Os09g28450 | paramyosin, putative, expressed |
| LOC_Os09g28560 | protein phosphotase protein, putative, expressed |
| LOC_Os09g28590 | AGAP010609-PA, putative, expressed |
| LOC_Os09g28600 | expressed protein |
| LOC_Os09g28840 | OsSCP43 - Putative Serine Carboxypeptidase homologue, expressed |
| LOC_Os09g28850 | expressed protein |
| LOC_Os09g29050 | mitochondrial carrier protein, putative, expressed |
| LOC_Os09g29060 | expressed protein |
| LOC_Os09g29100 | cyclin, putative, expressed |
| LOC_Os09g29110 | expressed protein |
| LOC_Os09g29200 | glutathione S-transferase, putative, expressed |
| LOC_Os09g29390 | plastocyanin-like domain containing protein, putative, expressed |
| LOC_Os09g29470 | AGAP005770-PA, putative, expressed |
| LOC_Os09g31360 | expressed protein |
| LOC_Os09g31410 | Os9bglu29 - beta-glucosidase homologue, similar to Os4bglu12 exoglucanase, expressed |
| LOC_Os09g31482 | splicing factor U2AF, putative, expressed |
| LOC_Os09g32010 | ternary complex factor MIP1, putative, expressed |
| LOC_Os09g32020 | ubiquitin fusion degradation protein, putative, expressed |
| LOC_Os09g32080 | CHIT13 - Chitinase family protein precursor, expressed |
| LOC_Os09g32095 | expressed protein |
| LOC_Os09g32150 | expressed protein |
| LOC_Os09g32220 | pob, putative, expressed |
| LOC_Os09g32300 | KH domain-containing protein, putative, expressed |
| LOC_Os09g32310 | expressed protein |
| LOC_Os09g32340 | expressed protein |
| LOC_Os09g32570 | alcohol dehydrogenase GroES-like domain containing protein, expressed |
| LOC_Os09g32590 | expressed protein |

| | |
|---|---|
| LOC_Os09g32740 | ubiquitin carboxyl-terminal hydrolase family protein, expressed |
| LOC_Os09g32800 | peptidase, T1 family, putative, expressed |
| LOC_Os09g32810 | ribulose-phosphate 3-epimerase, putative, expressed |
| LOC_Os09g32850 | expressed protein |
| LOC_Os09g33510 | hydroxyproline-rich glycoprotein family protein, putative, expressed |
| LOC_Os09g33640 | expressed protein |
| LOC_Os09g33690 | Os9bglu32 - beta-glucosidase homologue, similar to G |
| LOC_Os09g33820 | lecithine cholesterol acyltransferase, putative, expressed |
| LOC_Os09g33980 | bromodomain protein 103, putative, expressed |
| LOC_Os09g33986 | peptidase, T1 family, putative, expressed |
| LOC_Os09g33992 | expressed protein |
| LOC_Os09g34010 | expressed protein |
| LOC_Os09g34070 | RNA recognition motif containing protein, expressed |
| LOC_Os09g34150 | NBS-LRR disease resistance protein, putative, expressed |
| LOC_Os09g36890 | hydroxyproline-rich glycoprotein family protein, putative, expressed |
| LOC_Os09g38400 | prostatic spermine-binding protein precursor, putative, expressed |
| LOC_Os09g38410 | sialin, putative, expressed |
| LOC_Os09g39190 | OsFBO22 - F-box and other domain containing protein, expressed |
| LOC_Os09g39360 | expressed protein |
| LOC_Os09g39410 | male sterility protein, putative, expressed |
| LOC_Os09g39940 | plastocyanin-like domain containing protein, putative, expressed |
| LOC_Os10g01044 | isoflavone reductase, putative, expressed |
| LOC_Os10g01100 | receptor-like protein kinase, putative, expressed |
| LOC_Os10g01134 | OsSCP46 - Putative Serine Carboxypeptidase homologue, expressed |
| LOC_Os10g01240 | VQ domain containing protein, putative, expressed |
| LOC_Os10g03260 | expressed protein |
| LOC_Os10g03550 | expressed protein |
| LOC_Os10g03620 | OsFBX344 - F-box domain containing protein, expressed |
| LOC_Os10g03660 | OsFBX345 - F-box domain containing protein, expressed |

| | |
|---|---|
| LOC_Os10g06510 | cyclin-dependent kinase G-1, putative, expressed |
| LOC_Os10g06540 | vesicle-associated membrane protein, putative, expressed |
| LOC_Os10g06600 | expressed protein |
| LOC_Os10g06770 | expressed protein |
| LOC_Os10g06910 | expressed protein |
| LOC_Os10g07210 | hsp20/alpha crystallin family protein, putative, expressed |
| LOC_Os10g07430 | expressed protein |
| LOC_Os10g07440 | expressed protein |
| LOC_Os10g07450 | expressed protein |
| LOC_Os10g07546 | expressed protein |
| LOC_Os10g08400 | expressed protein |
| LOC_Os10g08580 | FAD binding domain of DNA photolyase domain containing protein, expressed |
| LOC_Os10g08590 | expressed protein |
| LOC_Os10g08710 | chalcone synthase, putative, expressed |
| LOC_Os10g08780 | expressed protein |
| LOC_Os10g08840 | hydrolase, alpha/beta fold family domain containing protein, expressed |
| LOC_Os10g09820 | no apical meristem protein, putative, expressed |
| LOC_Os10g09850 | EF hand family protein, putative, expressed |
| LOC_Os10g09910 | expressed protein |
| LOC_Os10g09920 | LTPL132 - Protease inhibitor/seed storage/LTP family protein precursor, putative, expressed |
| LOC_Os10g10000 | expressed protein |
| LOC_Os10g10020 | expressed protein |
| LOC_Os10g10180 | methyltransferase domain containing protein, putative, expressed |
| LOC_Os10g10244 | alanyl-tRNA synthetase, putative, expressed |
| LOC_Os10g10260 | expressed protein |
| LOC_Os10g10320 | expressed protein |
| LOC_Os10g10550 | hypothetical protein |
| LOC_Os10g13690 | retrotransposon protein, putative, unclassified, expressed |
| LOC_Os10g17690 | NB-ARC domain containing protein, expressed |

| | |
|---|---|
| LOC_Os10g17950 | TKL_IRAK_DUF26-lc |
| LOC_Os10g17990 | phosphatidate cytidylyltransferase, putative, expressed |
| LOC_Os10g18070 | expressed protein |
| LOC_Os10g18080 | expressed protein |
| LOC_Os10g18084 | expressed protein |
| LOC_Os10g21196 | chloroplast 30S ribosomal protein S16, putative, expressed |
| LOC_Os10g21398 | NADPH-dependent oxidoreductase, putative, expressed |
| LOC_Os10g21574 | expressed protein |
| LOC_Os10g21930 | OsFBK21 - F-box domain and kelch repeat containing protein, expressed |
| LOC_Os10g21940 | expressed protein |
| LOC_Os10g22410 | yippee zinc-binding protein, putative, expressed |
| LOC_Os10g22510 | MLA1, putative, expressed |
| LOC_Os10g22520 | cellulase, putative, expressed |
| LOC_Os10g22590 | POEI24 - Pollen Ole e I allergen and extensin family protein precursor, expressed |
| LOC_Os10g22740 | expressed protein |
| LOC_Os10g22860 | receptor kinase, putative, expressed |
| LOC_Os10g22880 | hypothetical protein |
| LOC_Os10g22890 | receptor-like kinase, putative, expressed |
| LOC_Os10g22950 | calmodulin-binding transcription activator, putative, expressed |
| LOC_Os10g23020 | hypothetical protein |
| LOC_Os10g23060 | expressed protein |
| LOC_Os10g23120 | pyridoxamine 5'-phosphate oxidase family protein, putative, expressed |
| LOC_Os10g23130 | cytochrome P450 88A1, putative, expressed |
| LOC_Os10g23230 | AT hook-containing MAR binding 1, putative, expressed |
| LOC_Os10g24490 | retrotransposon protein, putative, unclassified |
| LOC_Os10g27460 | expressed protein |
| LOC_Os10g28030 | OsPOP22 - Putative Prolyl Oligopeptidase homologue, expressed |
| LOC_Os10g28040 | histone acetyltransferase GCN5, putative, expressed |
| LOC_Os10g28080 | glycosyl hydrolase, putative, expressed |

| | |
|---|---|
| LOC_Os10g29660 | TATA-binding protein, putative, expressed |
| LOC_Os10g29670 | expressed protein |
| LOC_Os10g29850 | MBTB60 - Bric-a-Brac, Tramtrack, Broad Complex BTB domain with Meprin and TRAF Homology MATH domain, expressed |
| LOC_Os10g30054 | ENT domain containing protein, expressed |
| LOC_Os10g30550 | tRNA methyltransferase, putative, expressed |
| LOC_Os10g30560 | UDP-glucoronosyl and UDP-glucosyl transferase domain containing protein, expressed |
| LOC_Os10g30640 | translin, putative, expressed |
| LOC_Os10g30970 | expressed protein |
| LOC_Os10g31030 | EMB1691, putative, expressed |
| LOC_Os10g31050 | expressed protein |
| LOC_Os10g32080 | xyloglucan galactosyltransferase KATAMARI1, putative, expressed |
| LOC_Os10g33440 | NB-ARC domain containing protein, expressed |
| LOC_Os10g33450 | expressed protein |
| LOC_Os10g33460 | alcohol oxidase, putative, expressed |
| LOC_Os10g35780 | expressed protein |
| LOC_Os10g37690 | inner membrane protein, putative, expressed |
| LOC_Os10g37720 | hydrolase, alpha/beta fold family protein, putative, expressed |
| LOC_Os10g38370 | retrotransposon protein, putative, Ty3-gypsy subclass, expressed |
| LOC_Os10g39430 | expressed protein |
| LOC_Os10g40920 | pentatricopeptide, putative, expressed |
| LOC_Os10g40934 | flavonol synthase/flavanone 3-hydroxylase, putative, expressed |
| LOC_Os10g41030 | agenet domain containing protein, putative, expressed |
| LOC_Os10g41160 | expressed protein |
| LOC_Os10g41200 | MYB family transcription factor, putative, expressed |
| LOC_Os10g41440 | KH domain containing protein, putative, expressed |
| LOC_Os10g41510 | calcineurin B, putative, expressed |
| LOC_Os10g42500 | PAP fibrillin family domain containing protein, expressed |
| LOC_Os10g42510 | expressed protein |
| LOC_Os10g42520 | expressed protein |

| | |
|---|---|
| LOC_Os10g42540 | expressed protein |
| LOC_Os10g42690 | jmjC domain containing protein, expressed |
| LOC_Os10g42810 | hypothetical protein |
| LOC_Os10g42820 | early-responsive to dehydration protein-related, putative, expressed |
| LOC_Os10g42830 | transporter family protein, putative, expressed |
| LOC_Os10g42840 | NADH dehydrogenase 1 alpha subcomplex subunit 12, putative, expressed |
| LOC_Os10g43075 | expressed protein |
| LOC_Os11g01010 | autophagy-related protein 8D, putative, expressed |
| LOC_Os11g01040 | monoglyceride lipase, putative, expressed |
| LOC_Os11g01050 | exo70 exocyst complex subunit domain containing protein, expressed |
| LOC_Os11g01060 | expressed protein |
| LOC_Os11g01170 | tonneau 1b, putative, expressed |
| LOC_Os11g01180 | ulp1 protease family protein, putative, expressed |
| LOC_Os11g01200 | senescence-induced receptor-like serine/threonine-protein kinase precursor, putative, expressed |
| LOC_Os11g01260 | expressed protein |
| LOC_Os11g01270 | mitochondrial substrate carrier family protein, putative, expressed |
| LOC_Os11g01480 | MYB family transcription factor, putative, expressed |
| LOC_Os11g01490 | outer membrane protein, OMP85 family, putative, expressed |
| LOC_Os11g01530 | ferritin-1, chloroplast precursor, putative, expressed |
| LOC_Os11g01540 | expressed protein |
| LOC_Os11g01570 | PMR5, putative, expressed |
| LOC_Os11g01610 | prenylated rab acceptor, putative, expressed |
| LOC_Os11g01660 | expressed protein |
| LOC_Os11g01670 | expressed protein |
| LOC_Os11g01740 | serine/threonine-protein kinase, putative, expressed |
| LOC_Os11g01880 | expressed protein |
| LOC_Os11g01890 | expressed protein |
| LOC_Os11g02054 | ACT domain containing protein, expressed |
| LOC_Os11g02080 | expressed protein |

| | |
|---|---|
| LOC_Os11g02090 | expressed protein |
| LOC_Os11g02110 | expressed protein |
| LOC_Os11g02165 | LTPL57 - Protease inhibitor/seed storage/LTP family protein precursor, expressed |
| LOC_Os11g02180 | dual specificity protein phosphatase, putative, expressed |
| LOC_Os11g02190 | expressed protein |
| LOC_Os11g02220 | retrotransposon protein, putative, SINE subclass |
| LOC_Os11g02270 | expressed protein |
| LOC_Os11g02290 | expressed protein |
| LOC_Os11g02320 | nucleotidyltransferase, putative, expressed |
| LOC_Os11g03060 | SNF7 domain containing protein, putative, expressed |
| LOC_Os11g03900 | glucosamine--fructose-6-phosphate aminotransferase 1, putative, expressed |
| LOC_Os11g03910 | zinc finger DHHC domain-containing protein, putative, expressed |
| LOC_Os11g03970 | CAMK_KIN1/SNF1/Nim1_like 5 - CAMK includes calcium/calmodulin depedent protein kinases, expressed |
| LOC_Os11g03990 | WD domain, G-beta repeat domain containing protein, expressed |
| LOC_Os11g04130 | nodulin MtN21 family protein, putative, expressed |
| LOC_Os11g04180 | dual specificity protein phosphatase, putative, expressed |
| LOC_Os11g04210 | hydroxymethylglutaryl-CoA lyase, putative, expressed |
| LOC_Os11g04230 | transcription initiation factor, putative, expressed |
| LOC_Os11g04270 | zinc finger, C3HC4 type domain containing protein |
| LOC_Os11g04280 | expressed protein |
| LOC_Os11g04370 | ribosomal protein L24, putative, expressed |
| LOC_Os11g04390 | RNA recognition motif containing protein, expressed |
| LOC_Os11g05050 | stem-specific protein TSJT1, putative, expressed |
| LOC_Os11g05100 | nucleolar GTPase, putative, expressed |
| LOC_Os11g05160 | DNA binding protein, putative, expressed |
| LOC_Os11g05270 | hypothetical protein |
| LOC_Os11g05280 | expressed protein |
| LOC_Os11g05290 | stress responsive A/B Barrel domain containing protein, expressed |
| LOC_Os11g05340 | transposon protein, putative, unclassified, expressed |

| | |
|---|---|
| LOC_Os11g05580 | expressed protein |
| LOC_Os11g05780 | DUF623 domain containing protein, expressed |
| LOC_Os11g05800 | HVA22, putative, expressed |
| LOC_Os11g05810 | expressed protein |
| LOC_Os11g05850 | cyclin-T1-3, putative, expressed |
| LOC_Os11g05860 | harpin-induced protein 1 domain containing protein, expressed |
| LOC_Os11g05880 | exo70 exocyst complex subunit, putative, expressed |
| LOC_Os11g05960 | expressed protein |
| LOC_Os11g05970 | pyridine nucleotide-disulphide oxidoreductase family protein, putative, expressed |
| LOC_Os11g05980 | tetratricopeptide-like helical, putative, expressed |
| LOC_Os11g05990 | digalactosyldiacylglycerol synthase, chloroplast precursor, putative, expressed |
| LOC_Os11g06040 | PX domain containing protein, putative, expressed |
| LOC_Os11g06170 | bZIP transcriptional activator RSG, putative, expressed |
| LOC_Os11g06330 | expressed protein |
| LOC_Os11g06340 | ubiquinol-cytochrome C reductase hinge protein, putative, expressed |
| LOC_Os11g06410 | homeodomain, putative, expressed |
| LOC_Os11g06440 | expressed protein |
| LOC_Os11g06450 | hypothetical protein |
| LOC_Os11g06690 | serine esterase family protein, putative, expressed |
| LOC_Os11g06730 | ECAGL2 - ECA1 gametogenesis related family protein precursor, expressed |
| LOC_Os11g06740 | syntaxin 6, putative, expressed |
| LOC_Os11g06840 | ZOS11-01 - C2H2 zinc finger protein, expressed |
| LOC_Os11g06870 | glyoxal oxidase-related, putative, expressed |
| LOC_Os11g06900 | amidase family protein, putative, expressed |
| LOC_Os11g06990 | expressed protein |
| LOC_Os11g07030 | integrator complex subunit 3, putative, expressed |
| LOC_Os11g07050 | FHA domain containing protein, putative, expressed |
| LOC_Os11g07220 | retrotransposon protein, putative, unclassified, expressed |
| LOC_Os11g07970 | OsFBW2 - F-box domain and WD40 repeat containing protein, expressed |

| | |
|---|---|
| LOC_Os11g07980 | ion channel nompc, putative, expressed |
| LOC_Os11g08050 | expressed protein |
| LOC_Os11g08070 | ankyrin repeat family protein, putative, expressed |
| LOC_Os11g08090 | expressed protein |
| LOC_Os11g08100 | eukaryotic aspartyl protease domain containing protein, expressed |
| LOC_Os11g08170 | DEFL45 - Defensin and Defensin-like DEFL family, expressed |
| LOC_Os11g08220 | DEFL28 - Defensin and Defensin-like DEFL family, expressed |
| LOC_Os11g08314 | methyltransferase, putative, expressed |
| LOC_Os11g08350 | expressed protein |
| LOC_Os11g08360 | expressed protein |
| LOC_Os11g08370 | transporter, major facilitator family, putative, expressed |
| LOC_Os11g08400 | expressed protein |
| LOC_Os11g08680 | expressed protein |
| LOC_Os11g08707 | expressed protein |
| LOC_Os11g08750 | pectinesterase, putative, expressed |
| LOC_Os11g08760 | pectinesterase family protein, expressed |
| LOC_Os11g08940 | RNA polymerases N 8 kDa subunit, putative, expressed |
| LOC_Os11g08950 | protein kinase family protein, putative, expressed |
| LOC_Os11g08980 | preprotein translocase secA family protein, putative, expressed |
| LOC_Os11g09800 | expressed protein |
| LOC_Os11g10910 | chloroplast nucleoid DNA-binding protein, putative, expressed |
| LOC_Os11g10920 | carboxyl-terminal proteinase, putative, expressed |
| LOC_Os11g11180 | expressed protein |
| LOC_Os11g11720 | serpin domain containing protein, putative, expressed |
| LOC_Os11g12360 | transposon protein, putative, CACTA, En/Spm sub-class, expressed |
| LOC_Os11g12530 | receptor-like protein kinase 5 precursor, putative, expressed |
| LOC_Os11g12580 | conserved hypothetical protein |
| LOC_Os11g12600 | expressed protein |
| LOC_Os11g12760 | O-methyltransferase, putative, expressed |

| | |
|---|---|
| LOC_Os11g13410 | mla1, putative, expressed |
| LOC_Os11g13680 | expressed protein |
| LOC_Os11g13750 | expressed protein |
| LOC_Os11g13830 | expressed protein |
| LOC_Os11g13870 | plant-specific domain TIGR01627 family protein, expressed |
| LOC_Os11g13880 | expressed protein |
| LOC_Os11g13890 | chlorophyll A-B binding protein, putative, expressed |
| LOC_Os11g13934 | expressed protein |
| LOC_Os11g13940 | NBS-LRR disease resistance protein, putative, expressed |
| LOC_Os11g16450 | retrotransposon protein, putative, Ty3-gypsy subclass, expressed |
| LOC_Os11g16540 | tetratricopeptide repeat domain containing protein, expressed |
| LOC_Os11g16550 | uncharacterized protein ycf53, putative, expressed |
| LOC_Os11g17014 | NB-ARC domain containing protein, expressed |
| LOC_Os11g17940 | expressed protein |
| LOC_Os11g17970 | POT family protein, expressed |
| LOC_Os11g19250 | AARP2CN domain containing protein, expressed |
| LOC_Os11g19280 | hypothetical protein |
| LOC_Os11g19290 | lipase, putative, expressed |
| LOC_Os11g19300 | transposon protein, putative, Mutator sub-class, expressed |
| LOC_Os11g19454 | hypothetical protein |
| LOC_Os11g19480 | receptor protein kinase-like protein At4g34220 precursor, putative, expressed |
| LOC_Os11g19710 | expressed protein |
| LOC_Os11g19730 | expressed protein |
| LOC_Os11g19740 | expressed protein |
| LOC_Os11g19810 | PWWP domain containing protein, expressed |
| LOC_Os11g20050 | expressed protein |
| LOC_Os11g20100 | expressed protein |
| LOC_Os11g20330 | expressed protein |
| LOC_Os11g26340 | RALFL24 - Rapid ALkalinization Factor RALF family protein precursor, expressed |

| | |
|---|---|
| LOC_Os11g26350 | expressed protein |
| LOC_Os11g26580 | MEGL18 - Maternally expressed gene MEG family protein precursor, putative, expressed |
| LOC_Os11g26725 | expressed protein |
| LOC_Os11g31610 | expressed protein |
| LOC_Os11g31870 | retrotransposon protein, putative, unclassified, expressed |
| LOC_Os11g34000 | hypothetical protein |
| LOC_Os11g34110 | heparan-alpha-glucosaminide N-acetyltransferase, putative, expressed |
| LOC_Os11g34130 | expressed protein |
| LOC_Os11g34470 | expressed protein |
| LOC_Os11g34860 | palmitoyltransferase TIP1, putative, expressed |
| LOC_Os11g35030 | growth regulating factor protein, putative, expressed |
| LOC_Os11g35090 | kinesin motor domain containing protein, putative, expressed |
| LOC_Os11g35100 | conserved hypothetical protein |
| LOC_Os11g35320 | BSD domain-containing protein, putative, expressed |
| LOC_Os11g38840 | expressed protein |
| LOC_Os11g40650 | expressed protein |
| LOC_Os11g41820 | RNA recognition motif containing protein, expressed |
| LOC_Os11g43770 | Leucine Rich Repeat family protein, expressed |
| LOC_Os11g45410 | expressed protein |
| LOC_Os11g45600 | rust resistance protein Rp1-dp8, putative, expressed |
| LOC_Os11g47830 | RNA recognition motif containing protein, expressed |
| LOC_Os12g01180 | WRKY57, expressed |
| LOC_Os12g01590 | expressed protein |
| LOC_Os12g01600 | retrotransposon protein, putative, unclassified, expressed |
| LOC_Os12g01820 | ATCHX, putative, expressed |
| LOC_Os12g01955 | expressed protein |
| LOC_Os12g04390 | hypothetical protein |
| LOC_Os12g05715 | expressed protein |
| LOC_Os12g07890 | expressed protein |

| | |
|---|---|
| LOC_Os12g09455 | expressed protein |
| LOC_Os12g09510 | hypothetical protein |
| LOC_Os12g09540 | phosphoribosylamine--glycine ligase, putative, expressed |
| LOC_Os12g09580 | leafbladeless1, putative, expressed |
| LOC_Os12g09590 | leafbladeless1, putative, expressed |
| LOC_Os12g09700 | Jacalin-like lectin domain containing protein, putative, expressed |
| LOC_Os12g12170 | cytochrome b5-like Heme/Steroid binding domain containing protein, expressed |
| LOC_Os12g12220 | DEFL18 - Defensin and Defensin-like DEFL family, expressed |
| LOC_Os12g12260 | diacylglycerol kinase 1, putative, expressed |
| LOC_Os12g12300 | EDM2, putative, expressed |
| LOC_Os12g12370 | outer envelope protein, putative, expressed |
| LOC_Os12g12470 | NADP-dependent oxidoreductase, putative, expressed |
| LOC_Os12g12854 | expressed protein |
| LOC_Os12g12950 | expressed protein |
| LOC_Os12g13000 | expressed protein |
| LOC_Os12g13030 | osFTL7 FT-Like7 homologous to Flowering Locus T gene; contains Pfam profile PF01161: Phosphatidylethanolamine-binding protein, expressed |
| LOC_Os12g13120 | histidine triad family protein, putative, expressed |
| LOC_Os12g13250 | expressed protein |
| LOC_Os12g13360 | cullin-1, putative, expressed |
| LOC_Os12g14630 | retrotransposon protein, putative, LINE subclass, expressed |
| LOC_Os12g14939 | expressed protein |
| LOC_Os12g15520 | expressed protein |
| LOC_Os12g16080 | expressed protein |
| LOC_Os12g16120 | expressed protein |
| LOC_Os12g16160 | CCT motif family protein, expressed |
| LOC_Os12g16180 | lipase, putative, expressed |
| LOC_Os12g16190 | expressed protein |
| LOC_Os12g16200 | glutathione synthetase, chloroplast precursor, putative, expressed |
| LOC_Os12g16210 | zinc finger family protein, putative, expressed |

| | |
|---|---|
| LOC_Os12g17280 | expressed protein |
| LOC_Os12g17600 | ribulose bisphosphate carboxylase small chain, chloroplast precursor, putative, expressed |
| LOC_Os12g17880 | armadillo/beta-catenin repeat protein-related, putative, expressed |
| LOC_Os12g17930 | expressed protein |
| LOC_Os12g17950 | expressed protein |
| LOC_Os12g18120 | zinc finger C-x8-C-x5-C-x3-H type family protein, expressed |
| LOC_Os12g18900 | aminotransferase, putative, expressed |
| LOC_Os12g19030 | copine, putative, expressed |
| LOC_Os12g19040 | expressed protein |
| LOC_Os12g19110 | expressed protein |
| LOC_Os12g19170 | expressed protein |
| LOC_Os12g19260 | tetratricopeptide-like helical, putative, expressed |
| LOC_Os12g19270 | expressed protein |
| LOC_Os12g19590 | WD domain, G-beta repeat domain containing protein, expressed |
| LOC_Os12g21560 | retrotransposon protein, putative, unclassified, expressed |
| LOC_Os12g24910 | retrotransposon protein, putative, Ty3-gypsy subclass |
| LOC_Os12g25230 | hypothetical protein |
| LOC_Os12g27440 | expressed protein |
| LOC_Os12g27470 | expressed protein |
| LOC_Os12g27750 | OsFBX441 - F-box domain containing protein, expressed |
| LOC_Os12g27950 | expressed protein |
| LOC_Os12g28015 | expressed protein |
| LOC_Os12g29470 | hypothetical protein |
| LOC_Os12g29480 | SAM domain containing protein, putative, expressed |
| LOC_Os12g29660 | DEAD-box ATP-dependent RNA helicase, putative, expressed |
| LOC_Os12g29750 | protein of unknown function DUF502 domain containing protein, expressed |
| LOC_Os12g29760 | oxidoreductase, aldo/keto reductase family protein, putative, expressed |
| LOC_Os12g30110 | hypothetical protein |
| LOC_Os12g31420 | expressed protein |

| | |
|---|---|
| LOC_Os12g33946 | cytochrome c oxidase subunit 1, putative, expressed |
| LOC_Os12g38100 | membrane associated DUF588 domain containing protein, putative, expressed |
| LOC_Os12g38110 | importin subunit beta, putative, expressed |
| LOC_Os12g38120 | thaumatin family domain containing protein, expressed |
| LOC_Os12g38140 | expressed protein |
| LOC_Os12g38320 | expressed protein |
| LOC_Os12g40070 | B3 DNA binding domain containing protein, expressed |
| LOC_Os12g40100 | expressed protein |
| LOC_Os12g40160 | expressed protein |
| LOC_Os12g40330 | expressed protein |
| LOC_Os12g40340 | EXS family protein, putative, expressed |
| LOC_Os12g40390 | expressed protein |
| LOC_Os12g40540 | FAD dependent oxidoreductase, putative, expressed |
| LOC_Os12g40840 | expressed protein |
| LOC_Os12g40910 | expressed protein |
| LOC_Os12g41590 | oxidoreductase, short chain dehydrogenase/reductase family domain containing protein, expressed |
| LOC_Os12g41620 | WD domain, G-beta repeat domain containing protein, expressed |
| LOC_Os12g41800 | VTC2, putative, expressed |
| LOC_Os12g41900 | SET domain containing protein, expressed |
| LOC_Os12g42150 | WD domain, G-beta repeat domain containing protein, expressed |
| LOC_Os12g42160 | kinesin motor domain containing protein, putative, expressed |

**Private genes (DUPs) in japonica against rufipogon**

| | |
|---|---|
| LOC_Os01g01430 | No apical meristem protein, putative, expressed |
| LOC_Os01g01520 | transferase family protein, putative, expressed |
| LOC_Os01g01530 | transposon protein, putative, CACTA, En/Spm sub-class |
| LOC_Os01g02240 | transposon protein, putative, unclassified, expressed |
| LOC_Os01g03720 | MYB family transcription factor, putative, expressed |
| LOC_Os01g04750 | B3 DNA binding domain containing protein, expressed |

| | |
|---|---|
| LOC_Os01g06800 | expressed protein |
| LOC_Os01g07870 | ABC transporter family protein, putative, expressed |
| LOC_Os01g10030 | expressed protein |
| LOC_Os01g11190 | expressed protein |
| LOC_Os01g11330 | retrotransposon protein, putative, unclassified, expressed |
| LOC_Os01g13300 | B3 DNA binding domain containing protein, expressed |
| LOC_Os01g13840 | PE-PGRS family protein, putative, expressed |
| LOC_Os01g14910 | expressed protein |
| LOC_Os01g15420 | retrotransposon protein, putative, Ty3-gypsy subclass, expressed |
| LOC_Os01g17060 | expressed protein |
| LOC_Os01g18250 | subtilase, putative, expressed |
| LOC_Os01g19640 | retrotransposon protein, putative, Ty3-gypsy subclass |
| LOC_Os01g19840 | 60S ribosomal protein L29-2, putative, expressed |
| LOC_Os01g21320 | NAD dependent epimerase/dehydratase family domain containing protein, expressed |
| LOC_Os01g24080 | expressed protein |
| LOC_Os01g31629 | expressed protein |
| LOC_Os01g32530 | transposon protein, putative, CACTA, En/Spm sub-class |
| LOC_Os01g32550 | hypothetical protein |
| LOC_Os01g33900 | expressed protein |
| LOC_Os01g34620 | OsGrx_S15 1 - glutaredoxin subgroup II, expressed |
| LOC_Os01g36340 | expressed protein |
| LOC_Os01g37670 | OsFBX15 - F-box domain containing protein, expressed |
| LOC_Os01g37690 | sodium/calcium exchanger protein, putative, expressed |
| LOC_Os01g39280 | expressed protein |
| LOC_Os01g40180 | expressed protein |
| LOC_Os01g40499 | S-locus lectin protein kinase family protein, putative, expressed |
| LOC_Os01g50710 | hypothetical protein |
| LOC_Os01g63680 | MYB family transcription factor, putative, expressed |
| LOC_Os01g64010 | dual specificity protein phosphatase, putative, expressed |

| | |
|---|---|
| LOC_Os01g64860 | OsSub11 - Putative Subtilisin homologue, expressed |
| LOC_Os01g64870 | expressed protein |
| LOC_Os01g66600 | rhodanese-like, putative, expressed |
| LOC_Os01g67620 | expressed protein |
| LOC_Os01g69050 | lysine ketoglutarate reductase trans-splicing related 1, putative, expressed |
| LOC_Os01g70130 | protein kinase domain containing protein, expressed |
| LOC_Os01g71020 | expressed protein |
| LOC_Os01g71170 | classical arabinogalactan protein 26 precursor, putative, expressed |
| LOC_Os01g73390 | expressed protein |
| LOC_Os02g01010 | OsPDIL1-4 protein disulfide isomerase PDIL1-4, expressed |
| LOC_Os02g01020 | expressed protein |
| LOC_Os02g03280 | transmembrane BAX inhibitor motif-containing protein, putative, expressed |
| LOC_Os02g05140 | expressed protein |
| LOC_Os02g06100 | expressed protein |
| LOC_Os02g07320 | expressed protein |
| LOC_Os02g07650 | zinc-binding protein, putative, expressed |
| LOC_Os02g08040 | expressed protein |
| LOC_Os02g24380 | hypothetical protein |
| LOC_Os02g24390 | expressed protein |
| LOC_Os02g24400 | expressed protein |
| LOC_Os02g26400 | nuclease, EndA/NucM family protein, expressed |
| LOC_Os02g41860 | aquaporin protein, putative, expressed |
| LOC_Os02g47330 | expressed protein |
| LOC_Os02g48020 | expressed protein |
| LOC_Os02g51799 | MYB family transcription factor, putative, expressed |
| LOC_Os03g04340 | C-methyltransferase, putative, expressed |
| LOC_Os03g04350 | expressed protein |
| LOC_Os03g04990 | expressed protein |
| LOC_Os03g05210 | WD domain, G-beta repeat domain containing protein, expressed |

| | |
|---|---|
| LOC_Os03g05220 | expressed protein |
| LOC_Os03g05225 | expressed protein |
| LOC_Os03g06990 | oxidoreductase, putative, expressed |
| LOC_Os03g09840 | C2 domain containing protein, putative, expressed |
| LOC_Os03g10710 | expressed protein |
| LOC_Os03g13070 | thaumatin family domain containing protein, expressed |
| LOC_Os03g18050 | OsSAUR13 - Auxin-responsive SAUR gene family member, expressed |
| LOC_Os03g19380 | calvin cycle protein CP12, putative, expressed |
| LOC_Os03g19960 | ATP-dependent RNA helicase, putative, expressed |
| LOC_Os03g21990 | WD domain, G-beta repeat domain containing protein, expressed |
| LOC_Os03g25750 | SNARE associated Golgi protein, putative, expressed |
| LOC_Os03g25760 | calmodulin-binding protein, putative, expressed |
| LOC_Os03g41570 | transposon protein, putative, unclassified, expressed |
| LOC_Os03g41580 | transposon protein, putative, CACTA, En/Spm sub-class |
| LOC_Os03g55570 | upstream activation factor subunit spp27, putative, expressed |
| LOC_Os03g56270 | receptor protein kinase CLAVATA1 precursor, putative, expressed |
| LOC_Os03g57799 | transposon protein, putative, CACTA, En/Spm sub-class, expressed |
| LOC_Os04g01130 | chromatin modification-related protein EAF3, putative, expressed |
| LOC_Os04g01140 | cytochrome P450 93A2, putative, expressed |
| LOC_Os04g02660 | retrotransposon protein, putative, unclassified |
| LOC_Os04g19500 | retrotransposon protein, putative, unclassified, expressed |
| LOC_Os04g20220 | elongation factor Tu, putative, expressed |
| LOC_Os04g20230 | neighbor of COX4, putative, expressed |
| LOC_Os04g23440 | helix-loop-helix DNA-binding domain containing protein, expressed |
| LOC_Os04g32000 | expressed protein |
| LOC_Os04g38610 | RAN guanine nucleotide release factor, putative, expressed |
| LOC_Os04g44020 | expressed protein |
| LOC_Os04g44050 | expressed protein |
| LOC_Os04g44070 | expressed protein |

| | |
|---|---|
| LOC_Os04g45240 | expressed protein |
| LOC_Os04g59600 | pentatricopeptide, putative, expressed |
| LOC_Os05g02360 | hypothetical protein |
| LOC_Os05g04100 | expressed protein |
| LOC_Os05g05530 | expressed protein |
| LOC_Os05g19970 | ZOS5-06 - C2H2 zinc finger protein, expressed |
| LOC_Os05g28720 | PPR repeat containing protein, expressed |
| LOC_Os05g28730 | zinc finger, C3HC4 type domain containing protein, expressed |
| LOC_Os05g28740 | universal stress protein domain containing protein, putative, expressed |
| LOC_Os05g34290 | glutathione gamma-glutamylcysteinyltransferase 1, putative, expressed |
| LOC_Os05g41300 | expressed protein |
| LOC_Os05g46170 | expressed protein |
| LOC_Os06g01250 | cytochrome P450, putative, expressed |
| LOC_Os06g04660 | oxidoreductase, 2OG-Fe oxygenase family protein, putative, expressed |
| LOC_Os06g09240 | anthocyanidin 3-O-glucosyltransferase, putative, expressed |
| LOC_Os06g31270 | polygalacturonase, putative, expressed |
| LOC_Os06g33410 | expressed protein |
| LOC_Os06g33970 | VQ domain containing protein, putative, expressed |
| LOC_Os06g34570 | retrotransposon protein, putative, unclassified, expressed |
| LOC_Os06g35400 | hypothetical protein |
| LOC_Os07g01060 | expressed protein |
| LOC_Os07g03210 | expressed protein |
| LOC_Os07g03220 | bZIP transcription factor domain containing protein, expressed |
| LOC_Os07g04850 | pre-mRNA-processing factor 31, putative, expressed |
| LOC_Os07g04880 | retrotransposon protein, putative, unclassified |
| LOC_Os07g05670 | expressed protein |
| LOC_Os07g07820 | expressed protein |
| LOC_Os07g18250 | expressed protein |
| LOC_Os07g31450 | CHR4/MI-2-LIKE, putative, expressed |

| | |
|---|---|
| LOC_Os07g31460 | peptide-Nasparagine amidase, putative, expressed |
| LOC_Os07g32400 | STE_PAK_Ste20_MST_like 1 - STE kinases include homologs to sterile 7, sterile 11 and sterile 20 from yeast, expressed |
| LOC_Os08g03570 | NAD dependent epimerase/dehydratase family protein, putative, expressed |
| LOC_Os08g03590 | expressed protein |
| LOC_Os08g03600 | expressed protein |
| LOC_Os08g03610 | LSD1 zinc finger domain containing protein, expressed |
| LOC_Os08g09210 | phosphoribosylamine--glycine ligase, putative, expressed |
| LOC_Os08g13270 | expressed protein |
| LOC_Os08g15340 | expressed protein |
| LOC_Os08g29220 | expressed protein |
| LOC_Os08g30790 | hypersensitive-induced response protein, putative, expressed |
| LOC_Os08g34249 | inhibitor I family protein, putative, expressed |
| LOC_Os09g09350 | expressed protein |
| LOC_Os09g09360 | esterase/lipase/thioesterase, putative, expressed |
| LOC_Os09g22110 | retrotransposon protein, putative, Ty3-gypsy subclass, expressed |
| LOC_Os09g29470 | AGAP005770-PA, putative, expressed |
| LOC_Os09g33690 | Os9bglu32 - beta-glucosidase homologue, similar to G max hydroxyisourate hydrolase, expressed |
| LOC_Os09g39360 | expressed protein |
| LOC_Os10g17950 | TKL_IRAK_DUF26-lc 5 - DUF26 kinases have homology to DUF26 containing loci, expressed |
| LOC_Os10g17990 | phosphatidate cytidylyltransferase, putative, expressed |
| LOC_Os10g33440 | NB-ARC domain containing protein, expressed |
| LOC_Os11g01040 | monoglyceride lipase, putative, expressed |
| LOC_Os11g11720 | serpin domain containing protein, putative, expressed |
| LOC_Os11g17970 | POT family protein, expressed |
| LOC_Os11g19810 | PWWP domain containing protein, expressed |
| LOC_Os11g48090 | helicase conserved C-terminal domain containing protein, expressed |

**Private genes (DELs) in indica against rufipogon**

| | |
|---|---|
| LOC_Os01g01170 | expressed protein |

| | |
|---|---|
| LOC_Os01g01760 | expressed protein |
| LOC_Os01g01810 | expressed protein |
| LOC_Os01g02240 | transposon protein, putative, unclassified, expressed |
| LOC_Os01g07550 | hypothetical protein |
| LOC_Os01g07620 | expressed protein |
| LOC_Os01g07790 | polygalacturonase, putative, expressed |
| LOC_Os01g08490 | conserved hypothetical protein |
| LOC_Os01g08780 | endonuclease/exonuclease/phosphatase family domain containing protein, expressed |
| LOC_Os01g11054 | phosphoenolpyruvate carboxylase, putative, expressed |
| LOC_Os01g13300 | B3 DNA binding domain containing protein, expressed |
| LOC_Os01g13840 | PE-PGRS family protein, putative, expressed |
| LOC_Os01g15080 | hypothetical protein |
| LOC_Os01g17279 | exonuclease, putative, expressed |
| LOC_Os01g22080 | expressed protein |
| LOC_Os01g22490 | 40S ribosomal protein S27a, putative, expressed |
| LOC_Os01g23239 | retrotransposon protein, putative, Ty3-gypsy subclass, expressed |
| LOC_Os01g23590 | suppressor of Mek, putative, expressed |
| LOC_Os01g23740 | OsPDIL2-2 protein disulfide isomerase PDIL2-2, expressed |
| LOC_Os01g24090 | TPR repeat, putative, expressed |
| LOC_Os01g24160 | hypothetical protein |
| LOC_Os01g25300 | expressed protein |
| LOC_Os01g28270 | retrotransposon protein, putative, Ty1-copia subclass |
| LOC_Os01g29570 | retrotransposon protein, putative, Ty3-gypsy subclass, expressed |
| LOC_Os01g32670 | expressed protein |
| LOC_Os01g34614 | DNA directed RNA polymerase, 7 kDa subunit domain containing protein, expressed |
| LOC_Os01g39120 | RING zinc finger ankyrin protein, putative, expressed |
| LOC_Os01g42670 | hypothetical protein |
| LOC_Os01g43050 | CENP-C1, putative, expressed |
| LOC_Os01g45140 | anthocyanin 3-O-beta-glucosyltransferase, putative, expressed |

| | |
|---|---|
| LOC_Os01g46240 | lipase class 3 family protein, putative, expressed |
| LOC_Os01g53220 | HSF-type DNA-binding domain containing protein, expressed |
| LOC_Os01g61300 | expressed protein |
| LOC_Os01g63430 | expressed protein |
| LOC_Os01g64970 | CAMK_CAMK_like 11 - CAMK includes calcium/calmodulin depedent protein kinases, expressed |
| LOC_Os01g66020 | protein kinase family protein, putative, expressed |
| LOC_Os01g68120 | DCL3, putative, expressed |
| LOC_Os01g68310 | sas10/Utp3 family protein, expressed |
| LOC_Os01g72340 | beta-galactosidase, putative, expressed |
| LOC_Os01g74090 | transposon protein, putative, unclassified |
| LOC_Os01g74152 | haloacid dehalogenase-like hydrolase family protein, putative, expressed |
| LOC_Os01g74310 | expressed protein |
| LOC_Os02g01550 | OsFBL6 - F-box domain and LRR containing protein, expressed |
| LOC_Os02g07350 | inositol-1-monophosphatase, putative, expressed |
| LOC_Os02g07840 | bZIP transcription factor domain containing protein, expressed |
| LOC_Os02g07850 | DUF630/DUF632 domains containing protein, putative, expressed |
| LOC_Os02g08130 | FGGY family of carbohydrate kinases, putative, expressed |
| LOC_Os02g09660 | expressed protein |
| LOC_Os02g19180 | ZOS2-06 - C2H2 zinc finger protein, expressed |
| LOC_Os02g19890 | stripe rust resistance protein Yr10, putative, expressed |
| LOC_Os02g19960 | transposon protein, putative, unclassified, expressed |
| LOC_Os02g24040 | retrotransposon protein, putative, unclassified |
| LOC_Os02g26480 | OsSCP7 - Putative Serine Carboxypeptidase homologue, expressed |
| LOC_Os02g27110 | expressed protein |
| LOC_Os02g27210 | hypothetical protein |
| LOC_Os02g28980 | peptidyl-prolyl isomerase, putative, expressed |
| LOC_Os02g30690 | steroleosin, putative, expressed |
| LOC_Os02g34290 | expressed protein |
| LOC_Os02g35770 | homeobox associated leucine zipper, putative, expressed |

| | |
|---|---|
| LOC_Os02g43560 | WRKY34, expressed |
| LOC_Os02g43800 | retrotransposon protein, putative, Ty3-gypsy subclass, expressed |
| LOC_Os02g47770 | ZF-HD protein dimerisation region containing protein, expressed |
| LOC_Os02g48210 | lectin-like protein kinase, putative, expressed |
| LOC_Os02g51450 | mTERF family protein, expressed |
| LOC_Os02g51620 | glycosyl hydrolase family 3 protein, putative, expressed |
| LOC_Os02g51750 | annexin, putative, expressed |
| LOC_Os02g58554 | expressed protein |
| LOC_Os03g04300 | ankyrin repeat domain containing protein, expressed |
| LOC_Os03g04410 | aconitate hydratase protein, putative, expressed |
| LOC_Os03g04600 | expressed protein |
| LOC_Os03g05330 | HEAT repeat family protein, putative, expressed |
| LOC_Os03g05680 | histone demethylase JARID1C, putative, expressed |
| LOC_Os03g06190 | 3-5 exonuclease family protein, putative, expressed |
| LOC_Os03g06750 | expressed protein |
| LOC_Os03g07100 | LTPL82 - Protease inhibitor/seed storage/LTP family protein precursor, expressed |
| LOC_Os03g07220 | PPR repeat domain containing protein, putative, expressed |
| LOC_Os03g07410 | expressed protein |
| LOC_Os03g07580 | OsNucAP1 - Putative Nucleoporin Autopeptidase homologue, expressed |
| LOC_Os03g09940 | sulfate transporter, putative, expressed |
| LOC_Os03g09950 | retrotransposon protein, putative, unclassified |
| LOC_Os03g12900 | squalene monooxygenase, putative, expressed |
| LOC_Os03g17710 | splicing factor, arginine/serine-rich, putative, expressed |
| LOC_Os03g18020 | rhodanese-like family protein, putative, expressed |
| LOC_Os03g18190 | importin subunit beta, putative, expressed |
| LOC_Os03g21530 | AARP2CN domain containing protein, expressed |
| LOC_Os03g21630 | PRC-barrel domain protein, putative, expressed |
| LOC_Os03g26329 | retrotransposon protein, putative, unclassified, expressed |
| LOC_Os03g30300 | 6-phosphogluconolactonase, putative, expressed |

| | |
|---|---|
| LOC_Os03g30530 | expressed protein |
| LOC_Os03g43390 | F-box/LRR domain containing protein, putative, expressed |
| LOC_Os03g48730 | expressed protein |
| LOC_Os03g50230 | protein kinase, putative, expressed |
| LOC_Os03g57910 | expressed protein |
| LOC_Os03g63770 | RCD1, putative, expressed |
| LOC_Os04g01310 | serine/threonine-protein kinase receptor precursor, putative, expressed |
| LOC_Os04g02920 | leucine-rich repeat family protein, putative, expressed |
| LOC_Os04g10820 | transposon protein, putative, unclassified, expressed |
| LOC_Os04g18200 | dihydrodipicolinate synthase, chloroplast precursor, putative, expressed |
| LOC_Os04g24520 | ribosomal protein L51, putative, expressed |
| LOC_Os04g35480 | CDK5RAP1, putative, expressed |
| LOC_Os04g39780 | AMP-binding enzyme family protein, expressed |
| LOC_Os04g40010 | PPR repeat domain containing protein, putative, expressed |
| LOC_Os04g47680 | Ser/Thr-rich protein T10 in DGCR region, putative, expressed |
| LOC_Os04g48010 | WD-40 repeat family protein, putative, expressed |
| LOC_Os04g48940 | uncharacterized mscS family protein, putative, expressed |
| LOC_Os04g52361 | ribosomal protein S17, putative, expressed |
| LOC_Os04g52390 | potassium transporter, putative, expressed |
| LOC_Os04g52660 | expressed protein |
| LOC_Os05g01360 | expressed protein |
| LOC_Os05g02030 | OB-fold nucleic acid binding domain containing protein, putative, expressed |
| LOC_Os05g02564 | expressed protein |
| LOC_Os05g03100 | HECT-domain domain containing protein, expressed |
| LOC_Os05g05860 | retrotransposon protein, putative, unclassified, expressed |
| LOC_Os05g07030 | arginyl-tRNA synthetase, putative, expressed |
| LOC_Os05g08060 | expressed protein |
| LOC_Os05g08340 | expressed protein |
| LOC_Os05g10640 | hAT dimerisation domain-containing protein, putative, expressed |

| | |
|---|---|
| LOC_Os05g30250 | Os5bglu19 - beta-glucosidase homologue, similar to G |
| LOC_Os05g30470 | collagen alpha-1, putative, expressed |
| LOC_Os05g30790 | CW-type Zinc Finger, putative, expressed |
| LOC_Os05g31056 | tetratricopeptide repeat containing protein, putative, expressed |
| LOC_Os06g01700 | CWC15 homolog A, putative, expressed |
| LOC_Os06g01890 | MADS-box transcription factor, putative, expressed |
| LOC_Os06g02019 | cytochrome P450, putative, expressed |
| LOC_Os06g06740 | MYB family transcription factor, putative, expressed |
| LOC_Os06g12730 | expressed protein |
| LOC_Os06g17700 | embryogenesis transmembrane protein, putative, expressed |
| LOC_Os06g30310 | alpha-glucan water dikinase, chloroplast precursor, putative, expressed |
| LOC_Os06g51460 | white-brown complex homolog protein, putative, expressed |
| LOC_Os07g01940 | sister chromatid cohesion 2, putative, expressed |
| LOC_Os07g18250 | expressed protein |
| LOC_Os07g27180 | tetratricopeptide repeat domain containing protein, expressed |
| LOC_Os07g28920 | expressed protein |
| LOC_Os07g31340 | regulator of nonsense transcripts 1, putative, expressed |
| LOC_Os07g31460 | peptide-Nasparagine amidase, putative, expressed |
| LOC_Os07g39300 | expressed protein |
| LOC_Os07g42940 | CAMK_CAMK_like 7 - CAMK includes calcium/calmodulin depedent protein kinases, expressed |
| LOC_Os08g03560 | chloroplast unusual positioning protein, putative, expressed |
| LOC_Os08g03670 | galactosyltransferase family protein, putative, expressed |
| LOC_Os08g04420 | tyrosine protein kinase domain containing protein, putative, expressed |
| LOC_Os08g04780 | amine oxidase, putative, expressed |
| LOC_Os08g08084 | expressed protein |
| LOC_Os08g15030 | aspartate carbamoyltransferase, putative, expressed |
| LOC_Os08g19370 | expressed protein |
| LOC_Os08g23320 | para-hydroxybenzoate--polyprenyltransferase, mitochondrial precursor, putative, expressed |
| LOC_Os08g24380 | NB-ARC domain containing protein, expressed |

| | |
|---|---|
| LOC_Os08g33700 | pentatricopeptide, putative, expressed |
| LOC_Os08g35060 | protein binding protein, putative, expressed |
| LOC_Os08g35190 | auxin-repressed protein, putative, expressed |
| LOC_Os08g41390 | peptidyl-prolyl isomerase, putative, expressed |
| LOC_Os08g41500 | ubiquitin-associated protein, putative, expressed |
| LOC_Os08g42420 | expressed protein |
| LOC_Os09g01360 | expressed protein |
| LOC_Os09g03630 | ankyrin, putative, expressed |
| LOC_Os09g06620 | OsFBX307 - F-box domain containing protein, expressed |
| LOC_Os09g07800 | expressed protein |
| LOC_Os09g12550 | WD domain, G-beta repeat domain containing protein, expressed |
| LOC_Os09g13600 | expressed protein |
| LOC_Os09g20340 | DUF630/DUF632 domains containing protein, putative, expressed |
| LOC_Os09g30454 | OsWAK87 - OsWAK receptor-like protein kinase, expressed |
| LOC_Os09g31170 | expressed protein |
| LOC_Os09g33780 | expressed protein |
| LOC_Os09g36140 | RNA recognition motif 2 domain containing protein, expressed |
| LOC_Os09g39320 | CTR1-like protein kinase, putative, expressed |
| LOC_Os10g05069 | lysosomal alpha-mannosidase precursor, putative, expressed |
| LOC_Os10g05140 | hypothetical protein |
| LOC_Os10g06720 | aldose 1-epimerase, putative, expressed |
| LOC_Os10g10170 | PPR repeat domain containing protein, putative, expressed |
| LOC_Os10g14814 | ATFMN/FHY, putative, expressed |
| LOC_Os10g20550 | DEFL70 - Defensin and Defensin-like DEFL family, expressed |
| LOC_Os10g20590 | expressed protein |
| LOC_Os10g42439 | heat shock protein DnaJ, putative, expressed |
| LOC_Os11g27010 | hypothetical protein |
| LOC_Os11g27020 | expressed protein |
| LOC_Os11g35070 | expressed protein |

| | |
|---|---|
| LOC_Os12g05609 | OsFBO24 - F-box and other domain containing protein, expressed |
| LOC_Os12g07700 | nifU, putative, expressed |
| LOC_Os12g16640 | expressed protein |
| LOC_Os12g29660 | DEAD-box ATP-dependent RNA helicase, putative, expressed |
| LOC_Os12g40540 | FAD dependent oxidoreductase, putative, expressed |
| LOC_Os12g44090 | leucine-rich repeat family protein, putative, expressed |

**Private genes (DELs) in japonica against rufipogon**

| | |
|---|---|
| LOC_Os01g01190 | phosphoesterase family protein, putative, expressed |
| LOC_Os01g01560 | transposon protein, putative, unclassified, expressed |
| LOC_Os01g01810 | expressed protein |
| LOC_Os01g02240 | transposon protein, putative, unclassified, expressed |
| LOC_Os01g07410 | growth regulator related protein, putative, expressed |
| LOC_Os01g08010 | expressed protein |
| LOC_Os01g08390 | hypothetical protein |
| LOC_Os01g08490 | conserved hypothetical protein |
| LOC_Os01g09110 | retrotransposon protein, putative, unclassified |
| LOC_Os01g10030 | expressed protein |
| LOC_Os01g10430 | protein kinase family protein, putative, expressed |
| LOC_Os01g10560 | transposon protein, putative, unclassified |
| LOC_Os01g14926 | protein kinase domain containing protein, expressed |
| LOC_Os01g15080 | hypothetical protein |
| LOC_Os01g16300 | hypothetical protein |
| LOC_Os01g17279 | exonuclease, putative, expressed |
| LOC_Os01g20110 | expressed protein |
| LOC_Os01g22490 | 40S ribosomal protein S27a, putative, expressed |
| LOC_Os01g22930 | expressed protein |
| LOC_Os01g23610 | dihydrolipoyl dehydrogenase, putative, expressed |
| LOC_Os01g24090 | TPR repeat, putative, expressed |

| | |
|---|---|
| LOC_Os01g24160 | hypothetical protein |
| LOC_Os01g29570 | retrotransposon protein, putative, Ty3-gypsy subclass, expressed |
| LOC_Os01g32184 | tetratricopeptide repeat protein 5, putative, expressed |
| LOC_Os01g32520 | expressed protein |
| LOC_Os01g34910 | expressed protein |
| LOC_Os01g39120 | RING zinc finger ankyrin protein, putative, expressed |
| LOC_Os01g39280 | expressed protein |
| LOC_Os01g42360 | hypothetical protein |
| LOC_Os01g42980 | ribosomal protein L22, putative, expressed |
| LOC_Os01g50680 | OsSub2 - Putative Subtilisin homologue, expressed |
| LOC_Os01g52260 | serine acetyltransferase protein, putative, expressed |
| LOC_Os01g56530 | DUF260 domain containing protein, putative, expressed |
| LOC_Os01g61300 | expressed protein |
| LOC_Os01g61460 | leaf senescence related protein, putative, expressed |
| LOC_Os01g64970 | CAMK_CAMK_like 11 - CAMK includes calcium/calmodulin depedent protein kinases, expressed |
| LOC_Os01g68320 | DEAD-box ATP-dependent RNA helicase 30, putative, expressed |
| LOC_Os01g69830 | OsSPL2 - SBP-box gene family member, expressed |
| LOC_Os01g71240 | calcium-transporting ATPase, plasma membrane-type, putative, expressed |
| LOC_Os01g72990 | expressed protein |
| LOC_Os01g74090 | transposon protein, putative, unclassified |
| LOC_Os01g74310 | expressed protein |
| LOC_Os02g02050 | SNF2 family N-terminal domain containing protein, expressed |
| LOC_Os02g04670 | glucan endo-1,3-beta-glucosidase precursor, putative, expressed |
| LOC_Os02g07150 | urease accessory protein F, putative, expressed |
| LOC_Os02g07840 | bZIP transcription factor domain containing protein, expressed |
| LOC_Os02g07850 | DUF630/DUF632 domains containing protein, putative, expressed |
| LOC_Os02g07900 | ANTH/ENTH domain containing protein, putative, expressed |
| LOC_Os02g19180 | ZOS2-06 - C2H2 zinc finger protein, expressed |
| LOC_Os02g30530 | transposon protein, putative, unclassified, expressed |

| | |
|---|---|
| LOC_Os02g30690 | steroleosin, putative, expressed |
| LOC_Os02g32504 | heparan-alpha-glucosaminide N-acetyltransferase, putative, expressed |
| LOC_Os02g34290 | expressed protein |
| LOC_Os02g35770 | homeobox associated leucine zipper, putative, expressed |
| LOC_Os02g38690 | protein phosphatase 2C containing protein, expressed |
| LOC_Os02g40330 | retrotransposon protein, putative, Ty3-gypsy subclass, expressed |
| LOC_Os02g44134 | thioesterase family protein, putative, expressed |
| LOC_Os02g51750 | annexin, putative, expressed |
| LOC_Os03g03164 | homeobox protein knotted-1, putative, expressed |
| LOC_Os03g04770 | beta-amylase, putative, expressed |
| LOC_Os03g04890 | zinc finger family protein, putative, expressed |
| LOC_Os03g05020 | PIR, putative, expressed |
| LOC_Os03g05330 | HEAT repeat family protein, putative, expressed |
| LOC_Os03g05500 | DUF260 domain containing protein, putative, expressed |
| LOC_Os03g06740 | glutathione reductase, putative, expressed |
| LOC_Os03g06900 | DNA topoisomerase 3 protein, putative, expressed |
| LOC_Os03g07100 | LTPL82 - Protease inhibitor/seed storage/LTP family protein precursor, expressed |
| LOC_Os03g07410 | expressed protein |
| LOC_Os03g08380 | ABC transporter, ATP-binding protein, putative, expressed |
| LOC_Os03g10010 | expressed protein |
| LOC_Os03g10400 | peptidyl-prolyl cis-trans isomerase, putative, expressed |
| LOC_Os03g14254 | expressed protein |
| LOC_Os03g18020 | rhodanese-like family protein, putative, expressed |
| LOC_Os03g18120 | proteins of unknown function domain containing protein, expressed |
| LOC_Os03g18190 | importin subunit beta, putative, expressed |
| LOC_Os03g26329 | retrotransposon protein, putative, unclassified, expressed |
| LOC_Os03g30460 | vacuolar protein sorting-associated protein 52, putative, expressed |
| LOC_Os03g41780 | retrotransposon, putative, centromere-specific, expressed |
| LOC_Os03g42900 | KH domain containing protein, putative, expressed |

| | |
|---|---|
| LOC_Os03g43970 | amino acid permease family protein, putative, expressed |
| LOC_Os03g49790 | pattern formation protein EMB30, putative, expressed |
| LOC_Os03g57910 | expressed protein |
| LOC_Os03g62379 | MYB family transcription factor, putative, expressed |
| LOC_Os04g02730 | tRNA-dihydrouridine synthase 3-like, putative, expressed |
| LOC_Os04g06790 | 2-aminoethanethiol dioxygenase, putative, expressed |
| LOC_Os04g13220 | ABC transporter family protein, putative, expressed |
| LOC_Os04g18200 | dihydrodipicolinate synthase, chloroplast precursor, putative, expressed |
| LOC_Os04g32000 | expressed protein |
| LOC_Os04g34040 | hypothetical protein |
| LOC_Os04g35260 | DEAD/DEAH box helicase domain containing protein, expressed |
| LOC_Os04g35540 | amino acid permease family protein, putative, expressed |
| LOC_Os04g52660 | expressed protein |
| LOC_Os05g01470 | methionine S-methyltransferase, putative, expressed |
| LOC_Os05g05710 | serine/threonine-protein phosphatase 2A regulatory subunit B subunitgamma, putative, expressed |
| LOC_Os05g05790 | double-stranded RNA binding motif containing protein, expressed |
| LOC_Os05g08060 | expressed protein |
| LOC_Os05g10640 | hAT dimerisation domain-containing protein, putative, expressed |
| LOC_Os05g10770 | transcription factor jumonji, putative, expressed |
| LOC_Os05g19270 | MSH-like DNA mismatch repair protein, putative, expressed |
| LOC_Os05g19470 | expressed protein |
| LOC_Os05g24160 | double-stranded RNA binding motif containing protein, expressed |
| LOC_Os05g29040 | expressed protein |
| LOC_Os05g30470 | collagen alpha-1, putative, expressed |
| LOC_Os05g31480 | methyltransferase, putative, expressed |
| LOC_Os05g39600 | ATCHX15, putative, expressed |
| LOC_Os05g51690 | CCT motif family protein, expressed |
| LOC_Os05g51860 | expressed protein |
| LOC_Os06g37224 | cytochrome P450, putative, expressed |

| | |
|---|---|
| LOC_Os07g01260 | la domain containing protein, expressed |
| LOC_Os07g13050 | hypothetical protein |
| LOC_Os07g22370 | retrotransposon protein, putative, unclassified, expressed |
| LOC_Os07g31340 | regulator of nonsense transcripts 1, putative, expressed |
| LOC_Os07g31550 | powdery mildew resistant protein 5, putative, expressed |
| LOC_Os07g42810 | adaptor complexes medium subunit family domain containing protein, expressed |
| LOC_Os08g03676 | pentatricopeptide, putative, expressed |
| LOC_Os08g12840 | FabA-like domain containing protein, expressed |
| LOC_Os08g17970 | hypothetical protein |
| LOC_Os08g35990 | expressed protein |
| LOC_Os08g40840 | exo70 exocyst complex subunit, putative, expressed |
| LOC_Os09g07294 | purity of essence, putative, expressed |
| LOC_Os09g09370 | BTBN18 - Bric-a-Brac, Tramtrack, Broad Complex BTB domain with non-phototropic hypocotyl 3 NPH3 domain, expressed |
| LOC_Os09g12550 | WD domain, G-beta repeat domain containing protein, expressed |
| LOC_Os09g31350 | expressed protein |
| LOC_Os09g36200 | senescence-inducible chloroplast stay-green protein 1, putative, expressed |
| LOC_Os10g02920 | cytochrome b561, putative, expressed |
| LOC_Os10g05069 | lysosomal alpha-mannosidase precursor, putative, expressed |
| LOC_Os10g06720 | aldose 1-epimerase, putative, expressed |
| LOC_Os10g09850 | EF hand family protein, putative, expressed |
| LOC_Os10g10420 | OsFBX378 - F-box domain containing protein, expressed |
| LOC_Os10g13840 | expressed protein |
| LOC_Os10g13880 | expressed protein |
| LOC_Os10g34795 | expressed protein |
| LOC_Os11g34000 | hypothetical protein |
| LOC_Os11g35070 | expressed protein |
| LOC_Os12g07600 | expressed protein |
| LOC_Os12g15590 | heat shock protein DnaJ, putative, expressed |
| LOC_Os12g29660 | DEAD-box ATP-dependent RNA helicase, putative, expressed |

| | |
|---|---|
| LOC_Os12g44140 | expressed protein |

**Table S11.** GO analysis for fixed genes between each two species, rufipogon, indica and japonica, and private genes underlying SVs in each subspecies.

| GO term | Ontology | Description | Number in input list | Number in BG/Ref | p-value |
|---|---|---|---|---|---|
| **Fixed genes (SVs) in indica-japonica comparison** | | | | | |
| GO:0006437 | Biological Process | tyrosyl-tRNA aminoacylation | 1 | 7 | 0.0079 |
| GO:0016481 | Biological Process | negative regulation of transcription | 1 | 9 | 0.0099 |
| GO:0045934 | Biological Process | negative regulation of nucleobase, nucleoside, nucleotide and nucleic acid metabolic process | 1 | 11 | 0.012 |
| GO:0051172 | Biological Process | negative regulation of nitrogen compound metabolic process | 1 | 11 | 0.012 |
| GO:0010629 | Biological Process | negative regulation of gene expression | 1 | 12 | 0.013 |
| GO:0031327 | Biological Process | negative regulation of cellular biosynthetic process | 1 | 50 | 0.05 |
| GO:0009890 | Biological Process | negative regulation of biosynthetic process | 1 | 50 | 0.05 |
| GO:0010558 | Biological Process | negative regulation of macromolecule biosynthetic process | 1 | 50 | 0.05 |
| GO:0004831 | Molecular Function | tyrosine-tRNA ligase activity | 1 | 7 | 0.0079 |
| GO:0004091 | Molecular Function | carboxylesterase activity | 2 | 172 | 0.013 |
| GO:0010181 | Molecular Function | FMN binding | 1 | 35 | 0.035 |
| GO:0016874 | Molecular Function | ligase activity | 2 | 305 | 0.037 |
| GO:0016829 | Molecular Function | lyase activity | 2 | 309 | 0.038 |
| GO:0008375 | Molecular Function | acetylglucosaminyltransferase activity | 1 | 40 | 0.04 |
| GO:0030176 | Cellular Component | integral to endoplasmic reticulum membrane | 1 | 3 | 0.004 |
| GO:0031301 | Cellular Component | integral to organelle membrane | 1 | 17 | 0.018 |
| GO:0031227 | Cellular Component | intrinsic to endoplasmic reticulum membrane | 1 | 23 | 0.024 |
| GO:0031300 | Cellular Component | intrinsic to organelle membrane | 1 | 37 | 0.037 |
| GO:0005789 | Cellular Component | endoplasmic reticulum membrane | 1 | 47 | 0.047 |
| GO:0042175 | Cellular Component | nuclear envelope-endoplasmic reticulum network | 1 | 47 | 0.047 |
| GO:0044432 | Cellular Component | endoplasmic reticulum part | 1 | 49 | 0.049 |
| GO:0071212 | Cellular Component | subsynaptic reticulum | 1 | 49 | 0.049 |

**Private genes (DUPs) in indica against rufipogon**

| | | | | | |
|---|---|---|---|---|---|
| GO:0044281 | Biological Process | small molecule metabolic process | 111 | 1034 | 0.0000029 |
| GO:0044248 | Biological Process | cellular catabolic process | 46 | 337 | 0.000018 |
| GO:0019637 | Biological Process | organophosphate metabolic process | 19 | 84 | 0.000019 |
| GO:0006644 | Biological Process | phospholipid metabolic process | 17 | 77 | 0.000068 |
| GO:0009056 | Biological Process | catabolic process | 52 | 433 | 0.00011 |
| GO:0044265 | Biological Process | cellular macromolecule catabolic process | 35 | 254 | 0.00015 |
| GO:0009057 | Biological Process | macromolecule catabolic process | 40 | 322 | 0.00036 |
| GO:0046486 | Biological Process | glycerolipid metabolic process | 13 | 58 | 0.00042 |
| GO:0006650 | Biological Process | glycerophospholipid metabolic process | 13 | 58 | 0.00042 |
| GO:0055114 | Biological Process | oxidation reduction | 124 | 1351 | 0.00047 |
| GO:0010467 | Biological Process | gene expression | 204 | 2424 | 0.00065 |
| GO:0030384 | Biological Process | phosphoinositide metabolic process | 10 | 43 | 0.0015 |
| GO:0005975 | Biological Process | carbohydrate metabolic process | 80 | 833 | 0.0015 |
| GO:0016070 | Biological Process | RNA metabolic process | 100 | 1088 | 0.0016 |
| GO:0044262 | Biological Process | cellular carbohydrate metabolic process | 43 | 385 | 0.0016 |
| GO:0044255 | Biological Process | cellular lipid metabolic process | 29 | 229 | 0.0017 |
| GO:0044282 | Biological Process | small molecule catabolic process | 21 | 146 | 0.0018 |
| GO:0006066 | Biological Process | alcohol metabolic process | 26 | 202 | 0.0023 |
| GO:0019222 | Biological Process | regulation of metabolic process | 142 | 1664 | 0.0028 |
| GO:0046488 | Biological Process | phosphatidylinositol metabolic process | 6 | 18 | 0.003 |
| GO:0019318 | Biological Process | hexose metabolic process | 19 | 133 | 0.0031 |
| GO:0045449 | Biological Process | regulation of transcription | 130 | 1513 | 0.0033 |
| GO:0031323 | Biological Process | regulation of cellular metabolic process | 136 | 1593 | 0.0033 |
| GO:0019219 | Biological Process | regulation of nucleobase, nucleoside, nucleotide and nucleic acid metabolic process | 130 | 1520 | 0.0038 |
| GO:0051171 | Biological Process | regulation of nitrogen compound metabolic process | 130 | 1520 | 0.0038 |
| GO:0008654 | Biological Process | phospholipid biosynthetic process | 8 | 34 | 0.0041 |
| GO:0010468 | Biological Process | regulation of gene expression | 134 | 1577 | 0.0041 |

| | | | | | |
|---|---|---|---|---|---|
| GO:0044267 | Biological Process | cellular protein metabolic process | 229 | 2877 | 0.0043 |
| GO:0005996 | Biological Process | monosaccharide metabolic process | 20 | 150 | 0.005 |
| GO:0006350 | Biological Process | transcription | 137 | 1629 | 0.0052 |
| GO:0016051 | Biological Process | carbohydrate biosynthetic process | 17 | 121 | 0.0058 |
| GO:0060255 | Biological Process | regulation of macromolecule metabolic process | 137 | 1640 | 0.0064 |
| GO:0010556 | Biological Process | regulation of macromolecule biosynthetic process | 131 | 1564 | 0.007 |
| GO:0009889 | Biological Process | regulation of biosynthetic process | 131 | 1564 | 0.007 |
| GO:0031326 | Biological Process | regulation of cellular biosynthetic process | 131 | 1564 | 0.007 |
| GO:0050789 | Biological Process | regulation of biological process | 165 | 2027 | 0.0071 |
| GO:0034637 | Biological Process | cellular carbohydrate biosynthetic process | 16 | 114 | 0.0074 |
| GO:0044275 | Biological Process | cellular carbohydrate catabolic process | 14 | 94 | 0.0075 |
| GO:0046164 | Biological Process | alcohol catabolic process | 14 | 94 | 0.0075 |
| GO:0009250 | Biological Process | glucan biosynthetic process | 11 | 65 | 0.0075 |
| GO:0045859 | Biological Process | regulation of protein kinase activity | 5 | 16 | 0.0084 |
| GO:0032147 | Biological Process | activation of protein kinase activity | 5 | 16 | 0.0084 |
| GO:0051338 | Biological Process | regulation of transferase activity | 5 | 16 | 0.0084 |
| GO:0051347 | Biological Process | positive regulation of transferase activity | 5 | 16 | 0.0084 |
| GO:0033674 | Biological Process | positive regulation of kinase activity | 5 | 16 | 0.0084 |
| GO:0043549 | Biological Process | regulation of kinase activity | 5 | 16 | 0.0084 |
| GO:0007205 | Biological Process | activation of protein kinase C activity by G-protein coupled receptor protein signaling pathway | 5 | 16 | 0.0084 |
| GO:0045860 | Biological Process | positive regulation of protein kinase activity | 5 | 16 | 0.0084 |
| GO:0006519 | Biological Process | cellular amino acid and derivative metabolic process | 40 | 391 | 0.0086 |
| GO:0044106 | Biological Process | cellular amine metabolic process | 35 | 333 | 0.0094 |
| GO:0042180 | Biological Process | cellular ketone metabolic process | 47 | 480 | 0.0096 |
| GO:0005515 | Molecular Function | protein binding | 219 | 2460 | 0.000019 |
| GO:0000166 | Molecular Function | nucleotide binding | 323 | 3874 | 0.000026 |
| GO:0016853 | Molecular Function | isomerase activity | 24 | 161 | 0.00057 |
| GO:0016491 | Molecular Function | oxidoreductase activity | 146 | 1672 | 0.0011 |
| GO:0001883 | Molecular Function | purine nucleoside binding | 261 | 3247 | 0.0014 |

| | | | | | |
|---|---|---|---|---|---|
| GO:0030554 | Molecular Function | adenyl nucleotide binding | 261 | 3247 | 0.0014 |
| GO:0017076 | Molecular Function | purine nucleotide binding | 276 | 3456 | 0.0014 |
| GO:0001882 | Molecular Function | nucleoside binding | 261 | 3257 | 0.0016 |
| GO:0048037 | Molecular Function | cofactor binding | 56 | 543 | 0.0019 |
| GO:0016817 | Molecular Function | hydrolase activity, acting on acid anhydrides | 73 | 764 | 0.0027 |
| GO:0005524 | Molecular Function | ATP binding | 246 | 3085 | 0.0028 |
| GO:0032559 | Molecular Function | adenyl ribonucleotide binding | 246 | 3087 | 0.0029 |
| GO:0005319 | Molecular Function | lipid transporter activity | 6 | 18 | 0.003 |
| GO:0016307 | Molecular Function | phosphatidylinositol phosphate kinase activity | 6 | 18 | 0.003 |
| GO:0032553 | Molecular Function | ribonucleotide binding | 260 | 3292 | 0.0034 |
| GO:0032555 | Molecular Function | purine ribonucleotide binding | 260 | 3292 | 0.0034 |
| GO:0016818 | Molecular Function | hydrolase activity, acting on acid anhydrides, in phosphorus-containing anhydrides | 72 | 759 | 0.0034 |
| GO:0001727 | Molecular Function | lipid kinase activity | 6 | 20 | 0.0047 |
| GO:0016830 | Molecular Function | carbon-carbon lyase activity | 15 | 99 | 0.005 |
| GO:0003684 | Molecular Function | damaged DNA binding | 6 | 21 | 0.0057 |
| GO:0016857 | Molecular Function | racemase and epimerase activity, acting on carbohydrates and derivatives | 5 | 15 | 0.0067 |
| GO:0008137 | Molecular Function | NADH dehydrogenase (ubiquinone) activity | 7 | 30 | 0.0073 |
| GO:0050136 | Molecular Function | NADH dehydrogenase (quinone) activity | 7 | 30 | 0.0073 |
| GO:0003954 | Molecular Function | NADH dehydrogenase activity | 7 | 30 | 0.0073 |
| GO:0017111 | Molecular Function | nucleoside-triphosphatase activity | 67 | 726 | 0.008 |
| GO:0051536 | Molecular Function | iron-sulfur cluster binding | 14 | 95 | 0.0081 |
| GO:0051540 | Molecular Function | metal cluster binding | 14 | 95 | 0.0081 |
| GO:0004143 | Molecular Function | diacylglycerol kinase activity | 5 | 16 | 0.0084 |
| GO:0016462 | Molecular Function | pyrophosphatase activity | 68 | 743 | 0.0089 |
| GO:0016866 | Molecular Function | intramolecular transferase activity | 9 | 49 | 0.0096 |
| GO:0022892 | Molecular Function | substrate-specific transporter activity | 56 | 593 | 0.0097 |
| GO:0005623 | Cellular Component | cell | 464 | 5945 | 0.00014 |
| GO:0044464 | Cellular Component | cell part | 464 | 5945 | 0.00014 |

| | | | | | |
|---|---|---|---|---|---|
| GO:0016020 | Cellular Component | membrane | 186 | 2115 | 0.00016 |
| GO:0031224 | Cellular Component | intrinsic to membrane | 79 | 775 | 0.00034 |
| GO:0016021 | Cellular Component | integral to membrane | 77 | 752 | 0.00036 |
| GO:0044425 | Cellular Component | membrane part | 97 | 1007 | 0.00046 |
| GO:0005737 | Cellular Component | cytoplasm | 117 | 1263 | 0.0005 |

**Private genes (DUPs) in japonica against rufipogon**

| | | | | | |
|---|---|---|---|---|---|
| GO:0065007 | Biological Process | biological regulation | 13 | 2125 | 0.002 |
| GO:0050794 | Biological Process | regulation of cellular process | 12 | 1955 | 0.0029 |
| GO:0050789 | Biological Process | regulation of biological process | 12 | 2027 | 0.0039 |
| GO:0060548 | Biological Process | negative regulation of cell death | 1 | 1 | 0.005 |
| GO:0042981 | Biological Process | regulation of apoptosis | 1 | 1 | 0.005 |
| GO:0010941 | Biological Process | regulation of cell death | 1 | 1 | 0.005 |
| GO:0043067 | Biological Process | regulation of programmed cell death | 1 | 1 | 0.005 |
| GO:0043066 | Biological Process | negative regulation of apoptosis | 1 | 1 | 0.005 |
| GO:0043069 | Biological Process | negative regulation of programmed cell death | 1 | 1 | 0.005 |
| GO:0046937 | Biological Process | phytochelatin metabolic process | 1 | 2 | 0.0074 |
| GO:0046938 | Biological Process | phytochelatin biosynthetic process | 1 | 2 | 0.0074 |
| GO:0045449 | Biological Process | regulation of transcription | 9 | 1513 | 0.012 |
| GO:0019219 | Biological Process | regulation of nucleobase, nucleoside, nucleotide and nucleic acid metabolic process | 9 | 1520 | 0.013 |
| GO:0051171 | Biological Process | regulation of nitrogen compound metabolic process | 9 | 1520 | 0.013 |
| GO:0009228 | Biological Process | thiamin biosynthetic process | 1 | 5 | 0.015 |
| GO:0009889 | Biological Process | regulation of biosynthetic process | 9 | 1564 | 0.015 |
| GO:0010556 | Biological Process | regulation of macromolecule biosynthetic process | 9 | 1564 | 0.015 |
| GO:0031326 | Biological Process | regulation of cellular biosynthetic process | 9 | 1564 | 0.015 |
| GO:0010468 | Biological Process | regulation of gene expression | 9 | 1577 | 0.016 |
| GO:0031323 | Biological Process | regulation of cellular metabolic process | 9 | 1593 | 0.017 |
| GO:0080090 | Biological Process | regulation of primary metabolic process | 9 | 1626 | 0.019 |
| GO:0006350 | Biological Process | transcription | 9 | 1629 | 0.019 |

| | | | | | |
|---|---|---|---|---|---|
| GO:0060255 | Biological Process | regulation of macromolecule metabolic process | 9 | 1640 | 0.02 |
| GO:0019222 | Biological Process | regulation of metabolic process | 9 | 1664 | 0.022 |
| GO:0006772 | Biological Process | thiamin metabolic process | 1 | 8 | 0.022 |
| GO:0043043 | Biological Process | peptide biosynthetic process | 1 | 8 | 0.022 |
| GO:0042723 | Biological Process | thiamin and derivative metabolic process | 1 | 8 | 0.022 |
| GO:0042724 | Biological Process | thiamin and derivative biosynthetic process | 1 | 8 | 0.022 |
| GO:0009113 | Biological Process | purine base biosynthetic process | 1 | 10 | 0.027 |
| GO:0006012 | Biological Process | galactose metabolic process | 1 | 11 | 0.029 |
| GO:0006144 | Biological Process | purine base metabolic process | 1 | 11 | 0.029 |
| GO:0055085 | Biological Process | transmembrane transport | 5 | 701 | 0.031 |
| GO:0009875 | Biological Process | pollen-pistil interaction | 2 | 114 | 0.034 |
| GO:0008037 | Biological Process | cell recognition | 2 | 114 | 0.034 |
| GO:0048544 | Biological Process | recognition of pollen | 2 | 114 | 0.034 |
| GO:0022414 | Biological Process | reproductive process | 2 | 114 | 0.034 |
| GO:0009856 | Biological Process | pollination | 2 | 114 | 0.034 |
| GO:0051704 | Biological Process | multi-organism process | 2 | 120 | 0.037 |
| GO:0010038 | Biological Process | response to metal ion | 1 | 15 | 0.039 |
| GO:0006694 | Biological Process | steroid biosynthetic process | 1 | 15 | 0.039 |
| GO:0046112 | Biological Process | nucleobase biosynthetic process | 1 | 16 | 0.041 |
| GO:0045454 | Biological Process | cell redox homeostasis | 2 | 132 | 0.044 |
| GO:0000003 | Biological Process | reproduction | 2 | 133 | 0.045 |
| GO:0010035 | Biological Process | response to inorganic substance | 1 | 18 | 0.046 |
| GO:0009112 | Biological Process | nucleobase metabolic process | 1 | 18 | 0.046 |
| GO:0019725 | Biological Process | cellular homeostasis | 2 | 140 | 0.049 |
| GO:0016756 | Molecular Function | glutathione gamma-glutamylcysteinyltransferase activity | 1 | 2 | 0.0074 |
| GO:0004637 | Molecular Function | phosphoribosylamine-glycine ligase activity | 1 | 2 | 0.0074 |
| GO:0004386 | Molecular Function | helicase activity | 3 | 165 | 0.0085 |
| GO:0004867 | Molecular Function | serine-type endopeptidase inhibitor activity | 2 | 60 | 0.01 |
| GO:0004866 | Molecular Function | endopeptidase inhibitor activity | 2 | 72 | 0.015 |

| | | | | | |
|---|---|---|---|---|---|
| GO:0030414 | Molecular Function | peptidase inhibitor activity | 2 | 72 | 0.015 |
| GO:0016755 | Molecular Function | transferase activity, transferring amino-acyl groups | 1 | 6 | 0.017 |
| GO:0003978 | Molecular Function | UDP-glucose 4-epimerase activity | 1 | 8 | 0.022 |
| GO:0005215 | Molecular Function | transporter activity | 6 | 941 | 0.03 |
| GO:0070035 | Molecular Function | purine NTP-dependent helicase activity | 2 | 107 | 0.03 |
| GO:0008026 | Molecular Function | ATP-dependent helicase activity | 2 | 107 | 0.03 |
| GO:0042623 | Molecular Function | ATPase activity, coupled | 3 | 275 | 0.032 |
| GO:0017111 | Molecular Function | nucleoside-triphosphatase activity | 5 | 726 | 0.035 |
| GO:0016462 | Molecular Function | pyrophosphatase activity | 5 | 743 | 0.038 |
| GO:0016857 | Molecular Function | racemase and epimerase activity, acting on carbohydrates and derivatives | 1 | 15 | 0.039 |
| GO:0016818 | Molecular Function | hydrolase activity, acting on acid anhydrides, in phosphorus-containing anhydrides | 5 | 759 | 0.041 |
| GO:0031418 | Molecular Function | L-ascorbic acid binding | 1 | 16 | 0.041 |
| GO:0016817 | Molecular Function | hydrolase activity, acting on acid anhydrides | 5 | 764 | 0.042 |
| GO:0016854 | Molecular Function | racemase and epimerase activity | 1 | 17 | 0.044 |
| GO:0005783 | Cellular Component | endoplasmic reticulum | 2 | 123 | 0.039 |
| GO:0031224 | Cellular Component | intrinsic to membrane | 5 | 775 | 0.044 |

**Private genes (DELs) in indica against rufipogon**

| | | | | | |
|---|---|---|---|---|---|
| GO:0006970 | Biological Process | response to osmotic stress | 2 | 12 | 0.0013 |
| GO:0016070 | Biological Process | RNA metabolic process | 11 | 1088 | 0.0039 |
| GO:0005975 | Biological Process | carbohydrate metabolic process | 9 | 833 | 0.0059 |
| GO:0000956 | Biological Process | nuclear-transcribed mRNA catabolic process | 1 | 1 | 0.0078 |
| GO:0000184 | Biological Process | nuclear-transcribed mRNA catabolic process, nonsense-mediated decay | 1 | 1 | 0.0078 |
| GO:0016071 | Biological Process | mRNA metabolic process | 2 | 35 | 0.0093 |
| GO:0006420 | Biological Process | arginyl-tRNA aminoacylation | 1 | 2 | 0.012 |
| GO:0006401 | Biological Process | RNA catabolic process | 1 | 3 | 0.016 |
| GO:0019877 | Biological Process | diaminopimelate biosynthetic process | 1 | 3 | 0.016 |

| | | | | | |
|---|---|---|---|---|---|
| GO:0006402 | Biological Process | mRNA catabolic process | 1 | 3 | 0.016 |
| GO:0019318 | Biological Process | hexose metabolic process | 3 | 133 | 0.017 |
| GO:0051186 | Biological Process | cofactor metabolic process | 3 | 143 | 0.02 |
| GO:0005996 | Biological Process | monosaccharide metabolic process | 3 | 150 | 0.023 |
| GO:0006207 | Biological Process | 'de novo' pyrimidine base biosynthetic process | 1 | 5 | 0.023 |
| GO:0019856 | Biological Process | pyrimidine base biosynthetic process | 1 | 6 | 0.027 |
| GO:0006013 | Biological Process | mannose metabolic process | 1 | 6 | 0.027 |
| GO:0034641 | Biological Process | cellular nitrogen compound metabolic process | 5 | 435 | 0.03 |
| GO:0006206 | Biological Process | pyrimidine base metabolic process | 1 | 7 | 0.031 |
| GO:0009089 | Biological Process | lysine biosynthetic process via diaminopimelate | 1 | 7 | 0.031 |
| GO:0009085 | Biological Process | lysine biosynthetic process | 1 | 7 | 0.031 |
| GO:0006553 | Biological Process | lysine metabolic process | 1 | 7 | 0.031 |
| GO:0046451 | Biological Process | diaminopimelate metabolic process | 1 | 7 | 0.031 |
| GO:0000398 | Biological Process | nuclear mRNA splicing, via spliceosome | 1 | 8 | 0.035 |
| GO:0000375 | Biological Process | RNA splicing, via transesterification reactions | 1 | 8 | 0.035 |
| GO:0000377 | Biological Process | RNA splicing, via transesterification reactions with bulged adenosine as nucleophile | 1 | 8 | 0.035 |
| GO:0009628 | Biological Process | response to abiotic stimulus | 2 | 74 | 0.036 |
| GO:0006771 | Biological Process | riboflavin metabolic process | 1 | 9 | 0.039 |
| GO:0042727 | Biological Process | riboflavin and derivative biosynthetic process | 1 | 9 | 0.039 |
| GO:0042726 | Biological Process | riboflavin and derivative metabolic process | 1 | 9 | 0.039 |
| GO:0009231 | Biological Process | riboflavin biosynthetic process | 1 | 9 | 0.039 |
| GO:0006066 | Biological Process | alcohol metabolic process | 3 | 202 | 0.047 |
| GO:0043412 | Biological Process | macromolecule modification | 13 | 1979 | 0.048 |
| GO:0030246 | Molecular Function | carbohydrate binding | 5 | 219 | 0.0019 |
| GO:0004437 | Molecular Function | inositol or phosphatidylinositol phosphatase activity | 2 | 29 | 0.0066 |
| GO:0051536 | Molecular Function | iron-sulfur cluster binding | 3 | 95 | 0.0068 |
| GO:0051540 | Molecular Function | metal cluster binding | 3 | 95 | 0.0068 |
| GO:0051539 | Molecular Function | 4 iron, 4 sulfur cluster binding | 2 | 31 | 0.0075 |
| GO:0015925 | Molecular Function | galactosidase activity | 1 | 1 | 0.0078 |

| | | | | | |
|---|---|---|---|---|---|
| GO:0004565 | Molecular Function | beta-galactosidase activity | 1 | 1 | 0.0078 |
| GO:0004070 | Molecular Function | aspartate carbamoyltransferase activity | 1 | 1 | 0.0078 |
| GO:0008840 | Molecular Function | dihydrodipicolinate synthase activity | 1 | 1 | 0.0078 |
| GO:0008531 | Molecular Function | riboflavin kinase activity | 1 | 2 | 0.012 |
| GO:0004814 | Molecular Function | arginine-tRNA ligase activity | 1 | 2 | 0.012 |
| GO:0016743 | Molecular Function | carboxyl- or carbamoyltransferase activity | 1 | 2 | 0.012 |
| GO:0004506 | Molecular Function | squalene monooxygenase activity | 1 | 2 | 0.012 |
| GO:0004559 | Molecular Function | alpha-mannosidase activity | 1 | 3 | 0.016 |
| GO:0016597 | Molecular Function | amino acid binding | 2 | 47 | 0.016 |
| GO:0043176 | Molecular Function | amine binding | 2 | 47 | 0.016 |
| GO:0004527 | Molecular Function | exonuclease activity | 2 | 51 | 0.018 |
| GO:0004096 | Molecular Function | catalase activity | 1 | 4 | 0.02 |
| GO:0017057 | Molecular Function | 6-phosphogluconolactonase activity | 1 | 4 | 0.02 |
| GO:0004553 | Molecular Function | hydrolase activity, hydrolyzing O-glycosyl compounds | 5 | 424 | 0.027 |
| GO:0003899 | Molecular Function | DNA-directed RNA polymerase activity | 2 | 65 | 0.029 |
| GO:0034062 | Molecular Function | RNA polymerase activity | 2 | 65 | 0.029 |
| GO:0004386 | Molecular Function | helicase activity | 3 | 165 | 0.029 |
| GO:0008964 | Molecular Function | phosphoenolpyruvate carboxylase activity | 1 | 7 | 0.031 |
| GO:0031406 | Molecular Function | carboxylic acid binding | 2 | 71 | 0.034 |
| GO:0043565 | Molecular Function | sequence-specific DNA binding | 5 | 453 | 0.035 |
| GO:0015923 | Molecular Function | mannosidase activity | 1 | 9 | 0.039 |
| GO:0003950 | Molecular Function | NAD+ ADP-ribosyltransferase activity | 1 | 9 | 0.039 |
| GO:0016773 | Molecular Function | phosphotransferase activity, alcohol group as acceptor | 12 | 1726 | 0.04 |
| GO:0031177 | Molecular Function | phosphopantetheine binding | 1 | 10 | 0.042 |
| GO:0004611 | Molecular Function | phosphoenolpyruvate carboxykinase activity | 1 | 10 | 0.042 |
| GO:0016798 | Molecular Function | hydrolase activity, acting on glycosyl bonds | 5 | 480 | 0.043 |
| GO:0016301 | Molecular Function | kinase activity | 12 | 1750 | 0.043 |
| GO:0008289 | Molecular Function | lipid binding | 2 | 85 | 0.046 |

| | | | | | |
|---|---|---|---|---|---|
| GO:0005544 | Molecular Function | calcium-dependent phospholipid binding | 1 | 11 | 0.046 |
| GO:0015116 | Molecular Function | sulfate transmembrane transporter activity | 1 | 11 | 0.046 |
| GO:0008271 | Molecular Function | secondary active sulfate transmembrane transporter activity | 1 | 11 | 0.046 |
| GO:0000036 | Molecular Function | acyl carrier activity | 1 | 11 | 0.046 |
| GO:0017076 | Molecular Function | purine nucleotide binding | 20 | 3456 | 0.049 |
| GO:0001883 | Molecular Function | purine nucleoside binding | 19 | 3247 | 0.05 |
| GO:0030554 | Molecular Function | adenyl nucleotide binding | 19 | 3247 | 0.05 |
| GO:0044428 | Cellular Component | nuclear part | 3 | 97 | 0.0072 |
| GO:0009341 | Cellular Component | beta-galactosidase complex | 1 | 1 | 0.0078 |
| GO:0005665 | Cellular Component | DNA-directed RNA polymerase II, core complex | 1 | 2 | 0.012 |
| GO:0055029 | Cellular Component | nuclear DNA-directed RNA polymerase complex | 1 | 5 | 0.023 |
| GO:0000428 | Cellular Component | DNA-directed RNA polymerase complex | 1 | 5 | 0.023 |
| GO:0030880 | Cellular Component | RNA polymerase complex | 1 | 8 | 0.035 |
| GO:0005681 | Cellular Component | spliceosomal complex | 1 | 9 | 0.039 |
| **Private genes (DELs) in japonica against rufipogon** | | | | | |
| GO:0006790 | Biological Process | sulfur metabolic process | 3 | 63 | 0.00098 |
| GO:0006865 | Biological Process | amino acid transport | 2 | 27 | 0.0033 |
| GO:0015837 | Biological Process | amine transport | 2 | 27 | 0.0033 |
| GO:0015849 | Biological Process | organic acid transport | 2 | 33 | 0.0048 |
| GO:0046942 | Biological Process | carboxylic acid transport | 2 | 33 | 0.0048 |
| GO:0000956 | Biological Process | nuclear-transcribed mRNA catabolic process | 1 | 1 | 0.0059 |
| GO:0000184 | Biological Process | nuclear-transcribed mRNA catabolic process, nonsense-mediated decay | 1 | 1 | 0.0059 |
| GO:0006810 | Biological Process | transport | 11 | 1638 | 0.0084 |
| GO:0051234 | Biological Process | establishment of localization | 11 | 1638 | 0.0084 |
| GO:0051179 | Biological Process | localization | 11 | 1651 | 0.0089 |
| GO:0044272 | Biological Process | sulfur compound biosynthetic process | 2 | 48 | 0.0095 |
| GO:0044281 | Biological Process | small molecule metabolic process | 8 | 1034 | 0.011 |

| | | | | | |
|---|---|---|---|---|---|
| GO:0019877 | Biological Process | diaminopimelate biosynthetic process | 1 | 3 | 0.012 |
| GO:0006401 | Biological Process | RNA catabolic process | 1 | 3 | 0.012 |
| GO:0006402 | Biological Process | mRNA catabolic process | 1 | 3 | 0.012 |
| GO:0044283 | Biological Process | small molecule biosynthetic process | 4 | 320 | 0.015 |
| GO:0006268 | Biological Process | DNA unwinding during replication | 1 | 5 | 0.018 |
| GO:0032392 | Biological Process | DNA geometric change | 1 | 5 | 0.018 |
| GO:0032508 | Biological Process | DNA duplex unwinding | 1 | 5 | 0.018 |
| GO:0009228 | Biological Process | thiamin biosynthetic process | 1 | 5 | 0.018 |
| GO:0016192 | Biological Process | vesicle-mediated transport | 3 | 186 | 0.018 |
| GO:0006013 | Biological Process | mannose metabolic process | 1 | 6 | 0.02 |
| GO:0009628 | Biological Process | response to abiotic stimulus | 2 | 74 | 0.021 |
| GO:0006749 | Biological Process | glutathione metabolic process | 1 | 7 | 0.023 |
| GO:0009089 | Biological Process | lysine biosynthetic process via diaminopimelate | 1 | 7 | 0.023 |
| GO:0009085 | Biological Process | lysine biosynthetic process | 1 | 7 | 0.023 |
| GO:0006553 | Biological Process | lysine metabolic process | 1 | 7 | 0.023 |
| GO:0046451 | Biological Process | diaminopimelate metabolic process | 1 | 7 | 0.023 |
| GO:0006772 | Biological Process | thiamin metabolic process | 1 | 8 | 0.026 |
| GO:0042723 | Biological Process | thiamin and derivative metabolic process | 1 | 8 | 0.026 |
| GO:0042724 | Biological Process | thiamin and derivative biosynthetic process | 1 | 8 | 0.026 |
| GO:0016053 | Biological Process | organic acid biosynthetic process | 3 | 222 | 0.029 |
| GO:0046394 | Biological Process | carboxylic acid biosynthetic process | 3 | 222 | 0.029 |
| GO:0044271 | Biological Process | cellular nitrogen compound biosynthetic process | 3 | 231 | 0.032 |
| GO:0006265 | Biological Process | DNA topological change | 1 | 11 | 0.035 |
| GO:0006901 | Biological Process | vesicle coating | 1 | 12 | 0.038 |
| GO:0006900 | Biological Process | membrane budding | 1 | 12 | 0.038 |
| GO:0048268 | Biological Process | clathrin coat assembly | 1 | 12 | 0.038 |
| GO:0070838 | Biological Process | divalent metal ion transport | 1 | 12 | 0.038 |
| GO:0006816 | Biological Process | calcium ion transport | 1 | 12 | 0.038 |
| GO:0006970 | Biological Process | response to osmotic stress | 1 | 12 | 0.038 |
| GO:0016050 | Biological Process | vesicle organization | 1 | 13 | 0.04 |

| | | | | | |
|---|---|---|---|---|---|
| GO:0016070 | Biological Process | RNA metabolic process | 7 | 1088 | 0.042 |
| GO:0006298 | Biological Process | mismatch repair | 1 | 14 | 0.043 |
| GO:0004386 | Molecular Function | helicase activity | 5 | 165 | 0.00015 |
| GO:0042623 | Molecular Function | ATPase activity, coupled | 6 | 275 | 0.00018 |
| GO:0016668 | Molecular Function | oxidoreductase activity, acting on sulfur group of donors, NAD or NADP as acceptor | 2 | 9 | 0.00046 |
| GO:0022804 | Molecular Function | active transmembrane transporter activity | 6 | 333 | 0.00049 |
| GO:0016887 | Molecular Function | ATPase activity | 6 | 355 | 0.00068 |
| GO:0017111 | Molecular Function | nucleoside-triphosphatase activity | 8 | 726 | 0.0014 |
| GO:0016462 | Molecular Function | pyrophosphatase activity | 8 | 743 | 0.0016 |
| GO:0016818 | Molecular Function | hydrolase activity, acting on acid anhydrides, in phosphorus-containing anhydrides | 8 | 759 | 0.0018 |
| GO:0003725 | Molecular Function | double-stranded RNA binding | 2 | 20 | 0.0019 |
| GO:0016817 | Molecular Function | hydrolase activity, acting on acid anhydrides | 8 | 764 | 0.0019 |
| GO:0008289 | Molecular Function | lipid binding | 3 | 85 | 0.0022 |
| GO:0015171 | Molecular Function | amino acid transmembrane transporter activity | 2 | 27 | 0.0033 |
| GO:0005275 | Molecular Function | amine transmembrane transporter activity | 2 | 27 | 0.0033 |
| GO:0008026 | Molecular Function | ATP-dependent helicase activity | 3 | 107 | 0.0042 |
| GO:0070035 | Molecular Function | purine NTP-dependent helicase activity | 3 | 107 | 0.0042 |
| GO:0005543 | Molecular Function | phospholipid binding | 2 | 36 | 0.0056 |
| GO:0008840 | Molecular Function | dihydrodipicolinate synthase activity | 1 | 1 | 0.0059 |
| GO:0005342 | Molecular Function | organic acid transmembrane transporter activity | 2 | 39 | 0.0065 |
| GO:0046943 | Molecular Function | carboxylic acid transmembrane transporter activity | 2 | 39 | 0.0065 |
| GO:0004362 | Molecular Function | glutathione-disulfide reductase activity | 1 | 2 | 0.0088 |
| GO:0015038 | Molecular Function | glutathione disulfide oxidoreductase activity | 1 | 2 | 0.0088 |
| GO:0015037 | Molecular Function | peptide disulfide oxidoreductase activity | 1 | 2 | 0.0088 |
| GO:0042626 | Molecular Function | ATPase activity, coupled to transmembrane movement of substances | 3 | 142 | 0.009 |
| GO:0043492 | Molecular Function | ATPase activity, coupled to movement of substances | 3 | 142 | 0.009 |
| GO:0016820 | Molecular Function | hydrolase activity, acting on acid anhydrides, | 3 | 147 | 0.0099 |

| | | | | | |
|---|---|---|---|---|---|
| | | catalyzing transmembrane movement of substances | | | |
| GO:0050660 | Molecular Function | FAD binding | 3 | 156 | 0.012 |
| GO:0004559 | Molecular Function | alpha-mannosidase activity | 1 | 3 | 0.012 |
| GO:0022857 | Molecular Function | transmembrane transporter activity | 6 | 643 | 0.012 |
| GO:0016853 | Molecular Function | isomerase activity | 3 | 161 | 0.013 |
| GO:0016836 | Molecular Function | hydro-lyase activity | 2 | 58 | 0.014 |
| GO:0015399 | Molecular Function | primary active transmembrane transporter activity | 3 | 170 | 0.014 |
| GO:0015405 | Molecular Function | P-P-bond-hydrolysis-driven transmembrane transporter activity | 3 | 170 | 0.014 |
| GO:0016151 | Molecular Function | nickel ion binding | 1 | 4 | 0.015 |
| GO:0004148 | Molecular Function | dihydrolipoyl dehydrogenase activity | 1 | 4 | 0.015 |
| GO:0017150 | Molecular Function | tRNA dihydrouridine synthase activity | 1 | 4 | 0.015 |
| GO:0016651 | Molecular Function | oxidoreductase activity, acting on NADH or NADPH | 2 | 64 | 0.016 |
| GO:0003917 | Molecular Function | DNA topoisomerase type I activity | 1 | 5 | 0.018 |
| GO:0016412 | Molecular Function | serine O-acyltransferase activity | 1 | 6 | 0.02 |
| GO:0009001 | Molecular Function | serine O-acetyltransferase activity | 1 | 6 | 0.02 |
| GO:0005215 | Molecular Function | transporter activity | 7 | 941 | 0.021 |
| GO:0016667 | Molecular Function | oxidoreductase activity, acting on sulfur group of donors | 2 | 75 | 0.022 |
| GO:0016413 | Molecular Function | O-acetyltransferase activity | 1 | 7 | 0.023 |
| GO:0016835 | Molecular Function | carbon-oxygen lyase activity | 2 | 79 | 0.024 |
| GO:0001883 | Molecular Function | purine nucleoside binding | 16 | 3247 | 0.026 |
| GO:0030554 | Molecular Function | adenyl nucleotide binding | 16 | 3247 | 0.026 |
| GO:0001882 | Molecular Function | nucleoside binding | 16 | 3257 | 0.026 |
| GO:0015923 | Molecular Function | mannosidase activity | 1 | 9 | 0.029 |
| GO:0022892 | Molecular Function | substrate-specific transporter activity | 5 | 593 | 0.031 |
| GO:0005544 | Molecular Function | calcium-dependent phospholipid binding | 1 | 11 | 0.035 |
| GO:0016161 | Molecular Function | beta-amylase activity | 1 | 11 | 0.035 |
| GO:0003916 | Molecular Function | DNA topoisomerase activity | 1 | 11 | 0.035 |

| | | | | | |
|---|---|---|---|---|---|
| GO:0005545 | Molecular Function | phosphatidylinositol binding | 1 | 12 | 0.038 |
| GO:0005388 | Molecular Function | calcium-transporting ATPase activity | 1 | 12 | 0.038 |
| GO:0015085 | Molecular Function | calcium ion transmembrane transporter activity | 1 | 12 | 0.038 |
| GO:0030276 | Molecular Function | clathrin binding | 1 | 12 | 0.038 |
| GO:0047800 | Molecular Function | cysteamine dioxygenase activity | 1 | 13 | 0.04 |
| GO:0017076 | Molecular Function | purine nucleotide binding | 16 | 3456 | 0.042 |
| GO:0030983 | Molecular Function | mismatched DNA binding | 1 | 14 | 0.043 |
| GO:0015018 | Molecular Function | galactosylgalactosylxylosylprotein 3-beta-glucuronosyltransferase activity | 1 | 14 | 0.043 |
| GO:0003690 | Molecular Function | double-stranded DNA binding | 1 | 14 | 0.043 |
| GO:0015020 | Molecular Function | glucuronosyltransferase activity | 1 | 14 | 0.043 |
| GO:0044464 | Cellular Component | cell part | 32 | 5945 | 0.00015 |
| GO:0005623 | Cellular Component | cell | 32 | 5945 | 0.00015 |
| GO:0005622 | Cellular Component | intracellular | 23 | 4116 | 0.0013 |
| GO:0005737 | Cellular Component | cytoplasm | 10 | 1263 | 0.0039 |
| GO:0030118 | Cellular Component | clathrin coat | 2 | 32 | 0.0045 |
| GO:0044424 | Cellular Component | intracellular part | 19 | 3529 | 0.0058 |
| GO:0032300 | Cellular Component | mismatch repair complex | 1 | 1 | 0.0059 |
| GO:0032301 | Cellular Component | MutSalpha complex | 1 | 1 | 0.0059 |
| GO:0032302 | Cellular Component | MutSbeta complex | 1 | 1 | 0.0059 |
| GO:0032991 | Cellular Component | macromolecular complex | 9 | 1207 | 0.0091 |
| GO:0048475 | Cellular Component | coated membrane | 2 | 59 | 0.014 |
| GO:0030117 | Cellular Component | membrane coat | 2 | 59 | 0.014 |
| GO:0043234 | Cellular Component | protein complex | 6 | 706 | 0.018 |
| GO:0044425 | Cellular Component | membrane part | 7 | 1007 | 0.029 |
| GO:0044428 | Cellular Component | nuclear part | 2 | 97 | 0.034 |
| GO:0030119 | Cellular Component | AP-type membrane coat adaptor complex | 1 | 16 | 0.049 |
| GO:0030131 | Cellular Component | clathrin adaptor complex | 1 | 16 | 0.049 |

**Table S12.** Previously identified putative domestication or improvement genes with their ID, function and % ranking in $F_{ST}$ windows between the indica and japonica sample based on either SNPs or SVs. Cells are colored as to whether the gene is in a top 1% peak (dark grey), a top 10% peak (medium gray), or an extreme valley (≥90%) (light grey).

| **Locus** | **MSU or RAP ID** | **Function** | **$F_{ST}$ (i-j)** | |
|---|---|---|---|---|
| | | | SNPs | SVs |
| Sh4 | LOC_Os04g57530 | Shattering | 84.38 | 19.37 |
| qSW5 | Os05g0187500 | Grain width | 88.45 | 47.86 |
| qSH1 | LOC_Os01g62920 | Shattering | 15.63 | 82.11 |
| Sd1 | LOC_Os01g66100 | Semi-dwarfing | 8.96 | 5.83 |
| Wx | Os06g0133000 | Grain quality | 28.25 | 93.72 |
| Badh2.1 | LOC_Os04g39020 | Flavor or fragrance | 74.36 | 24.49 |
| Rc | LOC_Os07g11020 | Red pericarp | 56.27 | 59.57 |
| Bh4 | LOC_Os04g38660, LOC_Os04g38670, LOC_Os04g38680 | Hull color | 97.36 | 50.66 |
| PROG1 | LOC_Os07g05900 | Plant Architecture, tiller angle | 90.09 | 42.10 |
| OsCl | Os06g0205100 | Leaf sheath color & apiculus color | 88.70 | 57.16 |
| TAC1 | Os09g0529300 | Tiller angle | 19.78 | 90.34 |
| Dwarf4 | Os03g0227700 | Leaf architecture | 2.39 | 0.63 |
| SAG13 | Os03g0269100 | Senesence | 35.85 | 3.55 |
| TB1 | LOC_Os03g49880 | Tillering | 0.13 | 66.59 |
| Sh1 | LOC_Os03g44710 | Shattering | 73.87 | 47.69 |
